# *DRD2* and *FOXP2* are implicated in the associations between computerized device use and psychiatric disorders

**DOI:** 10.1101/497420

**Authors:** Frank R Wendt, Carolina Muniz Carvalho, Joel Gelernter, Renato Polimanti

**Author notes:** Corresponding author: Renato Polimanti, PhD. Yale University School of Medicine, Department of Psychiatry. VA CT 116A2, 950 Campbell Avenue, West Haven, CT 06516, USA.

## Abstract

The societal health effects of ubiquitous computerized device use (CDU) is mostly unknown. Epidemiological evidence supports associations between CDU and psychiatric traits, but the underlying biological mechanisms are unclear. We investigated genetic overlaps, causal relationships, and molecular pathways shared between these traits using genome-wide data regarding CDU (UK Biobank; up to N=361,194 individuals) and Psychiatric Genomics Consortium phenotypes (14,477<N<150,064). The strongest genetic correlations were between “*weekly usage of mobile phone in last 3 months*” (*PhoneUse*) *vs*. attention deficit hyperactivity disorder (ADHD) (rg=0.425, p=4.59x10^-11^) and “*plays computer games*” (*CompGaming*) *vs*. schizophrenia (SCZ) (rg=-0.271, p=7.16x10^-26^). Latent causal variable analysis did not support causal relationships between these traits, but the observed genetic overlap was related to shared molecular pathways, including: dopamine transport (Gene Ontology:0015872, p*_SCZvsCompGaming_*=2.74x10^-10^) and *DRD2* association (p_SCZ_=7.94x10^-8^; p*_CompGaming_*=3.98x10^-25^), and *FOXP2* association (p_ADHD_=9.32x10^-7^; p*_PhoneUse_*=9.00x10^-11^). Our results support epidemiological observations with genetic data, and uncover biological mechanisms underlying psychiatric disorders contribution to CDUs.

## 2. Introduction

The information and communication revolution over recent decades has contributed to the reliance of humans on technological means to store and transmit ideas to the point where in many societies computerized device use (CDU) is ubiquitous.^1^ These changes affect nearly every aspect of the human experience and are apparent in healthcare, recreation, infrastructure, culture, and more. While these developments are in many ways beneficial to individuals and to society, there is growing discussion about potential harms as well. Dependence on technology may contribute to brain changes over time.^2, 3^ A relationship between personal CDU (e.g., cellular phones, video/computer games, two-way pagers, desktop/laptop computers) and psychiatric traits (i.e., inattention,^4^ attention deficit hyperactivity disorder (ADHD),^5^ anxiety disorders,^6^ and schizophrenia (SCZ)) has been defined by epidemiological studies. While these relationships are often explored as being potentially detrimental to human health,^4-6^ there are data suggesting that CDUs also may be beneficial to certain psychiatric conditions; for example video games seem to have a therapeutic effect on SCZ.^7^ Though strong correlations between CDUs and psychiatric disorders have been documented, causality has been difficult to determine due to psychiatric disorder heterogeneity and sample size constraints^8^ – and more importantly, the inability to use conventional designs in a single-ascertainment context to draw such conclusions. Genetic information from large-scale genome-wide association studies (GWAS) can be used to detect genetic overlap, causal relationships, and molecular pathways shared between complex traits.^9, 10^ Elucidating the underlying mechanisms linking these traits will lead to improved understanding of the possibly long-term effects of CDUs, and also may conceivably contribute to developing nonpharmacological therapies for various disorders and provide biological evidence supporting the use of these therapies in combination with pharmacological intervention.

Due to the epidemiological evidence that CDU may be a contributing risk factor to various psychiatric disorders, or vice versa, we investigated the molecular mechanisms linking these traits using summary association data from GWASs (Table 1) generated by the Psychiatric Genomics Consortium (PGC)^11^ and the UK Biobank (UKB).^12^

**Table 1.**
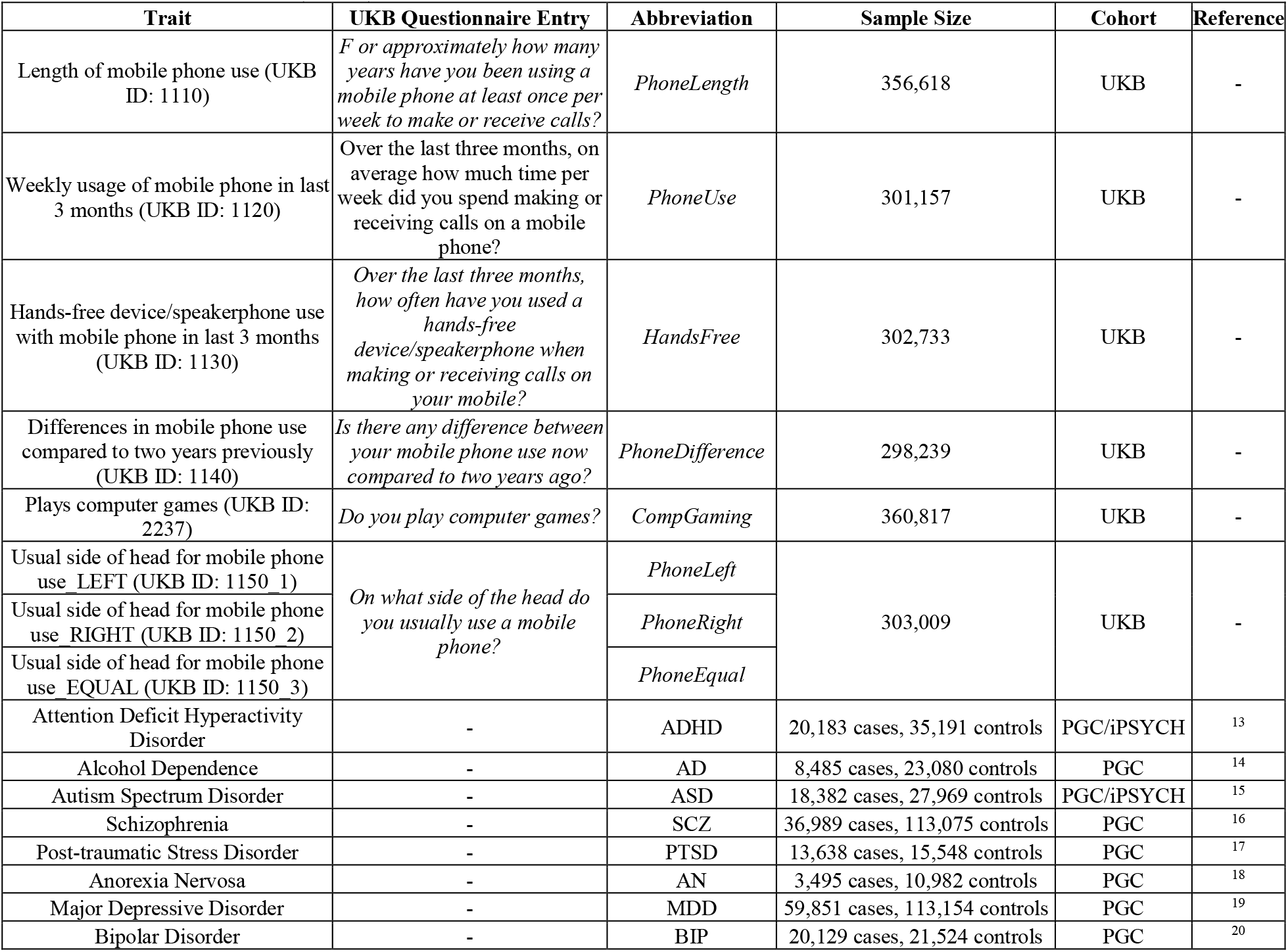
General information for computerized device UK Biobank (UKB) traits and Psychiatric Genomes Consortium (PGC) disorders.

| Trait | UKB Questionnaire Entry | Abbreviation | Sample Size | Cohort | Reference |
| --- | --- | --- | --- | --- | --- |
| Length of mobile phone use (UKB ID: 1110) | *F or approximately how many years have you been using a mobile phone at least once per week to make or receive calls?* | *PhoneLength* | 356,618 | UKB | - |
| Weekly usage of mobile phone in last 3 months (UKB ID: 1120) | *Over the last three months, on average how much time per week did you spend making or receiving calls on a mobile phone?* | *PhoneUse* | 301,157 | UKB | - |
| Hands-free device/speakerphone use with mobile phone in last 3 months (UKB ID: 1130) | *Over the last three months, how often have you used a hands-free device/speakerphone when making or receiving calls on your mobile?* | *HandsFree* | 302,733 | UKB | - |
| Differences in mobile phone use compared to two years previously (UKB ID: 1140) | *Is there any difference between your mobile phone use now compared to two years ago?* | *PhoneDifference* | 298,239 | UKB | - |
| Plays computer games (UKB ID: 2237) | *Do you play computer games?* | *CompGaming* | 360,817 | UKB | - |
| Usual side of head for mobile phone use_LEFT (UKB ID: 1150_1) | *On what side of the head do you usually use a mobile phone?* | *PhoneLeft* | 303,009 | UKB | - |
| Usual side of head for mobile phone use_RIGHT (UKB ID: 1150_2) |  | *PhoneRight* |  |  |  |
| Usual side of head for mobile phone use_EQUAL (UKB ID: 1150_3) |  | *PhoneEqual* |  |  |  |
| Attention Deficit Hyperactivity Disorder | - | ADHD | 20,183 cases, 35,191 controls | PGC/iPSYCH | <sup>13</sup> |
| Alcohol Dependence | - | AD | 8,485 cases, 23,080 controls | PGC | <sup>14</sup> |
| Autism Spectrum Disorder | - | ASD | 18,382 cases, 27,969 controls | PGC/iPSYCH | <sup>15</sup> |
| Schizophrenia | - | SCZ | 36,989 cases, 113,075 controls | PGC | <sup>16</sup> |
| Post-traumatic Stress Disorder | - | PTSD | 13,638 cases, 15,548 controls | PGC | <sup>17</sup> |
| Anorexia Nervosa | - | AN | 3,495 cases, 10,982 controls | PGC | <sup>18</sup> |
| Major Depressive Disorder | - | MDD | 59,851 cases, 113,154 controls | PGC | <sup>19</sup> |
| Bipolar Disorder | - | BIP | 20,129 cases, 21,524 controls | PGC | <sup>20</sup> |

## 3. Results

### 3.1 Genetic Correlation and Polygenic Risk Scores between Computerized Device Use and Psychiatric Disorders

Linkage disequilibrium score regression (LDSC)^21^ was used to estimate the genetic correlation between CDU traits and psychiatric disorders (Table 1 and Fig. 1). The strongest genetic correlations were observed between *PhoneUse* (UKB Field ID: 1120; “*weekly usage of mobile phone in last 3 months*”) *vs*. ADHD (r_g_=0.425, p=4.59x10^-11^) and *CompGaming* (UKB Field ID: 2237; “*plays computer games*”) *vs*. SCZ (r_g_=-0.271, p=7.16x10^-26^). After Bonferroni correction (p<8x10^-4^), 18 significant genetic correlations were observed between CDU *vs*. PGC traits (Table S1).

**Fig. 1.**
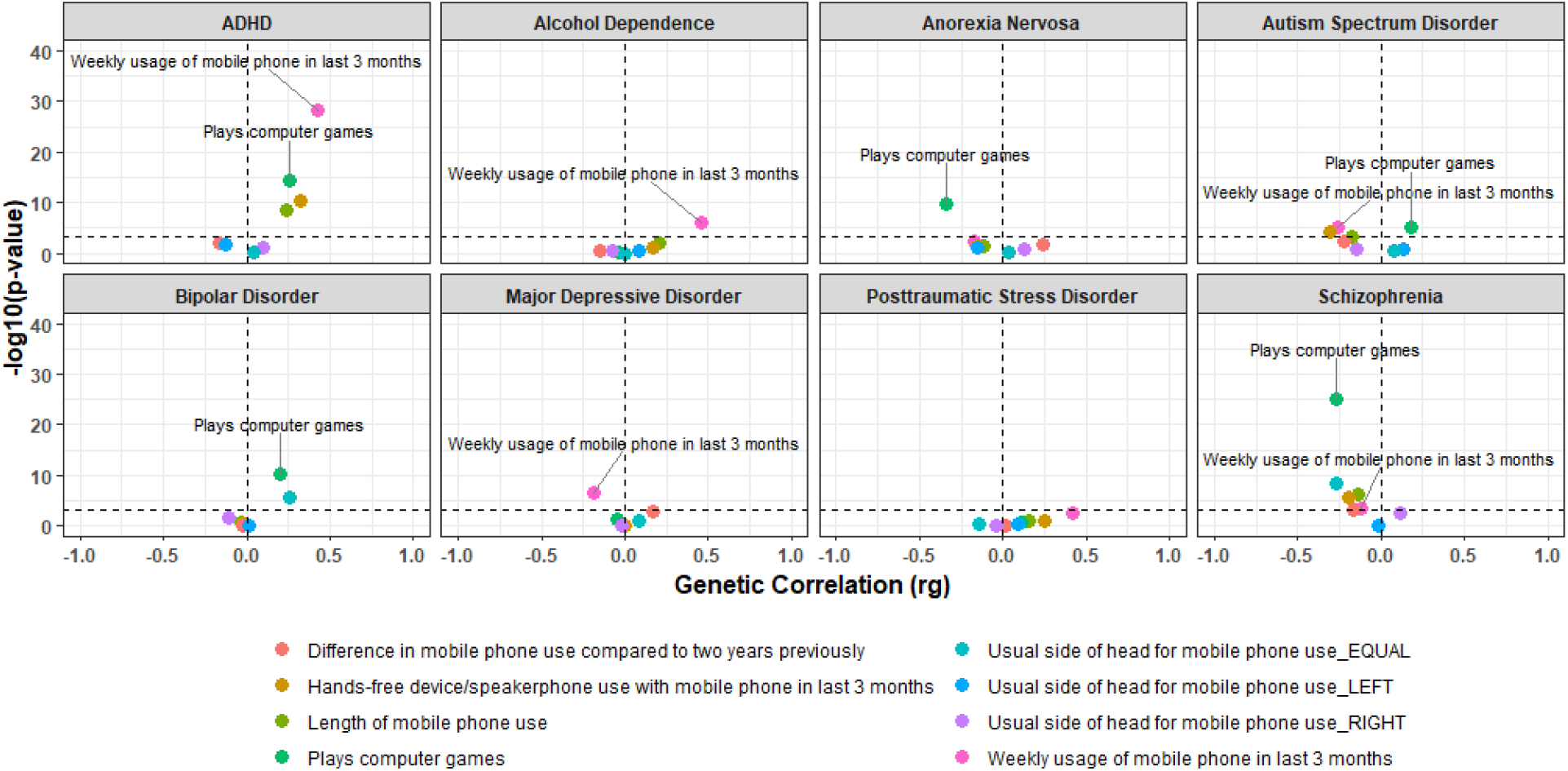
Linkage disequilibrium score regression for computerized device use and psychiatric disorder (attention deficit hyperactivity disorder (ADHD), alcohol dependence, autism spectrum disorder, bipolar disorder, anorexia nervosa, major depressive disorder, posttraumatic stress disorder, and schizophrenia) pairs. Note the horizontal dashed line represents the significance threshold after Bonferroni correction (p=7.81x10^-4^). Genetic correlations significant after Bonferroni correction are provided in Table S1.

Polygenic risk score (PRS) evaluation indicated substantial bidirectional prediction with maximum predictions occurring at: *PhoneUse*→ADHD inclusion threshold (PT)=0.535, r^2^=0.2%, p=9.62x10^-28^; ADHD→*PhoneUse* PT=1.00x10^-5^, r^2^=0.03%, p=6.95x10^-20^; *CompGaming*→SCZ PT=0.01, r^2^=0.2%, p=1.19x10^-20^; and SCZ→*CompGaming* PT=0.01, r^2^=0.05%, p=6.36x10^-37^, (Figs. S1 and S2). Common genetic variation accounted for 7.28% (SE=0.003), 4.82% (SE=0.003), 23.2% (SE=0.009), and 23.5% (SE=0.015) of the observed scale heritability of *CompGaming, PhoneUse*, SCZ, and ADHD, respectively.

Sex-stratified LDSC (available only for ADHD and PTSD; Table S1 and Fig. S3) supported a strong genetic correlation between ADHD and *PhoneUse* in females (r_g_=0.710, p=5.91x10^-13^), which is substantially greater than that observed in males (r_g_=0.322, p=9.51x10^-7^). PRS demonstrated similar sex differences (Figs. S4 and S5). *PhoneUse* in females predicted 0.22% of variance in ADHD at PT=0.13 (p=3.25x10^-12^) while the reverse prediction (ADHD→*PhoneUse*) was substantially less powerful (PT=0.1, r^2^=0.008%, p=2.08x10^-4^). Conversely, males demonstrated nearly negligible differences in explainable variance between *PhoneUse*→ADHD (PT=0.020, r^2^=0.07%, p=9.26x10^-7^) and ADHD→*PhoneUse* (PT=0.001, r^2^=0.02%, p=6.86x10^-8^). Common genetic variation accounted for 5.10% (SE=0.004), 5.09% (SE=0.005), 13.5% (SE=0.028), and 24.9% (SE=0.021) of the observed scale heritability in *PhoneUse_Female_*, *PhoneUse_Male_*, ADHD_Female_, and ADHD_Male_, respectively.

### 3.2 Mendelian Randomization

Based on the LDSC and PRS results described above, two-sample MR was performed to elucidate the causal relationships between ADHD *vs*. *PhoneUse* and SCZ *vs*. *CompGaming* using genetic instruments derived from large-scale GWAS from the UKB and PGC (Tables S2-S4). The PRS of CDU on psychiatric disorders was consistently more powerful than the reverse relationship so MR was performed testing this direction as the main hypothesis. Variants were included in the genetic instrument based on maximum R^2^ from PRS analyses (i.e., the number of variants included in the genetic instrument varies per exposure-outcome pair). Causal estimates were made using methods based on mean,^22^ median,^23^ and mode^24^ with various adjustments to accommodate weak instruments^25^ and the presence of horizontal pleiotropy^26^ (see MR methods). Unless otherwise noted, the inverse-variance weighted (IVW)^23^ causal estimate is discussed herein while all causal estimates are provided in Table S5. Specifically, we applied random-effect IVW, which is less affected by heterogeneity among the variants included in the genetic instrument than a fixed-effect model. Various sensitivity tests were used to detect whether the three assumptions underlying MR (i.e., the variants are associated with the risk factor, the variants are not associated with confounders of the risk factor and outcome relationship, and the variants are not associated with the outcome, except through the risk factor (i.e., horizontal pleiotropy^26^)) were violated by horizontal pleiotropy and/or heterogeneity (i.e., the compatibility of individual variants within the genetic instrument against risk factor and outcome). The causal effects (β; Fig. 2), not affected by heterogeneity and horizontal pleiotropy (Table S5), were bidirectional with CDUs having greater effects on psychiatric disorders (*PhoneUse*→ADHD β_IVW_=0.132, p_IVW_=1.89x10^-4^; *CompGaming*→SCZ β_IVW_=-0.194, p_IVW_=0.005) than the reverse relationship (ADHD→*PhoneUse* β_IVW_=0.084, p_IVW_=2.86x10^-10^; SCZ→*CompGaming* β_IVW_=-0.020, p_IVW_=6.46x10^-25^). When stratified by sex, the female *PhoneUse*→ADHD relationship was nearly 4-fold greater than that in males (β_IVW_female_=0.287, p_IVW_female_=1.09x10^-7^; β_IVW__male=0.080, p_IVW_male_=0.019).

**Fig. 2.**
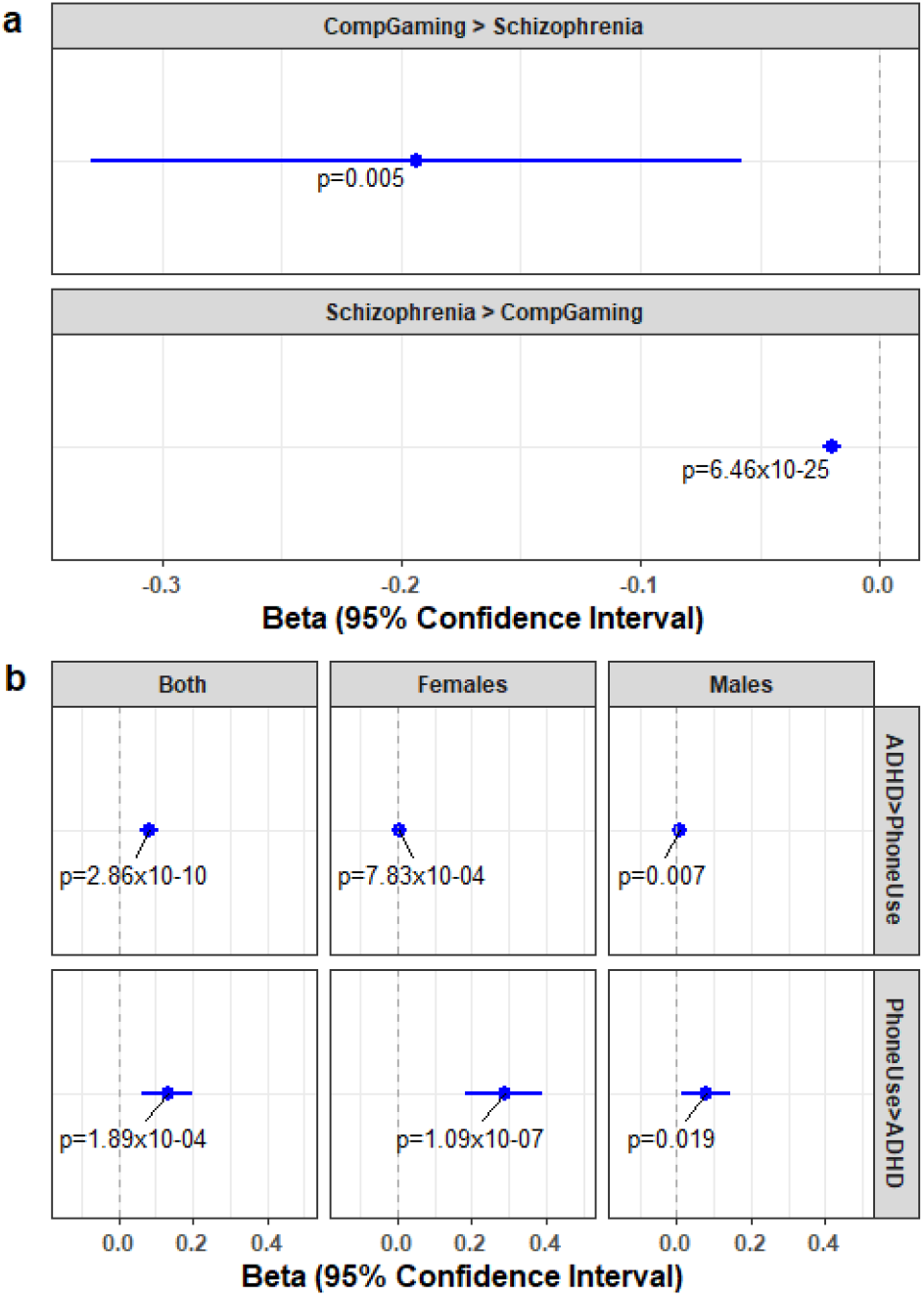
Causal estimates (beta and standard error) and p-values for the relationship between *CompGaming* (UK Biobank Field ID 2237 “*plays computer games*”) and schizophrenia (a) and *PhoneUse* (UK Biobank Field ID 1120 “*weekly usage of mobile phone in the last three months*”) and attention deficit hyperactivity disorder (ADHD; b) using the Mendelian randomization inverse-variance weighted tests.

Multivariable MR were performed to verify the causal estimates of the SCZ→*CompGaming* and ADHD→*PhoneUse* relationships given the genetic correlations of other psychiatric disorders with the same CDU trait. Multivariable MR is an extension of the two-sample MR whereby the genetic instruments of two risk factors are simultaneously used to estimate the causal effect of each risk factor on a single outcome (e.g., CDU trait).^27^ Similarly to SCZ (but with lower significance), ADHD, anorexia nervosa (AN), bipolar disorder (BIP), and autism spectrum disorder (ASD) were genetically correlated with *CompGaming* (Table S1). Conducting two-sample MR, ADHD, AN, and ASD, but not BIP, demonstrated significant effects on *CompGaming* (Table S5). After correcting these results for the SCZ→*CompGaming* effect using a multivariable MR, all estimates were not significant (Fig. 3 and Table S6). Conversely, the SCZ→*CompGaming* causal relationship was still significant after accounting for the effect of other psychiatric disorders (Fig. 3 and Table S6).

**Fig. 3.**
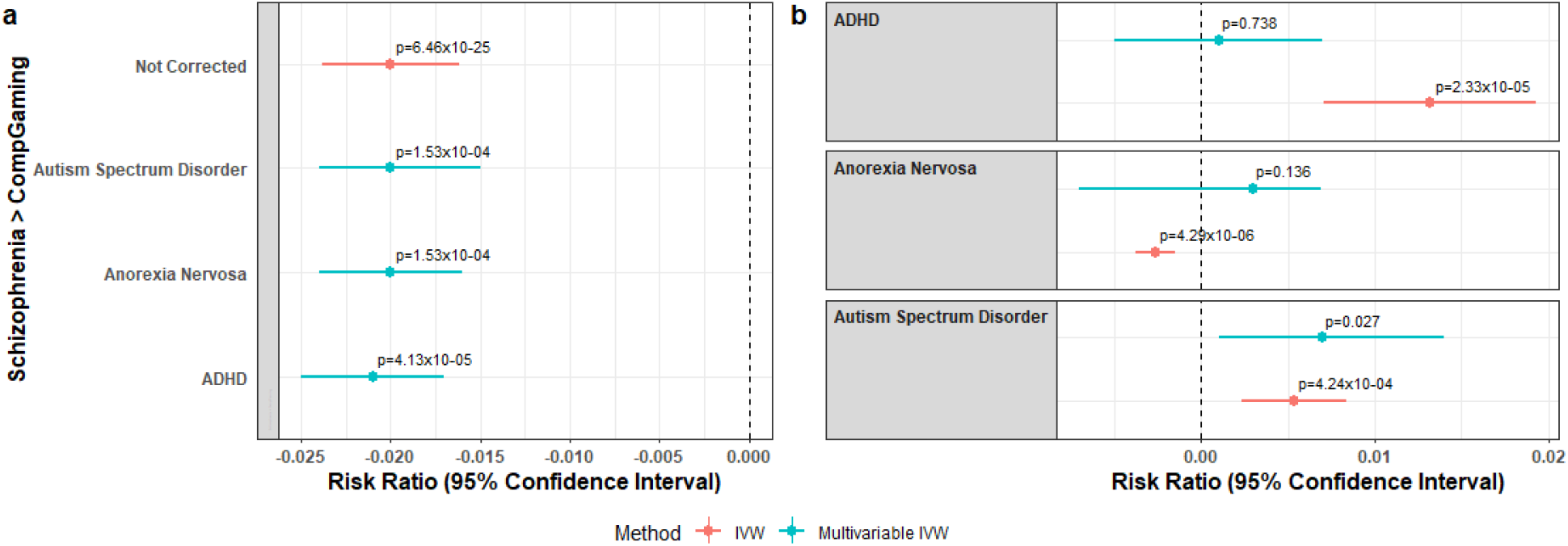
Causal estimates (beta and standard error) and p-values, using the inverse variance weighted (IVW) method, for the relationships between (a) schizophrenia and *CompGaming* (UK Biobank Field ID 2237 “*plays computer games*”) after correction for secondary psychiatric disorder diagnosis (y-axis) and (b) secondary psychiatric disorder diagnosis and *CompGaming* after correction for schizophrenia.

Similarly to ADHD (but with lower significance), ASD, alcohol dependence (AD), SCZ, and major depressive disorder (MDD) were genetically correlated with *PhoneUse*, and some of them demonstrated causal effects on this CDU trait (Table S5). After correction for these other psychiatric diagnoses using a multivariable MR, the ADHD→*PhoneUse* causal estimate remained significant (Fig. 4, Table S6), but, when adjusted for ADHD diagnosis, only the negative causal estimate between ASD_ADHD_→*PhoneUse* remained significant (β_IVW_=-0.019, p_IVW_=1.45x10^-4^; β_Multivariable_IVW_=-0.044, p_Multivariable_IVW_=2.40x10^-8^; Fig. S3 and Table S6).

**Fig. 4.**
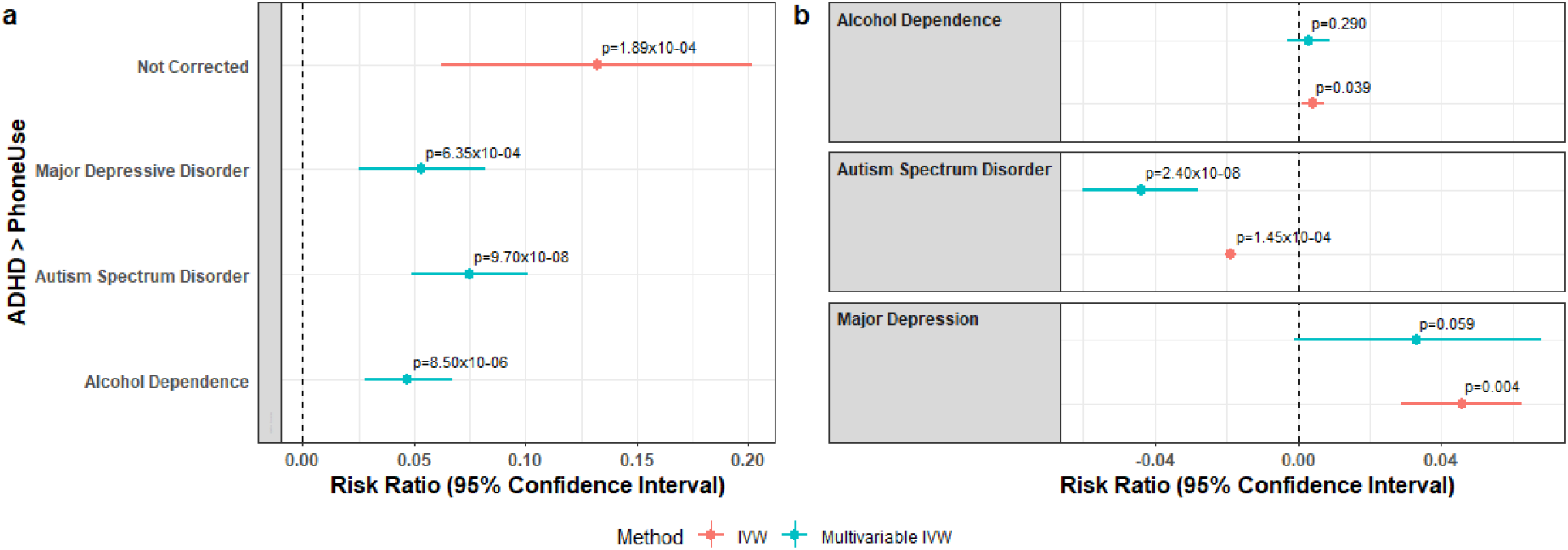
Causal estimates (beta and standard error) and p-values, using the inverse variance weighted (IVW) method, for the relationships between (a) attention deficit hyperactivity disorder (ADHD) and *PhoneUse* (UK Biobank Field ID 1120 “*weekly usage of mobile phone in the last three months*”) after correction for secondary psychiatric disorder diagnosis (y-axis) and (b) secondary psychiatric disorder diagnosis and *PhoneUse* after correction for ADHD.

The variants included in the MR genetic instruments may have effects on multiple traits contributing to observations of causation that are due to shared genetic mechanisms. To evaluate whether the causal estimates between CDU and psychiatric traits were influenced by genetic correlation, we used the latent causal variable (LCV) model.^28^ With LCV, the effects of each GWAS variant are corrected for the heritability of each trait and the genetic correlation between traits. Full GWAS summary data for both traits in each causal relationship (i.e., SCZ *vs*. *CompGaming*, ADHD *vs*. *PhoneUse*, and ASD *vs*. *PhoneUse*) were used to estimate the genetic causality proportion (gĉp) using regression weights. Z-scores for estimating significant trait heritability ranged from 6.42 (ASD) to 14.9 (*CompGaming*), indicating reliable power to detect causality. While partial causality estimates were not significant for any of the relationships, the gĉp estimates suggested the following effect directions: *CompGaming*→SCZ (gĉp=0.099, 0.565 SE, p_LCV_=0.478), *PhoneUse*→ADHD (gĉp=0.02, 0.590 SE, p_LCV_=0.492), and ASD→*PhoneUse* (gĉp=0.065, 0.399 SE, p_LCV_=0.280). Based on these LCV results, we concluded that the MR causal estimates between CDU traits and psychiatric disorders were not independent of the genetic correlations between them.

### 3.3 Genetic Similarities and Differences between Computerized Device use and Psychiatric Disorders

#### 3.3.1 Schizophrenia and CompGaming

SCZ and *CompGaming* have a negative genetic correlation suggesting shared genetic architecture with opposing effects, contributing to phenotype. FUnctional Mapping and Annotation (FUMA)^29^ was used to identify genetic overlap between CDUs and psychiatric disorder GWASs based on a genome-wide significance p<5x10^-8^, minor allele frequency ≥ 0.01, and LD blocks merged at < 250kb for LD r^2^≥0.6 with lead variant. In the SCZ GWAS (36,989 cases and 113,075 controls), 98 genomic risk loci were identified, representing 126 individual significant variants (LD r^2^<0.1; Table S7); in the *CompGaming* GWAS (301,157 subjects), 28 genomic risk loci were identified, representing 32 individual significant variants (LD r^2^<0.1; Table S8). One variant (rs62512616) met genome-wide significance in both *CompGaming* (p=3.01x10^-8^) and SCZ (2.30x10^-9^) GWAS; it maps to the intronic region of t-SNARE domain containing 1 (*TSNARE1*). The rs62512616 variant has opposite effects in SCZ (log_10_(odds ratio)=0.025, SE=0.011 and *CompGaming* (beta=-0.007, SE=0.001). Furthermore, one (rs2514218) and three (rs2734837, rs4648319, and rs4936271) lead variants from the SCZ and *CompGaming* GWAS, respectively, mapped to dopamine receptor D2 (*DRD2*) gene. In addition to *DRD2* and *TSNARE1*, 11 other genes met genome-wide significance for both traits in a gene-based GWAS (p<2.64x10^-6^; Fig 5). Of the 11 other genes associated with both SCZ and *CompGaming*, those with known function have been associated with SCZ and BIP (*FOXO6*)^30^ and various brain disorder diagnoses, all of which are characterized in part by impaired neurological development: congenital neutrophil defect (*VPS45*),^31^ *SATB2*-associated syndrome (*SATB2*),^32^ and ASD (*TCF20*).^33^ In the gene-based GWAS, all 13 genes had positive z-score converted effects in SCZ and *CompGaming* (Fig. 5).

**Fig. 5.**
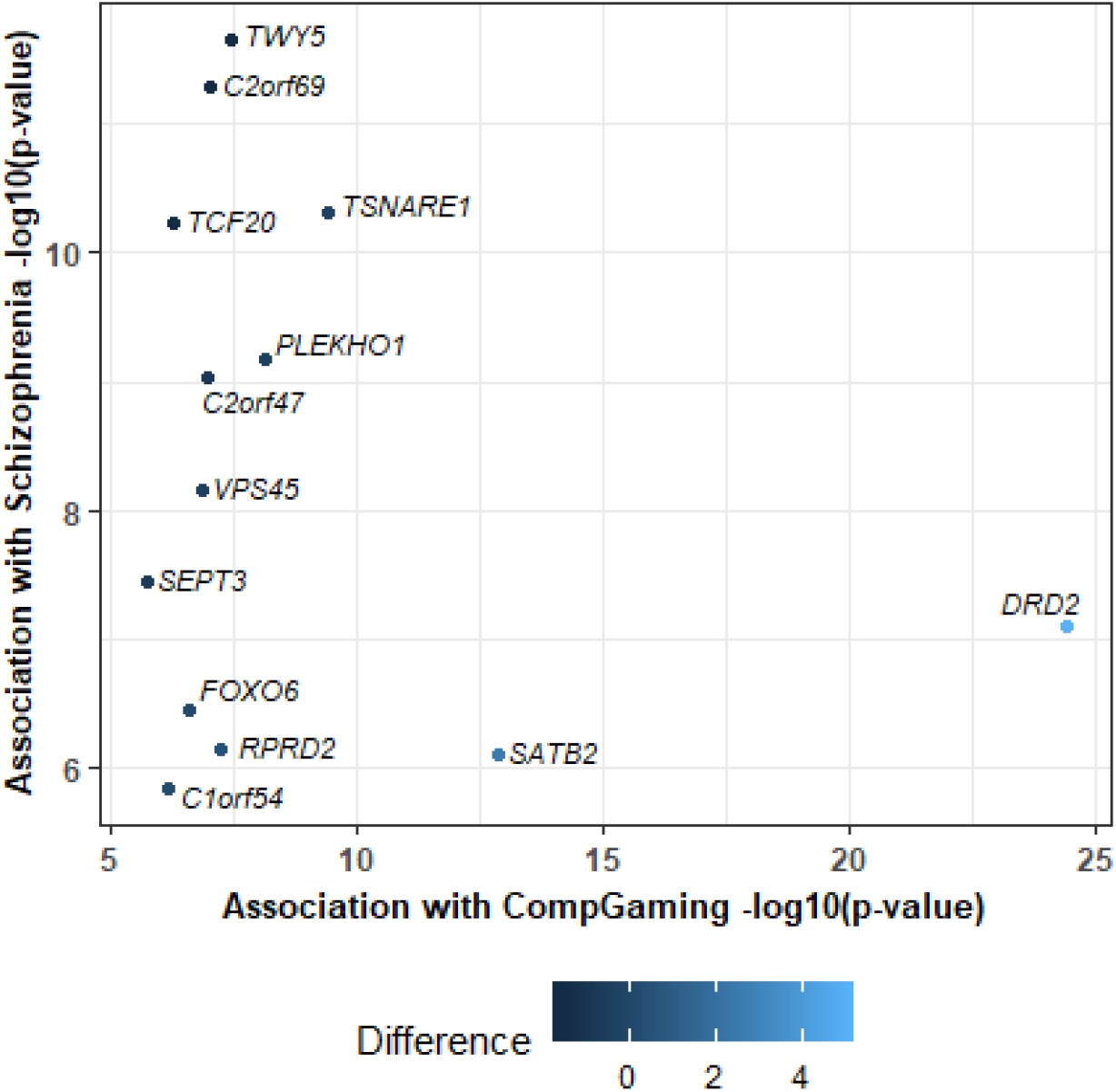
Genes significantly associated with both schizophrenia (SCZ) and UKB Field ID 2237 “*plays computer games*” (*CompGaming*) after Bonferroni correction (p<2.64x10^-6^). The difference in per-gene effects (z-score_*CompGaming*_ minus z-score_SCZ_) on each trait is color-coded.

Twenty-two molecular pathways showed differential enrichments in line with the negative association between SCZ and *CompGaming* (p<4.69x10^-6^; Fig. 6 and Table S9). These differential mechanisms include gene sets related to stress stimulus response (e.g., Systematic name M2492: MAPK11 (p*_SCZvsCompGaming_*=1.02x10^-13^) and MAPK14 Targets (p*_SCZvsCompGaming_*=1.25x10^-8^); and PID p38-Gamma and p38-delta Pathway p*_SCZvsCompGaming_*=1.96x10^-8^) and synapse structure, plasticity, function, and neurotransmitter signaling (e.g., Reactome Down Syndrome Cell Adhesion Molecule Interactions (R-HSA-376172) p*_SCZvsCompGaming_*=2.17x10^-13^; GO:0015872 Dopamine Transport p*_SCZvsCompGaming_*=2.74x10^-10^; GO:0007186 G-Protein Coupled Glutamate Receptor Signaling Pathway p*_SCZvsCompGaming_*=7.22x10^-7^; and GO:0006836 Neurotransmitter Transporter Activity p*_SCZvsCompGaming_*=2.40x10^-6^). The six and seven gene set enrichments surviving Bonferroni correction in SCZ and *CompGaming*, respectively, are provided in Tables S10 and S11, respectively.

**Fig. 6.**
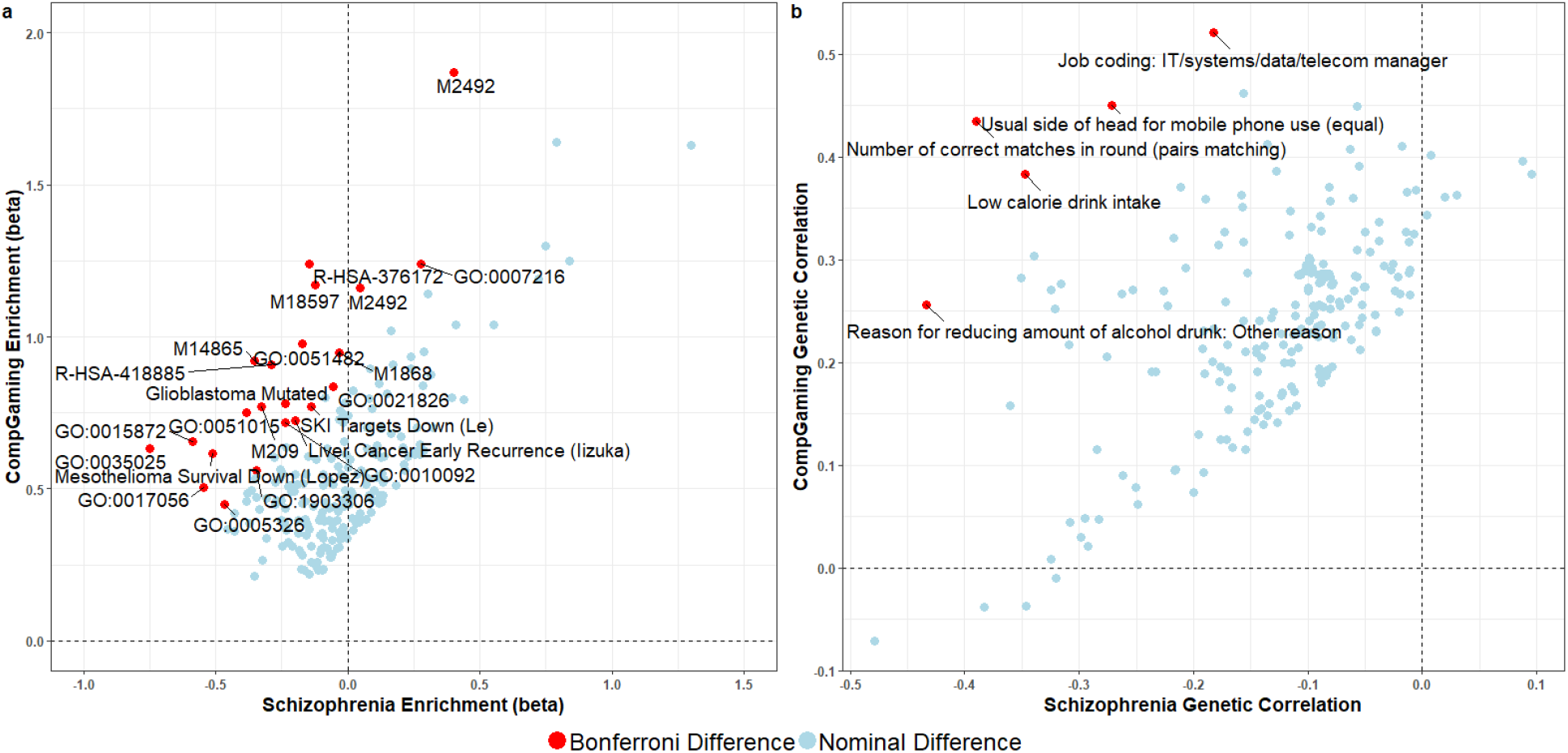
Scatterplots showing gene set enrichment (a; N=221 gene sets) and linkage disequilibrium score regression (b; N=224 traits) results for gene sets and UK Biobank traits with nominally significant differences between schizophrenia and *CompGaming* (UKB Field ID 2237 “*plays computer games*”). Gene sets are labeled with gene ontology (GO), Reactome (R), systematic identifiers (M), or specific study identifiers (last name of first author); the top five most significant differences in genetic correlation are labeled. Detailed results for gene set enrichment and genetic correlations surviving multiple testing correction are provided in Tables S6-S9.

Genetic correlation was calculated between 4,085 phenotypic traits and SCZ/*CompGaming* (Fig. 6 and Table S12). Five traits had significant differences between SCZ and *CompGaming* in line with the negative association between these traits (Table S9). Three of these significantly different traits also were significantly associated with SCZ and *CompGaming* after Bonferroni correction (p<1.22x10^-5^): “*number of correct matches in round (pairs matching*)” (UKB Field ID 398: rg_SCZ_=-0.390, p_SCZ_=4.33x10^-27^; rg*_CompGaming_*=0.435, p*_CompGaming_*=6.69x10^-25^), “*usual side of head for mobile phone use (equal)*” (UKB Field ID 1150_3: rg_SCZ_=-0.272, p_SCZ_=4.51x10^-9^; rg*_CompGaming_*=0.450, p*_CompGaming_*=2.49x10^-17^), and “*reason for reducing amount of alcohol drunk: other reason*” (UKB Field ID 2664_5: rg_SCZ_=-0.434, p_SCZ_=6.35x10^-27^; rg*_CompGaming_*=0.256, p*_CompGaming_*=1.09x10^-11^). Two hundred and sixty-three and 348 UKB traits were independently significantly associated with SCZ and *CompGaming*, respectively, after Bonferroni correction (p<1.14x10^-05^; Fig. 6 and Table S12).

#### 3.3.2 ADHD and PhoneUse

ADHD and *PhoneUse* showed a genetic correlation stronger in females than males. The shared genetic architecture of these traits was investigated using genome-wide data considering both sexes combined (ADHD: 19,099 cases and 34,194 controls; *PhoneUse:* 356,618 subjects), females (ADHD: 4,945 cases and 16,246 controls; *PhoneUse:* 191,522 subjects), and males (ADHD: 14,154 cases and 17,948 controls; *PhoneUse:* 165,096 subjects). In the ADHD GWAS for both sexes combined, 12 genomic risk loci were identified, representing 13 individual significant variants (LD r^2^<0.1; Table S13); in the *PhoneUse* GWAS, 6 genomic risk loci were identified, representing 6 individual significant variants (LD r^2^<0.1; Table S14). There was no overlap in genomic risk loci or individual significant variants between ADHD and *PhoneUse*; however, individual significant loci from both GWAS map to genes involved in fear recognition/consolidation (Sortilin Related VPS10 Domain Containing Receptor 3 (*SORCS3*) in ADHD^34^ and Hypocretin Receptor 2 (HCRTR2)^35^ in *PhoneUse*) and language/speech development/impairment (Semaphorin 6D (*SEMA6D*) in ADHD^36^ and forkhead box transcription factor (*FOXP2*) in ADHD and *PhoneUse*;^37^ Tables S13 and S14). *FOXP2* was the only gene-based genome-wide significant result in both ADHD (p=9.32x10^-7^) and *PhoneUse* (p=9.00x10^-11^). Conversely, *FOXP2* (p=0.084) was not significant in the ASD GWAS (18,382 cases and 27,969 controls), an observation in line with the independent associations of ADHD and ASD with *PhoneUse* (Tables S1 and S15).

GWAS at the variant and gene level for ADHD females and *PhoneUse* males were insufficiently powered to identify genome-wide significant variants at p<5x10^-8^ so these data were explored using a suggestive threshold of p<1x10^-5^. When stratified by sex there were 24 and 3 genomic risk loci, representing 27 and 3 individual significant variants (LD r^2^<0.1; Tables S16 and S17) for ADHD in females and males, respectively, and 3 and 53 genomic risk loci, representing 5 and 64 individual significant variants (LD r^2^<0.1; Tables S18 and S19) for *PhoneUse* in females and males, respectively. There were no significant overlapping variants or genes between ADHD and *PhoneUse* after Bonferroni correction in the sex-stratified cohorts.

Ninety-one, 29, and 39 gene sets were enriched in both ADHD and *PhoneUse* for both sexes, males, and females, respectively. None of these gene sets survived multiple testing correction in both GWAS (Tables S20-S22).

A total of 551 and 491 significant genetic correlations were detected for ADHD and *PhoneUse*, respectively. Three hundred and sixty-nine of these were significantly correlated with both traits after multiple testing correction (p<1.22x10^-5^; Fig. 7 and Table S23). Among the top ten traits correlated with ADHD and *PhoneUse*, three traits were shared: “*age at first sexual intercourse*” (UKB Field ID 2139: rg_ADHD_=-0.623, p_ADHD_=2.17x10^-128^; rg*_PhoneUse_*=-0.579, p*_PhoneUse_*=6.24x10^-132^), “*smoking status: never*” (UKB Field ID 20116_0; rg_ADHD_=-0.523, p_ADHD_=1.91x10^-79^; rg*_PhoneUse_*=-0.386, p*_PhoneUse_*=6.93x10^-56^), and “*qualifications: college or university degree*” (UKB Field ID 6138_1: rg_ADHD_=-0.510, p_ADHD_=7.52x10^-74^; rg*_PhoneUse_*=-0.380, p*_PhoneUse_*=1.70x10^-58^). These genetic correlations were replicated in females and males (Fig. 7 and Tables S24 and S25), with “*age at first sexual intercourse*” as the most significant genetic correlation with ADHD and *PhoneUse* in the sex-stratified cohorts (UKB Field ID 2139: rg_ADHD_Females_=-0.815, p_ADHD_Females_=3.25x10^-22^; rg_PhoneUse_Females_=-0.529, p_PhoneUse_Females_=1.31x10^-64^; rg_ADHD_Males_=-0.610, p_ADHD_Males_=6.56x10^-69^; rg_PhoneUse_Males_=-0.592, p_PhoneUse_Males_=5.03x10^-57^). Differential gene set enrichments and genetic correlations between ASD and *PhoneUse* were observed but did not immediately elucidate clear mechanistic insights into the observed genetic correlation between these traits (Tables S26 and S27).

**Fig. 7.**
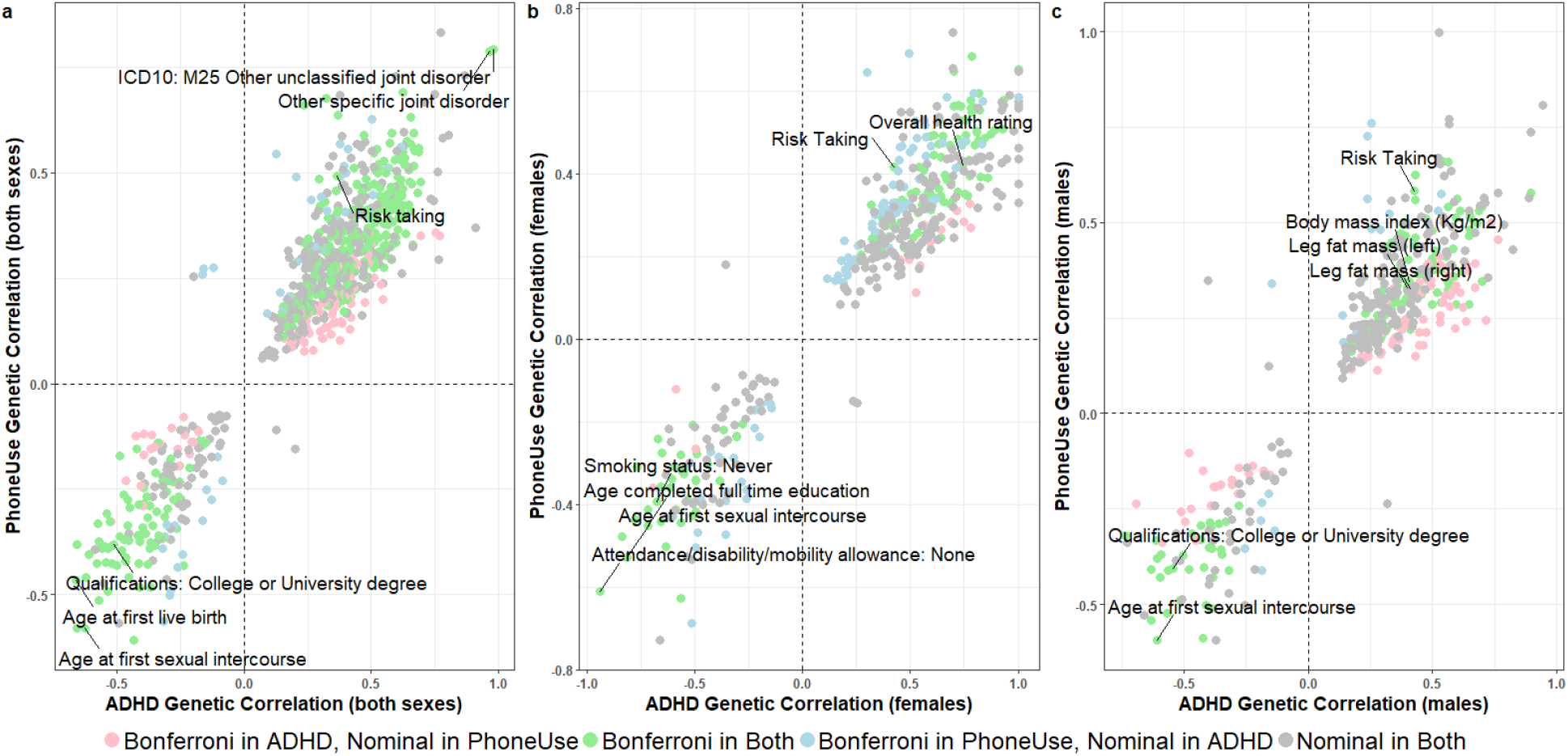
Scatterplots of genetic correlation results for attention deficit hyperactivity disorder (ADHD) and *PhoneUse*, for both sexes combined (a; N=841 traits), females only (b; N=478 traits), and males only (c; N=435 traits). Labeled traits are among the top ten most significant genetic correlations in ADHD and *PhoneUse* (UKB Field ID 1120 “*weekly usage of mobile phone in last three months*”). Detailed genetic correlation results surviving multiple testing correction are provided in Tables S19-S21.

## 4. Discussion

Increasing use of computerized devices has had vast consequences worldwide, and it is important to understand the relationship of this phenomenon to health and disease. Associations between CDUs and psychiatric disorders have been reported in epidemiological studies.^2, 3^ Using data from UKB and PGC, we investigated the underlying mechanisms responsible for these associations. We employed multiple methods based on large-scale genome-wide datasets and observed strong genetic correlation between *CompGaming vs*. SCZ and *PhoneUse vs*. ADHD that appear to be related to shared biological pathways rather than to causal relationships between these traits.

The genetic overlap identified herein suggest that SCZ and *CompGaming* share underlying opposing molecular mechanisms which contribute to the two phenotypes, thereby conferring risk on one trait and protective effects on the other (i.e., biological mechanisms for playing computer games mitigate, to some degree, SCZ symptoms). Previous research has improved our understanding of the neurobiological and psychiatric implications of computer and internet gaming.^38^ In particular, studies have focused on internet gaming disorders (IGDs; such as online gambling), implicating both behavioral (e.g., lack of impulse control,^38^ faulty decision making,^39^ and preference for immediate reward over long-term gain^39^) and biological (i.e., pathways involved with reward processing, sensory-motor coordination, and increased activity of several brain regions)^40, 41^ signatures of addiction. Grey matter density (GMD) in various brain regions of IGD patients was reported to be lower than that in controls^41^. Similarly, SCZ patients consistently show reduced GMD in temporal, prefrontal, and thalamus areas of the brain, overlapping those regions observed in IGD patients.^42^ Notably, decreased GMD has not been observed in professional gamers (computer and video) who lack addiction-related behavioral traits,^41^ who instead have increased GMD in the cingulate cortex, a region of the brain readily implicated in linking motivational outcomes to behavior.

Gene-based GWAS and pathway enrichment analyses implicated *DRD2* and dopamine transport in the SCZ and *CompGaming* protective relationship. Genetic variation in *DRD2* and imbalance of the dopamine transport system in various brain regions are readily observed in SCZ patients^16^ and most antipsychotic drugs are antagonists of D_2_ dopamine receptors (D_2_R), the protein product of DRD2.^16^ The involvement of DRD2 and dopamine transport in *CompGaming* is less obvious mechanistically, but there is evidence that video and computer game playing increases release of dopamine in the striatum.^43^ SCZ patients and healthy computer game users both exhibit increased dopamine levels in various brain regions suggesting that dopamine concentration *per se*, as opposed to alterations in its regulation, is an unlikely contributor to the *SCZ-CompGaming* negative relationship. However, D_2_R density demonstrates tissue- and allele-specific differences,^44^ notably in the cingulate cortex,^43^ which may contribute to GMD increases in non-addicted video and computer game users. Based on the findings of this study and the current literature, in SCZ patients lacking a secondary addiction diagnosis, the pairs-matching component of video and/or computer game playing may increase D_2_R density in the cingulate cortex, thereby increasing GMD in the region over time and providing protective effects against SCZ symptoms.

We observed strong evidence of genetic overlap between *PhoneUse* and ADHD with clear sex differences. *FOXP2* was the only gene overlapping the ADHD and *PhoneUse* GWASs. The *FOXP2* locus encodes a brain-expressed transcription factor strongly implicated in the evolution of speech and language but recent studies report no evidence of recent selection in humans.^45^ Nevertheless, *FOXP2* is consistently implicated in the human ability to communicate via complex speech.^37^ Additionally, we uncovered multiple genetically correlated traits shared between ADHD and *PhoneUse* in the combined and sex-stratified cohorts, including “age *at first sexual intercourse*” and “*risk taking*.” While these traits were detected by the PGC ADHD GWAS of both sexes combined,^13^ the connection to *PhoneUse* is perhaps best explored in the sex-stratified cohorts. The genetic correlation between males and female with ADHD is quite high, suggesting that in general, the same set of common variants contributes to ADHD in both sexes.^46^ We thus consider mechanistic differences which may contribute to higher effects in females than males.

Sex hormones are implicated in various aspects of human speech, including pitch, overall voice profile, and vocal structural anatomy, many of which are influenced by androgen/testosterone expression differences between males and females.^47^ *FOXP2* expression is dampened by androgen/testosterone expression in rats.^48^ Androgen receptor (AR) density in Purkinje neurons of rat brains is modulated by testosterone.^48^ These cell types also demonstrate an increase in *FOXP2*-expressing cells in males relative to females,^49^ although sex differences in FOXP2 concentration have not been observed. Additionally, *FOXP2* expression has been detected in the medial preoptic area (mPOA) of the brain, in which AR density is highly responsive to androgen/testosterone.^48^ This hypothesis also is supported by the “*age at first sexual intercourse*” and “*risk taking*” genetic correlations observed herein and in additional studies of how testosterone influences behavior.^50^ These data support the possibility that androgen/testosterone expression differences in males (upregulated) and females (downregulated) contribute to *FOXP2* expression differences (downregulated in males and upregulated in females), resulting in observations of stronger genetic correlation estimate between ADHD and *PhoneUse* in females.

Our findings demonstrate that CDUs have strong genetic overlap with specific psychiatric disorders. The epidemiological observations of mitigating SCZ symptoms through computer game playing may be generated through mechanisms involving D_2_R density in the cingulate cortex of the brain. Conversely, the length of time spent making and receiving calls on a mobile phone may differentially contribute to ADHD diagnoses in males and females due to sex-specific differences in androgen/testosterone expression. These results offer mechanistic insights into epidemiological observations of CDUs and psychiatric disorder correlations to identify potential pharmacological and nonpharmacological therapeutic targets for psychiatric disorders.

## 5. Materials and Methods

### 5.1 Genetic Data

GWAS data were obtained from PGC and the UKB. Sample sizes for each GWAS used here are provided in Table 1. Due to the limited representation of individuals of non-European descent in these previous studies, all analyses were restricted to GWAS summary association data for individuals of European ancestry. Details regarding quality control criteria and GWAS methods for the UKB and PGC are available at https://github.com/Nealelab/UK_Biobank_GWAS/tree/master/imputed-v2-gwas and https://www.med.unc.edu/pgc/results-and-downloads, respectively. Details regarding the original PGC analyses are described in previous articles.^13-20^

CDU GWAS were created from 6 questions in the UKB: “*length of mobile phone use*” (UKB Questionnaire Entry: For approximately how many years have you been using a mobile phone at least once per week to make or receive calls?), “*weekly usage of mobile phone in the last three months (PhoneUse*)” (UKB Questionnaire Entry: Over the last three months, on average how much time per week did you spend making or receiving calls on a mobile phone?), “*hands-free/speakerphone use*” (UKB Questionnaire Entry: Over the last three months, how often have you used a hands-free device/speakerphone when making or receiving calls on your mobile?), “*difference in mobile phone use compared to two year ago*” (UKB Questionnaire Entry: Is there any difference between your mobile phone use now compared to two years ago?), “*plays computer games (CompGaming*)” (UKB Questionnaire Entry: Do you play computer games?), and “*side of head on which mobile phone is typically held*” (UKB Questionnaire Entry: On what side of the head do you usually use a mobile phone?)).

### 5.2 Genetic Correlation

GWAS summary association data related to CDUs (i.e., “*length of mobile phone use*,” “*weekly usage of mobile phone in the last three months (PhoneUse*),” “*hands-free/speakerphone use*,” “*difference in mobile phone use compared to two year ago*,” “*plays computer games (CompGaming*),” and “*side of head on which mobile phone is typically held*”) and psychiatric disorders (i.e., ADHD,^13^ AD,^14^ AN,^18^ ASD,^15^ BIP,^20^ MDD,^19^ PTSD,^17^ and SCZ^16^) were used to calculate heritability estimates and genetic correlation using the LDSC method.^21, 51^ Subsequent genetic correlations were evaluated using GWAS summary statistics for the remaining UKB traits (4,085 for both sexes combined, 3,291 for females only, and 3,148 for males only). In this analysis, for quantitative traits we considered GWAS summary association data generated from a normalized outcome. Bonferroni correction was used accordingly to adjust for the number of tests performed.

### 5.3 Polygenic Risk Scoring and Genetic Instrument Selection

The polygenic architecture of CDU traits and psychiatric disorders was evaluated using PRS performed in PRSice v1.25^52^ with stringent clumping criteria (clump_kb_=10,000 and clump_r2_=0.001) to minimize correlation among genetic instruments which may bias downstream causal estimates between phenotypes. The region encoding the major histocompatibility complex was removed from PRS analyses. Genetic instruments were derived from the best-fit PRS p-value threshold (ranging from 1.0 to 5x10^-8^) for each relationship tested.

### 5.4 Mendelian Randomization

The causality between CDU and psychiatric disorders was evaluated using two-sample MR with genetic instruments included based on best-fit PRS estimates. Different MR methods have sensitivities to certain conditions but remain robust to others (e.g., many weak genetic instruments included in analysis) so multiple methods were used to verify the stability of causal estimate across methods. These included methods based on mean,^22^ median,^23^ and mode^24^ with various adjustments to accommodate weak instruments^25^ and the presence of horizontal pleiotropy.^53^ Appropriate sensitivity tests were used to evaluate the presence of pleiotropic effects (MR-Egger, MR robust adjusted profile score (RAPS) over dispersion and loss-of-function,^25^ and MR pleiotropy residual sum and outlier (PRESSO^53^)), heterogeneity, and pervasive effect of outliers in variants in the genetic instrument (leave-one-out). Two sample MR tests were conducted using the R package TwoSampleMR.^54^

Multivariable MR were performed using the MendelianRandomization R package^27, 55^ and genetic instruments contributing to the best-fit PRS analysis for each exposure (i.e., genetic instruments for multivariable MR included best-fit PRS variants from Disorder1→ComputerizedDeviceUse and best-fit PRS variants from Disorder2→ComputerizedDeviceUse).

The LCV model^28^ was used estimate the gĉp between traits using z-score converted per-variant effects and regression weights for genome-wide summary statistics.

### 5.5 Enrichment Analyses

Molecular pathways and gene ontology (GO) annotations contributing to the polygenic architecture of CDU and psychiatric disorders were evaluated using Multi-marker Analysis of GenoMic Annotation (MAGMA v1.06),^56^ implemented in FUMA v1.3.3c.^29^ Bonferroni correction was applied to adjust the enrichment analysis results for the 10,651 annotations tested.

## Supporting information

## 6. Acknowledgements

This study was supported by the Simons Foundation Autism Research Initiative (SFARI Explorer Award: 534858) and the American Foundation for Suicide Prevention (YIG-1-109-16). CMC was supported by a Fundação de Amparo à Pesquisa do Estado de São Paulo (FAPESP 2018/05995-4) international fellowship. The authors thank Luke O’Connor from the Department of Epidemiology, Harvard T.H. Chan School of Public Health for his assistance with interpreting the LCV model results.

## 7. Author contributions

FRW and RP conceived the analyses; RP received grant funding; FRW performed the analyses; CMC, JG, and RP provided critical analytic feedback; FRW and RP wrote the first draft and prepared materials for submission; FRW, CMC, JG, and RP reviewed, edited, and approved the manuscript for submission.

