## Supplementary material for "*DRD2* and *FOXP2* are implicated in the associations between computerized device use and psychiatric disorders"

**This PDF file includes:**

Figs. S1 to S5

Tables S1 and S5 to S28

Captions for Tables S2-S4

**Other supplementary materials for this manuscript include the following:**

Tables S2-S4

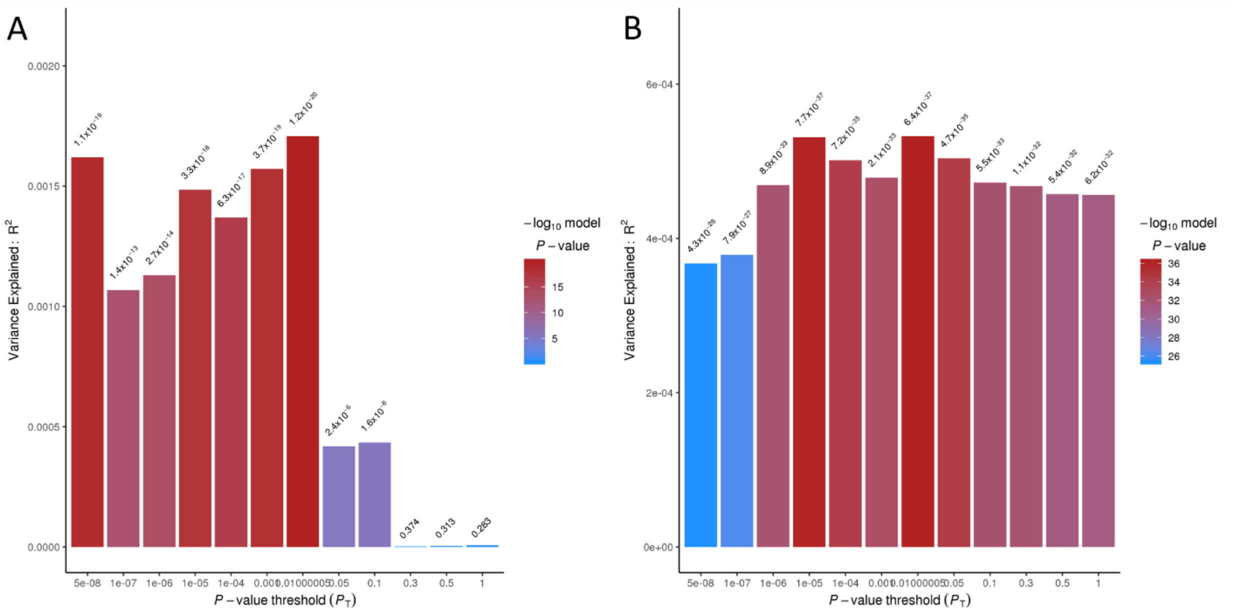

**Fig. S1.** Polygenic risk score barplots for *CompGaming* predicting schizophrenia (A) and schizophrenia predicting *CompGaming* (B) for both sexes combined.

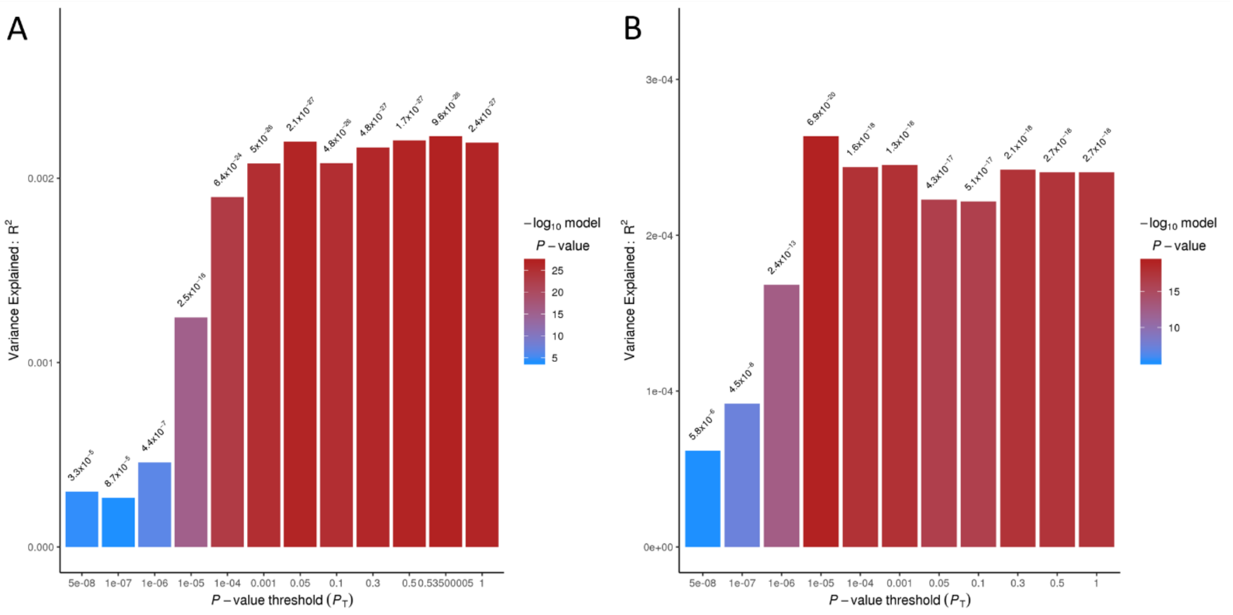

**Fig. S2.** Polygenic risk score barplots for *PhoneUse* predicting ADHD (A) and ADHD predicting *PhoneUse* (B) for both sexes combined.

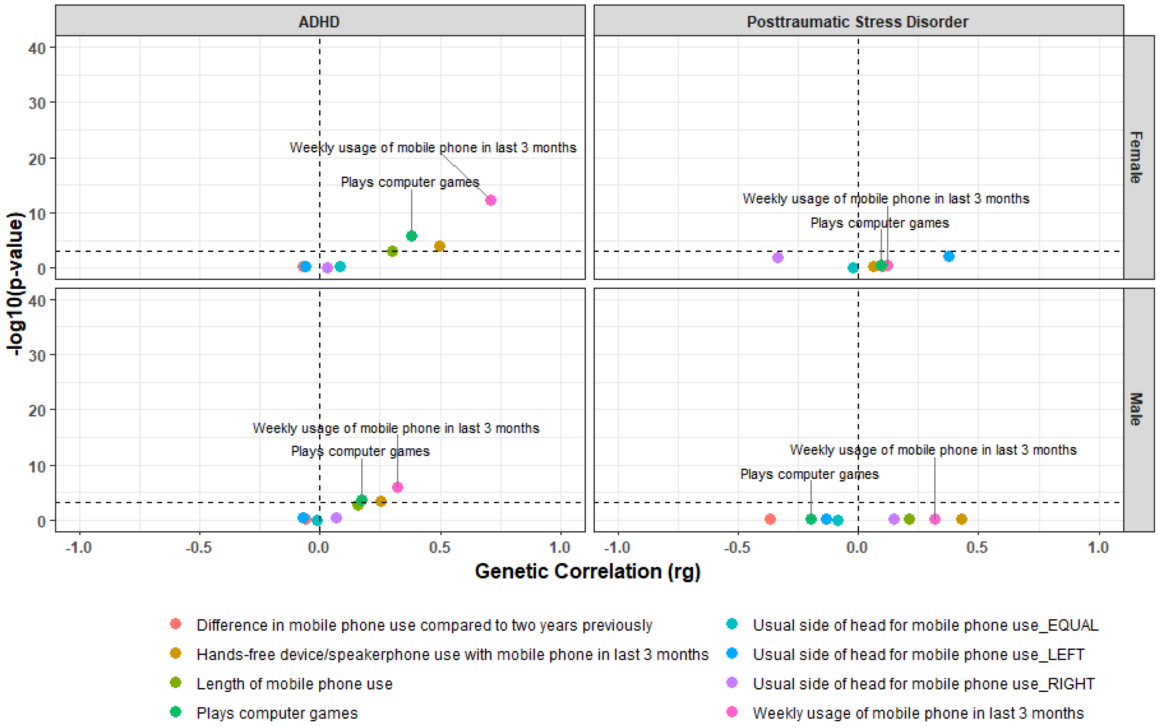

**Fig. S3.** LD score regression for computerized device use and attention deficit hyperactivity disorder (ADHD) and posttraumatic stress disorder stratified by sex.

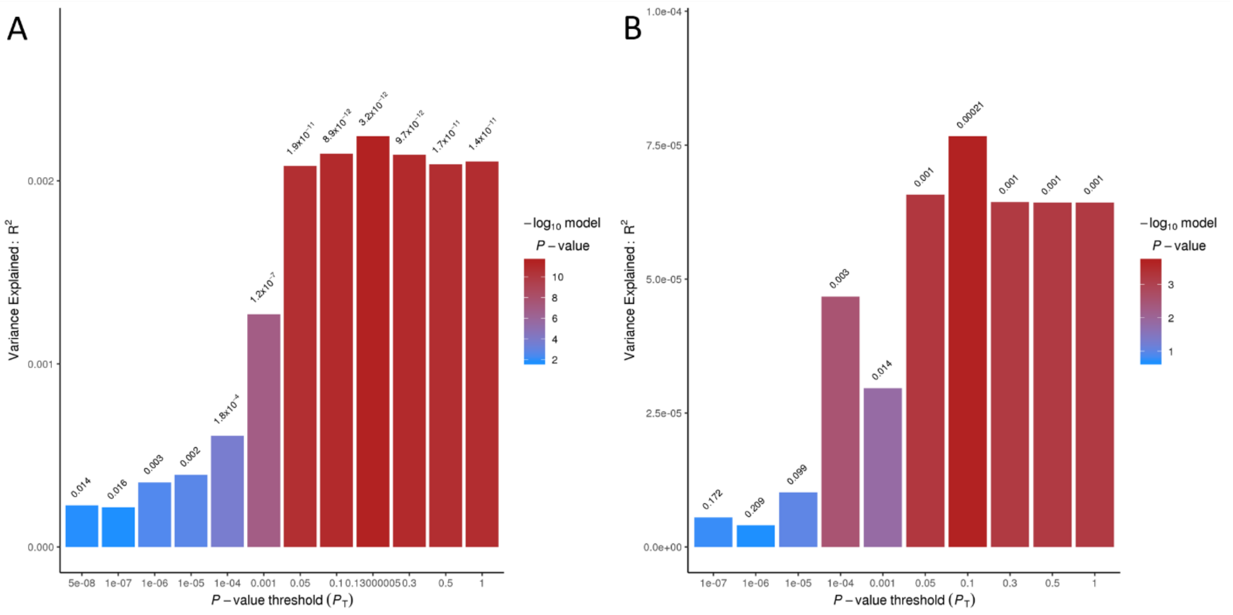

**Fig S4.** Polygenic risk score barplots for *PhoneUse* predicting ADHD (A) and ADHD predicting *PhoneUse* (B) for females.

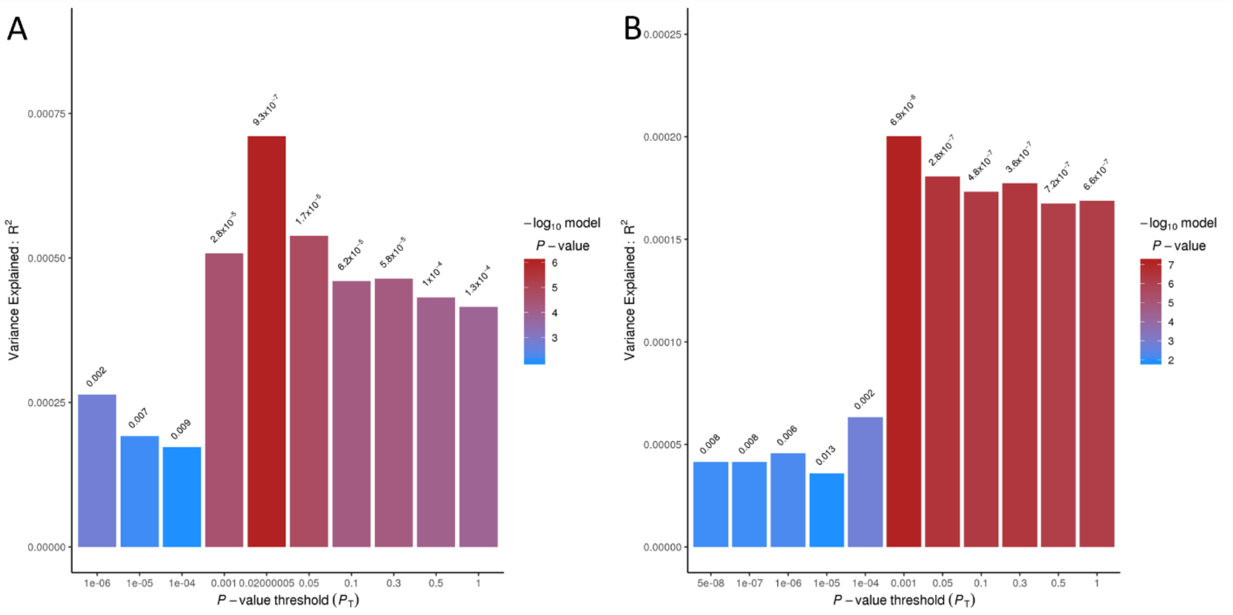

**Fig. S5.** Polygenic risk score barplots for *PhoneUse* predicting ADHD (A) and ADHD predicting *PhoneUse* (B) for males.

**Table S1.** Genetic correlations (rg) surviving Bonferroni correction (sexes combined $p<8x10^{-4}$; sex stratified p<0.002) between computerized device use traits and psychiatric disorders.

| UK Biobank Computerized Device Use Trait | Psychiatric Genomes Consortium Trait | Sex | rg | se | p |
|---|---|---|---|---|---|
| Length of mobile phone use | Attention deficit hyperactivity disorder | Both | 0.2359 | 0.0395 | 2.30x10-9 |
| Weekly usage of mobile phone in last 3 months | Attention deficit hyperactivity disorder | Both | 0.4252 | 0.038 | 4.59x10-29 |
| Hands-free device/speakerphone use with mobile phone in last 3 months | Attention deficit hyperactivity disorder | Both | 0.3204 | 0.0484 | 3.59x10-11 |
| Plays computer games | Attention deficit hyperactivity disorder | Both | 0.2565 | 0.0328 | 5.44x10-15 |
| Weekly usage of mobile phone in last 3 months | Alcohol dependence | Both | 0.456 | 0.0924 | 7.98x10-7 |
| Length of mobile phone use | Autism spectrum disorder | Both | -0.1726 | 0.0503 | 6.0x10-4 |
| Weekly usage of mobile phone in last 3 months | Autism spectrum disorder | Both | -0.2544 | 0.0563 | 6.15x10-6 |
| Hands-free device/speakerphone use with mobile phone in last 3 months | Autism spectrum disorder | Both | -0.3086 | 0.0775 | 6.79x10-5 |
| Plays computer games | Autism spectrum disorder | Both | 0.1788 | 0.0394 | 5.59x10-5 |
| Length of mobile phone use | Schizophrenia | Both | -0.1377 | 0.0273 | 4.38x10-7 |
| Weekly usage of mobile phone in last 3 months | Schizophrenia | Both | -0.1149 | 0.032 | 3.0x10-4 |
| Hands-free device/speakerphone use with mobile phone in last 3 months | Schizophrenia | Both | -0.1902 | 0.0406 | 2.88x10-6 |
| Difference in mobile phone use compared to two years previously | Schizophrenia | Both | -0.1679 | 0.0492 | 6.0x10-4 |
| Plays computer games | Schizophrenia | Both | -0.2706 | 0.0257 | 7.16x10-26 |
| Usual side of head for mobile phone use_EQUAL | Schizophrenia | Both | -0.2716 | 0.0463 | 4.51x10-9 |
| Plays computer games | Anorexia nervosa | Both | -0.336 | 0.0526 | 1.73x10-10 |
| Plays computer games | Bipolar disorder | Both | 0.2025 | 0.0308 | 4.87x10-11 |
| Usual side of head for mobile phone use_EQUAL | Bipolar disorder | Both | 0.2604 | 0.0557 | 2.93x10-6 |
| Weekly usage of mobile phone in last 3 months | Major depressive disorder | Both | -0.1847 | 0.0356 | 2.16x10-7 |
| Length of mobile phone use | Attention deficit hyperactivity disorder | Female | 0.3016 | 0.091 | 9.0x10-4 |
| Weekly usage of mobile phone in last 3 months | Attention deficit hyperactivity disorder | Female | 0.7095 | 0.0985 | 5.91x10-31 |
| Hands-free device/speakerphone use with mobile phone in last 3 months | Attention deficit hyperactivity disorder | Female | 0.4954 | 0.129 | 1.0x10-4 |
| Plays computer games | Attention deficit hyperactivity disorder | Female | 0.3801 | 0.0791 | 1.52x10-6 |
| Length of mobile phone use | Attention deficit hyperactivity disorder | Male | 0.1577 | 0.0495 | 0.001 |
| Weekly usage of mobile phone in last 3 months | Attention deficit hyperactivity disorder | Male | 0.3217 | 0.0656 | 9.51x10-7 |
| Hands-free device/speakerphone use with mobile phone in last 3 months | Attention deficit hyperactivity disorder | Male | 0.2522 | 0.0716 | 4.0x10-4 |
| Plays computer games | Attention deficit hyperactivity disorder | Male | 0.1709 | 0.0451 | 2.0x10-4 |

**Table S2. (separate file).** Variants included in the genetic instruments used to investigate the causal relationship between schizophrenia (SCZ) and *CompGaming*.

**Table S3. (separate file).** Variants included in the genetic instruments used to investigate the causal relationship between attention deficit hyperactivity disorder (ADHD) and *PhoneUse*.

**Table S4. (separate file).** Variants included in the genetic instruments used to investigate the causal relationship between autism spectrum disorder (ASD) and *PhoneUse*.

**Table S5.** Two-sample Mendelian randomization (MR) results for all causal estimates (β) between computerized device use traits *PhoneUse* (UK Biobank (UKB) Field ID 1120 "weekly usage of mobile phone in last 3 months") and *CompGaming* (UKB Field ID 2237 "plays computer games") and psychiatric disorders (attention deficit hyperactivity disorder (ADHD), schizophrenia (SCZ), autism spectrum disorder (ASD), bipolar disorder, alcohol dependence (AD), major depressive disorder (MDD), posttraumatic stress disorder, and anorexia nervosa, using the inverse-variance weighted (IVW) and MR with robust adjusted profile score (MR RAPS) methods. IVW estimates are based on the MR pleiotropy residual sum and outlier method (i.e., palindromic variants are included in the causal estimate). Sensitivity test statistics also are provided.

| Causal Relationship | Sex | Method | β | se | CI (lower upper) | | p | MR Egger | | | Heterogeneity | | | Over-dispersion | | | Global Test | |
|---|---|---|---|---|---|---|---|---|---|---|---|---|---|---|---|---|---|---|
| | | | | | | | | Intercept | se | p | Q | df | p | Variance | se | p | RSS | p |
| *CompGaming* → SCZ | Both | IVW | -0.173 | 0.043 | -0.256 | -0.09 | 4.80E-05 | -8.71E-04 | 0.001 | 0.394 | 456.8 | 1179 | 1 | 0 | 0 | NA | 471.1 | 1 |
| | | MR RAPS | -0.168 | 0.072 | -0.31 | -0.026 | 0.02 | | | | | | | | | | | |
| *SCZ→ CompGaming* | Both | IVW | -0.02 | 0.002 | -0.024 | -0.016 | 3.63E-26 | 9.55E-05 | 1.29E-04 | 0.47 | 924.9 | 1030 | 0.994 | 3.97E-08 | 1.09E-07 | 0.716 | 926.7 | 0.998 |
| | | MR RAPS | -0.028 | 0.003 | -0.034 | -0.022 | 3.47E-20 | | | | | | | | | | | |
| *PhoneUse* → ADHD | Both | IVW | 0.133 | 0.017 | 0.1 | 0.166 | 6.65E-15 | -1.44E-04 | 0.001 | 0.889 | 455.7 | 1931 | 1 | 0 | 0 | NA | 474.2 | 1 |
| | | MR RAPS | 0.134 | 0.037 | 0.061 | 0.207 | 2.91E-04 | | | | | | | | | | | |
| *ADHD→ PhoneUse* | Both | IVW | 0.084 | 0.013 | 0.058 | 0.11 | 1.86E-08 | 0.003 | 0.001 | 0.081 | 65.32 | 67 | 0.535 | 0 | 0 | NA | 67.22 | 0.755 |
| | | MR RAPS | 0.106 | 0.109 | -0.107 | 0.319 | 1.16E-08 | | | | | | | | | | | |
| *PhoneUse* → ADHD | Females | IVW | 0.287 | 0.048 | 0.193 | 0.381 | 2.50E-09 | 3.00E-03 | 0 | 0.29 | 1174 | 1498 | 1 | 0 | 0 | NA | 1175 | 1 |
| | | MR RAPS | 0.309 | 0.062 | 0.188 | 0.43 | 5.53E-07 | | | | | | | | | | | |
| *ADHD→ PhoneUse* | Females | IVW | 0.005 | 0.001 | 0.002 | 0.007 | 1.08E-04 | 6.06E-04 | 3.99E-04 | 0.13 | 1125 | 1503 | 1 | 0 | 0 | NA | 1127 | 1 |
| | | MR RAPS | 0.005 | 0.002 | 0.002 | 0.008 | 1.57E-03 | | | | | | | | | | | |
| *PhoneUse* → ADHD | Males | IVW | 0.08 | 0.031 | 0.02 | 0.14 | 8.83E-03 | 3.00E-03 | 0.002 | 0.114 | 829.9 | 1036 | 1 | 0 | 0 | NA | 831.5 | 1 |
| | | MR RAPS | 0.09 | 0.038 | 0.013 | 0.163 | 2.09E-02 | | | | | | | | | | | |
| *ADHD→ PhoneUse* | Males | IVW | 0.009 | 0.001 | 0.006 | 0.011 | 4.31E-03 | 0.001 | 7.66E-04 | 0.086 | 400.6 | 450 | 0.954 | 2.55E-06 | 3.29E-06 | 0.436 | 402.2 | 0.958 |
| | | MR RAPS | 0.009 | 0.004 | 0.002 | 0.017 | 1.12E-02 | | | | | | | | | | | |
| AD→ *PhoneUse* | Both | IVW | 0.004 | 1.71E-03 | 6.98E-04 | 0.007 | 1.80E-02 | 2.62E-04 | 2.25E-04 | 0.25 | 1750 | 2312 | 1 | 0 | 0 | NA | 1752 | 1 |
| | | MR RAPS | 0.013 | 0.008 | -0.003 | 0.028 | 1.14E-01 | | | | | | | | | | | |

| | | | | | | | | | | | | | | | | | | |
|---|---|---|---|---|---|---|---|---|---|---|---|---|---|---|---|---|---|---|
| *PhoneUse* → AD | Both | IVW | 0.078 | 0.028 | 0.022 | 0.134 | 6.07E-03 | 1.42E-04 | 0.003 | 0.592 | 238 | 1269 | 1 | 0 | 0 | NA | 247.9 | 1 |
| | | MR RAPS | 0.079 | 6.83E-02 | -0.054 | 0.213 | 2.45E-01 | | | | | | | | | | | |
| MDD→ *PhoneUse* | Both | IVW | 0.025 | 8.58E-02 | -0.144 | 0.193 | 4.37E-03 | 6.05E-04 | 3.9E-04 | 0.122 | 298.2 | 291 | 0.374 | 4.15E-05 | 2.1E-06 | 0.05 | 299.8 | 0.406 |
| | | MR RAPS | 0.027 | 0.012 | 0.004 | 0.051 | 2.26E-02 | | | | | | | | | | | |
| *PhoneUse* → MDD | Both | IVW | 0.131 | 0.035 | 0.062 | 0.2 | 8.66E-04 | 0 | 0.007 | 0.363 | 3.616 | 26 | 0.999 | 0 | 0 | NA | 4.11 | 1 |
| | | MR RAPS | 0.132 | 9.79E-02 | -0.06 | 0.324 | 1.79E-01 | | | | | | | | | | | |
| SCZ→ *PhoneUse* | Both | IVW | -0.009 | 0.011 | -0.03 | 0.012 | 3.76E-01 | -7.30E-04 | 0.001 | 0.547 | 139.8 | 130 | 0.263 | 2.51E-06 | 2.04E-06 | 0.217 | 141.9 | 0.265 |
| | | MR RAPS | -0.012 | 0.0134 | -0.038 | 0.015 | 3.88E-01 | | | | | | | | | | | |
| *PhoneUse* → SCZ | Both | IVW | -0.031 | 0.011 | -0.052 | -0.009 | 4.84E-03 | -1.54E-04 | 7.14E-04 | 0.83 | 242.4 | 641 | 1 | 0 | 0 | NA | 248.1 | 1 |
| | | MR RAPS | -0.031 | 0.019 | -0.068 | 0.006 | 9.56E-02 | | | | | | | | | | | |
| ASD→ *PhoneUse* | Both | IVW | -0.019 | 0.004 | -0.028 | -0.01 | 3.77E-05 | 2.63E-04 | 3.73E-04 | 0.481 | 711.4 | 846 | 1 | 0 | 0 | NA | 713.1 | 1 |
| | | MR RAPS | -0.027 | 0.008 | -0.043 | -0.011 | 9.16E-04 | | | | | | | | | | | |
| *PhoneUse* → ASD | Both | IVW | -0.029 | 0.0162 | -0.061 | 0.003 | 7.40E-02 | -2.89E-04 | 0.001 | 0.826 | 327 | 1545 | 1 | 0 | 0 | NA | 341.4 | 1 |
| | | MR RAPS | -0.09 | 0.037 | -0.162 | -0.018 | 4.20E-01 | | | | | | | | | | | |
| AN→ *CompGaming* | Both | IVW | -0.003 | 5.28E-04 | -0.004 | -0.002 | 4.99E-07 | -4.02E-04 | -8.35E-04 | 0.627 | 1486 | 1789 | 1 | 0 | 0 | NA | 1487 | 1 |
| | | MR RAPS | -0.006 | 0.001 | -0.009 | -0.003 | 4.94E-04 | | | | | | | | | | | |
| *CompGaming*→ AN | Both | IVW | -0.287 | 0.0848 | -0.453 | -0.121 | 4.19E-04 | -0.002 | 0.003 | 0.414 | 289.1 | 1465 | 1 | 0 | 0 | NA | 297.4 | 1 |
| | | MR RAPS | -0.289 | 2.00E-01 | -0.681 | 0.103 | 1.47E-01 | | | | | | | | | | | |
| BIP→ *CompGaming* | Both | IVW | -0.005 | 2.51E-03 | -0.009 | 3.80E-04 | 7.20E-02 | -2.34E-04 | 2.79E-04 | 0.403 | 215.4 | 233 | 0.79 | 3.37E-07 | 2.39E-07 | 0.158 | 217.2 | 0.796 |
| | | MR RAPS | -0.005 | 0.00379 | -0.013 | 0.002 | 1.25E-01 | | | | | | | | | | | |
| *CompGaming*→ BIP | Both | IVW | -0.149 | 0.053 | -0.252 | -0.046 | 0.005 | -3.99E-04 | 0.002 | 0.797 | 278.8 | 1012 | 1 | 0 | 0 | NA | 292 | 1 |
| | | MR RAPS | -0.146 | 0.104 | -0.35 | 0.058 | 0.161 | | | | | | | | | | | |
| ASD→ *CompGaming* | Both | IVW | 0.005 | 0.001 | 0.003 | 0.008 | 2.02E-04 | -1.36E-04 | 1.01E-04 | 0.179 | 1666 | 1860 | 0.999 | 9.49E-08 | 1.15E-07 | 0.408 | 1668 | 0.999 |
| | | MR RAPS | 0.014 | 0.004 | 0.006 | 0.023 | 9.67E-04 | | | | | | | | | | | |
| *CompGaming* → ASD | Both | IVW | 0.143 | 0.051 | 0.044 | 0.242 | 0.005 | 3.78E-04 | 0.002 | 0.857 | 154.8 | 691 | 1 | 0 | 0 | NA | 162.1 | 1 |

| | | | | | | | | | | | | | | | | | | |
|---|---|---|---|---|---|---|---|---|---|---|---|---|---|---|---|---|---|---|
| | | MR RAPS | 0.138 | 0.112 | -0.082 | 0.358 | 0.217 | | | | | | | | | | | |
| ADHD→ *CompGaming* | Both | IVW | 0.132 | 0.003 | 0.126 | 0.138 | 3.39E-05 | 1.87E-04 | 3.37E-04 | 0.579 | 253.2 | 225 | 0.095 | 5.22E-04 | 2.72E-04 | 0.055 | 255.5 | 0.142 |
| | | MR RAPS | 0.017 | 0.005 | 0.008 | 0.026 | 2.58E-04 | | | | | | | | | | | |
| *CompGaming* → ADHD | Both | IVW | 0.267 | 0.042 | 0.185 | 0.349 | 1.96E-10 | 0.001 | 0.001 | 0.373 | 362.7 | 1605 | 1 | 0 | 0 | NA | 435.5 | 1 |
| | | MR RAPS | 0.275 | 0.093 | 0.093 | 0.457 | 3.10E-03 | | | | | | | | | | | |

**Table S6.** Multivariable Mendelian randomization results using the inverse-variance weighted estimate of the causal relationship between computerized device use traits *PhoneUse* (UK Biobank (UKB) Field ID 1120 "weekly usage of mobile phone in last 3 months") and *CompGaming* (UKB Field ID 2237 "plays computer games") and psychiatric disorders (attention deficit hyperactivity disorder (ADHD), schizophrenia (SCZ), autism spectrum disorder (ASD), bipolar disorder (BIP), alcohol dependence (AD), major depressive disorder (MDD), and anorexia nervosa (ED).

| Exposure | Outcome | Multivariate | Method | β | se | Lower CI | Upper CI | P |
|---|---|---|---|---|---|---|---|---|
| SCZ | *CompGaming* | Uncorrected | IVW | -0.02 | 0.002 | -0.024 | -0.016 | 6.46E-25 |
| SCZ | *CompGaming* | ADHD | Multivariable IVW | -0.021 | 0.002 | -0.025 | -0.017 | 4.13E-05 |
| SCZ | *CompGaming* | ASD | Multivariable IVW | -0.02 | 0.002 | -0.024 | -0.015 | 1.53E-04 |
| SCZ | *CompGaming* | ED | Multivariable IVW | -0.02 | 0.002 | -0.024 | -0.016 | 1.53E-04 |
| ADHD | *CompGaming* | SCZ | Multivariable IVW | 0.001 | 0.003 | -0.005 | 0.007 | 7.38E-01 |
| ADHD | *CompGaming* | Uncorrected | IVW | 0.013 | 0.00311 | 0.007 | 0.019 | 2.33E-05 |
| ASD | *CompGaming* | SCZ | Multivariable IVW | 0.007 | 0.003 | 0.001 | 0.014 | 0.027 |
| ASD | *CompGaming* | Uncorrected | IVW | 0.005 | 0.00152 | 0.002 | 0.008 | 4.24E-04 |
| AN | *CompGaming* | SCZ | Multivariable IVW | 0.003 | 0.002 | -0.007 | 0 | 0.033 |
| AN | *CompGaming* | Uncorrected | IVW | -0.003 | 5.79E-04 | -0.004 | -0.002 | 4.29E-06 |
| ADHD | *PhoneUse* | Uncorrected | IVW | 0.132 | 0.036 | 0.062 | 0.202 | 1.89E-04 |
| ADHD | *PhoneUse* | MDD | Multivariable IVW | 0.053 | 0.015 | 0.025 | 0.082 | 6.35E-04 |
| ADHD | *PhoneUse* | AD | Multivariable IVW | 0.047 | 0.01 | 0.028 | 0.067 | 9.70E-08 |
| ADHD | *PhoneUse* | ASD | Multivariable IVW | 0.075 | 0.013 | 0.049 | 0.101 | 8.50E-06 |
| MDD | *PhoneUse* | ADHD | Multivariable IVW | 0.033 | 0.018 | -0.001 | 0.068 | 0.059 |
| MDD | *PhoneUse* | Uncorrected | IVW | 0.046 | 8.58E-03 | 0.029 | 0.062 | 0.004 |
| AD | *PhoneUse* | ADHD | Multivariable IVW | 0.003 | 0.003 | -0.003 | 0.009 | 0.290 |
| AD | *PhoneUse* | Uncorrected | IVW | 0.004 | 1.67E-03 | 7.77E-04 | 0.007 | 0.039 |
| ASD | *PhoneUse* | ADHD | Multivariable IVW | -0.044 | 0.008 | -0.06 | -0.028 | 2.40E-08 |
| ASD | *PhoneUse* | Uncorrected | IVW | -0.019 | 4.93E-04 | -0.02 | -0.018 | 1.45E-04 |

**Table S7.** Latent Causal Variable (LCV) heritability z-scores. For all analyses, trait 1 is considered

| Trait 1 | Trait 2 | Sex | Trait 1 Heritability z-score | Trait 2 Heritability z-score |
|---|---|---|---|---|
| SCZ | *CompGaming* | Both | 12.9 | 14.9 |
| ADHD and | *PhoneUse* | Both | 7.19 | 7.3 |
| ASD and | *PhoneUse* | Both | 6.42 | 11.69 |
| ADHD and Females | *PhoneUse* | Females | 2.66 | 12.32 |
| ADHD and Males | *PhoneUse* | Males | 5.37 | 4.8 |

**Table S8.** Genomic risk loci and independent variants (LD $r^2$<0.1) identified in schizophrenia GWAS.

| rsID | Chromosome | Position | Alleles | p | Consequence | Gene Symbol | Region Biotype |
|---|---|---|---|---|---|---|---|
| rs9922678 | 16 | 9946319 | A/G | 6.72E-09 | intronic | *GRIN2A* | protein_coding |
| rs9841616 | 3 | 181167585 | A/T | 1.65E-08 | ncRNA_intronic | - | - |
| rs9636107 | 18 | 53200117 | A/G | 9.09E-13 | intronic | - | - |
| rs9607782 | 22 | 41587556 | A/T | 6.76E-12 | ncRNA_intronic | - | - |
| rs950169 | 15 | 84706461 | C/T | 7.62E-11 | exonic | - | - |
| rs9461856 | 6 | 33395199 | A/G | 5.36E-12 | intronic | - | - |
| rs9398171 | 6 | 108983527 | C/T | 3.37E-08 | intronic | *FOXO3* | protein_coding |
| rs832187 | 3 | 63833050 | C/T | 2.58E-08 | intronic | *C3orf49* | protein_coding |
| rs8082590 | 17 | 17958402 | A/G | 6.84E-09 | intronic | *RAI1* | protein_coding |
| rs8044995 | 16 | 68189340 | A/G | 3.27E-08 | intronic | *RANBP10* | protein_coding |
| rs8042374 | 15 | 78908032 | A/G | 1.87E-12 | intronic | - | - |
| rs80256351 | 2 | 58071593 | A/C | 3.20E-08 | intergenic | - | - |
| rs79780963 | 10 | 104952499 | C/T | 7.51E-16 | intronic | - | - |
| rs7951870 | 11 | 46373311 | C/T | 8.25E-11 | intronic | *CREB3L1* | protein_coding |
| rs79212538 | 5 | 151993104 | G/T | 3.84E-08 | intergenic | - | - |
| rs7893279 | 10 | 18745105 | G/T | 3.56E-11 | intronic | - | - |
| rs78322266 | 18 | 53063676 | G/T | 1.10E-08 | intronic | - | - |
| rs7819570 | 8 | 89588626 | G/T | 1.90E-08 | ncRNA_intronic | *MMP16* | protein_coding |
| rs7801375 | 7 | 131567263 | A/G | 2.26E-08 | intergenic | - | - |
| rs77502336 | 11 | 123394636 | C/G | 2.01E-09 | intergenic | *GRAMD1B* | protein_coding |
| rs77149735 | 1 | 243555105 | A/G | 4.40E-09 | intronic | - | - |
| rs764284 | 6 | 27312078 | A/G | 3.27E-11 | intergenic | - | - |
| rs76355118 | 2 | 149412005 | A/G | 2.78E-08 | intronic | *EPC2* | protein_coding |
| rs75968099 | 3 | 36858583 | C/T | 3.39E-12 | intergenic | - | - |
| rs75059851 | 11 | 133822569 | A/G | 1.23E-11 | intronic | - | - |
| rs74979580 | 6 | 25247680 | C/T | 1.12E-13 | ncRNA_intronic | *FAM65B* | protein_coding |
| rs7432375 | 3 | 136288405 | A/G | 5.27E-11 | intronic | *SLC35G2* | protein_coding |
| rs7405404 | 16 | 13749859 | C/T | 3.93E-10 | intergenic | - | - |
| rs73229090 | 8 | 27442127 | A/C | 1.95E-08 | ncRNA_intronic | *EPHX2* | protein_coding |
| rs7312955 | 12 | 123747783 | A/C | 4.68E-10 | intronic | - | - |
| rs72986630 | 19 | 11849736 | C/T | 4.64E-08 | UTR5 | *ZNF823* | protein_coding |
| rs72934570 | 18 | 53533189 | C/T | 3.67E-12 | intergenic | - | - |
| rs715170 | 18 | 53795514 | C/T | 3.47E-08 | ncRNA_intronic | - | - |
| rs6984242 | 8 | 60700469 | A/G | 1.76E-09 | intergenic | - | - |
| rs6940116 | 6 | 27708732 | A/G | 1.36E-16 | intergenic | - | - |
| rs6918586 | 6 | 26097384 | C/T | 6.61E-10 | UTR3 | - | - |
| rs6918354 | 6 | 25444614 | A/T | 8.47E-10 | intronic | - | - |
| rs6913724 | 6 | 27254843 | A/T | 3.01E-09 | intronic | - | - |
| rs6716963 | 2 | 194367540 | A/G | 2.33E-08 | intergenic | - | - |
| rs6704768 | 2 | 233592501 | A/G | 3.15E-12 | intronic | - | - |
| rs6704641 | 2 | 200164252 | A/G | 3.40E-08 | intronic | - | - |
| rs6466055 | 7 | 104929064 | A/C | 2.46E-09 | intronic | - | - |
| rs6434928 | 2 | 198304577 | A/G | 1.48E-11 | intergenic | - | - |
| rs6430095 | 2 | 146439945 | A/G | 1.21E-08 | intergenic | - | - |
| rs6065094 | 20 | 37453194 | A/G | 5.52E-11 | intronic | - | - |
| rs6002655 | 22 | 42603814 | C/T | 1.48E-09 | intronic | - | - |
| rs59979824 | 2 | 193848340 | A/C | 1.08E-08 | intergenic | - | - |
| rs5995756 | 22 | 40000313 | C/T | 2.84E-11 | intronic | - | - |
| rs5937157 | 23 | 68377126 | G/T | 5.74E-11 | intergenic | *PJA1* | protein_coding |
| rs58120505 | 7 | 2029867 | C/T | 6.43E-14 | intronic | - | - |
| rs56205728 | 15 | 40567237 | A/G | 4.92E-08 | intronic | *PAK6* | protein_coding |
| rs55661361 | 11 | 124613957 | A/G | 3.68E-12 | intronic | - | - |
| rs4919680 | 10 | 104570118 | C/T | 3.05E-09 | intronic | - | - |
| rs4766428 | 12 | 110723245 | C/T | 7.09E-10 | intronic | *ATP2A2* | protein_coding |
| rs4711350 | 6 | 33741716 | A/G | 1.50E-08 | intronic | *ITPR3* | protein_coding |
| rs4702 | 15 | 91426560 | A/G | 2.30E-12 | UTR3 | - | - |
| rs4648845 | 1 | 2387101 | C/T | 4.03E-09 | intronic | - | - |
| rs4523957 | 17 | 2208899 | G/T | 1.04E-09 | intronic | *SMG6* | protein_coding |
| rs4391122 | 5 | 60598543 | A/G | 1.73E-13 | intergenic | - | - |
| rs427230 | 20 | 62155111 | C/T | 4.03E-08 | intergenic | *PPDPF* | protein_coding |
| rs4129585 | 8 | 143312933 | A/C | 2.03E-13 | intronic | - | - |
| rs41266839 | 6 | 26409890 | C/G | 8.39E-27 | exonic | *HIST1H3G* | protein_coding |
| rs3849046 | 5 | 137851192 | C/T | 4.83E-09 | intronic | *ETF1* | protein_coding |
| rs3798869 | 6 | 84328660 | A/G | 1.15E-09 | intronic | - | - |
| rs3735025 | 7 | 137074844 | C/T | 7.75E-09 | UTR3 | *DGKI* | protein_coding |

| | | | | | | | |
|---|---|---|---|---|---|---|---|
| rs36068923 | 8 | 111485761 | A/G | 1.05E-10 | intergenic | - | - |
| rs35225200 | 4 | 103146888 | A/C | 9.56E-13 | intergenic | - | - |
| rs34796896 | 3 | 180623255 | A/G | 6.23E-11 | intronic | *DNAJC19* | protein_coding |
| rs34038921 | 6 | 33195383 | C/G | 4.41E-08 | intergenic | *VPS52* | protein_coding |
| rs3131855 | 6 | 29610281 | A/C | 2.22E-12 | intergenic | - | - |
| rs301797 | 1 | 8487323 | A/C | 2.72E-09 | ncRNA_intronic | *SLC45A1* | protein_coding |
| rs2973155 | 5 | 152608619 | C/T | 1.02E-09 | intergenic | - | - |
| rs2945232 | 8 | 8098038 | C/T | 2.03E-08 | ncRNA_exonic | *SGK223* | protein_coding |
| rs2909457 | 2 | 162845855 | A/G | 4.38E-08 | intergenic | *SLC4A10* | protein_coding |
| rs2905426 | 19 | 19478022 | G/T | 6.92E-09 | intergenic | *NCAN* | protein_coding |
| rs2851447 | 12 | 123665113 | C/G | 2.19E-14 | intronic | - | - |
| rs281768 | 2 | 200825240 | A/T | 3.47E-14 | intronic | - | - |
| rs2693698 | 14 | 99719219 | A/G | 1.38E-08 | intronic | *BCL11B* | protein_coding |
| rs2535627 | 3 | 52845105 | C/T | 3.96E-11 | intergenic | - | - |
| rs2514218 | 11 | 113392994 | C/T | 4.09E-10 | intergenic | *DRD2* | protein_coding |
| rs2332700 | 14 | 72417326 | C/G | 1.69E-09 | intronic | - | - |
| rs2239063 | 12 | 2511831 | A/C | 5.39E-09 | intronic | - | - |
| rs215411 | 4 | 23423603 | A/T | 1.22E-08 | intergenic | - | - |
| rs2068012 | 14 | 30190316 | C/T | 4.14E-08 | intronic | - | - |
| rs2053079 | 19 | 30987423 | A/G | 3.79E-09 | intronic | *ZNF536* | protein_coding |
| rs2021078 | 6 | 29412968 | A/G | 1.01E-09 | intronic | - | - |
| rs2018916 | 16 | 63700508 | C/T | 1.26E-08 | intergenic | - | - |
| rs2007044 | 12 | 2344960 | A/G | 2.63E-17 | intronic | - | - |
| rs17194490 | 3 | 2547786 | G/T | 4.87E-11 | intronic | - | - |
| rs1702294 | 1 | 98501984 | C/T | 2.79E-17 | ncRNA_intronic | - | - |
| rs16867576 | 5 | 88746331 | A/G | 1.36E-08 | ncRNA_intronic | - | - |
| rs1501357 | 5 | 45364875 | C/T | 1.24E-08 | intronic | - | - |
| rs1498232 | 1 | 30433951 | C/T | 1.28E-09 | intergenic | - | - |
| rs140505938 | 1 | 150031490 | C/T | 9.34E-10 | intergenic | - | - |
| rs1378559 | 23 | 21380266 | C/T | 1.68E-11 | intergenic | *CNKSR2* | protein_coding |
| rs1353545 | 3 | 60287845 | C/G | 4.26E-08 | intronic | *FHIT* | protein_coding |
| rs133047 | 22 | 41027819 | C/T | 9.73E-10 | intronic | *MKL1* | protein_coding |
| rs13240464 | 7 | 110898915 | C/T | 6.16E-13 | intronic | - | - |
| rs13170232 | 5 | 152899532 | A/T | 3.82E-08 | intronic | *GRIA1* | protein_coding |
| rs12903146 | 15 | 61854663 | A/G | 1.04E-09 | ncRNA_intronic | - | - |
| rs12891702 | 14 | 72448232 | G/T | 2.20E-08 | intronic | - | - |
| rs12887734 | 14 | 104046834 | G/T | 1.17E-13 | intronic | *CKB* | protein_coding |
| rs12826178 | 12 | 57622371 | G/T | 5.30E-12 | upstream:downstream | *LRP1* | protein_coding |
| rs12704290 | 7 | 86427626 | A/G | 1.04E-10 | intronic | *GRM3* | protein_coding |
| rs12691307 | 16 | 29939877 | A/G | 1.30E-10 | ncRNA_exonic | *ASPHD1* | protein_coding |
| rs1233578 | 6 | 28712247 | A/G | 3.86E-32 | intergenic | - | - |
| rs12325245 | 16 | 58681393 | A/T | 1.15E-08 | intergenic | *CNOT1* | protein_coding |
| rs12129573 | 1 | 73768366 | A/C | 2.35E-10 | intergenic | - | - |
| rs11874716 | 18 | 52750688 | G/T | 4.13E-11 | intergenic | - | - |
| rs11740474 | 5 | 153680747 | A/T | 3.94E-08 | intronic | *GALNT10* | protein_coding |
| rs117074560 | 6 | 96459651 | C/T | 1.66E-08 | intergenic | - | - |
| rs11693094 | 2 | 185601420 | C/T | 7.13E-12 | intronic | - | - |
| rs11685299 | 2 | 225391296 | A/C | 1.11E-08 | intronic | *CUL3* | protein_coding |
| rs11682175 | 2 | 57987593 | C/T | 2.54E-12 | ncRNA_intronic | - | - |
| rs11210892 | 1 | 44100084 | A/G | 4.97E-10 | intergenic | *PTPRF* | protein_coding |
| rs11191419 | 10 | 104612335 | A/T | 9.24E-18 | intergenic | - | - |
| rs11139497 | 9 | 84739941 | A/T | 3.09E-09 | intergenic | - | - |
| rs111294930 | 5 | 152177121 | A/G | 1.31E-09 | ncRNA_intronic | - | - |
| rs1106568 | 4 | 176861301 | A/G | 1.15E-08 | intronic | *GPM6A* | protein_coding |
| rs11027857 | 11 | 24403620 | A/G | 3.21E-09 | intergenic | - | - |
| rs10946817 | 6 | 26364056 | C/T | 2.42E-12 | intergenic | - | - |
| rs10803138 | 1 | 243555219 | A/G | 1.79E-08 | intronic | - | - |
| rs10791097 | 11 | 130718630 | G/T | 2.88E-12 | ncRNA_intronic | - | - |
| rs10520163 | 4 | 170626552 | C/T | 1.02E-08 | intronic | *C4orf27* | protein_coding |
| rs10503253 | 8 | 4180844 | A/C | 2.69E-08 | intronic | *CSMD1* | protein_coding |
| rs10043984 | 5 | 137712121 | C/T | 2.18E-08 | intronic | *GFRA3* | protein_coding |

**Table S9.** Genomic risk loci and independent variants (LD $r^2$<0.1) identified in *CompGaming* GWAS.

| rsID | Chromosome | Position | Alleles | p | Consequence | Gene Symbol | Region Biotype |
|---|---|---|---|---|---|---|---|
| rs10119540 | 9 | 71582658 | C/T | 4.65E-08 | intronic | *PIP5K1B* | protein_coding |
| rs11223780 | 11 | 134261150 | A/G | 1.86E-09 | UTR5 | *GLB1L2* | protein_coding |
| rs112365747 | 1 | 27200284 | A/G | 3.29E-10 | intergenic | - | - |
| rs11283226 | 16 | 31114916 | C/G | 1.13E-12 | intergenic | - | - |
| rs116518031 | 7 | 32894682 | C/T | 2.55E-09 | intronic | *LSM5* | protein_coding |
| rs116966595 | 9 | 95526685 | A/G | 4.90E-09 | intronic | *IARS* | protein_coding |
| rs11743441 | 5 | 88065637 | G/T | 3.31E-10 | intronic | - | - |
| rs1190736 | 23 | 136113464 | A/C | 1.37E-08 | exonic | *ARHGEF6* | protein_coding |
| rs13026471 | 2 | 137564022 | C/T | 1.29E-14 | intronic | - | - |
| rs13198347 | 5 | 12992383 | C/T | 3.23E-10 | ncRNA_intronic | - | - |
| rs13262595 | 8 | 143316970 | A/G | 5.21E-15 | intronic | - | - |
| rs1406069 | 17 | 44225886 | A/G | 2.16E-17 | intronic | - | - |
| rs1441188 | 1 | 163790947 | C/G | 3.09E-08 | intergenic | - | - |
| rs16838536 | 2 | 155889400 | C/T | 3.39E-11 | intergenic | - | - |
| rs17184707 | 2 | 166183577 | C/T | 7.70E-10 | intronic | *SCN2A* | protein_coding |
| rs17790917 | 3 | 70571159 | A/G | 1.92E-08 | intergenic | - | - |
| rs2014653 | 20 | 50788476 | A/G | 2.09E-10 | intronic | - | - |
| rs2352741 | 16 | 10172705 | C/T | 3.05E-09 | intronic | - | - |
| rs2734837 | 11 | 113286829 | C/T | 1.13E-17 | intronic | *DRD2* | - |
| rs4017425 | 3 | 44028764 | C/T | 2.40E-09 | intergenic | - | - |
| rs4113587 | 6 | 128311640 | G/T | 3.08E-08 | intronic | *PTPRK* | protein_coding |
| rs4146338 | 3 | 117493964 | C/T | 1.44E-08 | ncRNA_intronic | *LSAMP* | protein_coding |
| rs4936271 | 11 | 113318705 | C/T | 3.42E-10 | intronic | *DRD2* | - |
| rs55938136 | 17 | 43798360 | A/G | 1.09E-15 | ncRNA_intronic | - | - |
| rs6021740 | 20 | 50761157 | C/T | 5.04E-09 | intronic | - | - |
| rs61554893 | 8 | 141610141 | A/AAT | 3.04E-08 | intronic | *AGO2* | protein_coding |
| rs62512616 | 8 | 143297663 | C/T | 3.01E-08 | intronic | *TSNARE1* | - |
| rs62584692 | 23 | 22548983 | A/G | 3.59E-08 | ncRNA_intronic | - | - |
| rs6627903 | 23 | 23214048 | C/T | 3.45E-08 | ncRNA_intronic | - | - |
| rs7103627 | 11 | 66648083 | A/G | 3.59E-11 | intronic | *CTSF* | protein_coding |
| rs72836318 | 17 | 44121579 | C/T | 1.39E-08 | intronic | - | - |
| rs75620859 | 2 | 175294443 | C/T | 4.15E-08 | downstream | *SCRN3* | protein_coding |
| rs7571383 | 2 | 200228383 | C/T | 7.18E-11 | intronic | - | - |

**Table S10.** Differential gene set enrichment between schizophrenia and *CompGaming* surviving Bonferroni correction ($p<4.69x10^{-6}$).

| Enrichment | N genes | Schizophrenia | | *CompGaming* | | Difference | |
|---|---|---|---|---|---|---|---|
| | | beta | p | beta | p | z | p |
| MAPK11 Targets (Turjanski) | 3 | 0.402 | 0.179 | 1.87 | 3.79E-05 | -2.28 | 1.02E-13 |
| Reactome Down Syndrome Cell Adhesion Molecule Interactions | 7 | -0.144 | 0.663 | 1.24 | 6.00E-05 | -2.96 | 2.17E-12 |
| GO Positive Regulation of Rho Protein Signal Transduction | 9 | -0.748 | 0.991 | 0.632 | 0.018 | -3.15 | 2.50E-11 |
| Targets of PAX3-FOXO1 Fusion and PAX3 (Begum) | 5 | -0.123 | 0.608 | 1.17 | 0.004 | -2.05 | 4.90E-11 |
| Gamma Secretase Sensitivity Down (Palomero) | 3 | -0.355 | 0.729 | 0.923 | 0.049 | -1.59 | 8.03E-11 |
| GO Dopamine Transport | 7 | -0.586 | 0.959 | 0.654 | 0.014 | -2.75 | 2.74E-10 |
| Reactome DCC Mediated Attactive Signaling | 9 | -0.288 | 0.844 | 0.907 | 9.83E-04 | -2.92 | 1.12E-09 |
| GO Positive Regulation of Cytosolic Calcium Ion Concentration Involved in Phospholipase-C Activating G-Protein Coupled Signaling Pathway | 9 | -0.174 | 0.707 | 0.977 | 7.83E-04 | -2.59 | 4.22E-09 |
| GO Filamen Binding | 9 | -0.382 | 0.915 | 0.751 | 0.003 | -2.91 | 7.16E-09 |
| Mesothelioma Survival Down (Lopez) | 8 | -0.512 | 0.938 | 0.615 | 0.026 | -2.45 | 8.53E-09 |
| MAPK14 Targets (Turjanski) | 7 | 0.046 | 0.446 | 1.16 | 8.29E-05 | -2.44 | 1.25E-08 |
| PID p38-Gamma and p38-delta Pathway | 7 | -0.328 | 0.854 | 0.770 | 0.010 | -2.41 | 1.96E-08 |
| GO Structural Constituent of Nuclear Pore | 9 | -0.547 | 0.957 | 0.505 | 0.033 | -2.50 | 7.04E-08 |
| TCGA Glioblastoma Mutated | 5 | -0.236 | 0.742 | 0.779 | 0.015 | -1.99 | 1.90E-07 |
| Nuclear receptors whose expression correlated with that of aromatase (CYP19A1; Miki) | 4 | -0.032 | 0.532 | 0.946 | 0.012 | -1.68 | 4.92E-07 |
| GO G-Protein Coupled Glutamate Receptor Signaling Pathway | 9 | 0.277 | 0.188 | 1.24 | 2.08E-05 | -2.21 | 7.22E-07 |
| GO Specification of Organ Identity | 9 | -0.236 | 0.784 | 0.718 | 0.008 | -2.25 | 9.04E-07 |
| Liver Cancer Early Recurrence (Iizuka) | 7 | -0.198 | 0.720 | 0.724 | 0.015 | -1.94 | 1.98E-06 |
| GO Neurotransmitter Transporter Activity | 17 | -0.465 | 0.985 | 0.449 | 0.016 | -3.06 | 2.40E-06 |
| SKI Targets Down (Le) | 5 | -0.139 | 0.625 | 0.771 | 0.027 | -1.53 | 2.64E-06 |
| GO Negative Regulation of Regulated Secretory Pathway | 14 | -0.347 | 0.928 | 0.561 | 0.006 | -2.80 | 2.77E-06 |
| GO Substrate-Independent Telencephalic Tangential Migration | 13 | -0.054 | 0.568 | 0.837 | 0.004 | -2.00 | 4.12E-06 |

**Table S11.** Gene set enrichments surviving Bonferroni correction for schizophrenia ($p<4.69x10^{-06}$).

| Enrichment | N genes | Schizophrenia | | *CompGaming* | | Difference | |
|---|---|---|---|---|---|---|---|
| | | beta | p | beta | p | z | p |
| GO Neuron Projection | 939 | 0.182 | 7.36E-08 | 0.156 | 1.43E-06 | 0.542 | 0.552 |
| GO Postsynapse | 375 | 0.286 | 1.33E-07 | 0.233 | 6.66E-06 | 0.688 | 0.605 |
| GO Neuron Spine | 121 | 0.468 | 5.64E-07 | 0.244 | 0.003 | 1.70 | 0.869 |
| GO Neuron Part | 1256 | 0.147 | 5.76E-07 | 0.113 | 4.93E-05 | 0.811 | 0.568 |
| Genes with promoters bound by FOXP3 and which are down-regulated only in mature regulatory CD4+ T-lymphocytes (Zheng) | 36 | 0.931 | 6.48E-07 | 0.26 | 0.075 | 2.54 | 1 |
| GO High Voltage Gated Calcium Channel Activity | 11 | 1.45 | 1.83E-06 | 0.25 | 0.208 | 2.74 | 1 |

**Table S12.** Gene set enrichments surviving Bonferroni correction for *CompGaming* ($p<4.69x10^{-06}$).

| **Enrichment** | **N genes** | **Schizophrenia** | | ***CompGaming*** | | **Difference** | |
|---|---|---|---|---|---|---|---|
| | | **beta** | **p** | **beta** | **p** | **z** | **p** |
| GO Positive Regulation of Long Term Synaptic Potentiation | 14 | 1.30 | 1.82E-05 | 1.63 | 2.69E-09 | -0.786 | 0.049 |
| GO Regulation of Long Term Synaptic Potentiation | 19 | 0.838 | 7.75E-04 | 1.25 | 5.95E-08 | -1.16 | 0.020 |
| GO Synapse | 749 | 0.165 | 1.35E-05 | 0.185 | 3.45E-07 | -0.369 | 0.460 |
| GO Neuron Projection | 939 | 0.182 | 7.36E-08 | 0.156 | 1.43E-06 | 0.542 | 0.552 |
| GO Modulation of Synaptic Transmission | 296 | 0.248 | 1.41E-05 | 0.257 | 2.84E-06 | -0.110 | 0.482 |
| GO Regulation of Synaptic Structure or Activity | 232 | 0.277 | 2.92E-05 | 0.298 | 2.89E-06 | -0.221 | 0.458 |
| GO Synapse Part | 608 | 0.191 | 4.41E-06 | 0.184 | 3.75E-06 | 0.118 | 0.514 |

**Table S13.** Linkage disequilibrium score regression (rg) results for 1,045 UK Biobank traits demonstrating nominally significant genetic correlations in schizophrenia and/or *CompGaming*. Traits are listed in ascending order by most significant difference between schizophrenia and *CompGaming*.

| Field ID | Trait | Schizophrenia | | *CompGaming* | | Difference | |
|---|---|---|---|---|---|---|---|
| | | rg | p | rg | p | z | p |
| 398 | Number of correct matches in round | -0.39 | 4.33E-27 | 0.435 | 6.69E-25 | -14.8383 | 1.63E-07 |
| 100160 | Low calorie drink intake | -0.3475 | 0.0072 | 0.3829 | 0.0021 | -4.0755 | 3.06E-06 |
| 1150_3 | Usual side of head for mobile phone use: Equally left and right | -0.2716 | 4.51E-09 | 0.4502 | 2.49E-17 | -10.2345 | 3.93E-06 |
| 22601_11363057 | Job coding: ict or it manager, systems/data processing manager, telecom manager | -0.1826 | 0.2038 | 0.5209 | 0.0409 | -2.4049 | 6.64E-06 |
| 2664_5 | Reason for reducing amount of alcohol drunk: Other reason | -0.4336 | 6.35E-27 | 0.2562 | 1.09E-11 | -12.4833 | 9.74E-06 |
| 6157_100 | Why stopped smoking: None of the above | -0.3393 | 9.42E-11 | 0.3039 | 6.46E-08 | -8.3708 | 3.41E-05 |
| 20128 | Number of fluid intelligence questions attempted within time limit | -0.3509 | 6.34E-27 | 0.2821 | 4.43E-17 | -13.501 | 4.44E-05 |
| 22617_1136 | Job SOC coding: Information and communication technology managers | -0.1563 | 0.2307 | 0.4621 | 0.017 | -2.6484 | 6.44E-05 |
| M18 | Diagnoses - main ICD10: M18 Arthrosis of first carpometacarpal joint | -0.3241 | 0.063 | 0.2709 | 0.0267 | -2.7951 | 0.000115 |
| 100250 | Instant coffee intake | -0.3155 | 0.0004 | 0.2769 | 0.0019 | -4.6961 | 0.000122 |
| 22601_71113328 | Job coding: sales assistant, retail/shop assistant, counter assistant, sub post office assistant, takeaway food worker | -0.2111 | 0.103 | 0.3706 | 0.0137 | -2.932 | 0.000158 |
| I24 | Diagnoses - main ICD10: I24 Other acute ischaemic heart diseases | -0.3211 | 0.0151 | 0.2523 | 0.1177 | -2.7494 | 0.000193 |
| 22617_7111 | Job SOC coding: Sales and retail assistants | -0.1896 | 0.0862 | 0.359 | 0.0089 | -3.1141 | 0.000341 |
| 3160 | Weight, manual entry | -0.1352 | 0.1795 | 0.4127 | 0.0037 | -3.1459 | 0.000347 |
| 20003_1140909708 | Treatment/medication code: furosemide | -0.2176 | 0.0179 | 0.3212 | 0.0004 | -4.1751 | 0.000425 |
| 5779 | FI11 : antonym | -0.2627 | 0.0024 | 0.2668 | 0.0041 | -4.1712 | 0.000522 |
| 5251 | 6mm index of best keratometry results (right) | -0.3086 | 0.0114 | 0.2173 | 0.0807 | -3.0183 | 0.000565 |
| 1468_5 | Cereal type: Other (e.g. Cornflakes, Frosties) | -0.2529 | 7.33E-20 | 0.2706 | 1.45E-19 | -12.8438 | 0.000595 |
| 22604_3 | Work hours - lumped category: 30 to 40 hours | -0.158 | 0.2568 | 0.3626 | 0.0094 | -2.6379 | 0.000634 |
| 6162_1 | Types of transport used (excluding work): Car/motor vehicle | -0.3602 | 2.33E-26 | 0.1583 | 9.15E-06 | -10.532 | 0.000663 |
| T81 | Diagnoses - main ICD10: T81 Complications of procedures, not elsewhere classified | -0.1572 | 0.0523 | 0.3516 | 2.97E-05 | -4.3548 | 0.000816 |
| 1080 | Time spent using computer | -0.0569 | 0.0126 | 0.4494 | 1.4E-132 | -17.3178 | 0.00086 |
| M25 | Diagnoses - main ICD10: M25 Other joint disorders, not elsewhere classified | -0.1726 | 0.0373 | 0.327 | 0.0001 | -4.1749 | 0.00099 |
| 6153_3 | Medication for cholesterol, blood pressure, diabetes, or take exogenous hormones: Insulin | -0.2069 | 0.006 | 0.2921 | 0.0002 | -4.5904 | 0.001003 |
| 777 | Frequency of travelling from home to job workplace | -0.2259 | 2.61E-06 | 0.2698 | 1.2E-07 | -7.0709 | 0.001074 |
| M13_JOINTOTH | Other specific joint derangements/joint disorders | -0.1785 | 0.0244 | 0.3146 | 0.0002 | -4.2525 | 0.001133 |
| 4294_1 | Final attempt correct: yes | -0.2759 | 1.03E-05 | 0.2055 | 0.0006 | -5.574 | 0.001439 |
| 20003_1140860840 | Treatment/medication code: nitrolingual 400micrograms spray | -0.2223 | 0.0643 | 0.2554 | 0.043 | -2.7409 | 0.00155 |
| I27 | Diagnoses - main ICD10: I27 Other pulmonary heart diseases | -0.1565 | 0.2844 | 0.317 | 0.041 | -2.2215 | 0.001686 |
| 22617_2131 | Job SOC coding: IT strategy and planning professionals | -0.0634 | 0.5653 | 0.4075 | 0.0072 | -2.5107 | 0.001776 |
| 1448_4 | Bread type: Other type of bread | -0.1156 | 0.1408 | 0.3469 | 0.0005 | -3.6561 | 0.002095 |
| 100170 | Fizzy drink intake | -0.081 | 0.3742 | 0.371 | 0.0036 | -2.8845 | 0.002567 |
| 22601_21313058 | Job coding: it strategy or planning professional, computer consultant, software consultant | -0.0554 | 0.5648 | 0.3915 | 0.0039 | -2.6871 | 0.00283 |

| | | | | | | | |
|---|---|---|---|---|---|---|---|
| 3751 | Chest pain or discomfort when walking uphill or hurrying | -0.1165 | 0.0972 | 0.3244 | 3.89E-07 | -4.641 | 0.003169 |
| M13_ADHCAPSULITIS | Adhesive capsulitis of shoulder | -0.1529 | 0.1192 | 0.2868 | 0.0025 | -3.2215 | 0.003241 |
| 20003_1140923350 | Treatment/medication code: co-dydramol | -0.0809 | 0.4784 | 0.3575 | 0.0099 | -2.4431 | 0.003321 |
| 1598 | Average weekly spirits intake | -0.1723 | 1.12E-07 | 0.2591 | 2.54E-13 | -8.977 | 0.003782 |
| I9_UAP | Unstable angina pectoris | -0.0892 | 0.118 | 0.3421 | 2.81E-08 | -5.1391 | 0.003789 |
| E4_DM2NOCOMP | Type 2 diabetes without complications | -0.0972 | 0.3578 | 0.3322 | 0.0226 | -2.3863 | 0.003924 |
| 22601_21323059 | Job coding: software professional, analyst-programmer, computer programmer, software analyst or engineer, systems designer or programmer | -0.0177 | 0.81 | 0.4104 | 2.57E-05 | -3.5061 | 0.004019 |
| 20016 | Fluid intelligence score | -0.2366 | 1.03E-20 | 0.1913 | 4.15E-13 | -11.6801 | 0.004033 |
| 4440 | Average monthly spirits intake | -0.1909 | 0.0056 | 0.2311 | 0.0011 | -4.2717 | 0.004491 |
| 20002_1081 | Non-cancer illness code, self-reported: stroke | -0.0605 | 0.4919 | 0.3604 | 6.95E-05 | -3.3325 | 0.004581 |
| 6138_5 | Qualifications: NVQ or HND or HNC or equivalent | -0.0884 | 0.032 | 0.3275 | 7.13E-14 | -6.9164 | 0.005012 |
| 6143_1 | Transport type for commuting to job workplace: Car/motor vehicle | -0.1823 | 5.75E-06 | 0.2325 | 3.01E-08 | -7.1347 | 0.005112 |
| 20111_101 | Illnesses of siblings: None of the above (group 2) | -0.4783 | 8.74E-15 | -0.0711 | 0.1618 | -5.095 | 0.005849 |
| M13_ARTHROSIS_OTH | Other arthrosis | -0.1918 | 0.0059 | 0.2137 | 0.0025 | -4.0815 | 0.006027 |
| 48 | Waist circumference | -0.0988 | 1.75E-08 | 0.3016 | 1.31E-51 | -15.0666 | 0.006587 |
| 100360 | Decaffeinated coffee | -0.2848 | 0.0158 | 0.1151 | 0.2947 | -2.481 | 0.006645 |
| I9_PULMOTHHD | Other pulmonary heart/vessel disease | -0.1567 | 0.1826 | 0.24 | 0.0283 | -2.4688 | 0.007023 |
| 22617_2132 | Job SOC coding: Software professionals | 0.0071 | 0.9238 | 0.4013 | 2.23E-05 | -3.2872 | 0.007331 |
| 21001 | Body mass index (BMI) | -0.1063 | 3.32E-09 | 0.2864 | 3.26E-47 | -14.6755 | 0.007522 |
| 23104 | Body mass index (BMI) | -0.1055 | 5.72E-09 | 0.2859 | 1.28E-46 | -14.5501 | 0.00769 |
| 23112 | Leg fat mass (right) | -0.0966 | 1.58E-07 | 0.2918 | 9.41E-46 | -14.0618 | 0.008093 |
| 23116 | Leg fat mass (left) | -0.0974 | 9.07E-08 | 0.2901 | 1.3E-45 | -14.1355 | 0.008217 |
| 20118_12 | Home area population density - urban or rural: Scotland - Other Urban Area | -0.1702 | 0.013 | 0.2168 | 0.0037 | -3.8184 | 0.008287 |
| 1389 | Pork intake | -0.1836 | 1.05E-08 | 0.2007 | 1.84E-09 | -8.2958 | 0.008672 |
| 23124 | Arm fat mass (left) | -0.097 | 6.1E-08 | 0.287 | 3.84E-45 | -14.1489 | 0.008716 |
| 23100 | Whole body fat mass | -0.0953 | 1.3E-07 | 0.2878 | 1.7E-45 | -14.0859 | 0.008849 |
| 20003_1140883066 | Treatment/medication code: insulin product | -0.1421 | 0.016 | 0.2404 | 7.51E-05 | -4.5186 | 0.008938 |
| I20 | Diagnoses - main ICD10: I20 Angina pectoris | -0.0865 | 0.0649 | 0.2956 | 2.11E-10 | -5.7855 | 0.008998 |
| 1130 | Hands-free device/speakerphone use with mobile phone in last 3 month | -0.1902 | 2.88E-06 | 0.1893 | 1.9E-06 | -6.6832 | 0.009397 |
| 3393 | Hearing aid user | -0.1298 | 0.0062 | 0.2493 | 1.06E-06 | -5.4391 | 0.009459 |
| 20003_1141168318 | Treatment/medication code: clopidogrel | -0.0132 | 0.8611 | 0.3654 | 0.0003 | -3.0047 | 0.009538 |
| 23120 | Arm fat mass (right) | -0.0908 | 4.76E-07 | 0.2856 | 7.86E-45 | -13.8734 | 0.009892 |
| K81 | Diagnoses - main ICD10: K81 Cholecystitis | -0.0498 | 0.5728 | 0.3266 | 0.0063 | -2.5343 | 0.009892 |
| 22617_3513 | Job SOC coding: Ship and hovercraft officers | -0.1317 | 0.1189 | 0.2432 | 0.0253 | -2.723 | 0.01014 |
| I9_PAD | Peripheral artery disease | -0.0057 | 0.9444 | 0.3678 | 0.0074 | -2.3435 | 0.010376 |
| 1100 | Drive faster than motorway speed limit | -0.1768 | 2.19E-11 | 0.1959 | 7.56E-11 | -9.3089 | 0.010513 |
| 23111 | Leg fat percentage (right) | -0.0859 | 6.98E-06 | 0.2865 | 1.08E-47 | -13.5719 | 0.010565 |
| 23115 | Leg fat percentage (left) | -0.0863 | 5.24E-06 | 0.2835 | 1.54E-46 | -13.4759 | 0.011024 |
| 49 | Hip circumference | -0.0998 | 2.95E-07 | 0.2698 | 4.22E-41 | -13.1978 | 0.01106 |
| 23123 | Arm fat percentage (left) | -0.0872 | 1.15E-06 | 0.2804 | 6.44E-47 | -13.8874 | 0.011425 |
| 23099 | Body fat percentage | -0.0836 | 5.15E-06 | 0.2835 | 1.29E-46 | -13.6156 | 0.011518 |
| 23128 | Trunk fat mass | -0.0886 | 6.6E-07 | 0.2784 | 3.11E-43 | -13.6312 | 0.011537 |
| PRIM_KNEEARTHROSIS | Primary gonarthrosis, bilateral | -0.1112 | 0.1225 | 0.2557 | 0.0044 | -3.1877 | 0.011555 |
| 6150_3 | Vascular/heart problems diagnosed by doctor: Stroke | -0.0567 | 0.5108 | 0.3091 | 7.09E-05 | -3.1523 | 0.011763 |
| 1070 | Time spent watching television (TV) | -0.1548 | 1.96E-12 | 0.2101 | 4.66E-22 | -11.7818 | 0.011935 |
| M75 | Diagnoses - main ICD10: M75 Shoulder lesions | -0.1703 | 0.0011 | 0.1919 | 0.0001 | -5.0207 | 0.012465 |

| | | | | | | | |
|---|---|---|---|---|---|---|---|
| M13_SHOULDER | Shoulder lesions | -0.1701 | 0.0011 | 0.1914 | 0.0001 | -5.011 | 0.012605 |
| 22601_82112603 | Job coding: heavy goods vehicle (hgv) driver, lorry or truck driver, tanker driver, haulage driver | -0.1453 | 0.0756 | 0.2136 | 0.0086 | -3.1119 | 0.013139 |
| M19 | Diagnoses - main ICD10: M19 Other arthrosis | -0.1774 | 0.0354 | 0.1812 | 0.0297 | -3.024 | 0.013202 |
| 20002_1220 | Non-cancer illness code, self-reported: diabetes | -0.1111 | 0.0005 | 0.2454 | 2.92E-16 | -8.1682 | 0.01365 |
| 20003_1141152998 | Treatment/medication code: irbesartan | -0.0377 | 0.6382 | 0.3178 | 0.0046 | -2.5792 | 0.013867 |
| 23119 | Arm fat percentage (right) | -0.0783 | 1.46E-05 | 0.277 | 1.54E-46 | -13.4281 | 0.013911 |
| 1359 | Poultry intake | -0.1454 | 2.96E-05 | 0.2091 | 3.8E-11 | -7.5415 | 0.014088 |
| 23127 | Trunk fat percentage | -0.0792 | 1.46E-05 | 0.2752 | 1.26E-42 | -13.0377 | 0.01411 |
| 1110 | Length of mobile phone use | -0.1377 | 4.38E-07 | 0.2163 | 2.05E-16 | -9.3385 | 0.0142 |
| 6141_6 | How are people in household related to participant: Grandchild | -0.0453 | 0.5637 | 0.3077 | 6.28E-05 | -3.2123 | 0.014425 |
| 20110_101 | Illnesses of mother: None of the above (group 2) | -0.3079 | 5.5E-07 | 0.0447 | 0.4545 | -4.1138 | 0.014516 |
| 3496 | Wants to stop smoking | -0.2616 | 0.0002 | 0.0898 | 0.2304 | -3.4049 | 0.014791 |
| OTHER_ILD_CVD_COMORB | Other ILD-related CVD-co-morbidities | -0.123 | 0.1248 | 0.2277 | 0.001 | -3.3131 | 0.014954 |
| 21002 | Weight | -0.0954 | 9.12E-08 | 0.2546 | 5.89E-36 | -12.932 | 0.015119 |
| 23098 | Weight | -0.0949 | 1.25E-07 | 0.2539 | 1.12E-35 | -12.8208 | 0.015404 |
| E4_DM2 | Type 2 diabetes | -0.0899 | 0.2387 | 0.2565 | 0.0024 | -3.0406 | 0.015989 |
| 20126_0 | Bipolar and major depression status: No Bipolar or Depression | -0.3829 | 1.15E-20 | -0.0378 | 0.3384 | -6.054 | 0.016314 |
| 22617_8211 | Job SOC coding: Heavy goods vehicle drivers | -0.1405 | 0.0881 | 0.2045 | 0.0128 | -2.966 | 0.016339 |
| 20433 | Age at first episode of depression | -0.2952 | 4.51E-06 | 0.0482 | 0.4087 | -3.9531 | 0.016747 |
| 110001 | Invitation to complete online 24-hour recall dietary questionnaire, acceptance | -0.1666 | 1.79E-06 | 0.1757 | 1.11E-06 | -6.8171 | 0.017033 |
| 22617_9219 | Job SOC coding: Elementary office occupations n.e.c. | -0.0584 | 0.5245 | 0.2838 | 0.0043 | -2.5304 | 0.017059 |
| 20003_1140851812 | Treatment/medication code: gtn 400micrograms spray | -0.0802 | 0.2974 | 0.2618 | 0.0061 | -2.7878 | 0.017112 |
| 20003_1140910766 | Treatment/medication code: nicorandil | -0.0146 | 0.8347 | 0.3265 | 0.0007 | -2.8637 | 0.017349 |
| E11 | Diagnoses - main ICD10: E11 Non-insulin-dependent diabetes mellitus | -0.0866 | 0.2491 | 0.2541 | 0.002 | -3.0559 | 0.017456 |
| I73 | Diagnoses - main ICD10: I73 Other peripheral vascular diseases | 0.0198 | 0.8546 | 0.3605 | 0.032 | -1.7035 | 0.017456 |
| 20003_1140874744 | Treatment/medication code: gliclazide | -0.0942 | 0.0512 | 0.2463 | 3.76E-07 | -4.9746 | 0.017509 |
| 20455 | Age when last took cannabis | -0.0567 | 0.5507 | 0.2834 | 0.032 | -2.0884 | 0.017617 |
| 2443 | Diabetes diagnosed by doctor | -0.0991 | 0.0008 | 0.2407 | 9.41E-18 | -8.3248 | 0.017697 |
| 22617_5242 | Job SOC coding: Telecommunications engineers | 0.004 | 0.9699 | 0.3437 | 0.0303 | -1.7836 | 0.017724 |
| M13_MUSCLE | Disorders of muscles | -0.1062 | 0.2444 | 0.2331 | 0.0358 | -2.3605 | 0.017833 |
| 20003_1140884600 | Treatment/medication code: metformin | -0.1125 | 0.001 | 0.2267 | 2.94E-13 | -7.3378 | 0.01786 |
| 4429 | Average monthly beer plus cider intake | -0.1211 | 0.2102 | 0.2173 | 0.0327 | -2.4114 | 0.018079 |
| 20111_2 | Illnesses of siblings: Stroke | -0.107 | 0.2004 | 0.2292 | 0.0134 | -2.6947 | 0.018693 |
| 20152 | Reproduciblity of spirometry measurement using ERS/ATS criteria | -0.144 | 0.0015 | 0.1907 | 3.83E-05 | -5.1671 | 0.019122 |
| 1448_1 | Bread type: White | -0.0807 | 0.0025 | 0.2522 | 2.07E-21 | -8.8494 | 0.019647 |
| 22601_52422945 | Job coding: telecommunication engineer, telephone engineer, telephone technician | -0.0076 | 0.9414 | 0.3253 | 0.034 | -1.7968 | 0.019647 |
| 6156_100 | Manic/hyper symptoms: None of the above | -0.3242 | 0.0002 | 0.0082 | 0.9165 | -2.8646 | 0.019795 |
| 6164_100 | Types of physical activity in last 4 weeks: None of the above | 0.0304 | 0.397 | 0.3625 | 4.86E-30 | -6.9247 | 0.019885 |
| 20003_1140864752 | Treatment/medication code: lansoprazole | -0.0728 | 0.0987 | 0.259 | 2.95E-08 | -5.1657 | 0.019974 |
| 20527 | Been involved in combat or exposed to war-zone | -0.0119 | 0.874 | 0.3175 | 4.67E-06 | -3.2304 | 0.020704 |
| 20111_1 | Illnesses of siblings: Heart disease | -0.0598 | 0.1395 | 0.2695 | 3.43E-09 | -5.3994 | 0.020735 |
| 1090 | Time spent driving | -0.2504 | 3.11E-14 | 0.0781 | 0.0139 | -7.168 | 0.020983 |

| | | | | | | | |
|---|---|---|---|---|---|---|---|
| 6141_1 | How are people in household related to participant: Husband, wife or partner | -0.2983 | 1.13E-16 | 0.0299 | 0.3854 | -6.5913 | 0.021077 |
| K11_GASTRODUOULC | Gastroduodenal ulcer | -0.0335 | 0.7635 | 0.2942 | 0.0408 | -1.8007 | 0.021234 |
| 20002_1223 | Non-cancer illness code, self-reported: type 2 diabetes | -0.0721 | 0.2804 | 0.2552 | 0.0009 | -3.2203 | 0.021361 |
| 2844 | Had other major operations | -0.0649 | 0.1172 | 0.2614 | 2.97E-09 | -5.3945 | 0.02168 |
| 22617_4150 | Job SOC coding: General office assistants/clerks | -0.0916 | 0.2912 | 0.2347 | 0.033 | -2.3274 | 0.02168 |
| 20003_1140874420 | Treatment/medication code: quinine | -0.0973 | 0.1333 | 0.2287 | 0.0005 | -3.53 | 0.021776 |
| M13_GANGLION | Ganglion | -0.1952 | 0.0136 | 0.1302 | 0.0996 | -2.9089 | 0.02197 |
| 20003_1140883568 | Treatment/medication code: oxybutynin | -0.1141 | 0.1416 | 0.2106 | 0.0354 | -2.5636 | 0.022198 |
| IV_ENDOCRIN_NUTRIT | Endocrine, nutritional and metabolic diseases | -0.0884 | 0.2725 | 0.2362 | 0.0064 | -2.7437 | 0.022231 |
| 20003_1140923346 | Treatment/medication code: co-codamol | -0.0512 | 0.2576 | 0.2733 | 9.34E-09 | -4.9435 | 0.022264 |
| 2207 | Wears glasses or contact lenses | -0.1694 | 2.42E-06 | 0.154 | 6.36E-05 | -6.1435 | 0.022628 |
| 22601_92193312 | Job coding: other general office clerical tasks including: office junior, office worker, photocopy/print room operator, office machinist | -0.0496 | 0.5828 | 0.2736 | 0.0044 | -2.4536 | 0.022694 |
| M13_SOFTTISSUEOTH | Other soft tissue disorders, not elsewhere classified | -0.108 | 0.1149 | 0.2146 | 0.0028 | -3.2533 | 0.022895 |
| M79 | Diagnoses - main ICD10: M79 Other soft tissue disorders, not elsewhere classified | -0.108 | 0.1149 | 0.2146 | 0.0028 | -3.2533 | 0.022895 |
| K11_GALLBILPANC | Disorders of gallbladder, biliary tract and pancreas | -0.0256 | 0.5292 | 0.2958 | 2.21E-12 | -5.4887 | 0.023301 |
| 3839 | Number of spontaneous miscarriages | -0.1208 | 0.1698 | 0.1995 | 0.0236 | -2.5708 | 0.023679 |
| BRONCHITIS | Bronchitis | -0.0271 | 0.8051 | 0.2897 | 0.0247 | -1.8687 | 0.024915 |
| 6148_1 | Eye problems/disorders: Diabetes related eye disease | -0.091 | 0.1745 | 0.2251 | 0.0006 | -3.3711 | 0.025169 |
| I26 | Diagnoses - main ICD10: I26 Pulmonary embolism | -0.1146 | 0.1709 | 0.1999 | 0.0053 | -2.8553 | 0.025757 |
| 100260 | Added milk to instant coffee | -0.2924 | 0.0372 | 0.0212 | 0.8379 | -1.7981 | 0.026093 |
| M13_ENTESOPATHYOTH | Other enthesopathies | -0.2167 | 0.0245 | 0.0946 | 0.3503 | -2.2272 | 0.026967 |
| M77 | Diagnoses - main ICD10: M77 Other enthesopathies | -0.2158 | 0.0231 | 0.0953 | 0.3403 | -2.2567 | 0.027045 |
| 20107_101 | Illnesses of father: None of the above (group 2) | -0.249 | 6.82E-08 | 0.0614 | 0.2165 | -4.5743 | 0.027316 |
| 20491 | Someone to take to doctor when needed as a child | -0.3205 | 2.24E-12 | -0.0105 | 0.8455 | -4.3821 | 0.027472 |
| 1369 | Beef intake | -0.1425 | 5.28E-06 | 0.1674 | 4.66E-09 | -7.3092 | 0.027512 |
| 670_1 | Type of accommodation lived in: A house or bungalow | -0.3461 | 4.9E-13 | -0.0374 | 0.4118 | -4.6726 | 0.027985 |
| 22601_33113434 | Job coding: non-commissioned officers or other rank of armed forces | 0.0878 | 0.3287 | 0.396 | 0.0002 | -2.1943 | 0.028185 |
| I9_CHD | Major coronary heart disease event | -0.0523 | 0.1122 | 0.2557 | 1.66E-11 | -6.1277 | 0.028265 |
| I9_CHD_NOREV | Major coronary heart disease event excluding revascularizations | -0.0523 | 0.1122 | 0.2557 | 1.66E-11 | -6.1277 | 0.028265 |
| 22601_41503307 | Job coding: general office assistant/clerk, clerical officer, clerk-typist, office supervisor, press/newspaper corrector/reader | -0.0782 | 0.3481 | 0.2291 | 0.0338 | -2.2534 | 0.028547 |
| 20422 | More irritable than usual during worst period of anxiety | -0.0214 | 0.7625 | 0.2834 | 0.0025 | -2.5918 | 0.029572 |
| 20003_1141177526 | Treatment/medication code: esomeprazole | -0.0677 | 0.4709 | 0.2356 | 0.0311 | -2.1058 | 0.030202 |
| M13_SOFTTISSUENAS | Other specified/unspecified soft tissue disorders | -0.0779 | 0.3193 | 0.224 | 0.0247 | -2.3826 | 0.0308 |
| 6155_100 | Vitamin and mineral supplements: None of the above | -0.1762 | 8.01E-08 | 0.1253 | 0.0001 | -6.5097 | 0.030973 |
| K80 | Diagnoses - main ICD10: K80 Cholelithiasis | -0.0105 | 0.8062 | 0.2899 | 3.9E-11 | -4.8885 | 0.031452 |
| 20003_1140909786 | Treatment/medication code: beclometasone | -0.0115 | 0.8828 | 0.2872 | 0.0008 | -2.5746 | 0.032204 |
| 6138_4 | Qualifications: CSEs or equivalent | -0.1434 | 6.2E-05 | 0.1534 | 4.15E-05 | -5.7328 | 0.033062 |

| | | | | | | | |
|---|---|---|---|---|---|---|---|
| 2405 | Number of children fathered | -0.1712 | 5.16E-06 | 0.125 | 0.001 | -5.5408 | 0.033337 |
| 680_2 | Own or rent accommodation lived in: Own with a mortgage | -0.141 | 0.0028 | 0.1549 | 0.0006 | -4.5278 | 0.033475 |
| 20003_1141146234 | Treatment/medication code: atorvastatin | -0.0527 | 0.2136 | 0.243 | 1.17E-06 | -4.515 | 0.033567 |
| I9_K_CARDIAC | Death due to cardiac causes | -0.0241 | 0.74 | 0.2699 | 0.0044 | -2.4638 | 0.03436 |
| 30530 | Sodium in urine | -0.1336 | 3E-08 | 0.1595 | 2.37E-09 | -8.1489 | 0.034787 |
| 20003_1140860806 | Treatment/medication code: ramipril | -0.1224 | 0.007 | 0.1702 | 0.0001 | -4.6026 | 0.035025 |
| 136 | Number of operations, self-reported | -0.0554 | 0.0665 | 0.2369 | 6.76E-19 | -7.2512 | 0.035169 |
| 4282 | Maximum digits remembered correctly | -0.1224 | 0.01 | 0.1695 | 0.0001 | -4.48 | 0.035362 |
| 1120 | Weekly usage of mobile phone in last 3 months | -0.1149 | 0.0003 | 0.1764 | 5.52E-09 | -6.6101 | 0.035652 |
| M13_IMPINGEMENT | Impingement syndrome of shoulder | -0.1467 | 0.0242 | 0.1442 | 0.0164 | -3.2833 | 0.035847 |
| 20003_1140864992 | Treatment/medication code: tramadol | -0.0413 | 0.4344 | 0.2466 | 4.34E-07 | -4.0002 | 0.037335 |
| 22617_3311 | Job SOC coding: NCOs and other ranks | 0.0957 | 0.2801 | 0.3829 | 0.0003 | -2.0777 | 0.03769 |
| 20003_1140870570 | Treatment/medication code: vitamin b12 preparation | -0.0197 | 0.8402 | 0.2671 | 0.0232 | -1.8749 | 0.037893 |
| 20086_10 | Type of special diet followed: Low calorie | -0.1528 | 0.0234 | 0.1325 | 0.0635 | -2.9057 | 0.038666 |
| I9_IHD | Ischaemic heart disease, wide definition | -0.0621 | 0.0293 | 0.2223 | 3.29E-12 | -6.6485 | 0.039135 |
| 6139_1 | Gas or solid-fuel cooking/heating: A gas hob or gas cooker | -0.191 | 0.0246 | 0.0933 | 0.2461 | -2.4299 | 0.039187 |
| K42 | Diagnoses - main ICD10: K42 Umbilical hernia | -0.1358 | 0.0439 | 0.1483 | 0.0216 | -3.0431 | 0.039292 |
| 20002_1162 | Non-cancer illness code, self-reported: cholelithiasis/gall stones | -0.0897 | 0.1186 | 0.1937 | 0.0002 | -3.6429 | 0.039662 |
| 20003_1140861958 | Treatment/medication code: simvastatin | -0.112 | 4.48E-05 | 0.1712 | 1.42E-07 | -6.6621 | 0.039768 |
| 20003_1140884444 | Treatment/medication code: codeine | -0.1656 | 0.0316 | 0.1176 | 0.1205 | -2.6193 | 0.039768 |
| 3647 | Duration of other exercises | -0.0899 | 0.0443 | 0.1879 | 0.0003 | -4.0378 | 0.042721 |
| 22608_1 | Workplace very hot: Sometimes | -0.0107 | 0.8282 | 0.2654 | 1.07E-05 | -3.5477 | 0.043686 |
| 20003_1140866738 | Treatment/medication code: atenolol | -0.0786 | 0.0581 | 0.1961 | 2.51E-07 | -4.8819 | 0.044494 |
| 22601_21113020 | Job coding: chemist (analytic, developmental, industrial, research) | -0.2003 | 0.0238 | 0.0738 | 0.4085 | -2.1789 | 0.044844 |
| 6149_6 | Mouth/teeth dental problems: Dentures | -0.0403 | 0.1773 | 0.2306 | 2.4E-16 | -6.6022 | 0.046749 |
| 6146_3 | Attendance/disability/mobility allowance: Blue badge | -0.0408 | 0.2872 | 0.23 | 9.78E-10 | -5.0455 | 0.046809 |
| K30 | Diagnoses - main ICD10: K30 Dyspepsia | -0.0837 | 0.2825 | 0.1869 | 0.0348 | -2.2937 | 0.046931 |
| M13_SYNOTEND | Disorders of synovium and tendon | -0.1542 | 0.024 | 0.1156 | 0.0745 | -2.8657 | 0.047418 |
| E4_OBESITY | Obesity | -0.1163 | 0.0565 | 0.1534 | 0.0197 | -3.0058 | 0.047479 |
| 6159_7 | Pain type(s) experienced in last month: Knee pain | -0.0202 | 0.4883 | 0.2488 | 9.29E-20 | -6.7301 | 0.04791 |
| XIII_MUSCULOSKELET | Diseases of the musculoskeletal system and connective tissue | -0.1266 | 5.99E-05 | 0.1415 | 2.77E-05 | -5.8027 | 0.048468 |
| E66 | Diagnoses - main ICD10: E66 Obesity | -0.11 | 0.0688 | 0.1579 | 0.0182 | -2.9723 | 0.048593 |
| 22601_41413308 | Job coding: telephonist, telephone/switchboard operator | -0.0544 | 0.544 | 0.2119 | 0.0457 | -1.9176 | 0.049599 |
| 20002_1331 | Non-cancer illness code, self-reported: pernicious anaemia | -0.0467 | 0.5516 | 0.2189 | 0.0045 | -2.4154 | 0.050045 |
| 23118 | Leg predicted mass (left) | -0.0823 | 4.98E-06 | 0.1823 | 8.59E-19 | -9.6724 | 0.050687 |
| I9_CORATHER | Coronary atherosclerosis | -0.0702 | 0.025 | 0.1937 | 2.77E-09 | -5.8393 | 0.051141 |
| 23117 | Leg fat-free mass (left) | -0.0814 | 6.8E-06 | 0.1824 | 8.4E-19 | -9.62 | 0.051206 |
| 22606_1 | Workplace very noisy: Sometimes | 0.0017 | 0.977 | 0.265 | 1.13E-05 | -3.1573 | 0.051532 |
| 22609_1 | Workplace very dusty: Sometimes | 0.0699 | 0.2395 | 0.3332 | 1.01E-05 | -2.7391 | 0.051532 |
| 1707_3 | Handedness (chirality/laterality): Use both right and left hands equally | 0.0117 | 0.8802 | 0.2746 | 0.0021 | -2.2181 | 0.051794 |
| 6177_1 | Medication for cholesterol, blood pressure or diabetes: Cholesterol lowering medication | -0.0875 | 0.0024 | 0.1751 | 2.4E-08 | -6.1632 | 0.051992 |
| 23122 | Arm predicted mass (right) | -0.0883 | 2.08E-06 | 0.1741 | 1.2E-17 | -9.505 | 0.052123 |
| 23126 | Arm predicted mass (left) | -0.0816 | 1.19E-05 | 0.1807 | 2.47E-18 | -9.4254 | 0.05219 |
| 23105 | Basal metabolic rate | -0.08 | 1.1E-05 | 0.182 | 2E-19 | -9.636 | 0.052388 |
| 23121 | Arm fat-free mass (right) | -0.0881 | 1.89E-06 | 0.1738 | 9.78E-18 | -9.5357 | 0.052454 |

| | | | | | | | |
|---|---|---|---|---|---|---|---|
| 22607_1 | Workplace very cold: Sometimes | -0.0049 | 0.9079 | 0.2569 | 3.15E-08 | -4.1607 | 0.052521 |
| HEARTFAIL | Heart failure | -0.1108 | 0.0803 | 0.1507 | 0.0405 | -2.6959 | 0.05272 |
| I9_HEARTFAIL | Heart failure,strict | -0.1108 | 0.0803 | 0.1507 | 0.0405 | -2.6959 | 0.05272 |
| I9_HEARTFAIL_NS | Heart failure, not strict | -0.1108 | 0.0803 | 0.1507 | 0.0405 | -2.6959 | 0.05272 |
| N35 | Diagnoses - main ICD10: N35 Urethral stricture | 0.0014 | 0.9911 | 0.2623 | 0.0194 | -1.528 | 0.053121 |
| I25 | Diagnoses - main ICD10: I25 Chronic ischaemic heart disease | -0.0676 | 0.0337 | 0.1932 | 5.02E-09 | -5.6908 | 0.053188 |
| 23125 | Arm fat-free mass (left) | -0.0773 | 3.12E-05 | 0.1816 | 1.81E-18 | -9.3033 | 0.054475 |
| 1538_2 | Major dietary changes in the last 5 years: Yes, because of other reasons | -0.0561 | 0.1455 | 0.2026 | 4.76E-07 | -4.6477 | 0.054611 |
| 1170 | Getting up in morning | -0.2445 | 1.13E-21 | 0.0125 | 0.6269 | -7.0848 | 0.055786 |
| 22704 | Home location - north co-ordinate (rounded) | -0.0871 | 0.0455 | 0.1697 | 2.8E-05 | -4.3207 | 0.055925 |
| 22660_105 | Gap coding: Looking after the home and/or family | 0.0303 | 0.7362 | 0.2857 | 0.0027 | -1.9505 | 0.056909 |
| 20002_1075 | Non-cancer illness code, self-reported: heart attack/myocardial infarction | -0.0468 | 0.1849 | 0.2077 | 3.38E-08 | -4.9347 | 0.057548 |
| 6153_5 | Medication for cholesterol, blood pressure, diabetes, or take exogenous hormones: Oral contraceptive pill or minipill | -0.2284 | 0.0235 | 0.026 | 0.7622 | -1.919 | 0.05762 |
| 2654_8 | Non-butter spread type details: Other low or reduced fat spread | -0.0756 | 0.2396 | 0.1781 | 0.006 | -2.7813 | 0.058122 |
| 4957 | FI3 : word interpolation | -0.1607 | 3.43E-06 | 0.092 | 0.0042 | -5.3541 | 0.058844 |
| 30510 | Creatinine (enzymatic) in urine | -0.0836 | 0.0007 | 0.1685 | 6.09E-11 | -7.0445 | 0.059282 |
| 6150_1 | Vascular/heart problems diagnosed by doctor: Heart attack | -0.04 | 0.266 | 0.2112 | 2.28E-08 | -4.8123 | 0.059942 |
| I9_MI_STRICT | Myocardial infarction, strict | -0.0447 | 0.2013 | 0.2058 | 1.71E-07 | -4.7533 | 0.06046 |
| R06 | Diagnoses - main ICD10: R06 Abnormalities of breathing | 0.0119 | 0.8926 | 0.262 | 0.0178 | -1.7672 | 0.060757 |
| 30300 | High light scatter reticulocyte count | -0.0757 | 0.001 | 0.1737 | 5.94E-16 | -7.9215 | 0.06128 |
| 20003_1140872228 | Treatment/medication code: gabapentin | -0.0084 | 0.9108 | 0.2405 | 0.0059 | -2.1577 | 0.061656 |
| 20003_1140888552 | Treatment/medication code: enalapril | -0.06 | 0.3605 | 0.1889 | 0.009 | -2.5478 | 0.061656 |
| 22617_2111 | Job SOC coding: Chemists | -0.1812 | 0.0184 | 0.0667 | 0.4164 | -2.2052 | 0.062413 |
| 23114 | Leg predicted mass (right) | -0.0793 | 1.04E-05 | 0.1686 | 7.29E-17 | -9.1624 | 0.062413 |
| 23113 | Leg fat-free mass (right) | -0.0794 | 1.03E-05 | 0.1681 | 9.82E-17 | -9.1476 | 0.062718 |
| R07 | Diagnoses - main ICD10: R07 Pain in throat and chest | 0.0224 | 0.5229 | 0.2694 | 3.06E-12 | -4.7343 | 0.0631 |
| 767 | Length of working week for main job | -0.1341 | 0.0022 | 0.1123 | 0.0077 | -4.051 | 0.063562 |
| 2877_3 | Type of tobacco previously smoked: Cigars or pipes | -0.2783 | 0.0312 | -0.0321 | 0.7403 | -1.5245 | 0.063716 |
| E4_OBESITYNAS | Obesity, other/unspecified | -0.104 | 0.0796 | 0.1405 | 0.0264 | -2.8189 | 0.065041 |
| III_BLOOD_IMMUN | Diseases of the blood and blood-forming organs and certain disorders involving the immune mechanism | -0.0669 | 0.3045 | 0.1774 | 0.0035 | -2.7403 | 0.065198 |
| 30290 | High light scatter reticulocyte percentage | -0.0756 | 0.0011 | 0.168 | 5.39E-15 | -7.7193 | 0.065751 |
| 6177_3 | Medication for cholesterol, blood pressure or diabetes: Insulin | -0.0793 | 0.2838 | 0.164 | 0.033 | -2.2798 | 0.065989 |
| M13_OTHERJOINT | #Other joint disorders | -0.1203 | 0.0012 | 0.123 | 0.0025 | -4.4071 | 0.065989 |
| 20003_1140865634 | Treatment/medication code: omeprazole | -0.0214 | 0.5868 | 0.2209 | 7.09E-09 | -4.4152 | 0.066787 |
| 20111_9 | Illnesses of siblings: Diabetes | -0.0284 | 0.4528 | 0.2126 | 7.01E-08 | -4.4139 | 0.067835 |
| 102250 | Sweet snack consumers | -0.0608 | 0.4107 | 0.18 | 0.0288 | -2.1757 | 0.067998 |
| 6146_1 | Attendance/disability/mobility allowance: Attendance allowance | 0.0636 | 0.227 | 0.304 | 8.65E-07 | -2.9599 | 0.068323 |
| M13_ROTATORCUFF | Rotator cuff syndrome | -0.1214 | 0.0289 | 0.1175 | 0.04 | -2.9975 | 0.069556 |
| 30250 | Reticulocyte count | -0.0822 | 0.0005 | 0.1563 | 5.08E-15 | -7.7288 | 0.069887 |
| IX_CIRCULATORY | Diseases of the circulatory system | -0.0334 | 0.2339 | 0.2049 | 6.36E-09 | -5.2889 | 0.070053 |
| 20003_1140871310 | Treatment/medication code: ibuprofen | -0.0713 | 0.1005 | 0.1655 | 5.09E-05 | -3.9754 | 0.07131 |
| 1548 | Variation in diet | 0.0306 | 0.297 | 0.267 | 6.49E-20 | -5.7149 | 0.071648 |

| | | | | | | | |
|---|---|---|---|---|---|---|---|
| 4283 | Number of rounds of numeric memory test performed | -0.103 | 0.0418 | 0.1332 | 0.007 | -3.3401 | 0.071817 |
| 20002_1474 | Non-cancer illness code, self-reported: hiatus hernia | -0.0115 | 0.8301 | 0.2247 | 6.47E-05 | -3.0468 | 0.071817 |
| 4803_12 | Tinnitus: Yes, now a lot of the time | -0.0403 | 0.6171 | 0.1953 | 0.0196 | -2.0288 | 0.072327 |
| 20075 | Home location at assessment - north co-ordinate (rounded) | -0.0762 | 0.0928 | 0.1576 | 0.0002 | -3.7637 | 0.073874 |
| 20002_1138 | Non-cancer illness code, self-reported: gastro-oesophageal reflux (gord) / gastric reflux | -0.0797 | 0.0796 | 0.154 | 0.0009 | -3.5961 | 0.073961 |
| 20002_1465 | Non-cancer illness code, self-reported: osteoarthritis | -0.0561 | 0.1386 | 0.1767 | 4.48E-06 | -4.3091 | 0.074744 |
| 20110_9 | Illnesses of mother: Diabetes | -0.0708 | 0.0493 | 0.1614 | 4.18E-06 | -4.6182 | 0.07527 |
| 30240 | Reticulocyte percentage | -0.0821 | 0.0004 | 0.1496 | 3.58E-14 | -7.5748 | 0.07571 |
| 2877_1 | Type of tobacco previously smoked: Manufactured cigarettes | -0.161 | 0.0472 | 0.0706 | 0.3743 | -2.0393 | 0.075799 |
| 20003_1140879778 | Treatment/medication code: doxazosin | -0.0617 | 0.3141 | 0.1678 | 0.007 | -2.628 | 0.077672 |
| 24011 | Traffic intensity on the nearest major road | 0.0336 | 0.732 | 0.2631 | 0.0239 | -1.5062 | 0.077672 |
| 20002_1563 | Non-cancer illness code, self-reported: urticaria | 0.0253 | 0.8549 | 0.2545 | 0.0487 | -1.2119 | 0.077942 |
| 6153_1 | Medication for cholesterol, blood pressure, diabetes, or take exogenous hormones: Cholesterol lowering medication | -0.05 | 0.1162 | 0.1792 | 4.06E-07 | -4.8166 | 0.077942 |
| I21 | Diagnoses - main ICD10: I21 Acute myocardial infarction | -0.0287 | 0.4252 | 0.2002 | 9.56E-07 | -4.201 | 0.078213 |
| M13_DORSALGIANAS | Other/unspecified dorsalgia | -0.0654 | 0.3838 | 0.1634 | 0.0159 | -2.2614 | 0.078304 |
| 728 | Number of vehicles in household | -0.287 | 6.11E-19 | -0.0585 | 0.0703 | -5.0023 | 0.078576 |
| 4260 | Round of numeric memory test | -0.0996 | 0.0486 | 0.1281 | 0.0094 | -3.2264 | 0.079305 |
| KNEE_ARTHROSIS | Gonarthrosis [arthrosis of knee](FG) | -0.0977 | 0.0087 | 0.129 | 0.0002 | -4.4563 | 0.080223 |
| M17 | Diagnoses - main ICD10: M17 Gonarthrosis [arthrosis of knee] | -0.0977 | 0.0087 | 0.129 | 0.0002 | -4.4563 | 0.080223 |
| 5474 | Leg pain when walking uphill or hurrying | -0.0597 | 0.3416 | 0.1669 | 0.0257 | -2.3201 | 0.080315 |
| 22611_1 | Workplace had a lot of cigarette smoke from other people smoking: Sometimes | 0.0285 | 0.6641 | 0.2546 | 0.0018 | -2.1624 | 0.080778 |
| J45 | Diagnoses - main ICD10: J45 Asthma | -0.0256 | 0.7321 | 0.1999 | 0.0164 | -2.0142 | 0.081336 |
| S82 | Diagnoses - main ICD10: S82 Fracture of lower leg, including ankle | -0.033 | 0.6831 | 0.1924 | 0.0456 | -1.7941 | 0.081429 |
| 20003_1140860954 | Treatment/medication code: isosorbide mononitrate | -0.012 | 0.8749 | 0.2131 | 0.0132 | -1.9602 | 0.081709 |
| 20533 | Trouble falling asleep | -0.1632 | 0.0154 | 0.0617 | 0.3382 | -2.4126 | 0.081896 |
| 23102 | Whole body water mass | -0.0734 | 6.23E-05 | 0.1512 | 4.31E-14 | -8.2851 | 0.082178 |
| 2834 | Bilateral oophorectomy (both ovaries removed) | -0.0939 | 0.0356 | 0.1306 | 0.0022 | -3.6317 | 0.082272 |
| 6154_5 | Medication for pain relief, constipation, heartburn: Omeprazole (e.g. Zanprol) | -0.0052 | 0.8859 | 0.2187 | 2.12E-10 | -4.4706 | 0.082837 |
| 23101 | Whole body fat-free mass | -0.0728 | 7.03E-05 | 0.1509 | 5.6E-14 | -8.2295 | 0.083026 |
| K76 | Diagnoses - main ICD10: K76 Other diseases of liver | -0.04 | 0.5639 | 0.1834 | 0.0286 | -2.0544 | 0.083311 |
| J84 | Diagnoses - main ICD10: J84 Other interstitial pulmonary diseases | -0.0222 | 0.7632 | 0.2006 | 0.0232 | -1.9358 | 0.083881 |
| 22617_9149 | Job SOC coding: Other goods handling and storage occupations n.e.c. | -0.0067 | 0.9375 | 0.2155 | 0.02 | -1.7591 | 0.084455 |
| 6150_2 | Vascular/heart problems diagnosed by doctor: Angina | -0.0022 | 0.9474 | 0.22 | 1.73E-10 | -4.5939 | 0.084455 |
| G6_NERPLEX | Nerve, nerve root and plexus disorders | -0.1441 | 0.0003 | 0.0778 | 0.0593 | -3.869 | 0.084743 |
| 20002_1087 | Non-cancer illness code, self-reported: leg claudication/ intermittent claudication | -0.0334 | 0.6421 | 0.1876 | 0.0222 | -2.0277 | 0.085612 |
| 20002_1093 | Non-cancer illness code, self-reported: pulmonary embolism +/- dvt | -0.0327 | 0.5791 | 0.1883 | 0.0018 | -2.6196 | 0.085612 |
| 5012 | FI8 : chained arithmetic | -0.1068 | 0.0027 | 0.1141 | 0.0022 | -4.2902 | 0.085709 |
| 4803_11 | Tinnitus: Yes, now most or all of the time | 0.0127 | 0.7917 | 0.2329 | 9.08E-06 | -3.0984 | 0.086389 |
| 20002_1311 | Non-cancer illness code, self-reported: spine arthritis/spondylitis | 0.0168 | 0.8311 | 0.2365 | 0.0058 | -1.8894 | 0.086878 |
| R51 | Diagnoses - main ICD10: R51 Headache | 0.063 | 0.3696 | 0.282 | 0.0037 | -1.8278 | 0.087565 |
| 22601_35393271 | Job coding: management information officer, conference/events co-ordinator/organiser, exhibition officer, | 0.0411 | 0.5811 | 0.26 | 0.0014 | -1.9838 | 0.087664 |

| | | | | | | | |
|---|---|---|---|---|---|---|---|
| | work study engineer/officer/analyst, contract adviser/agent, election agent, business system analyst | | | | | | |
| 1883 | Number of full sisters | 0.0947 | 0.016 | 0.3119 | 9.78E-14 | -3.7809 | 0.089352 |
| 20111_5 | Illnesses of siblings: Breast cancer | -0.009 | 0.8984 | 0.2081 | 0.0132 | -1.9774 | 0.089452 |
| 20002_1074 | Non-cancer illness code, self-reported: angina | 0.0087 | 0.7987 | 0.2257 | 6.78E-11 | -4.4798 | 0.089552 |
| 6152_7 | Blood clot, DVT, bronchitis, emphysema, asthma, rhinitis, eczema, allergy diagnosed by doctor: Blood clot in the lung | -0.0406 | 0.4768 | 0.1764 | 0.0027 | -2.6476 | 0.089552 |
| 2415 | Had major operations | -0.0168 | 0.7134 | 0.1999 | 2.97E-06 | -3.461 | 0.089853 |
| 5855_3 | Which eye(s) affected by astigmatism: Both eyes | -0.1929 | 0.0399 | 0.0237 | 0.8147 | -1.569 | 0.089954 |
| 2887 | Number of cigarettes previously smoked daily | 0.0626 | 0.0973 | 0.2791 | 2.06E-13 | -4.0446 | 0.090054 |
| 20536_1 | Weight change during worst episode of depression: Gained weight | 0.0566 | 0.2178 | 0.2723 | 2.08E-07 | -3.0965 | 0.090861 |
| 4728 | Leg pain on walking | 0.0568 | 0.1476 | 0.2711 | 2.92E-12 | -3.8854 | 0.092287 |
| 1508_2 | Coffee type: Instant coffee | -0.0843 | 0.0084 | 0.1297 | 0.0001 | -4.6117 | 0.092595 |
| 30280 | Immature reticulocyte fraction | -0.0441 | 0.0567 | 0.1676 | 9.4E-12 | -6.2734 | 0.094979 |
| 709 | Number in household | -0.2243 | 3.16E-06 | -0.0127 | 0.7902 | -3.1204 | 0.095084 |
| 20003_1140868226 | Treatment/medication code: aspirin | -0.0089 | 0.7978 | 0.2027 | 5.52E-08 | -4.1535 | 0.095084 |
| 20489 | Felt loved as a child | -0.3361 | 7.23E-28 | -0.1248 | 0.0008 | -4.3809 | 0.095398 |
| 2188 | Long-standing illness, disability or infirmity | 0.0838 | 0.001 | 0.2944 | 1.93E-29 | -5.7826 | 0.096135 |
| Z42 | Diagnoses - main ICD10: Z42 Follow-up care involving plastic surgery | -0.0152 | 0.8357 | 0.1952 | 0.0208 | -1.8799 | 0.096346 |
| I9_DVTANDPULM | DVT of lower extremities and pulmonary embolism | -0.0808 | 0.1144 | 0.1287 | 0.0025 | -3.1454 | 0.097301 |
| 137 | Number of treatments/medications taken | -0.0055 | 0.8189 | 0.2024 | 8.91E-19 | -6.2809 | 0.099016 |
| 1508_1 | Coffee type: Decaffeinated coffee (any type) | -0.1176 | 0.0044 | 0.0901 | 0.072 | -3.202 | 0.099232 |
| I9_VTE | Venous thromboembolism | -0.0747 | 0.1203 | 0.1324 | 0.0011 | -3.2902 | 0.099882 |
| 20003_1141192736 | Treatment/medication code: ezetimibe | -0.0597 | 0.3349 | 0.1464 | 0.0375 | -2.1986 | 0.100972 |
| 5001 | FI7 : synonym | -0.1416 | 0.0011 | 0.064 | 0.1578 | -3.2737 | 0.10152 |
| 20003_1140888560 | Treatment/medication code: perindopril | 0.0327 | 0.6451 | 0.2374 | 0.0005 | -2.0764 | 0.102512 |
| 20003_1141194794 | Treatment/medication code: bendroflumethiazide | -0.0709 | 0.0569 | 0.132 | 0.0005 | -3.8258 | 0.104518 |
| 22617_3539 | Job SOC coding: Business and related associate professionals n.e.c. | 0.0523 | 0.5296 | 0.2545 | 0.0037 | -1.6736 | 0.105306 |
| 20107_9 | Illnesses of father: Diabetes | -0.0927 | 0.0273 | 0.1091 | 0.0156 | -3.2745 | 0.105758 |
| 20126_5 | Bipolar and major depression status: Single Probable major depression episode | 0.0455 | 0.6107 | 0.2472 | 0.0271 | -1.4096 | 0.105871 |
| 20003_1140879406 | Treatment/medication code: ranitidine | 0.0128 | 0.8325 | 0.2137 | 0.0006 | -2.3078 | 0.10678 |
| 20003_1140884488 | Treatment/medication code: diclofenac | -0.0555 | 0.3115 | 0.1448 | 0.0091 | -2.5658 | 0.107465 |
| 41248_5000 | Destinations on discharge from hospital (recoded): Transfer to other NHS provider | 0.0198 | 0.7662 | 0.2192 | 0.0047 | -1.9499 | 0.108499 |
| M13_ARTHROSIS | #Arthrosis | -0.0951 | 0.006 | 0.1042 | 0.0041 | -3.98 | 0.108614 |
| 1349 | Processed meat intake | -0.047 | 0.1356 | 0.1522 | 2.23E-06 | -4.4222 | 0.10873 |
| 20003_1140926606 | Treatment/medication code: salbutamol 100micrograms spacehaler | 0.0748 | 0.3162 | 0.2725 | 0.001 | -1.7763 | 0.110471 |
| 22601_31152695 | Job coding: quality assurance technician/co-ordinator | 0.1044 | 0.3989 | 0.3016 | 0.0487 | -1.002 | 0.111056 |
| 6154_1 | Medication for pain relief, constipation, heartburn: Aspirin | -0.0132 | 0.703 | 0.1838 | 1.17E-06 | -3.8443 | 0.111291 |
| AB1_INFECTIONS | Certain infectious and parasitic diseases | 0.1117 | 0.2358 | 0.3085 | 0.0045 | -1.3696 | 0.111526 |
| 20118_5 | Home area population density - urban or rural: England/Wales - Urban - less sparse | 0.1081 | 0.0818 | 0.3035 | 1.7E-05 | -2.0782 | 0.113181 |
| K11_OTHGASTR | Other gastritis (incl. Duodenitis) | -0.023 | 0.669 | 0.1694 | 0.0028 | -2.4593 | 0.116785 |
| 6142_1 | Current employment status: In paid employment or self-employed | -0.2099 | 8.29E-07 | -0.0189 | 0.6576 | -3.1666 | 0.118495 |

| | | | | | | | |
|---|---|---|---|---|---|---|---|
| 20002_1094 | Non-cancer illness code, self-reported: deep venous thrombosis (dvt) | -0.0059 | 0.9014 | 0.1841 | 0.0002 | -2.744 | 0.119727 |
| 6177_2 | Medication for cholesterol, blood pressure or diabetes: Blood pressure medication | -0.0596 | 0.0363 | 0.1301 | 9.33E-07 | -4.8745 | 0.120098 |
| 4717 | Shortness of breath walking on level ground | 0.0592 | 0.1085 | 0.2488 | 1.55E-10 | -3.5362 | 0.120222 |
| 6151_7 | Fractured bone site(s): Other bones | -0.052 | 0.3185 | 0.1357 | 0.0076 | -2.577 | 0.122595 |
| 1279 | Exposure to tobacco smoke outside home | 0.0922 | 0.0042 | 0.2787 | 7.96E-15 | -3.8673 | 0.12411 |
| 2296 | Falls in the last year | 0.05 | 0.0972 | 0.2365 | 2.05E-15 | -4.3957 | 0.12411 |
| 23129 | Trunk fat-free mass | -0.0629 | 0.0008 | 0.1233 | 4.51E-10 | -6.8369 | 0.124491 |
| 1468_1 | Cereal type: Bran cereal (e.g. All Bran, Branflakes) | -0.1803 | 0.0002 | 0.0059 | 0.9085 | -2.6378 | 0.124491 |
| 5103 | 3mm weak meridian angle (left) | -0.0725 | 0.1665 | 0.1132 | 0.0165 | -2.6332 | 0.125128 |
| 6159_6 | Pain type(s) experienced in last month: Hip pain | 0.0559 | 0.1178 | 0.2414 | 2.91E-13 | -3.8046 | 0.125383 |
| 41215_0 | Detention categories: Informal, not formally detained | -0.037 | 0.5898 | 0.1484 | 0.0303 | -1.9124 | 0.125511 |
| 23130 | Trunk predicted mass | -0.0619 | 0.001 | 0.1228 | 6.22E-10 | -6.7647 | 0.126408 |
| 5112 | 3mm cylindrical power angle (left) | -0.0714 | 0.1731 | 0.1131 | 0.017 | -2.6112 | 0.126665 |
| 22617_5245 | Job SOC coding: Computer engineers, installation and maintenance | 0.1049 | 0.1686 | 0.2876 | 0.0188 | -1.2672 | 0.128995 |
| VI_NERVOUS | Diseases of the nervous system | -0.0643 | 0.1095 | 0.1184 | 0.0059 | -3.1037 | 0.128995 |
| 3143 | Ankle spacing width | -0.0348 | 0.0351 | 0.1477 | 2.02E-13 | -7.0179 | 0.129256 |
| 21021 | Pulse wave Arterial Stiffness index | -0.0431 | 0.349 | 0.139 | 0.005 | -2.6948 | 0.129778 |
| 22502 | Cough on most days | 0.0453 | 0.2806 | 0.2269 | 8.4E-07 | -2.912 | 0.130433 |
| 6152_5 | Blood clot, DVT, bronchitis, emphysema, asthma, rhinitis, eczema, allergy diagnosed by doctor: Blood clot in the leg (DVT) | -0.0023 | 0.9621 | 0.1785 | 0.0004 | -2.5929 | 0.131486 |
| 20003_1140881856 | Treatment/medication code: salbutamol | 0.0203 | 0.7548 | 0.2003 | 0.0012 | -2.0085 | 0.132545 |
| 6154_2 | Medication for pain relief, constipation, heartburn: Ibuprofen (e.g. Nurofen) | -0.0494 | 0.2133 | 0.1292 | 0.0004 | -3.3159 | 0.134413 |
| N39 | Diagnoses - main ICD10: N39 Other disorders of urinary system | 0.0394 | 0.4686 | 0.218 | 3.03E-05 | -2.369 | 0.134413 |
| 5113 | 6mm cylindrical power angle (left) | -0.0713 | 0.1166 | 0.1067 | 0.0482 | -2.5208 | 0.135219 |
| 5102 | 6mm weak meridian angle (left) | -0.0727 | 0.1075 | 0.1052 | 0.0484 | -2.5456 | 0.135353 |
| G56 | Diagnoses - main ICD10: G56 Mononeuropathies of upper limb | -0.117 | 0.0028 | 0.0604 | 0.1216 | -3.2082 | 0.136028 |
| 20427 | Frequent trouble falling or staying asleep during worst period of anxiety | -0.1699 | 0.0054 | 0.0071 | 0.9155 | -1.96 | 0.136569 |
| 874 | Duration of walks | -0.1483 | 8.86E-07 | 0.0285 | 0.3494 | -4.1191 | 0.13684 |
| 699 | Length of time at current address | -0.243 | 3.9E-13 | -0.0672 | 0.0774 | -3.4703 | 0.138202 |
| 4119 | Ankle spacing width (right) | -0.0437 | 0.0181 | 0.1319 | 6.97E-08 | -5.7198 | 0.138475 |
| 22615_1 | Workplace had a lot of diesel exhaust: Sometimes | 0.0882 | 0.1111 | 0.2637 | 0.0002 | -1.9677 | 0.138612 |
| 2463 | Fractured/broken bones in last 5 years | -0.0695 | 0.0555 | 0.1058 | 0.0088 | -3.2276 | 0.138886 |
| ASTHMA_CHILD | Childhood asthma (age<16) | 0.001 | 0.9881 | 0.1763 | 0.0238 | -1.6823 | 0.138886 |
| J10_ASTHMA | Asthma | 0.001 | 0.9881 | 0.1763 | 0.0238 | -1.6823 | 0.138886 |
| J10_ASTHMA_MAIN | Asthma | 0.001 | 0.9881 | 0.1763 | 0.0238 | -1.6823 | 0.138886 |
| 3741 | Stomach/abdominal pain for 3+ months | 0.0712 | 0.5167 | 0.2464 | 0.0289 | -1.113 | 0.139023 |
| I84 | Diagnoses - main ICD10: I84 Haemorrhoids | 0.0042 | 0.9245 | 0.1791 | 0.0003 | -2.6226 | 0.139435 |
| 2654_4 | Non-butter spread type details: Soft (tub) margarine | -0.0009 | 0.9833 | 0.1736 | 0.0003 | -2.6432 | 0.139985 |
| 1960 | Fed-up feelings | -0.0009 | 0.974 | 0.1728 | 4.69E-12 | -4.5909 | 0.14109 |
| 4100 | Ankle spacing width (left) | -0.0459 | 0.0146 | 0.1273 | 1.37E-07 | -5.6519 | 0.141784 |
| 2907 | Ever stopped smoking for 6+ months | 0.0705 | 0.3018 | 0.2432 | 0.0006 | -1.7581 | 0.14248 |
| 20002_1473 | Non-cancer illness code, self-reported: high cholesterol | -0.0507 | 0.051 | 0.1217 | 0.0002 | -4.119 | 0.142899 |
| 6179_100 | Mineral and other dietary supplements: None of the above | -0.0042 | 0.9048 | 0.1676 | 5.2E-08 | -3.6672 | 0.143739 |
| 1508_4 | Coffee type: Other type of coffee | 0.0275 | 0.7425 | 0.1992 | 0.0369 | -1.3522 | 0.143879 |
| XXI_HEALTHFACTORS | Factors influencing health status and contact with health services | -0.0236 | 0.6805 | 0.1478 | 0.0047 | -2.2072 | 0.144301 |
| 20003_1141156836 | Treatment/medication code: candesartan cilexetil | 0.0016 | 0.9745 | 0.1728 | 0.0017 | -2.3257 | 0.144583 |

| | | | | | | | |
|---|---|---|---|---|---|---|---|
| 4968 | FI4 : positional arithmetic | -0.1161 | 0.002 | 0.0539 | 0.1565 | -3.1801 | 0.146279 |
| 20003_1140879802 | Treatment/medication code: amlodipine | -0.0405 | 0.2192 | 0.1292 | 7.54E-05 | -3.6583 | 0.146706 |
| 100890 | Milk added to cereal | -0.2394 | 0.0124 | -0.0698 | 0.4478 | -1.2769 | 0.146848 |
| 6153_2 | Medication for cholesterol, blood pressure, diabetes, or take exogenous hormones: Blood pressure medication | -0.0317 | 0.2573 | 0.1367 | 3.79E-06 | -4.133 | 0.148562 |
| G6_CARPTU | Carpal tunnel syndrome | -0.121 | 0.0019 | 0.0466 | 0.2322 | -3.0426 | 0.149713 |
| N92 | Diagnoses - main ICD10: N92 Excessive, frequent and irregular menstruation | 0.0257 | 0.6575 | 0.1925 | 0.0022 | -1.9512 | 0.150869 |
| ASTHMA_HOSPITAL1 | Asthma, hospital admissions 1 | 0.0036 | 0.9587 | 0.1701 | 0.0303 | -1.5931 | 0.151304 |
| 4079 | Diastolic blood pressure, automated reading | -0.0366 | 0.0992 | 0.1287 | 2.59E-08 | -5.1595 | 0.153053 |
| R10 | Diagnoses - main ICD10: R10 Abdominal and pelvic pain | 0.0162 | 0.7296 | 0.1813 | 1.1E-05 | -2.6479 | 0.153346 |
| 3773 | Knee pain for 3+ months | 0.0286 | 0.638 | 0.1932 | 0.0034 | -1.8373 | 0.15408 |
| 826 | Job involves shift work | -0.0305 | 0.4287 | 0.1338 | 0.001 | -2.9403 | 0.154521 |
| 1707_1 | Handedness (chirality/laterality): Right-handed | -0.1493 | 0.0016 | 0.0149 | 0.7648 | -2.3858 | 0.154668 |
| M23 | Diagnoses - main ICD10: M23 Internal derangement of knee | -0.1041 | 0.0245 | 0.0593 | 0.206 | -2.4794 | 0.15585 |
| 2247_1 | Hearing difficulty/problems: Yes | 0.0602 | 0.028 | 0.2233 | 6.56E-15 | -4.1105 | 0.156294 |
| 20002_1478 | Non-cancer illness code, self-reported: cervical spondylosis | 0.0178 | 0.8238 | 0.1805 | 0.0274 | -1.4237 | 0.156888 |
| 22610_1 | Workplace full of chemical or other fumes: Sometimes | 0.0751 | 0.1759 | 0.2372 | 0.0002 | -1.9101 | 0.157782 |
| K21 | Diagnoses - main ICD10: K21 Gastro-oesophageal reflux disease | 0.0007 | 0.9883 | 0.162 | 0.0006 | -2.4626 | 0.158979 |
| K85 | Diagnoses - main ICD10: K85 Acute pancreatitis | 0.0493 | 0.516 | 0.2099 | 0.0105 | -1.4373 | 0.160032 |
| 6141_4 | How are people in household related to participant: Mother and/or father | 0.0305 | 0.7119 | 0.1901 | 0.0409 | -1.2831 | 0.161543 |
| 20003_1141188442 | Treatment/medication code: glucosamine product | -0.2152 | 0.0007 | -0.0564 | 0.2667 | -1.949 | 0.162759 |
| 4803_13 | Tinnitus: Yes, now some of the time | 0.1344 | 0.0975 | 0.2927 | 0.0007 | -1.3392 | 0.163522 |
| 20528 | Diagnosed with life-threatening illness | 0.1121 | 0.0889 | 0.2696 | 7.5E-05 | -1.662 | 0.164747 |
| 3064 | Peak expiratory flow (PEF) | -0.1227 | 6.39E-09 | 0.0345 | 0.1879 | -4.673 | 0.165208 |
| 93 | Systolic blood pressure, manual reading | -0.1337 | 0.0021 | 0.0235 | 0.6044 | -2.5002 | 0.165208 |
| 20003_2038460150 | Treatment/medication code: paracetamol | 0.0163 | 0.5912 | 0.1733 | 7.47E-09 | -3.6759 | 0.165516 |
| 22599 | Number of jobs held | 0.1615 | 0.0014 | 0.3172 | 4.42E-08 | -2.0263 | 0.167526 |
| M13_LOWBACKPAIN | Low back pain | -0.0109 | 0.8287 | 0.1444 | 0.0039 | -2.1897 | 0.168147 |
| 1498 | Coffee intake | -0.0862 | 0.001 | 0.0683 | 0.0089 | -4.1697 | 0.169395 |
| 20003_1140865354 | Treatment/medication code: gaviscon liquid | 0.069 | 0.3909 | 0.2233 | 0.015 | -1.2644 | 0.169708 |
| 1379 | Lamb/mutton intake | -0.0537 | 0.1021 | 0.1005 | 0.0008 | -3.4633 | 0.169864 |
| N20 | Diagnoses - main ICD10: N20 Calculus of kidney and ureter | -0.0447 | 0.3428 | 0.1086 | 0.0224 | -2.2917 | 0.171277 |
| 1418_6 | Milk type used: Never/rarely have milk | -0.0254 | 0.5391 | 0.1273 | 0.0028 | -2.5706 | 0.172223 |
| 6145_2 | Illness, injury, bereavement, stress in last 2 years: Serious illness, injury or assault of a close relative | 0.0056 | 0.9173 | 0.158 | 0.0006 | -2.1445 | 0.172698 |
| 22618_1 | Breathing problems improved/stopped away from workplace or on holiday: Yes | 0.1094 | 0.1857 | 0.2616 | 0.0015 | -1.3045 | 0.173014 |
| 6138_3 | Qualifications: O levels/GCSEs or equivalent | -0.1214 | 9.39E-05 | 0.0302 | 0.3171 | -3.4971 | 0.173967 |
| 20471 | Ever seen an un-real vision | 0.1891 | 0.0078 | 0.3399 | 0.0001 | -1.3247 | 0.175242 |
| 6159_4 | Pain type(s) experienced in last month: Back pain | 0.0292 | 0.3266 | 0.1796 | 6.85E-11 | -3.709 | 0.175881 |
| 3466 | Time from waking to first cigarette | -0.2103 | 0.0002 | -0.061 | 0.3068 | -1.8316 | 0.177648 |
| 2492 | Taking other prescription medications | 0.0431 | 0.0977 | 0.1914 | 3.52E-14 | -4.0879 | 0.179264 |
| 20003_1140883548 | Treatment/medication code: ipratropium | 0.0725 | 0.4125 | 0.2204 | 0.0344 | -1.0818 | 0.179913 |
| 6154_4 | Medication for pain relief, constipation, heartburn: Ranitidine (e.g. Zantac) | 0.0383 | 0.5214 | 0.1861 | 0.0036 | -1.6888 | 0.180075 |
| 6150_4 | Vascular/heart problems diagnosed by doctor: High blood pressure | -0.0114 | 0.6124 | 0.136 | 6.53E-09 | -4.5503 | 0.180726 |

| | | | | | | | |
|---|---|---|---|---|---|---|---|
| K02 | Diagnoses - main ICD10: K02 Dental caries | 0.1459 | 0.2355 | 0.2932 | 0.0448 | -0.7715 | 0.180889 |
| 4674 | Private healthcare | 0.0087 | 0.8582 | 0.1559 | 0.0008 | -2.186 | 0.181052 |
| 20002_1065 | Non-cancer illness code, self-reported: hypertension | -0.0127 | 0.5671 | 0.1342 | 7.93E-09 | -4.5646 | 0.181542 |
| I_INFECT_PARASIT | Certain infectious and parasitic diseases | 0.0946 | 0.2624 | 0.2414 | 0.0062 | -1.2018 | 0.181705 |
| 20111_8 | Illnesses of siblings: High blood pressure | -0.0185 | 0.5794 | 0.128 | 7.96E-05 | -3.1532 | 0.182196 |
| XIV_GENITOURINARY | Diseases of the genitourinary system | 0.0154 | 0.672 | 0.1619 | 5.56E-06 | -2.8774 | 0.182196 |
| 6164_5 | Types of physical activity in last 4 weeks: Heavy DIY (eg: weeding, lawn mowing, carpentry, digging) | -0.199 | 2.1E-10 | -0.0539 | 0.0672 | -3.3736 | 0.184496 |
| 41248_5001 | Destinations on discharge from hospital (recoded): Transfer to other NHS provider: General ward, young physically disabled, A&E | 0.1258 | 0.1669 | 0.2704 | 0.011 | -1.0328 | 0.185322 |
| 5107 | 3mm strong meridian angle (right) | -0.0492 | 0.3017 | 0.0939 | 0.0344 | -2.1984 | 0.187814 |
| K29 | Diagnoses - main ICD10: K29 Gastritis and duodenitis | 0.0205 | 0.6915 | 0.1623 | 0.0026 | -1.8986 | 0.189991 |
| 4825 | Noisy workplace | 0.0623 | 0.1075 | 0.2035 | 1.01E-06 | -2.4851 | 0.191 |
| XII_SKIN_SUBCUTAN | Diseases of the skin and subcutaneous tissue | 0.0075 | 0.9105 | 0.1475 | 0.044 | -1.4088 | 0.19303 |
| 20110_1 | Illnesses of mother: Heart disease | 0.0224 | 0.6249 | 0.1622 | 0.0001 | -2.2399 | 0.193369 |
| 41248_1000 | Destinations on discharge from hospital (recoded): Usual Place of residence | 0.0404 | 0.2071 | 0.1793 | 3.44E-07 | -2.9198 | 0.194901 |
| ICDMAIN_ANY_ENTRY | Any ICDMAIN event in hilmo or causes of death | 0.0181 | 0.564 | 0.1569 | 3.78E-07 | -3.1507 | 0.195072 |
| 129 | Place of birth in UK - north co-ordinate | 0.0057 | 0.8897 | 0.1423 | 0.0006 | -2.333 | 0.19885 |
| 1418_3 | Milk type used: Skimmed | -0.0586 | 0.1048 | 0.0779 | 0.0368 | -2.6296 | 0.199023 |
| 41235 | Spells in hospital | 0.0405 | 0.2334 | 0.1767 | 1.17E-07 | -2.8577 | 0.199542 |
| 22601_32223079 | Job coding: occupational therapist | -0.2241 | 0.0394 | -0.0882 | 0.3912 | -0.9075 | 0.200062 |
| 4979 | FI5 : family relationship calculation | -0.0217 | 0.6435 | 0.1138 | 0.0121 | -2.0804 | 0.200756 |
| 20003_1140879616 | Treatment/medication code: amitriptyline | 0.0365 | 0.5147 | 0.1715 | 0.0035 | -1.6626 | 0.201626 |
| 6154_3 | Medication for pain relief, constipation, heartburn: Paracetamol | 0.0092 | 0.7527 | 0.1434 | 4.41E-07 | -3.3004 | 0.203022 |
| 20003_1140860696 | Treatment/medication code: lisinopril | -0.0247 | 0.5708 | 0.1091 | 0.0098 | -2.2051 | 0.203722 |
| 4990 | FI6 : conditional arithmetic | -0.057 | 0.0821 | 0.0767 | 0.0296 | -2.7746 | 0.203897 |
| 22617_3222 | Job SOC coding: Occupational therapists | -0.2196 | 0.0308 | -0.0866 | 0.3682 | -0.9501 | 0.205127 |
| 135 | Number of self-reported non-cancer illnesses | 0.0727 | 0.002 | 0.2056 | 1.96E-20 | -4.111 | 0.205303 |
| M13_DORSALGIA | Dorsalgia | -0.0245 | 0.6037 | 0.1075 | 0.0127 | -2.0675 | 0.206891 |
| M54 | Diagnoses - main ICD10: M54 Dorsalgia | -0.0245 | 0.6037 | 0.1075 | 0.0127 | -2.0675 | 0.206891 |
| M13_MENISCUSDERANGEMENTS | Meniscus derangement | -0.0943 | 0.0412 | 0.0347 | 0.4715 | -1.9321 | 0.212238 |
| 2335 | Chest pain or discomfort | 0.1199 | 4.01E-05 | 0.2485 | 8.34E-16 | -3.0249 | 0.212957 |
| 22613_2 | Worked with paints, thinners or glues: Often | 0.0752 | 0.4221 | 0.203 | 0.0243 | -0.9831 | 0.214399 |
| 6160_100 | Leisure/social activities: None of the above | -0.0114 | 0.7137 | 0.1161 | 0.0009 | -2.727 | 0.214941 |
| 4080 | Systolic blood pressure, automated reading | -0.0424 | 0.0587 | 0.085 | 0.0002 | -3.977 | 0.215122 |
| 2704 | Years since last cervical smear test | 0.0217 | 0.7339 | 0.1484 | 0.0062 | -1.5135 | 0.216391 |
| XVIII_MISCFINDINGS | Symptoms, signs and abnormal clinical and laboratory findings, not elsewhere classified | 0.0973 | 0.0025 | 0.2232 | 4.91E-13 | -2.8257 | 0.217846 |
| 20522 | Been in a confiding relationship as an adult | -0.1417 | 0.0006 | -0.0165 | 0.7077 | -2.077 | 0.219124 |
| 41231_1 | Hospital episode type: General episode | 0.0169 | 0.6489 | 0.139 | 0.0002 | -2.3115 | 0.224836 |
| 1628 | Alcohol intake versus 10 years previously | 0.0332 | 0.2718 | 0.1545 | 4.55E-06 | -2.6806 | 0.226324 |
| 20003_1140875408 | Treatment/medication code: allopurinol | -0.0459 | 0.1818 | 0.0752 | 0.0356 | -2.4391 | 0.226697 |
| COPD_EXCL | COPD differential diagnosis | 0.0321 | 0.4093 | 0.1525 | 0.0007 | -2.0215 | 0.228004 |
| ILD_DIFF_DG | ILD differential diagnosis | 0.0321 | 0.4093 | 0.1525 | 0.0007 | -2.0215 | 0.228004 |

| | | | | | | | |
|---|---|---|---|---|---|---|---|
| 20548_7 | Manifestations of mania or irritability: I was easily distracted | 0.1757 | 0.0027 | 0.2953 | 1.18E-06 | -1.4175 | 0.229504 |
| M48 | Diagnoses - main ICD10: M48 Other spondylopathies | -0.0047 | 0.9379 | 0.114 | 0.0367 | -1.4684 | 0.231197 |
| 20517 | Trouble falling or staying asleep, or sleeping too much | 0.0777 | 0.0237 | 0.1953 | 1.06E-06 | -2.2291 | 0.233276 |
| 3090 | Used an inhaler for chest within last hour | 0.0386 | 0.6176 | 0.1551 | 0.0364 | -1.0872 | 0.235366 |
| 20002_1294 | Non-cancer illness code, self-reported: back problem | 0.0343 | 0.6468 | 0.1499 | 0.0363 | -1.1164 | 0.237083 |
| G6_SLEEPAP NO | Sleep apnoea | 0.081 | 0.2046 | 0.1953 | 0.0144 | -1.1172 | 0.239576 |
| COPD_EARL YANDLATER | COPD, early/later onset | 0.078 | 0.1146 | 0.1918 | 0.001 | -1.4922 | 0.240538 |
| 1200 | Sleeplessness / insomnia | 0.0023 | 0.9273 | 0.1158 | 1.15E-05 | -3.0864 | 0.241117 |
| N32 | Diagnoses - main ICD10: N32 Other disorders of bladder | 0.0675 | 0.3367 | 0.1803 | 0.0123 | -1.1217 | 0.242469 |
| 41248_1001 | Destinations on discharge from hospital (recoded): Usual Place of residence: Living with relatives | 0.0485 | 0.4219 | 0.1611 | 0.0275 | -1.1875 | 0.242857 |
| 4101 | Heel broadband ultrasound attenuation (left) | -0.0209 | 0.3713 | 0.0916 | 0.001 | -3.1015 | 0.24305 |
| 20003_99999 | Treatment/medication code: Free-text entry, unable to be coded | 0.158 | 0.0493 | 0.2693 | 0.0096 | -0.8476 | 0.245382 |
| PULMONAR YDG | Other pulmonary diagnosis | 0.0316 | 0.435 | 0.1427 | 0.0018 | -1.8194 | 0.245772 |
| X_RESPIRAT ORY | Diseases of the respiratory system | 0.0316 | 0.435 | 0.1427 | 0.0018 | -1.8194 | 0.245772 |
| J44 | Diagnoses - main ICD10: J44 Other chronic obstructive pulmonary disease | 0.0719 | 0.1448 | 0.1809 | 0.0016 | -1.4434 | 0.249884 |
| 1787 | Maternal smoking around birth | 0.0973 | 0.001 | 0.2059 | 5.96E-12 | -2.5855 | 0.250672 |
| 2634 | Duration of heavy DIY | -0.0103 | 0.7944 | 0.0981 | 0.021 | -1.8661 | 0.251066 |
| 6159_3 | Pain type(s) experienced in last month: Neck or shoulder pain | 0.0972 | 0.0014 | 0.205 | 1.14E-12 | -2.5698 | 0.25225 |
| 4501 | Non-accidental death in close genetic family | 0.0322 | 0.6115 | 0.139 | 0.0306 | -1.1827 | 0.254231 |
| 30130 | Monocyte count | -0.0163 | 0.413 | 0.0904 | 1.72E-05 | -3.6881 | 0.25443 |
| M13_SPINST ENOSIS | Spinal stenosis | -0.0002 | 0.9975 | 0.1042 | 0.0478 | -1.3222 | 0.259018 |
| 20511 | Recent poor appetite or overeating | 0.1475 | 0.0002 | 0.2516 | 1.1E-09 | -1.8304 | 0.25962 |
| 20003_114117 6832 | Treatment/medication code: seretide 50 evohaler | 0.0142 | 0.7487 | 0.1175 | 0.0065 | -1.6713 | 0.261228 |
| 20110_2 | Illnesses of mother: Stroke | 0.0784 | 0.2727 | 0.1807 | 0.0051 | -1.0624 | 0.263245 |
| 20110_8 | Illnesses of mother: High blood pressure | -0.0124 | 0.7173 | 0.0898 | 0.0067 | -2.1409 | 0.263447 |
| 1150_1 | Usual side of head for mobile phone use: Left | -0.0144 | 0.7446 | 0.0877 | 0.0249 | -1.7323 | 0.26365 |
| 2316 | Wheeze or whistling in the chest in last year | 0.0976 | 6.22E-05 | 0.1993 | 1.16E-13 | -2.8003 | 0.264459 |
| 30070 | Red blood cell (erythrocyte) distribution width | -0.019 | 0.3648 | 0.0827 | 0.0001 | -3.3999 | 0.264459 |
| 2814 | Ever used hormone-replacement therapy (HRT) | 0.0168 | 0.5818 | 0.1174 | 0.0002 | -2.3058 | 0.266692 |
| 6145_3 | Illness, injury, bereavement, stress in last 2 years: Death of a close relative | 0.0242 | 0.6927 | 0.1246 | 0.0319 | -1.1887 | 0.267099 |
| 2178 | Overall health rating | 0.1389 | 2.57E-10 | 0.2386 | 4.13E-25 | -3.1254 | 0.268527 |
| 22606_2 | Workplace very noisy: Often | 0.113 | 0.0126 | 0.2124 | 8.53E-06 | -1.511 | 0.269139 |
| G6_EPIPARO X | Episodal and paroxysmal disorders | 0.0902 | 0.1332 | 0.1893 | 0.0033 | -1.1241 | 0.269753 |
| 20003_114119 1044 | Treatment/medication code: levothyroxine sodium | -0.0209 | 0.544 | 0.0778 | 0.0138 | -2.113 | 0.270572 |
| 20002_1197 | Non-cancer illness code, self-reported: kidney stone/ureter stone/bladder stone | 0.0082 | 0.8774 | 0.1062 | 0.0376 | -1.3285 | 0.272008 |
| R55 | Diagnoses - main ICD10: R55 Syncope and collapse | 0.0377 | 0.4761 | 0.1355 | 0.0296 | -1.1966 | 0.27242 |
| 102 | Pulse rate, automated reading | -0.0067 | 0.7501 | 0.0903 | 4.15E-06 | -3.3596 | 0.274067 |
| 6149_4 | Mouth/teeth dental problems: Loose teeth | 0.1739 | 4.19E-05 | 0.2695 | 3.53E-09 | -1.5337 | 0.276962 |
| 1140 | Difference in mobile phone use compared to two years previously | -0.1679 | 0.0006 | -0.0725 | 0.1468 | -1.36 | 0.277377 |
| 30000 | White blood cell (leukocyte) count | 0.0105 | 0.6178 | 0.1057 | 1.79E-07 | -3.2672 | 0.277792 |
| 24005 | Particulate matter air pollution (pm10); 2010 | 0.0974 | 0.2057 | 0.1925 | 0.0224 | -0.8329 | 0.277999 |

| | | | | | | | |
|---|---|---|---|---|---|---|---|
| 6149_5 | Mouth/teeth dental problems: Toothache | 0.1345 | 0.0103 | 0.2292 | 7.62E-05 | -1.2127 | 0.278831 |
| 1687 | Comparative body size at age 10 | -0.0869 | 6.57E-05 | 0.0072 | 0.7332 | -3.1016 | 0.28008 |
| 1930 | Miserableness | 0.0659 | 0.0194 | 0.1597 | 5.65E-09 | -2.3856 | 0.280706 |
| M13_HALLUXVALGUS | Hallux valgus (acquired) | -0.0847 | 0.0337 | 0.0087 | 0.8413 | -1.5882 | 0.281541 |
| 20459 | General happiness with own health | 0.1334 | 5.93E-05 | 0.2267 | 1.13E-11 | -1.9812 | 0.28175 |
| 20519 | Recent feelings of tiredness or low energy | 0.1539 | 6.74E-05 | 0.2464 | 1.06E-11 | -1.7457 | 0.283425 |
| 4120 | Heel broadband ultrasound attenuation (right) | -0.0207 | 0.4038 | 0.0717 | 0.0091 | -2.4952 | 0.283634 |
| 816 | Job involves heavy manual or physical work | -0.0195 | 0.4874 | 0.0707 | 0.0122 | -2.2658 | 0.288266 |
| 30140 | Neutrophill count | -0.0125 | 0.5578 | 0.0766 | 0.0002 | -2.9926 | 0.290595 |
| 6145_1 | Illness, injury, bereavement, stress in last 2 years: Serious illness, injury or assault to yourself | 0.1431 | 0.0007 | 0.2321 | 2.01E-08 | -1.5091 | 0.290807 |
| XI_DIGESTIVE | Diseases of the digestive system | 0.0719 | 0.0201 | 0.1607 | 6.8E-08 | -2.0686 | 0.291232 |
| 1269 | Exposure to tobacco smoke at home | 0.1928 | 0.0002 | 0.2795 | 2.18E-08 | -1.2163 | 0.295707 |
| 20002_1385 | Non-cancer illness code, self-reported: allergy or anaphylactic reaction to food | 0.2156 | 0.0449 | 0.3022 | 0.085 | -0.4208 | 0.295921 |
| SLEEP | Sleep disorders (combined) | 0.0745 | 0.2019 | 0.161 | 0.0125 | -0.9941 | 0.296135 |
| 20107_3 | Illnesses of father: Lung cancer | 0.0443 | 0.2939 | 0.1302 | 0.001 | -1.4843 | 0.29742 |
| 4548 | Health satisfaction | 0.1434 | 2.51E-06 | 0.2292 | 1.01E-10 | -1.8332 | 0.297635 |
| G47 | Diagnoses - main ICD10: G47 Sleep disorders | 0.0752 | 0.2054 | 0.1607 | 0.0141 | -0.9677 | 0.298278 |
| 680_3 | Own or rent accommodation lived in: Rent - from local authority, local council, housing association | 0.1399 | 1.21E-06 | 0.2248 | 6.63E-13 | -1.9961 | 0.299568 |
| M20 | Diagnoses - main ICD10: M20 Acquired deformities of fingers and toes | -0.083 | 0.0339 | 0.0016 | 0.9693 | -1.4744 | 0.300214 |
| 20002_1277 | Non-cancer illness code, self-reported: glaucoma | -0.096 | 0.0414 | -0.0115 | 0.8072 | -1.2686 | 0.300429 |
| 2395_4 | Hair/balding pattern: Pattern 4 | -0.0699 | 0.0017 | 0.0139 | 0.5963 | -2.4403 | 0.301939 |
| 1518 | Hot drink temperature | 0.0245 | 0.2957 | 0.1081 | 9.65E-06 | -2.4678 | 0.30237 |
| 3571 | Back pain for 3+ months | 0.0388 | 0.3975 | 0.1211 | 0.0086 | -1.2665 | 0.305185 |
| 30500 | Microalbumin in urine | 0.0242 | 0.6291 | 0.1056 | 0.0252 | -1.1813 | 0.30714 |
| 20405_0 | Ever had known person concerned about, or recommend reduction of, alcohol consumption: No | -0.1597 | 0.002 | -0.0791 | 0.119 | -1.1131 | 0.308882 |
| 3606 | Chest pain or discomfort walking normally | 0.04 | 0.3619 | 0.1203 | 0.0169 | -1.2014 | 0.309537 |
| 20003_1140861998 | Treatment/medication code: ventolin 100micrograms inhaler | 0.0256 | 0.477 | 0.1058 | 0.0073 | -1.5046 | 0.309755 |
| XIX_INJURY_POISON | Injury, poisoning and certain other consequences of external causes | 0.1188 | 0.0024 | 0.1989 | 1.04E-07 | -1.4804 | 0.309974 |
| 4194 | Pulse rate | -0.0062 | 0.8228 | 0.0722 | 0.0078 | -2.0232 | 0.313696 |
| 20002_1226 | Non-cancer illness code, self-reported: hypothyroidism/myxoedema | -0.0123 | 0.6851 | 0.0659 | 0.0312 | -1.8189 | 0.314136 |
| 20548_2 | Manifestations of mania or irritability: I was more restless than usual | 0.14 | 0.0018 | 0.218 | 4.61E-06 | -1.1908 | 0.314575 |
| 3144 | Heel Broadband ultrasound attenuation, direct entry | -0.0048 | 0.8466 | 0.0723 | 0.0061 | -2.1326 | 0.316556 |
| 6160_5 | Leisure/social activities: Other group activity | 0.0234 | 0.523 | 0.1001 | 0.0071 | -1.4697 | 0.317438 |
| 6145_6 | Illness, injury, bereavement, stress in last 2 years: Financial difficulties | 0.1577 | 8.88E-07 | 0.233 | 6.07E-15 | -1.7165 | 0.320534 |
| 30010 | Red blood cell (erythrocyte) count | -0.0147 | 0.4544 | 0.0604 | 0.0076 | -2.5104 | 0.320977 |
| 22615_2 | Workplace had a lot of diesel exhaust: Often | 0.1293 | 0.0561 | 0.2037 | 0.0203 | -0.6715 | 0.32253 |
| 4104 | Heel quantitative ultrasound index (QUI), direct entry (left) | -0.0183 | 0.4076 | 0.055 | 0.0335 | -2.1529 | 0.324978 |
| 4106 | Heel bone mineral density (BMD) T-score, automated (left) | -0.0183 | 0.4077 | 0.055 | 0.0336 | -2.1529 | 0.324978 |
| 22128 | Doctor diagnosed emphysema | 0.1466 | 0.2151 | 0.2197 | 0.037 | -0.4618 | 0.325423 |
| 6179_2 | Mineral and other dietary supplements: Glucosamine | -0.1313 | 0.0003 | -0.0584 | 0.0813 | -1.4715 | 0.325869 |
| 4105 | Heel bone mineral density (BMD) (left) | -0.018 | 0.4148 | 0.0543 | 0.0343 | -2.133 | 0.327209 |
| 20003_1140884464 | Treatment/medication code: dihydrocodeine | 0.1421 | 0.0784 | 0.2143 | 0.0238 | -0.5799 | 0.327433 |
| 47 | Hand grip strength (right) | -0.0593 | 0.0049 | 0.0128 | 0.5289 | -2.4566 | 0.327656 |

| | | | | | | | |
|---|---|---|---|---|---|---|---|
| COLITNONINFNAS | Noninfectious colitis NAS | 0.0662 | 0.2912 | 0.1383 | 0.0273 | -0.8131 | 0.327656 |
| 22619_0 | Breathing problems responsible for leaving job: No | 0.1097 | 0.0252 | 0.1808 | 0.0006 | -0.9911 | 0.329895 |
| 20003_1140909674 | Treatment/medication code: cod liver oil capsule | -0.1981 | 0.0054 | -0.1284 | 0.0527 | -0.7164 | 0.33304 |
| M13_SPONDYLOPATHY | #Spondylopathies | 0.0766 | 0.1848 | 0.1458 | 0.0159 | -0.8277 | 0.334165 |
| K52 | Diagnoses - main ICD10: K52 Other non-infective gastro-enteritis and colitis | 0.0724 | 0.2535 | 0.1414 | 0.0203 | -0.7849 | 0.334616 |
| 22660_106 | Gap coding: Unable to work due to sickness or disability | 0.2237 | 0.0106 | 0.2922 | 0.0013 | -0.5423 | 0.335744 |
| 1747_2 | Hair colour (natural, before greying): Red | -0.049 | 0.0201 | 0.0194 | 0.4595 | -2.0333 | 0.33597 |
| 6159_1 | Pain type(s) experienced in last month: Headache | 0.0794 | 0.0079 | 0.1468 | 2.86E-07 | -1.629 | 0.338231 |
| 1050 | Time spend outdoors in summer | -0.0549 | 0.03 | 0.0121 | 0.6515 | -1.8215 | 0.339137 |
| 20002_1112 | Non-cancer illness code, self-reported: chronic obstructive airways disease/copd | 0.0658 | 0.2682 | 0.1316 | 0.0289 | -0.778 | 0.341861 |
| 1239 | Current tobacco smoking | 0.1217 | 9.11E-06 | 0.1849 | 1.14E-11 | -1.637 | 0.347791 |
| 1608 | Average weekly fortified wine intake | -0.1297 | 0.0137 | -0.067 | 0.2829 | -0.7683 | 0.348935 |
| 20411_0 | Ever been injured or injured someone else through drinking alcohol: No | -0.2329 | 0.0177 | -0.1703 | 0.0825 | -0.451 | 0.349165 |
| 1468_2 | Cereal type: Biscuit cereal (e.g. Weetabix) | 0.0519 | 0.358 | 0.1144 | 0.0283 | -0.8132 | 0.349394 |
| 1873 | Number of full brothers | 0.1723 | 0.0001 | 0.2328 | 7.96E-08 | -0.9733 | 0.353988 |
| 1538_1 | Major dietary changes in the last 5 years: Yes, because of illness | 0.1386 | 2.31E-05 | 0.1988 | 6.85E-09 | -1.2703 | 0.354679 |
| 20002_1453 | Non-cancer illness code, self-reported: psoriasis | 0.1018 | 0.0719 | 0.1614 | 0.007 | -0.7244 | 0.356063 |
| 22616_1 | Breathing problems during period of job: Yes | 0.1265 | 0.0182 | 0.1861 | 0.0004 | -0.7921 | 0.356063 |
| 30090 | Platelet crit | -0.0071 | 0.7304 | 0.0524 | 0.0076 | -2.0979 | 0.356293 |
| 20002_1111 | Non-cancer illness code, self-reported: asthma | 0.0412 | 0.0902 | 0.0962 | 0.0002 | -1.5391 | 0.366731 |
| 20111_6 | Illnesses of siblings: Chronic bronchitis/emphysema | 0.1675 | 0.0244 | 0.2223 | 0.0022 | -0.5275 | 0.367198 |
| 30120 | Lymphocyte count | 0.0504 | 0.0097 | 0.1042 | 1.12E-06 | -1.8583 | 0.369532 |
| 30030 | Haematocrit percentage | -0.0008 | 0.9676 | 0.0529 | 0.0224 | -1.7345 | 0.369766 |
| 20107_6 | Illnesses of father: Chronic bronchitis/emphysema | 0.1074 | 0.0079 | 0.16 | 0.0005 | -0.8571 | 0.372339 |
| 6152_8 | Blood clot, DVT, bronchitis, emphysema, asthma, rhinitis, eczema, allergy diagnosed by doctor: Asthma | 0.0441 | 0.0733 | 0.0967 | 0.0002 | -1.4636 | 0.372339 |
| 5264 | Corneal hysteresis (left) | 0.0113 | 0.6969 | 0.0598 | 0.0442 | -1.1664 | 0.381982 |
| 3456 | Number of cigarettes currently smoked daily (current cigarette smokers) | 0.1147 | 0.0355 | 0.1627 | 0.0066 | -0.5922 | 0.383163 |
| 22612_1 | Worked with materials containing asbestos: Sometimes | 0.1401 | 0.0208 | 0.1875 | 0.0063 | -0.5178 | 0.384582 |
| 20513 | Recent thoughts of suicide or self-harm | 0.1972 | 0.004 | 0.2438 | 0.0005 | -0.4762 | 0.386476 |
| 22620_1 | Job involved shift work: Yes | 0.1341 | 0.0096 | 0.1796 | 0.0011 | -0.6011 | 0.389085 |
| 22613_1 | Worked with paints, thinners or glues: Sometimes | 0.1131 | 0.0332 | 0.1562 | 0.0117 | -0.528 | 0.394794 |
| 6164_4 | Types of physical activity in last 4 weeks: Light DIY (eg: pruning, watering the lawn) | -0.0803 | 0.008 | -0.0398 | 0.1371 | -1.0012 | 0.401004 |
| 20553_5 | Methods of self-harm used: Self-injury such as self-cutting, scratching or hitting, etc. | 0.2169 | 0.024 | 0.2572 | 0.0131 | -0.2852 | 0.401483 |
| 20514 | Recent lack of interest or pleasure in doing things | 0.1415 | 0.0017 | 0.1816 | 3.06E-05 | -0.64 | 0.401962 |
| 20510 | Recent feelings of depression | 0.1421 | 0.0005 | 0.1807 | 2.69E-05 | -0.6535 | 0.405559 |
| 6146_2 | Attendance/disability/mobility allowance: Disability living allowance | 0.1439 | 1.41E-05 | 0.178 | 2.01E-07 | -0.7165 | 0.416395 |
| 1259 | Smoking/smokers in household | 0.1859 | 0.0005 | 0.2194 | 7.79E-06 | -0.4623 | 0.417845 |
| 20460 | Belief that own life is meaningful | -0.1491 | 1.78E-05 | -0.1157 | 0.0021 | -0.6529 | 0.418087 |
| 22506_114 | Tobacco smoking: Never smoked | -0.1896 | 2.68E-07 | -0.1571 | 7.68E-05 | -0.5996 | 0.420264 |
| 189 | Townsend deprivation index at recruitment | 0.2093 | 7.88E-11 | 0.2385 | 1.48E-14 | -0.6533 | 0.428267 |
| 20116_2 | Smoking status: Current | 0.1317 | 1.87E-06 | 0.16 | 9.46E-09 | -0.7211 | 0.430456 |
| 6144_4 | Never eat eggs, dairy, wheat, sugar: Sugar or foods/drinks containing sugar | 0.056 | 0.0397 | 0.0841 | 0.0044 | -0.699 | 0.430942 |

| | | | | | | | |
|---|---|---|---|---|---|---|---|
| 20117_2 | Alcohol drinker status: Current | -0.0932 | 0.0093 | -0.0656 | 0.0814 | -0.5302 | 0.432159 |
| 680_4 | Own or rent accommodation lived in: Rent - from private landlord or letting agency | 0.1966 | 0.0013 | 0.2214 | 8.06E-05 | -0.2987 | 0.438984 |
| 22618_0 | Breathing problems improved/stopped away from workplace or on holiday: No | 0.1338 | 0.0531 | 0.1586 | 0.0243 | -0.2512 | 0.438984 |
| 20502 | Ever had period extreme irritability | 0.1861 | 7.41E-07 | 0.2094 | 3.04E-07 | -0.4194 | 0.442648 |
| 20417 | Tense, sore, or aching muscles during worst period of anxiety | 0.0981 | 0.0734 | 0.1198 | 0.0332 | -0.2762 | 0.446562 |
| 20116_1 | Smoking status: Previous | 0.1419 | 3.07E-07 | 0.163 | 5E-09 | -0.5367 | 0.448031 |
| 30200 | Neutrophill percentage | -0.0495 | 0.0107 | -0.0296 | 0.1969 | -0.663 | 0.450971 |
| R31 | Diagnoses - main ICD10: R31 Unspecified haematuria | 0.1039 | 0.0791 | 0.1231 | 0.034 | -0.2315 | 0.452688 |
| 4581 | Financial situation satisfaction | 0.1282 | 0.0003 | 0.1457 | 9.64E-05 | -0.3394 | 0.45686 |
| 20403 | Amount of alcohol drunk on a typical drinking day | 0.1004 | 0.005 | 0.1169 | 0.0084 | -0.2893 | 0.459316 |
| 6152_100 | Blood clot, DVT, bronchitis, emphysema, asthma, rhinitis, eczema, allergy diagnosed by doctor: None of the above | -0.0669 | 0.0048 | -0.0512 | 0.0253 | -0.4764 | 0.461282 |
| 24006 | Particulate matter air pollution (pm2.5); 2010 | 0.1489 | 0.0026 | 0.1637 | 0.0008 | -0.2129 | 0.463495 |
| 6162_2 | Types of transport used (excluding work): Walk | -0.0805 | 0.023 | -0.0682 | 0.0323 | -0.2585 | 0.469649 |
| 6144_5 | Never eat eggs, dairy, wheat, sugar: I eat all of the above | -0.0958 | 0.0012 | -0.0836 | 0.0048 | -0.2914 | 0.469895 |
| 845 | Age completed full time education | -0.0998 | 3.5E-05 | -0.0895 | 0.0006 | -0.2911 | 0.474576 |
| 6160_3 | Leisure/social activities: Religious group | -0.058 | 0.04 | -0.0483 | 0.1412 | -0.2242 | 0.476056 |
| 6157_1 | Why stopped smoking: Illness or ill health | 0.1628 | 0.0487 | 0.172 | 0.0387 | -0.0785 | 0.477288 |
| 20160 | Ever smoked | 0.1545 | 1.5E-10 | 0.159 | 1.88E-10 | -0.1296 | 0.488887 |
| 20416 | Frequency of consuming six or more units of alcohol | 0.0836 | 0.0195 | 0.0855 | 0.029 | -0.0358 | 0.495307 |
| 2267 | Use of sun/uv protection | -0.0701 | 0.0091 | -0.0689 | 0.007 | -0.0323 | 0.497036 |
| 3731 | Former alcohol drinker | 0.1489 | 0.0076 | 0.1436 | 0.0104 | 0.067 | 0.513089 |
| 20111_3 | Illnesses of siblings: Lung cancer | 0.2375 | 0.0094 | 0.231 | 0.0574 | 0.0427 | 0.516051 |
| 22607_2 | Workplace very cold: Often | 0.141 | 0.0078 | 0.132 | 0.0197 | 0.1161 | 0.522218 |
| 20508 | Recent trouble concentrating on things | 0.1817 | 0.0001 | 0.1692 | 0.0004 | 0.1863 | 0.530844 |
| 20490 | Sexually molested as a child | 0.264 | 1.51E-05 | 0.2511 | 1.35E-05 | 0.1536 | 0.531829 |
| 20110_6 | Illnesses of mother: Chronic bronchitis/emphysema | 0.1764 | 0.0002 | 0.1629 | 0.0002 | 0.2099 | 0.533306 |
| 1180 | Morning/evening person (chronotype) | 0.1075 | 5.76E-06 | 0.0889 | 0.0003 | 0.5445 | 0.54584 |
| 4653 | Ever highly irritable/argumentative for 2 days | 0.1494 | 0.0001 | 0.1241 | 0.0025 | 0.4487 | 0.562236 |
| 2080 | Frequency of tiredness / lethargy in last 2 weeks | 0.2158 | 2.48E-16 | 0.1895 | 2.51E-12 | 0.6964 | 0.564675 |
| 24004 | Nitrogen oxides air pollution; 2010 | 0.1482 | 0.0019 | 0.1215 | 0.0111 | 0.395 | 0.565649 |
| 20116_0 | Smoking status: Never | -0.1607 | 2.7E-12 | -0.1905 | 1.06E-16 | 0.9162 | 0.57319 |
| 30180 | Lymphocyte percentage | 0.0547 | 0.0036 | 0.0248 | 0.276 | 1.0118 | 0.573433 |
| 20463 | Ever heard an un-real voice | 0.3114 | 0.0005 | 0.2813 | 0.0062 | 0.2217 | 0.573918 |
| 2020 | Loneliness, isolation | 0.1733 | 1.93E-09 | 0.1406 | 4.35E-06 | 0.7769 | 0.58022 |
| 6159_2 | Pain type(s) experienced in last month: Facial pain | 0.2091 | 0.0012 | 0.1756 | 0.0032 | 0.3808 | 0.582155 |
| 3526 | Mother's age at death | -0.1215 | 0.0059 | -0.1556 | 0.003 | 0.4979 | 0.583605 |
| 20110_3 | Illnesses of mother: Lung cancer | 0.2264 | 0.0074 | 0.191 | 0.0257 | 0.294 | 0.586742 |
| 1940 | Irritability | 0.087 | 0.0059 | 0.0512 | 0.0748 | 0.8386 | 0.587707 |
| 1588 | Average weekly beer plus cider intake | 0.1017 | 0.0018 | 0.0652 | 0.0465 | 0.7905 | 0.589393 |
| 5992 | ECG, phase duration | -0.1359 | 0.0369 | -0.1727 | 0.0227 | 0.3683 | 0.590115 |
| 20084_473 | Vitamin and/or mineral supplement use: Glucosamine/chondroitin | -0.1587 | 0.1412 | -0.1956 | 0.0439 | 0.2542 | 0.590356 |
| 20419 | Difficulty concentrating during worst period of anxiety | 0.2641 | 0.0038 | 0.2271 | 0.035 | 0.2619 | 0.590596 |
| 22609_2 | Workplace very dusty: Often | 0.1725 | 0.0007 | 0.1337 | 0.0103 | 0.5322 | 0.594921 |
| 1220 | Daytime dozing / sleeping (narcolepsy) | 0.0834 | 0.0021 | 0.0437 | 0.1384 | 0.9911 | 0.59708 |
| 1249 | Past tobacco smoking | -0.1547 | 1.61E-10 | -0.1947 | 1.08E-15 | 1.1664 | 0.597799 |
| 22506_113 | Tobacco smoking: Ex-smoker | 0.1952 | 5.44E-07 | 0.1551 | 0.0003 | 0.6925 | 0.598038 |
| 22601_22113066 | Job coding: medical doctor, general practitioner, hospital consultant | -0.0905 | 0.1076 | -0.1306 | 0.0143 | 0.5177 | 0.598038 |
| 20429 | Easily tired during worst period of anxiety | 0.2454 | 0.0015 | 0.2039 | 0.005 | 0.3919 | 0.601387 |
| S09 | Diagnoses - main ICD10: S09 Other and unspecified injuries of head | 0.2468 | 0.0006 | 0.2048 | 0.0062 | 0.4053 | 0.602582 |
| 2257 | Hearing difficulty/problems with background noise | 0.213 | 3.4E-13 | 0.1685 | 2.91E-09 | 1.0906 | 0.608539 |

| | | | | | | | |
|---|---|---|---|---|---|---|---|
| 3063 | Forced expiratory volume in 1-second (FEV1) | -0.0407 | 0.0498 | -0.0858 | 0.0001 | 1.4861 | 0.609965 |
| 6149_2 | Mouth/teeth dental problems: Painful gums | 0.2027 | 0.0013 | 0.1572 | 0.0267 | 0.4797 | 0.610915 |
| 20531 | Victim of sexual assault | 0.271 | 3.12E-09 | 0.2246 | 2.01E-06 | 0.7055 | 0.61305 |
| 738 | Average total household income before tax | -0.0885 | 6.82E-05 | -0.1377 | 7.11E-10 | 1.5636 | 0.61967 |
| 2473 | Other serious medical condition/disability diagnosed by doctor | 0.2027 | 6.57E-09 | 0.1522 | 1.68E-05 | 1.0159 | 0.622731 |
| 6142_4 | Current employment status: Unable to work because of sickness or disability | 0.2262 | 3.54E-10 | 0.1757 | 2.51E-06 | 0.9729 | 0.622731 |
| 30050 | Mean corpuscular haemoglobin | 0.0115 | 0.5604 | -0.0392 | 0.0443 | 1.8291 | 0.623202 |
| 6152_9 | Blood clot, DVT, bronchitis, emphysema, asthma, rhinitis, eczema, allergy diagnosed by doctor: Hayfever, allergic rhinitis or eczema | 0.0492 | 0.0327 | -0.0024 | 0.9207 | 1.549 | 0.625316 |
| 3761 | Age hay fever, rhinitis or eczema diagnosed | 0.1046 | 0.0074 | 0.0513 | 0.1549 | 1.0016 | 0.629299 |
| 22617_2211 | Job SOC coding: Medical practitioners | -0.0802 | 0.162 | -0.1347 | 0.0103 | 0.7013 | 0.632103 |
| 1920 | Mood swings | 0.1225 | 1.25E-05 | 0.0671 | 0.0128 | 1.4243 | 0.634201 |
| 20117_1 | Alcohol drinker status: Previous | 0.1843 | 4.25E-05 | 0.1257 | 0.0031 | 0.9467 | 0.641627 |
| 1060 | Time spent outdoors in winter | -0.0238 | 0.4119 | -0.0832 | 0.0025 | 1.4863 | 0.643476 |
| 6149_3 | Mouth/teeth dental problems: Bleeding gums | 0.1723 | 9.07E-06 | 0.1105 | 0.0034 | 1.1409 | 0.649001 |
| 1418_2 | Milk type used: Semi-skimmed | -0.0792 | 0.0584 | -0.1416 | 0.0012 | 1.0331 | 0.650377 |
| 20154 | Forced expiratory volume in 1-second (FEV1), predicted percentage | -0.0469 | 0.06 | -0.1097 | 0.0002 | 1.6268 | 0.651294 |
| 20505 | Recent easy annoyance or irritability | 0.1884 | 5.62E-06 | 0.1244 | 0.0028 | 1.0879 | 0.654038 |
| 1031 | Frequency of friend/family visits | 0.0789 | 0.0054 | 0.0147 | 0.5946 | 1.6241 | 0.654494 |
| 30260 | Mean reticulocyte volume | 0.0414 | 0.043 | -0.023 | 0.2769 | 2.1891 | 0.654951 |
| 20150 | Forced expiratory volume in 1-second (FEV1), Best measure | -0.0227 | 0.2947 | -0.0888 | 8.69E-05 | 2.1097 | 0.658821 |
| 2139 | Age first had sexual intercourse | -0.0981 | 2.21E-05 | -0.1654 | 6.34E-13 | 2.0646 | 0.661542 |
| 20553_4 | Methods of self-harm used: Ingesting a medication in excess of the normal dose | 0.3361 | 8.27E-05 | 0.2652 | 0.0014 | 0.595 | 0.669657 |
| 20521 | Belittlement by partner or ex-partner as an adult | 0.2888 | 1.64E-13 | 0.2174 | 4.01E-07 | 1.2287 | 0.670777 |
| 22127 | Doctor diagnosed asthma | 0.0849 | 0.017 | 0.0135 | 0.7201 | 1.3751 | 0.670777 |
| 3436 | Age started smoking in current smokers | -0.0931 | 0.0614 | -0.1652 | 0.0013 | 1.0074 | 0.672344 |
| 4836 | Loud music exposure frequency | 0.2686 | 5.06E-08 | 0.196 | 0.0002 | 1.0091 | 0.673461 |
| 20485 | Ever contemplated self-harm | 0.2993 | 3.21E-11 | 0.2262 | 8.55E-08 | 1.1835 | 0.674577 |
| 4270 | Volume level set by participant (left) | -0.0427 | 0.4008 | -0.1163 | 0.0337 | 0.9841 | 0.675691 |
| 20544_11 | Mental health problems ever diagnosed by a professional: Depression | 0.2349 | 2.69E-08 | 0.1612 | 9.23E-05 | 1.2496 | 0.675913 |
| 20002_99999 | Non-cancer illness code, self-reported: unclassifiable | 0.3636 | 0.0053 | 0.2898 | 0.0085 | 0.4321 | 0.676136 |
| 20480 | Ever self-harmed | 0.2941 | 8.43E-07 | 0.2197 | 0.0004 | 0.8673 | 0.67747 |
| 981 | Duration walking for pleasure | -0.0299 | 0.4452 | -0.1046 | 0.0105 | 1.3186 | 0.678136 |
| 20405_2 | Ever had known person concerned about, or recommend reduction of, alcohol consumption: Yes, during the last year | 0.1637 | 0.0045 | 0.0888 | 0.1555 | 0.8812 | 0.67858 |
| 20532 | Did your sleep change? | 0.2156 | 0.0103 | 0.1363 | 0.0645 | 0.7096 | 0.688277 |
| 2060 | Frequency of unenthusiasm / disinterest in last 2 weeks | 0.2466 | 1.31E-15 | 0.166 | 6.14E-08 | 1.8504 | 0.691118 |
| K40 | Diagnoses - main ICD10: K40 Inguinal hernia | -0.0085 | 0.8082 | -0.0904 | 0.0073 | 1.6881 | 0.693947 |
| 1190 | Nap during day | 0.1502 | 1.2E-10 | 0.066 | 0.0061 | 2.5118 | 0.698924 |
| 1408 | Cheese intake | -0.0063 | 0.8286 | -0.0912 | 0.0006 | 2.149 | 0.700432 |
| C_SKIN | NA | 0.0161 | 0.6521 | -0.0708 | 0.0297 | 1.7975 | 0.704721 |
| C3_SKIN | Malignant neoplasm of skin | 0.0161 | 0.6521 | -0.0708 | 0.0297 | 1.7975 | 0.704721 |
| 20536_0 | Weight change during worst episode of depression: Stayed about the same or was on a diet | -0.1122 | 0.0155 | -0.2001 | 0.0001 | 1.2652 | 0.706854 |
| 6159_8 | Pain type(s) experienced in last month: Pain all over the body | 0.1757 | 0.0001 | 0.0871 | 0.0751 | 1.325 | 0.708343 |
| 4277 | Volume level set by participant (right) | -0.0323 | 0.5492 | -0.1225 | 0.0174 | 1.21 | 0.711734 |
| 20479 | Ever thought that life not worth living | 0.2647 | 2.96E-13 | 0.1741 | 8.12E-06 | 1.7005 | 0.712579 |
| 6146_100 | Attendance/disability/mobility allowance: None of the above | -0.0992 | 0.0014 | -0.1934 | 7.67E-09 | 2.0608 | 0.720128 |
| 6159_5 | Pain type(s) experienced in last month: Stomach or abdominal pain | 0.2667 | 1.17E-11 | 0.1713 | 4.34E-06 | 1.7607 | 0.722623 |

| | | | | | | | |
|---|---|---|---|---|---|---|---|
| 3062 | Forced vital capacity (FVC) | -0.0091 | 0.6457 | -0.105 | 2.3E-07 | 3.3735 | 0.72366 |
| 1289 | Cooked vegetable intake | 0.0924 | 0.0028 | -0.0046 | 0.8799 | 2.2377 | 0.725933 |
| 4803_0 | Tinnitus: No, never | -0.1201 | 0.0004 | -0.2175 | 1.2E-10 | 2.0256 | 0.726757 |
| 1618 | Alcohol usually taken with meals | -0.0658 | 0.0164 | -0.1659 | 4.83E-10 | 2.6212 | 0.732289 |
| 1578 | Average weekly champagne plus white wine intake | -0.0707 | 0.0585 | -0.1715 | 1.67E-05 | 1.848 | 0.733714 |
| 20524 | Sexual interference by partner or ex-partner without consent as an adult | 0.2753 | 4.21E-05 | 0.1712 | 0.0138 | 1.0768 | 0.74038 |
| 6145_100 | Illness, injury, bereavement, stress in last 2 years: None of the above | -0.1248 | 0.0012 | -0.2294 | 3.06E-11 | 2.0263 | 0.741382 |
| 22130 | Doctor diagnosed COPD (chronic obstructive pulmonary disease) | 0.3128 | 0.0161 | 0.2065 | 0.1103 | 0.5798 | 0.744775 |
| 5993 | ECG, number of stages in a phase | -0.0688 | 0.237 | -0.1761 | 0.0108 | 1.1877 | 0.74676 |
| 22610_2 | Workplace full of chemical or other fumes: Often | 0.2795 | 0.0009 | 0.1722 | 0.0386 | 0.907 | 0.74676 |
| 30270 | Mean sphered cell volume | 0.0536 | 0.0072 | -0.0537 | 0.0144 | 3.6089 | 0.74676 |
| 6139_100 | Gas or solid-fuel cooking/heating: None of the above | 0.1697 | 0.0164 | 0.0617 | 0.36 | 1.1057 | 0.748145 |
| 6157_2 | Why stopped smoking: Doctor's advice | 0.1661 | 0.0295 | 0.0577 | 0.4132 | 1.0428 | 0.748934 |
| 1458 | Cereal intake | -0.0398 | 0.1291 | -0.1522 | 4.32E-09 | 3.051 | 0.756756 |
| 2664_1 | Reason for reducing amount of alcohol drunk: Illness or ill health | 0.2455 | 0.0007 | 0.1328 | 0.0967 | 1.0465 | 0.757337 |
| 20107_100 | Illnesses of father: None of the above (group 1) | 0.0034 | 0.9358 | -0.1099 | 0.0151 | 1.8262 | 0.758497 |
| 6162_3 | Types of transport used (excluding work): Public transport | 0.1208 | 0.0004 | 0.0063 | 0.8599 | 2.3091 | 0.760809 |
| 1807 | Father's age at death | -0.0683 | 0.063 | -0.1837 | 4.48E-06 | 2.1232 | 0.762535 |
| 20151 | Forced vital capacity (FVC), Best measure | 0.0129 | 0.538 | -0.1072 | 9.58E-08 | 4.1418 | 0.771434 |
| 100580 | Alcohol consumed | -0.0363 | 0.6051 | -0.1586 | 0.0448 | 1.1572 | 0.775535 |
| 2644 | Light smokers, at least 100 smokes in lifetime | 0.1042 | 0.0117 | -0.0197 | 0.6008 | 2.2184 | 0.778491 |
| 20544_6 | Mental health problems ever diagnosed by a professional: Panic attacks | 0.154 | 0.0048 | 0.0299 | 0.5969 | 1.5795 | 0.778858 |
| 20435 | Difficulty concentrating during worst depression | 0.2077 | 0.0044 | 0.0792 | 0.2552 | 1.2749 | 0.786863 |
| 1528 | Water intake | 0.0227 | 0.4214 | -0.1068 | 1.07E-05 | 3.4717 | 0.788659 |
| 20546_3 | Substances taken for depression: Medication prescribed to you (for at least two weeks) | 0.24 | 1.73E-09 | 0.1103 | 0.006 | 2.2928 | 0.789017 |
| 2159 | Ever had same-sex intercourse | 0.1899 | 4.65E-05 | 0.0588 | 0.2424 | 1.912 | 0.791513 |
| 3404 | Neck/shoulder pain for 3+ months | 0.2021 | 0.0095 | 0.0708 | 0.3249 | 1.2378 | 0.791868 |
| 20003_1193 | Treatment/medication code: omega-3/fish oil supplement | -0.0432 | 0.5846 | -0.1752 | 0.0318 | 1.1615 | 0.793109 |
| 6141_2 | How are people in household related to participant: Son and/or daughter (include step-children) | -0.0085 | 0.8951 | -0.1409 | 0.0199 | 1.4972 | 0.793815 |
| 20543 | Number of things worried about during worst period of anxiety | 0.2033 | 0.0081 | 0.0683 | 0.4011 | 1.2071 | 0.798374 |
| 20516 | Recent restlessness | 0.1839 | 0.0003 | 0.0487 | 0.2619 | 2.0212 | 0.798723 |
| T39 | Diagnoses - main ICD10: T39 Poisoning by nonopioid analgesics, antipyretics and antirheumatics | 0.2155 | 0.0329 | 0.0798 | 0.3797 | 0.9992 | 0.799592 |
| 104670 | Vitamin supplement user | -0.0215 | 0.7796 | -0.1581 | 0.031 | 1.2858 | 0.80115 |
| 20488 | Physically abused by family as a child | 0.2859 | 3.98E-15 | 0.1487 | 0.0002 | 2.5648 | 0.802185 |
| I9_DISVEINLYMPH | Diseases of veins, lymphatic vessels and lymph nodes, not elsewhere classified | 0.0577 | 0.1423 | -0.0799 | 0.0229 | 2.6114 | 0.802873 |
| 6153_4 | Medication for cholesterol, blood pressure, diabetes, or take exogenous hormones: Hormone replacement therapy | 0.1542 | 0.0024 | 0.0158 | 0.7754 | 1.8411 | 0.804244 |
| 20003_1140863202 | Treatment/medication code: temazepam | 0.1986 | 0.0368 | 0.0596 | 0.4853 | 1.0875 | 0.805269 |
| 20523 | Physical violence by partner or ex-partner as an adult | 0.3314 | 1.6E-11 | 0.1922 | 5.02E-05 | 2.0375 | 0.80561 |
| 1950 | Sensitivity / hurt feelings | 0.1543 | 5.07E-08 | 0.0146 | 0.6126 | 3.4599 | 0.806461 |
| 6159_100 | Pain type(s) experienced in last month: None of the above | -0.0868 | 0.0005 | -0.2268 | 3.59E-20 | 3.9916 | 0.80697 |
| 20512 | Recent feelings of foreboding | 0.2421 | 5.35E-09 | 0.1009 | 0.0185 | 2.3656 | 0.809 |
| 20002_1225 | Non-cancer illness code, self-reported: hyperthyroidism/thyrotoxicosis | 0.1358 | 0.0366 | -0.0057 | 0.9226 | 1.622 | 0.809505 |
| 2000 | Worry too long after embarrassment | 0.1397 | 1.65E-06 | -0.0026 | 0.9263 | 3.5056 | 0.810848 |

| | | | | | | | |
|---|---|---|---|---|---|---|---|
| 20001_1061 | Cancer code, self-reported: basal cell carcinoma | 0.112 | 0.0093 | -0.0303 | 0.5761 | 2.0549 | 0.810848 |
| 20450 | Feelings of worthlessness during worst period of depression | 0.2265 | 9.03E-05 | 0.0833 | 0.1704 | 1.707 | 0.812352 |
| K11_BARRET | Barret oesophagus | 0.1321 | 0.0389 | -0.0125 | 0.8314 | 1.6678 | 0.814678 |
| 22504 | Bring up phlegm/sputum/mucus on most days | 0.2719 | 1.23E-05 | 0.1271 | 0.0294 | 1.6972 | 0.815008 |
| 20002_1113 | Non-cancer illness code, self-reported: emphysema/chronic bronchitis | 0.2448 | 5.51E-08 | 0.0993 | 0.067 | 2.0635 | 0.816163 |
| 4609 | Longest period of depression | 0.3073 | 0.0026 | 0.1605 | 0.1208 | 1.0107 | 0.818295 |
| 1418_5 | Milk type used: Other type of milk | 0.2425 | 0.0359 | 0.0944 | 0.2742 | 1.0266 | 0.820412 |
| 1737 | Childhood sunburn occasions | 0.1155 | 0.0022 | -0.035 | 0.1606 | 3.3311 | 0.824279 |
| 6152_6 | Blood clot, DVT, bronchitis, emphysema, asthma, rhinitis, eczema, allergy diagnosed by doctor: Emphysema/chronic bronchitis | 0.2379 | 6.1E-10 | 0.0856 | 0.0796 | 2.4495 | 0.827144 |
| 6032 | Maximum workload during fitness test | -0.0267 | 0.5543 | -0.1803 | 0.0005 | 2.2294 | 0.829195 |
| 22611_2 | Workplace had a lot of cigarette smoke from other people smoking: Often | 0.2608 | 4.47E-06 | 0.1068 | 0.0972 | 1.7934 | 0.829823 |
| 2050 | Frequency of depressed mood in last 2 weeks | 0.2564 | 6.66E-15 | 0.0999 | 0.0005 | 3.5792 | 0.833713 |
| 6149_100 | Mouth/teeth dental problems: None of the above | -0.0787 | 0.0132 | -0.2358 | 1.15E-12 | 3.4173 | 0.834638 |
| 20534 | Sleeping too much | 0.2441 | 0.0008 | 0.0866 | 0.1847 | 1.6105 | 0.835253 |
| 3872 | Age of primiparous women at birth of child | -0.0197 | 0.7457 | -0.1779 | 0.0041 | 1.8218 | 0.836326 |
| 2040 | Risk taking | 0.2461 | 5.16E-23 | 0.0877 | 0.0019 | 4.2105 | 0.836631 |
| 2654_6 | Non-butter spread type details: Olive oil based spread (eg: Bertolli) | -0.0011 | 0.985 | -0.1614 | 0.0025 | 2.0333 | 0.839516 |
| 6144_1 | Never eat eggs, dairy, wheat, sugar: Eggs or foods containing eggs | 0.1532 | 0.0058 | -0.0092 | 0.8581 | 2.139 | 0.842665 |
| 2247_0 | Hearing difficulty/problems: No | -0.0604 | 0.027 | -0.2237 | 6.27E-15 | 4.1227 | 0.844002 |
| 24003 | Nitrogen dioxide air pollution; 2010 | 0.2148 | 1.39E-05 | 0.0492 | 0.3165 | 2.3776 | 0.847385 |
| 20002_1452 | Non-cancer illness code, self-reported: eczema/dermatitis | 0.1371 | 0.0348 | -0.0292 | 0.6193 | 1.8988 | 0.848405 |
| 20518 | Recent changes in speed/amount of moving or speaking | 0.2633 | 3.6E-06 | 0.0963 | 0.0563 | 2.1992 | 0.849421 |
| 6153_100 | Medication for cholesterol, blood pressure, diabetes, or take exogenous hormones: None of the above | 0.0282 | 0.3616 | -0.1404 | 1.13E-05 | 3.7901 | 0.851724 |
| 6150_100 | Vascular/heart problems diagnosed by doctor: None of the above | 0.016 | 0.474 | -0.1527 | 2.54E-11 | 5.2663 | 0.851868 |
| 1767 | Adopted as a child | 0.1972 | 0.0024 | 0.0254 | 0.6785 | 1.9243 | 0.856261 |
| K62 | Diagnoses - main ICD10: K62 Other diseases of anus and rectum | 0.1576 | 0.0164 | -0.0145 | 0.8551 | 1.6724 | 0.856681 |
| 20554_1 | Actions taken following self-harm: See anyone from psychiatric or mental health services, including liaison services | 0.3444 | 3.37E-05 | 0.1719 | 0.0267 | 1.5172 | 0.857241 |
| 20487 | Felt hated by family member as a child | 0.3385 | 3.19E-21 | 0.1658 | 9.43E-05 | 3.1079 | 0.85752 |
| 20420 | Longest period spent worried or anxious | 0.2539 | 0.0015 | 0.0811 | 0.2582 | 1.6107 | 0.857659 |
| 20127 | Neuroticism score | 0.1847 | 1.93E-10 | 0.0114 | 0.6644 | 4.4342 | 0.858355 |
| R04 | Diagnoses - main ICD10: R04 Haemorrhage from respiratory passages | 0.264 | 0.0024 | 0.0905 | 0.2456 | 1.4857 | 0.858632 |
| 6143_4 | Transport type for commuting to job workplace: Cycle | 0.0439 | 0.2638 | -0.13 | 0.0013 | 3.0894 | 0.859187 |
| 6154_100 | Medication for pain relief, constipation, heartburn: None of the above | 0.0033 | 0.9023 | -0.173 | 5.77E-12 | 4.7634 | 0.86248 |
| 20535 | Waking too early | 0.0426 | 0.5627 | -0.1356 | 0.0387 | 1.8075 | 0.86505 |
| 2030 | Guilty feelings | 0.2137 | 5.29E-13 | 0.0353 | 0.2063 | 4.3858 | 0.865319 |
| 1428_0 | Spread type: Never/rarely use spread | 0.1261 | 0.0002 | -0.0528 | 0.1439 | 3.6175 | 0.865989 |
| 6179_1 | Mineral and other dietary supplements: Fish oil (including cod liver oil) | -0.0196 | 0.5885 | -0.1989 | 1.5E-10 | 3.7621 | 0.866523 |
| 20494 | Felt irritable or had angry outbursts in past month | 0.3018 | 0.0001 | 0.121 | 0.0934 | 1.6963 | 0.868514 |
| 5984 | ECG, load | -0.0263 | 0.6 | -0.2071 | 0.0002 | 2.4086 | 0.868514 |
| 6160_1 | Leisure/social activities: Sports club or gym | -0.0162 | 0.5753 | -0.1982 | 4.46E-11 | 4.3544 | 0.870091 |
| 6038 | Number of trend entries | -0.016 | 0.7617 | -0.1982 | 0.0028 | 2.1509 | 0.870353 |
| 24017 | Nitrogen dioxide air pollution; 2006 | 0.1861 | 0.0002 | 0.002 | 0.9671 | 2.6756 | 0.87282 |

| | | | | | | | |
|---|---|---|---|---|---|---|---|
| 20110_100 | Illnesses of mother: None of the above (group 1) | -0.0115 | 0.7641 | -0.1958 | 1.05E-07 | 3.4699 | 0.873078 |
| 894 | Duration of moderate activity | 0.0825 | 0.0148 | -0.1023 | 0.0052 | 3.7094 | 0.873721 |
| 24019 | Particulate matter air pollution (pm10); 2007 | 0.1583 | 0.0006 | -0.0275 | 0.5028 | 3.0116 | 0.875 |
| 20003_1140921600 | Treatment/medication code: citalopram | 0.194 | 0.0103 | 0.0055 | 0.9344 | 1.8673 | 0.878408 |
| 6144_2 | Never eat eggs, dairy, wheat, sugar: Dairy products | 0.2934 | 0.0014 | 0.1049 | 0.2077 | 1.5187 | 0.878408 |
| 4196 | Pulse wave peak to peak time | 0.0502 | 0.2769 | -0.1385 | 0.0043 | 2.8171 | 0.878658 |
| 1568 | Average weekly red wine intake | 0.0366 | 0.197 | -0.1522 | 1.02E-06 | 4.4828 | 0.878783 |
| 943 | Frequency of stair climbing in last 4 weeks | -0.0372 | 0.2494 | -0.2261 | 7.17E-16 | 4.419 | 0.878907 |
| 12340 | QRS duration | 0.0533 | 0.4425 | -0.1369 | 0.0362 | 1.996 | 0.88052 |
| 20548_3 | Manifestations of mania or irritability: My thoughts were racing | 0.2879 | 1.94E-08 | 0.0975 | 0.0719 | 2.5513 | 0.880767 |
| S42 | Diagnoses - main ICD10: S42 Fracture of shoulder and upper arm | 0.2139 | 0.0224 | 0.023 | 0.7919 | 1.4906 | 0.881382 |
| 680_5 | Own or rent accommodation lived in: Pay part rent and part mortgage (shared ownership) | 0.2063 | 0.022 | 0.0148 | 0.8398 | 1.6487 | 0.882118 |
| 4803_14 | Tinnitus: Yes, but not now, but have in the past | 0.2419 | 6.23E-05 | 0.0497 | 0.3935 | 2.2895 | 0.882971 |
| 1011 | Frequency of light DIY in last 4 weeks | 0.1249 | 0.0058 | -0.0683 | 0.1516 | 2.9402 | 0.884184 |
| 3536 | Age started hormone-replacement therapy (HRT) | 0.067 | 0.1384 | -0.1277 | 0.0024 | 3.152 | 0.885985 |
| 3476 | Difficulty not smoking for 1 day | 0.202 | 7.78E-05 | 0.0062 | 0.9042 | 2.6988 | 0.887294 |
| 1309 | Fresh fruit intake | -0.0057 | 0.8228 | -0.2016 | 1.71E-15 | 5.4428 | 0.887412 |
| 24016 | Nitrogen dioxide air pollution; 2005 | 0.1924 | 0.0001 | -0.0043 | 0.9289 | 2.8323 | 0.888356 |
| 20507 | Recent feelings of inadequacy | 0.2829 | 2.69E-09 | 0.0854 | 0.0607 | 2.9993 | 0.889295 |
| 6143_3 | Transport type for commuting to job workplace: Public transport | 0.136 | 0.0013 | -0.063 | 0.1314 | 3.3503 | 0.891039 |
| 2754 | Age at first live birth | -0.0307 | 0.2374 | -0.2299 | 1.54E-18 | 5.3967 | 0.89127 |
| 4642 | Ever manic/hyper for 2 days | 0.3946 | 1.97E-08 | 0.1935 | 0.0024 | 2.1213 | 0.893448 |
| 1538_0 | Major dietary changes in the last 5 years: No | -0.0367 | 0.2338 | -0.2396 | 1.19E-14 | 4.6431 | 0.895482 |
| 23107 | Impedance of leg (right) | 0.0826 | 1.17E-05 | -0.1207 | 1.68E-08 | 7.1371 | 0.89593 |
| 6164_3 | Types of physical activity in last 4 weeks: Strenuous sports | 0.0182 | 0.6192 | -0.1859 | 7.15E-08 | 4.0638 | 0.896822 |
| 20437 | Thoughts of death during worst depression | 0.3443 | 0.0005 | 0.1391 | 0.0718 | 1.6386 | 0.89804 |
| 20453 | Ever taken cannabis | 0.2164 | 4.66E-10 | 0.0107 | 0.7579 | 4.1857 | 0.89859 |
| 20548_1 | Manifestations of mania or irritability: I was more talkative than usual | 0.3196 | 8.07E-09 | 0.1135 | 0.0374 | 2.652 | 0.899028 |
| 6156_15 | Manic/hyper symptoms: All of the above | 0.3822 | 0.005 | 0.1758 | 0.1384 | 1.1433 | 0.899356 |
| 24012 | Inverse distance to the nearest major road | 0.2818 | 0.0166 | 0.0752 | 0.419 | 1.3767 | 0.899574 |
| 4537 | Work/job satisfaction | 0.1689 | 0.0001 | -0.0388 | 0.3902 | 3.2818 | 0.900768 |
| 100008 | Total sugars | 0.0967 | 0.0412 | -0.1112 | 0.032 | 2.9578 | 0.900984 |
| 6158_1 | Why reduced smoking: Illness or ill health | 0.2761 | 0.0008 | 0.068 | 0.3341 | 1.9147 | 0.901199 |
| 2365 | Ever had prostate specific antigen (PSA) test | 0.0465 | 0.2067 | -0.1626 | 0.0006 | 3.4708 | 0.902272 |
| 100270 | Filtered coffee intake | 0.119 | 0.0357 | -0.091 | 0.1023 | 2.6421 | 0.90323 |
| 2664_2 | Reason for reducing amount of alcohol drunk: Doctor's advice | 0.1926 | 0.0335 | -0.0179 | 0.8546 | 1.5798 | 0.903759 |
| 20441 | Ever had prolonged loss of interest in normal activities | 0.3108 | 1.11E-17 | 0.0992 | 0.0067 | 4.1049 | 0.904916 |
| 24018 | Nitrogen dioxide air pollution; 2007 | 0.1927 | 7.55E-05 | -0.0192 | 0.6785 | 3.1534 | 0.90523 |
| 23108 | Impedance of leg (left) | 0.0837 | 9.09E-06 | -0.1286 | 3E-09 | 7.3775 | 0.905647 |
| 20526 | Been in serious accident believed to be life-threatening | 0.3381 | 4.14E-07 | 0.125 | 0.0563 | 2.2778 | 0.906478 |
| 680_1 | Own or rent accommodation lived in: Own outright (by you or someone in your household) | -0.0398 | 0.1745 | -0.2533 | 1.03E-15 | 4.9543 | 0.906891 |
| 20539 | Frequency of inability to stop worrying during worst period of anxiety | 0.1181 | 0.0308 | -0.0973 | 0.1103 | 2.629 | 0.908834 |
| 20515 | Recent trouble relaxing | 0.2312 | 5.18E-08 | 0.0136 | 0.7457 | 3.6417 | 0.911047 |
| 20529 | Victim of physically violent crime | 0.3159 | 2.13E-10 | 0.0966 | 0.064 | 3.0426 | 0.91273 |
| 2217 | Age started wearing glasses or contact lenses | 0.0964 | 0.0003 | -0.1231 | 2.12E-07 | 6.1354 | 0.912926 |
| 20500 | Ever suffered mental distress preventing usual activities | 0.3496 | 8.76E-20 | 0.13 | 0.0005 | 4.0914 | 0.913024 |

| | | | | | | | |
|---|---|---|---|---|---|---|---|
| 2654_2 | Non-butter spread type details: Flora Pro-Active or Benecol | 0.1432 | 0.0109 | -0.0764 | 0.1888 | 2.712 | 0.913024 |
| I83 | Diagnoses - main ICD10: I83 Varicose veins of lower extremities | 0.0877 | 0.023 | -0.132 | 4.96E-05 | 4.3539 | 0.913122 |
| 1468_4 | Cereal type: Muesli | 0.0586 | 0.0284 | -0.1612 | 1.73E-09 | 5.8102 | 0.91322 |
| 20552_2 | Behavioural and miscellaneous addictions: A behaviour | 0.4936 | 0.0353 | 0.2719 | 0.0281 | 0.8361 | 0.915064 |
| 100015 | Vitamin C | 0.0061 | 0.9088 | -0.2158 | 0.0004 | 2.7511 | 0.915257 |
| 670_4 | Type of accommodation lived in: Sheltered accommodation | 0.2594 | 0.0052 | 0.0374 | 0.6688 | 1.7396 | 0.915353 |
| 22617_2314 | Job SOC coding: Secondary education teaching professionals | 0.127 | 0.0426 | -0.0963 | 0.0908 | 2.6396 | 0.916594 |
| 6155_7 | Vitamin and mineral supplements: Multivitamins +/- minerals | 0.1326 | 0.0002 | -0.0916 | 0.0069 | 4.5473 | 0.917446 |
| 2345 | Ever had bowel cancer screening | 0.1547 | 0.0003 | -0.0703 | 0.1104 | 3.6696 | 0.918197 |
| 20003_1140876592 | Treatment/medication code: multivitamin+mineral preparations | 0.1615 | 0.0339 | -0.0635 | 0.4479 | 1.9878 | 0.918197 |
| 20084_472 | Vitamin and/or mineral supplement use: Fish oil | 0.0062 | 0.9471 | -0.2188 | 0.0049 | 1.851 | 0.918197 |
| 20509 | Recent inability to stop or control worrying | 0.2408 | 1.25E-09 | 0.014 | 0.742 | 3.9092 | 0.919869 |
| 6138_1 | Qualifications: College or University degree | 0.0897 | 7E-06 | -0.1377 | 1.59E-10 | 7.7442 | 0.920421 |
| 6177_100 | Medication for cholesterol, blood pressure or diabetes: None of the above | 0.0704 | 0.0136 | -0.157 | 4.86E-09 | 5.8019 | 0.920421 |
| 5100 | 3mm weak meridian angle (right) | 0.0885 | 0.1043 | -0.1393 | 0.0109 | 2.9502 | 0.920787 |
| 2149 | Lifetime number of sexual partners | 0.2754 | 1.9E-22 | 0.0473 | 0.0989 | 5.6692 | 0.921061 |
| 5115 | 3mm cylindrical power angle (right) | 0.0911 | 0.0967 | -0.1373 | 0.0122 | 2.9445 | 0.921334 |
| 22660_108 | Gap coding: Retired | -0.0023 | 0.9832 | -0.2309 | 0.0481 | 1.4278 | 0.921515 |
| 6143_2 | Transport type for commuting to job workplace: Walk | 0.1256 | 0.0084 | -0.1037 | 0.0312 | 3.3885 | 0.922148 |
| 20498 | Felt very upset when reminded of stressful experience in past month | 0.3637 | 1.59E-18 | 0.1343 | 0.0009 | 3.9657 | 0.922238 |
| 2227 | Other eye problems | 0.2765 | 6.15E-07 | 0.046 | 0.3606 | 3.0773 | 0.923225 |
| 100280 | Added milk to filtered coffee | 0.1038 | 0.0668 | -0.1269 | 0.0343 | 2.7969 | 0.923403 |
| 757 | Time employed in main current job | -0.0368 | 0.5611 | -0.2682 | 0.0004 | 2.3468 | 0.924024 |
| 6142_7 | Current employment status: Full or part-time student | 0.3144 | 0.0257 | 0.0826 | 0.5139 | 1.2237 | 0.924378 |
| 100012 | Vitamin B6 | 0.0668 | 0.3268 | -0.1651 | 0.0307 | 2.2659 | 0.924466 |
| 102800 | Cheese consumers | 0.0683 | 0.4057 | -0.1649 | 0.042 | 2.0208 | 0.925605 |
| 1707_2 | Handedness (chirality/laterality): Left-handed | 0.1475 | 0.0011 | -0.0868 | 0.0662 | 3.5812 | 0.926558 |
| 23106 | Impedance of whole body | 0.0932 | 9.72E-07 | -0.1411 | 3.25E-11 | 8.2087 | 0.926558 |
| 2867 | Age started smoking in former smokers | 0.0421 | 0.2127 | -0.1922 | 3.57E-06 | 4.3776 | 0.926558 |
| 20118_8 | Home area population density - urban or rural: England/Wales - Hamlet and Isolated Dwelling - less sparse | 0.0159 | 0.8186 | -0.2186 | 0.004 | 2.2815 | 0.926731 |
| 5855_1 | Which eye(s) affected by astigmatism: Right eye | 0.1887 | 0.003 | -0.0467 | 0.5272 | 2.4144 | 0.927502 |
| 23110 | Impedance of arm (left) | 0.0906 | 2.8E-06 | -0.145 | 8.63E-12 | 8.2178 | 0.927673 |
| 20493 | Severity of problems due to mania or irritability | 0.1303 | 0.0444 | -0.1057 | 0.1691 | 2.3486 | 0.928013 |
| 1448_3 | Bread type: Wholemeal or wholegrain | 0.0349 | 0.2352 | -0.2012 | 4.77E-13 | 5.8351 | 0.928098 |
| 2794 | Age started oral contraceptive pill | 0.0034 | 0.9166 | -0.234 | 1.48E-13 | 5.1962 | 0.929195 |
| 6145_5 | Illness, injury, bereavement, stress in last 2 years: Marital separation/divorce | 0.3316 | 0.0013 | 0.0928 | 0.2784 | 1.78 | 0.930362 |
| 20546_4 | Substances taken for depression: Drugs or alcohol (more than once) | 0.3292 | 1.66E-08 | 0.09 | 0.1098 | 2.9514 | 0.930692 |
| 20111_100 | Illnesses of siblings: None of the above (group 1) | 0.0199 | 0.5342 | -0.2203 | 2.98E-12 | 5.341 | 0.931513 |
| 22601_23143401 | Job coding: secondary school teacher or teaching professional (including head teacher) | 0.1352 | 0.0334 | -0.1065 | 0.0682 | 2.7992 | 0.932731 |
| 971 | Frequency of walking for pleasure in last 4 weeks | 0.1228 | 0.0003 | -0.1192 | 0.0003 | 5.1079 | 0.932972 |
| 4244 | Mean signal-to-noise ratio (SNR), (right) | 0.0897 | 0.2799 | -0.1526 | 0.0493 | 2.1324 | 0.933213 |
| 20401 | Ever addicted to any substance or behaviour | 0.4518 | 1.71E-09 | 0.2068 | 0.0052 | 2.3253 | 0.935351 |

| | | | | | | | |
|---|---|---|---|---|---|---|---|
| 6139_3 | Gas or solid-fuel cooking/heating: An open solid fuel fire that you use regularly in winter time | 0.0166 | 0.6594 | -0.2286 | 8.16E-09 | 4.473 | 0.935507 |
| 924 | Usual walking pace | 0.0127 | 0.5822 | -0.2332 | 1.38E-24 | 7.5762 | 0.936051 |
| 22501 | Year ended full time education | 0.0231 | 0.4467 | -0.2253 | 8.32E-14 | 5.8064 | 0.937967 |
| 2070 | Frequency of tenseness / restlessness in last 2 weeks | 0.2981 | 3.16E-23 | 0.0459 | 0.1359 | 5.8657 | 0.940792 |
| 6017 | Able to walk or cycle unaided for 10 minutes | -0.0884 | 0.2976 | -0.3417 | 0.0125 | 1.5733 | 0.94159 |
| 6164_2 | Types of physical activity in last 4 weeks: Other exercises (eg: swimming, cycling, keep fit, bowling) | 0.0387 | 0.1372 | -0.2154 | 1.56E-15 | 6.779 | 0.942166 |
| 20548_5 | Manifestations of mania or irritability: I needed less sleep than usual | 0.4385 | 7.32E-10 | 0.1824 | 0.0097 | 2.5559 | 0.943585 |
| 20003_1140916282 | Treatment/medication code: venlafaxine | 0.277 | 0.0004 | 0.0199 | 0.7618 | 2.5245 | 0.944284 |
| 23109 | Impedance of arm (right) | 0.1066 | 4.14E-08 | -0.1516 | 1.02E-12 | 8.962 | 0.945045 |
| 3637 | Frequency of other exercises in last 4 weeks | 0.0838 | 0.0804 | -0.1769 | 0.0005 | 3.7338 | 0.946745 |
| 20002_1408 | Non-cancer illness code, self-reported: alcohol dependency | 0.1994 | 0.0241 | -0.0624 | 0.4601 | 2.142 | 0.947479 |
| 5201 | logMAR, final (right) | 0.2643 | 2.23E-05 | 0.0019 | 0.975 | 3.0508 | 0.947877 |
| 3894 | Age heart attack diagnosed | -0.0325 | 0.7174 | -0.2951 | 0.0348 | 1.5809 | 0.948008 |
| age | Age at recruitment | 0.046 | 0.4179 | -0.2191 | 0.0008 | 3.0631 | 0.949635 |
| 20499 | Ever sought or received professional help for mental distress | 0.3157 | 6.83E-17 | 0.0495 | 0.2033 | 4.9078 | 0.950337 |
| 4570 | Friendships satisfaction | 0.31 | 2.15E-17 | 0.0427 | 0.2532 | 5.1149 | 0.951032 |
| 20439 | Frequency of depressed days during worst episode of depression | 0.3074 | 3.91E-05 | 0.0396 | 0.5451 | 2.6973 | 0.951345 |
| 20074 | Home location at assessment - east co-ordinate (rounded) | 0.1317 | 0.004 | -0.1365 | 0.0047 | 4.0293 | 0.951594 |
| 24007 | Particulate matter air pollution (pm2.5) absorbance; 2010 | 0.268 | 3.6E-05 | -0.0002 | 0.9967 | 3.1359 | 0.951594 |
| 2090 | Seen doctor (GP) for nerves, anxiety, tension or depression | 0.3066 | 3.96E-32 | 0.0381 | 0.1329 | 7.387 | 0.951781 |
| 20096_1 | Size of red wine glass drunk: small (125ml) | 0.207 | 0.0267 | -0.062 | 0.5272 | 1.9859 | 0.95209 |
| 22702 | Home location - east co-ordinate (rounded) | 0.1176 | 0.0104 | -0.1523 | 0.0015 | 4.0639 | 0.952643 |
| 5375 | Longest period of unenthusiasm / disinterest | 0.2577 | 0.0088 | -0.0145 | 0.8759 | 2.0125 | 0.954033 |
| 20458 | General happiness | 0.2956 | 7.13E-18 | 0.0221 | 0.5793 | 5.198 | 0.954803 |
| 22601_34123157 | Job coding: author, writer, biographer, book editor, novelist, dramatist, playwright, poet | 0.0644 | 0.429 | -0.2092 | 0.0295 | 2.1724 | 0.954862 |
| 6164_1 | Types of physical activity in last 4 weeks: Walking for pleasure (not as a means of transport) | -0.0279 | 0.3826 | -0.3017 | 6.69E-31 | 6.6429 | 0.95498 |
| 22617_2311 | Job SOC coding: Higher education teaching professionals | 0.1306 | 0.0199 | -0.1434 | 0.041 | 3.0491 | 0.955097 |
| 6179_5 | Mineral and other dietary supplements: Iron | 0.3132 | 1.77E-06 | 0.0386 | 0.5062 | 3.1363 | 0.955448 |
| 20520 | Recent worrying too much about different things | 0.2507 | 1.2E-08 | -0.0249 | 0.5794 | 4.379 | 0.956027 |
| 1299 | Salad / raw vegetable intake | 0.0754 | 0.0076 | -0.203 | 2.29E-11 | 6.714 | 0.957616 |
| 2129 | Answered sexual history questions | 0.1645 | 1.73E-06 | -0.1155 | 0.0007 | 5.7723 | 0.958503 |
| 2010 | Suffer from 'nerves' | 0.2016 | 3.52E-10 | -0.0796 | 0.0119 | 6.2331 | 0.959158 |
| 104400 | Fruit consumers | 0.0127 | 0.835 | -0.2692 | 0.0004 | 2.8918 | 0.959537 |
| 5208 | logMAR, final (left) | 0.2401 | 2.4E-05 | -0.0434 | 0.4317 | 3.5794 | 0.960391 |
| 20446 | Ever had prolonged feelings of sadness or depression | 0.3475 | 8.01E-17 | 0.062 | 0.1313 | 4.8762 | 0.961438 |
| 6142_5 | Current employment status: Unemployed | 0.2738 | 0.0003 | -0.0118 | 0.8634 | 2.7953 | 0.96149 |
| 20544_1 | Mental health problems ever diagnosed by a professional: Social anxiety or social phobia | 0.24 | 0.0074 | -0.0458 | 0.5527 | 2.4178 | 0.961593 |
| 1210 | Snoring | 0.1107 | 2.27E-07 | -0.1753 | 1.52E-12 | 8.731 | 0.961696 |
| 1438 | Bread intake | 0.1369 | 1.11E-07 | -0.1511 | 1.12E-07 | 7.4915 | 0.962715 |
| 6160_4 | Leisure/social activities: Adult education class | 0.1658 | 9.03E-06 | -0.1227 | 0.0012 | 5.4326 | 0.962966 |
| 4526 | Happiness | 0.3033 | 4.61E-17 | 0.013 | 0.7321 | 5.531 | 0.963859 |

| | | | | | | | |
|---|---|---|---|---|---|---|---|
| 20544_3 | Mental health problems ever diagnosed by a professional: Any other type of psychosis or psychotic illness | 0.2961 | 0.0045 | 0.0057 | 0.9533 | 2.0301 | 0.963908 |
| 3446_2 | Type of tobacco currently smoked: Hand-rolled cigarettes | 0.43 | 0.0054 | 0.1394 | 0.2231 | 1.5118 | 0.964006 |
| 6155_6 | Vitamin and mineral supplements: Folic acid or Folate (Vit B9) | 0.3254 | 0.0055 | 0.0347 | 0.7359 | 1.8647 | 0.964055 |
| 1319 | Dried fruit intake | 0.0858 | 0.0015 | -0.2052 | 4.65E-17 | 7.98 | 0.964202 |
| 20540 | Multiple worries during worst period of anxiety | 0.2525 | 0.0149 | -0.039 | 0.6767 | 2.0887 | 0.964445 |
| 20506 | Recent feelings or nervousness or anxiety | 0.2821 | 1.01E-10 | -0.0095 | 0.8374 | 4.5798 | 0.964493 |
| 864 | Number of days/week walked 10+ minutes | 0.0435 | 0.1176 | -0.2487 | 2.48E-19 | 7.4456 | 0.964783 |
| 1508_3 | Coffee type: Ground coffee (include espresso, filter etc) | 0.1255 | 3.89E-06 | -0.1668 | 2.01E-10 | 7.7397 | 0.964831 |
| 20547_1 | Activities undertaken to treat depression: Talking therapies, such as psychotherapy, counselling, group therapy or CBT | 0.3275 | 2.73E-14 | 0.0347 | 0.42 | 4.8093 | 0.96507 |
| 22601_23113399 | Job coding: higher education teaching professional, university lecturer/professor (including college/university head/vice chancellor) | 0.1273 | 0.0369 | -0.168 | 0.0187 | 3.142 | 0.966248 |
| 104450 | Apple intake | 0.0069 | 0.925 | -0.2935 | 0.0034 | 2.4228 | 0.968548 |
| 20024_4113 | Job code - deduced: Local government clerical officers and assistants | 0.0203 | 0.8423 | -0.2809 | 0.0435 | 1.746 | 0.968897 |
| 100024 | Calcium | 0.1887 | 0.0024 | -0.1151 | 0.0615 | 3.4703 | 0.970009 |
| 1488 | Tea intake | 0.1987 | 6.18E-13 | -0.1056 | 2.58E-05 | 8.1568 | 0.970219 |
| 20501 | Ever had period of mania / excitability | 0.3929 | 1.88E-07 | 0.087 | 0.1526 | 3.1582 | 0.970883 |
| 6179_3 | Mineral and other dietary supplements: Calcium | 0.13 | 0.0021 | -0.1765 | 7.47E-05 | 4.9918 | 0.971128 |
| 20002_1482 | Non-cancer illness code, self-reported: chronic fatigue syndrome | 0.2061 | 0.0035 | -0.101 | 0.144 | 3.1064 | 0.971372 |
| 100004 | Fat | 0.1963 | 0.0229 | -0.111 | 0.1622 | 2.6205 | 0.971453 |
| 2764 | Age at last live birth | 0.0839 | 0.0103 | -0.2248 | 1.62E-12 | 6.7678 | 0.972015 |
| 22660_103 | Gap coding: Full-time or part-time education | 0.2703 | 0.0002 | -0.039 | 0.5513 | 3.1823 | 0.972252 |
| 100460 | Added milk to standard tea | 0.0561 | 0.3462 | -0.2546 | 0.0001 | 3.5023 | 0.9728 |
| 20537 | Frequency of difficulty controlling worry during worst period of anxiety | 0.1797 | 0.0415 | -0.1313 | 0.1093 | 2.5824 | 0.972917 |
| S02 | Diagnoses - main ICD10: S02 Fracture of skull and facial bones | 0.1032 | 0.2922 | -0.2177 | 0.0405 | 2.2195 | 0.976528 |
| 20546_1 | Substances taken for depression: Unprescribed medication (more than once) | 0.2314 | 0.0078 | -0.091 | 0.3284 | 2.5315 | 0.977038 |
| 20548_9 | Manifestations of mania or irritability: I was more active than usual | 0.3743 | 8.11E-08 | 0.0519 | 0.4068 | 3.4386 | 0.977038 |
| 2926 | Number of unsuccessful stop-smoking attempts | 0.3674 | 4.65E-13 | 0.0439 | 0.3603 | 4.6287 | 0.977406 |
| 6142_3 | Current employment status: Looking after home and/or family | 0.0759 | 0.2758 | -0.2503 | 0.0015 | 3.1051 | 0.978288 |
| 670_2 | Type of accommodation lived in: A flat, maisonette or apartment | 0.35 | 2.44E-12 | 0.0237 | 0.6115 | 4.774 | 0.97832 |
| 20425 | Ever worried more than most people would in similar situation | 0.2981 | 3.95E-10 | -0.0309 | 0.4893 | 5.0384 | 0.979172 |
| 100014 | Folate | 0.1288 | 0.0225 | -0.2012 | 0.0017 | 3.8587 | 0.979481 |
| 6141_8 | How are people in household related to participant: Other unrelated | 0.3209 | 0.022 | -0.0109 | 0.9128 | 1.9328 | 0.980026 |
| 6158_3 | Why reduced smoking: Health precaution | 0.1356 | 0.147 | -0.2033 | 0.0417 | 2.4781 | 0.982058 |
| 20003_1140871024 | Treatment/medication code: vitamin b compound tablet | 0.2925 | 0.0207 | -0.0466 | 0.6204 | 2.15 | 0.982113 |
| 20002_1286 | Non-cancer illness code, self-reported: depression | 0.3192 | 3.62E-14 | -0.0202 | 0.6104 | 5.8648 | 0.982194 |
| 1160 | Sleep duration | 0.1609 | 1.76E-09 | -0.1802 | 1.01E-11 | 9.0674 | 0.982651 |
| F5_ALCOHOLAC | Acute alcohol intoxication | 0.3025 | 0.0337 | -0.0402 | 0.7433 | 1.8225 | 0.983071 |
| 6162_4 | Types of transport used (excluding work): Cycle | 0.0989 | 0.0019 | -0.2453 | 3.18E-13 | 7.4175 | 0.983458 |
| T43 | Diagnoses - main ICD10: T43 Poisoning by psychotropic drugs, not elsewhere classified | 0.4112 | 0.0399 | 0.0663 | 0.5856 | 1.473 | 0.983635 |
| 20496 | Felt distant from other people in past month | 0.4188 | 6.86E-09 | 0.0696 | 0.2988 | 3.5426 | 0.984691 |

| | | | | | | | |
|---|---|---|---|---|---|---|---|
| H7_CONVER STRAB | Convergent concomitant strabismus | 0.121 | 0.3081 | -0.2297 | 0.046 | 2.1211 | 0.985046 |
| 6179_4 | Mineral and other dietary supplements: Zinc | 0.2623 | 1.37E-08 | -0.0892 | 0.0356 | 5.5994 | 0.985232 |
| 104460 | Banana intake | 0.0574 | 0.3894 | -0.2951 | 6.03E-05 | 3.5489 | 0.985462 |
| 1150_2 | Usual side of head for mobile phone use: Right | 0.1122 | 0.0039 | -0.2406 | 3.41E-12 | 6.7766 | 0.98553 |
| 20418 | Impact on normal roles during worst period of anxiety | 0.2717 | 2.77E-06 | -0.0836 | 0.1905 | 4.1172 | 0.986089 |
| 1980 | Worrier / anxious feelings | 0.1892 | 3.16E-09 | -0.1677 | 3.04E-08 | 8.112 | 0.986436 |
| 20002_1287 | Non-cancer illness code, self-reported: anxiety/panic attacks | 0.2201 | 0.0005 | -0.1396 | 0.0259 | 4.0501 | 0.987027 |
| SPONDYLOP ATHY_FG | Spondylopathies (FG) | 0.2846 | 0.013 | -0.0777 | 0.3446 | 2.5689 | 0.987555 |
| 20021 | Speech-reception-threshold (SRT) estimate (right) | 0.1307 | 0.0565 | -0.2406 | 0.0004 | 3.8384 | 0.989243 |
| 20549_4 | Substances taken for anxiety: Drugs or alcohol (more than once) | 0.3789 | 1.01E-05 | 0.0065 | 0.9311 | 3.266 | 0.989435 |
| 4631 | Ever unenthusiastic/disinterested for a whole week | 0.386 | 1.34E-21 | 0.0125 | 0.7649 | 6.425 | 0.989624 |
| 20126_4 | Bipolar and major depression status: Probable Recurrent major depression (moderate) | 0.3346 | 1.62E-08 | -0.0394 | 0.4931 | 4.5318 | 0.989709 |
| 100670 | White wine intake | 0.0523 | 0.5929 | -0.3226 | 0.0068 | 2.4305 | 0.98986 |
| M45 | Diagnoses - main ICD10: M45 Ankylosing spondylitis | 0.362 | 0.0331 | -0.0153 | 0.8735 | 1.9339 | 0.990254 |
| 20468 | Ever believed in an un-real conspiracy against self | 0.6352 | 0.0441 | 0.2569 | 0.1436 | 1.0477 | 0.990414 |
| 5663 | Length of longest manic/irritable episode | 0.2719 | 0.0096 | -0.1066 | 0.2602 | 2.6769 | 0.990446 |
| 4559 | Family relationship satisfaction | 0.3934 | 6.54E-26 | 0.0132 | 0.741 | 6.9429 | 0.990712 |
| 102130 | Dessert consumers | 0.0541 | 0.5735 | -0.3284 | 0.0308 | 2.1266 | 0.991062 |
| 20002_1067 | Non-cancer illness code, self-reported: peripheral vascular disease | 0.2785 | 0.0152 | -0.1049 | 0.2179 | 2.6833 | 0.991196 |
| 6144_3 | Never eat eggs, dairy, wheat, sugar: Wheat products | 0.3736 | 0.0008 | -0.0112 | 0.893 | 2.7719 | 0.9914 |
| 904 | Number of days/week of vigorous physical activity 10+ minutes | 0.1233 | 1.64E-05 | -0.2692 | 6.34E-20 | 9.5527 | 0.992453 |
| 20406 | Ever addicted to alcohol | 0.2168 | 0.0211 | -0.1797 | 0.0354 | 3.122 | 0.992953 |
| 6155_3 | Vitamin and mineral supplements: Vitamin C | 0.2548 | 1.74E-09 | -0.1424 | 0.0007 | 6.6634 | 0.993038 |
| 4598 | Ever depressed for a whole week | 0.3848 | 2.14E-27 | -0.0197 | 0.5997 | 7.8223 | 0.993867 |
| 6179_6 | Mineral and other dietary supplements: Selenium | 0.2534 | 0.000039 | -0.1547 | 0.0076 | 4.8234 | 0.994243 |
| 100390 | Tea consumed | 0.1249 | 0.0306 | -0.2833 | 2.64E-06 | 4.887 | 0.994253 |
| 6155_5 | Vitamin and mineral supplements: Vitamin E | 0.2279 | 0.0005 | -0.1811 | 0.0017 | 4.682 | 0.994333 |
| 20548_6 | Manifestations of mania or irritability: I was more creative or had more ideas than usual | 0.5073 | 2.56E-06 | 0.0976 | 0.2262 | 3.0407 | 0.994403 |
| 100016 | Potassium | 0.1822 | 0.0006 | -0.2281 | 0.0001 | 5.1438 | 0.994462 |
| 3669 | Lifetime number of same-sex sexual partners | 0.1594 | 0.0732 | -0.2541 | 0.0116 | 3.0768 | 0.994768 |
| 20019 | Speech-reception-threshold (SRT) estimate (left) | 0.2247 | 8.04E-05 | -0.1892 | 5.47E-05 | 5.6073 | 0.994805 |
| 1418_4 | Milk type used: Soya | 0.3046 | 7.31E-09 | -0.1134 | 0.0191 | 5.8418 | 0.995173 |
| 100003 | Protein | 0.2494 | 0.0048 | -0.1688 | 0.0439 | 3.4312 | 0.99519 |
| 100011 | Iron | 0.1916 | 0.0011 | -0.2272 | 0.0007 | 4.6897 | 0.995242 |
| 100005 | Carbohydrate | 0.2185 | 0.0003 | -0.2004 | 0.0011 | 4.8717 | 0.99525 |
| 1970 | Nervous feelings | 0.2645 | 4.22E-18 | -0.1552 | 3.05E-07 | 9.7622 | 0.995318 |
| 1339 | Non-oily fish intake | 0.1825 | 2.03E-06 | -0.2379 | 7.42E-15 | 8.5619 | 0.995377 |
| 20544_15 | Mental health problems ever diagnosed by a professional: Anxiety, nerves or generalized anxiety disorder | 0.2967 | 5.44E-10 | -0.126 | 0.0086 | 6.2399 | 0.995566 |
| 100002 | Energy | 0.2422 | 0.0003 | -0.1829 | 0.0047 | 4.5465 | 0.995755 |
| 20550_1 | Activities undertaken to treat anxiety: Talking therapies, such as psychotherapy, counselling, group therapy or CBT | 0.3743 | 2.69E-13 | -0.0543 | 0.2942 | 5.8847 | 0.996018 |
| F31 | Diagnoses - main ICD10: F31 Bipolar affective disorder | 0.2971 | 8.5E-07 | -0.1332 | 0.0265 | 5.0585 | 0.996141 |

| | | | | | | | |
|---|---|---|---|---|---|---|---|
| 20495 | Avoided activities or situations because of previous stressful experience in past month | 0.5194 | 2.7E-25 | 0.0855 | 0.0863 | 6.1486 | 0.996389 |
| 20003_1140867504 | Treatment/medication code: priadel 200mg m/r tablet | 0.3185 | 0.0094 | -0.1222 | 0.2389 | 2.7421 | 0.996819 |
| 100025 | Vitamin E | 0.24 | 0.0007 | -0.2009 | 0.0039 | 4.4251 | 0.996831 |
| 6155_4 | Vitamin and mineral supplements: Vitamin D | 0.2614 | 0.0001 | -0.1827 | 0.0039 | 4.7803 | 0.997016 |
| 3849 | Number of pregnancy terminations | 0.2936 | 1.94E-08 | -0.1522 | 0.0058 | 5.8626 | 0.997111 |
| 20549_3 | Substances taken for anxiety: Medication prescribed to you (for at least two weeks) | 0.3603 | 4.57E-12 | -0.0879 | 0.0786 | 6.2068 | 0.99724 |
| 20497 | Repeated disturbing thoughts of stressful experience in past month | 0.4669 | 2.13E-33 | 0.0171 | 0.6897 | 7.7762 | 0.997322 |
| 20003_1140867888 | Treatment/medication code: paroxetine | 0.2855 | 0.0208 | -0.1651 | 0.1065 | 2.8098 | 0.997363 |
| 20003_1140863152 | Treatment/medication code: diazepam | 0.397 | 0.0014 | -0.0553 | 0.4917 | 3.0588 | 0.997448 |
| 884 | Number of days/week of moderate physical activity 10+ minutes | 0.1905 | 9.03E-11 | -0.2626 | 9.71E-17 | 10.4978 | 0.997486 |
| 20550_3 | Activities undertaken to treat anxiety: Other therapeutic activities such as mindfulness, yoga or art classes | 0.3395 | 3.69E-08 | -0.1214 | 0.0327 | 5.4958 | 0.997839 |
| 4241 | Signal-to-noise-ratio (SNR) of triplet (right) | 0.2174 | 0.0003 | -0.2456 | 6.87E-06 | 5.671 | 0.997925 |
| 4294_0 | Final attempt correct: no | 0.2697 | 3.27E-05 | -0.1992 | 0.001 | 5.2767 | 0.998153 |
| 20023 | Mean time to correctly identify matches | 0.2258 | 5.68E-26 | -0.2436 | 6.48E-24 | 14.5303 | 0.998171 |
| 1448_2 | Bread type: Brown | 0.1854 | 2.24E-05 | -0.2856 | 1.28E-07 | 6.7726 | 0.998228 |
| 20428 | Professional informed about anxiety | 0.2965 | 4.67E-05 | -0.1766 | 0.0204 | 4.4892 | 0.9983 |
| 100009 | Englyst dietary fibre | 0.2294 | 1.26E-05 | -0.2442 | 3.9E-05 | 5.9741 | 0.998317 |
| 4233 | Mean signal-to-noise ratio (SNR), (left) | 0.2585 | 7.93E-05 | -0.2181 | 0.0003 | 5.3492 | 0.998415 |
| 1329 | Oily fish intake | 0.2217 | 3.69E-18 | -0.2642 | 2.08E-25 | 13.5003 | 0.998687 |
| 6155_2 | Vitamin and mineral supplements: Vitamin B | 0.361 | 4.5E-10 | -0.1249 | 0.0123 | 6.357 | 0.998687 |
| 20003_1140883476 | Treatment/medication code: procyclidine | 0.4486 | 0.0158 | -0.0381 | 0.6953 | 2.3201 | 0.998708 |
| 1468_3 | Cereal type: Oat cereal (e.g. Ready Brek, porridge) | 0.3209 | 2.53E-16 | -0.1724 | 1.41E-05 | 8.8418 | 0.998872 |
| 404 | Duration to first press of snap-button in each round | 0.2423 | 6.78E-31 | -0.2553 | 3.21E-25 | 15.3844 | 0.998968 |
| 20421 | Ever felt worried, tense, or anxious for most of a month or longer | 0.3889 | 6.11E-19 | -0.1115 | 0.0052 | 8.4563 | 0.999026 |
| 1990 | Tense / 'highly strung' | 0.3479 | 2.11E-30 | -0.154 | 3.4E-07 | 11.7127 | 0.999056 |
| 41248_5003 | Destinations on discharge from hospital (recoded): Transfer to other NHS provider: Psychiatry, learning disabilities | 0.5126 | 0.0305 | 0.0022 | 0.9889 | 1.7858 | 0.999211 |
| 20126_1 | Bipolar and major depression status: Bipolar I Disorder | 0.4765 | 0.0012 | -0.034 | 0.7131 | 2.9298 | 0.999213 |
| 20442 | Lifetime number of depressed periods | 0.4575 | 2.75E-05 | -0.0581 | 0.5172 | 3.6505 | 0.999294 |
| 4230 | Signal-to-noise-ratio (SNR) of triplet (left) | 0.263 | 5.68E-06 | -0.2573 | 2.09E-07 | 6.8245 | 0.999362 |
| 20126_3 | Bipolar and major depression status: Probable Recurrent major depression (severe) | 0.5219 | 1.67E-07 | -0.0004 | 0.9959 | 4.3279 | 0.999389 |
| 102700 | Starchy food consumers | 0.2321 | 0.0025 | -0.2903 | 0.0017 | 4.353 | 0.99939 |
| 20547_3 | Activities undertaken to treat depression: Other therapeutic activities such as mindfulness, yoga or art classes | 0.3957 | 7.93E-13 | -0.1345 | 0.0112 | 6.9285 | 0.999486 |
| 2877_2 | Type of tobacco previously smoked: Hand-rolled cigarettes | 0.4411 | 0.0012 | -0.0893 | 0.3742 | 3.1335 | 0.999488 |
| 6155_1 | Vitamin and mineral supplements: Vitamin A | 0.3307 | 5.41E-06 | -0.202 | 0.0184 | 4.7401 | 0.999513 |
| 2100 | Seen a psychiatrist for nerves, anxiety, tension or depression | 0.4821 | 1.82E-53 | -0.0537 | 0.1102 | 11.6681 | 0.999546 |
| 4291 | Number of attempts | 0.2612 | 2.18E-12 | -0.2759 | 3.42E-13 | 10.1137 | 0.999558 |
| KRA_PSY_ANYMENTAL | Any mental disorder | 0.4836 | 1.22E-13 | -0.0568 | 0.3683 | 5.9559 | 0.99959 |
| V_MENTAL_BEHAV | Mental and behavioural disorders | 0.5054 | 2.29E-14 | -0.0492 | 0.4515 | 5.9643 | 0.999702 |
| 5386 | Number of unenthusiastic/disinterested episodes | 0.3818 | 2.7E-05 | -0.1768 | 0.0281 | 4.598 | 0.999728 |

| | | | | | | | |
|---|---|---|---|---|---|---|---|
| T42 | Diagnoses - main ICD10: T42 Poisoning by antiepileptic, sedative-hypnotic and anti-Parkinsonism drugs | 0.5076 | 0.039 | -0.0609 | 0.6195 | 2.0683 | 0.999784 |
| 22617_2312 | Job SOC coding: Further education teaching professionals | 0.4665 | 0.0033 | -0.1035 | 0.3643 | 2.9174 | 0.999791 |
| XV_PREGNANCY_BIRTH | Pregnancy, childbirth and the puerperium | 0.2135 | 0.0562 | -0.3577 | 0.0013 | 3.6224 | 0.999797 |
| 100023 | Starch | 0.3279 | 0.0034 | -0.2544 | 0.0092 | 3.9197 | 0.999844 |
| 100017 | Magnesium | 0.2904 | 2.61E-08 | -0.2944 | 1.23E-06 | 7.3047 | 0.999853 |
| F5_DEPRESSIO | Depression | 0.4496 | 2.22E-07 | -0.1382 | 0.0369 | 5.3846 | 0.999863 |
| 20118_7 | Home area population density - urban or rural: England/Wales - Village - less sparse | 0.0382 | 0.7091 | -0.5551 | 0.0006 | 3.1021 | 0.99988 |
| 104340 | Fresh tomato intake | 0.3041 | 0.0101 | -0.2949 | 0.0079 | 3.6958 | 0.999896 |
| 103140 | Fish consumer | 0.2875 | 0.0073 | -0.3147 | 0.0171 | 3.543 | 0.999904 |
| 20466 | Ever prescribed a medication for unusual or psychotic experiences | 0.5658 | 0.0185 | -0.0615 | 0.624 | 2.3158 | 0.999949 |
| 2664_3 | Reason for reducing amount of alcohol drunk: Health precaution | 0.3424 | 2.46E-20 | -0.2895 | 1.57E-13 | 11.7078 | 0.999954 |
| 6157_3 | Why stopped smoking: Health precaution | 0.2628 | 1.24E-11 | -0.3709 | 1.43E-16 | 10.6788 | 0.999956 |
| 20111_12 | Illnesses of siblings: Severe depression | 0.5346 | 7.84E-31 | -0.1027 | 0.0234 | 9.8387 | 0.99996 |
| 20002_1291 | Non-cancer illness code, self-reported: mania/bipolar disorder/manic depression | 0.5121 | 2.59E-09 | -0.1269 | 0.0861 | 5.6354 | 0.999962 |
| F32 | Diagnoses - main ICD10: F32 Depressive episode | 0.483 | 0.0002 | -0.1579 | 0.0657 | 4.1368 | 0.999964 |
| 20474 | Ever believed in un-real communications or signs | 0.809 | 0.0156 | 0.155 | 0.1671 | 1.8527 | 0.999974 |
| 20548_8 | Manifestations of mania or irritability: I was more confident than usual | 0.6805 | 0.0272 | 0.0253 | 0.8479 | 1.9552 | 0.999975 |
| FIBRO_COMORB | Fibromyalgia related co-morbidities | 0.4862 | 0.0001 | -0.1884 | 0.0235 | 4.4777 | 0.999985 |
| 4620 | Number of depression episodes | 0.4872 | 4.53E-06 | -0.2004 | 0.0218 | 4.9993 | 0.99999 |
| F33 | Diagnoses - main ICD10: F33 Recurrent depressive disorder | 0.5184 | 0.0066 | -0.1725 | 0.1058 | 3.1617 | 0.999991 |
| F5_MOOD | Mood [affective] disorders | 0.5095 | 5.22E-11 | -0.1838 | 0.0041 | 6.8925 | 0.999991 |
| KRA_PSY_MOOD | Mood disorders | 0.5095 | 5.22E-11 | -0.1838 | 0.0041 | 6.8925 | 0.999991 |
| 4290 | Duration screen displayed | 0.3179 | 5.34E-26 | -0.3772 | 2.44E-33 | 16.007 | 0.999992 |
| 22601_23123400 | Job coding: further education teaching professional, college lecturer/professor | 0.5542 | 0.033 | -0.1456 | 0.3115 | 2.3549 | 0.999993 |
| 20110_12 | Illnesses of mother: Severe depression | 0.6117 | 7.83E-23 | -0.1088 | 0.0574 | 8.5195 | 0.999996 |
| 20090_353 | Type of fat/oil used in cooking: Olive oil | 0.3764 | 0.0006 | -0.3557 | 3.39E-05 | 5.2657 | 0.999997 |
| 20107_12 | Illnesses of father: Severe depression | 0.5466 | 4.94E-13 | -0.186 | 0.003 | 7.4638 | 0.999997 |
| 4288 | Time to answer | 0.3181 | 1.52E-14 | -0.4554 | 1.61E-24 | 12.7109 | 0.999999 |
| 5674 | Severity of manic/irritable episodes | 0.667 | 0.0403 | -0.1156 | 0.3756 | 2.2328 | 0.999999 |
| 399 | Number of incorrect matches in round | 0.3243 | 1.72E-36 | -0.5381 | 3.33E-93 | 23.4526 | 1 |
| 20477 | Ever talked to a health professional about unusual or psychotic experiences | 0.7207 | 0.04 | -0.1568 | 0.211 | 2.3549 | 1 |
| 400 | Time to complete round | 0.3449 | 7.53E-59 | -0.6092 | 1.1E-177 | 31.5995 | 1 |
| F20 | Diagnoses - main ICD10: F20 Schizophrenia | 0.8824 | 0.0007 | -0.1066 | 0.3494 | 3.4685 | 1 |
| 4924 | Attempted fluid intelligence (FI) test. | 0.4138 | 0.0016 | -0.5836 | 0.0017 | 4.3788 | 1 |
| F5_SCHIZO | Schizophrenia, schizotypal and delusional disorders | 1.1918 | 0.0099 | -0.1203 | 0.3289 | 2.7436 | 1 |
| KRA_PSY_SCHIZODEL | Schizophrenia or delusion | 1.1918 | 0.0099 | -0.1203 | 0.3289 | 2.7436 | 1 |

**Table S14.** Genomic risk loci and independent variants (LD $r^2$<0.1) identified in attention deficit hyperactivity disorder GWAS.

| rsID | Chromosome | Position | Alleles | p | Consequence | Gene Symbol | Region Biotype |
|---|---|---|---|---|---|---|---|
| rs11420276 | 1 | 44184192 | G/GT | 6.45E-13 | ncRNA_intronic | - | - |
| rs3952787 | 1 | 44323244 | C/T | 7.06E-09 | intronic | *ARTN* | protein_coding |
| rs1222063 | 1 | 96602440 | A/G | 3.07E-08 | intergenic | - | - |
| rs4858241 | 3 | 20669071 | G/T | 8.17E-09 | intergenic | - | - |
| rs28411770 | 4 | 31151456 | C/T | 1.15E-08 | intergenic | *PCDH7* | protein_coding |
| rs4916723 | 5 | 87854395 | A/C | 1.81E-08 | ncRNA_intronic | - | - |
| rs304132 | 5 | 88215594 | A/G | 3.05E-08 | ncRNA_intronic | - | - |
| rs5886709 | 7 | 114086133 | G/GTC | 2.06E-08 | intronic | *FOXP2* | - |
| rs74760947 | 8 | 34352610 | A/G | 1.39E-08 | intergenic | - | - |
| rs11591402 | 10 | 106747354 | A/T | 1.76E-08 | intronic | *SORCS3* | protein_coding |
| rs1427829 | 12 | 89760744 | A/G | 1.35E-09 | upstream | *DUSP6* | protein_coding |
| rs8039398 | 15 | 47730870 | C/T | 2.99E-09 | intronic | *SEMA6D* | protein_coding |
| rs212178 | 16 | 72578131 | A/G | 1.20E-08 | ncRNA_intronic | - | - |

**Table S15.** Genomic risk loci and independent variants (LD $r^2$<0.1) identified in *PhoneUse* GWAS.

| rsID | Chromosome | Position | Alleles | p | Consequence | Gene Symbol | Region Biotype |
|---|---|---|---|---|---|---|---|
| rs13425125 | 2 | 31968142 | A/T | 7.05E-09 | intergenic | *MEMO1* | protein_coding |
| rs13062093 | 3 | 35667057 | G/T | 4.42E-12 | intergenic | - | - |
| rs78648104 | 6 | 50683009 | C/T | 2.64E-08 | exonic | *TFAP2D* | protein_coding |
| rs76899638 | 6 | 55147508 | A/ATG | 1.54E-08 | downstream | *HCRTR2* | protein_coding |
| rs10228494 | 7 | 114083550 | C/G | 2.10E-09 | intronic | *FOXP2* | protein_coding |
| rs4638225 | 10 | 118676964 | A/G | 2.04E-11 | intronic | - | - |

**Table S16.** Genomic risk loci and independent variants (LD $r^2$<0.1) identified in autism spectrum disorder GWAS.

| rsID | Chromosome | Position | Alleles | p | Consequence | Gene Symbol | Region Biotype |
|---|---|---|---|---|---|---|---|
| rs10099100 | 8 | 10576775 | C/G | 1.07E-08 | intergenic | *RP1L1* | protein_coding |
| rs71190156 | 20 | 14836243 | G/GTTTTTTT | 2.75E-08 | intronic | *MACROD2* | protein_coding |
| rs11475262 | 20 | 21145353 | C/CT | 2.28E-08 | ncRNA_intronic | - | - |
| rs12625304 | 20 | 21182388 | C/T | 8.16E-09 | ncRNA_intronic | - | - |
| rs910805 | 20 | 21248116 | A/G | 2.04E-09 | intergenic | *NKX2-4* | protein_coding |

**Table S17.** Genomic risk loci and independent variants (LD $r^2$<0.1) identified in attention deficit hyperactivity disorder GWAS with female subjects.

| rsID | Chromosome | Position | Alleles | p | Consequence | Gene Symbol | Region Biotype |
|---|---|---|---|---|---|---|---|
| rs222903 | 1 | 96485694 | A/C | 4.87E-06 | intronic | *RP11-147C23.1* | protein_coding |
| rs56757467 | 2 | 20732527 | G/GA | 3.93E-06 | intergenic | - | - |
| rs150844750 | 2 | 109359283 | C/T | 1.46E-06 | intronic | *SULT1C4* | protein_coding |
| rs12052540 | 2 | 118666061 | C/T | 3.14E-06 | ncRNA_intronic | *CCDC93* | protein_coding |
| rs79891548 | 2 | 166337818 | C/T | 1.10E-06 | intronic | *SCN2A* | protein_coding |
| rs58520674 | 2 | 172443949 | C/G | 4.10E-06 | downstream | - | - |
| rs76384310 | 3 | 53768055 | A/G | 4.75E-06 | intronic | *CACNA1D* | protein_coding |
| rs17815990 | 4 | 89798892 | C/T | 3.94E-06 | intronic | *TIGD2* | protein_coding |
| rs17588356 | 4 | 151210677 | G/T | 2.33E-06 | intronic | *DCLK2* | protein_coding |
| rs114247285 | 5 | 75339982 | A/G | 7.75E-07 | intergenic | *SV2C* | protein_coding |
| rs142295881 | 6 | 53899088 | A/G | 1.03E-06 | ncRNA_intronic | *MLIP* | protein_coding |
| rs12535840 | 7 | 81669553 | C/G | 3.03E-06 | intronic | *CACNA2D1* | protein_coding |
| rs35678938 | 7 | 116459931 | A/C | 3.45E-06 | intronic | *MET* | protein_coding |
| rs28600876 | 11 | 69705480 | C/G | 6.55E-07 | intergenic | - | - |
| rs71450738 | 12 | 24198831 | T/TCCATAG | 1.65E-06 | intergenic | - | - |
| rs770082 | 12 | 89776485 | A/G | 1.79E-06 | intergenic | *DUSP6* | protein_coding |
| rs11849175 | 14 | 33382322 | C/G | 4.33E-06 | intergenic | *NPAS3* | protein_coding |
| rs5022358 | 15 | 24414283 | A/G | 4.14E-06 | ncRNA_intronic | - | - |
| rs7166121 | 15 | 29885746 | A/G | 4.29E-06 | intergenic | *FAM189A1* | protein_coding |
| rs12594695 | 15 | 40220100 | C/G | 3.86E-06 | intergenic | *GPR176* | protein_coding |
| rs7181782 | 15 | 80686993 | C/G | 9.34E-08 | intronic | *ARNT2* | protein_coding |
| rs4984687 | 16 | 754314 | C/T | 1.89E-07 | intronic | *WDR90* | protein_coding |
| rs9941217 | 16 | 18050926 | C/G | 9.35E-07 | ncRNA_intronic | - | - |
| rs12949493 | 17 | 62020678 | A/T | 2.81E-06 | intronic | *DCAF7* | protein_coding |

**Table S18.** Genomic risk loci and independent variants (LD $r^2$<0.1) identified in attention deficit hyperactivity disorder GWAS with male subjects.

| rsID | Chromosome | Position | Alleles | p | Consequence | Gene Symbol | Region Biotype |
|---|---|---|---|---|---|---|---|
| rs3047819 | 5 | 88175199 | T/TTA | 1.47E-08 | intronic | *MEF2C* | protein_coding |
| rs200508662 | 5 | 120391182 | C/T | 3.09E-08 | intergenic | - | - |
| rs8039398 | 15 | 47730870 | C/T | 2.55E-08 | intronic | *SEMA6D* | protein_coding |

**Table S19.** Genomic risk loci and independent variants (LD $r^2$<0.1) identified in *PhoneUse* GWAS with female subjects.

| rsID | Chromosome | Position | Alleles | p | Consequence | Gene Symbol | Region Biotype |
|---|---|---|---|---|---|---|---|
| rs13094224 | 3 | 35784581 | A/G | 9.35E-11 | intronic | - | - |
| rs6974757 | 7 | 114124660 | C/G | 6.54E-09 | intronic | *FOXP2* | protein_coding |
| rs6585422 | 10 | 118695141 | C/T | 2.22E-08 | intronic | *ENO4* | protein_coding |

**Table S20.** Genomic risk loci and independent variants (LD $r^2$<0.1) identified in *PhoneUse* GWAS with male subjects.

| rsID | Chromosome | Position | Alleles | p | Consequence | Gene Symbol | Region Biotype |
|---|---|---|---|---|---|---|---|
| rs5774341 | 1 | 57839849 | G/GA | 5.47E-06 | intronic | *DAB1* | protein_coding |
| rs3860292 | 1 | 207698607 | C/T | 1.28E-06 | intronic | *CR1* | protein_coding |
| rs148481064 | 1 | 241215760 | A/G | 7.05E-06 | intronic | *RGS7* | protein_coding |
| rs139071744 | 1 | 174708968 | A/G | 8.45E-06 | intronic | *RABGAP1L* | protein_coding |
| rs288324 | 2 | 183701588 | A/G | 5.03E-06 | intronic | *FRZB* | protein_coding |
| rs212699 | 2 | 32443519 | A/G | 5.64E-07 | intronic | *YIPF4* | protein_coding |
| rs112662433 | 2 | 151539798 | C/T | 2.17E-06 | intergenic | - | - |
| rs75542933 | 3 | 120534887 | A/G | 5.11E-06 | intergenic | - | - |
| rs746696365 | 3 | 103020660 | C/CT | 9.26E-06 | intergenic | - | - |
| rs73002363 | 3 | 1295742 | A/G | 1.36E-06 | intronic | *CNTN6* | protein_coding |
| rs6783978 | 3 | 24888172 | C/T | 9.78E-06 | ncRNA_intronic | - | - |
| rs161894 | 3 | 7853659 | G/T | 3.98E-06 | intergenic | - | - |
| rs13080060 | 3 | 32398234 | C/T | 4.14E-06 | intronic | *CMTM8* | protein_coding |
| rs12054250 | 3 | 20424345 | A/G | 7.14E-06 | intergenic | - | - |
| rs77607787 | 4 | 34925914 | G/T | 5.73E-06 | intergenic | - | - |
| rs7376072 | 4 | 132277631 | C/T | 4.00E-06 | intergenic | - | - |
| rs56672563 | 4 | 188803346 | T/TG | 1.91E-06 | intergenic | - | - |
| rs13150201 | 4 | 130987179 | A/C | 1.78E-06 | intergenic | - | - |
| rs10001603 | 4 | 152647577 | A/T | 5.63E-06 | intronic | *FAM160A1* | protein_coding |
| rs79955751 | 5 | 107274714 | C/T | 3.87E-07 | intronic | *FBXL17* | protein_coding |
| rs7717164 | 5 | 75942970 | A/G | 8.25E-06 | intronic | *IQGAP2* | protein_coding |
| rs56047718 | 5 | 176138484 | A/T | 3.24E-06 | ncRNA_intronic | - | - |
| rs3066289 | 5 | 62960761 | C/T | 6.74E-06 | intergenic | - | - |
| rs10077638 | 5 | 124237394 | G/T | 6.67E-06 | intergenic | - | - |
| rs10044452 | 5 | 66302605 | C/T | 8.55E-06 | intronic | *MAST4* | protein_coding |
| rs9349536 | 6 | 50322852 | G/T | 2.35E-06 | intergenic | - | - |
| rs7341218 | 6 | 29012116 | A/G | 7.01E-06 | exonic | *ZNF311* | protein_coding |
| rs370705844 | 6 | 55144318 | A/AAC | 6.20E-07 | intronic | *HCRTR2* | protein_coding |
| rs141547796 | 6 | 50615935 | A/G | 5.16E-07 | intergenic | *TFAP2D* | protein_coding |
| rs138799337 | 6 | 55517145 | A/AATCTGGTAGAC | 6.43E-07 | intergenic | - | - |
| rs13212282 | 6 | 6134032 | A/C | 7.64E-06 | intergenic | *F13A1* | protein_coding |
| rs57635750 | 7 | 153484439 | A/G | 4.62E-06 | intergenic | - | - |
| rs55751112 | 7 | 113284977 | G/T | 4.18E-06 | intergenic | *TSRM* | protein_coding |
| rs4313070 | 7 | 39942647 | C/T | 6.43E-06 | intergenic | - | - |
| rs373152299 | 7 | 69803819 | C/CTTAT | 2.53E-06 | intronic | *AUTS2* | protein_coding |
| rs35309425 | 7 | 68888708 | C/CA | 5.18E-07 | ncRNA_intronic | - | - |
| rs7460351 | 8 | 131052733 | C/T | 5.21E-06 | intergenic | *ASAP1* | protein_coding |
| rs3739241 | 8 | 142221361 | C/G | 4.51E-07 | UTR3 | - | - |
| rs189269915 | 8 | 18427711 | A/C | 3.72E-06 | intronic | *PSD3* | protein_coding |
| rs141917978 | 8 | 42086951 | C/T | 8.87E-06 | intergenic | *SLC20A2* | protein_coding |
| rs10090800 | 8 | 10306432 | C/T | 2.34E-07 | intergenic | *MSRA* | protein_coding |
| rs75740715 | 9 | 15071508 | C/T | 5.71E-06 | ncRNA_intronic | *FREM1* | protein_coding |
| rs11795275 | 9 | 32261513 | C/G | 3.65E-06 | intergenic | - | - |
| rs10120311 | 9 | 38708965 | A/T | 7.64E-06 | intergenic | - | - |
| rs148304049 | 10 | 107809763 | A/AT | 8.30E-06 | intergenic | - | - |
| rs77007523 | 11 | 8633428 | G/T | 7.85E-06 | downstream | *TRIM66* | protein_coding |
| rs1106305 | 11 | 12065087 | C/T | 9.61E-06 | ncRNA_intronic | - | - |
| rs57291280 | 12 | 44146799 | C/T | 2.29E-06 | intronic | *PUS7L* | protein_coding |
| rs55775958 | 14 | 84397578 | G/T | 6.87E-06 | intergenic | - | - |
| rs8063688 | 16 | 80637015 | A/G | 2.42E-06 | UTR3 | *CDYL2* | protein_coding |
| rs190318326 | 16 | 10772298 | A/G | 2.24E-06 | intronic | *TEKT5* | protein_coding |
| rs6146325 | 18 | 53029668 | A/G | 1.87E-06 | intronic | - | - |
| rs117402026 | 20 | 16572060 | C/T | 2.70E-06 | intergenic | *KIF16B* | protein_coding |
| rs62234000 | 22 | 24808961 | C/T | 5.01E-06 | intronic | *GGT5* | protein_coding |

**Table S21.** Gene set enrichments meeting nominal significant in attention deficit hyperactivity disorder (ADHD) and *PhoneUse*. Traits are listed in ascending order by *PhoneUse* significance.

| Enrichment | N genes | ADHD | | *PhoneUse* | | Difference | |
|---|---|---|---|---|---|---|---|
| | | beta | p | beta | p | z | p |
| go_neuron_projection | 912 | 0.0503 | 0.049399 | 0.117 | 6.19E-05 | -1.5489 | 0.0141 |
| go_negative_regulation_of_nitrogen_compound_metabolic_process | 1414 | 0.0555 | 0.011086 | 0.0871 | 0.000191 | -0.9176 | 0.0958 |
| mikkelsen_ips_with_hcp_h3k27me3 | 98 | 0.157 | 0.046864 | 0.342 | 0.000275 | -1.3565 | 0.0244 |
| go_negative_regulation_of_transcription_from_rna_polymerase_ii_promoter | 702 | 0.058 | 0.041554 | 0.117 | 0.000317 | -1.2324 | 0.0391 |
| go_postsynapse | 364 | 0.108 | 0.012069 | 0.164 | 0.000419 | -0.8156 | 0.1222 |
| go_single_organism_behavior | 374 | 0.125 | 0.004061 | 0.158 | 0.000465 | -0.4923 | 0.2422 |
| go_negative_regulation_of_gene_expression | 1400 | 0.0694 | 0.002145 | 0.0796 | 0.000585 | -0.2956 | 0.3373 |
| go_synapse_organization | 138 | 0.129 | 0.040965 | 0.259 | 0.000601 | -1.193 | 0.0399 |
| go_regulation_of_viral_transcription | 58 | 0.252 | 0.014594 | 0.361 | 0.000735 | -0.6731 | 0.1716 |
| go_cognition | 242 | 0.13 | 0.012903 | 0.185 | 0.000803 | -0.6648 | 0.1732 |
| go_regulation_of_potassium_ion_transmembrane_transport | 59 | 0.271 | 0.011031 | 0.384 | 0.000805 | -0.6658 | 0.1691 |
| go_regulation_of_transcription_from_rna_polymerase_ii_promoter | 1694 | 0.0503 | 0.012106 | 0.0682 | 0.001186 | -0.5663 | 0.2111 |
| go_rna_polymerase_ii_transcription_factor_activity_sequence_specific_dna_binding | 601 | 0.113 | 0.001243 | 0.111 | 0.001531 | 0.0378 | 0.5213 |
| go_membrane_depolarization_during_cardiac_muscle_cell_action_potential | 14 | 0.495 | 0.030251 | 0.821 | 0.001585 | -0.8503 | 0.1084 |
| go_somatodendritic_compartment | 629 | 0.0642 | 0.037731 | 0.107 | 0.001759 | -0.8337 | 0.1179 |
| go_cell_morphogenesis_involved_in_neuron_differentiation | 359 | 0.142 | 0.002244 | 0.146 | 0.001919 | -0.0563 | 0.468 |
| go_regulation_of_potassium_ion_transport | 80 | 0.244 | 0.007473 | 0.301 | 0.002137 | -0.3931 | 0.2843 |
| go_negative_regulation_of_cell_projection_organization | 139 | 0.127 | 0.041098 | 0.209 | 0.002446 | -0.7883 | 0.1307 |
| go_axon | 406 | 0.0738 | 0.049682 | 0.127 | 0.002594 | -0.8332 | 0.1175 |
| reactome_nuclear_events_kinase_and_transcription_factor_activation | 23 | 0.67 | 8.44E-05 | 0.504 | 0.002716 | 0.6539 | 0.8245 |
| go_neuron_projection_morphogenesis | 391 | 0.0831 | 0.040245 | 0.129 | 0.003738 | -0.6783 | 0.1669 |
| reactome_erk_mapk_targets | 20 | 0.703 | 0.000111 | 0.504 | 0.004732 | 0.7328 | 0.8525 |
| go_synapse | 729 | 0.0962 | 0.002391 | 0.0876 | 0.005189 | 0.1781 | 0.5996 |
| go_ion_antiporter_activity | 46 | 0.304 | 0.010831 | 0.335 | 0.005287 | -0.1667 | 0.4072 |
| go_cell_morphogenesis_involved_in_differentiation | 499 | 0.114 | 0.003432 | 0.107 | 0.005871 | 0.1169 | 0.566 |
| go_regulation_of_neuron_projection_development | 391 | 0.0791 | 0.039588 | 0.115 | 0.006173 | -0.5567 | 0.213 |
| go_gaba_receptor_binding | 15 | 0.471 | 0.020904 | 0.545 | 0.006413 | -0.2325 | 0.3744 |
| go_cation_cation_antiporter_activity | 24 | 0.612 | 0.00077 | 0.464 | 0.006438 | 0.5507 | 0.7784 |
| go_synapse_assembly | 65 | 0.327 | 0.001437 | 0.29 | 0.006578 | 0.2304 | 0.6317 |
| go_regulation_of_potassium_ion_transmembrane_transporter_activity | 39 | 0.264 | 0.038369 | 0.378 | 0.006675 | -0.5338 | 0.2221 |
| go_behavior | 499 | 0.0977 | 0.009354 | 0.102 | 0.007063 | -0.0731 | 0.4587 |
| go_transcriptional_activator_activity_rna_polymerase_ii_transcription_regulatory_region_sequence_specific_binding | 305 | 0.171 | 0.000465 | 0.129 | 0.007568 | 0.5683 | 0.7926 |
| go_regulation_of_cell_projection_organization | 538 | 0.0885 | 0.010116 | 0.0948 | 0.007973 | -0.1151 | 0.4343 |
| reactome_mapk_targets_nuclear_events_mediated_by_map_kinases | 29 | 0.497 | 0.000936 | 0.392 | 0.008968 | 0.4554 | 0.7442 |
| geiss_response_to_dsrna_up | 35 | 0.229 | 0.045407 | 0.33 | 0.009637 | -0.5174 | 0.2272 |
| go_estrous_cycle | 19 | 0.376 | 0.033167 | 0.507 | 0.010605 | -0.4356 | 0.2614 |
| go_heart_formation | 12 | 0.846 | 0.00089 | 0.619 | 0.011134 | 0.5923 | 0.7989 |
| go_transcription_factor_activity_rna_polymerase_ii_core_promoter_proximal_region_sequence_specific_binding | 321 | 0.147 | 0.0016 | 0.117 | 0.011301 | 0.4196 | 0.7265 |
| myllykangas_amplification_hot_spot_16 | 9 | 1.32 | 0.000123 | 0.622 | 0.011753 | 1.5408 | 0.9737 |
| kim_all_disorders_oligodendrocyte_number_corr_up | 725 | 0.0548 | 0.041081 | 0.0695 | 0.012161 | -0.3337 | 0.3204 |
| go_transcription_from_rna_polymerase_ii_promoter | 691 | 0.0677 | 0.020173 | 0.0757 | 0.012298 | -0.1696 | 0.4042 |
| reactome_regulatory_rna_pathways | 25 | 0.394 | 0.008138 | 0.389 | 0.012634 | 0.0209 | 0.5122 |
| go_neuron_differentiation | 847 | 0.071 | 0.013496 | 0.0735 | 0.013512 | -0.0541 | 0.469 |
| go_camera_type_eye_photoreceptor_cell_differentiation | 15 | 0.496 | 0.015572 | 0.519 | 0.013884 | -0.0698 | 0.4602 |
| go_sequence_specific_dna_binding | 979 | 0.0742 | 0.006496 | 0.0657 | 0.014345 | 0.2007 | 0.6119 |
| go_solute_cation_antiporter_activity | 30 | 0.452 | 0.005282 | 0.354 | 0.015293 | 0.4061 | 0.7101 |
| go_regulation_of_chromosome_organization | 266 | 0.114 | 0.01393 | 0.111 | 0.01541 | 0.0411 | 0.5231 |
| reactome_elongation_arrest_and_recovery | 30 | 0.355 | 0.00855 | 0.333 | 0.015531 | 0.1027 | 0.5587 |
| go_peptidyl_proline_modification | 49 | 0.22 | 0.026889 | 0.242 | 0.015855 | -0.1371 | 0.4235 |
| rampon_enriched_learning_environment_late_up | 22 | 0.31 | 0.043755 | 0.384 | 0.016295 | -0.2891 | 0.3422 |
| go_nucleic_acid_binding_transcription_factor_activity | 1137 | 0.0686 | 0.006834 | 0.0597 | 0.016687 | 0.2256 | 0.6256 |

| | | | | | | | |
|---|---|---|---|---|---|---|---|
| go_transcriptional_repressor_activity_rna_polymerase_ii_core_promoter_proximal_region_sequence_specific_binding | 104 | 0.152 | 0.040033 | 0.187 | 0.018353 | -0.281 | 0.343 |
| chin_breast_cancer_copy_number_up | 26 | 0.404 | 0.010087 | 0.35 | 0.019801 | 0.222 | 0.6219 |
| go_synapse_part | 591 | 0.0987 | 0.004116 | 0.0771 | 0.020322 | 0.4073 | 0.7182 |
| go_dendrite_membrane | 20 | 0.331 | 0.044059 | 0.399 | 0.020466 | -0.2472 | 0.363 |
| go_neuronal_postsynaptic_density | 52 | 0.211 | 0.038208 | 0.259 | 0.020771 | -0.2758 | 0.3433 |
| go_excitatory_synapse | 190 | 0.17 | 0.004247 | 0.138 | 0.021128 | 0.3422 | 0.6904 |
| kegg_bladder_cancer | 41 | 0.236 | 0.037255 | 0.283 | 0.02183 | -0.2434 | 0.3619 |
| go_regulation_of_neuron_differentiation | 534 | 0.0806 | 0.02016 | 0.0805 | 0.022228 | 0.0018 | 0.501 |
| go_negative_regulation_of_cell_morphogenesis_involved_in_differentiation | 113 | 0.153 | 0.036535 | 0.168 | 0.022994 | -0.1251 | 0.4301 |
| go_cd40_receptor_complex | 11 | 0.46 | 0.040111 | 0.551 | 0.023829 | -0.2378 | 0.3647 |
| go_double_stranded_dna_binding | 733 | 0.0755 | 0.011392 | 0.0661 | 0.02533 | 0.1984 | 0.6115 |
| ikeda_mir1_targets_up | 53 | 0.358 | 0.005636 | 0.248 | 0.025527 | 0.5797 | 0.7823 |
| go_membrane_depolarization_during_action_potential | 39 | 0.248 | 0.045467 | 0.304 | 0.026811 | -0.2595 | 0.3516 |
| go_regulation_of_synaptic_transmission_gabaergic | 29 | 0.304 | 0.033192 | 0.351 | 0.028013 | -0.1902 | 0.3885 |
| go_positive_regulation_of_biosynthetic_process | 1730 | 0.048 | 0.015501 | 0.0429 | 0.028254 | 0.1613 | 0.5908 |
| go_camera_type_eye_morphogenesis | 100 | 0.212 | 0.009963 | 0.186 | 0.028261 | 0.1946 | 0.6122 |
| go_regulation_of_multicellular_organismal_development | 1608 | 0.047 | 0.021246 | 0.0453 | 0.0289 | 0.051 | 0.5292 |
| st_erk1_erk2_mapk_pathway | 31 | 0.377 | 0.006881 | 0.323 | 0.02987 | 0.2346 | 0.6379 |
| go_rna_polymerase_ii_activating_transcription_factor_binding | 35 | 0.462 | 0.001059 | 0.294 | 0.029954 | 0.7763 | 0.8686 |
| sabates_colorectal_adenoma_size_dn | 13 | 0.514 | 0.02469 | 0.501 | 0.03095 | 0.0347 | 0.5198 |
| reactome_erks_are_inactivated | 12 | 0.48 | 0.015998 | 0.457 | 0.031274 | 0.0693 | 0.5409 |
| reactome_activation_of_nmda_receptor_upon_glutamate_binding_and_postsynaptic_events | 34 | 0.353 | 0.015878 | 0.311 | 0.032017 | 0.1789 | 0.6011 |
| biocarta_pitx2_pathway | 15 | 0.823 | 0.000599 | 0.428 | 0.03293 | 1.146 | 0.94 |
| go_regulation_of_intracellular_estrogen_receptor_signaling_pathway | 29 | 0.328 | 0.018511 | 0.299 | 0.035361 | 0.1273 | 0.5733 |
| go_protein_heterodimerization_activity | 424 | 0.0957 | 0.016034 | 0.0793 | 0.035369 | 0.2621 | 0.6435 |
| go_neurogenesis | 1358 | 0.0626 | 0.006513 | 0.0469 | 0.036438 | 0.4327 | 0.7334 |
| reactome_microrna_mirna_biogenesis | 22 | 0.419 | 0.007984 | 0.316 | 0.040516 | 0.4102 | 0.7231 |
| iritani_mad1_targets_up | 12 | 0.562 | 0.016421 | 0.497 | 0.041431 | 0.167 | 0.5976 |
| go_regulatory_region_nucleic_acid_binding | 784 | 0.081 | 0.00645 | 0.0574 | 0.042002 | 0.5072 | 0.7654 |
| go_gtp_metabolic_process | 22 | 0.395 | 0.029229 | 0.33 | 0.042445 | 0.2302 | 0.6227 |
| amit_egf_response_40_mcf10a | 19 | 0.371 | 0.047254 | 0.389 | 0.043276 | -0.0567 | 0.4677 |
| pid_reg_gr_pathway | 82 | 0.153 | 0.045043 | 0.167 | 0.044508 | -0.1048 | 0.4385 |
| go_specification_of_organ_identity | 14 | 0.599 | 0.007705 | 0.465 | 0.045074 | 0.3632 | 0.7063 |
| reactome_post_nmda_receptor_activation_events | 31 | 0.36 | 0.017629 | 0.299 | 0.045456 | 0.2479 | 0.6394 |
| reactome_formation_of_rna_pol_ii_elongation_complex_ | 39 | 0.211 | 0.036041 | 0.223 | 0.045457 | -0.068 | 0.4592 |
| go_regulation_of_cardiac_muscle_cell_differentiation | 18 | 0.371 | 0.038656 | 0.385 | 0.04594 | -0.0452 | 0.4734 |
| myllykangas_amplification_hot_spot_14 | 3 | 1.14 | 0.046872 | 0.798 | 0.046316 | 0.4134 | 0.693 |
| go_neural_retina_development | 50 | 0.398 | 0.001436 | 0.233 | 0.047027 | 0.8577 | 0.8926 |
| reactome_developmental_biology | 373 | 0.0926 | 0.025091 | 0.08 | 0.047965 | 0.187 | 0.605 |
| go_synaptic_signaling | 407 | 0.08 | 0.035239 | 0.0754 | 0.049293 | 0.0724 | 0.5414 |

**Table S22.** Gene set enrichments meeting nominal significant in attention deficit hyperactivity disorder (ADHD) and *PhoneUse* in females. Traits are listed in ascending order by *PhoneUse* significance.

| Enrichment | N genes | ADHD Females | | *PhoneUse* Females | | Difference | |
|---|---|---|---|---|---|---|---|
| | | beta | p | beta | p | z | p |
| caffarel_response_to_thc_8hr_3_dn | 9 | 0.492 | 0.0173 | 0.648 | 0.0038 | -0.4634 | 0.1755 |
| go_positive_regulation_of_biomineral_tissue_development | 36 | 0.25 | 0.0433 | 0.387 | 0.007 | -0.6368 | 0.2064 |
| go_regulation_of_skeletal_muscle_tissue_development | 46 | 0.261 | 0.0171 | 0.39 | 0.001 | -0.7296 | 0.2203 |
| go_negative_regulation_of_smooth_muscle_cell_migration | 15 | 0.43 | 0.0151 | 0.548 | 0.0085 | -0.3888 | 0.2403 |
| go_adrenergic_receptor_activity | 14 | 0.46 | 0.0274 | 0.531 | 0.0147 | -0.2074 | 0.3356 |
| go_negative_regulation_of_gliogenesis | 35 | 0.247 | 0.0403 | 0.318 | 0.0183 | -0.3425 | 0.3356 |
| go_cardiac_ventricle_morphogenesis | 60 | 0.193 | 0.0457 | 0.255 | 0.0131 | -0.3829 | 0.3555 |
| go_adrenergic_receptor_signaling_pathway | 21 | 0.356 | 0.0384 | 0.411 | 0.0228 | -0.1911 | 0.3712 |
| go_synaptic_signaling | 399 | 0.0813 | 0.0264 | 0.122 | 0.0026 | -0.6715 | 0.4039 |
| gaussmann_mll_af4_fusion_targets_a_dn | 85 | 0.166 | 0.0412 | 0.206 | 0.0134 | -0.3005 | 0.4055 |
| go_regulation_of_chromosome_organization | 261 | 0.0961 | 0.0257 | 0.129 | 0.0051 | -0.4676 | 0.422 |
| go_central_nervous_system_development | 813 | 0.0625 | 0.02 | 0.0926 | 0.0016 | -0.6898 | 0.4286 |
| go_transcriptional_activator_activity_rna_polymerase_ii_transcription_regulatory_region_sequence_specific_binding | 300 | 0.0982 | 0.022 | 0.127 | 0.0062 | -0.4092 | 0.4317 |
| go_positive_regulation_of_chromatin_modification | 81 | 0.15 | 0.0443 | 0.173 | 0.0283 | -0.1816 | 0.4453 |
| go_regulation_of_muscle_tissue_development | 98 | 0.151 | 0.0377 | 0.173 | 0.0267 | -0.1778 | 0.4477 |
| ginestier_breast_cancer_znf217_amplified_up | 72 | 0.161 | 0.0431 | 0.181 | 0.0373 | -0.145 | 0.4524 |
| ren_alveolar_rhabdomyosarcoma_up | 94 | 0.142 | 0.0481 | 0.158 | 0.0389 | -0.1296 | 0.4619 |
| go_regulation_of_catabolic_process | 690 | 0.0566 | 0.0327 | 0.0683 | 0.0144 | -0.2673 | 0.4721 |
| reactome_chondroitin_sulfate_dermatan_sulfate_metabolism | 42 | 0.242 | 0.0219 | 0.248 | 0.0297 | -0.0336 | 0.4857 |
| go_transcription_factor_activity_rna_polymerase_ii_core_promoter_proximal_region_sequence_specific_binding | 316 | 0.108 | 0.0114 | 0.113 | 0.0106 | -0.0733 | 0.4881 |
| bild_hras_oncogenic_signature | 242 | 0.102 | 0.0351 | 0.105 | 0.0301 | -0.0379 | 0.4928 |
| go_regulation_of_embryonic_development | 107 | 0.171 | 0.0213 | 0.172 | 0.0192 | -0.0084 | 0.4976 |
| go_membrane_depolarization_during_cardiac_muscle_cell_action_potential | 14 | 0.453 | 0.0342 | 0.444 | 0.0454 | 0.0248 | 0.5215 |
| go_positive_regulation_of_ossification | 80 | 0.2 | 0.0204 | 0.17 | 0.0475 | 0.2123 | 0.5712 |
| yan_escape_from_anoikis | 20 | 0.45 | 0.0125 | 0.411 | 0.0089 | 0.1471 | 0.5922 |
| go_transcriptional_activator_activity_rna_polymerase_ii_core_promoter_proximal_region_sequence_specific_binding | 216 | 0.17 | 0.0016 | 0.127 | 0.0168 | 0.5174 | 0.6014 |
| mootha_pyr | 6 | 0.828 | 0.0025 | 0.765 | 0.0132 | 0.1388 | 0.6468 |
| go_prepulse_inhibition | 11 | 0.547 | 0.0368 | 0.449 | 0.0416 | 0.2445 | 0.721 |
| go_regulation_of_rna_polymerase_ii_transcriptional_preinitiation_complex_assembly | 13 | 0.639 | 0.0075 | 0.385 | 0.0185 | 0.7899 | 0.9355 |

**Table S23.** Gene set enrichments meeting nominal significant in attention deficit hyperactivity disorder (ADHD) and *PhoneUse* males. Traits are listed in ascending order by *PhoneUse* significance.

| Enrichment | N genes | ADHD Males | | *PhoneUse* Males | | Difference | |
|---|---|---|---|---|---|---|---|
| | | beta | p | beta | p | z | p |
| go_cation_cation_antiporter_activity | 23 | 0.367 | 0.026 | 0.538 | 0.0006 | -0.6816 | 0.1581 |
| go_alkali_metal_ion_binding | 21 | 0.468 | 0.015 | 0.525 | 0.0011 | -0.2069 | 0.3692 |
| go_ion_antiporter_activity | 45 | 0.219 | 0.0367 | 0.346 | 0.0017 | -0.7483 | 0.2284 |
| go_solute_cation_antiporter_activity | 29 | 0.321 | 0.0232 | 0.449 | 0.0018 | -0.5745 | 0.2266 |
| chow_rassf1_targets_dn | 29 | 0.254 | 0.0496 | 0.381 | 0.0026 | -0.6161 | 0.2284 |
| thillainadesan_znf217_targets_up | 43 | 0.224 | 0.0331 | 0.3 | 0.0065 | -0.4423 | 0.328 |
| dawson_methylated_in_lymphoma_tcl1 | 57 | 0.306 | 0.0085 | 0.32 | 0.0069 | -0.0767 | 0.4673 |
| go_rna_polymerase_ii_transcription_factor_activity_sequence_specific_dna_binding | 590 | 0.0681 | 0.0299 | 0.0821 | 0.0098 | -0.2773 | 0.4673 |
| pid_telomerase_pathway | 67 | 0.193 | 0.0359 | 0.231 | 0.0114 | -0.2583 | 0.4119 |
| yagi_aml_with_11q23_rearranged | 323 | 0.123 | 0.0055 | 0.103 | 0.0125 | 0.3005 | 0.5467 |
| go_negative_regulation_of_gene_expression | 1370 | 0.0411 | 0.0394 | 0.0516 | 0.0128 | -0.3193 | 0.4755 |
| go_nucleic_acid_binding_transcription_factor_activity | 1115 | 0.0538 | 0.023 | 0.0574 | 0.0139 | -0.0959 | 0.4916 |
| myllykangas_amplification_hot_spot_16 | 9 | 0.705 | 0.0162 | 0.67 | 0.0141 | 0.078 | 0.5813 |
| go_regulation_of_peptide_secretion | 193 | 0.106 | 0.0428 | 0.13 | 0.0177 | -0.2753 | 0.4441 |
| mishra_carcinoma_associated_fibroblast_dn | 20 | 0.414 | 0.0134 | 0.359 | 0.0187 | 0.2159 | 0.6264 |
| go_negative_regulation_of_interleukin_17_production | 10 | 0.471 | 0.0387 | 0.517 | 0.0209 | -0.1251 | 0.3937 |
| go_neuron_migration | 102 | 0.274 | 0.0013 | 0.179 | 0.0209 | 0.753 | 0.7111 |
| go_negative_regulation_of_humoral_immune_response | 13 | 0.719 | 0.0021 | 0.521 | 0.0216 | 0.5512 | 0.8771 |
| pid_erbb1_downstream_pathway | 104 | 0.149 | 0.0431 | 0.171 | 0.0237 | -0.1797 | 0.4487 |
| go_regulation_of_lipid_kinase_activity | 50 | 0.227 | 0.0272 | 0.255 | 0.0241 | -0.1602 | 0.4348 |
| go_mechanosensory_behavior | 12 | 0.784 | 0.0013 | 0.519 | 0.0242 | 0.7166 | 0.9398 |
| go_negative_regulation_of_cytosolic_calcium_ion_concentration | 11 | 0.467 | 0.0449 | 0.52 | 0.0272 | -0.1375 | 0.3781 |
| reactome_nuclear_events_kinase_and_transcription_factor_activation | 22 | 0.537 | 0.0006 | 0.332 | 0.0275 | 0.8526 | 0.8852 |
| go_solute_proton_antiporter_activity | 17 | 0.49 | 0.008 | 0.373 | 0.028 | 0.4157 | 0.7535 |
| go_monovalent_cation_proton_antiporter_activity | 10 | 0.651 | 0.0102 | 0.422 | 0.0306 | 0.6361 | 0.9102 |
| go_mitotic_cell_cycle_checkpoint | 134 | 0.126 | 0.0378 | 0.131 | 0.0324 | -0.0499 | 0.4883 |
| bafna_muc4_targets_up | 6 | 0.577 | 0.0483 | 0.66 | 0.0328 | -0.166 | 0.3133 |
| go_neurotransmitter_receptor_activity | 62 | 0.213 | 0.0335 | 0.198 | 0.0343 | 0.0942 | 0.535 |
| geiss_response_to_dsrna_up | 34 | 0.295 | 0.0116 | 0.242 | 0.0351 | 0.2839 | 0.6219 |
| go_activating_transcription_factor_binding | 55 | 0.376 | 0.0004 | 0.202 | 0.0387 | 1.0839 | 0.8461 |
| go_regulation_of_mrna_splicing_via_spliceosome | 58 | 0.277 | 0.0091 | 0.172 | 0.0392 | 0.6883 | 0.7308 |
| biocarta_cdmac_pathway | 15 | 0.381 | 0.04 | 0.387 | 0.0397 | -0.0194 | 0.486 |
| go_c2h2_zinc_finger_domain_binding | 10 | 0.769 | 0.0006 | 0.535 | 0.0419 | 0.6 | 0.9149 |
| ramjaun_apoptosis_by_tgfb1_via_smad4_up | 7 | 0.64 | 0.0416 | 0.529 | 0.044 | 0.2303 | 0.7423 |
| go_gtp_metabolic_process | 22 | 0.349 | 0.0298 | 0.301 | 0.044 | 0.188 | 0.6108 |
| mikkelsen_ips_with_hcp_h3k27me3 | 95 | 0.164 | 0.0459 | 0.16 | 0.0447 | 0.0296 | 0.5094 |
| reactome_glycolysis | 25 | 0.266 | 0.0433 | 0.236 | 0.0456 | 0.1436 | 0.5698 |
| reactome_hdl_mediated_lipid_transport | 15 | 0.375 | 0.0451 | 0.389 | 0.0456 | -0.0439 | 0.4673 |
| go_cognition | 235 | 0.131 | 0.0093 | 0.0916 | 0.0491 | 0.5011 | 0.5913 |

**Table S24.** Linkage disequilibrium score regression (rg) results for 841 UK Biobank traits demonstrating nominally significant genetic correlations with attention deficit hyperactivity disorder (ADHD) and *PhoneUse*.

| Field ID | Trait | ADHD | | *PhoneUse* | | Difference | |
|---|---|---|---|---|---|---|---|
| | | rg | p | rg | p | z | p |
| M25 | Diagnoses - main ICD10: M25 Other joint disorders, not elsewhere classified | 0.9787 | 7.8838E-08 | 0.7912 | 1.9443E-08 | 0.8138 | 0.7441 |
| M13_JOINTOTH | Other specific joint derangements/joint disorders | 0.9635 | 5.908E-09 | 0.7876 | 9.6275E-09 | 0.8177 | 0.7309 |
| 22613_2 | Worked with paints, thinners or glues: Often | 0.9093 | 0.0167 | 0.3696 | 0.0032 | 1.3488 | 0.9705 |
| M13_LIMBPAIN | Pain in limb | 0.8668 | 0.0178 | 0.7301 | 0.0017 | 0.3152 | 0.6838 |
| 22617_3311 | Job SOC coding: NCOs and other ranks | 0.8043 | 0.0007 | 0.588 | 1.6358E-05 | 0.792 | 0.7755 |
| 22601_33113434 | Job coding: non-commissioned officers or other rank of armed forces | 0.7798 | 0.0022 | 0.5821 | 3.3333E-05 | 0.6807 | 0.7555 |
| R13 | Diagnoses - main ICD10: R13 Dysphagia | 0.7732 | 0.0013 | 0.8313 | 0.0047 | -0.1528 | 0.4194 |
| E4_OBESITY | Obesity | 0.7681 | 7.4882E-07 | 0.3507 | 0.0002 | 2.3089 | 0.928 |
| G6_ULLNLE | Lesion of ulnar nerve | 0.763 | 0.0259 | 0.2932 | 0.0416 | 1.2646 | 0.9499 |
| 3741 | Stomach/abdominal pain for 3+ months | 0.76 | 0.0032 | 0.5019 | 0.0013 | 0.8568 | 0.8168 |
| E66 | Diagnoses - main ICD10: E66 Obesity | 0.7508 | 2.9348E-07 | 0.3587 | 0.0002 | 2.2495 | 0.915 |
| 20003_1140923350 | Treatment/medication code: co-dydramol | 0.7492 | 0.0077 | 0.6852 | 0.0012 | 0.1819 | 0.5886 |
| 6141_6 | How are people in household related to participant: Grandchild | 0.7431 | 0.00006416 | 0.4323 | 1.7359E-05 | 1.4704 | 0.8616 |
| AB1_INFECTIONS | Certain infectious and parasitic diseases | 0.7323 | 0.0005 | 0.4376 | 0.0017 | 1.1628 | 0.8488 |
| 6145_5 | Illness, injury, bereavement, stress in last 2 years: Marital separation/divorce | 0.7086 | 6.3829E-05 | 0.6656 | 1.3713E-05 | 0.1836 | 0.5598 |
| E4_OBESITYNAS | Obesity, other/unspecified | 0.6955 | 3.605E-07 | 0.3516 | 0.0001 | 2.0977 | 0.8856 |
| 22620_1 | Job involved shift work: Yes | 0.6927 | 1.9627E-17 | 0.4786 | 6.7541E-12 | 1.9965 | 0.7732 |
| 1259 | Smoking/smokers in household | 0.6847 | 1.0586E-17 | 0.5584 | 6.0991E-19 | 1.2428 | 0.6708 |
| 41231_1 | Hospital episode type: General episode | 0.6841 | 2.5349E-35 | 0.5497 | 4.4175E-34 | 1.8838 | 0.681 |
| 6142_4 | Current employment status: Unable to work because of sickness or disability | 0.6816 | 5.7946E-50 | 0.4522 | 2.8913E-27 | 3.6952 | 0.789 |
| ICDMAIN_ANY_ENTRY | Any ICDMAIN event in hilmo or causes of death | 0.6705 | 3.5265E-49 | 0.5508 | 1.3078E-50 | 2.0455 | 0.6624 |
| 41235 | Spells in hospital | 0.6668 | 1.5482E-47 | 0.5149 | 1.4351E-40 | 2.5296 | 0.7025 |
| 20110_3 | Illnesses of mother: Lung cancer | 0.6664 | 1.1962E-06 | 0.5925 | 2.6801E-06 | 0.3964 | 0.602 |
| 6146_3 | Attendance/disability/mobility allowance: Blue badge | 0.6662 | 4.2263E-43 | 0.5558 | 3.0264E-37 | 1.6948 | 0.6504 |
| 4825 | Noisy workplace | 0.6627 | 1.2634E-31 | 0.3296 | 6.2403E-12 | 4.4923 | 0.8781 |
| I_INFECT_PARASIT | Certain infectious and parasitic diseases | 0.6623 | 6.5426E-05 | 0.324 | 0.0057 | 1.666 | 0.8818 |
| 6159_8 | Pain type(s) experienced in last month: Pain all over the body | 0.6578 | 3.0301E-19 | 0.4727 | 1.8712E-14 | 1.9319 | 0.7414 |
| 41248_1000 | Destinations on discharge from hospital (recoded): Usual Place of residence | 0.6559 | 3.5331E-45 | 0.5678 | 3.5591E-43 | 1.4181 | 0.6211 |
| 22607_2 | Workplace very cold: Often | 0.6558 | 1.1422E-13 | 0.4101 | 7.3695E-09 | 2.1697 | 0.8051 |
| R51 | Diagnoses - main ICD10: R51 Headache | 0.6545 | 1.8894E-06 | 0.6305 | 9.2513E-07 | 0.1276 | 0.5335 |

| | | | | | | | |
|---|---|---|---|---|---|---|---|
| 41215_0 | Detention categories: Informal, not formally detained | 0.6488 | 7.4558E-08 | 0.4784 | 6.4642E-09 | 1.1666 | 0.7245 |
| 20002_1112 | Non-cancer illness code, self-reported: chronic obstructive airways disease/copd | 0.646 | 1.0351E-08 | 0.424 | 1.1129E-05 | 1.4955 | 0.7814 |
| XXI_HEALTHFACTORS | Factors influencing health status and contact with health services | 0.6457 | 3.7721E-11 | 0.507 | 3.143E-11 | 1.119 | 0.6863 |
| 22610_2 | Workplace full of chemical or other fumes: Often | 0.6445 | 0.0004 | 0.5143 | 4.9234E-05 | 0.5906 | 0.6757 |
| M47 | Diagnoses - main ICD10: M47 Spondylosis | 0.6437 | 0.0004 | 0.5107 | 1.2307E-05 | 0.6148 | 0.6792 |
| 6146_2 | Attendance/disability/mobility allowance: Disability living allowance | 0.6399 | 7.3108E-50 | 0.5751 | 3.5405E-53 | 1.1343 | 0.5897 |
| 20003_1140879616 | Treatment/medication code: amitriptyline | 0.6333 | 2.0035E-09 | 0.3549 | 7.2848E-07 | 2.1821 | 0.835 |
| XVIII_MISCFINDINGS | Symptoms, signs and abnormal clinical and laboratory findings, not elsewhere classified | 0.6314 | 4.3034E-43 | 0.5343 | 1.2368E-58 | 1.7159 | 0.633 |
| 22615_2 | Workplace had a lot of diesel exhaust: Often | 0.6297 | 6.1201E-06 | 0.5171 | 6.6131E-06 | 0.6241 | 0.6532 |
| COPD_EARLYANDLATER | COPD, early/later onset | 0.6281 | 1.2417E-11 | 0.3941 | 8.6212E-09 | 2.0301 | 0.7936 |
| 20111_6 | Illnesses of siblings: Chronic bronchitis/emphysema | 0.6275 | 2.0008E-07 | 0.4219 | 9.2888E-06 | 1.3374 | 0.7641 |
| J18 | Diagnoses - main ICD10: J18 Pneumonia, organism unspecified | 0.6274 | 0.0019 | 0.5621 | 2.2539E-05 | 0.2698 | 0.5904 |
| 20003_1140884464 | Treatment/medication code: dihydrocodeine | 0.6263 | 1.4326E-05 | 0.5959 | 2.1122E-05 | 0.1511 | 0.5424 |
| IV_ENDOCRIN_NUTRIT | Endocrine, nutritional and metabolic diseases | 0.6235 | 3.3245E-06 | 0.4606 | 0.00000888 | 0.961 | 0.7157 |
| PULMONARYDG | Other pulmonary diagnosis | 0.6234 | 2.2271E-24 | 0.5488 | 9.4511E-22 | 0.8898 | 0.603 |
| X_RESPIRATORY | Diseases of the respiratory system | 0.6234 | 2.2271E-24 | 0.5488 | 9.4511E-22 | 0.8898 | 0.603 |
| M13_SOFTTISSUEOTH | Other soft tissue disorders, not elsewhere classified | 0.623 | 2.064E-07 | 0.6912 | 2.965E-11 | -0.4295 | 0.4057 |
| M79 | Diagnoses - main ICD10: M79 Other soft tissue disorders, not elsewhere classified | 0.623 | 2.064E-07 | 0.6912 | 2.965E-11 | -0.4295 | 0.4057 |
| 6164_100 | Types of physical activity in last 4 weeks: None of the above | 0.6204 | 1.4003E-40 | 0.4446 | 2.72E-30 | 2.8998 | 0.7308 |
| G43 | Diagnoses - main ICD10: G43 Migraine | 0.6194 | 0.0183 | 0.2602 | 0.0369 | 1.236 | 0.8956 |
| 22606_2 | Workplace very noisy: Often | 0.619 | 7.0542E-19 | 0.4466 | 2.1996E-18 | 1.9929 | 0.7269 |
| 22609_1 | Workplace very dusty: Sometimes | 0.6172 | 1.4298E-09 | 0.5308 | 4.6513E-09 | 0.6333 | 0.6188 |
| 22615_1 | Workplace had a lot of diesel exhaust: Sometimes | 0.6168 | 5.3038E-11 | 0.4819 | 4.3526E-09 | 1.0809 | 0.6816 |
| ASTHMA_PNEUMONIA | Asthma-related pneumonia | 0.6165 | 0.0002 | 0.5603 | 1.5688E-06 | 0.2767 | 0.578 |
| PNEUMONIA | Pneumonias (Asthma/COPD co-morbidities) | 0.6165 | 0.0002 | 0.5603 | 1.5688E-06 | 0.2767 | 0.578 |
| 20523 | Physical violence by partner or ex-partner as an adult | 0.6139 | 1.3162E-15 | 0.4241 | 6.1265E-12 | 1.9266 | 0.7467 |
| 6152_6 | Blood clot, DVT, bronchitis, emphysema, asthma, rhinitis, eczema, allergy diagnosed by doctor: Emphysema/chronic bronchitis | 0.6137 | 1.0862E-23 | 0.4087 | 1.8287E-12 | 2.4313 | 0.7634 |
| COPD_EXCL | COPD differential diagnosis | 0.611 | 2.0542E-24 | 0.5446 | 4.9611E-23 | 0.8158 | 0.5919 |
| ILD_DIFF_DG | ILD differential diagnosis | 0.611 | 2.0542E-24 | 0.5446 | 4.9611E-23 | 0.8158 | 0.5919 |
| 20003_1140923346 | Treatment/medication code: co-codamol | 0.6108 | 9.69E-16 | 0.4748 | 5.9235E-16 | 1.4151 | 0.6829 |
| J44 | Diagnoses - main ICD10: J44 Other chronic obstructive pulmonary disease | 0.6107 | 5.9913E-11 | 0.3916 | 9.0574E-09 | 1.8968 | 0.7784 |
| 4717 | Shortness of breath walking on level ground | 0.607 | 2.3997E-31 | 0.4218 | 1.4041E-16 | 2.5402 | 0.7416 |

| | | | | | | | |
|---|---|---|---|---|---|---|---|
| 680_3 | Own or rent accommodation lived in: Rent - from local authority, local council, housing association | 0.606 | 2.8841E-58 | 0.3944 | 8.0619E-40 | 4.3976 | 0.7705 |
| 20003_1140865634 | Treatment/medication code: omeprazole | 0.6054 | 1.1141E-21 | 0.4644 | 2.4335E-18 | 1.7052 | 0.6892 |
| 22610_1 | Workplace full of chemical or other fumes: Sometimes | 0.6052 | 1.9149E-09 | 0.3637 | 1.863E-06 | 1.9103 | 0.801 |
| 4728 | Leg pain on walking | 0.6019 | 2.2021E-29 | 0.4358 | 5.9281E-23 | 2.3935 | 0.7195 |
| 20002_1113 | Non-cancer illness code, self-reported: emphysema/chronic bronchitis | 0.6019 | 2.6714E-17 | 0.4088 | 1.1823E-09 | 1.9738 | 0.7504 |
| 20003_1140864992 | Treatment/medication code: tramadol | 0.6019 | 1.59E-14 | 0.4891 | 3.3182E-12 | 1.0719 | 0.6535 |
| R06 | Diagnoses - main ICD10: R06 Abnormalities of breathing | 0.6016 | 0.0024 | 0.5044 | 0.0038 | 0.3687 | 0.6331 |
| E4_DM2NOCOMP | Type 2 diabetes without complications | 0.6006 | 0.0051 | 0.5521 | 0.0059 | 0.1651 | 0.5674 |
| 20488 | Physically abused by family as a child | 0.5968 | 2.0144E-31 | 0.347 | 3.0464E-12 | 3.5008 | 0.809 |
| 1873 | Number of full brothers | 0.5957 | 6.5934E-25 | 0.3352 | 1.2172E-10 | 3.3477 | 0.819 |
| M24 | Diagnoses - main ICD10: M24 Other specific joint derangements | 0.5948 | 0.0125 | 0.7276 | 0.0071 | -0.3686 | 0.3211 |
| M18 | Diagnoses - main ICD10: M18 Arthrosis of first carpometacarpal joint | 0.5938 | 0.0065 | 0.4667 | 0.0174 | 0.4334 | 0.6718 |
| R07 | Diagnoses - main ICD10: R07 Pain in throat and chest | 0.5917 | 4.3472E-28 | 0.5289 | 2.1844E-34 | 0.9093 | 0.587 |
| XI_DIGESTIVE | Diseases of the digestive system | 0.5906 | 1.6519E-48 | 0.4505 | 1.4799E-39 | 2.6506 | 0.688 |
| 20003_1140864752 | Treatment/medication code: lansoprazole | 0.5888 | 7.5639E-18 | 0.4288 | 5.1847E-16 | 1.8504 | 0.7122 |
| M13_MUSCULOSKELEOTH | Other disorders of the musculoskeletal system and connective tissue | 0.5864 | 0.0312 | 0.3481 | 0.0188 | 0.7692 | 0.7979 |
| COLITNONINFNAS | Noninfectious colitis NAS | 0.5857 | 1.9277E-08 | 0.4524 | 3.9272E-08 | 1.0039 | 0.6796 |
| 100160 | Low calorie drink intake | 0.5853 | 0.0314 | 0.3826 | 0.008 | 0.6584 | 0.761 |
| 20117_1 | Alcohol drinker status: Previous | 0.5834 | 1.0423E-23 | 0.3321 | 8.8247E-11 | 3.2451 | 0.8104 |
| 22607_1 | Workplace very cold: Sometimes | 0.583 | 1.1422E-20 | 0.3839 | 1.6282E-11 | 2.3537 | 0.757 |
| 1269 | Exposure to tobacco smoke at home | 0.582 | 2.6583E-20 | 0.5925 | 4.2704E-25 | -0.1233 | 0.4853 |
| 20003_1140909708 | Treatment/medication code: furosemide | 0.5818 | 1.6207E-05 | 0.3284 | 0.0015 | 1.4903 | 0.8124 |
| 189 | Townsend deprivation index at recruitment | 0.5794 | 9.758E-46 | 0.2934 | 2.8867E-17 | 5.3398 | 0.8416 |
| 22609_2 | Workplace very dusty: Often | 0.5794 | 1.0564E-14 | 0.4125 | 3.8988E-10 | 1.673 | 0.7204 |
| 1508_4 | Coffee type: Other type of coffee | 0.5768 | 0.0107 | 0.2962 | 0.0067 | 1.1173 | 0.837 |
| XIV_GENITOURINARY | Diseases of the genitourinary system | 0.5757 | 1.0084E-27 | 0.4655 | 2.5853E-26 | 1.6049 | 0.6501 |
| K21 | Diagnoses - main ICD10: K21 Gastro-oesophageal reflux disease | 0.5756 | 6.9386E-14 | 0.3255 | 8.198E-10 | 2.6779 | 0.8093 |
| N39 | Diagnoses - main ICD10: N39 Other disorders of urinary system | 0.5738 | 5.9024E-11 | 0.5707 | 9.7104E-14 | 0.0266 | 0.5043 |
| M13_IMPINGEMENT | Impingement syndrome of shoulder | 0.5729 | 1.9309E-09 | 0.4299 | 2.1449E-07 | 1.1314 | 0.6916 |
| M13_DORSALGIANAS | Other/unspecified dorsalgia | 0.5712 | 3.6252E-05 | 0.4479 | 2.7171E-05 | 0.7059 | 0.667 |
| N35 | Diagnoses - main ICD10: N35 Urethral stricture | 0.5687 | 0.0347 | 0.322 | 0.0129 | 0.8256 | 0.806 |
| 6145_6 | Illness, injury, bereavement, stress in last 2 years: Financial difficulties | 0.5681 | 5.6011E-42 | 0.5104 | 8.9585E-52 | 1.0746 | 0.58 |
| 3404 | Neck/shoulder pain for 3+ months | 0.5667 | 2.3559E-06 | 0.4178 | 3.3026E-06 | 0.9929 | 0.6989 |
| 6138_100 | Qualifications: None of the above | 0.5662 | 4.9467E-74 | 0.3776 | 1.5096E-53 | 4.7637 | 0.7454 |
| 20111_2 | Illnesses of siblings: Stroke | 0.5642 | 0.0004 | 0.3936 | 0.0028 | 0.8227 | 0.7248 |

| | | | | | | | |
|---|---|---|---|---|---|---|---|
| K52 | Diagnoses - main ICD10: K52 Other non-infective gastro-enteritis and colitis | 0.5628 | 7.9488E-08 | 0.4496 | 9.1288E-08 | 0.8424 | 0.654 |
| 826 | Job involves shift work | 0.5602 | 1.0207E-22 | 0.4014 | 1.6265E-16 | 2.116 | 0.7108 |
| 22606_1 | Workplace very noisy: Sometimes | 0.5594 | 3.7413E-08 | 0.3865 | 1.0736E-06 | 1.3413 | 0.7274 |
| 20527 | Been involved in combat or exposed to war-zone | 0.5594 | 1.7367E-05 | 0.5117 | 2.4014E-07 | 0.2915 | 0.5663 |
| 2178 | Overall health rating | 0.5579 | 3.183E-100 | 0.3516 | 2.219E-36 | 5.3805 | 0.7648 |
| 4548 | Health satisfaction | 0.5574 | 1.3175E-33 | 0.3042 | 1.595E-13 | 4.0953 | 0.8122 |
| 20002_1465 | Non-cancer illness code, self-reported: osteoarthritis | 0.5562 | 8.0882E-29 | 0.3289 | 9.9467E-16 | 3.5195 | 0.7868 |
| 20002_1465 | Non-cancer illness code, self-reported: osteoarthritis | 0.5562 | 8.0882E-29 | 0.3289 | 9.9467E-16 | 3.5195 | 0.7868 |
| 20511 | Recent poor appetite or overeating | 0.5558 | 3.2305E-27 | 0.3524 | 1.4357E-12 | 2.842 | 0.7617 |
| XIII_MUSCULOSKELET | Diseases of the musculoskeletal system and connective tissue | 0.5557 | 9.2245E-58 | 0.4998 | 7.9285E-51 | 1.1623 | 0.5776 |
| 4642 | Ever manic/hyper for 2 days | 0.5551 | 3.2273E-10 | 0.4444 | 2.3009E-08 | 0.9317 | 0.6508 |
| 5474 | Leg pain when walking uphill or hurrying | 0.5549 | 1.8605E-05 | 0.3212 | 0.001 | 1.4421 | 0.7933 |
| 1239 | Current tobacco smoking | 0.5513 | 3.6641E-59 | 0.4342 | 3.2529E-43 | 2.5265 | 0.659 |
| 20107_6 | Illnesses of father: Chronic bronchitis/emphysema | 0.5504 | 4.5241E-18 | 0.2739 | 3.1785E-07 | 3.3274 | 0.8334 |
| 22506_113 | Tobacco smoking: Ex-smoker | 0.5496 | 3.1837E-22 | 0.3777 | 5.0486E-16 | 2.3422 | 0.7263 |
| 22608_1 | Workplace very hot: Sometimes | 0.5486 | 2.0438E-14 | 0.3277 | 5.6584E-07 | 2.2746 | 0.7803 |
| 20116_2 | Smoking status: Current | 0.5476 | 6.4371E-54 | 0.429 | 5.4191E-40 | 2.4714 | 0.661 |
| R10 | Diagnoses - main ICD10: R10 Abdominal and pelvic pain | 0.5474 | 7.1061E-20 | 0.5273 | 3.1561E-22 | 0.2482 | 0.528 |
| J45 | Diagnoses - main ICD10: J45 Asthma | 0.5454 | 0.0003 | 0.2382 | 0.0157 | 1.7106 | 0.8588 |
| 20003_1140879540 | Treatment/medication code: fluoxetine | 0.5441 | 4.8471E-06 | 0.1995 | 0.0154 | 2.3817 | 0.8861 |
| T81 | Diagnoses - main ICD10: T81 Complications of procedures, not elsewhere classified | 0.5428 | 5.4284E-05 | 0.4759 | 2.6586E-05 | 0.3804 | 0.5926 |
| 816 | Job involves heavy manual or physical work | 0.542 | 1.3984E-48 | 0.304 | 2.6911E-23 | 4.9569 | 0.7976 |
| 1787 | Maternal smoking around birth | 0.5397 | 2.8018E-49 | 0.4369 | 9.8514E-34 | 1.9997 | 0.6405 |
| XIX_INJURY_POISON | Injury, poisoning and certain other consequences of external causes | 0.5372 | 1.0005E-27 | 0.4288 | 3.7287E-20 | 1.5996 | 0.6478 |
| 20490 | Sexually molested as a child | 0.5367 | 1.416E-12 | 0.2345 | 0.0007 | 2.9444 | 0.8549 |
| M13_ENTESOPATHYOTH | Other enthesopathies | 0.5365 | 0.0016 | 0.3066 | 0.0268 | 1.048 | 0.7895 |
| M13_SHOULDER | Shoulder lesions | 0.5351 | 1.6629E-13 | 0.4442 | 1.1219E-10 | 0.9082 | 0.6248 |
| M75 | Diagnoses - main ICD10: M75 Shoulder lesions | 0.5351 | 1.7664E-13 | 0.4455 | 9.7342E-11 | 0.8952 | 0.6231 |
| K29 | Diagnoses - main ICD10: K29 Gastritis and duodenitis | 0.5341 | 6.1517E-16 | 0.4119 | 7.3105E-13 | 1.3971 | 0.6656 |
| 20110_6 | Illnesses of mother: Chronic bronchitis/emphysema | 0.5333 | 6.7546E-14 | 0.4644 | 1.7866E-15 | 0.7482 | 0.5953 |
| M77 | Diagnoses - main ICD10: M77 Other enthesopathies | 0.5321 | 0.0015 | 0.3049 | 0.0241 | 1.054 | 0.7867 |
| M13_LOWBACKPAIN | Low back pain | 0.5311 | 1.5665E-16 | 0.5202 | 2.1364E-17 | 0.1226 | 0.5152 |
| 6154_5 | Medication for pain relief, constipation, heartburn: Omeprazole (e.g. Zanprol) | 0.5295 | 3.9834E-19 | 0.4151 | 3.4456E-19 | 1.5209 | 0.6556 |
| 2188 | Long-standing illness, disability or infirmity | 0.5292 | 7.8346E-56 | 0.3137 | 2.7928E-28 | 4.8983 | 0.7746 |
| 6159_6 | Pain type(s) experienced in last month: Hip pain | 0.5291 | 1.1832E-31 | 0.3453 | 3.5703E-17 | 3.0119 | 0.74 |
| M13_ARTHRITISNAS | Ohter specific/unspecified arthritis | 0.5287 | 0.0039 | 0.2836 | 0.0201 | 1.114 | 0.8045 |

| | | | | | | | |
|---|---|---|---|---|---|---|---|
| VI_NERVOUS | Diseases of the nervous system | 0.5261 | 1.0806E-21 | 0.5002 | 1.7943E-23 | 0.3481 | 0.5361 |
| M13_DORSALGIA | Dorsalgia | 0.5251 | 1.6884E-22 | 0.5077 | 9.8511E-24 | 0.2356 | 0.5243 |
| M54 | Diagnoses - main ICD10: M54 Dorsalgia | 0.5251 | 1.6884E-22 | 0.5077 | 9.8511E-24 | 0.2356 | 0.5243 |
| 6159_3 | Pain type(s) experienced in last month: Neck or shoulder pain | 0.5233 | 7.9345E-39 | 0.3962 | 6.8498E-31 | 2.4052 | 0.6718 |
| 6146_1 | Attendance/disability/mobility allowance: Attendance allowance | 0.5221 | 2.1123E-10 | 0.5552 | 1.5866E-13 | -0.2971 | 0.4539 |
| 20422 | More irritable than usual during worst period of anxiety | 0.5213 | 0.0001 | 0.3036 | 0.0041 | 1.268 | 0.7769 |
| J34 | Diagnoses - main ICD10: J34 Other disorders of nose and nasal sinuses | 0.5203 | 0.0002 | 0.4236 | 0.0046 | 0.4742 | 0.6325 |
| XII_SKIN_SUBCUTAN | Diseases of the skin and subcutaneous tissue | 0.5166 | 3.7818E-06 | 0.5029 | 4.3315E-07 | 0.0915 | 0.5191 |
| 4653 | Ever highly irritable/argumentative for 2 days | 0.5157 | 4.6039E-22 | 0.3573 | 1.806E-10 | 2.0471 | 0.7103 |
| 20487 | Felt hated by family member as a child | 0.513 | 1.6059E-21 | 0.2815 | 5.1729E-09 | 3.2049 | 0.7911 |
| 2316 | Wheeze or whistling in the chest in last year | 0.5129 | 1.7435E-56 | 0.3823 | 4.7708E-36 | 2.935 | 0.6762 |
| R04 | Diagnoses - main ICD10: R04 Haemorrhage from respiratory passages | 0.511 | 0.0031 | 0.5122 | 8.8296E-05 | -0.0055 | 0.4983 |
| 1767 | Adopted as a child | 0.5105 | 3.495E-07 | 0.5463 | 3.781E-11 | -0.2757 | 0.4501 |
| 20003_1140881856 | Treatment/medication code: salbutamol | 0.5102 | 1.9867E-06 | 0.2514 | 0.0014 | 1.9449 | 0.8175 |
| 6149_4 | Mouth/teeth dental problems: Loose teeth | 0.5095 | 4.5917E-18 | 0.2759 | 1.0753E-07 | 2.9785 | 0.7932 |
| 6138_5 | Qualifications: NVQ or HND or HNC or equivalent | 0.5094 | 2.8888E-14 | 0.2911 | 3.1908E-10 | 2.6805 | 0.7776 |
| 3606 | Chest pain or discomfort walking normally | 0.5089 | 2.7909E-15 | 0.4738 | 1.4721E-21 | 0.4315 | 0.5489 |
| 2296 | Falls in the last year | 0.5088 | 4.8929E-34 | 0.3344 | 2.9585E-24 | 3.2785 | 0.7292 |
| 1883 | Number of full sisters | 0.5054 | 1.4132E-15 | 0.3506 | 4.6089E-12 | 1.9087 | 0.706 |
| VIII_EAR_MASTOID | Diseases of the ear and mastoid process | 0.5048 | 3.2858E-06 | 0.4118 | 4.2058E-06 | 0.6612 | 0.6276 |
| 6157_2 | Why stopped smoking: Doctor's advice | 0.5046 | 0.0012 | 0.4481 | 2.4309E-06 | 0.3103 | 0.5784 |
| 20002_1311 | Non-cancer illness code, self-reported: spine arthritis/spondylitis | 0.504 | 9.1345E-05 | 0.6261 | 5.5141E-06 | -0.6473 | 0.3346 |
| 6150_3 | Vascular/heart problems diagnosed by doctor: Stroke | 0.5038 | 0.0018 | 0.4178 | 2.9765E-05 | 0.4536 | 0.6183 |
| 2492 | Taking other prescription medications | 0.5033 | 1.3744E-44 | 0.338 | 7.8151E-34 | 3.6356 | 0.7185 |
| 2335 | Chest pain or discomfort | 0.5023 | 3.8507E-43 | 0.3116 | 4.7828E-18 | 3.7198 | 0.7477 |
| 20111_3 | Illnesses of siblings: Lung cancer | 0.5015 | 0.0085 | 0.5509 | 0.0001 | -0.2062 | 0.4314 |
| M15 | Diagnoses - main ICD10: M15 Polyarthrosis | 0.5012 | 0.0009 | 0.3114 | 0.0092 | 0.9836 | 0.7467 |
| M13_POLYARTHROPATHIES | #Polyarthropathies | 0.5 | 2.6748E-06 | 0.3301 | 0.0003 | 1.2157 | 0.7239 |
| T39 | Diagnoses - main ICD10: T39 Poisoning by nonopioid analgesics, antipyretics and antirheumatics | 0.5 | 0.0117 | 0.3157 | 0.0072 | 0.7993 | 0.7405 |
| K11_OTHGASTR | Other gastritis (incl. Duodenitis) | 0.499 | 7.2701E-12 | 0.3996 | 1.0225E-09 | 1.0157 | 0.636 |
| 2060 | Frequency of unenthusiasm / disinterest in last 2 weeks | 0.4947 | 1.1331E-35 | 0.2887 | 2.7959E-13 | 3.6784 | 0.7645 |
| 20516 | Recent restlessness | 0.4945 | 8.7257E-13 | 0.3659 | 1.4657E-10 | 1.4334 | 0.6737 |
| 22608_2 | Workplace very hot: Often | 0.4929 | 4.7395E-09 | 0.3518 | 3.969E-07 | 1.2931 | 0.6893 |
| 20003_1141177600 | Treatment/medication code: rosiglitazone | 0.4929 | 0.0001 | 0.2354 | 0.0212 | 1.5683 | 0.8163 |
| 2814 | Ever used hormone-replacement therapy (HRT) | 0.4926 | 2.6624E-26 | 0.3937 | 1.6893E-21 | 1.5921 | 0.6354 |
| M51 | Diagnoses - main ICD10: M51 Other intervertebral disk disorders | 0.4918 | 5.4743E-07 | 0.3633 | 1.2784E-06 | 1.0399 | 0.6735 |

| | | | | | | | |
|---|---|---|---|---|---|---|---|
| 20003_1141162764 | Treatment/medication code: tolterodine l-tartrate | 0.4918 | 0.0305 | 0.3019 | 0.0393 | 0.7025 | 0.7468 |
| 22613_1 | Worked with paints, thinners or glues: Sometimes | 0.4904 | 1.7862E-08 | 0.2871 | 8.1105E-05 | 1.7909 | 0.7616 |
| 22611_2 | Workplace had a lot of cigarette smoke from other people smoking: Often | 0.4902 | 1.347E-07 | 0.3325 | 5.0638E-06 | 1.3346 | 0.7095 |
| K43 | Diagnoses - main ICD10: K43 Ventral hernia | 0.4893 | 5.7122E-06 | 0.3565 | 0.0001 | 0.9301 | 0.6789 |
| 6159_4 | Pain type(s) experienced in last month: Back pain | 0.4891 | 1.1294E-32 | 0.3667 | 7.6268E-32 | 2.3721 | 0.6658 |
| 6159_7 | Pain type(s) experienced in last month: Knee pain | 0.4886 | 6.0568E-39 | 0.3378 | 9.2019E-25 | 3.0274 | 0.7012 |
| 806 | Job involves mainly walking or standing | 0.4863 | 1.1064E-32 | 0.3146 | 1.8809E-22 | 3.2995 | 0.726 |
| 20548_9 | Manifestations of mania or irritability: I was more active than usual | 0.4862 | 2.7931E-06 | 0.3046 | 3.4353E-05 | 1.4278 | 0.7375 |
| M13_SPONDYLOPATHY | #Spondylopathies | 0.4856 | 2.5885E-10 | 0.3699 | 1.6901E-06 | 1.0618 | 0.6572 |
| 6153_4 | Medication for cholesterol, blood pressure, diabetes, or take exogenous hormones: Hormone replacement therapy | 0.4846 | 8.442E-10 | 0.4654 | 1.2516E-09 | 0.1745 | 0.5268 |
| K81 | Diagnoses - main ICD10: K81 Cholecystitis | 0.4833 | 0.0225 | 0.3565 | 0.0198 | 0.4853 | 0.6714 |
| D17 | Diagnoses - main ICD10: D17 Benign lipomatous neoplasm | 0.4827 | 0.0008 | 0.2547 | 0.0231 | 1.2499 | 0.7875 |
| 20003_1140874420 | Treatment/medication code: quinine | 0.4825 | 2.0291E-08 | 0.412 | 4.8538E-07 | 0.5936 | 0.5974 |
| M13_OTHERJOINT | #Other joint disorders | 0.4789 | 5.1526E-21 | 0.486 | 2.6121E-22 | -0.0995 | 0.4901 |
| 3090 | Used an inhaler for chest within last hour | 0.4765 | 0.0007 | 0.2261 | 0.0346 | 1.4128 | 0.8096 |
| K63 | Diagnoses - main ICD10: K63 Other diseases of intestine | 0.4764 | 0.0002 | 0.5728 | 0.0002 | -0.4793 | 0.3679 |
| 1279 | Exposure to tobacco smoke outside home | 0.4763 | 1.3245E-27 | 0.5064 | 7.0821E-46 | -0.534 | 0.4581 |
| 20002_1081 | Non-cancer illness code, self-reported: stroke | 0.4751 | 0.003 | 0.4434 | 0.00009672 | 0.1616 | 0.5442 |
| 135 | Number of self-reported non-cancer illnesses | 0.474 | 6.8236E-47 | 0.2794 | 7.5984E-26 | 4.5912 | 0.7521 |
| 20118_5 | Home area population density - urban or rural: England/Wales - Urban - less sparse | 0.4737 | 4.5841E-07 | 0.1916 | 0.0152 | 2.3001 | 0.8382 |
| M13_SOFTOVERUSE | Soft tissue disorders related to use, overuse and pressure | 0.4731 | 0.0439 | 0.5069 | 0.0156 | -0.1074 | 0.4529 |
| 20111_9 | Illnesses of siblings: Diabetes | 0.473 | 2.3078E-17 | 0.2519 | 2.117E-07 | 2.9879 | 0.7805 |
| 3773 | Knee pain for 3+ months | 0.4706 | 4.2937E-05 | 0.3192 | 3.9347E-06 | 1.128 | 0.7019 |
| 1538_1 | Major dietary changes in the last 5 years: Yes, because of illness | 0.4705 | 2.9708E-26 | 0.3163 | 2.5685E-16 | 2.621 | 0.7053 |
| 3571 | Back pain for 3+ months | 0.4704 | 6.2192E-12 | 0.3496 | 1.7669E-09 | 1.346 | 0.6638 |
| M70 | Diagnoses - main ICD10: M70 Soft tissue disorders related to use, overuse and pressure | 0.4696 | 0.0413 | 0.5032 | 0.015 | -0.1086 | 0.4532 |
| 2277 | Frequency of solarium/sunlamp use | 0.4693 | 9.562E-20 | 0.5888 | 1.2248E-49 | -1.8338 | 0.3379 |
| 20111_1 | Illnesses of siblings: Heart disease | 0.4689 | 5.8192E-12 | 0.2896 | 4.9335E-10 | 2.1729 | 0.7348 |
| 2080 | Frequency of tiredness / lethargy in last 2 weeks | 0.4685 | 4.1265E-52 | 0.2685 | 4.5296E-19 | 4.6441 | 0.758 |
| ASTHMA_HOSPITAL1 | Asthma, hospital admissions 1 | 0.4678 | 0.0001 | 0.2649 | 0.0038 | 1.3371 | 0.7612 |
| ASTHMA_CHILD | Childhood asthma (age<16) | 0.4674 | 8.7168E-05 | 0.2536 | 0.0056 | 1.4235 | 0.7728 |
| J10_ASTHMA | Asthma | 0.4674 | 8.7168E-05 | 0.2536 | 0.0056 | 1.4235 | 0.7728 |
| J10_ASTHMA_MAIN | Asthma | 0.4674 | 8.7168E-05 | 0.2536 | 0.0056 | 1.4235 | 0.7728 |

| | | | | | | | |
|---|---|---|---|---|---|---|---|
| K11_GASTRODUOULC | Gastroduodenal ulcer | 0.4672 | 0.0089 | 0.4632 | 0.0165 | 0.0152 | 0.5056 |
| 137 | Number of treatments/medications taken | 0.4664 | 4.7927E-41 | 0.3173 | 1.7283E-31 | 3.3757 | 0.6991 |
| G6_EPIPAROX | Episodal and paroxysmal disorders | 0.4651 | 1.5319E-07 | 0.3467 | 3.4463E-06 | 1.0217 | 0.6607 |
| 20003_1140883548 | Treatment/medication code: ipratropium | 0.4651 | 0.0021 | 0.283 | 0.0294 | 0.9142 | 0.738 |
| 20003_1140884444 | Treatment/medication code: codeine | 0.4648 | 6.2363E-05 | 0.1617 | 0.0489 | 2.1316 | 0.8556 |
| 20003_1141168318 | Treatment/medication code: clopidogrel | 0.4645 | 0.0003 | 0.4005 | 0.0002 | 0.3773 | 0.5886 |
| 136 | Number of operations, self-reported | 0.4629 | 1.5173E-28 | 0.4549 | 1.7561E-48 | 0.1535 | 0.5112 |
| K20 | Diagnoses - main ICD10: K20 Oesophagitis | 0.4621 | 0.0067 | 0.518 | 0.0002 | -0.2525 | 0.4224 |
| 20003_1140879406 | Treatment/medication code: ranitidine | 0.462 | 7.4634E-05 | 0.3407 | 0.0004 | 0.8035 | 0.6644 |
| 20003_1140872228 | Treatment/medication code: gabapentin | 0.462 | 0.0003 | 0.432 | 0.0005 | 0.1679 | 0.5418 |
| 20002_1142 | Non-cancer illness code, self-reported: gastric/stomach ulcers | 0.4579 | 0.0244 | 0.5614 | 0.0232 | -0.3232 | 0.3586 |
| C_STROKE | STROKE | 0.4575 | 0.0189 | 0.4411 | 0.0059 | 0.0651 | 0.5229 |
| 20002_99999 | Non-cancer illness code, self-reported: unclassifiable | 0.4572 | 0.0251 | 0.3139 | 0.0306 | 0.5721 | 0.692 |
| 6145_1 | Illness, injury, bereavement, stress in last 2 years: Serious illness, injury or assault to yourself | 0.4564 | 1.364E-15 | 0.3154 | 2.4049E-09 | 1.813 | 0.6892 |
| T40 | Diagnoses - main ICD10: T40 Poisoning by narcotics and psychodysleptics [hallucinogens] | 0.4548 | 0.0114 | 0.4373 | 0.021 | 0.067 | 0.5244 |
| 20548_2 | Manifestations of mania or irritability: I was more restless than usual | 0.4541 | 1.274E-11 | 0.342 | 3.6364E-08 | 1.2261 | 0.6526 |
| 6158_1 | Why reduced smoking: Illness or ill health | 0.4528 | 0.0004 | 0.2291 | 0.0202 | 1.3881 | 0.7832 |
| 1050 | Time spend outdoors in summer | 0.4521 | 4.1671E-48 | 0.4211 | 7.9142E-45 | 0.7186 | 0.5432 |
| 6154_4 | Medication for pain relief, constipation, heartburn: Ranitidine (e.g. Zantac) | 0.4506 | 3.5906E-05 | 0.3291 | 0.0007 | 0.8304 | 0.6647 |
| 20003_99999 | Treatment/medication code: Free-text entry, unable to be coded | 0.4505 | 0.0094 | 0.4275 | 0.0016 | 0.1046 | 0.5321 |
| M13 | Diagnoses - main ICD10: M13 Other arthritis | 0.4489 | 0.001 | 0.3389 | 0.0029 | 0.6191 | 0.6499 |
| RHEU_ARTHRITIS_OTH | Other arthritis (FG) | 0.4489 | 0.001 | 0.3389 | 0.0029 | 0.6191 | 0.6499 |
| 6145_3 | Illness, injury, bereavement, stress in last 2 years: Death of a close relative | 0.4471 | 1.7098E-06 | 0.4301 | 2.721E-08 | 0.1401 | 0.5237 |
| E4_DM2 | Type 2 diabetes | 0.4464 | 0.0001 | 0.4195 | 3.8515E-05 | 0.1729 | 0.5375 |
| 20532 | Did your sleep change? | 0.4455 | 0.0004 | 0.3597 | 0.0002 | 0.5433 | 0.618 |
| 680_4 | Own or rent accommodation lived in: Rent - from private landlord or letting agency | 0.4449 | 1.3077E-07 | 0.4702 | 2.5568E-08 | -0.2121 | 0.4647 |
| R31 | Diagnoses - main ICD10: R31 Unspecified haematuria | 0.4441 | 1.9763E-06 | 0.2836 | 0.0004 | 1.3093 | 0.7128 |
| 2844 | Had other major operations | 0.444 | 6.396E-14 | 0.407 | 9.182E-17 | 0.4819 | 0.5515 |
| I9_PAD | Peripheral artery disease | 0.4437 | 0.0108 | 0.4006 | 0.0038 | 0.1936 | 0.5599 |
| 20003_1140860834 | Treatment/medication code: glyceryl trinitrate | 0.4432 | 0.0155 | 0.5679 | 0.037 | -0.38 | 0.3313 |
| KRA_PSY_PERSON | Personality disorders | 0.4426 | 0.0123 | 0.3097 | 0.0219 | 0.5971 | 0.6791 |
| 20417 | Tense, sore, or aching muscles during worst period of anxiety | 0.4425 | 1.3936E-08 | 0.4732 | 1.0777E-08 | -0.2699 | 0.4572 |
| 20003_1140863152 | Treatment/medication code: diazepam | 0.4405 | 0.0006 | 0.4323 | 0.0018 | 0.0435 | 0.5114 |
| KNEE_ARTHROSIS | Gonarthrosis [arthrosis of knee](FG) | 0.4398 | 1.0661E-20 | 0.3482 | 5.684E-18 | 1.4777 | 0.6257 |

| | | | | | | | |
|---|---|---|---|---|---|---|---|
| M17 | Diagnoses - main ICD10: M17 Gonarthrosis [arthrosis of knee] | 0.4398 | 1.0661E-20 | 0.3482 | 5.684E-18 | 1.4777 | 0.6257 |
| 6138_4 | Qualifications: CSEs or equivalent | 0.4362 | 1.3046E-14 | 0.2224 | 5.7968E-07 | 2.9695 | 0.7728 |
| 20521 | Belittlement by partner or ex-partner as an adult | 0.4342 | 6.0014E-15 | 0.2655 | 4.055E-08 | 2.2885 | 0.7225 |
| K44 | Diagnoses - main ICD10: K44 Diaphragmatic hernia | 0.4318 | 6.2267E-07 | 0.512 | 1.0441E-13 | -0.7247 | 0.3895 |
| IBD_PULM | Respiratory diseases principally affecting the interstitium, IBD co-morbidites | 0.4313 | 0.0492 | 0.3763 | 0.0274 | 0.198 | 0.5763 |
| M13_ARTHROSIS | #Arthrosis | 0.4309 | 9.1959E-25 | 0.3838 | 4.5645E-23 | 0.8248 | 0.5655 |
| 2227 | Other eye problems | 0.4309 | 8.7617E-09 | 0.2279 | 0.0006 | 2.0281 | 0.7613 |
| N92 | Diagnoses - main ICD10: N92 Excessive, frequent and irregular menstruation | 0.4301 | 1.188E-07 | 0.4127 | 1.6403E-07 | 0.1538 | 0.5243 |
| 2664_1 | Reason for reducing amount of alcohol drunk: Illness or ill health | 0.4297 | 0.0006 | 0.2382 | 0.0231 | 1.1773 | 0.7486 |
| E11 | Diagnoses - main ICD10: E11 Non-insulin-dependent diabetes mellitus | 0.4295 | 0.0002 | 0.4015 | 3.5566E-05 | 0.1873 | 0.539 |
| 20548_3 | Manifestations of mania or irritability: My thoughts were racing | 0.428 | 5.7549E-10 | 0.3613 | 4.6423E-09 | 0.72 | 0.5923 |
| FIBRO_COMORB | Fibromyalgia related co-morbidities | 0.428 | 0.0104 | 0.3338 | 0.0038 | 0.4639 | 0.6292 |
| 2834 | Bilateral oophorectomy (both ovaries removed) | 0.4269 | 1.9422E-12 | 0.3529 | 9.776E-11 | 0.908 | 0.6022 |
| 20526 | Been in serious accident believed to be life-threatening | 0.4269 | 1.8148E-05 | 0.2621 | 0.0008 | 1.2995 | 0.7179 |
| G6_NERPLEX | Nerve, nerve root and plexus disorders | 0.4263 | 8.97E-16 | 0.4113 | 7.6669E-21 | 0.218 | 0.5209 |
| 22612_1 | Worked with materials containing asbestos: Sometimes | 0.4261 | 5.9973E-06 | 0.2945 | 0.0002 | 1.0733 | 0.6774 |
| 4581 | Financial situation satisfaction | 0.4257 | 2.0431E-19 | 0.398 | 2.3719E-22 | 0.4435 | 0.5386 |
| 20536_1 | Weight change during worst episode of depression: Gained weight | 0.425 | 1.2779E-09 | 0.2836 | 2.5975E-06 | 1.5305 | 0.6896 |
| C34 | Diagnoses - main ICD10: C34 Malignant neoplasm of bronchus and lung | 0.4249 | 0.0018 | 0.3777 | 0.0073 | 0.2414 | 0.5656 |
| 20530 | Witnessed sudden violent death | 0.4242 | 2.7842E-08 | 0.2741 | 2.5995E-05 | 1.4944 | 0.7003 |
| IX_CIRCULATORY | Diseases of the circulatory system | 0.4238 | 1.5223E-23 | 0.3519 | 5.1015E-22 | 1.2852 | 0.5993 |
| 2664_4 | Reason for reducing amount of alcohol drunk: Financial reasons | 0.4211 | 0.0007 | 0.3575 | 0.0001 | 0.4091 | 0.5881 |
| 2070 | Frequency of tenseness / restlessness in last 2 weeks | 0.421 | 1.956E-28 | 0.265 | 3.7592E-12 | 2.8952 | 0.7074 |
| 6149_6 | Mouth/teeth dental problems: Dentures | 0.4208 | 4.3583E-32 | 0.3593 | 3.1925E-25 | 1.237 | 0.5852 |
| 20548_5 | Manifestations of mania or irritability: I needed less sleep than usual | 0.4206 | 2.5513E-05 | 0.3995 | 1.3258E-05 | 0.1556 | 0.5294 |
| 6159_5 | Pain type(s) experienced in last month: Stomach or abdominal pain | 0.4205 | 1.2158E-12 | 0.2725 | 1.1425E-07 | 1.8877 | 0.6978 |
| 22504 | Bring up phlegm/sputum/mucus on most days | 0.4192 | 8.7447E-06 | 0.1889 | 0.013 | 1.9015 | 0.7899 |
| 6144_3 | Never eat eggs, dairy, wheat, sugar: Wheat products | 0.4192 | 0.0063 | 0.534 | 1.8043E-05 | -0.5806 | 0.3439 |
| 2050 | Frequency of depressed mood in last 2 weeks | 0.4191 | 1.4108E-20 | 0.2058 | 3.6847E-06 | 3.3666 | 0.7723 |
| 777 | Frequency of travelling from home to job workplace | 0.4187 | 8.5264E-10 | 0.1565 | 0.0026 | 3.0594 | 0.8206 |
| 3456 | Number of cigarettes currently smoked daily (current cigarette smokers) | 0.4165 | 2.2233E-06 | 0.2545 | 0.0003 | 1.4319 | 0.7146 |
| 4631 | Ever unenthusiastic/disinterested for a whole week | 0.4163 | 1.4446E-13 | 0.1295 | 0.0056 | 3.9174 | 0.8422 |
| 4836 | Loud music exposure frequency | 0.4157 | 1.8142E-08 | 0.2177 | 0.0003 | 2.0873 | 0.7558 |
| 23115 | Leg fat percentage (left) | 0.415 | 5.1723E-58 | 0.321 | 2.3164E-42 | 2.6935 | 0.6289 |

| | | | | | | | |
|---|---|---|---|---|---|---|---|
| 20502 | Ever had period extreme irritability | 0.4141 | 2.0574E-15 | 0.2043 | 0.0002 | 2.7907 | 0.7686 |
| 23111 | Leg fat percentage (right) | 0.4128 | 7.0231E-60 | 0.3139 | 1.0626E-39 | 2.8473 | 0.6354 |
| 20003_1140926606 | Treatment/medication code: salbutamol 100micrograms spacehaler | 0.4127 | 7.7342E-05 | 0.2212 | 0.0088 | 1.4265 | 0.7486 |
| 2887 | Number of cigarettes previously smoked daily | 0.4125 | 2.0859E-16 | 0.2822 | 2.7099E-08 | 1.8244 | 0.6758 |
| Z47 | Diagnoses - main ICD10: Z47 Other orthopaedic follow-up care | 0.4116 | 0.001 | 0.2923 | 0.0027 | 0.7531 | 0.6618 |
| 6157_1 | Why stopped smoking: Illness or ill health | 0.4112 | 1.5772E-05 | 0.3654 | 0.0003 | 0.3319 | 0.5637 |
| M13_ARTHROSIS_OTH | Other arthrosis | 0.41 | 0.00002583 | 0.5567 | 1.0466E-09 | -1.0988 | 0.3038 |
| 20003_1140871462 | Treatment/medication code: naproxen | 0.4097 | 0.0206 | 0.4563 | 0.0364 | -0.1659 | 0.4352 |
| 20002_1138 | Non-cancer illness code, self-reported: gastro-oesophageal reflux (gord) / gastric reflux | 0.4093 | 1.4151E-09 | 0.3395 | 5.4008E-10 | 0.8027 | 0.5965 |
| 20518 | Recent changes in speed/amount of moving or speaking | 0.408 | 1.4777E-08 | 0.312 | 6.9012E-06 | 0.96 | 0.6316 |
| 20519 | Recent feelings of tiredness or low energy | 0.4078 | 1.3874E-19 | 0.2646 | 9.0557E-10 | 2.2956 | 0.6919 |
| 2020 | Loneliness, isolation | 0.4061 | 8.6437E-29 | 0.2372 | 8.0454E-10 | 3.1793 | 0.7228 |
| 2100 | Seen a psychiatrist for nerves, anxiety, tension or depression | 0.4056 | 6.2434E-22 | 0.1992 | 5.2861E-09 | 3.8097 | 0.765 |
| 20002_1223 | Non-cancer illness code, self-reported: type 2 diabetes | 0.4056 | 0.0001 | 0.1691 | 0.0181 | 1.8533 | 0.7961 |
| 1920 | Mood swings | 0.4035 | 3.5098E-31 | 0.2354 | 3.1054E-09 | 3.1881 | 0.7218 |
| 22617_8211 | Job SOC coding: Heavy goods vehicle drivers | 0.4019 | 0.0026 | 0.3752 | 0.0005 | 0.1556 | 0.5372 |
| 20401 | Ever addicted to any substance or behaviour | 0.4016 | 3.0867E-05 | 0.2686 | 0.0029 | 1.0085 | 0.6792 |
| E87 | Diagnoses - main ICD10: E87 Other disorders of fluid, electrolyte and acid-base balance | 0.3996 | 0.0169 | 0.2979 | 0.0167 | 0.4877 | 0.639 |
| 6152_7 | Blood clot, DVT, bronchitis, emphysema, asthma, rhinitis, eczema, allergy diagnosed by doctor: Blood clot in the lung | 0.3995 | 9.3795E-05 | 0.5001 | 1.1614E-09 | -0.7666 | 0.3624 |
| 6142_5 | Current employment status: Unemployed | 0.3992 | 0.0002 | 0.1672 | 0.0281 | 1.7463 | 0.7916 |
| 23112 | Leg fat mass (right) | 0.3991 | 3.1772E-54 | 0.3059 | 2.794E-39 | 2.6867 | 0.6279 |
| F5_PERSONALITY | Specific personality disorders | 0.398 | 0.0067 | 0.2525 | 0.0286 | 0.7792 | 0.6947 |
| 21001 | Body mass index (BMI) | 0.3968 | 8.6725E-55 | 0.31 | 1.0833E-40 | 2.5178 | 0.6193 |
| 23104 | Body mass index (BMI) | 0.3957 | 1.3917E-53 | 0.3104 | 9.2281E-40 | 2.4494 | 0.6173 |
| 23116 | Leg fat mass (left) | 0.3954 | 4.566E-53 | 0.307 | 1.1412E-39 | 2.5429 | 0.6215 |
| 20514 | Recent lack of interest or pleasure in doing things | 0.3954 | 1.5471E-10 | 0.2072 | 0.0001 | 2.3024 | 0.7449 |
| 20116_1 | Smoking status: Previous | 0.3949 | 1.3354E-32 | 0.2823 | 9.5226E-25 | 2.6119 | 0.6532 |
| T84 | Diagnoses - main ICD10: T84 Complications of internal orthopaedic prosthetic devices, implants and grafts | 0.3949 | 1.1962E-05 | 0.4079 | 6.8256E-06 | -0.1016 | 0.4819 |
| 2090 | Seen doctor (GP) for nerves, anxiety, tension or depression | 0.3942 | 2.6916E-26 | 0.1716 | 3.5901E-07 | 4.4347 | 0.782 |
| 20002_1093 | Non-cancer illness code, self-reported: pulmonary embolism +/- dvt | 0.3936 | 0.0001 | 0.5164 | 2.3904E-09 | -0.9114 | 0.3337 |
| 20002_1202 | Non-cancer illness code, self-reported: urinary frequency / incontinence | 0.3917 | 0.0093 | 0.293 | 0.0074 | 0.53 | 0.6351 |
| K59 | Diagnoses - main ICD10: K59 Other functional intestinal disorders | 0.3913 | 0.0173 | 0.4714 | 0.0061 | -0.3368 | 0.3896 |

| | | | | | | | |
|---|---|---|---|---|---|---|---|
| 41248_6000 | Destinations on discharge from hospital (recoded): Transfer within NHS provider | 0.3908 | 0.0225 | 0.309 | 0.0233 | 0.3738 | 0.6127 |
| K11_GALLBILPANC | Disorders of gallbladder, biliary tract and pancreas | 0.3902 | 1.7736E-09 | 0.2068 | 0.0004 | 2.1038 | 0.7395 |
| 20459 | General happiness with own health | 0.3883 | 3.1823E-17 | 0.1978 | 2.9484E-06 | 3.0484 | 0.7475 |
| 20002_1406 | Non-cancer illness code, self-reported: muscle or soft tissue injuries | 0.3871 | 0.0309 | 0.5053 | 0.0265 | -0.4078 | 0.3396 |
| 3591 | Ever had hysterectomy (womb removed) | 0.387 | 9.6077E-10 | 0.36 | 1.7738E-09 | 0.3101 | 0.5376 |
| G56 | Diagnoses - main ICD10: G56 Mononeuropathies of upper limb | 0.3865 | 2.9842E-13 | 0.3413 | 2.0906E-14 | 0.6525 | 0.5628 |
| 20160 | Ever smoked | 0.3861 | 1.5606E-35 | 0.2597 | 5.5563E-22 | 3.0796 | 0.6709 |
| 100250 | Instant coffee intake | 0.3859 | 0.0003 | 0.2526 | 0.0089 | 0.9314 | 0.6796 |
| 22506_111 | Tobacco smoking: Smokes on most or all days | 0.3853 | 0.0091 | 0.4675 | 0.0025 | -0.3843 | 0.3868 |
| 48 | Waist circumference | 0.385 | 1.5471E-48 | 0.269 | 8.4928E-35 | 3.3894 | 0.6576 |
| 22605 | Work hours per week - exact value | 0.385 | 0.0039 | 0.5865 | 0.0008 | -0.915 | 0.2403 |
| 20002_1286 | Non-cancer illness code, self-reported: depression | 0.3849 | 8.9971E-12 | 0.1428 | 0.0056 | 3.1699 | 0.8016 |
| 20003_1140884488 | Treatment/medication code: diclofenac | 0.3847 | 8.4731E-07 | 0.3363 | 6.3158E-07 | 0.4689 | 0.5673 |
| 20003_2038460150 | Treatment/medication code: paracetamol | 0.3844 | 2.0063E-23 | 0.317 | 7.064E-15 | 1.203 | 0.5932 |
| 20003_1140921600 | Treatment/medication code: citalopram | 0.3838 | 0.001 | 0.2218 | 0.0228 | 1.0652 | 0.7146 |
| 894 | Duration of moderate activity | 0.3819 | 7.3462E-13 | 0.2094 | 2.7974E-07 | 2.5729 | 0.727 |
| 20126_3 | Bipolar and major depression status: Probable Recurrent major depression (severe) | 0.3796 | 0.0003 | 0.2529 | 0.0016 | 0.9552 | 0.6713 |
| 20513 | Recent thoughts of suicide or self-harm | 0.3794 | 0.0003 | 0.2996 | 0.0013 | 0.5677 | 0.61 |
| 20512 | Recent feelings of foreboding | 0.3792 | 2.2846E-12 | 0.1487 | 0.0052 | 3.0407 | 0.7901 |
| 20528 | Diagnosed with life-threatening illness | 0.3788 | 0.0002 | 0.2824 | 0.0019 | 0.7144 | 0.6321 |
| 20003_1140865716 | Treatment/medication code: senna | 0.3783 | 0.0499 | 0.6834 | 0.0022 | -1.034 | 0.1428 |
| 20086_10 | Type of special diet followed: Low calorie | 0.3779 | 0.0002 | 0.4892 | 5.8094E-06 | -0.7534 | 0.3484 |
| 1548 | Variation in diet | 0.3776 | 5.6389E-26 | 0.2773 | 1.1255E-16 | 2.0486 | 0.6372 |
| Z03 | Diagnoses - main ICD10: Z03 Medical observation and evaluation for suspected diseases and conditions | 0.3767 | 0.0006 | 0.3841 | 0.0014 | -0.0455 | 0.4897 |
| 20546_3 | Substances taken for depression: Medication prescribed to you (for at least two weeks) | 0.3759 | 1.5568E-14 | 0.1019 | 0.0391 | 3.9419 | 0.8312 |
| M19 | Diagnoses - main ICD10: M19 Other arthrosis | 0.3754 | 0.0012 | 0.5687 | 3.2932E-08 | -1.2484 | 0.2494 |
| 20498 | Felt very upset when reminded of stressful experience in past month | 0.3742 | 5.3818E-13 | 0.2565 | 5.0716E-09 | 1.7315 | 0.6598 |
| 22611_1 | Workplace had a lot of cigarette smoke from other people smoking: Sometimes | 0.373 | 0.0005 | 0.2685 | 0.004 | 0.7368 | 0.6427 |
| 22601_82112603 | Job coding: heavy goods vehicle (hgv) driver, lorry or truck driver, tanker driver, haulage driver | 0.373 | 0.0036 | 0.3657 | 0.0006 | 0.044 | 0.5102 |
| 20508 | Recent trouble concentrating on things | 0.3728 | 2.0368E-10 | 0.2212 | 1.2485E-05 | 1.9581 | 0.7021 |
| 20448 | Professional informed about depression | 0.3725 | 1.0352E-06 | 0.212 | 0.0014 | 1.587 | 0.7128 |
| 6154_6 | Medication for pain relief, constipation, heartburn: Laxatives (e.g. Dulcolax, Senokot) | 0.371 | 7.0177E-12 | 0.4177 | 1.9857E-12 | -0.5813 | 0.4351 |

| | | | | | | | |
|---|---|---|---|---|---|---|---|
| 1070 | Time spent watching television (TV) | 0.37 | 1.9114E-31 | 0.2251 | 7.5343E-15 | 3.3726 | 0.694 |
| 20553_4 | Methods of self-harm used: Ingesting a medication in excess of the normal dose | 0.3697 | 0.0007 | 0.2298 | 0.0152 | 0.9699 | 0.6878 |
| K62 | Diagnoses - main ICD10: K62 Other diseases of anus and rectum | 0.3692 | 0.0002 | 0.4209 | 4.3399E-05 | -0.3633 | 0.4282 |
| 20002_1408 | Non-cancer illness code, self-reported: alcohol dependency | 0.369 | 0.0063 | 0.3396 | 0.0186 | 0.1488 | 0.541 |
| 6144_2 | Never eat eggs, dairy, wheat, sugar: Dairy products | 0.3688 | 0.0019 | 0.3523 | 0.0018 | 0.1008 | 0.523 |
| 20544_6 | Mental health problems ever diagnosed by a professional: Panic attacks | 0.3685 | 6.5015E-06 | 0.2781 | 1.8573E-05 | 0.8664 | 0.6241 |
| 6162_100 | Types of transport used (excluding work): None of the above | 0.3677 | 0.0009 | 0.4209 | 3.3265E-05 | -0.3533 | 0.4262 |
| 6152_5 | Blood clot, DVT, bronchitis, emphysema, asthma, rhinitis, eczema, allergy diagnosed by doctor: Blood clot in the leg (DVT) | 0.3673 | 6.5964E-07 | 0.4084 | 3.7564E-09 | -0.4057 | 0.4428 |
| 6143_1 | Transport type for commuting to job workplace: Car/motor vehicle | 0.3672 | 2.2644E-09 | 0.6369 | 1.0069E-34 | -3.3573 | 0.1726 |
| 100370 | Intake of sugar added to coffee | 0.3665 | 8.4568E-05 | 0.2006 | 0.0105 | 1.3622 | 0.7192 |
| 6151_7 | Fractured bone site(s): Other bones | 0.3651 | 4.8607E-07 | 0.2856 | 5.7531E-05 | 0.7829 | 0.6096 |
| N17 | Diagnoses - main ICD10: N17 Acute renal failure | 0.365 | 0.028 | 0.2545 | 0.0469 | 0.5268 | 0.6505 |
| Risk taking | #N/A | 0.3645 | 3.8514E-25 | 0.491 | 1.0664E-58 | -2.7198 | 0.329 |
| 2443 | Diabetes diagnosed by doctor | 0.3635 | 2.1028E-19 | 0.1875 | 3.4636E-09 | 3.4326 | 0.731 |
| 6144_4 | Never eat eggs, dairy, wheat, sugar: Sugar or foods/drinks containing sugar | 0.363 | 6.847E-22 | 0.2733 | 2.4019E-15 | 1.7527 | 0.6232 |
| 20003_1141174520 | Treatment/medication code: symbicort 100/6 turbohaler | 0.363 | 0.0018 | 0.2283 | 0.0181 | 0.8919 | 0.6813 |
| 2644 | Light smokers, at least 100 smokes in lifetime | 0.3627 | 1.3707E-11 | 0.4562 | 6.058E-32 | -1.4113 | 0.3718 |
| M13_ROTATORCUFF | Rotator cuff syndrome | 0.3621 | 2.5902E-05 | 0.3107 | 2.1861E-05 | 0.4548 | 0.5714 |
| S09 | Diagnoses - main ICD10: S09 Other and unspecified injuries of head | 0.3615 | 0.0005 | 0.2922 | 0.0033 | 0.4822 | 0.5958 |
| 6151_1 | Fractured bone site(s): Ankle | 0.3614 | 0.0001 | 0.24 | 0.0038 | 0.9704 | 0.6645 |
| 2473 | Other serious medical condition/disability diagnosed by doctor | 0.361 | 9.9073E-17 | 0.2114 | 1.5972E-07 | 2.5228 | 0.6997 |
| 20548_1 | Manifestations of mania or irritability: I was more talkative than usual | 0.3602 | 1.5063E-05 | 0.1969 | 0.0031 | 1.5332 | 0.7162 |
| 20002_1094 | Non-cancer illness code, self-reported: deep venous thrombosis (dvt) | 0.36 | 3.4154E-07 | 0.4007 | 3.5417E-09 | -0.4155 | 0.4434 |
| 20544_11 | Mental health problems ever diagnosed by a professional: Depression | 0.3599 | 4.7574E-13 | 0.1462 | 0.0025 | 3.0773 | 0.7727 |
| 20003_1140884600 | Treatment/medication code: metformin | 0.3585 | 3.2389E-14 | 0.211 | 1.3137E-08 | 2.4569 | 0.6971 |
| 23120 | Arm fat mass (right) | 0.3584 | 9.42E-44 | 0.2755 | 1.838E-32 | 2.3893 | 0.6141 |
| 4294_0 | Final attempt correct: no | 0.358 | 2.8595E-05 | 0.3877 | 2.4074E-06 | -0.2503 | 0.4586 |
| K80 | Diagnoses - main ICD10: K80 Cholelithiasis | 0.3579 | 1.5452E-08 | 0.1834 | 0.0022 | 2.0023 | 0.7293 |
| R69 | Diagnoses - main ICD10: R69 Unknown and unspecified causes of morbidity | 0.3578 | 0.0024 | 0.3702 | 0.0005 | -0.0781 | 0.4827 |
| 6015 | Chest pain felt during physical activity | 0.3578 | 0.0093 | 0.3797 | 0.0045 | -0.1141 | 0.4695 |
| 23124 | Arm fat mass (left) | 0.3576 | 1.1439E-42 | 0.2797 | 5.63E-33 | 2.2223 | 0.6074 |

| | | | | | | | |
|---|---|---|---|---|---|---|---|
| 1538_2 | Major dietary changes in the last 5 years: Yes, because of other reasons | 0.3567 | 6.1341E-11 | 0.3525 | 1.4303E-15 | 0.0599 | 0.5059 |
| 20507 | Recent feelings of inadequacy | 0.3561 | 2.3443E-08 | 0.1945 | 0.0007 | 1.8801 | 0.7141 |
| 20003_1140868226 | Treatment/medication code: aspirin | 0.3557 | 2.136E-12 | 0.2681 | 5.9423E-08 | 1.2375 | 0.6204 |
| 6159_2 | Pain type(s) experienced in last month: Facial pain | 0.3541 | 0.0003 | 0.2485 | 0.0042 | 0.8118 | 0.6441 |
| 2415 | Had major operations | 0.3535 | 2.1696E-08 | 0.3595 | 1.9294E-11 | -0.0724 | 0.4916 |
| 20548_7 | Manifestations of mania or irritability: I was easily distracted | 0.3533 | 1.021E-06 | 0.2702 | 0.0004 | 0.7878 | 0.6144 |
| 3731 | Former alcohol drinker | 0.353 | 6.3964E-06 | 0.1922 | 0.0069 | 1.5214 | 0.7132 |
| 23100 | Whole body fat mass | 0.3528 | 1.7114E-42 | 0.2769 | 8.1003E-34 | 2.2044 | 0.6047 |
| 20548_6 | Manifestations of mania or irritability: I was more creative or had more ideas than usual | 0.3524 | 0.0043 | 0.2412 | 0.0179 | 0.6952 | 0.6514 |
| 23099 | Body fat percentage | 0.352 | 3.2366E-43 | 0.274 | 1.1889E-32 | 2.2714 | 0.6076 |
| 20497 | Repeated disturbing thoughts of stressful experience in past month | 0.3504 | 1.652E-11 | 0.2129 | 1.1856E-05 | 1.9318 | 0.6848 |
| 20510 | Recent feelings of depression | 0.3503 | 4.3907E-09 | 0.1766 | 0.0005 | 2.2122 | 0.7284 |
| 41248_5001 | Destinations on discharge from hospital (recoded): Transfer to other NHS provider: General ward, young physically disabled, A&E | 0.3497 | 0.0073 | 0.259 | 0.0418 | 0.4979 | 0.6245 |
| 20500 | Ever suffered mental distress preventing usual activities | 0.3494 | 1.464E-11 | 0.1429 | 0.0007 | 3.1031 | 0.7651 |
| 23119 | Arm fat percentage (right) | 0.3489 | 1.3851E-42 | 0.2635 | 1.3551E-30 | 2.4917 | 0.6175 |
| K30 | Diagnoses - main ICD10: K30 Dyspepsia | 0.3488 | 0.0084 | 0.5391 | 1.5839E-05 | -1.0455 | 0.2527 |
| 2704 | Years since last cervical smear test | 0.3471 | 0.0002 | 0.2012 | 0.005 | 1.25 | 0.6952 |
| 20003_1140874744 | Treatment/medication code: gliclazide | 0.345 | 1.914E-06 | 0.2621 | 3.1627E-06 | 0.9037 | 0.6141 |
| 1960 | Fed-up feelings | 0.3449 | 2.0829E-21 | 0.163 | 0.0000741 | 3.3172 | 0.7378 |
| 20003_1140876404 | Treatment/medication code: aqueous cream bp | 0.3448 | 0.0261 | 0.3056 | 0.0227 | 0.1912 | 0.5546 |
| 20002_1267 | Non-cancer illness code, self-reported: spinal injury | 0.3434 | 0.0357 | 0.3657 | 0.0407 | -0.0921 | 0.4689 |
| R32 | Diagnoses - main ICD10: R32 Unspecified urinary incontinence | 0.3429 | 0.0236 | 0.3183 | 0.0073 | 0.1279 | 0.5343 |
| 1448_1 | Bread type: White | 0.3428 | 2.8284E-20 | 0.2429 | 3.6837E-12 | 1.9559 | 0.6367 |
| 6154_1 | Medication for pain relief, constipation, heartburn: Aspirin | 0.3427 | 7.8312E-12 | 0.2709 | 5.4817E-08 | 1.0164 | 0.5992 |
| C_RESPIRATORY_INTRATHORACIC | NA | 0.3426 | 0.0006 | 0.3895 | 0.0006 | -0.3117 | 0.4348 |
| C3_RESPIRATORY_INTRATHORACIC | Malignant neoplasm of respiratory system and intrathoracic organs | 0.3426 | 0.0006 | 0.3895 | 0.0006 | -0.3117 | 0.4348 |
| 20107_3 | Illnesses of father: Lung cancer | 0.342 | 2.1719E-09 | 0.3476 | 1.0318E-11 | -0.0731 | 0.4922 |
| 1930 | Miserableness | 0.3417 | 3.2117E-17 | 0.1167 | 0.0148 | 3.587 | 0.7845 |
| 2654_4 | Non-butter spread type details: Soft (tub) margarine | 0.3417 | 6.4497E-09 | 0.2471 | 9.1072E-07 | 1.2214 | 0.6297 |
| LUNG_CANCER | Lung cancer and mesothelioma | 0.3417 | 0.0007 | 0.3582 | 0.0006 | -0.1134 | 0.477 |
| LUNG_CANCER_MESOT | Lung cancer and mesothelioma | 0.3417 | 0.0007 | 0.3582 | 0.0006 | -0.1134 | 0.477 |
| 1777 | Part of a multiple birth | 0.3414 | 0.0406 | 0.3834 | 0.0385 | -0.1686 | 0.4416 |
| 20003_1140860954 | Treatment/medication code: isosorbide mononitrate | 0.341 | 0.002 | 0.2728 | 0.0072 | 0.4545 | 0.5943 |
| G6_SLEEPAPNO | Sleep apnoea | 0.3409 | 0.0005 | 0.2642 | 0.0029 | 0.5823 | 0.6058 |
| 23123 | Arm fat percentage (left) | 0.3403 | 6.3768E-39 | 0.2663 | 3.1802E-30 | 2.1151 | 0.6022 |

| | | | | | | | |
|---|---|---|---|---|---|---|---|
| 1747_6 | Hair colour (natural, before greying): Other | 0.339 | 0.0093 | 0.3046 | 0.0129 | 0.1922 | 0.5479 |
| 1747_6 | Hair colour (natural, before greying): Other | 0.339 | 0.0093 | 0.3046 | 0.0129 | 0.1922 | 0.5479 |
| 6153_1 | Medication for cholesterol, blood pressure, diabetes, or take exogenous hormones: Cholesterol lowering medication | 0.3377 | 2.8605E-12 | 0.2197 | 1.7869E-08 | 1.9008 | 0.6602 |
| I20 | Diagnoses - main ICD10: I20 Angina pectoris | 0.3366 | 2.4631E-07 | 0.316 | 3.3138E-06 | 0.2187 | 0.5287 |
| 20003_1141146234 | Treatment/medication code: atorvastatin | 0.3354 | 8.0842E-08 | 0.1934 | 0.0003 | 1.7233 | 0.6904 |
| 6153_3 | Medication for cholesterol, blood pressure, diabetes, or take exogenous hormones: Insulin | 0.335 | 0.0006 | 0.1869 | 0.0295 | 1.1384 | 0.6979 |
| 20002_1190 | Non-cancer illness code, self-reported: peritonitis | 0.3344 | 0.0164 | 0.2905 | 0.0186 | 0.2358 | 0.5611 |
| G47 | Diagnoses - main ICD10: G47 Sleep disorders | 0.3338 | 9.8547E-05 | 0.3127 | 9.1931E-05 | 0.1801 | 0.5294 |
| 6154_3 | Medication for pain relief, constipation, heartburn: Paracetamol | 0.3332 | 9.1163E-19 | 0.2768 | 2.0236E-13 | 1.0578 | 0.5782 |
| G6_CARPTU | Carpal tunnel syndrome | 0.3331 | 1.399E-09 | 0.3259 | 2.8295E-13 | 0.1017 | 0.5101 |
| 20447 | Depression possibly related to stressful or traumatic event | 0.3328 | 0.0053 | 0.5146 | 8.8077E-05 | -1.0248 | 0.2623 |
| M48 | Diagnoses - main ICD10: M48 Other spondylopathies | 0.332 | 2.1338E-05 | 0.2147 | 0.0024 | 1.1142 | 0.6593 |
| 20002_1478 | Non-cancer illness code, self-reported: cervical spondylosis | 0.3311 | 0.0134 | 0.3048 | 0.0033 | 0.1554 | 0.5367 |
| 20002_1478 | Non-cancer illness code, self-reported: cervical spondylosis | 0.3311 | 0.0134 | 0.3048 | 0.0033 | 0.1554 | 0.5367 |
| 2149 | Lifetime number of sexual partners | 0.3291 | 5.6024E-20 | 0.3942 | 3.2367E-52 | -1.4679 | 0.4099 |
| SLEEP | Sleep disorders (combined) | 0.3291 | 9.1934E-05 | 0.3105 | 8.2695E-05 | 0.1613 | 0.526 |
| 2306 | Weight change compared with 1 year ago | 0.3277 | 0.0002 | 0.4099 | 1.4215E-10 | -0.7514 | 0.3868 |
| 20002_1220 | Non-cancer illness code, self-reported: diabetes | 0.3256 | 7.5166E-14 | 0.1864 | 6.3603E-08 | 2.5072 | 0.6869 |
| 1588 | Average weekly beer plus cider intake | 0.3244 | 1.0583E-09 | 0.1328 | 0.0032 | 2.7472 | 0.7487 |
| 20455 | Age when last took cannabis | 0.3244 | 0.0077 | 0.4521 | 0.009 | -0.6035 | 0.3275 |
| 20002_1074 | Non-cancer illness code, self-reported: angina | 0.3233 | 4.0044E-10 | 0.313 | 5.5011E-13 | 0.1526 | 0.5144 |
| 23128 | Trunk fat mass | 0.3224 | 1.5426E-35 | 0.254 | 1.4393E-29 | 1.9937 | 0.5946 |
| 2624 | Frequency of heavy DIY in last 4 weeks | 0.3219 | 2.4598E-05 | 0.2111 | 0.0034 | 1.0555 | 0.6509 |
| 6150_2 | Vascular/heart problems diagnosed by doctor: Angina | 0.3208 | 1.2671E-09 | 0.3118 | 2.2203E-12 | 0.1305 | 0.5126 |
| 1130 | Hands-free device/speakerphone use with mobile phone in last 3 month | 0.3204 | 3.5823E-11 | 0.6755 | 3.129E-105 | -6.1782 | 0.107 |
| 6149_2 | Mouth/teeth dental problems: Painful gums | 0.3203 | 9.5419E-05 | 0.2848 | 0.0006 | 0.3048 | 0.5494 |
| 2907 | Ever stopped smoking for 6+ months | 0.3194 | 0.0005 | 0.368 | 2.1209E-05 | -0.3867 | 0.4325 |
| 20441 | Ever had prolonged loss of interest in normal activities | 0.3192 | 6.1979E-11 | 0.1346 | 0.0014 | 2.8613 | 0.7409 |
| 20505 | Recent easy annoyance or irritability | 0.3187 | 3.981E-09 | 0.1915 | 0.0009 | 1.6112 | 0.6719 |
| M23 | Diagnoses - main ICD10: M23 Internal derangement of knee | 0.3172 | 8.7624E-08 | 0.4317 | 1.7786E-12 | -1.3436 | 0.3443 |
| 5463 | Leg pain in calf/calves | 0.3159 | 0.0093 | 0.3378 | 0.0017 | -0.1348 | 0.4695 |
| 20546_4 | Substances taken for depression: Drugs or alcohol (more than once) | 0.3153 | 0.0000171 | 0.1539 | 0.0316 | 1.5751 | 0.7139 |
| 20539 | Frequency of inability to stop worrying during worst period of anxiety | 0.3147 | 0.0008 | 0.2537 | 0.001 | 0.502 | 0.5845 |
| 20524 | Sexual interference by partner or ex-partner without consent as an adult | 0.3141 | 0.0006 | 0.2178 | 0.0015 | 0.8387 | 0.6319 |

| | | | | | | | |
|---|---|---|---|---|---|---|---|
| M13_SPINSTENOSIS | Spinal stenosis | 0.3129 | 6.0021E-05 | 0.2021 | 0.0036 | 1.0606 | 0.6509 |
| 971 | Frequency of walking for pleasure in last 4 weeks | 0.312 | 4.8708E-13 | 0.2783 | 2.3758E-11 | 0.5613 | 0.5469 |
| 20531 | Victim of sexual assault | 0.3117 | 1.4928E-07 | 0.1085 | 0.0439 | 2.5378 | 0.7615 |
| 4803_14 | Tinnitus: Yes, but not now, but have in the past | 0.3116 | 0.0007 | 0.163 | 0.0248 | 1.2654 | 0.6985 |
| PRIM_KNEEARTHROSIS | Primary gonarthrosis, bilateral | 0.3116 | 0.0073 | 0.2016 | 0.0453 | 0.7157 | 0.6499 |
| 23127 | Trunk fat percentage | 0.3115 | 2.0025E-32 | 0.246 | 2.9591E-27 | 1.8853 | 0.5907 |
| F5_DEPRESSIO | Depression | 0.3106 | 0.0049 | 0.2173 | 0.0098 | 0.6723 | 0.628 |
| F32 | Diagnoses - main ICD10: F32 Depressive episode | 0.3094 | 0.0278 | 0.274 | 0.0117 | 0.1993 | 0.5493 |
| 1468_2 | Cereal type: Biscuit cereal (e.g. Weetabix) | 0.3086 | 0.00006504 | 0.3219 | 7.5942E-07 | -0.1316 | 0.4814 |
| 1468_2 | Cereal type: Biscuit cereal (e.g. Weetabix) | 0.3086 | 0.00006504 | 0.3219 | 7.5942E-07 | -0.1316 | 0.4814 |
| 1628 | Alcohol intake versus 10 years previously | 0.308 | 5.8766E-15 | 0.2404 | 1.253E-10 | 1.2427 | 0.5935 |
| 1200 | Sleeplessness / insomnia | 0.3077 | 2.8066E-18 | 0.2121 | 2.8683E-11 | 2.0093 | 0.631 |
| 21002 | Weight | 0.3073 | 7.4002E-35 | 0.236 | 2.3144E-27 | 2.1544 | 0.5985 |
| 23098 | Weight | 0.3068 | 2.1615E-34 | 0.2366 | 5.3972E-27 | 2.1033 | 0.597 |
| 1060 | Time spent outdoors in winter | 0.3064 | 2.489E-13 | 0.3216 | 1.6142E-17 | -0.2694 | 0.4788 |
| 6160_100 | Leisure/social activities: None of the above | 0.3055 | 3.2519E-12 | 0.1083 | 0.0044 | 3.4008 | 0.7549 |
| K22 | Diagnoses - main ICD10: K22 Other diseases of oesophagus | 0.3055 | 0.0001 | 0.2374 | 0.0018 | 0.6163 | 0.5942 |
| Z42 | Diagnoses - main ICD10: Z42 Follow-up care involving plastic surgery | 0.3051 | 0.0058 | 0.2272 | 0.0111 | 0.5475 | 0.6074 |
| V_MENTAL_BEHAV | Mental and behavioural disorders | 0.3048 | 0.0006 | 0.2557 | 0.0005 | 0.4245 | 0.5682 |
| 991 | Frequency of strenuous sports in last 4 weeks | 0.3047 | 0.0135 | 0.2791 | 0.004 | 0.1631 | 0.5357 |
| KRA_PSY_ANYMENTAL | Any mental disorder | 0.3046 | 0.0005 | 0.2345 | 0.0016 | 0.6122 | 0.5969 |
| 20003_1140883468 | Treatment/medication code: clonidine | 0.3038 | 0.0339 | 0.2707 | 0.0426 | 0.1691 | 0.5461 |
| 20501 | Ever had period of mania / excitability | 0.3034 | 0.0005 | 0.2882 | 0.0002 | 0.1313 | 0.5212 |
| 6149_5 | Mouth/teeth dental problems: Toothache | 0.3028 | 6.0216E-05 | 0.2579 | 0.0001 | 0.4442 | 0.5624 |
| 20434 | Age at last episode of depression | 0.3022 | 0.0011 | 0.4436 | 0.00000196 | -1.0745 | 0.3104 |
| 3751 | Chest pain or discomfort when walking uphill or hurrying | 0.3007 | 0.0002 | 0.3175 | 4.8338E-06 | -0.1589 | 0.4766 |
| 6148_1 | Eye problems/disorders: Diabetes related eye disease | 0.2999 | 0.0002 | 0.2869 | 4.3148E-05 | 0.1206 | 0.5181 |
| D12 | Diagnoses - main ICD10: D12 Benign neoplasm of colon, rectum, anus and anal canal | 0.2997 | 5.8956E-07 | 0.1249 | 0.019 | 2.1781 | 0.7296 |
| 20517 | Trouble falling or staying asleep, or sleeping too much | 0.2983 | 1.454E-08 | 0.215 | 4.7648E-06 | 1.1809 | 0.6147 |
| M13_MENISCUSDERANGEMENTS | Meniscus derangement | 0.2975 | 5.6169E-07 | 0.4208 | 4.0605E-11 | -1.4145 | 0.333 |
| 20003_1140871310 | Treatment/medication code: ibuprofen | 0.2956 | 2.1385E-07 | 0.3586 | 4.0383E-11 | -0.8003 | 0.4127 |
| 20003_1140861998 | Treatment/medication code: ventolin 100micrograms inhaler | 0.2934 | 4.2773E-08 | 0.1711 | 0.0002 | 1.7267 | 0.6657 |
| 2405 | Number of children fathered | 0.293 | 2.6278E-08 | 0.4855 | 1.1493E-25 | -2.7416 | 0.2503 |
| J84 | Diagnoses - main ICD10: J84 Other interstitial pulmonary diseases | 0.293 | 0.0225 | 0.3308 | 0.0014 | -0.2294 | 0.4474 |
| M13_SYNOTEND | Disorders of synovium and tendon | 0.2922 | 0.0053 | 0.4106 | 8.1607E-05 | -0.8015 | 0.3393 |
| II_NEOPLASM | Neoplasms | 0.292 | 2.0815E-07 | 0.1842 | 0.0006 | 1.3905 | 0.647 |

| | | | | | | | |
|---|---|---|---|---|---|---|---|
| 1598 | Average weekly spirits intake | 0.2898 | 2.8277E-08 | 0.3192 | 3.7271E-13 | -0.431 | 0.459 |
| 20419 | Difficulty concentrating during worst period of anxiety | 0.2896 | 0.0061 | 0.3341 | 0.0102 | -0.2655 | 0.4381 |
| 20002_1201 | Non-cancer illness code, self-reported: bladder problem (not cancer) | 0.2888 | 0.0172 | 0.2955 | 0.008 | -0.0407 | 0.4906 |
| 22618_0 | Breathing problems improved/stopped away from workplace or on holiday: No | 0.2885 | 0.005 | 0.2563 | 0.0054 | 0.2332 | 0.5449 |
| 20002_1075 | Non-cancer illness code, self-reported: heart attack/myocardial infarction | 0.2883 | 3.1459E-08 | 0.2916 | 2.2475E-11 | -0.0486 | 0.4954 |
| 20003_1140863202 | Treatment/medication code: temazepam | 0.288 | 0.0282 | 0.2647 | 0.0318 | 0.1294 | 0.5325 |
| 20003_1140860840 | Treatment/medication code: nitrolingual 400micrograms spray | 0.2858 | 0.0282 | 0.3486 | 0.0259 | -0.3085 | 0.413 |
| 6150_1 | Vascular/heart problems diagnosed by doctor: Heart attack | 0.2857 | 9.3437E-08 | 0.2859 | 8.4095E-11 | -0.0029 | 0.4997 |
| 22502 | Cough on most days | 0.2843 | 2.0246E-05 | 0.1499 | 0.0232 | 1.4323 | 0.681 |
| 49 | Hip circumference | 0.2841 | 1.2486E-27 | 0.2108 | 6.7553E-20 | 2.103 | 0.6012 |
| 914 | Duration of vigorous activity | 0.2825 | 3.2709E-07 | 0.2525 | 1.7951E-07 | 0.4082 | 0.5418 |
| K85 | Diagnoses - main ICD10: K85 Acute pancreatitis | 0.2817 | 0.0248 | 0.2248 | 0.0202 | 0.359 | 0.5789 |
| C_BRONCHUS_LUNG | Malignant neoplasm of bronchus and lung | 0.281 | 0.0007 | 0.3127 | 0.0007 | -0.2549 | 0.4558 |
| C3_BRONCHUS_LUNG | Malignant neoplasm of bronchus and lung | 0.281 | 0.0007 | 0.3127 | 0.0007 | -0.2549 | 0.4558 |
| I84 | Diagnoses - main ICD10: I84 Haemorrhoids | 0.2807 | 5.9339E-06 | 0.2783 | 1.7942E-07 | 0.0294 | 0.5034 |
| 3849 | Number of pregnancy terminations | 0.2791 | 0.0002 | 0.2584 | 2.4913E-05 | 0.2159 | 0.5289 |
| 20002_1474 | Non-cancer illness code, self-reported: hiatus hernia | 0.2791 | 0.0006 | 0.3248 | 1.5721E-06 | -0.4322 | 0.4365 |
| 20428 | Professional informed about anxiety | 0.2791 | 0.0105 | 0.2995 | 0.0024 | -0.1388 | 0.4715 |
| 20454 | Maximum frequency of taking cannabis | 0.2791 | 0.0124 | 0.3981 | 0.0003 | -0.7587 | 0.3385 |
| 874 | Duration of walks | 0.2784 | 9.2975E-11 | 0.2507 | 2.0543E-12 | 0.4956 | 0.5386 |
| 30530 | Sodium in urine | 0.2765 | 2.6377E-16 | 0.2705 | 4.2577E-16 | 0.1265 | 0.5084 |
| 20003_1141192736 | Treatment/medication code: ezetimibe | 0.2735 | 0.0031 | 0.1883 | 0.0111 | 0.718 | 0.6172 |
| 20515 | Recent trouble relaxing | 0.2724 | 8.0399E-07 | 0.1818 | 0.0004 | 1.1979 | 0.6244 |
| I9_HYPTENS | Hypertension | 0.2715 | 0.0265 | 0.3031 | 0.0117 | -0.1841 | 0.456 |
| 2463 | Fractured/broken bones in last 5 years | 0.27 | 4.2401E-09 | 0.252 | 5.9874E-07 | 0.2635 | 0.5251 |
| 1289 | Cooked vegetable intake | 0.2696 | 2.3166E-10 | 0.0799 | 0.0367 | 3.3158 | 0.7466 |
| 20002_1162 | Non-cancer illness code, self-reported: cholelithiasis/gall stones | 0.2688 | 0.0004 | 0.1711 | 0.0194 | 0.924 | 0.6338 |
| 30280 | Immature reticulocyte fraction | 0.2684 | 1.8412E-11 | 0.1743 | 3.5675E-12 | 1.9963 | 0.629 |
| 1478 | Salt added to food | 0.2669 | 2.2871E-15 | 0.2385 | 1.197E-19 | 0.6644 | 0.5396 |
| 22616_1 | Breathing problems during period of job: Yes | 0.2664 | 0.001 | 0.2426 | 0.0006 | 0.2215 | 0.5332 |
| M13_SOFTTISSUENAS | Other specified/unspecified soft tissue disorders | 0.2642 | 0.0062 | 0.4354 | 8.2478E-06 | -1.2467 | 0.2745 |
| L03 | Diagnoses - main ICD10: L03 Cellulitis | 0.2641 | 0.0028 | 0.2901 | 0.0004 | -0.2154 | 0.4637 |
| K60 | Diagnoses - main ICD10: K60 Fissure and fistula of anal and rectal regions | 0.2627 | 0.0022 | 0.1781 | 0.0074 | 0.7782 | 0.6164 |
| HEARTFAIL | Heart failure | 0.2625 | 0.0009 | 0.1736 | 0.0415 | 0.7665 | 0.6221 |
| I9_HEARTFAIL | Heart failure,strict | 0.2625 | 0.0009 | 0.1736 | 0.0415 | 0.7665 | 0.6221 |

| | | | | | | | |
|---|---|---|---|---|---|---|---|
| I9_HEARTFAIL_NS | Heart failure, not strict | 0.2625 | 0.0009 | 0.1736 | 0.0415 | 0.7665 | 0.6221 |
| 30290 | High light scatter reticulocyte percentage | 0.2603 | 1.1208E-12 | 0.1746 | 2.0697E-13 | 1.963 | 0.6179 |
| 20002_1464 | Non-cancer illness code, self-reported: rheumatoid arthritis | 0.26 | 0.0063 | 0.2762 | 0.0006 | -0.1297 | 0.4774 |
| 20429 | Easily tired during worst period of anxiety | 0.2588 | 0.0034 | 0.1895 | 0.032 | 0.554 | 0.5958 |
| 30300 | High light scatter reticulocyte count | 0.2583 | 4.9366E-12 | 0.1722 | 1.0739E-13 | 1.9563 | 0.6184 |
| 20003_1140860806 | Treatment/medication code: ramipril | 0.2572 | 7.6212E-06 | 0.1198 | 0.0443 | 1.6591 | 0.6847 |
| 2237 | Plays computer games | 0.2565 | 5.436E-15 | 0.1764 | 5.5205E-09 | 1.7938 | 0.6104 |
| I9_IHD | Ischaemic heart disease, wide definition | 0.2551 | 6.4484E-11 | 0.2288 | 9.959E-11 | 0.4993 | 0.5367 |
| 20002_1442 | Non-cancer illness code, self-reported: helicobacter pylori | 0.2538 | 0.0322 | 0.3325 | 0.0064 | -0.4629 | 0.3915 |
| 20403 | Amount of alcohol drunk on a typical drinking day | 0.2531 | 3.0849E-06 | 0.1639 | 0.0002 | 1.274 | 0.6225 |
| 6159_1 | Pain type(s) experienced in last month: Headache | 0.2529 | 1.9221E-09 | 0.2027 | 1.5242E-06 | 0.8422 | 0.5697 |
| 1359 | Poultry intake | 0.2527 | 8.5933E-08 | 0.3434 | 2.5716E-18 | -1.4767 | 0.3755 |
| 6160_2 | Leisure/social activities: Pub or social club | 0.2526 | 3.2105E-08 | 0.2193 | 9.3015E-09 | 0.5591 | 0.5464 |
| 20418 | Impact on normal roles during worst period of anxiety | 0.2511 | 0.0021 | 0.2263 | 0.0044 | 0.218 | 0.5346 |
| 20547_1 | Activities undertaken to treat depression: Talking therapies, such as psychotherapy, counselling, group therapy or CBT | 0.2492 | 9.8316E-06 | 0.1295 | 0.0137 | 1.5521 | 0.6624 |
| 6177_1 | Medication for cholesterol, blood pressure or diabetes: Cholesterol lowering medication | 0.2483 | 5.7473E-08 | 0.1646 | 8.7745E-06 | 1.4216 | 0.6152 |
| M67 | Diagnoses - main ICD10: M67 Other disorders of synovium and tendon | 0.2477 | 0.023 | 0.2455 | 0.029 | 0.014 | 0.5031 |
| M13_TRIGGERFINGER | Trigger finger | 0.2472 | 0.0143 | 0.3169 | 0.0008 | -0.5039 | 0.4036 |
| 6154_2 | Medication for pain relief, constipation, heartburn: Ibuprofen (e.g. Nurofen) | 0.2465 | 5.182E-07 | 0.31 | 3.1415E-10 | -0.9126 | 0.4121 |
| I50 | Diagnoses - main ICD10: I50 Heart failure | 0.2464 | 0.0021 | 0.1909 | 0.0313 | 0.4652 | 0.577 |
| F5_MOOD | Mood [affective] disorders | 0.2464 | 0.0175 | 0.1763 | 0.0184 | 0.5482 | 0.5969 |
| KRA_PSY_MOOD | Mood disorders | 0.2464 | 0.0175 | 0.1763 | 0.0184 | 0.5482 | 0.5969 |
| 20437 | Thoughts of death during worst depression | 0.2463 | 0.0304 | 0.2986 | 0.0021 | -0.3501 | 0.4274 |
| I9_CHD | Major coronary heart disease event | 0.2454 | 1.085E-06 | 0.2361 | 7.6054E-08 | 0.1393 | 0.513 |
| I9_CHD_NOREV | Major coronary heart disease event excluding revascularizations | 0.2454 | 1.085E-06 | 0.2361 | 7.6054E-08 | 0.1393 | 0.513 |
| 20003_1140861958 | Treatment/medication code: simvastatin | 0.2442 | 1.1384E-07 | 0.1496 | 0.0002 | 1.5519 | 0.6297 |
| 4291 | Number of attempts | 0.2442 | 3.4983E-06 | 0.3135 | 5.7207E-13 | -1.0153 | 0.4042 |
| 20002_1154 | Non-cancer illness code, self-reported: irritable bowel syndrome | 0.2441 | 0.0189 | 0.2513 | 0.0092 | -0.0507 | 0.4899 |
| 23126 | Arm predicted mass (left) | 0.2437 | 1.3121E-23 | 0.1851 | 3.3784E-17 | 1.7877 | 0.5812 |
| K11_HERNIA | Hernia | 0.2436 | 6.7681E-06 | 0.2196 | 1.0275E-06 | 0.3411 | 0.5335 |
| 23125 | Arm fat-free mass (left) | 0.2431 | 9.781E-23 | 0.1805 | 2.5695E-16 | 1.8883 | 0.5867 |
| S02 | Diagnoses - main ICD10: S02 Fracture of skull and facial bones | 0.2428 | 0.0494 | 0.2786 | 0.0247 | -0.2044 | 0.4501 |
| 20002_1473 | Non-cancer illness code, self-reported: high cholesterol | 0.2421 | 4.8429E-07 | 0.1332 | 0.0001 | 1.8361 | 0.6484 |
| M65 | Diagnoses - main ICD10: M65 Synovitis and tenosynovitis | 0.2389 | 0.02 | 0.311 | 0.001 | -0.5161 | 0.4004 |
| J47 | Diagnoses - main ICD10: J47 Bronchiectasis | 0.2389 | 0.0475 | 0.2726 | 0.027 | -0.1955 | 0.4531 |

| | | | | | | | |
|---|---|---|---|---|---|---|---|
| PULM_MEDICATIO_COMORB | Medication related adverse effects (Asthma/COPD) | 0.2365 | 0.0002 | 0.1223 | 0.0184 | 1.3859 | 0.6553 |
| 20002_1294 | Non-cancer illness code, self-reported: back problem | 0.2364 | 0.0108 | 0.221 | 0.0098 | 0.122 | 0.5215 |
| 1120 | Weekly usage of mobile phone in last 3 months | 0.2359 | 2.3035E-09 | 0.6604 | 2.143E-147 | -9.0288 | 0.0687 |
| 1468_5 | Cereal type: Other (e.g. Cornflakes, Frosties) | 0.2357 | 1.5556E-07 | 0.3346 | 5.6651E-17 | -1.6447 | 0.3646 |
| 1468_5 | Cereal type: Other (e.g. Cornflakes, Frosties) | 0.2357 | 1.5556E-07 | 0.3346 | 5.6651E-17 | -1.6447 | 0.3646 |
| 22704 | Home location - north co-ordinate (rounded) | 0.2354 | 4.5542E-05 | 0.1628 | 0.002 | 0.929 | 0.6003 |
| 2247_1 | Hearing difficulty/problems: Yes | 0.2346 | 1.1608E-09 | 0.076 | 0.03 | 3.0438 | 0.7106 |
| 6152_8 | Blood clot, DVT, bronchitis, emphysema, asthma, rhinitis, eczema, allergy diagnosed by doctor: Asthma | 0.2345 | 9.9384E-10 | 0.099 | 0.0013 | 2.7491 | 0.6823 |
| R55 | Diagnoses - main ICD10: R55 Syncope and collapse | 0.2339 | 0.0024 | 0.1782 | 0.0157 | 0.5215 | 0.5773 |
| 20110_1 | Illnesses of mother: Heart disease | 0.2328 | 0.0005 | 0.1361 | 0.0109 | 1.1287 | 0.6325 |
| 134 | Number of self-reported cancers | 0.2323 | 0.0088 | 0.2072 | 0.0034 | 0.2212 | 0.535 |
| 20075 | Home location at assessment - north co-ordinate (rounded) | 0.2316 | 0.0002 | 0.1716 | 0.0015 | 0.7278 | 0.5832 |
| 21021 | Pulse wave Arterial Stiffness index | 0.2314 | 0.0001 | 0.1273 | 0.0169 | 1.2995 | 0.6422 |
| 20002_1111 | Non-cancer illness code, self-reported: asthma | 0.2302 | 1.4217E-09 | 0.1019 | 0.0009 | 2.6229 | 0.6733 |
| 30240 | Reticulocyte percentage | 0.2295 | 3.2924E-11 | 0.1596 | 2.8956E-11 | 1.66 | 0.5966 |
| 30250 | Reticulocyte count | 0.2261 | 2.0609E-10 | 0.1557 | 1.8376E-11 | 1.6568 | 0.5973 |
| 3476 | Difficulty not smoking for 1 day | 0.2248 | 0.0017 | 0.1894 | 0.0052 | 0.3587 | 0.5493 |
| 2453 | Cancer diagnosed by doctor | 0.2245 | 0.0122 | 0.2003 | 0.0081 | 0.2064 | 0.5337 |
| 6155_2 | Vitamin and mineral supplements: Vitamin B | 0.2237 | 0.003 | 0.2955 | 0.00001595 | -0.7048 | 0.4008 |
| 23122 | Arm predicted mass (right) | 0.2235 | 6.3613E-20 | 0.1784 | 1.3359E-16 | 1.3808 | 0.5627 |
| 23121 | Arm fat-free mass (right) | 0.2227 | 2.0326E-19 | 0.1776 | 2.0734E-16 | 1.3745 | 0.5627 |
| 20107_9 | Illnesses of father: Diabetes | 0.2226 | 0.00003115 | 0.1484 | 0.0028 | 1.0171 | 0.6024 |
| ASTHMA_OPPORTUNIST_INFECTIONS | NA | 0.2219 | 0.0479 | 0.2308 | 0.0499 | -0.0548 | 0.4876 |
| 20111_8 | Illnesses of siblings: High blood pressure | 0.2191 | 3.8716E-07 | 0.1668 | 6.0725E-05 | 0.8721 | 0.5726 |
| VII_EYE_ADNEXA | Diseases of the eye and adnexa | 0.2176 | 0.0004 | 0.1259 | 0.0178 | 1.1265 | 0.6259 |
| 23105 | Basal metabolic rate | 0.2173 | 1.4443E-18 | 0.1655 | 3.6822E-14 | 1.5692 | 0.5719 |
| 129 | Place of birth in UK - north co-ordinate | 0.2173 | 0.0002 | 0.1674 | 0.0019 | 0.6238 | 0.5693 |
| 4440 | Average monthly spirits intake | 0.2171 | 0.0138 | 0.338 | 4.5595E-05 | -0.9994 | 0.3361 |
| I21 | Diagnoses - main ICD10: I21 Acute myocardial infarction | 0.2162 | 7.8334E-05 | 0.2087 | 0.00002178 | 0.1019 | 0.5105 |
| I9_MI_STRICT | Myocardial infarction, strict | 0.2142 | 7.7936E-05 | 0.1918 | 6.8434E-05 | 0.3088 | 0.5312 |
| K57 | Diagnoses - main ICD10: K57 Diverticular disease of intestine | 0.2124 | 1.5567E-06 | 0.1972 | 1.737E-06 | 0.2516 | 0.5212 |
| 2634 | Duration of heavy DIY | 0.2106 | 0.0002 | 0.2893 | 9.4966E-08 | -1.0061 | 0.3915 |
| 20003_1140879802 | Treatment/medication code: amlodipine | 0.2105 | 5.5726E-05 | 0.1143 | 0.0117 | 1.3919 | 0.6318 |
| 20003_1140888266 | Treatment/medication code: warfarin | 0.2064 | 0.0262 | 0.2051 | 0.0166 | 0.0103 | 0.5018 |
| 20003_1141176832 | Treatment/medication code: seretide 50 evohaler | 0.2063 | 0.002 | 0.1926 | 0.0016 | 0.1517 | 0.5191 |
| 5208 | logMAR, final (left) | 0.2055 | 0.0087 | 0.1611 | 0.0385 | 0.402 | 0.5617 |
| 3393 | Hearing aid user | 0.2054 | 0.0027 | 0.1933 | 0.0009 | 0.1348 | 0.5169 |

| | | | | | | | |
|---|---|---|---|---|---|---|---|
| 767 | Length of working week for main job | 0.2052 | 0.0008 | 0.4886 | 1.1603E-20 | -3.5175 | 0.1606 |
| 23117 | Leg fat-free mass (left) | 0.2 | 3.2216E-15 | 0.1569 | 5.0136E-12 | 1.2652 | 0.5599 |
| 23118 | Leg predicted mass (left) | 0.2 | 3.4276E-15 | 0.1556 | 8.0842E-12 | 1.3008 | 0.5617 |
| 2724 | Had menopause | 0.1999 | 3.0974E-05 | 0.0968 | 0.0456 | 1.5125 | 0.6409 |
| 4674 | Private healthcare | 0.1994 | 0.0059 | -0.1539 | 0.0193 | 3.6112 | 0.8919 |
| 4290 | Duration screen displayed | 0.199 | 2.6858E-06 | 0.1898 | 6.2167E-08 | 0.1671 | 0.5128 |
| 6150_4 | Vascular/heart problems diagnosed by doctor: High blood pressure | 0.1982 | 1.175E-09 | 0.0928 | 0.0026 | 2.3501 | 0.6439 |
| 670_2 | Type of accommodation lived in: A flat, maisonette or apartment | 0.197 | 0.0009 | 0.0909 | 0.0483 | 1.4108 | 0.6448 |
| 2030 | Guilty feelings | 0.1965 | 2.5789E-06 | 0.1133 | 0.0058 | 1.4193 | 0.6145 |
| 1021 | Duration of light DIY | 0.1954 | 0.0001 | 0.109 | 0.0051 | 1.3404 | 0.6188 |
| COX_ARTHROSIS | Coxarthrosis [arthrosis of hip](FG) | 0.1949 | 0.0008 | 0.177 | 0.0007 | 0.2292 | 0.525 |
| M16 | Diagnoses - main ICD10: M16 Coxarthrosis [arthrosis of hip] | 0.1949 | 0.0008 | 0.177 | 0.0007 | 0.2292 | 0.525 |
| 20416 | Frequency of consuming six or more units of alcohol | 0.1927 | 4.4163E-05 | 0.1841 | 6.7243E-05 | 0.1302 | 0.512 |
| I9_UAP | Unstable angina pectoris | 0.1924 | 0.0101 | 0.2829 | 0.0002 | -0.847 | 0.3757 |
| 22599 | Number of jobs held | 0.1918 | 0.0056 | 0.2328 | 0.001 | -0.4144 | 0.443 |
| 30000 | White blood cell (leukocyte) count | 0.1908 | 7.2623E-11 | 0.1368 | 7.4527E-09 | 1.4329 | 0.5749 |
| 20002_1065 | Non-cancer illness code, self-reported: hypertension | 0.1893 | 2.7336E-09 | 0.0996 | 0.0012 | 2.0294 | 0.6232 |
| 30510 | Creatinine (enzymatic) in urine | 0.1871 | 9.8535E-08 | 0.1843 | 4.0744E-09 | 0.0595 | 0.5039 |
| I9_CORATHER | Coronary atherosclerosis | 0.1869 | 4.4046E-06 | 0.2063 | 1.6766E-07 | -0.3425 | 0.4729 |
| 1508_2 | Coffee type: Instant coffee | 0.1862 | 7.3414E-05 | 0.1102 | 0.0071 | 1.2185 | 0.6049 |
| 1508_2 | Coffee type: Instant coffee | 0.1862 | 7.3414E-05 | 0.1102 | 0.0071 | 1.2185 | 0.6049 |
| 1950 | Sensitivity / hurt feelings | 0.1844 | 2.0146E-06 | 0.0857 | 0.0398 | 1.7328 | 0.6351 |
| 3786 | Age asthma diagnosed | 0.1835 | 0.0046 | 0.1871 | 0.0019 | -0.0407 | 0.495 |
| 6177_2 | Medication for cholesterol, blood pressure or diabetes: Blood pressure medication | 0.1834 | 1.1281E-05 | 0.1358 | 0.0002 | 0.8537 | 0.5662 |
| 6153_2 | Medication for cholesterol, blood pressure, diabetes, or take exogenous hormones: Blood pressure medication | 0.1826 | 5.9574E-06 | 0.0936 | 0.016 | 1.589 | 0.6223 |
| 23113 | Leg fat-free mass (right) | 0.1825 | 7.4059E-13 | 0.1456 | 1.2655E-10 | 1.0853 | 0.5514 |
| 23114 | Leg predicted mass (right) | 0.1823 | 8.1167E-13 | 0.146 | 1.2737E-10 | 1.0633 | 0.5505 |
| 2217 | Age started wearing glasses or contact lenses | 0.1822 | 3.892E-07 | 0.1842 | 2.8676E-08 | -0.0409 | 0.4972 |
| 23102 | Whole body water mass | 0.1819 | 2.0072E-13 | 0.1383 | 2.0694E-10 | 1.3204 | 0.5606 |
| 23101 | Whole body fat-free mass | 0.1804 | 3.6191E-13 | 0.1381 | 2.3345E-10 | 1.2811 | 0.5588 |
| 41248_1001 | Destinations on discharge from hospital (recoded): Usual Place of residence: Living with relatives | 0.18 | 0.0353 | 0.2019 | 0.0219 | -0.1784 | 0.4695 |
| H26 | Diagnoses - main ICD10: H26 Other cataract | 0.1794 | 0.0115 | 0.1259 | 0.0089 | 0.6238 | 0.5743 |
| 20003_1141194794 | Treatment/medication code: bendroflumethiazide | 0.1793 | 0.0001 | 0.1218 | 0.0109 | 0.8569 | 0.5797 |
| I25 | Diagnoses - main ICD10: I25 Chronic ischaemic heart disease | 0.1792 | 1.7331E-05 | 0.2071 | 3.5478E-07 | -0.4788 | 0.4611 |
| 30500 | Microalbumin in urine | 0.1792 | 0.0141 | 0.1533 | 0.0483 | 0.2431 | 0.5361 |
| I9_DISVEINLYMPH | Diseases of veins, lymphatic vessels and lymph nodes, not elsewhere classified | 0.1786 | 0.0009 | 0.1818 | 0.0002 | -0.0438 | 0.4955 |

| | | | | | | | |
|---|---|---|---|---|---|---|---|
| 5201 | logMAR, final (right) | 0.1776 | 0.0361 | 0.2983 | 0.0003 | -1.0214 | 0.3364 |
| 20003_1141145660 | Treatment/medication code: valsartan | 0.1739 | 0.0475 | 0.3163 | 0.0001 | -1.1807 | 0.3091 |
| M20 | Diagnoses - main ICD10: M20 Acquired deformities of fingers and toes | 0.1737 | 0.0018 | 0.1696 | 0.002 | 0.0524 | 0.5057 |
| 4233 | Mean signal-to-noise ratio (SNR), (left) | 0.1722 | 0.049 | 0.1908 | 0.0083 | -0.164 | 0.474 |
| 20003_1140916356 | Treatment/medication code: losartan | 0.1703 | 0.0175 | 0.2001 | 0.0062 | -0.291 | 0.4585 |
| 1389 | Pork intake | 0.1691 | 0.0005 | 0.1504 | 7.2397E-05 | 0.3038 | 0.5261 |
| 904 | Number of days/week of vigorous physical activity 10+ minutes | 0.1664 | 4.3761E-05 | 0.0819 | 0.0211 | 1.5646 | 0.6163 |
| 30120 | Lymphocyte count | 0.1643 | 4.9378E-07 | 0.1365 | 1.509E-09 | 0.6994 | 0.5388 |
| 30140 | Neutrophill count | 0.1598 | 4.2172E-08 | 0.1064 | 1.9415E-05 | 1.3915 | 0.5741 |
| 5084 | Spherical power (right) | 0.159 | 7.8069E-06 | 0.1731 | 1.9908E-10 | -0.3147 | 0.4803 |
| 23130 | Trunk predicted mass | 0.1547 | 5.2125E-10 | 0.1118 | 1.8496E-07 | 1.304 | 0.5597 |
| 23129 | Trunk fat-free mass | 0.1547 | 5.4383E-10 | 0.1124 | 1.4882E-07 | 1.2884 | 0.5588 |
| 3143 | Ankle spacing width | 0.1536 | 2.9946E-08 | 0.182 | 1.0595E-14 | -0.7818 | 0.4604 |
| 20003_1140875408 | Treatment/medication code: allopurinol | 0.1522 | 0.0034 | 0.1701 | 0.0002 | -0.2598 | 0.475 |
| 5085 | Spherical power (left) | 0.151 | 1.7261E-05 | 0.1711 | 1.0003E-10 | -0.457 | 0.472 |
| 4100 | Ankle spacing width (left) | 0.1508 | 2.5733E-07 | 0.157 | 3.7187E-09 | -0.1567 | 0.4913 |
| 4119 | Ankle spacing width (right) | 0.1465 | 1.9513E-06 | 0.1626 | 9.9555E-10 | -0.3956 | 0.4775 |
| 2345 | Ever had bowel cancer screening | 0.1444 | 0.0245 | 0.1476 | 0.0064 | -0.0381 | 0.4955 |
| I9_VTE | Venous thromboembolism | 0.1431 | 0.0232 | 0.1277 | 0.009 | 0.1931 | 0.5215 |
| 3761 | Age hay fever, rhinitis or eczema diagnosed | 0.1404 | 0.0211 | 0.2165 | 1.0672E-06 | -1.0097 | 0.395 |
| 20002_1466 | Non-cancer illness code, self-reported: gout | 0.1351 | 0.0086 | 0.1108 | 0.0102 | 0.3623 | 0.5339 |
| I9_DVTANDPULM | DVT of lower extremities and pulmonary embolism | 0.1349 | 0.0394 | 0.1282 | 0.0103 | 0.0814 | 0.5094 |
| 1418_1 | Milk type used: Full cream | 0.1262 | 0.0323 | -0.1096 | 0.0429 | 2.9484 | 0.7954 |
| 1418_1 | Milk type used: Full cream | 0.1262 | 0.0323 | -0.1096 | 0.0429 | 2.9484 | 0.7954 |
| Time spent driving | #N/A | 0.1248 | 0.0044 | 0.5442 | 2.6405E-59 | -7.6058 | 0.0711 |
| 1220 | Daytime dozing / sleeping (narcolepsy) | 0.1206 | 0.0012 | 0.0742 | 0.0256 | 0.9306 | 0.5645 |
| 30130 | Monocyte count | 0.1183 | 5.3027E-05 | 0.0615 | 0.0236 | 1.4207 | 0.5788 |
| I83 | Diagnoses - main ICD10: I83 Varicose veins of lower extremities | 0.1175 | 0.0256 | 0.1578 | 0.0014 | -0.5579 | 0.4439 |
| 6149_3 | Mouth/teeth dental problems: Bleeding gums | 0.1124 | 0.0194 | 0.1599 | 0.0012 | -0.6896 | 0.434 |
| 30520 | Potassium in urine | 0.1039 | 0.0101 | 0.0774 | 0.0261 | 0.497 | 0.5369 |
| 5256 | Corneal hysteresis (right) | 0.1004 | 0.0183 | 0.08 | 0.0122 | 0.3833 | 0.5285 |
| 1747_1 | Hair colour (natural, before greying): Blonde | 0.0981 | 0.0007 | 0.0684 | 0.0129 | 0.7445 | 0.5414 |
| 1747_1 | Hair colour (natural, before greying): Blonde | 0.0981 | 0.0007 | 0.0684 | 0.0129 | 0.7445 | 0.5414 |
| 3144 | Heel Broadband ultrasound attenuation, direct entry | 0.097 | 0.0009 | 0.0771 | 0.0367 | 0.4223 | 0.5278 |
| 1369 | Beef intake | 0.0923 | 0.0347 | 0.1667 | 1.1108E-07 | -1.3826 | 0.3973 |
| 5096 | 3mm weak meridian (left) | 0.0901 | 0.0069 | 0.0734 | 0.0345 | 0.3472 | 0.5233 |
| 5099 | 3mm weak meridian (right) | 0.0825 | 0.0125 | 0.0698 | 0.0394 | 0.2684 | 0.5177 |
| 5133 | 6mm strong meridian (right) | 0.0749 | 0.0285 | 0.0683 | 0.0496 | 0.1353 | 0.5092 |
| 30070 | Red blood cell (erythrocyte) distribution width | 0.0735 | 0.0227 | 0.0601 | 0.0467 | 0.3035 | 0.5187 |

| | | | | | | | |
|---|---|---|---|---|---|---|---|
| 6152_9 | Blood clot, DVT, bronchitis, emphysema, asthma, rhinitis, eczema, allergy diagnosed by doctor: Hayfever, allergic rhinitis or eczema | -0.07 | 0.047 | -0.0773 | 0.0028 | 0.167 | 0.5102 |
| 1747_4 | Hair colour (natural, before greying): Dark brown | -0.0782 | 0.0005 | -0.1041 | 0.0046 | 0.5997 | 0.5361 |
| 1747_4 | Hair colour (natural, before greying): Dark brown | -0.0782 | 0.0005 | -0.1041 | 0.0046 | 0.5997 | 0.5361 |
| 1737 | Childhood sunburn occasions | -0.0857 | 0.0061 | -0.2307 | 3.1237E-12 | 3.1877 | 0.6941 |
| 3064 | Peak expiratory flow (PEF) | -0.0911 | 0.0104 | -0.0842 | 0.0105 | -0.1423 | 0.4904 |
| 5254 | Intra-ocular pressure, corneal-compensated (right) | -0.0933 | 0.0445 | -0.1397 | 8.6262E-05 | 0.7934 | 0.5645 |
| 2744 | Birth weight of first child | -0.0952 | 0.0101 | -0.0741 | 0.023 | -0.4279 | 0.4706 |
| 6155_100 | Vitamin and mineral supplements: None of the above | -0.0967 | 0.0265 | -0.1467 | 0.0003 | 0.8421 | 0.5695 |
| 50 | Standing height | -0.0983 | 4.4916E-05 | -0.0768 | 0.0001 | -0.6851 | 0.47 |
| 1160 | Sleep duration | -0.1027 | 0.0099 | -0.173 | 3.014E-07 | 1.3463 | 0.5972 |
| 2395_3 | Hair/balding pattern: Pattern 3 | -0.111 | 0.0144 | -0.0889 | 0.0301 | -0.3613 | 0.4692 |
| 20002_1387 | Non-cancer illness code, self-reported: hayfever/allergic rhinitis | -0.1131 | 0.0305 | -0.1114 | 0.006 | -0.0257 | 0.4976 |
| 6162_1 | Types of transport used (excluding work): Car/motor vehicle | -0.1205 | 0.0081 | 0.2746 | 1.0472E-11 | -6.4933 | 0.0834 |
| 1150_1 | Usual side of head for mobile phone use: Left | -0.1266 | 0.0248 | -0.1724 | 0.0072 | 0.5364 | 0.5637 |
| 1150_1 | Usual side of head for mobile phone use: Left | -0.1266 | 0.0248 | -0.1724 | 0.0072 | 0.5364 | 0.5637 |
| 1438 | Bread intake | -0.1272 | 0.0003 | -0.2756 | 8.5827E-20 | 3.2056 | 0.6982 |
| 1438 | Bread intake | -0.1272 | 0.0003 | -0.2756 | 8.5827E-20 | 3.2056 | 0.6982 |
| 20153 | Forced expiratory volume in 1-second (FEV1), predicted | -0.1276 | 1.7299E-05 | -0.0781 | 0.0034 | -1.2394 | 0.4312 |
| 5262 | Intra-ocular pressure, corneal-compensated (left) | -0.1345 | 0.0009 | -0.1302 | 0.0004 | -0.0786 | 0.494 |
| 2664_3 | Reason for reducing amount of alcohol drunk: Health precaution | -0.1473 | 0.0164 | -0.2512 | 6.0155E-08 | 1.35 | 0.6419 |
| 3062 | Forced vital capacity (FVC) | -0.1513 | 2.2297E-07 | -0.1408 | 2.323E-08 | -0.2722 | 0.4853 |
| 3063 | Forced expiratory volume in 1-second (FEV1) | -0.1578 | 1.7567E-07 | -0.1358 | 6.5635E-07 | -0.5404 | 0.4693 |
| 100270 | Filtered coffee intake | -0.1591 | 0.0363 | -0.214 | 0.0019 | 0.5355 | 0.5762 |
| 728 | Number of vehicles in household | -0.1596 | 0.0002 | 0.2716 | 4.5664E-14 | -7.6785 | 0.0656 |
| 1140 | Difference in mobile phone use compared to two years previously | -0.1628 | 0.0118 | 0.2587 | 6.602E-06 | -4.8775 | 0.0701 |
| 5983 | ECG, heart rate | -0.1633 | 0.006 | -0.1801 | 0.0002 | 0.2198 | 0.5234 |
| 699 | Length of time at current address | -0.1636 | 0.0013 | -0.3238 | 3.1491E-14 | 2.4192 | 0.7125 |
| 23108 | Impedance of leg (left) | -0.1652 | 5.9782E-10 | -0.139 | 3.9272E-08 | -0.7123 | 0.4635 |
| 23107 | Impedance of leg (right) | -0.1675 | 3.3284E-10 | -0.1466 | 4.3496E-09 | -0.5714 | 0.4708 |
| 4260 | Round of numeric memory test | -0.1731 | 0.0167 | -0.1749 | 0.0022 | 0.0196 | 0.5025 |
| 20150 | Forced expiratory volume in 1-second (FEV1), Best measure | -0.1748 | 2.7893E-08 | -0.1347 | 1.0768E-06 | -0.9575 | 0.4442 |
| 20151 | Forced vital capacity (FVC), Best measure | -0.1749 | 6.6933E-09 | -0.1435 | 1.4876E-08 | -0.797 | 0.4562 |
| 100008 | Total sugars | -0.1758 | 0.0307 | -0.1884 | 0.0013 | 0.1256 | 0.5176 |
| 20154 | Forced expiratory volume in 1-second (FEV1), predicted percentage | -0.1773 | 6.9759E-07 | -0.1143 | 0.0005 | -1.2941 | 0.4127 |
| 4283 | Number of rounds of numeric memory test performed | -0.1781 | 0.014 | -0.1771 | 0.0019 | -0.0109 | 0.4986 |
| 1210 | Snoring | -0.1792 | 8.8567E-07 | -0.1505 | 5.8141E-07 | -0.6076 | 0.46 |
| 2207 | Wears glasses or contact lenses | -0.1798 | 0.0009 | -0.1527 | 0.0012 | -0.3782 | 0.4622 |

| | | | | | | | |
|---|---|---|---|---|---|---|---|
| 1309 | Fresh fruit intake | -0.1819 | 8.533E-08 | -0.1097 | 0.0008 | -1.5305 | 0.4003 |
| 4282 | Maximum digits remembered correctly | -0.1877 | 0.0027 | -0.1993 | 0.0003 | 0.1388 | 0.5162 |
| 796 | Distance between home and job workplace | -0.198 | 0.0283 | 0.2546 | 0.0018 | -3.7208 | 0.0566 |
| 20128 | Number of fluid intelligence questions attempted within time limit | -0.1985 | 4.4621E-06 | -0.1391 | 0.0006 | -1.0019 | 0.4177 |
| 398 | Number of correct matches in round | -0.2005 | 0.0007 | -0.1011 | 0.0423 | -1.2824 | 0.364 |
| 100016 | Potassium | -0.205 | 0.0135 | -0.2234 | 0.0006 | 0.1748 | 0.5257 |
| 943 | Frequency of stair climbing in last 4 weeks | -0.2057 | 1.6014E-06 | -0.1369 | 7.2419E-05 | -1.2497 | 0.4049 |
| 1508_3 | Coffee type: Ground coffee (include espresso, filter etc) | -0.2116 | 7.1292E-09 | -0.1982 | 6.2953E-10 | -0.2753 | 0.4813 |
| 1508_3 | Coffee type: Ground coffee (include espresso, filter etc) | -0.2116 | 7.1292E-09 | -0.1982 | 6.2953E-10 | -0.2753 | 0.4813 |
| 3581 | Age at menopause (last menstrual period) | -0.2151 | 7.5448E-09 | -0.1658 | 1.4076E-05 | -0.9246 | 0.4315 |
| 22601_23143401 | Job coding: secondary school teacher or teaching professional (including head teacher) | -0.2191 | 0.0099 | -0.1564 | 0.0432 | -0.5461 | 0.4132 |
| 6147_4 | Reason for glasses/contact lenses: For 'astigmatism' | -0.22 | 0.0058 | -0.2416 | 0.0001 | 0.2124 | 0.5301 |
| 102800 | Cheese consumers | -0.2267 | 0.0199 | -0.284 | 0.0045 | 0.4107 | 0.5795 |
| 6034 | Target heart rate achieved | -0.2306 | 0.0069 | -0.325 | 0.0000651 | 0.8001 | 0.6294 |
| 1677 | Breastfed as a baby | -0.2308 | 1.2366E-05 | -0.1116 | 0.0225 | -1.6563 | 0.3383 |
| 6150_100 | Vascular/heart problems diagnosed by doctor: None of the above | -0.2316 | 2.7768E-13 | -0.1211 | 5.3486E-05 | -2.5318 | 0.3495 |
| 2247_0 | Hearing difficulty/problems: No | -0.2355 | 1.1438E-09 | -0.078 | 0.0263 | -3.0146 | 0.2907 |
| 1031 | Frequency of friend/family visits | -0.2381 | 7.581E-10 | -0.4317 | 1.292E-41 | 3.8602 | 0.751 |
| 6162_3 | Types of transport used (excluding work): Public transport | -0.2392 | 2.3245E-06 | -0.2921 | 3.0005E-13 | 0.8191 | 0.5734 |
| 100024 | Calcium | -0.24 | 0.0012 | -0.292 | 7.4069E-05 | 0.4969 | 0.5722 |
| 6142_2 | Current employment status: Retired | -0.2417 | 0.0014 | -0.4051 | 5.772E-11 | 1.6697 | 0.7163 |
| 100004 | Fat | -0.2443 | 0.0075 | -0.3193 | 0.0003 | 0.5895 | 0.6035 |
| 670_1 | Type of accommodation lived in: A house or bungalow | -0.248 | 1.6482E-05 | -0.1087 | 0.0147 | -1.9122 | 0.313 |
| 6020_33 | Completion status of test: Heart rate reached safety level | -0.25 | 0.0154 | -0.2719 | 0.0089 | 0.1495 | 0.5305 |
| 1319 | Dried fruit intake | -0.2514 | 3.3594E-13 | -0.2324 | 5.8241E-14 | -0.4102 | 0.4735 |
| 23106 | Impedance of whole body | -0.2537 | 1.6453E-21 | -0.1953 | 1.5334E-16 | -1.6392 | 0.419 |
| 20074 | Home location at assessment - east co-ordinate (rounded) | -0.2557 | 0.0004 | -0.2403 | 0.0002 | -0.1585 | 0.4785 |
| 6157_3 | Why stopped smoking: Health precaution | -0.2561 | 1.7563E-06 | -0.267 | 6.1412E-08 | 0.1497 | 0.5152 |
| 757 | Time employed in main current job | -0.2571 | 0.0022 | -0.284 | 6.1594E-05 | 0.2451 | 0.5375 |
| 4979 | FI5 : family relationship calculation | -0.2583 | 7.7349E-06 | -0.1951 | 0.0004 | -0.7949 | 0.4125 |
| 6177_100 | Medication for cholesterol, blood pressure or diabetes: None of the above | -0.2592 | 3.7862E-10 | -0.1547 | 2.0171E-05 | -1.8979 | 0.3573 |
| 6143_2 | Transport type for commuting to job workplace: Walk | -0.2602 | 0.0002 | -0.436 | 1.9181E-10 | 1.7884 | 0.7308 |
| 4196 | Pulse wave peak to peak time | -0.2609 | 1.2046E-05 | -0.1654 | 0.0022 | -1.1874 | 0.3691 |
| 981 | Duration walking for pleasure | -0.2638 | 3.8764E-07 | -0.3427 | 8.7634E-14 | 1.1365 | 0.6088 |
| 100002 | Energy | -0.2707 | 0.0007 | -0.3353 | 3.947E-06 | 0.5976 | 0.5894 |
| 100460 | Added milk to standard tea | -0.2751 | 0.0015 | -0.1976 | 0.0155 | -0.6517 | 0.3931 |
| 6162_4 | Types of transport used (excluding work): Cycle | -0.2756 | 2.4272E-09 | -0.2747 | 9.767E-13 | -0.015 | 0.4987 |

| | | | | | | | |
|---|---|---|---|---|---|---|---|
| 23109 | Impedance of arm (right) | -0.2793 | 3.5968E-24 | -0.2334 | 1.0733E-23 | -1.2735 | 0.4362 |
| 22702 | Home location - east co-ordinate (rounded) | -0.2822 | 0.0001 | -0.2256 | 0.0002 | -0.5877 | 0.4215 |
| 6139_3 | Gas or solid-fuel cooking/heating: An open solid fuel fire that you use regularly in winter time | -0.2855 | 3.4334E-08 | -0.104 | 0.0239 | -2.6228 | 0.2627 |
| 6156_100 | Manic/hyper symptoms: None of the above | -0.2863 | 0.015 | -0.3699 | 0.002 | 0.4982 | 0.6151 |
| 22617_2321 | Job SOC coding: Scientific researchers | -0.2866 | 0.0215 | -0.4886 | 1.8575E-06 | 1.252 | 0.7602 |
| 1568 | Average weekly red wine intake | -0.2873 | 2.2393E-11 | -0.1783 | 1.9232E-08 | -2.0434 | 0.3514 |
| 5855_3 | Which eye(s) affected by astigmatism: Both eyes | -0.2898 | 0.0336 | -0.3424 | 0.0154 | 0.2677 | 0.573 |
| 100280 | Added milk to filtered coffee | -0.2925 | 0.0006 | -0.213 | 0.0031 | -0.7151 | 0.3904 |
| 22601_23213026 | Job coding: scientific researcher, scientific officer, medical research associate, experimental officer | -0.2934 | 0.0293 | -0.5019 | 3.4548E-06 | 1.2072 | 0.7672 |
| 22601_23113399 | Job coding: higher education teaching professional, university lecturer/professor (including college/university head/vice chancellor) | -0.2962 | 0.0008 | -0.3394 | 7.3671E-05 | 0.3517 | 0.5601 |
| 23110 | Impedance of arm (left) | -0.2979 | 3.321E-27 | -0.2302 | 1.932E-23 | -1.881 | 0.4064 |
| 4407 | Average monthly red wine intake | -0.2995 | 0.001 | -0.1911 | 0.0381 | -0.8363 | 0.3522 |
| 6038 | Number of trend entries | -0.3004 | 0.0002 | -0.2207 | 0.0037 | -0.7261 | 0.3902 |
| 6033 | Maximum heart rate during fitness test | -0.3009 | 1.8792E-06 | -0.2956 | 2.5586E-07 | -0.0621 | 0.4926 |
| 6153_5 | Medication for cholesterol, blood pressure, diabetes, or take exogenous hormones: Oral contraceptive pill or minipill | -0.3019 | 0.0092 | -0.4627 | 0.0005 | 0.9154 | 0.7132 |
| 6147_1 | Reason for glasses/contact lenses: For short-sightedness, i.e. only or mainly for distance viewing such as driving, cinema etc (called 'myopia') | -0.3035 | 1.5455E-08 | -0.2551 | 5.093E-10 | -0.7164 | 0.4327 |
| 22610_0 | Workplace full of chemical or other fumes: Rarely/never | -0.3043 | 0.0044 | -0.2052 | 0.0318 | -0.6914 | 0.3644 |
| 100017 | Magnesium | -0.3045 | 7.4225E-05 | -0.3331 | 2.9122E-07 | 0.2843 | 0.5399 |
| 6153_100 | Medication for cholesterol, blood pressure, diabetes, or take exogenous hormones: None of the above | -0.3077 | 4.9061E-13 | -0.2293 | 3.8811E-09 | -1.359 | 0.3919 |
| 6164_5 | Types of physical activity in last 4 weeks: Heavy DIY (eg: weeding, lawn mowing, carpentry, digging) | -0.312 | 7.3012E-12 | -0.315 | 9.5585E-20 | 0.0525 | 0.5042 |
| 22617_2311 | Job SOC coding: Higher education teaching professionals | -0.3124 | 0.0005 | -0.3625 | 5.9504E-06 | 0.4148 | 0.5696 |
| 6142_3 | Current employment status: Looking after home and/or family | -0.3134 | 0.0014 | -0.5637 | 2.2124E-07 | 1.707 | 0.8095 |
| 22606_0 | Workplace very noisy: Rarely/never | -0.3164 | 1.1924E-05 | -0.2631 | 0.0002 | -0.5274 | 0.426 |
| 22617_2211 | Job SOC coding: Medical practitioners | -0.318 | 0.0003 | -0.1471 | 0.0339 | -1.5367 | 0.2749 |
| 3496 | Wants to stop smoking | -0.3189 | 0.0036 | -0.2203 | 0.0121 | -0.7025 | 0.365 |
| 6160_5 | Leisure/social activities: Other group activity | -0.3195 | 2.2266E-10 | -0.2309 | 3.7044E-06 | -1.2492 | 0.3783 |
| 100005 | Carbohydrate | -0.3197 | 0.0002 | -0.3907 | 9.7321E-07 | 0.6071 | 0.5981 |
| 5001 | FI7 : synonym | -0.3206 | 1.2464E-08 | -0.3677 | 3.6236E-11 | 0.5952 | 0.5655 |
| 100009 | Englyst dietary fibre | -0.3206 | 0.0003 | -0.2409 | 0.0008 | -0.7006 | 0.3902 |
| 100390 | Tea consumed | -0.3211 | 0.0001 | -0.1603 | 0.0308 | -1.4482 | 0.2868 |
| 6032 | Maximum workload during fitness test | -0.3326 | 6.4664E-07 | -0.1404 | 0.0156 | -2.171 | 0.2506 |
| 100007 | Polyunsaturated fat | -0.3331 | 0.0319 | -0.3689 | 0.0034 | 0.1791 | 0.5499 |

| | | | | | | | |
|---|---|---|---|---|---|---|---|
| 1807 | Father's age at death | -0.3332 | 5.2519E-14 | -0.3594 | 7.4939E-11 | 0.3702 | 0.5365 |
| 1807 | Father's age at death | -0.3332 | 5.2519E-14 | -0.3594 | 7.4939E-11 | 0.3702 | 0.5365 |
| 6164_2 | Types of physical activity in last 4 weeks: Other exercises (eg: swimming, cycling, keep fit, bowling) | -0.3404 | 1.9038E-16 | -0.1396 | 7.3835E-06 | -3.8734 | 0.2411 |
| 6143_4 | Transport type for commuting to job workplace: Cycle | -0.3426 | 2.2038E-09 | -0.3021 | 4.4222E-13 | -0.5715 | 0.4436 |
| 5012 | FI8 : chained arithmetic | -0.346 | 4.6538E-12 | -0.2115 | 1.7305E-07 | -2.0903 | 0.3189 |
| 6160_4 | Leisure/social activities: Adult education class | -0.3491 | 1.4391E-09 | -0.383 | 1.0238E-15 | 0.4528 | 0.5472 |
| 20126_0 | Bipolar and major depression status: No Bipolar or Depression | -0.3504 | 1.5097E-09 | -0.1496 | 0.0038 | -2.5866 | 0.2411 |
| 2654_6 | Non-butter spread type details: Olive oil based spread (eg: Bertolli) | -0.3507 | 7.5376E-05 | -0.2448 | 0.0042 | -0.8596 | 0.3555 |
| 924 | Usual walking pace | -0.3571 | 1.6617E-27 | -0.3368 | 2.5223E-33 | -0.4699 | 0.4717 |
| 20118_11 | Home area population density - urban or rural: Scotland - Large Urban Area | -0.3572 | 1.1744E-05 | -0.1833 | 0.008 | -1.6265 | 0.2714 |
| 103990 | Vegetable consumers | -0.3579 | 0.0071 | -0.4187 | 0.0001 | 0.3521 | 0.5843 |
| 100018 | Retinol | -0.3582 | 0.0391 | -0.2667 | 0.0153 | -0.4453 | 0.3744 |
| 1578 | Average weekly champagne plus white wine intake | -0.3636 | 5.8623E-13 | -0.1215 | 0.0012 | -3.8489 | 0.1984 |
| 5984 | ECG, load | -0.3647 | 2.5494E-07 | -0.155 | 0.0125 | -2.2267 | 0.2315 |
| 6162_2 | Types of transport used (excluding work): Walk | -0.3653 | 2.8149E-18 | -0.4826 | 2.8072E-41 | 2.1259 | 0.6593 |
| 4294_1 | Final attempt correct: yes | -0.3666 | 6.7349E-06 | -0.376 | 1.3752E-06 | 0.0834 | 0.5131 |
| 1408 | Cheese intake | -0.3695 | 2.3872E-26 | -0.2901 | 6.7141E-17 | -1.6157 | 0.3906 |
| 1468_4 | Cereal type: Muesli | -0.3715 | 3.4241E-27 | -0.3903 | 6.2918E-34 | 0.3996 | 0.5262 |
| 1468_4 | Cereal type: Muesli | -0.3715 | 3.4241E-27 | -0.3903 | 6.2918E-34 | 0.3996 | 0.5262 |
| 6144_5 | Never eat eggs, dairy, wheat, sugar: I eat all of the above | -0.376 | 1.0847E-23 | -0.3168 | 4.7185E-19 | -1.1464 | 0.4179 |
| 20536_0 | Weight change during worst episode of depression: Stayed about the same or was on a diet | -0.3763 | 5.7739E-07 | -0.4285 | 2.3156E-11 | 0.5279 | 0.5725 |
| 1448_3 | Bread type: Wholemeal or wholegrain | -0.3802 | 3.0695E-22 | -0.2668 | 1.1717E-13 | -2.1307 | 0.3457 |
| 1448_3 | Bread type: Wholemeal or wholegrain | -0.3802 | 3.0695E-22 | -0.2668 | 1.1717E-13 | -2.1307 | 0.3457 |
| 6164_4 | Types of physical activity in last 4 weeks: Light DIY (eg: pruning, watering the lawn) | -0.3838 | 9.4831E-23 | -0.3554 | 1.0334E-24 | -0.5439 | 0.4604 |
| 100014 | Folate | -0.3879 | 3.2974E-05 | -0.2371 | 0.0005 | -1.3073 | 0.2988 |
| 20016 | Fluid intelligence score | -0.3927 | 5.1487E-30 | -0.2852 | 2.7968E-20 | -2.3211 | 0.3534 |
| 20111_101 | Illnesses of siblings: None of the above (group 2) | -0.3941 | 7.5034E-08 | -0.1185 | 0.0479 | -2.9114 | 0.1674 |
| 22608_0 | Workplace very hot: Rarely/never | -0.3947 | 5.3738E-07 | -0.168 | 0.0074 | -2.253 | 0.2138 |
| 20111_100 | Illnesses of siblings: None of the above (group 1) | -0.395 | 2.6312E-19 | -0.2476 | 2.2302E-10 | -2.507 | 0.303 |
| 22607_0 | Workplace very cold: Rarely/never | -0.3957 | 2.1996E-09 | -0.2899 | 1.7469E-05 | -1.1199 | 0.3556 |
| 1458 | Cereal intake | -0.3967 | 1.4332E-28 | -0.3402 | 5.9828E-29 | -1.2013 | 0.4216 |
| 4935 | FI1 : numeric addition test | -0.405 | 0.0019 | -0.2191 | 0.0499 | -1.0837 | 0.2577 |
| 5699 | FI10 : arithmetic sequence recognition | -0.4051 | 1.2649E-08 | -0.2408 | 0.0001 | -1.7282 | 0.2826 |
| 4990 | FI6 : conditional arithmetic | -0.406 | 1.1187E-16 | -0.3608 | 3.4505E-19 | -0.7124 | 0.4372 |
| 6154_100 | Medication for pain relief, constipation, heartburn: None of the above | -0.4072 | 1.9226E-31 | -0.3734 | 2.2818E-29 | -0.7017 | 0.4529 |

| | | | | | | | |
|---|---|---|---|---|---|---|---|
| 3536 | Age started hormone-replacement therapy (HRT) | -0.4077 | 1.3999E-14 | -0.4826 | 1.344E-22 | 1.0348 | 0.6034 |
| 6143_3 | Transport type for commuting to job workplace: Public transport | -0.4122 | 2.0224E-13 | -0.4268 | 1.4514E-16 | 0.1914 | 0.5204 |
| 6160_3 | Leisure/social activities: Religious group | -0.4123 | 1.2398E-30 | -0.2862 | 8.5033E-19 | -2.6152 | 0.3295 |
| 100025 | Vitamin E | -0.4161 | 0.0002 | -0.3603 | 2.9705E-05 | -0.3993 | 0.4226 |
| 6149_100 | Mouth/teeth dental problems: None of the above | -0.4174 | 1.7748E-29 | -0.3597 | 1.9964E-21 | -1.0908 | 0.42 |
| 1618 | Alcohol usually taken with meals | -0.4201 | 5.549E-28 | -0.1539 | 7.4014E-06 | -5.1776 | 0.1758 |
| 20489 | Felt loved as a child | -0.4236 | 8.7201E-22 | -0.1236 | 0.0026 | -4.9705 | 0.1469 |
| 680_1 | Own or rent accommodation lived in: Own outright (by you or someone in your household) | -0.4316 | 4.2403E-25 | -0.6071 | 2.6952E-79 | 3.3311 | 0.7305 |
| 1249 | Past tobacco smoking | -0.4487 | 9.1303E-54 | -0.2932 | 5.8352E-33 | -4.0878 | 0.2932 |
| 100760 | Breakfast cereal consumed | -0.4511 | 1.8311E-05 | -0.3562 | 0.0002 | -0.6695 | 0.3699 |
| 20491 | Someone to take to doctor when needed as a child | -0.4519 | 1.0801E-12 | -0.3711 | 1.1645E-09 | -0.9176 | 0.3887 |
| 5507 | Leg pain on walking : action taken | -0.4532 | 1.0429E-05 | -0.4582 | 1.7646E-06 | 0.0356 | 0.507 |
| 6138_6 | Qualifications: Other professional qualifications eg: nursing, teaching | -0.4567 | 5.2043E-40 | -0.2751 | 8.7955E-17 | -3.7983 | 0.2625 |
| 6164_1 | Types of physical activity in last 4 weeks: Walking for pleasure (not as a means of transport) | -0.4578 | 1.2512E-29 | -0.3367 | 5.8216E-25 | -2.3293 | 0.3359 |
| 1608 | Average weekly fortified wine intake | -0.4629 | 3.0106E-09 | -0.2309 | 0.0016 | -2.1716 | 0.2084 |
| 3872 | Age of primiparous women at birth of child | -0.4636 | 7.1709E-07 | -0.4114 | 3.774E-07 | -0.422 | 0.4275 |
| 110001 | Invitation to complete online 24-hour recall dietary questionnaire, acceptance | -0.4646 | 8.8157E-27 | -0.4214 | 6.7945E-25 | -0.7244 | 0.4399 |
| 20117_2 | Alcohol drinker status: Current | -0.4716 | 4.7735E-21 | -0.2622 | 9.4854E-09 | -3.0879 | 0.2318 |
| 3436 | Age started smoking in current smokers | -0.472 | 1.1367E-09 | -0.3682 | 5.6843E-08 | -1.008 | 0.3582 |
| 1538_0 | Major dietary changes in the last 5 years: No | -0.4813 | 3.1135E-31 | -0.3956 | 5.252E-30 | -1.5846 | 0.3821 |
| 1538_0 | Major dietary changes in the last 5 years: No | -0.4813 | 3.1135E-31 | -0.3956 | 5.252E-30 | -1.5846 | 0.3821 |
| 738 | Average total household income before tax | -0.4881 | 1.3791E-61 | -0.1705 | 8.0953E-11 | -8.0497 | 0.1332 |
| 100011 | Iron | -0.4885 | 2.5785E-07 | -0.3613 | 6.9833E-07 | -1.0642 | 0.3281 |
| 100023 | Starch | -0.4896 | 0.0003 | -0.5666 | 2.6278E-05 | 0.4036 | 0.6062 |
| 4957 | FI3 : word interpolation | -0.4975 | 5.2256E-28 | -0.4306 | 6.7962E-26 | -1.0961 | 0.4074 |
| 6138_1 | Qualifications: College or University degree | -0.5103 | 7.5189E-74 | -0.3801 | 1.699E-58 | -3.5481 | 0.3243 |
| 3526 | Mother's age at death | -0.517 | 3.8314E-13 | -0.4264 | 5.0545E-12 | -0.961 | 0.3756 |
| 104400 | Fruit consumers | -0.5207 | 3.0161E-07 | -0.3259 | 0.0002 | -1.4527 | 0.2477 |
| 20116_0 | Smoking status: Never | -0.5228 | 1.9097E-79 | -0.3863 | 6.9316E-56 | -3.6912 | 0.3164 |
| 6142_6 | Current employment status: Doing unpaid or voluntary work | -0.5265 | 5.9178E-11 | -0.327 | 1.8924E-06 | -1.8876 | 0.2425 |
| 6159_100 | Pain type(s) experienced in last month: None of the above | -0.5332 | 1.0018E-53 | -0.3898 | 4.9949E-47 | -3.2628 | 0.3079 |
| 6145_100 | Illness, injury, bereavement, stress in last 2 years: None of the above | -0.5348 | 4.2324E-34 | -0.4924 | 4.8283E-29 | -0.6822 | 0.441 |
| 4968 | FI4 : positional arithmetic | -0.5391 | 2.001E-31 | -0.3082 | 3.0707E-14 | -3.7542 | 0.2095 |
| 5556 | FI9 : concept interpolation | -0.5413 | 2.6166E-20 | -0.4229 | 5.3681E-15 | -1.4846 | 0.3393 |
| 22506_114 | Tobacco smoking: Never smoked | -0.5478 | 8.8979E-27 | -0.3866 | 3.4557E-19 | -2.4091 | 0.2863 |
| 6017 | Able to walk or cycle unaided for 10 minutes | -0.5512 | 0.0034 | -0.4731 | 0.0013 | -0.327 | 0.3923 |

| | | | | | | | |
|---|---|---|---|---|---|---|---|
| 2764 | Age at last live birth | -0.5583 | 1.2146E-45 | -0.3901 | 2.3952E-24 | -3.0611 | 0.278 |
| 2794 | Age started oral contraceptive pill | -0.5709 | 6.316E-38 | -0.5123 | 4.3708E-43 | -1.013 | 0.4188 |
| 20525 | Able to pay rent/mortgage as an adult | -0.5713 | 3.3373E-11 | -0.3746 | 8.1481E-07 | -1.7116 | 0.2456 |
| 6138_3 | Qualifications: O levels/GCSEs or equivalent | -0.572 | 5.2421E-49 | -0.3307 | 1.2513E-23 | -4.7303 | 0.1992 |
| 22501 | Year ended full time education | -0.5773 | 1.9606E-48 | -0.4257 | 2.6062E-33 | -2.8581 | 0.2979 |
| 100890 | Milk added to cereal | -0.5862 | 2.3733E-05 | -0.3921 | 0.001 | -1.0602 | 0.2485 |
| 6138_2 | Qualifications: A levels/AS levels or equivalent | -0.609 | 7.9549E-89 | -0.4058 | 2.1449E-50 | -4.9723 | 0.2385 |
| 2867 | Age started smoking in former smokers | -0.6185 | 4.8085E-32 | -0.4579 | 6.8526E-24 | -2.3139 | 0.287 |
| 2139 | Age first had sexual intercourse | -0.6231 | 2.173E-128 | -0.5791 | 6.238E-132 | -1.2533 | 0.4388 |
| 6146_100 | Attendance/disability/mobility allowance: None of the above | -0.6547 | 1.4243E-55 | -0.5784 | 3.1485E-56 | -1.3752 | 0.3947 |
| 845 | Age completed full time education | -0.6549 | 4.2654E-98 | -0.3804 | 1.6693E-47 | -6.727 | 0.1684 |
| 2754 | Age at first live birth | -0.6612 | 4.2769E-92 | -0.4658 | 2.5192E-56 | -4.4518 | 0.247 |

**Table S25.** Linkage disequilibrium score regression (rg) results for 478 normalized UK Biobank traits demonstrating nominally significant genetic correlations with both attention deficit hyperactivity disorder (ADHD) and *PhoneUse* in female subjects.

| Field ID | Trait | ADHD | | *PhoneUse* | | Difference | |
|---|---|---|---|---|---|---|---|
| | | rg | p | rg | p | z | p |
| 2139 | Age at first sexual intercourse | -0.8147 | 3.2497E-22 | -0.5286 | 1.3125E-64 | -3.1907 | 0.2279 |
| 2178 | Overall health rating | 0.7439 | 6.712E-19 | 0.4216 | 3.139E-28 | 3.498 | 0.798 |
| 6138_1 | Qualifications: College or University degree | -0.6177 | 2.69E-17 | -0.3589 | 2.7532E-22 | -3.1622 | 0.2631 |
| 23115 | Leg fat percentage (left) | 0.4599 | 6.2702E-16 | 0.3374 | 4.5074E-26 | 1.8765 | 0.6268 |
| 6138_2 | Qualifications: A levels/AS levels or equivalent | -0.7156 | 9.3415E-16 | -0.4134 | 2.4659E-25 | -3.0968 | 0.2291 |
| 20116_0 | Smoking status: Never | -0.6755 | 1.7241E-15 | -0.3931 | 2.9213E-32 | -3.0978 | 0.2304 |
| 23111 | Leg fat percentage (right) | 0.4419 | 4.0731E-15 | 0.3307 | 1.4991E-24 | 1.7132 | 0.6158 |
| 845 | Age completed full time education | -0.7178 | 9.4227E-15 | -0.452 | 8.3378E-32 | -2.648 | 0.2452 |
| 738 | Average total household income before tax | -0.6516 | 1.4161E-14 | -0.2742 | 3.7952E-10 | -3.9579 | 0.1638 |
| 6146_100 | Attendance/disability/mobility allowance: None | -0.9383 | 1.625E-14 | -0.6091 | 7.6861E-30 | -2.4663 | 0.2058 |
| 137 | Number of treatments/medications taken | 0.6009 | 3.487E-14 | 0.3412 | 1.5092E-14 | 2.8575 | 0.7498 |
| 6138_100 | Qualifications: None of the above | 0.609 | 4.303E-14 | 0.3793 | 1.6321E-22 | 2.5666 | 0.7136 |
| XI_DIGESTIVE | Diseases of the digestive system | 0.8729 | 9.6121E-14 | 0.4973 | 2.102E-20 | 2.9135 | 0.8693 |
| 2492 | Taking other prescription medications | 0.6393 | 1.0708E-13 | 0.3467 | 1.0982E-14 | 3.016 | 0.7754 |
| 1239 | Current tobacco smoking | 0.7496 | 1.3778E-13 | 0.5293 | 9.4263E-32 | 1.9867 | 0.7159 |
| 6145_6 | Illness, injury, bereavement, stress in last 2 years: Financial difficulties | 0.7792 | 1.5497E-13 | 0.5787 | 1.6725E-26 | 1.6898 | 0.6914 |
| 1249 | Past tobacco smoking | -0.5641 | 1.8727E-13 | -0.2808 | 5.837E-14 | -3.32 | 0.2314 |
| 6146_3 | Attendance/disability/mobility allowance: Blue badge | 1 | 1.9616E-13 | 0.6529 | 1.1911E-20 | 2.1588 | 0.8083 |
| 21001 | Body mass index (BMI) | 0.4426 | 2.2871E-13 | 0.3281 | 4.4877E-25 | 1.6786 | 0.6174 |
| 23116 | Leg fat mass (left) | 0.4244 | 3.0492E-13 | 0.3174 | 1.0265E-23 | 1.6157 | 0.6112 |
| 6146_2 | Attendance/disability/mobility allowance: Disability living allowance | 0.9309 | 3.3157E-13 | 0.5789 | 5.4027E-23 | 2.502 | 0.8114 |
| 680_3 | Own or rent accommodation lived in: Rent - from local authority, local council, housing association | 0.8188 | 4.359E-13 | 0.4829 | 2.4399E-21 | 2.7103 | 0.8056 |
| 20116_2 | Smoking status: Current | 0.7522 | 4.3702E-13 | 0.5289 | 2.3341E-37 | 1.9982 | 0.7204 |
| 23104 | Body mass index (BMI) | 0.4325 | 5.6158E-13 | 0.3309 | 1.6371E-25 | 1.4972 | 0.606 |
| XVIII_MISCFINDINGS | Symptoms, signs and abnormal clinical and laboratory findings, not elsewhere classified | 0.7336 | 6.3667E-13 | 0.5629 | 2.5879E-27 | 1.491 | 0.7308 |
| 23112 | Leg fat mass (right) | 0.4155 | 7.6314E-13 | 0.3148 | 2.9324E-23 | 1.5235 | 0.6051 |
| 23119 | Arm fat percentage (right) | 0.4007 | 7.8974E-13 | 0.304 | 1.0122E-21 | 1.5048 | 0.6005 |
| 23099 | Body fat percentage | 0.3844 | 8.5458E-13 | 0.2882 | 9.3784E-20 | 1.5427 | 0.6007 |
| 2188 | Long-standing illness, disability or infirmity | 0.6342 | 1.1634E-12 | 0.3643 | 2.7049E-14 | 2.6657 | 0.7577 |
| 6164_1 | Types of physical activity in last 4 weeks: Walking for pleasure (not as a means of transport) | -0.6558 | 1.1718E-12 | -0.3569 | 4.5917E-12 | -2.8266 | 0.2239 |
| 23123 | Arm fat percentage (left) | 0.4014 | 1.2004E-12 | 0.3054 | 1.5335E-21 | 1.4785 | 0.5995 |
| 6142_4 | Current employment status: Unable to work because of sickness or disability | 0.9274 | 1.3184E-12 | 0.5089 | 1.7141E-13 | 2.829 | 0.85 |
| 6164_100 | Types of physical activity in last 4 weeks: None of the above | 0.8912 | 2.6659E-12 | 0.5064 | 1.9277E-13 | 2.6568 | 0.8367 |
| 2060 | Frequency of unenthusiasm / disinterest in last 2 weeks | 0.751 | 2.7666E-12 | 0.3526 | 1.6983E-11 | 3.3314 | 0.8488 |
| 23120 | Arm fat mass (right) | 0.4067 | 2.9621E-12 | 0.308 | 3.199E-22 | 1.4862 | 0.6025 |
| 23100 | Whole body fat mass | 0.382 | 4.1118E-12 | 0.285 | 5.8552E-20 | 1.5319 | 0.6015 |
| 48 | Waist circumference | 0.4123 | 4.344E-12 | 0.2758 | 7.2798E-20 | 2.0457 | 0.6334 |
| 23124 | Arm fat mass (left) | 0.3991 | 5.9754E-12 | 0.3078 | 4.0816E-22 | 1.3803 | 0.5947 |
| 1618 | Alcohol usually taken with meals | -0.6699 | 8.2063E-12 | -0.2416 | 7.0099E-06 | -3.8311 | 0.1335 |
| XIII_MUSCULOSKELET | Diseases of the musculoskeletal system and connective tissue | 0.634 | 1.094E-11 | 0.4732 | 1.8504E-22 | 1.5292 | 0.7016 |
| 6154_100 | Medication for pain relief, constipation, heartburn: None of the above | -0.5341 | 1.3158E-11 | -0.3874 | 1.6253E-18 | -1.623 | 0.3569 |
| 1787 | Maternal smoking around birth | 0.7389 | 1.689E-11 | 0.4702 | 7.456E-21 | 2.2256 | 0.757 |
| 1050 | Time spend outdoors in summer | 0.5627 | 1.9642E-11 | 0.409 | 2.1829E-20 | 1.6208 | 0.655 |
| 189 | Townsend deprivation index at recruitment | 0.6899 | 2.1519E-11 | 0.3812 | 2.2349E-14 | 2.6972 | 0.7893 |
| 189 | Townsend deprivation index at recruitment | 0.6899 | 2.1519E-11 | 0.3812 | 2.2349E-14 | 2.6972 | 0.7894 |
| 2316 | Wheeze or whistling in the chest in last year | 0.6109 | 2.607E-11 | 0.445 | 2.175E-23 | 1.6277 | 0.6662 |

| | | | | | | | |
|---|---|---|---|---|---|---|---|
| 6159_100 | Pain type(s) experienced in last month: None of the above | -0.5876 | 4.108E-11 | -0.4117 | 6.0693E-22 | -1.7811 | 0.3293 |
| 2080 | Frequency of tiredness / lethargy in last 2 weeks | 0.5965 | 4.8857E-11 | 0.3124 | 9.0862E-13 | 2.8219 | 0.769 |
| 6159_7 | Pain type(s) experienced in last month: Knee pain | 0.5797 | 6.157E-11 | 0.3575 | 1.6055E-15 | 2.237 | 0.7118 |
| 20489 | Felt loved as a child | -0.5862 | 8.4533E-11 | -0.1197 | 0.0211 | -4.479 | 0.1122 |
| 2070 | Frequency of tenseness / restlessness in last 2 weeks | 0.5984 | 9.9499E-11 | 0.326 | 1.7515E-12 | 2.6345 | 0.7597 |
| 2149 | Lifetime number of sexual partners | 0.5239 | 1.0172E-10 | 0.3559 | 4.2031E-23 | 1.8953 | 0.6682 |
| 20488 | Physically abused by family as a child | 0.7685 | 1.4126E-10 | 0.3479 | 1.2713E-06 | 3.0114 | 0.8633 |
| 6149_6 | Mouth/teeth dental problems: Dentures | 0.591 | 1.8699E-10 | 0.3765 | 1.7544E-14 | 2.0431 | 0.7049 |
| 2090 | Seen doctor (GP) for nerves, anxiety, tension or depression | 0.5347 | 2.4758E-10 | 0.2587 | 1.9873E-09 | 2.9096 | 0.7626 |
| 4548 | Health satisfaction | 0.6714 | 3.1578E-10 | 0.3853 | 2.4819E-12 | 2.3833 | 0.7702 |
| 2814 | Ever used hormone-replacement therapy (HRT) | 0.562 | 3.3036E-10 | 0.3858 | 1.3235E-15 | 1.734 | 0.6755 |
| 23128 | Trunk fat mass | 0.3303 | 3.4513E-10 | 0.2489 | 2.3847E-16 | 1.3399 | 0.5843 |
| 2100 | Seen a psychiatrist for nerves, anxiety, tension or depression | 0.6394 | 4.4751E-10 | 0.2466 | 1.4205E-06 | 3.4296 | 0.8454 |
| 22506_114 | Tobacco smoking: Never smoked | -0.7698 | 4.5191E-10 | -0.3093 | 1.5674E-07 | -3.3645 | 0.1141 |
| 23127 | Trunk fat percentage | 0.321 | 5.3201E-10 | 0.2432 | 5.7678E-15 | 1.2895 | 0.5807 |
| 4717 | Shortness of breath walking on level ground | 0.735 | 5.7914E-10 | 0.4249 | 2.6208E-09 | 2.2401 | 0.7887 |
| 2020 | Loneliness, isolation | 0.5542 | 6.9751E-10 | 0.3239 | 6.6373E-11 | 2.243 | 0.7245 |
| ICDMAIN_ANY_ ENTRY | Any ICDMAIN event in hilmo or causes of death | 0.796 | 7.6171E-10 | 0.5602 | 3.3219E-20 | 1.6493 | 0.7278 |
| 21002 | Weight | 0.344 | 8.4155E-10 | 0.2488 | 1.5382E-16 | 1.4974 | 0.5981 |
| 6160_3 | Leisure/social activities: Religious group | -0.5406 | 1.0852E-09 | -0.3087 | 4.7257E-12 | -2.3347 | 0.2798 |
| 1408 | Cheese intake | -0.4953 | 1.104E-09 | -0.3433 | 8.6732E-14 | -1.6272 | 0.3469 |
| 136 | Number of operations, self-reported | 0.6423 | 1.1452E-09 | 0.44 | 8.0274E-28 | 1.7919 | 0.7002 |
| R07 | Diagnoses - main ICD10: R07 Pain in throat and chest | 0.8226 | 1.4825E-09 | 0.5543 | 2.9409E-15 | 1.753 | 0.775 |
| 23098 | Weight | 0.3352 | 1.6155E-09 | 0.2522 | 7.3826E-17 | 1.3118 | 0.5872 |
| 20160 | Ever smoked | 0.473 | 1.6503E-09 | 0.2619 | 6.5813E-12 | 2.4193 | 0.7094 |
| 2050 | Frequency of depressed mood in last 2 weeks | 0.613 | 1.6911E-09 | 0.2807 | 1.0214E-08 | 2.9436 | 0.8051 |
| VI_NERVOUS | Diseases of the nervous system | 0.7579 | 1.7412E-09 | 0.479 | 5.8742E-12 | 1.9387 | 0.7812 |
| 6154_5 | Medication for pain relief, constipation, heartburn: Omeprazole (e.g. Zanprol) | 0.7206 | 2.2749E-09 | 0.485 | 2.8426E-14 | 1.7268 | 0.7222 |
| 4728 | Leg pain on walking | 0.8075 | 2.4005E-09 | 0.5043 | 2.7227E-12 | 1.9777 | 0.7836 |
| 2277 | Frequency of solarium/sunlamp use | 0.7854 | 2.7649E-09 | 0.6844 | 1.0661E-32 | 0.701 | 0.6031 |
| 49 | Hip circumference | 0.3263 | 4.1805E-09 | 0.2256 | 1.3405E-12 | 1.5743 | 0.5992 |
| 41248_1000 | Destinations on discharge from hospital (recoded): Usual Place of residence | 0.8158 | 4.3085E-09 | 0.5946 | 3.6032E-18 | 1.4287 | 0.7106 |
| 2335 | Chest pain or discomfort | 0.6556 | 4.5914E-09 | 0.3326 | 3.5E-11 | 2.6336 | 0.7984 |
| XIV_GENITOUR INARY | Diseases of the genitourinary system | 0.6824 | 5.9824E-09 | 0.4708 | 8.116E-17 | 1.6252 | 0.758 |
| 20116_1 | Smoking status: Previous | 0.5104 | 6.1629E-09 | 0.2452 | 3.5545E-08 | 2.6942 | 0.756 |
| 6159_3 | Pain type(s) experienced in last month: Neck or shoulder pain | 0.5734 | 8.8407E-09 | 0.3802 | 6.7444E-12 | 1.6939 | 0.6867 |
| 20003_114086563 4 | Treatment/medication code: omeprazole | 0.7854 | 1.1124E-08 | 0.5495 | 1.2342E-13 | 1.5103 | 0.7301 |
| M54 | Diagnoses - main ICD10: M54 Dorsalgia | 0.8464 | 1.1804E-08 | 0.4869 | 1.9679E-12 | 2.1955 | 0.803 |
| M13_DORSALGI A | Dorsalgia | 0.8464 | 1.1804E-08 | 0.4869 | 1.9679E-12 | 2.1955 | 0.805 |
| 1960 | Fed-up feelings | 0.4538 | 1.1994E-08 | 0.1932 | 3.3646E-05 | 2.8253 | 0.7503 |
| 20521 | Belittlement by partner or ex-partner as an adult | 0.7004 | 1.2319E-08 | 0.3397 | 4.9992E-08 | 2.6161 | 0.8261 |
| 20523 | Physical violence by partner or ex-partner as an adult | 0.7976 | 1.3474E-08 | 0.4689 | 9.52E-10 | 2.0552 | 0.8039 |
| 6138_6 | Qualifications: Other professional qualifications eg: nursing, teaching | -0.4961 | 1.4101E-08 | -0.3406 | 2.2709E-14 | -1.5833 | 0.3506 |
| 924 | Usual walking pace | -0.3858 | 1.5432E-08 | -0.3434 | 1.6913E-16 | -0.5304 | 0.4562 |
| 1920 | Mood swings | 0.4704 | 1.5517E-08 | 0.2207 | 5.6183E-07 | 2.6517 | 0.7413 |
| 1070 | Time spent watching television (TV) | 0.3734 | 1.7324E-08 | 0.216 | 8.7903E-07 | 1.9815 | 0.6585 |
| 20016 | Fluid intelligence score | -0.4894 | 2.0187E-08 | -0.2777 | 3.3344E-09 | -2.1371 | 0.2907 |
| 6159_4 | Pain type(s) experienced in last month: Back pain | 0.5941 | 3.3421E-08 | 0.4925 | 4.8646E-22 | 0.8532 | 0.6009 |
| 6149_100 | Mouth/teeth dental problems: None of the above | -0.5515 | 3.4754E-08 | -0.325 | 5.3655E-09 | -1.9788 | 0.2847 |

| | | | | | | | |
|---|---|---|---|---|---|---|---|
| 1930 | Miserableness | 0.5254 | 3.9506E-08 | 0.113 | 0.0276 | 3.798 | 0.8575 |
| 41231_1 | Hospital episode type: General episode | 0.8155 | 4.9709E-08 | 0.5598 | 7.5435E-15 | 1.5401 | 0.7393 |
| 20002_1465 | Non-cancer illness code, self-reported: osteoarthritis | 0.5841 | 6.3546E-08 | 0.3168 | 2.8227E-07 | 2.149 | 0.7565 |
| 1200 | Sleeplessness / insomnia | 0.4142 | 6.8752E-08 | 0.2775 | 1.2905E-09 | 1.5296 | 0.6384 |
| 680_1 | Own or rent accommodation lived in: Own outright (by you or someone in your household) | -0.5669 | 9.2286E-08 | -0.6266 | 1.4438E-30 | 0.5005 | 0.5607 |
| 22501 | Year ended full time education | -0.5671 | 9.7359E-08 | -0.4448 | 2.6055E-13 | -0.998 | 0.3748 |
| 22506_113 | Tobacco smoking: Ex-smoker | 0.7782 | 1.0599E-07 | 0.2693 | 3.0016E-05 | 3.181 | 0.9084 |
| 6144_4 | Never eat eggs, dairy, wheat, sugar: Sugar or foods/drinks containing sugar | 0.4806 | 1.3069E-07 | 0.331 | 2.2247E-12 | 1.4581 | 0.6449 |
| 1468_4 | Cereal type: Muesli | -0.5116 | 1.617E-07 | -0.427 | 4.3293E-15 | -0.7565 | 0.4126 |
| 20459 | General happiness with own health | 0.5577 | 1.6573E-07 | 0.2375 | 6.0648E-05 | 2.626 | 0.7976 |
| 6138_3 | Qualifications: O levels/GCSEs or equivalent | -0.6041 | 1.7989E-07 | -0.3368 | 1.1534E-11 | -2.1218 | 0.2552 |
| 20487 | Felt hated by family member as a child | 0.6819 | 2.3138E-07 | 0.3271 | 8.6473E-07 | 2.4034 | 0.8222 |
| 6159_8 | Pain type(s) experienced in last month: Pain all over the body | 0.8269 | 2.3532E-07 | 0.3914 | 6.7671E-06 | 2.3912 | 0.8632 |
| IX_CIRCULATORY | Diseases of the circulatory system | 0.6308 | 2.5309E-07 | 0.3454 | 1.1389E-08 | 2.0903 | 0.7682 |
| 1269 | Exposure to tobacco smoke at home | 0.7028 | 2.6008E-07 | 0.5638 | 1.9035E-10 | 0.8544 | 0.6408 |
| 2774 | Ever had stillbirth, spontaneous miscarriage or termination | 0.7252 | 2.8677E-07 | 0.5669 | 2.6394E-12 | 0.9719 | 0.6587 |
| 110001 | Invitation to complete online 24-hour recall dietary questionnaire, acceptance | -0.6565 | 2.8982E-07 | -0.4406 | 9.3803E-11 | -1.4896 | 0.2867 |
| 20110_6 | Illnesses of mother: Chronic bronchitis/emphysema | 0.7765 | 3.0787E-07 | 0.3959 | 4.9979E-07 | 2.2264 | 0.8393 |
| 1558 | Alcohol intake frequency. | 0.3728 | 3.3033E-07 | 0.2108 | 1.1355E-08 | 1.9805 | 0.6626 |
| 20117_1 | Alcohol drinker status: Previous | 0.719 | 3.4042E-07 | 0.357 | 1.7229E-06 | 2.2693 | 0.8281 |
| 4968 | FI4 : positional arithmetic | -0.5944 | 3.8571E-07 | -0.3079 | 4.9508E-06 | -2.1205 | 0.229 |
| 1538_0 | Major dietary changes in the last 5 years: No | -0.4936 | 3.8917E-07 | -0.4178 | 3.3233E-19 | -0.7026 | 0.4216 |
| 6164_2 | Types of physical activity in last 4 weeks: Other exercises (eg: swimming, cycling, keep fit, bowling) | -0.5111 | 4.1823E-07 | -0.208 | 1.8082E-06 | -2.7552 | 0.2196 |
| 6159_6 | Pain type(s) experienced in last month: Hip pain | 0.5988 | 4.2319E-07 | 0.3217 | 1.1622E-07 | 2.0826 | 0.7571 |
| 2296 | Falls in the last year | 0.5047 | 4.4718E-07 | 0.3516 | 4.2045E-12 | 1.3655 | 0.6541 |
| Risk Taking | #N/A | 0.4262 | 5.0328E-07 | 0.4164 | 1.5117E-20 | 0.1022 | 0.5101 |
| 1279 | Exposure to tobacco smoke outside home | 0.6071 | 5.0348E-07 | 0.4933 | 2.4151E-13 | 0.8227 | 0.6161 |
| 23109 | Impedance of arm (right) | -0.3082 | 5.6103E-07 | -0.2333 | 8.617E-16 | -1.1001 | 0.4215 |
| M13_SHOULDER | Shoulder lesions | 1 | 6.3655E-07 | 0.5621 | 4.627E-06 | 1.6936 | 0.8473 |
| M75 | Diagnoses - main ICD10: M75 Shoulder lesions | 1 | 6.4208E-07 | 0.5653 | 3.8483E-06 | 1.685 | 0.8487 |
| 23110 | Impedance of arm (left) | -0.3047 | 7.1231E-07 | -0.2355 | 3.0856E-16 | -1.019 | 0.4274 |
| 6160_100 | Leisure/social activities: None of the above | 0.4921 | 8.019E-07 | 0.1937 | 0.0001 | 2.6743 | 0.7739 |
| 20511 | Recent poor appetite or overeating | 0.5956 | 8.8275E-07 | 0.3492 | 2.0573E-07 | 1.778 | 0.7394 |
| 23125 | Arm fat-free mass (left) | 0.2738 | 9.0056E-07 | 0.1993 | 2.1504E-11 | 1.1793 | 0.5774 |
| 6144_5 | Never eat eggs, dairy, wheat, sugar: I eat all of the above | -0.4662 | 9.3281E-07 | -0.3928 | 1.0443E-15 | -0.6867 | 0.4277 |
| PULMONARYDG | Other pulmonary diagnosis | 0.8099 | 9.4739E-07 | 0.501 | 1.6294E-06 | 1.5802 | 0.8101 |
| X_RESPIRATORY | Diseases of the respiratory system | 0.8099 | 9.4739E-07 | 0.501 | 1.6294E-06 | 1.5802 | 0.8185 |
| K29 | Diagnoses - main ICD10: K29 Gastritis and duodenitis | 0.7743 | 1.1449E-06 | 0.3269 | 0.0002 | 2.468 | 0.8697 |
| M13_LOWBACKPAIN | Low back pain | 0.8606 | 1.3065E-06 | 0.3905 | 1.2418E-06 | 2.4086 | 0.867 |
| 23126 | Arm predicted mass (left) | 0.2706 | 1.3188E-06 | 0.2049 | 6.1518E-12 | 1.0357 | 0.5683 |
| 2844 | Had other major operations | 0.5931 | 1.3418E-06 | 0.3405 | 1.0703E-09 | 1.874 | 0.7429 |
| ILD_DIFF_DG | ILD differential diagnosis | 0.7976 | 1.3828E-06 | 0.4689 | 2.0343E-06 | 1.7081 | 0.8003 |
| COPD_EXCL | COPD differential diagnosis | 0.7976 | 1.3828E-06 | 0.4689 | 2.0343E-06 | 1.7081 | 0.8057 |
| 20003_2038460150 | Treatment/medication code: paracetamol | 0.5367 | 1.3836E-06 | 0.3069 | 2.1952E-07 | 1.8241 | 0.7249 |
| 6154_6 | Medication for pain relief, constipation, heartburn: Laxatives (e.g. Dulcolax, Senokot) | 0.6498 | 1.4457E-06 | 0.4908 | 9.9699E-11 | 1.0278 | 0.6546 |
| 2237 | Plays computer games | 0.3801 | 1.5198E-06 | 0.2098 | 2.9636E-08 | 1.9426 | 0.6704 |

| | | | | | | | |
|---|---|---|---|---|---|---|---|
| 20003_1140923346 | Treatment/medication code: co-codamol | 0.7783 | 1.8081E-06 | 0.5596 | 8.9499E-10 | 1.1706 | 0.7158 |
| 20107_6 | Illnesses of father: Chronic bronchitis/emphysema | 0.6895 | 2.1342E-06 | 0.2838 | 0.0004 | 2.4475 | 0.8548 |
| 30530 | Sodium in urine | 0.319 | 2.2796E-06 | 0.3249 | 1.2919E-13 | -0.0733 | 0.494 |
| 6145_1 | Illness, injury, bereavement, stress in last 2 years: Serious illness, injury or assault to yourself | 0.7172 | 2.2864E-06 | 0.3104 | 0.0002 | 2.3402 | 0.8441 |
| 1458 | Cereal intake | -0.44 | 2.596E-06 | -0.3014 | 2.0678E-11 | -1.3345 | 0.3597 |
| 6154_3 | Medication for pain relief, constipation, heartburn: Paracetamol | 0.4537 | 2.6847E-06 | 0.282 | 1.5471E-07 | 1.5523 | 0.6664 |
| 2473 | Other serious medical condition/disability diagnosed by doctor | 0.5 | 2.8288E-06 | 0.203 | 0.0014 | 2.3923 | 0.7788 |
| 6153_100 | Medication for cholesterol, blood pressure, diabetes, or take exogenous hormones: None of the above | -0.3682 | 2.935E-06 | -0.2206 | 2.2961E-07 | -1.6477 | 0.356 |
| N39 | Diagnoses - main ICD10: N39 Other disorders of urinary system | 0.6343 | 3.2988E-06 | 0.5294 | 2.7668E-11 | 0.6644 | 0.5982 |
| 2867 | Age started smoking in former smokers | -0.8371 | 3.7909E-06 | -0.4782 | 9.4726E-08 | -1.7763 | 0.1771 |
| 20117_2 | Alcohol drinker status: Current | -0.4977 | 3.885E-06 | -0.2662 | 1.5311E-05 | -1.8645 | 0.2724 |
| 20519 | Recent feelings of tiredness or low energy | 0.5615 | 3.9434E-06 | 0.2769 | 0.00001615 | 2.0684 | 0.7706 |
| K11_OTHGASTR | Other gastritis (incl. Duodenitis) | 0.7485 | 4.0462E-06 | 0.2843 | 0.0035 | 2.4513 | 0.8795 |
| 23105 | Basal metabolic rate | 0.2517 | 4.0535E-06 | 0.1882 | 1.576E-10 | 1.024 | 0.5667 |
| R10 | Diagnoses - main ICD10: R10 Abdominal and pelvic pain | 0.7015 | 4.1803E-06 | 0.647 | 2.0644E-17 | 0.3199 | 0.5612 |
| 23106 | Impedance of whole body | -0.276 | 4.2816E-06 | -0.2046 | 2.1294E-11 | -1.0601 | 0.4252 |
| 1448_1 | Bread type: White | 0.3883 | 4.285E-06 | 0.2755 | 4.0443E-07 | 1.1224 | 0.6158 |
| 6145_100 | Illness, injury, bereavement, stress in last 2 years: None of the above | -0.6325 | 4.4604E-06 | -0.5025 | 5.7863E-10 | -0.813 | 0.3733 |
| 4631 | Ever unenthusiastic/disinterested for a whole week | 0.6405 | 4.6467E-06 | 0.2655 | 0.0003 | 2.3785 | 0.8337 |
| 20491 | Someone to take to doctor when needed as a child | -0.6929 | 5.8209E-06 | -0.3601 | 0.00003733 | -1.8902 | 0.1928 |
| 6152_6 | Blood clot, DVT, bronchitis, emphysema, asthma, rhinitis, eczema, allergy diagnosed by doctor: Emphysema/chronic bronchitis | 0.772 | 5.8783E-06 | 0.4342 | 1.4288E-06 | 1.7525 | 0.8014 |
| 1259 | Smoking/smokers in household | 0.7574 | 6.5065E-06 | 0.4942 | 2.7088E-09 | 1.4043 | 0.7525 |
| G6_NERPLEX | Nerve, nerve root and plexus disorders | 0.5411 | 6.5627E-06 | 0.4156 | 7.5676E-11 | 0.9234 | 0.6276 |
| 6153_1 | Medication for cholesterol, blood pressure, diabetes, or take exogenous hormones: Cholesterol lowering medication | 0.3956 | 8.1623E-06 | 0.2475 | 9.9091E-08 | 1.4788 | 0.6446 |
| 4653 | Ever highly irritable/argumentative for 2 days | 0.6579 | 9.0955E-06 | 0.3979 | 0.00001055 | 1.4974 | 0.7493 |
| 20003_1140864752 | Treatment/medication code: lansoprazole | 0.591 | 9.3908E-06 | 0.3337 | 3.6801E-06 | 1.6968 | 0.748 |
| 1548 | Variation in diet | 0.379 | 9.624E-06 | 0.3084 | 4.244E-11 | 0.7237 | 0.5726 |
| 6160_2 | Leisure/social activities: Pub or social club | 0.494 | 1.0066E-05 | 0.2891 | 1.7466E-06 | 1.6107 | 0.697 |
| 2734 | Number of live births | 0.3678 | 1.0364E-05 | 0.2941 | 5.1328E-09 | 0.7567 | 0.5755 |
| 20002_1286 | Non-cancer illness code, self-reported: depression | 0.5929 | 1.1471E-05 | 0.2326 | 0.002 | 2.3288 | 0.8257 |
| 5556 | FI9 : concept interpolation | -0.779 | 1.1925E-05 | -0.4358 | 5.9748E-07 | -1.7319 | 0.1858 |
| 1598 | Average weekly spirits intake | 0.471 | 1.2554E-05 | 0.3237 | 7.5861E-08 | 1.193 | 0.6486 |
| 4990 | FI6 : conditional arithmetic | -0.5584 | 0.00001299 | -0.3186 | 4.0633E-07 | -1.6803 | 0.2669 |
| 20502 | Ever had period extreme irritability | 0.543 | 1.3654E-05 | 0.1781 | 0.0139 | 2.5291 | 0.8292 |
| 23121 | Arm fat-free mass (right) | 0.2393 | 1.4854E-05 | 0.1763 | 1.0911E-09 | 1.0111 | 0.5658 |
| XIX_INJURY_POISON | Injury, poisoning and certain other consequences of external causes | 0.6165 | 1.5121E-05 | 0.3395 | 1.3194E-05 | 1.7056 | 0.8245 |
| G56 | Diagnoses - main ICD10: G56 Mononeuropathies of upper limb | 0.492 | 1.5988E-05 | 0.3279 | 3.0055E-08 | 1.2775 | 0.665 |
| 20524 | Sexual interference by partner or ex-partner without consent as an adult | 0.6994 | 0.00001619 | 0.2459 | 0.0014 | 2.527 | 0.8812 |
| 23122 | Arm predicted mass (right) | 0.2355 | 1.7003E-05 | 0.1756 | 1.3261E-09 | 0.9675 | 0.5625 |
| 971 | Frequency of walking for pleasure in last 4 weeks | 0.443 | 1.7566E-05 | 0.3285 | 1.3133E-08 | 0.968 | 0.6168 |
| M13_OTHERJOINT | #Other joint disorders | 0.4464 | 1.8196E-05 | 0.4346 | 8.4671E-11 | 0.0953 | 0.5111 |
| 20077 | Number of diet questionnaires completed | -0.6037 | 1.9557E-05 | -0.4299 | 4.1035E-08 | -1.075 | 0.325 |
| 4825 | Noisy workplace | 0.7019 | 1.9869E-05 | 0.2792 | 0.0014 | 2.2686 | 0.863 |
| 2443 | Diabetes diagnosed by doctor | 0.3999 | 2.0374E-05 | 0.2213 | 1.1375E-05 | 1.6773 | 0.678 |
| 6138_4 | Qualifications: CSEs or equivalent | 0.5336 | 2.1448E-05 | 0.2224 | 0.0003 | 2.2246 | 0.7788 |

| | | | | | | | |
|---|---|---|---|---|---|---|---|
| 20003_1140864992 | Treatment/medication code: tramadol | 0.7126 | 2.8794E-05 | 0.4141 | 6.3211E-06 | 1.5426 | 0.7809 |
| 1448_3 | Bread type: Wholemeal or wholegrain | -0.3752 | 3.1302E-05 | -0.292 | 6.5931E-09 | -0.8063 | 0.4141 |
| 6138_5 | Qualifications: NVQ or HND or HNC or equivalent | 0.6509 | 3.1373E-05 | 0.4753 | 2.111E-09 | 1.0016 | 0.6676 |
| 20497 | Repeated disturbing thoughts of stressful experience in past month | 0.5107 | 3.2241E-05 | 0.2042 | 0.0059 | 2.135 | 0.7879 |
| 4598 | Ever depressed for a whole week | 0.5311 | 3.5016E-05 | 0.2043 | 0.004 | 2.2266 | 0.8008 |
| 20544_11 | Mental health problems ever diagnosed by a professional: Depression | 0.4561 | 0.00003827 | 0.1559 | 0.0149 | 2.3461 | 0.7845 |
| 20126_0 | Bipolar and major depression status: No Bipolar or Depression | -0.6195 | 3.8786E-05 | -0.2163 | 0.0092 | -2.3441 | 0.1452 |
| 4957 | FI3 : word interpolation | -0.4939 | 0.00003949 | -0.5036 | 7.7859E-17 | 0.0721 | 0.51 |
| K11_HERNIA | Hernia | 0.8305 | 4.0221E-05 | 0.437 | 9.2971E-06 | 1.7485 | 0.8402 |
| 1359 | Poultry intake | 0.4206 | 4.2099E-05 | 0.3293 | 6.238E-10 | 0.7891 | 0.5936 |
| 20002_1113 | Non-cancer illness code, self-reported: emphysema/chronic bronchitis | 0.8629 | 4.3768E-05 | 0.4148 | 6.0186E-05 | 1.9056 | 0.8782 |
| 6160_1 | Leisure/social activities: Sports club or gym | -0.4025 | 5.1582E-05 | -0.1152 | 0.0337 | -2.5365 | 0.2349 |
| 4581 | Financial situation satisfaction | 0.512 | 5.3746E-05 | 0.4622 | 2.5878E-11 | 0.3446 | 0.5512 |
| 20498 | Felt very upset when reminded of stressful experience in past month | 0.4501 | 5.8881E-05 | 0.2442 | 2.4311E-05 | 1.6331 | 0.7043 |
| 20548_2 | Manifestations of mania or irritability: I was more restless than usual | 0.6397 | 6.1588E-05 | 0.295 | 0.0028 | 1.835 | 0.8164 |
| 20111_9 | Illnesses of siblings: Diabetes | 0.5762 | 6.4164E-05 | 0.3195 | 5.0432E-05 | 1.5621 | 0.7487 |
| 6164_4 | Types of physical activity in last 4 weeks: Light DIY (eg: pruning, watering the lawn) | -0.3574 | 6.7883E-05 | -0.3897 | 4.7181E-15 | 0.315 | 0.533 |
| 3591 | Ever had hysterectomy (womb removed) | 0.4696 | 0.00007343 | 0.3684 | 3.4229E-08 | 0.7444 | 0.6031 |
| M13_JOINTOTH | Other specific joint derangements/joint disorders | 0.8353 | 0.00007683 | 0.5752 | 5.3046E-06 | 1.057 | 0.7307 |
| 6150_100 | Vascular/heart problems diagnosed by doctor: None of the above | -0.2791 | 7.6913E-05 | -0.088 | 0.0363 | -2.3263 | 0.3158 |
| 2644 | Light smokers, at least 100 smokes in lifetime | 0.4979 | 7.7405E-05 | 0.4524 | 6.9052E-12 | 0.3199 | 0.5468 |
| 20002_1111 | Non-cancer illness code, self-reported: asthma | 0.2908 | 8.1684E-05 | 0.1879 | 5.3561E-05 | 1.1797 | 0.6055 |
| 6154_1 | Medication for pain relief, constipation, heartburn: Aspirin | 0.4764 | 8.6306E-05 | 0.2545 | 0.0008 | 1.5508 | 0.7106 |
| I9_IHD | Ischaemic heart disease, wide definition | 0.4295 | 8.8191E-05 | 0.2677 | 0.0002 | 1.2346 | 0.6624 |
| K44 | Diagnoses - main ICD10: K44 Diaphragmatic hernia | 0.8331 | 9.1117E-05 | 0.4721 | 1.1525E-05 | 1.5133 | 0.8174 |
| 20514 | Recent lack of interest or pleasure in doing things | 0.5532 | 9.4916E-05 | 0.2329 | 0.0044 | 1.9576 | 0.7978 |
| 1873 | Number of full brothers | 0.5992 | 9.6407E-05 | 0.3348 | 7.6337E-06 | 1.5468 | 0.7535 |
| 23102 | Whole body water mass | 0.2093 | 9.9294E-05 | 0.1547 | 9.8598E-08 | 0.8934 | 0.5575 |
| 1130 | Hands-free device/speakerphone use with mobile phone in last 3 month | 0.4954 | 0.0001 | 0.6921 | 5.6138E-28 | -1.3697 | 0.305 |
| 1319 | Dried fruit intake | -0.3213 | 0.0001 | -0.2863 | 1.7523E-11 | -0.3744 | 0.4638 |
| 23118 | Leg predicted mass (left) | 0.2065 | 0.0001 | 0.1585 | 1.2536E-07 | 0.7737 | 0.5503 |
| 23101 | Whole body fat-free mass | 0.2061 | 0.0001 | 0.1537 | 1.3102E-07 | 0.8579 | 0.5552 |
| 23117 | Leg fat-free mass (left) | 0.2059 | 0.0001 | 0.1573 | 1.767E-07 | 0.7839 | 0.551 |
| G6_CARPTU | Carpal tunnel syndrome | 0.4352 | 0.0001 | 0.3047 | 2.591E-07 | 1.0135 | 0.6324 |
| 2834 | Bilateral oophorectomy (both ovaries removed) | 0.4678 | 0.0001 | 0.3252 | 6.8669E-07 | 1.0214 | 0.6437 |
| 20002_1074 | Non-cancer illness code, self-reported: angina | 0.8113 | 0.0001 | 0.4396 | 9.0434E-06 | 1.5862 | 0.8331 |
| 20003_1140868226 | Treatment/medication code: aspirin | 0.4612 | 0.0001 | 0.2608 | 0.0008 | 1.3939 | 0.6989 |
| 6153_4 | Medication for cholesterol, blood pressure, diabetes, or take exogenous hormones: Hormone replacement therapy | 0.5309 | 0.0002 | 0.4855 | 2.4865E-08 | 0.273 | 0.5452 |
| 23114 | Leg predicted mass (right) | 0.205 | 0.0002 | 0.1564 | 1.427E-07 | 0.7841 | 0.5511 |
| 23113 | Leg fat-free mass (right) | 0.2061 | 0.0002 | 0.1555 | 1.587E-07 | 0.8152 | 0.5532 |
| 23107 | Impedance of leg (right) | -0.2211 | 0.0002 | -0.1724 | 2.1442E-07 | -0.7157 | 0.4488 |
| 699 | Length of time at current address | -0.4247 | 0.0002 | -0.2736 | 2.6641E-07 | -1.1857 | 0.3488 |
| 1060 | Time spent outdoors in winter | 0.3691 | 0.0002 | 0.2792 | 3.1887E-07 | 0.7909 | 0.5922 |
| 20111_100 | Illnesses of siblings: None of the above (group 1) | -0.3907 | 0.0002 | -0.2842 | 3.9705E-07 | -0.8999 | 0.3907 |
| 826 | Job involves shift work | 0.5964 | 0.0002 | 0.4669 | 9.5008E-07 | 0.696 | 0.6316 |
| M25 | Diagnoses - main ICD10: M25 Other joint disorders, not elsewhere classified | 0.7941 | 0.0002 | 0.5939 | 4.6144E-06 | 0.8032 | 0.6805 |

| | | | | | | | |
|---|---|---|---|---|---|---|---|
| 20490 | Sexually molested as a child | 0.7295 | 0.0002 | 0.4715 | 5.7365E-06 | 1.1752 | 0.7493 |
| 20111_6 | Illnesses of siblings: Chronic bronchitis/emphysema | 0.7513 | 0.0002 | 0.4797 | 7.1366E-06 | 1.1854 | 0.7609 |
| 6150_2 | Vascular/heart problems diagnosed by doctor: Angina | 0.8281 | 0.0002 | 0.4417 | 3.1481E-05 | 1.5584 | 0.834 |
| 6152_8 | Blood clot, DVT, bronchitis, emphysema, asthma, rhinitis, eczema, allergy diagnosed by doctor: Asthma | 0.2767 | 0.0002 | 0.1892 | 0.00005356 | 1.0032 | 0.5868 |
| 6160_5 | Leisure/social activities: Other group activity | -0.4425 | 0.0002 | -0.2414 | 8.6455E-05 | -1.5194 | 0.3056 |
| 20111_1 | Illnesses of siblings: Heart disease | 0.5972 | 0.0002 | 0.3196 | 0.0001 | 1.5169 | 0.7652 |
| 20516 | Recent restlessness | 0.6157 | 0.0002 | 0.289 | 0.0003 | 1.7997 | 0.8024 |
| R51 | Diagnoses - main ICD10: R51 Headache | 1 | 0.0002 | 0.5761 | 0.0014 | 1.2577 | 0.882 |
| 4979 | FI5 : family relationship calculation | -0.6093 | 0.0002 | -0.2478 | 0.0044 | -1.9688 | 0.1743 |
| 6160_4 | Leisure/social activities: Adult education class | -0.4732 | 0.0003 | -0.3886 | 1.6763E-09 | -0.5758 | 0.4155 |
| 30280 | Immature reticulocyte fraction | 0.2525 | 0.0003 | 0.1925 | 3.8338E-07 | 0.7571 | 0.5608 |
| 2887 | Number of cigarettes previously smoked daily | 0.409 | 0.0003 | 0.2908 | 1.0402E-05 | 0.8991 | 0.6199 |
| 20002_1220 | Non-cancer illness code, self-reported: diabetes | 0.3602 | 0.0003 | 0.254 | 2.0576E-05 | 0.9082 | 0.6089 |
| 6159_5 | Pain type(s) experienced in last month: Stomach or abdominal pain | 0.5073 | 0.0003 | 0.3175 | 2.4918E-05 | 1.1822 | 0.6835 |
| 20531 | Victim of sexual assault | 0.5387 | 0.0003 | 0.3075 | 8.2545E-05 | 1.3867 | 0.7266 |
| 20416 | Frequency of consuming six or more units of alcohol | 0.4771 | 0.0003 | 0.2513 | 0.0018 | 1.4702 | 0.7223 |
| K21 | Diagnoses - main ICD10: K21 Gastro-oesophageal reflux disease | 0.6985 | 0.0003 | 0.3201 | 0.0019 | 1.73 | 0.83 |
| 20002_1201 | Non-cancer illness code, self-reported: bladder problem (not cancer) | 0.827 | 0.0003 | 0.4375 | 0.002 | 1.447 | 0.8445 |
| 20153 | Forced expiratory volume in 1-second (FEV1), predicted | -0.1995 | 0.0003 | -0.0943 | 0.0036 | -1.6436 | 0.3912 |
| 1950 | Sensitivity / hurt feelings | 0.3079 | 0.0003 | 0.1408 | 0.0038 | 1.7149 | 0.6675 |
| 30300 | High light scatter reticulocyte count | 0.2101 | 0.0004 | 0.179 | 2.9295E-09 | 0.4648 | 0.5315 |
| 1468_5 | Cereal type: Other (e.g. Cornflakes, Frosties) | 0.3383 | 0.0004 | 0.3008 | 1.9734E-06 | 0.3266 | 0.539 |
| 1883 | Number of full sisters | 0.5659 | 0.0004 | 0.3275 | 1.6892E-05 | 1.3421 | 0.7318 |
| 4290 | Duration screen displayed | 0.3234 | 0.0004 | 0.1941 | 0.0003 | 1.223 | 0.6269 |
| 6152_100 | Blood clot, DVT, bronchitis, emphysema, asthma, rhinitis, eczema, allergy diagnosed by doctor: None of the above | -0.2446 | 0.0004 | -0.1415 | 0.0004 | -1.2873 | 0.398 |
| 20441 | Ever had prolonged loss of interest in normal activities | 0.4208 | 0.0004 | 0.2104 | 0.0007 | 1.58 | 0.7083 |
| 3571 | Back pain for 3+ months | 0.7833 | 0.0004 | 0.4008 | 0.0008 | 1.5191 | 0.8384 |
| J44 | Diagnoses - main ICD10: J44 Other chronic obstructive pulmonary disease | 0.9019 | 0.0004 | 0.3576 | 0.002 | 1.9501 | 0.9167 |
| COPD_EARLYANDLATER | COPD, early/later onset | 0.9238 | 0.0004 | 0.3479 | 0.0029 | 2.0016 | 0.9343 |
| 1588 | Average weekly beer plus cider intake | 0.4411 | 0.0004 | 0.1541 | 0.0228 | 2.0189 | 0.7714 |
| 1031 | Frequency of friend/family visits | -0.2587 | 0.0005 | -0.3872 | 2.1028E-19 | 1.5014 | 0.6306 |
| 1538_1 | Major dietary changes in the last 5 years: Yes, because of illness | 0.4407 | 0.0005 | 0.4074 | 5.8402E-12 | 0.2389 | 0.5346 |
| 23129 | Trunk fat-free mass | 0.1877 | 0.0005 | 0.1367 | 2.1252E-06 | 0.8382 | 0.553 |
| 23130 | Trunk predicted mass | 0.1877 | 0.0005 | 0.1363 | 2.443E-06 | 0.8429 | 0.5534 |
| 30510 | Creatinine (enzymatic) in urine | 0.238 | 0.0005 | 0.1936 | 3.7452E-06 | 0.5553 | 0.5449 |
| 3849 | Number of pregnancy terminations | 0.5039 | 0.0005 | 0.211 | 0.0021 | 1.8239 | 0.7755 |
| 20512 | Recent feelings of foreboding | 0.4428 | 0.0005 | 0.1761 | 0.0263 | 1.7732 | 0.7563 |
| 20002_1065 | Non-cancer illness code, self-reported: hypertension | 0.2438 | 0.0005 | 0.0832 | 0.0476 | 1.9591 | 0.662 |
| 1478 | Salt added to food | 0.2594 | 0.0006 | 0.2271 | 2.1486E-10 | 0.3866 | 0.5334 |
| 20002_1473 | Non-cancer illness code, self-reported: high cholesterol | 0.3369 | 0.0006 | 0.26 | 3.5939E-07 | 0.6952 | 0.5793 |
| M17 | Diagnoses - main ICD10: M17 Gonarthrosis [arthrosis of knee] | 0.4645 | 0.0006 | 0.3151 | 4.2961E-07 | 1.003 | 0.6381 |
| KNEE_ARTHROSIS | Gonarthrosis [arthrosis of knee](FG) | 0.4645 | 0.0006 | 0.3151 | 4.2961E-07 | 1.003 | 0.6452 |
| 20003_1140861998 | Treatment/medication code: ventolin 100micrograms inhaler | 0.503 | 0.0006 | 0.2317 | 0.0017 | 1.644 | 0.7594 |
| K11_GALLBILPANC | Disorders of gallbladder, biliary tract and pancreas | 0.5062 | 0.0006 | 0.2097 | 0.003 | 1.8167 | 0.7737 |

| | | | | | | | |
|---|---|---|---|---|---|---|---|
| 1210 | Snoring | -0.234 | 0.0006 | -0.1143 | 0.0094 | -1.481 | 0.3782 |
| 4836 | Loud music exposure frequency | 0.6257 | 0.0006 | 0.2409 | 0.0107 | 1.8712 | 0.8404 |
| 20510 | Recent feelings of depression | 0.4578 | 0.0006 | 0.1697 | 0.0156 | 1.9179 | 0.7734 |
| 6162_2 | Types of transport used (excluding work): Walk | -0.3563 | 0.0007 | -0.473 | 1.1448E-18 | 0.9869 | 0.6161 |
| 1538_2 | Major dietary changes in the last 5 years: Yes, because of other reasons | 0.3971 | 0.0007 | 0.3136 | 2.7793E-08 | 0.6445 | 0.5862 |
| 5012 | FI8 : chained arithmetic | -0.4289 | 0.0007 | -0.2595 | 6.6961E-05 | -1.1937 | 0.33 |
| 30000 | White blood cell (leukocyte) count | 0.1909 | 0.0007 | 0.124 | 8.4322E-05 | 1.0342 | 0.5691 |
| 1628 | Alcohol intake versus 10 years previously | 0.3255 | 0.0007 | 0.2142 | 0.0003 | 0.9855 | 0.6136 |
| 30290 | High light scatter reticulocyte percentage | 0.1981 | 0.0008 | 0.1694 | 2.3887E-08 | 0.4324 | 0.5291 |
| 20500 | Ever suffered mental distress preventing usual activities | 0.3977 | 0.0008 | 0.2432 | 4.8913E-05 | 1.1652 | 0.6564 |
| I9_DISVEINLYMPH | Diseases of veins, lymphatic vessels and lymph nodes, not elsewhere classified | 0.3917 | 0.0008 | 0.1751 | 0.0099 | 1.609 | 0.7123 |
| 1110 | Length of mobile phone use | 0.3016 | 0.0009 | 0.6445 | 1.5001E-53 | -3.4242 | 0.187 |
| 20001_1041 | Cancer code, self-reported: cervical cancer | 0.6679 | 0.0009 | 0.5865 | 1.331E-06 | 0.3476 | 0.5839 |
| 6147_1 | Reason for glasses/contact lenses: For short-sightedness, i.e. only or mainly for distance viewing such as driving, cinema etc (called 'myopia') | -0.366 | 0.0009 | -0.2503 | 7.6765E-05 | -0.9079 | 0.3855 |
| 3731 | Former alcohol drinker | 0.4893 | 0.0009 | 0.2036 | 0.0149 | 1.6851 | 0.7699 |
| 23108 | Impedance of leg (left) | -0.1947 | 0.001 | -0.1587 | 2.2598E-06 | -0.5295 | 0.4621 |
| 20002_1138 | Non-cancer illness code, self-reported: gastro-oesophageal reflux (gord) / gastric reflux | 0.542 | 0.001 | 0.3884 | 0.0000118 | 0.8223 | 0.6552 |
| K80 | Diagnoses - main ICD10: K80 Cholelithiasis | 0.4535 | 0.001 | 0.186 | 0.0086 | 1.7217 | 0.7485 |
| 20507 | Recent feelings of inadequacy | 0.54 | 0.001 | 0.1706 | 0.0266 | 2.0384 | 0.832 |
| 6143_1 | Transport type for commuting to job workplace: Car/motor vehicle | 0.5209 | 0.0011 | 0.5264 | 3.2114E-08 | -0.0296 | 0.4946 |
| I20 | Diagnoses - main ICD10: I20 Angina pectoris | 0.5897 | 0.0011 | 0.3651 | 0.0024 | 1.0345 | 0.7226 |
| 3143 | Ankle spacing width | 0.2099 | 0.0012 | 0.197 | 1.4165E-09 | 0.1777 | 0.513 |
| M13_ARTHROSIS | #Arthrosis | 0.3646 | 0.0012 | 0.3066 | 1.8037E-07 | 0.4568 | 0.5557 |
| 1767 | Adopted as a child | 1 | 0.0012 | 0.558 | 5.9318E-05 | 1.2629 | 0.874 |
| 20548_3 | Manifestations of mania or irritability: My thoughts were racing | 0.5138 | 0.0012 | 0.341 | 0.0005 | 0.9255 | 0.6744 |
| 3606 | Chest pain or discomfort walking normally | 1 | 0.0014 | 0.5866 | 9.0018E-06 | 1.1465 | 0.8572 |
| 2217 | Age started wearing glasses or contact lenses | 0.2662 | 0.0014 | 0.1155 | 0.0116 | 1.5868 | 0.6531 |
| 3526 | Mother's age at death | -0.4876 | 0.0015 | -0.4679 | 3.3784E-07 | -0.1099 | 0.4801 |
| 4119 | Ankle spacing width (right) | 0.2282 | 0.0015 | 0.1882 | 1.9097E-06 | 0.4866 | 0.54 |
| M23 | Diagnoses - main ICD10: M23 Internal derangement of knee | 0.5715 | 0.0016 | 0.4438 | 1.0261E-05 | 0.6162 | 0.6173 |
| 6143_4 | Transport type for commuting to job workplace: Cycle | -0.4485 | 0.0016 | -0.3316 | 0.0001 | -0.7045 | 0.3858 |
| M13_SPONDYLOPATHY | #Spondylopathies | 1 | 0.0017 | 0.3745 | 0.0088 | 1.6369 | 0.9302 |
| 20002_1075 | Non-cancer illness code, self-reported: heart attack/myocardial infarction | 0.5532 | 0.0018 | 0.2741 | 0.0055 | 1.3745 | 0.7661 |
| 816 | Job involves heavy manual or physical work | 0.3233 | 0.0019 | 0.2799 | 2.9175E-08 | 0.376 | 0.5448 |
| 4291 | Number of attempts | 0.3803 | 0.0019 | 0.3308 | 1.2881E-07 | 0.3603 | 0.5509 |
| 806 | Job involves mainly walking or standing | 0.3205 | 0.0019 | 0.2686 | 1.7081E-07 | 0.4505 | 0.5536 |
| 50 | Standing height | -0.1316 | 0.002 | -0.1033 | 3.0887E-05 | -0.5741 | 0.4719 |
| 6150_1 | Vascular/heart problems diagnosed by doctor: Heart attack | 0.5462 | 0.002 | 0.2293 | 0.0188 | 1.5692 | 0.7868 |
| G6_EPIPAROX | Episodal and paroxysmal disorders | 1 | 0.002 | 0.3305 | 0.0474 | 1.6405 | 0.9586 |
| 20536_1 | Weight change during worst episode of depression: Gained weight | 0.4498 | 0.0021 | 0.2467 | 0.0007 | 1.2402 | 0.7029 |
| 20003_1140884600 | Treatment/medication code: metformin | 0.3455 | 0.0023 | 0.2238 | 0.0001 | 0.9554 | 0.6246 |
| 6141_6 | How are people in household related to participant: Grandchild | 0.8007 | 0.0023 | 0.4336 | 0.0012 | 1.2448 | 0.8176 |
| 20003_1141192736 | Treatment/medication code: ezetimibe | 0.6124 | 0.0023 | 0.27 | 0.0128 | 1.4973 | 0.8135 |
| I83 | Diagnoses - main ICD10: I83 Varicose veins of lower extremities | 0.3399 | 0.0024 | 0.1492 | 0.0319 | 1.4449 | 0.6902 |

| | | | | | | | |
|---|---|---|---|---|---|---|---|
| 20002_1094 | Non-cancer illness code, self-reported: deep venous thrombosis (dvt) | 0.4632 | 0.0025 | 0.4676 | 1.3263E-06 | -0.0243 | 0.4954 |
| 6152_5 | Blood clot, DVT, bronchitis, emphysema, asthma, rhinitis, eczema, allergy diagnosed by doctor: Blood clot in the leg (DVT) | 0.4509 | 0.0026 | 0.464 | 5.7274E-07 | -0.0743 | 0.4869 |
| I25 | Diagnoses - main ICD10: I25 Chronic ischaemic heart disease | 0.3773 | 0.0027 | 0.2054 | 0.0037 | 1.1912 | 0.6741 |
| 20003_1140926606 | Treatment/medication code: salbutamol 100micrograms spacehaler | 0.6053 | 0.0027 | 0.2806 | 0.0073 | 1.4288 | 0.8018 |
| 5084 | Spherical power (right) | 0.2442 | 0.0029 | 0.1716 | 9.1285E-05 | 0.7813 | 0.5718 |
| 20003_1140879616 | Treatment/medication code: amitriptyline | 0.6047 | 0.0029 | 0.3954 | 0.0005 | 0.9004 | 0.7071 |
| 6159_1 | Pain type(s) experienced in last month: Headache | 0.2726 | 0.0032 | 0.2315 | 1.6184E-05 | 0.3849 | 0.5411 |
| 3090 | Used an inhaler for chest within last hour | 0.5813 | 0.0032 | 0.2465 | 0.0435 | 1.4424 | 0.8064 |
| 20446 | Ever had prolonged feelings of sadness or depression | 0.3443 | 0.0032 | 0.1252 | 0.0483 | 1.6486 | 0.716 |
| 30140 | Neutrophill count | 0.1734 | 0.0033 | 0.0842 | 0.0068 | 1.3357 | 0.5907 |
| 22606_2 | Workplace very noisy: Often | 0.5673 | 0.0034 | 0.5173 | 5.5398E-08 | 0.2315 | 0.5522 |
| 20003_1140884488 | Treatment/medication code: diclofenac | 0.485 | 0.0034 | 0.3003 | 0.0009 | 0.9801 | 0.6851 |
| 20018 | Prospective memory result | 0.3675 | 0.0036 | 0.3219 | 5.3768E-07 | 0.3223 | 0.5474 |
| Z46 | Diagnoses - main ICD10: Z46 Fitting and adjustment of other devices | 0.5946 | 0.0036 | 0.2177 | 0.0404 | 1.6394 | 0.9426 |
| 20518 | Recent changes in speed/amount of moving or speaking | 0.602 | 0.0037 | 0.275 | 0.0269 | 1.3536 | 0.8026 |
| 981 | Duration walking for pleasure | -0.3463 | 0.0039 | -0.3689 | 1.1087E-09 | 0.1683 | 0.5234 |
| 20003_1140861958 | Treatment/medication code: simvastatin | 0.304 | 0.0041 | 0.2188 | 6.4113E-05 | 0.7153 | 0.5875 |
| 20126_3 | Bipolar and major depression status: Probable Recurrent major depression (severe) | 0.905 | 0.0041 | 0.32 | 0.0224 | 1.6967 | 0.9377 |
| 6149_4 | Mouth/teeth dental problems: Loose teeth | 0.4292 | 0.0046 | 0.2028 | 0.0081 | 1.335 | 0.7154 |
| 104400 | Fruit consumers | -0.6092 | 0.0047 | -0.433 | 0.0001 | -0.7225 | 0.3234 |
| M13_MENISCUSDERANGEMENTS | Meniscus derangement | 0.5766 | 0.0047 | 0.4239 | 0.0002 | 0.6561 | 0.6413 |
| N92 | Diagnoses - main ICD10: N92 Excessive, frequent and irregular menstruation | 0.4524 | 0.005 | 0.3762 | 2.0129E-05 | 0.4145 | 0.5719 |
| 6033 | Maximum heart rate during fitness test | -0.3741 | 0.0051 | -0.1876 | 0.0122 | -1.2174 | 0.326 |
| 20003_1140874420 | Treatment/medication code: quinine | 0.5727 | 0.0052 | 0.4891 | 6.2436E-06 | 0.3609 | 0.5861 |
| 1608 | Average weekly fortified wine intake | -0.5422 | 0.0052 | -0.3938 | 0.00009014 | -0.6785 | 0.3504 |
| 20517 | Trouble falling or staying asleep, or sleeping too much | 0.3645 | 0.0053 | 0.2339 | 0.001 | 0.877 | 0.6331 |
| M13_ROTATORCUFF | Rotator cuff syndrome | 0.6956 | 0.0054 | 0.3951 | 0.0023 | 1.0682 | 0.7598 |
| 20548_7 | Manifestations of mania or irritability: I was easily distracted | 0.4943 | 0.0054 | 0.2891 | 0.0056 | 0.9955 | 0.7043 |
| 6142_6 | Current employment status: Doing unpaid or voluntary work | -0.462 | 0.0057 | -0.3999 | 0.00002382 | -0.3231 | 0.4389 |
| 20002_1112 | Non-cancer illness code, self-reported: chronic obstructive airways disease/copd | 1 | 0.0058 | 0.4635 | 0.0074 | 1.2324 | 0.9184 |
| 1309 | Fresh fruit intake | -0.2259 | 0.0059 | -0.1093 | 0.0199 | -1.2332 | 0.3812 |
| 6159_2 | Pain type(s) experienced in last month: Facial pain | 0.582 | 0.0059 | 0.2601 | 0.0304 | 1.323 | 0.7913 |
| 20540 | Multiple worries during worst period of anxiety | 0.8114 | 0.006 | 0.3182 | 0.0201 | 1.5153 | 0.9005 |
| 3063 | Forced expiratory volume in 1-second (FEV1) | -0.159 | 0.0062 | -0.1533 | 4.2963E-06 | -0.0851 | 0.4942 |
| 30250 | Reticulocyte count | 0.1637 | 0.0064 | 0.1546 | 7.3043E-08 | 0.1368 | 0.5093 |
| 41215_0 | Detention categories: Informal, not formally detained | 0.9347 | 0.0066 | 0.5652 | 0.0016 | 0.9526 | 0.8228 |
| 6162_3 | Types of transport used (excluding work): Public transport | -0.3458 | 0.0068 | -0.2445 | 0.00007089 | -0.7142 | 0.3987 |
| I9_CORATHER | Coronary atherosclerosis | 0.3281 | 0.0068 | 0.2295 | 0.0007 | 0.7099 | 0.6008 |
| 22620_1 | Job involved shift work: Yes | 0.4681 | 0.0069 | 0.2836 | 0.0028 | 0.9338 | 0.6869 |
| 20530 | Witnessed sudden violent death | 0.5948 | 0.007 | 0.2518 | 0.0317 | 1.3731 | 0.8143 |
| 20002_1311 | Non-cancer illness code, self-reported: spine arthritis/spondylitis | 0.9793 | 0.0071 | 0.5044 | 0.0038 | 1.1773 | 0.8916 |

| | | | | | | | |
|---|---|---|---|---|---|---|---|
| 1807 | Father's age at death | -0.2846 | 0.0072 | -0.3442 | 1.6267E-06 | 0.4658 | 0.5617 |
| 1807 | Father's age at death | -0.2846 | 0.0072 | -0.3442 | 1.6267E-06 | 0.4658 | 0.5617 |
| 20151 | Forced vital capacity (FVC), Best measure | -0.1575 | 0.0072 | -0.1501 | 6.4982E-06 | -0.1098 | 0.4922 |
| 20547_1 | Activities undertaken to treat depression: Talking therapies, such as psychotherapy, counselling, group therapy or CBT | 0.3434 | 0.0073 | 0.2087 | 0.0069 | 0.9008 | 0.6379 |
| IV_ENDOCRIN_NUTRIT | Endocrine, nutritional and metabolic diseases | 0.8666 | 0.0074 | 0.4442 | 0.0093 | 1.1548 | 0.8607 |
| 20150 | Forced expiratory volume in 1-second (FEV1), Best measure | -0.1586 | 0.0075 | -0.1397 | 5.4614E-05 | -0.2753 | 0.4802 |
| 6141_1 | How are people in household related to participant: Husband, wife or partner | -0.3096 | 0.0076 | -0.1528 | 0.0145 | -1.1892 | 0.3494 |
| 1160 | Sleep duration | -0.2215 | 0.0077 | -0.2147 | 1.002E-06 | -0.0724 | 0.493 |
| 20401 | Ever addicted to any substance or behaviour | 0.9466 | 0.0081 | 0.4351 | 0.0157 | 1.2776 | 0.9093 |
| 670_1 | Type of accommodation lived in: A house or bungalow | -0.3628 | 0.0081 | -0.1696 | 0.0438 | -1.2008 | 0.3106 |
| I9_UAP | Unstable angina pectoris | 0.5599 | 0.0082 | 0.3414 | 0.0201 | 0.8479 | 0.7134 |
| 4100 | Ankle spacing width (left) | 0.1805 | 0.0086 | 0.1894 | 2.3949E-06 | -0.1119 | 0.491 |
| 20525 | Able to pay rent/mortgage as an adult | -0.5015 | 0.0089 | -0.2119 | 0.0226 | -1.3589 | 0.2253 |
| 6162_4 | Types of transport used (excluding work): Cycle | -0.2843 | 0.0095 | -0.3458 | 3.2352E-09 | 0.4952 | 0.562 |
| 20007 | Interpolated Age of participant when cancer first diagnosed | -0.6617 | 0.0098 | -0.7278 | 0.0271 | 0.1584 | 0.5682 |
| 20003_1140884444 | Treatment/medication code: codeine | 0.7384 | 0.01 | 0.4303 | 0.0099 | 0.9288 | 0.7892 |
| 3062 | Forced vital capacity (FVC) | -0.1488 | 0.0102 | -0.158 | 5.3727E-07 | 0.1396 | 0.5093 |
| 20110_3 | Illnesses of mother: Lung cancer | 0.4781 | 0.0103 | 0.2614 | 0.0343 | 0.9699 | 0.7139 |
| 30240 | Reticulocyte percentage | 0.1525 | 0.0105 | 0.1425 | 9.4744E-07 | 0.1508 | 0.5102 |
| 777 | Frequency of travelling from home to job workplace | 0.4198 | 0.0106 | 0.3726 | 6.546E-06 | 0.2568 | 0.5488 |
| M51 | Diagnoses - main ICD10: M51 Other intervertebral disk disorders | 0.8209 | 0.0106 | 0.3675 | 0.0029 | 1.317 | 0.8591 |
| 20543 | Number of things worried about during worst period of anxiety | 0.6879 | 0.0108 | 0.3726 | 0.0141 | 1.0182 | 0.7943 |
| 680_4 | Own or rent accommodation lived in: Rent - from private landlord or letting agency | 0.9252 | 0.0109 | 0.5537 | 0.0003 | 0.9415 | 0.83 |
| 5085 | Spherical power (left) | 0.2143 | 0.0112 | 0.1523 | 0.0007 | 0.6476 | 0.5613 |
| 6145_3 | Illness, injury, bereavement, stress in last 2 years: Death of a close relative | 0.5118 | 0.0113 | 0.4131 | 0.0001 | 0.4306 | 0.5972 |
| 1438 | Bread intake | -0.2019 | 0.0118 | -0.2357 | 1.3569E-08 | 0.3747 | 0.5352 |
| XXI_HEALTHFACTORS | Factors influencing health status and contact with health services | 0.3454 | 0.0118 | 0.3918 | 3.3306E-05 | -0.2788 | 0.432 |
| 1707_2 | Handedness (chirality/laterality): Left-handed | -0.3703 | 0.0122 | -0.1891 | 0.0242 | -1.0662 | 0.3182 |
| 1707_2 | Handedness (chirality/laterality): Left-handed | -0.3703 | 0.0122 | -0.1891 | 0.0242 | -1.0662 | 0.3183 |
| 5001 | FI7 : synonym | -0.3839 | 0.0127 | -0.2983 | 0.0008 | -0.4812 | 0.4121 |
| 22504 | Bring up phlegm/sputum/mucus on most days | 0.5739 | 0.0127 | 0.2487 | 0.0236 | 1.2744 | 0.8015 |
| E66 | Diagnoses - main ICD10: E66 Obesity | 0.6093 | 0.0131 | 0.3867 | 0.0009 | 0.8186 | 0.7171 |
| 20403 | Amount of alcohol drunk on a typical drinking day | 0.409 | 0.0136 | 0.2127 | 0.0169 | 1.0432 | 0.6961 |
| E4_OBESITY | Obesity | 0.6057 | 0.0137 | 0.3903 | 0.0007 | 0.7934 | 0.711 |
| 22609_1 | Workplace very dusty: Sometimes | 0.9257 | 0.0139 | 0.573 | 0.0213 | 0.7819 | 0.8227 |
| 6142_3 | Current employment status: Looking after home and/or family | -0.5134 | 0.0143 | -0.6872 | 2.5938E-06 | 0.6799 | 0.6662 |
| 1568 | Average weekly red wine intake | -0.2406 | 0.0145 | -0.1712 | 0.0003 | -0.6341 | 0.4287 |
| 20118_5 | Home area population density - urban or rural: England/Wales - Urban - less sparse | 0.774 | 0.0164 | 0.432 | 0.0099 | 0.9415 | 0.8148 |
| 6164_5 | Types of physical activity in last 4 weeks: Heavy DIY (eg: weeding, lawn mowing, carpentry, digging) | -0.2613 | 0.0169 | -0.3644 | 2.2951E-11 | 0.8435 | 0.6043 |
| E4_OBESITYNAS | Obesity, other/unspecified | 0.5501 | 0.0174 | 0.3913 | 0.0005 | 0.6174 | 0.6591 |
| 20154 | Forced expiratory volume in 1-second (FEV1), predicted percentage | -0.1706 | 0.0175 | -0.1025 | 0.0175 | -0.8132 | 0.4291 |
| 22609_2 | Workplace very dusty: Often | 0.6217 | 0.0185 | 0.564 | 0.0122 | 0.1663 | 0.5602 |
| 2227 | Other eye problems | 0.3316 | 0.0189 | 0.2208 | 0.0114 | 0.6671 | 0.6129 |
| 6143_2 | Transport type for commuting to job workplace: Walk | -0.385 | 0.0196 | -0.3951 | 0.0005 | 0.0503 | 0.51 |

| | | | | | | | |
|---|---|---|---|---|---|---|---|
| 22702 | Home location - east co-ordinate (rounded) | -0.5214 | 0.0199 | -0.2981 | 0.0075 | -0.8929 | 0.2767 |
| 5507 | Leg pain on walking : action taken | -0.5356 | 0.0207 | -0.5256 | 0.0009 | -0.0356 | 0.4896 |
| 20003_1141146234 | Treatment/medication code: atorvastatin | 0.3469 | 0.0209 | 0.2007 | 0.0227 | 0.8396 | 0.6486 |
| 100011 | Iron | -0.5907 | 0.021 | -0.4153 | 0.0093 | -0.5815 | 0.3232 |
| 1289 | Cooked vegetable intake | 0.1983 | 0.0214 | 0.1227 | 0.0202 | 0.7479 | 0.5777 |
| 20448 | Professional informed about depression | 0.4706 | 0.0214 | 0.2266 | 0.0491 | 1.0396 | 0.7375 |
| 22601_33113434 | Job coding: non-commissioned officers or other rank of armed forces | 0.6704 | 0.0218 | 0.4353 | 0.0143 | 0.6875 | 0.7316 |
| 20458 | General happiness | 0.237 | 0.0222 | -0.1507 | 0.0182 | 3.1865 | 0.8435 |
| 20118_11 | Home area population density - urban or rural: Scotland - Large Urban Area | -0.5071 | 0.0224 | -0.3987 | 0.0008 | -0.4299 | 0.3883 |
| 1418_6 | Milk type used: Never/rarely have milk | 0.2687 | 0.0226 | 0.1913 | 0.0118 | 0.5518 | 0.5801 |
| 22502 | Cough on most days | 0.3262 | 0.0227 | 0.1707 | 0.0413 | 0.9375 | 0.6573 |
| 46 | Hand grip strength (left) | -0.1424 | 0.0229 | -0.1668 | 4.2331E-06 | 0.3372 | 0.5243 |
| M13_ARTHROSIS_OTH | Other arthrosis | 0.4639 | 0.023 | 0.4886 | 0.0001 | -0.1027 | 0.4763 |
| 6145_5 | Illness, injury, bereavement, stress in last 2 years: Marital separation/divorce | 0.923 | 0.023 | 0.4268 | 0.0054 | 1.1434 | 0.8917 |
| 22617_3311 | Job SOC coding: NCOs and other ranks | 0.6888 | 0.0232 | 0.4379 | 0.0203 | 0.7022 | 0.7463 |
| 100025 | Vitamin E | -0.4987 | 0.0235 | -0.3936 | 0.0039 | -0.4058 | 0.3914 |
| 1418_2 | Milk type used: Semi-skimmed | -0.2633 | 0.0242 | -0.1281 | 0.0377 | -1.0239 | 0.3619 |
| 20417 | Tense, sore, or aching muscles during worst period of anxiety | 0.4544 | 0.0247 | 0.5484 | 5.6435E-05 | -0.3854 | 0.403 |
| 20549_3 | Substances taken for anxiety: Medication prescribed to you (for at least two weeks) | 0.3527 | 0.0257 | 0.2031 | 0.0423 | 0.7997 | 0.6522 |
| 20074 | Home location at assessment - east co-ordinate (rounded) | -0.4458 | 0.0268 | -0.2927 | 0.0089 | -0.6649 | 0.3452 |
| 6154_4 | Medication for pain relief, constipation, heartburn: Ranitidine (e.g. Zantac) | 0.6206 | 0.0268 | 0.3795 | 0.0153 | 0.7511 | 0.7268 |
| 2664_3 | Reason for reducing amount of alcohol drunk: Health precaution | -0.266 | 0.027 | -0.1943 | 0.0015 | -0.5316 | 0.4266 |
| 30120 | Lymphocyte count | 0.1158 | 0.0276 | 0.1467 | 1.2435E-05 | -0.4951 | 0.4683 |
| 1468_2 | Cereal type: Biscuit cereal (e.g. Weetabix) | 0.7147 | 0.0277 | 0.4439 | 0.0034 | 0.7556 | 0.7601 |
| S52 | Diagnoses - main ICD10: S52 Fracture of forearm | -0.3818 | 0.0284 | -0.1879 | 0.0424 | -0.9829 | 0.2986 |
| 6017 | Able to walk or cycle unaided for 10 minutes | -0.6341 | 0.0294 | -0.3291 | 0.0382 | -0.9195 | 0.2131 |
| 20002_1398 | Non-cancer illness code, self-reported: pneumonia | 0.5666 | 0.0296 | 0.3145 | 0.0306 | 0.8451 | 0.7444 |
| 20003_1140883066 | Treatment/medication code: insulin product | 0.3773 | 0.0296 | 0.2303 | 0.031 | 0.7215 | 0.6492 |
| 2030 | Guilty feelings | 0.2059 | 0.0297 | 0.1159 | 0.0182 | 0.8437 | 0.5921 |
| 20553_4 | Methods of self-harm used: Ingesting a medication in excess of the normal dose | 0.3706 | 0.0301 | 0.246 | 0.0241 | 0.6145 | 0.6275 |
| 6143_3 | Transport type for commuting to job workplace: Public transport | -0.3028 | 0.0302 | -0.3044 | 0.0007 | 0.0096 | 0.5016 |
| 2654_4 | Non-butter spread type details: Soft (tub) margarine | 0.4518 | 0.0305 | 0.3675 | 0.0026 | 0.3484 | 0.586 |
| T39 | Diagnoses - main ICD10: T39 Poisoning by nonopioid analgesics, antipyretics and antirheumatics | 0.8294 | 0.0305 | 0.4402 | 0.0255 | 0.9031 | 0.8643 |
| 22127 | Doctor diagnosed asthma | 0.2881 | 0.0316 | 0.1732 | 0.0239 | 0.7438 | 0.6177 |
| 20003_1140871688 | Treatment/medication code: solpadol caplet | 0.5766 | 0.0316 | 0.2575 | 0.048 | 1.07 | 0.7966 |
| 4570 | Friendships satisfaction | 0.2543 | 0.0321 | -0.1556 | 0.0203 | 3.0081 | 0.8554 |
| 20003_1140883548 | Treatment/medication code: ipratropium | 0.4725 | 0.0342 | 0.2761 | 0.025 | 0.7706 | 0.6956 |
| G6_ULLNLE | Lesion of ulnar nerve | 0.7121 | 0.0351 | 0.538 | 0.032 | 0.4137 | 0.6738 |
| 6155_100 | Vitamin and mineral supplements: None of the above | -0.1989 | 0.0354 | -0.1518 | 0.0083 | -0.4258 | 0.4531 |
| M13_IMPINGEMENT | Impingement syndrome of shoulder | 1 | 0.0356 | 0.648 | 0.013 | 0.4888 | 0.7965 |
| 100017 | Magnesium | -0.4161 | 0.0358 | -0.3867 | 0.0003 | -0.1307 | 0.4693 |
| 20426 | Restless during period of worst anxiety | 0.6992 | 0.0361 | 0.7414 | 0.0345 | -0.0872 | 0.4562 |
| 6153_3 | Medication for cholesterol, blood pressure, diabetes, or take exogenous hormones: Insulin | 0.3776 | 0.0365 | 0.2869 | 0.0098 | 0.4278 | 0.5898 |
| K52 | Diagnoses - main ICD10: K52 Other non-infective gastro-enteritis and colitis | 0.802 | 0.0365 | 0.4322 | 0.0461 | 0.8397 | 0.8243 |

| | | | | | | | |
|---|---|---|---|---|---|---|---|
| 2966 | Age high blood pressure diagnosed | -0.3575 | 0.0372 | 0.1803 | 0.0392 | -2.7927 | 0.0812 |
| 3404 | Neck/shoulder pain for 3+ months | 0.564 | 0.0373 | 0.5293 | 0.0088 | 0.1027 | 0.5357 |
| 20003_1140884464 | Treatment/medication code: dihydrocodeine | 0.6975 | 0.0381 | 0.6565 | 0.0114 | 0.0965 | 0.5426 |
| 20539 | Frequency of inability to stop worrying during worst period of anxiety | 0.4101 | 0.0382 | 0.2891 | 0.005 | 0.5424 | 0.6237 |
| AB1_INFECTIONS | Certain infectious and parasitic diseases | 0.7462 | 0.0383 | 0.4072 | 0.0336 | 0.8309 | 0.8094 |
| 20414 | Frequency of drinking alcohol | -0.1924 | 0.0384 | -0.1731 | 0.0045 | -0.1737 | 0.4799 |
| 3456 | Number of cigarettes currently smoked daily (current cigarette smokers) | 0.5051 | 0.0384 | 0.2181 | 0.0478 | 1.072 | 0.7708 |
| 2306 | Weight change compared with 1 year ago | 0.4032 | 0.0386 | 0.2999 | 0.0052 | 0.4643 | 0.6054 |
| T81 | Diagnoses - main ICD10: T81 Complications of procedures, not elsewhere classified | 0.8109 | 0.0387 | 0.5584 | 0.0433 | 0.5262 | 0.7617 |
| 6157_3 | Why stopped smoking: Health precaution | -0.3487 | 0.0396 | -0.3773 | 0.0002 | 0.1446 | 0.5286 |
| 100005 | Carbohydrate | -0.5129 | 0.0401 | -0.5332 | 0.0004 | 0.0697 | 0.5211 |
| 20003_1140879406 | Treatment/medication code: ranitidine | 0.7097 | 0.0406 | 0.3435 | 0.0301 | 0.9611 | 0.8294 |
| K02 | Diagnoses - main ICD10: K02 Dental caries | 0.6302 | 0.0407 | 0.5372 | 0.0048 | 0.2568 | 0.5935 |
| 20003_1140881856 | Treatment/medication code: salbutamol | 0.4247 | 0.041 | 0.32 | 0.0114 | 0.4304 | 0.6074 |
| 20002_1478 | Non-cancer illness code, self-reported: cervical spondylosis | 0.4989 | 0.0411 | 0.5488 | 0.0002 | -0.1746 | 0.4484 |
| XII_SKIN_SUBCUTAN | Diseases of the skin and subcutaneous tissue | 0.9615 | 0.0411 | 0.59 | 0.014 | 0.7031 | 0.8742 |
| 6034 | Target heart rate achieved | -0.3159 | 0.043 | -0.2086 | 0.0351 | -0.5805 | 0.3897 |
| 6149_2 | Mouth/teeth dental problems: Painful gums | 0.4085 | 0.0432 | 0.2481 | 0.0427 | 0.6791 | 0.6568 |
| 100009 | Englyst dietary fibre | -0.4018 | 0.0433 | -0.2142 | 0.0464 | -0.8301 | 0.3119 |
| M47 | Diagnoses - main ICD10: M47 Spondylosis | 1 | 0.0447 | 0.4368 | 0.0173 | 0.8227 | 0.9063 |
| 3761 | Age hay fever, rhinitis or eczema diagnosed | 0.2757 | 0.0453 | 0.1985 | 0.0032 | 0.5038 | 0.5775 |
| 2704 | Years since last cervical smear test | 0.331 | 0.0455 | 0.3086 | 0.0003 | 0.1205 | 0.5231 |
| 20003_1141162764 | Treatment/medication code: tolterodine l-tartrate | 0.6936 | 0.0455 | 0.4611 | 0.0326 | 0.5692 | 0.7277 |
| 22601_71113328 | Job coding: sales assistant, retail/shop assistant, counter assistant, sub post office assistant, takeaway food worker | 0.5689 | 0.0455 | 0.3797 | 0.0407 | 0.557 | 0.6905 |
| 20554_1 | Actions taken following self-harm: See anyone from psychiatric or mental health services, including liaison services | 0.3799 | 0.0476 | 0.2741 | 0.0321 | 0.4589 | 0.6087 |
| 22607_1 | Workplace very cold: Sometimes | 0.2926 | 0.0481 | 0.382 | 2.2577E-05 | -0.5156 | 0.4072 |
| 22608_2 | Workplace very hot: Often | 0.6226 | 0.0485 | 0.5248 | 0.0082 | 0.2624 | 0.6013 |
| 874 | Duration of walks | 0.1941 | 0.0492 | 0.2375 | 2.7439E-06 | -0.3913 | 0.4551 |
| 20002_1474 | Non-cancer illness code, self-reported: hiatus hernia | 0.3786 | 0.0495 | 0.3625 | 0.0011 | 0.0724 | 0.5167 |

**Table S26.** Linkage disequilibrium score regression (rg) results for 435 UK Biobank traits demonstrating nominally significant genetic correlations with attention deficit hyperactivity disorder (ADHD) and *PhoneUse* in male subjects.

| Field ID | Trait | ADHD | | *PhoneUse* | | Difference | |
|---|---|---|---|---|---|---|---|
| | | rg | p | rg | p | z | p |
| 2139 | Age at first sexual intercourse | -0.6103 | 6.56E-69 | -0.5923 | 5.03E-57 | -0.35336 | 0.475964 |
| 6138_100 | Qualifications: None of the above | 0.6209 | 1.18E-42 | 0.3714 | 7.89E-21 | 4.136985 | 0.798314 |
| 6138_1 | Qualifications: College or University degree | -0.5455 | 3.83E-40 | -0.4048 | 5.49E-30 | -2.58762 | 0.31874 |
| 845 | Age completed full time education | -0.7382 | 1.35E-38 | -0.3202 | 7.96E-11 | -5.55768 | 0.080765 |
| 6138_2 | Qualifications: A levels/AS levels or equivalent | -0.6078 | 2.31E-38 | -0.3786 | 8.14E-18 | -3.56406 | 0.221355 |
| 816 | Job involves heavy manual or physical work | 0.595 | 8.1E-36 | 0.2861 | 1.23E-09 | 4.61293 | 0.849561 |
| 806 | Job involves mainly walking or standing | 0.6495 | 5.23E-35 | 0.2854 | 3.69E-08 | 4.931993 | 0.888659 |
| 23112 | Leg fat mass (right) | 0.4108 | 1.51E-32 | 0.3263 | 7.01E-23 | 1.764723 | 0.611413 |
| 23111 | Leg fat percentage (right) | 0.4135 | 7.93E-32 | 0.3101 | 4.68E-21 | 2.146055 | 0.635442 |
| 21001 | Body mass index (BMI) | 0.4036 | 2.26E-31 | 0.3454 | 4.15E-24 | 1.198036 | 0.577273 |
| 1050 | Time spend outdoors in summer | 0.4988 | 3.38E-31 | 0.4049 | 5.86E-20 | 1.522678 | 0.623424 |
| 23104 | Body mass index (Kg/m2) | 0.4073 | 4.51E-31 | 0.3455 | 2.08E-23 | 1.252105 | 0.581986 |
| 6138_3 | Qualifications: O levels/GCSEs or equivalent | -0.6137 | 1.03E-30 | -0.3319 | 9.81E-08 | -3.43705 | 0.172638 |
| 23116 | Leg fat mass (left) | 0.4005 | 2.51E-30 | 0.3305 | 1.66E-23 | 1.453105 | 0.592679 |
| 23115 | Leg fat percentage (left) | 0.4078 | 6.88E-30 | 0.3225 | 5.96E-23 | 1.75658 | 0.612439 |
| 738 | Average total household income before tax | -0.4787 | 7.82E-28 | -0.1032 | 0.0138 | -6.19494 | 0.104267 |
| 2316 | Wheeze or whistling in the chest in last year | 0.5183 | 1.1E-27 | 0.3161 | 2.04E-09 | 2.849997 | 0.75086 |
| 6159_100 | Pain type(s) experienced in last month: None of the above | -0.5936 | 6.91E-27 | -0.368 | 5.64E-10 | -2.78231 | 0.224954 |
| 20116_0 | Smoking status: Never | -0.4825 | 1.97E-26 | -0.4091 | 1.91E-18 | -1.12696 | 0.402908 |
| XIII_MUSCULOSKELET | Diseases of the musculoskeletal system and connective tissue | 0.5948 | 6.11E-26 | 0.5131 | 2.41E-15 | 0.950303 | 0.607814 |
| 23110 | Impedance of arm (left) | -0.3616 | 3.53E-25 | -0.2887 | 8.35E-16 | -1.45602 | 0.403556 |
| 4825 | Noisy workplace | 0.6992 | 6.73E-24 | 0.3467 | 3.6E-07 | 3.625367 | 0.881115 |
| 48 | Waist circumference | 0.3512 | 7E-24 | 0.2969 | 1.01E-18 | 1.120846 | 0.572153 |
| 6146_100 | Attendance/disability/mobility allowance: None of the above | -0.6331 | 7.38E-24 | -0.5404 | 5E-18 | -1.04543 | 0.378103 |
| 23100 | Whole body fat mass | 0.3453 | 4.44E-22 | 0.2908 | 9.29E-18 | 1.107024 | 0.572416 |
| 680_3 | Own or rent accommodation lived in: Rent - from local authority, local council, housing association | 0.5645 | 5.96E-22 | 0.3426 | 7.8E-09 | 2.661649 | 0.771313 |
| 23120 | Arm fat mass (right) | 0.3323 | 2.85E-21 | 0.2952 | 2.22E-18 | 0.762449 | 0.549443 |
| 23109 | Impedance of arm (right) | -0.3326 | 3.42E-21 | -0.282 | 2.26E-14 | -0.99222 | 0.432715 |
| 1239 | Current tobacco smoking | 0.511 | 1.79E-20 | 0.3897 | 2.7E-11 | 1.509395 | 0.657721 |
| 1458 | Cereal intake | -0.4321 | 2.87E-20 | -0.3294 | 4.58E-11 | -1.49959 | 0.365439 |
| 6159_3 | Pain type(s) experienced in last month: Neck or shoulder pain | 0.5616 | 4.23E-20 | 0.3256 | 1.15E-07 | 2.7223 | 0.785353 |
| 23099 | Body fat percentage | 0.3359 | 4.78E-20 | 0.2721 | 3.51E-16 | 1.28761 | 0.5846 |
| 1787 | Maternal smoking around birth | 0.569 | 4.92E-20 | 0.3487 | 2.85E-09 | 2.578046 | 0.769688 |
| 23124 | Arm fat mass (left) | 0.3255 | 8.76E-20 | 0.2921 | 1.3E-17 | 0.674606 | 0.544533 |
| 6142_4 | Current employment status: Unable to work because of sickness or disability | 0.6267 | 9.06E-20 | 0.3988 | 1.06E-09 | 2.399031 | 0.777349 |
| 189 | Townsend deprivation index at recruitment | 0.533 | 1.31E-19 | 0.2391 | 0.0001 | 3.442422 | 0.83752 |
| 6159_7 | Pain type(s) experienced in last month: Knee pain | 0.5336 | 2.27E-19 | 0.2904 | 1.13E-05 | 2.738694 | 0.792325 |
| 23128 | Trunk fat mass | 0.3226 | 3.17E-19 | 0.2791 | 1.61E-16 | 0.880913 | 0.557916 |
| 2188 | Long-standing illness, disability or infirmity | 0.4998 | 4.49E-19 | 0.2857 | 8.61E-08 | 2.766889 | 0.76333 |
| ICDMAIN_ANY_ENTRY | Any ICDMAIN event in hilmo or causes of death | 0.6883 | 5.06E-19 | 0.5313 | 2.82E-16 | 1.556681 | 0.700492 |
| 6146_3 | Attendance/disability/mobility allowance: Blue badge | 0.6502 | 1.64E-18 | 0.5134 | 1.47E-12 | 1.319597 | 0.676583 |
| 1249 | Past tobacco smoking | -0.401 | 2.15E-18 | -0.3512 | 7.7E-14 | -0.75886 | 0.433768 |
| 2040 | Risk Taking | 0.4273 | 2.76E-18 | 0.5833 | 2.67E-29 | -2.18559 | 0.300672 |
| 20116_2 | Smoking status: Current | 0.5083 | 3.65E-18 | 0.3772 | 6.65E-10 | 1.549829 | 0.669696 |
| 22501 | Year ended full time education | -0.5985 | 3.92E-18 | -0.4292 | 4.78E-10 | -1.73749 | 0.285353 |
| 23098 | Weight | 0.2845 | 1.13E-17 | 0.2559 | 2.79E-14 | 0.605476 | 0.538155 |

| | | | | | | | |
|---|---|---|---|---|---|---|---|
| 20016 | Fluid intelligence score | -0.431 | 1.66E-17 | -0.2941 | 1.11E-08 | -1.89616 | 0.323297 |
| 21002 | Weight | 0.2779 | 2.46E-17 | 0.2557 | 1.51E-14 | 0.474957 | 0.529635 |
| 1538_0 | Major dietary changes in the last 5 years: No | -0.503 | 3.04E-17 | -0.3714 | 9.47E-10 | -1.54699 | 0.329697 |
| 23119 | Arm fat percentage (right) | 0.3055 | 4.91E-17 | 0.2556 | 3.24E-14 | 1.005948 | 0.566363 |
| 1548 | Variation in diet | 0.4645 | 5.3E-17 | 0.1784 | 0.0009 | 3.70479 | 0.831017 |
| 41248_1000 | Destinations on discharge from hospital (recoded): Usual Place of residence | 0.6803 | 8.32E-17 | 0.4984 | 3.8E-11 | 1.636148 | 0.728808 |
| 2080 | Frequency of tiredness / lethargy in last 2 weeks | 0.4466 | 8.99E-17 | 0.1798 | 0.0011 | 3.467656 | 0.81422 |
| 23106 | Impedance of whole body | -0.2816 | 1.29E-16 | -0.2413 | 3.47E-11 | -0.80909 | 0.446317 |
| 6146_2 | Attendance/disability/mobility allowance: Disability living allowance | 0.5974 | 2.14E-16 | 0.5608 | 9.09E-18 | 0.374536 | 0.54878 |
| 2060 | Frequency of unenthusiasm / disinterest in last 2 weeks | 0.4644 | 2.82E-16 | 0.2462 | 0.0005 | 2.405991 | 0.767545 |
| 23127 | Trunk fat percentage | 0.3043 | 5.79E-16 | 0.2571 | 2.29E-14 | 0.9348 | 0.562803 |
| 1070 | Time spent watching television (TV) | 0.3719 | 1.08E-15 | 0.2859 | 1.86E-11 | 1.3653 | 0.613337 |
| 6145_6 | Illness, injury, bereavement, stress in last 2 years: Financial difficulties | 0.4988 | 1.8E-15 | 0.4777 | 1.17E-14 | 0.239481 | 0.528169 |
| 137 | Number of treatments/medications taken | 0.4114 | 6.69E-15 | 0.3052 | 1.36E-10 | 1.495316 | 0.63896 |
| 6138_6 | Qualifications: Other professional qualifications eg: nursing, teaching | -0.5126 | 4.21E-14 | -0.2571 | 4.75E-05 | -2.75438 | 0.196079 |
| 2492 | Taking other prescription medications | 0.4537 | 4.72E-14 | 0.3128 | 1.51E-08 | 1.723668 | 0.681499 |
| 2867 | Age started smoking in former smokers | -0.6336 | 4.74E-14 | -0.407 | 2.58E-07 | -1.96509 | 0.223951 |
| 135 | Number of self-reported non-cancer illnesses | 0.4124 | 6.38E-14 | 0.2203 | 1.08E-05 | 2.582071 | 0.740009 |
| 23123 | Arm fat percentage (left) | 0.2813 | 8.88E-14 | 0.2477 | 6.18E-13 | 0.658362 | 0.544799 |
| 23126 | Arm predicted mass (left) | 0.2468 | 2.14E-13 | 0.2144 | 3.25E-10 | 0.676799 | 0.543205 |
| 23125 | Arm fat-free mass (left) | 0.2474 | 2.17E-13 | 0.217 | 1.47E-10 | 0.635975 | 0.540548 |
| 6143_3 | Transport type for commuting to job workplace: Public transport | -0.5685 | 2.35E-13 | -0.5237 | 3.03E-11 | -0.40508 | 0.440366 |
| 2887 | Number of cigarettes previously smoked daily | 0.4472 | 5.02E-13 | 0.2138 | 0.0085 | 2.285924 | 0.782802 |
| 894 | Duration of moderate activity | 0.4914 | 5.54E-13 | 0.3417 | 2.29E-07 | 1.578537 | 0.691944 |
| XI_DIGESTIVE | Diseases of the digestive system | 0.4632 | 6.21E-13 | 0.3247 | 1.47E-06 | 1.48572 | 0.678625 |
| 2296 | Falls in the last year | 0.5345 | 6.49E-13 | 0.3765 | 1.04E-07 | 1.539495 | 0.701655 |
| 1920 | Mood swings | 0.3912 | 8.65E-13 | 0.246 | 0.0012 | 1.553343 | 0.686622 |
| 23121 | Arm fat-free mass (right) | 0.2412 | 8.7E-13 | 0.2204 | 1.13E-10 | 0.433208 | 0.527769 |
| 6164_1 | Types of physical activity in last 4 weeks: Walking for pleasure (not as a means of transport) | -0.4694 | 1.02E-12 | -0.3151 | 2.28E-06 | -1.64562 | 0.302657 |
| 23122 | Arm predicted mass (right) | 0.2405 | 1.1E-12 | 0.2185 | 1.86E-10 | 0.456856 | 0.529368 |
| 924 | Usual walking pace | -0.3201 | 1.28E-12 | -0.4117 | 6.19E-21 | 1.455395 | 0.620496 |
| 6159_4 | Pain type(s) experienced in last month: Back pain | 0.5317 | 1.34E-12 | 0.2563 | 0.0002 | 2.702339 | 0.821829 |
| 826 | Job involves shift work | 0.5865 | 1.36E-12 | 0.3592 | 1.29E-06 | 2.045762 | 0.77675 |
| 1448_3 | Bread type: Wholemeal or wholegrain | -0.3818 | 1.39E-12 | -0.1919 | 0.0018 | -2.32217 | 0.262387 |
| 1448_1 | Bread type: White | 0.3676 | 1.63E-12 | 0.1954 | 0.0002 | 2.330381 | 0.717937 |
| 41235 | Spells in hospital | 0.6101 | 2.2E-12 | 0.531 | 4.9E-11 | 0.666609 | 0.604464 |
| 4968 | FI4 : positional arithmetic | -0.6941 | 3.12E-12 | -0.2356 | 0.0122 | -3.34786 | 0.06232 |
| 6149_6 | Mouth/teeth dental problems: Dentures | 0.3714 | 3.26E-12 | 0.3887 | 3.27E-12 | -0.22419 | 0.476898 |
| 6145_100 | Illness, injury, bereavement, stress in last 2 years: None of the above | -0.5188 | 3.71E-12 | -0.4897 | 2.59E-10 | -0.27035 | 0.46118 |
| 874 | Duration of walks | 0.3983 | 4.88E-12 | 0.2324 | 6.5E-05 | 2.026035 | 0.710766 |
| 49 | Hip circumference | 0.2427 | 5.6E-12 | 0.2194 | 9.68E-10 | 0.463426 | 0.5311 |
| 2149 | Lifetime number of sexual partners | 0.3132 | 7.81E-12 | 0.4362 | 1.13E-18 | -1.82588 | 0.34019 |
| 6149_100 | Mouth/teeth dental problems: None of the above | -0.42 | 7.9E-12 | -0.4026 | 9.07E-10 | -0.1935 | 0.476765 |
| 41231_1 | Hospital episode type: General episode | 0.6999 | 8.15E-12 | 0.5424 | 6.78E-13 | 1.237972 | 0.701074 |
| 2443 | Diabetes diagnosed by doctor | 0.3804 | 1E-11 | 0.2193 | 4.8E-05 | 2.072755 | 0.705246 |
| 4957 | FI3 : word interpolation | -0.5664 | 1.59E-11 | -0.4098 | 2.51E-08 | -1.40302 | 0.299973 |
| 30290 | High light scatter reticulocyte percentage | 0.3177 | 2.07E-11 | 0.2255 | 2.83E-08 | 1.477306 | 0.62126 |
| 4653 | Ever highly irritable/argumentative for 2 days | 0.4756 | 2.59E-11 | 0.3503 | 4.05E-05 | 1.127057 | 0.662629 |
| 20116_1 | Smoking status: Previous | 0.3693 | 2.7E-11 | 0.3536 | 8.82E-11 | 0.202024 | 0.520967 |
| 20002_1465 | Non-cancer illness code, self-reported: osteoarthritis | 0.6462 | 2.8E-11 | 0.4658 | 6.53E-06 | 1.272466 | 0.72714 |
| 680_1 | Own or rent accommodation lived in: Own outright (by you or someone in your household) | -0.4226 | 3.29E-11 | -0.5891 | 1.82E-20 | 1.85115 | 0.711453 |

| | | | | | | | |
|---|---|---|---|---|---|---|---|
| XVIII_MISCFINDINGS | Symptoms, signs and abnormal clinical and laboratory findings, not elsewhere classified | 0.5623 | 5.08E-11 | 0.4226 | 7.45E-09 | 1.241056 | 0.680064 |
| 22620_1 | Job involved shift work: Yes | 0.8969 | 5.91E-11 | 0.578 | 9.9E-06 | 1.683614 | 0.85725 |
| 1578 | Average weekly champagne plus white wine intake | -0.4242 | 8.11E-11 | -0.149 | 0.0132 | -3.10094 | 0.178346 |
| 30240 | Reticulocyte percentage | 0.2995 | 9.26E-11 | 0.2085 | 2.44E-07 | 1.482746 | 0.61973 |
| 1468_4 | Cereal type: Muesli | -0.3487 | 9.7E-11 | -0.3691 | 1.87E-12 | 0.271374 | 0.527236 |
| 22606_2 | Workplace very noisy: Often | 0.6368 | 1.01E-10 | 0.2414 | 0.0017 | 3.162568 | 0.907291 |
| 4728 | Leg pain on walking | 0.5636 | 1.22E-10 | 0.3624 | 4.32E-05 | 1.614842 | 0.749797 |
| 20488 | Physically abused by family as a child | 0.5941 | 1.38E-10 | 0.3346 | 0.0003 | 1.986936 | 0.807605 |
| M13_ARTHROSIS | #Arthrosis | 0.4268 | 1.87E-10 | 0.4401 | 5.36E-10 | -0.13634 | 0.482235 |
| 20111_100 | Illnesses of siblings: None of the above (group 1) | -0.4738 | 2.17E-10 | -0.2492 | 0.0002 | -2.24293 | 0.225959 |
| 1478 | Salt added to food | 0.2999 | 2.25E-10 | 0.2318 | 8.24E-08 | 1.063086 | 0.590207 |
| 30300 | High light scatter reticulocyte count | 0.3103 | 2.43E-10 | 0.2164 | 9.82E-08 | 1.475613 | 0.623424 |
| 20160 | Ever smoked | 0.3012 | 2.83E-10 | 0.3237 | 1.18E-10 | -0.32459 | 0.469966 |
| 6154_100 | Medication for pain relief, constipation, heartburn: None of the above | -0.3503 | 3.25E-10 | -0.3555 | 1.49E-10 | 0.066132 | 0.506947 |
| 6164_4 | Types of physical activity in last 4 weeks: Light DIY (eg: pruning, watering the lawn) | -0.393 | 3.9E-10 | -0.2974 | 4.18E-07 | -1.11123 | 0.374416 |
| 1628 | Alcohol intake versus 10 years previously | 0.3749 | 4.47E-10 | 0.2351 | 2.48E-05 | 1.704669 | 0.680183 |
| 1807 | Father's age at death | -0.4532 | 5.66E-10 | -0.3329 | 7.15E-05 | -1.08181 | 0.34351 |
| 23117 | Leg fat-free mass (left) | 0.2056 | 5.96E-10 | 0.1974 | 1.21E-08 | 0.171004 | 0.510955 |
| 22607_1 | Workplace very cold: Sometimes | 0.5776 | 6.26E-10 | 0.3877 | 4.13E-06 | 1.510133 | 0.737613 |
| 23118 | Leg predicted mass (left) | 0.2049 | 6.84E-10 | 0.1971 | 1.21E-08 | 0.162662 | 0.510421 |
| 22615_1 | Workplace had a lot of diesel exhaust: Sometimes | 0.5615 | 7.36E-10 | 0.3854 | 2.41E-05 | 1.364619 | 0.722333 |
| 2335 | Chest pain or discomfort | 0.4021 | 1.08E-09 | 0.242 | 0.0002 | 1.740171 | 0.70409 |
| 110001 | Invitation to complete online 24-hour recall dietary questionnaire, acceptance | -0.3892 | 1.16E-09 | -0.3544 | 8.34E-08 | -0.37823 | 0.453608 |
| 30250 | Reticulocyte count | 0.2888 | 1.54E-09 | 0.1937 | 1.35E-06 | 1.524217 | 0.624949 |
| 20003_1140884600 | Treatment/medication code: metformin | 0.3941 | 1.59E-09 | 0.2441 | 0.0002 | 1.620561 | 0.692297 |
| 1873 | Number of full brothers | 0.5804 | 1.61E-09 | 0.3185 | 0.0002 | 2.02442 | 0.809796 |
| 2070 | Frequency of tenseness / restlessness in last 2 weeks | 0.3566 | 1.83E-09 | 0.2275 | 0.0011 | 1.410733 | 0.667266 |
| 2020 | Loneliness, isolation | 0.4295 | 2.3E-09 | 0.1507 | 0.0477 | 2.663 | 0.824782 |
| 23105 | Basal metabolic rate | 0.1984 | 2.35E-09 | 0.1852 | 4.16E-08 | 0.27861 | 0.517631 |
| 1060 | Time spent outdoors in winter | 0.3604 | 2.41E-09 | 0.3344 | 6.34E-11 | 0.328362 | 0.534695 |
| 22613_1 | Worked with paints, thinners or glues: Sometimes | 0.5713 | 2.48E-09 | 0.28 | 0.003 | 2.167005 | 0.835371 |
| 6160_3 | Leisure/social activities: Religious group | -0.3962 | 2.6E-09 | -0.2863 | 1.18E-05 | -1.17829 | 0.35641 |
| XIX_INJURY_POISON | Injury, poisoning and certain other consequences of external causes | 0.6513 | 3.08E-09 | 0.4823 | 1.53E-06 | 1.135838 | 0.714305 |
| 6159_6 | Pain type(s) experienced in last month: Hip pain | 0.4455 | 3.5E-09 | 0.3643 | 2.81E-06 | 0.749478 | 0.60717 |
| 1538_1 | Major dietary changes in the last 5 years: Yes, because of illness | 0.3872 | 3.53E-09 | 0.2913 | 2.44E-06 | 1.064072 | 0.625964 |
| 6149_4 | Mouth/teeth dental problems: Loose teeth | 0.6122 | 4.98E-09 | 0.3156 | 0.0018 | 2.039806 | 0.839732 |
| 22610_1 | Workplace full of chemical or other fumes: Sometimes | 0.7173 | 5.3E-09 | 0.2435 | 0.0263 | 2.877251 | 0.943724 |
| 20003_1140864752 | Treatment/medication code: lansoprazole | 0.6697 | 5.69E-09 | 0.3936 | 7.45E-05 | 1.816394 | 0.822439 |
| 22609_2 | Workplace very dusty: Often | 0.6347 | 5.79E-09 | 0.3891 | 3.53E-05 | 1.705563 | 0.794619 |
| M13_DORSALGIA | Dorsalgia | 0.431 | 6.44E-09 | 0.5584 | 8.68E-08 | -0.99486 | 0.334805 |
| M54 | Diagnoses - main ICD10: M54 Dorsalgia | 0.431 | 6.44E-09 | 0.5584 | 8.68E-08 | -0.99486 | 0.334805 |
| 6162_2 | Types of transport used (excluding work): Walk | -0.3904 | 7.4E-09 | -0.5111 | 4.85E-13 | 1.234803 | 0.656983 |
| 6152_6 | Blood clot, DVT, bronchitis, emphysema, asthma, rhinitis, eczema, allergy diagnosed by doctor: Emphysema/chronic bronchitis | 0.5438 | 8.33E-09 | 0.2911 | 0.0029 | 1.861053 | 0.801316 |
| M13_OTHERJOINT | #Other joint disorders | 0.5647 | 8.79E-09 | 0.5404 | 3.31E-08 | 0.175333 | 0.532432 |
| 1408 | Cheese intake | -0.3085 | 8.92E-09 | -0.1745 | 0.0048 | -1.6352 | 0.326793 |
| 20002_1220 | Non-cancer illness code, self-reported: diabetes | 0.3485 | 1.3E-08 | 0.1994 | 0.0006 | 1.769675 | 0.691237 |

| | | | | | | | |
|---|---|---|---|---|---|---|---|
| IX_CIRCULATORY | Diseases of the circulatory system | 0.3928 | 1.45E-08 | 0.3 | 1.4E-06 | 0.996564 | 0.622024 |
| 6164_100 | Types of physical activity in last 4 weeks: None of the above | 0.585 | 1.56E-08 | 0.4329 | 1.61E-08 | 1.181244 | 0.694766 |
| 20117_1 | Alcohol drinker status: Previous | 0.5551 | 1.86E-08 | 0.4114 | 1.05E-05 | 1.057497 | 0.684839 |
| 1279 | Exposure to tobacco smoke outside home | 0.3894 | 1.95E-08 | 0.4652 | 5.07E-15 | -0.83047 | 0.3998 |
| KNEE_ARTHROSIS | Gonarthrosis [arthrosis of knee](FG) | 0.402 | 2.15E-08 | 0.4036 | 9.59E-10 | -0.01641 | 0.497862 |
| M17 | Diagnoses - main ICD10: M17 Gonarthrosis [arthrosis of knee] | 0.402 | 2.15E-08 | 0.4036 | 9.59E-10 | -0.01641 | 0.497862 |
| 6177_100 | Medication for cholesterol, blood pressure or diabetes: None of the above | -0.2819 | 2.69E-08 | -0.1419 | 0.0042 | -1.97386 | 0.319577 |
| 30280 | Immature reticulocyte fraction | 0.2827 | 3.2E-08 | 0.2247 | 7.62E-07 | 0.848515 | 0.57701 |
| 6177_1 | Medication for cholesterol, blood pressure or diabetes: Cholesterol lowering medication | 0.292 | 3.3E-08 | 0.1546 | 0.0017 | 1.901915 | 0.677304 |
| 5556 | FI9 : concept interpolation | -0.5875 | 3.87E-08 | -0.3376 | 0.0002 | -1.77187 | 0.201309 |
| 1558 | Alcohol intake frequency. | 0.2435 | 4.78E-08 | 0.1499 | 0.0008 | 1.47899 | 0.623042 |
| 136 | Number of operations, self-reported | 0.3327 | 5.01E-08 | 0.4465 | 1.19E-12 | -1.29984 | 0.351552 |
| 4717 | Shortness of breath walking on level ground | 0.495 | 1.18E-07 | 0.3364 | 0.0003 | 1.202649 | 0.702352 |
| 20111_9 | Illnesses of siblings: Diabetes | 0.5906 | 1.26E-07 | 0.2227 | 0.0106 | 2.595893 | 0.891054 |
| 6164_5 | Types of physical activity in last 4 weeks: Heavy DIY (eg: weeding, lawn mowing, carpentry, digging) | -0.3527 | 1.36E-07 | -0.1856 | 0.0021 | -1.85395 | 0.287861 |
| 1588 | Average weekly beer plus cider intake | 0.2935 | 1.43E-07 | 0.2138 | 0.0008 | 0.939479 | 0.605238 |
| 6150_100 | Vascular/heart problems diagnosed by doctor: None of the above | -0.2404 | 1.45E-07 | -0.1575 | 0.0005 | -1.29256 | 0.390643 |
| 22506_114 | Tobacco smoking: Never smoked | -0.4106 | 1.46E-07 | -0.5026 | 9.9E-11 | 0.835091 | 0.621005 |
| 6143_1 | Transport type for commuting to job workplace: Car/motor vehicle | 0.4291 | 1.52E-07 | 0.6258 | 1.12E-15 | -1.74033 | 0.255019 |
| 1883 | Number of full sisters | 0.5713 | 1.59E-07 | 0.3093 | 0.0008 | 1.833524 | 0.809887 |
| COPD_EXCL | COPD differential diagnosis | 0.54 | 1.63E-07 | 0.6603 | 1.69E-09 | -0.79949 | 0.34351 |
| ILD_DIFF_DG | ILD differential diagnosis | 0.54 | 1.63E-07 | 0.6603 | 1.69E-09 | -0.79949 | 0.34351 |
| 4990 | FI6 : conditional arithmetic | -0.3859 | 1.67E-07 | -0.4285 | 9.94E-09 | 0.405682 | 0.556726 |
| 4581 | Financial situation satisfaction | 0.5099 | 1.84E-07 | 0.4257 | 3.69E-06 | 0.627088 | 0.611028 |
| 20117_2 | Alcohol drinker status: Current | -0.3907 | 1.87E-07 | -0.2407 | 0.0045 | -1.32587 | 0.307703 |
| 1269 | Exposure to tobacco smoke at home | 0.6534 | 1.92E-07 | 0.5164 | 1.46E-05 | 0.791512 | 0.676823 |
| 23102 | Whole body water mass | 0.1743 | 2.29E-07 | 0.1637 | 1.15E-06 | 0.222413 | 0.51416 |
| PULMONARYDG | Other pulmonary diagnosis | 0.5631 | 2.29E-07 | 0.6594 | 8.87E-09 | -0.60913 | 0.373528 |
| X_RESPIRATORY | Diseases of the respiratory system | 0.5631 | 2.29E-07 | 0.6594 | 8.87E-09 | -0.60913 | 0.373528 |
| 1259 | Smoking/smokers in household | 0.7645 | 2.39E-07 | 0.4551 | 0.0001 | 1.639978 | 0.849952 |
| 1538_2 | Major dietary changes in the last 5 years: Yes, because of other reasons | 0.4529 | 2.76E-07 | 0.3178 | 0.0003 | 1.089272 | 0.674535 |
| 20511 | Recent poor appetite or overeating | 0.4763 | 3.34E-07 | 0.3837 | 8.21E-05 | 0.686554 | 0.62177 |
| 23101 | Whole body fat-free mass | 0.1714 | 3.45E-07 | 0.1649 | 9.95E-07 | 0.136588 | 0.508684 |
| 22607_2 | Workplace very cold: Often | 0.6226 | 3.58E-07 | 0.2788 | 0.0034 | 2.21828 | 0.875224 |
| 23114 | Leg predicted mass (right) | 0.1705 | 3.92E-07 | 0.1781 | 3.64E-07 | -0.15664 | 0.489847 |
| 23113 | Leg fat-free mass (right) | 0.1699 | 4.75E-07 | 0.1773 | 4.07E-07 | -0.1523 | 0.490114 |
| R07 | Diagnoses - main ICD10: R07 Pain in throat and chest | 0.3709 | 5.53E-07 | 0.4705 | 5.14E-09 | -0.91032 | 0.36935 |
| 2473 | Other serious medical condition/disability diagnosed by doctor | 0.3427 | 7.21E-07 | 0.1918 | 0.0056 | 1.540829 | 0.693357 |
| 20003_2038460150 | Treatment/medication code: paracetamol | 0.3416 | 8.79E-07 | 0.3273 | 2.55E-05 | 0.137174 | 0.519099 |
| 2415 | Had major operations | 0.35 | 8.81E-07 | 0.3049 | 1.06E-05 | 0.454235 | 0.560031 |
| 904 | Number of days/week of vigorous physical activity 10+ minutes | 0.278 | 9.16E-07 | 0.1139 | 0.0408 | 2.066474 | 0.708702 |
| 20487 | Felt hated by family member as a child | 0.6725 | 1.29E-06 | 0.331 | 0.0034 | 1.908535 | 0.873633 |
| 6159_5 | Pain type(s) experienced in last month: Stomach or abdominal pain | 0.4736 | 1.58E-06 | 0.2126 | 0.0215 | 1.931494 | 0.808976 |
| 6154_3 | Medication for pain relief, constipation, heartburn: Paracetamol | 0.313 | 1.62E-06 | 0.2898 | 2.3E-05 | 0.245145 | 0.530967 |
| 20002_1065 | Non-cancer illness code, self-reported: hypertension | 0.2175 | 1.69E-06 | 0.1478 | 0.0015 | 1.071329 | 0.592289 |
| G6_NERPLEX | Nerve, nerve root and plexus disorders | 0.4815 | 1.71E-06 | 0.3919 | 5.28E-05 | 0.641475 | 0.617944 |
| 6162_4 | Types of transport used (excluding work): Cycle | -0.3071 | 1.71E-06 | -0.1861 | 0.0089 | -1.2631 | 0.342648 |

| | | | | | | | |
|---|---|---|---|---|---|---|---|
| 1309 | Fresh fruit intake | -0.241 | 1.84E-06 | -0.1568 | 0.0028 | -1.15701 | 0.388972 |
| 30000 | White blood cell (leukocyte) count | 0.1972 | 1.99E-06 | 0.1354 | 0.001 | 1.056806 | 0.581986 |
| G56 | Diagnoses - main ICD10: G56 Mononeuropathies of upper limb | 0.4456 | 2.16E-06 | 0.3419 | 0.0002 | 0.786741 | 0.63582 |
| 6150_4 | Vascular/heart problems diagnosed by doctor: High blood pressure | 0.2154 | 2.28E-06 | 0.1489 | 0.0011 | 1.028938 | 0.588123 |
| 22506_113 | Tobacco smoking: Ex-smoker | 0.3818 | 2.67E-06 | 0.4933 | 4.59E-09 | -0.95263 | 0.354414 |
| 6144_4 | Never eat eggs, dairy, wheat, sugar: Sugar or foods/drinks containing sugar | 0.2925 | 2.75E-06 | 0.1609 | 0.0061 | 1.537345 | 0.670303 |
| 1508_3 | Coffee type: Ground coffee (include espresso, filter etc) | -0.2234 | 3.11E-06 | -0.137 | 0.0164 | -1.15925 | 0.38615 |
| 6147_1 | Reason for glasses/contact lenses: For short-sightedness, i.e. only or mainly for distance viewing such as driving, cinema etc (called 'myopia') | -0.3707 | 3.64E-06 | -0.1922 | 0.0128 | -1.60558 | 0.274979 |
| 22612_1 | Worked with materials containing asbestos: Sometimes | 0.5288 | 3.65E-06 | 0.4098 | 0.0002 | 0.7561 | 0.654887 |
| 20111_1 | Illnesses of siblings: Heart disease | 0.6447 | 4.19E-06 | 0.3739 | 0.0002 | 1.564223 | 0.817784 |
| 2644 | Light smokers, at least 100 smokes in lifetime | 0.4289 | 4.48E-06 | 0.4613 | 4.66E-08 | -0.25723 | 0.456795 |
| 22611_2 | Workplace had a lot of cigarette smoke from other people smoking: Often | 0.5516 | 4.99E-06 | 0.3098 | 0.0212 | 1.337507 | 0.79098 |
| 6146_1 | Attendance/disability/mobility allowance: Attendance allowance | 0.5306 | 5.24E-06 | 0.3887 | 0.0002 | 0.909028 | 0.682694 |
| 23108 | Impedance of leg (left) | -0.154 | 5.63E-06 | -0.1593 | 2.49E-05 | 0.104383 | 0.507081 |
| 22609_1 | Workplace very dusty: Sometimes | 0.7351 | 5.67E-06 | 0.5002 | 0.0005 | 1.083411 | 0.784276 |
| 6144_5 | Never eat eggs, dairy, wheat, sugar: I eat all of the above | -0.2823 | 6.04E-06 | -0.2111 | 0.0002 | -0.84178 | 0.405762 |
| 20003_1140864992 | Treatment/medication code: tramadol | 0.4957 | 6.11E-06 | 0.3471 | 0.0009 | 0.98173 | 0.690647 |
| 2217 | Age started wearing glasses or contact lenses | 0.2236 | 6.7E-06 | 0.1403 | 0.0067 | 1.160382 | 0.609872 |
| 20002_1113 | Non-cancer illness code, self-reported: emphysema/chronic bronchitis | 0.4372 | 7.49E-06 | 0.2652 | 0.0091 | 1.220862 | 0.717711 |
| 914 | Duration of vigorous activity | 0.3392 | 7.83E-06 | 0.1821 | 0.0284 | 1.395884 | 0.700609 |
| 30140 | Neutrophill count | 0.1734 | 8.88E-06 | 0.1161 | 0.0056 | 1.001019 | 0.576092 |
| 30530 | Sodium in urine | 0.2297 | 9.27E-06 | 0.2611 | 2.14E-06 | -0.4152 | 0.458123 |
| 20151 | Forced vital capacity (FVC), Best measure | -0.1805 | 9.5E-06 | -0.1569 | 2.38E-05 | -0.42796 | 0.4685 |
| 22606_0 | Workplace very noisy: Rarely/never | -0.4984 | 1.3E-05 | -0.283 | 0.0332 | -1.22935 | 0.235329 |
| 20150 | Forced expiratory volume in 1-second (FEV1), Best measure | -0.1812 | 1.42E-05 | -0.1498 | 0.0003 | -0.53501 | 0.458123 |
| VI_NERVOUS | Diseases of the nervous system | 0.4734 | 1.43E-05 | 0.5089 | 2.21E-06 | -0.23178 | 0.452679 |
| 23107 | Impedance of leg (right) | -0.1439 | 1.79E-05 | -0.1682 | 6.41E-06 | 0.484689 | 0.532432 |
| 20003_1140874744 | Treatment/medication code: gliclazide | 0.4095 | 1.81E-05 | 0.2882 | 0.0019 | 0.909968 | 0.657721 |
| 3062 | Forced vital capacity (FVC) | -0.1726 | 1.84E-05 | -0.1619 | 2.53E-05 | -0.19222 | 0.485707 |
| 3063 | Forced expiratory volume in 1-second (FEV1) | -0.1773 | 1.86E-05 | -0.1596 | 0.0001 | -0.30086 | 0.476365 |
| 20003_1140923346 | Treatment/medication code: co-codamol | 0.5415 | 1.94E-05 | 0.5143 | 3.36E-05 | 0.153428 | 0.536292 |
| XIV_GENITOURINARY | Diseases of the genitourinary system | 0.4234 | 1.95E-05 | 0.2376 | 0.0043 | 1.435914 | 0.733119 |
| 4642 | Ever manic/hyper for 2 days | 0.5743 | 2.36E-05 | 0.3899 | 0.0036 | 0.967618 | 0.731575 |
| 6143_4 | Transport type for commuting to job workplace: Cycle | -0.3256 | 3.03E-05 | -0.2567 | 4.86E-05 | -0.68579 | 0.408752 |
| 6154_1 | Medication for pain relief, constipation, heartburn: Aspirin | 0.2952 | 3.23E-05 | 0.2587 | 0.0007 | 0.349961 | 0.548647 |
| 3143 | Ankle spacing width | 0.1433 | 3.93E-05 | 0.1867 | 1.35E-07 | -0.87305 | 0.442216 |
| 5084 | Spherical power (right) | 0.2118 | 3.99E-05 | 0.1838 | 0.0002 | 0.393106 | 0.537357 |
| 20502 | Ever had period extreme irritability | 0.3621 | 4.25E-05 | 0.2033 | 0.0274 | 1.242557 | 0.702584 |
| M13_LOWBACKPAIN | Low back pain | 0.575 | 4.25E-05 | 0.6357 | 6.24E-05 | -0.28628 | 0.419453 |
| 2624 | Frequency of heavy DIY in last 4 weeks | 0.4256 | 4.25E-05 | 0.2656 | 0.015 | 1.061008 | 0.703974 |
| 6162_3 | Types of transport used (excluding work): Public transport | -0.2565 | 4.4E-05 | -0.3544 | 1.37E-07 | 1.063557 | 0.6285 |
| 6138_5 | Qualifications: NVQ or HND or HNC or equivalent | 0.3594 | 4.42E-05 | 0.1703 | 0.0309 | 1.599947 | 0.736739 |
| 5001 | FI7 : synonym | -0.3953 | 4.51E-05 | -0.4313 | 3.52E-05 | 0.252869 | 0.547984 |
| 2405 | Number of children fathered | 0.2533 | 5.67E-05 | 0.483 | 2.88E-17 | -2.70175 | 0.220857 |
| 20516 | Recent restlessness | 0.5341 | 6.25E-05 | 0.575 | 9.39E-07 | -0.23025 | 0.445523 |

| | | | | | | | |
|---|---|---|---|---|---|---|---|
| 22608_1 | Workplace very hot: Sometimes | 0.6267 | 6.41E-05 | 0.3983 | 0.0008 | 1.16174 | 0.777848 |
| 1090 | Time spent driving | 0.2362 | 6.71E-05 | 0.563 | 7.99E-24 | -4.0101 | 0.136867 |
| 20525 | Able to pay rent/mortgage as an adult | -0.6602 | 6.99E-05 | -0.5286 | 0.006 | -0.51788 | 0.329697 |
| 3606 | Chest pain or discomfort walking normally | 0.3068 | 7.18E-05 | 0.3411 | 6.47E-05 | -0.29777 | 0.454272 |
| 6157_3 | Why stopped smoking: Health precaution | -0.3083 | 7.49E-05 | -0.2773 | 0.0013 | -0.26714 | 0.458655 |
| 20527 | Been involved in combat or exposed to war-zone | 0.5244 | 7.5E-05 | 0.5318 | 2.81E-06 | -0.04243 | 0.490114 |
| 23130 | Trunk predicted mass | 0.1356 | 7.86E-05 | 0.1287 | 0.0001 | 0.144756 | 0.509218 |
| 20003_1140868226 | Treatment/medication code: aspirin | 0.2772 | 8.16E-05 | 0.2369 | 0.0019 | 0.387911 | 0.553683 |
| 23129 | Trunk fat-free mass | 0.1351 | 8.62E-05 | 0.1286 | 0.0001 | 0.135961 | 0.508684 |
| 4290 | Duration screen displayed | 0.2442 | 9.29E-05 | 0.1865 | 0.0026 | 0.655942 | 0.576617 |
| 3773 | Knee pain for 3+ months | 0.42 | 0.0001 | 0.3035 | 0.0003 | 0.842441 | 0.651797 |
| 20548_3 | Manifestations of mania or irritability: My thoughts were racing | 0.487 | 0.0001 | 0.4904 | 0.0006 | -0.01768 | 0.495457 |
| G6_CARPTU | Carpal tunnel syndrome | 0.3587 | 0.0001 | 0.3157 | 0.0007 | 0.327643 | 0.557255 |
| 20107_3 | Illnesses of father: Lung cancer | 0.4402 | 0.0001 | 0.3001 | 0.0034 | 0.917422 | 0.680543 |
| 20414 | Frequency of drinking alcohol | -0.28 | 0.0001 | -0.183 | 0.0154 | -0.92363 | 0.372641 |
| R10 | Diagnoses - main ICD10: R10 Abdominal and pelvic pain | 0.3543 | 0.0001 | 0.2458 | 0.0463 | 0.707196 | 0.641841 |
| 1319 | Dried fruit intake | -0.1887 | 0.0002 | -0.2336 | 5.28E-06 | 0.620703 | 0.559766 |
| 5085 | Spherical power (left) | 0.1929 | 0.0002 | 0.211 | 1.2E-05 | -0.25687 | 0.475831 |
| 2463 | Fractured/broken bones in last 5 years | 0.3608 | 0.0002 | 0.4553 | 2.07E-05 | -0.65281 | 0.375813 |
| 6142_2 | Current employment status: Retired | -0.5097 | 0.0002 | -0.487 | 7.24E-05 | -0.12283 | 0.469699 |
| 6159_8 | Pain type(s) experienced in last month: Pain all over the body | 0.5789 | 0.0002 | 0.4898 | 0.0005 | 0.422932 | 0.617305 |
| COPD_EARLYANDLATER | COPD, early/later onset | 0.6482 | 0.0002 | 0.5306 | 0.0006 | 0.503162 | 0.653158 |
| J44 | Diagnoses - main ICD10: J44 Other chronic obstructive pulmonary disease | 0.5856 | 0.0002 | 0.4661 | 0.0006 | 0.580815 | 0.655504 |
| 6150_2 | Vascular/heart problems diagnosed by doctor: Angina | 0.2868 | 0.0002 | 0.2245 | 0.0009 | 0.611301 | 0.58264 |
| 22608_0 | Workplace very hot: Rarely/never | -0.5311 | 0.0002 | -0.3855 | 0.001 | -0.78914 | 0.312904 |
| 22607_0 | Workplace very cold: Rarely/never | -0.5154 | 0.0002 | -0.3846 | 0.0011 | -0.72671 | 0.330668 |
| 20002_1074 | Non-cancer illness code, self-reported: angina | 0.2792 | 0.0002 | 0.2193 | 0.0011 | 0.597828 | 0.5795 |
| 4100 | Ankle spacing width (left) | 0.1549 | 0.0002 | 0.1455 | 0.0014 | 0.151652 | 0.512557 |
| 3571 | Back pain for 3+ months | 0.3983 | 0.0002 | 0.3219 | 0.0014 | 0.523065 | 0.600976 |
| 20018 | Prospective memory result | 0.282 | 0.0002 | 0.2237 | 0.0042 | 0.532939 | 0.577404 |
| 6177_2 | Medication for cholesterol, blood pressure or diabetes: Blood pressure medication | 0.1875 | 0.0002 | 0.1365 | 0.0045 | 0.734285 | 0.567812 |
| 20519 | Recent feelings of tiredness or low energy | 0.2818 | 0.0002 | 0.2116 | 0.0065 | 0.645455 | 0.592939 |
| 30130 | Monocyte count | 0.1381 | 0.0002 | 0.0931 | 0.0133 | 0.85646 | 0.559899 |
| 41215_0 | Detention categories: Informal, not formally detained | 0.7257 | 0.0002 | 0.3567 | 0.0145 | 1.506222 | 0.891741 |
| 2634 | Duration of heavy DIY | 0.2649 | 0.0002 | 0.1985 | 0.0162 | 0.60743 | 0.587992 |
| 2237 | Plays computer games | 0.1709 | 0.0002 | 0.1218 | 0.0265 | 0.691068 | 0.565309 |
| 2277 | Frequency of solarium/sunlamp use | 0.3282 | 0.0003 | 0.5241 | 4E-09 | -1.53303 | 0.25588 |
| 1210 | Snoring | -0.1929 | 0.0003 | -0.2799 | 5.77E-08 | 1.170471 | 0.614618 |
| 20003_1140865634 | Treatment/medication code: omeprazole | 0.4315 | 0.0003 | 0.3338 | 0.0014 | 0.614645 | 0.628246 |
| 20548_2 | Manifestations of mania or irritability: I was more restless than usual | 0.4283 | 0.0003 | 0.3334 | 0.0027 | 0.582675 | 0.624695 |
| 6145_1 | Illness, injury, bereavement, stress in last 2 years: Serious illness, injury or assault to yourself | 0.3122 | 0.0003 | 0.2167 | 0.0086 | 0.799416 | 0.625457 |
| 3436 | Age started smoking in current smokers | -0.5329 | 0.0003 | -0.3148 | 0.0172 | -1.10442 | 0.232557 |
| 20111_8 | Illnesses of siblings: High blood pressure | 0.2813 | 0.0003 | 0.1619 | 0.0225 | 1.133539 | 0.655381 |
| 6160_2 | Leisure/social activities: Pub or social club | 0.2081 | 0.0003 | 0.1334 | 0.0298 | 0.889459 | 0.598777 |
| 1130 | Hands-free device/speakerphone use with mobile phone in last 3 month | 0.2522 | 0.0004 | 0.7605 | 4.4E-54 | -5.85477 | 0.044343 |
| 20077 | Number of diet questionnaires completed | -0.3823 | 0.0004 | -0.3317 | 0.0003 | -0.35748 | 0.432715 |
| 20505 | Recent easy annoyance or irritability | 0.3002 | 0.0004 | 0.2357 | 0.0123 | 0.509965 | 0.585514 |
| 22610_2 | Workplace full of chemical or other fumes: Often | 0.6328 | 0.0004 | 0.4257 | 0.0198 | 0.809699 | 0.756037 |
| 20128 | Number of fluid intelligence questions attempted within time limit | -0.2458 | 0.0004 | -0.1423 | 0.0437 | -1.04919 | 0.364432 |
| I9_IHD | Ischaemic heart disease, wide definition | 0.2136 | 0.0005 | 0.1805 | 0.0006 | 0.410224 | 0.544135 |
| 22615_2 | Workplace had a lot of diesel exhaust: Often | 0.7879 | 0.0005 | 0.4942 | 0.003 | 1.040515 | 0.837355 |

| | | | | | | | |
|---|---|---|---|---|---|---|---|
| 4119 | Ankle spacing width (right) | 0.1488 | 0.0006 | 0.1595 | 0.0008 | -0.16625 | 0.485707 |
| 22610_0 | Workplace full of chemical or other fumes: Rarely/never | -0.4018 | 0.0006 | -0.3074 | 0.0191 | -0.53784 | 0.37594 |
| M23 | Diagnoses - main ICD10: M23 Internal derangement of knee | 0.3672 | 0.0007 | 0.4376 | 0.0001 | -0.44633 | 0.406802 |
| 20003_1140871310 | Treatment/medication code: ibuprofen | 0.3963 | 0.0008 | 0.4982 | 2.94E-05 | -0.6083 | 0.366447 |
| 971 | Frequency of walking for pleasure in last 4 weeks | 0.2365 | 0.0008 | 0.1953 | 0.0107 | 0.395498 | 0.554874 |
| 50 | Standing height | -0.1165 | 0.0008 | -0.0741 | 0.0227 | -0.88909 | 0.443538 |
| 20002_1075 | Non-cancer illness code, self-reported: heart attack/myocardial infarction | 0.2161 | 0.0009 | 0.2065 | 0.001 | 0.106135 | 0.512825 |
| M13_SHOULDER | Shoulder lesions | 0.373 | 0.0009 | 0.3941 | 0.0014 | -0.12614 | 0.471831 |
| M75 | Diagnoses - main ICD10: M75 Shoulder lesions | 0.373 | 0.0009 | 0.3941 | 0.0014 | -0.12614 | 0.471831 |
| 1727 | Ease of skin tanning | -0.1354 | 0.001 | -0.1771 | 0.0017 | 0.596342 | 0.555536 |
| 2207 | Wears glasses or contact lenses | -0.2721 | 0.001 | -0.2102 | 0.006 | -0.55051 | 0.417883 |
| 6151_7 | Fractured bone site(s): Other bones | 0.4619 | 0.001 | 0.4379 | 0.013 | 0.106644 | 0.532032 |
| VII_EYE_ADNEXA | Diseases of the eye and adnexa | 0.294 | 0.0011 | 0.359 | 0.0003 | -0.48618 | 0.413833 |
| 6150_1 | Vascular/heart problems diagnosed by doctor: Heart attack | 0.2152 | 0.0011 | 0.21 | 0.0008 | 0.057164 | 0.506947 |
| 20002_1387 | Non-cancer illness code, self-reported: hayfever/allergic rhinitis | -0.2343 | 0.0011 | -0.1764 | 0.013 | -0.57421 | 0.423121 |
| 1468_5 | Cereal type: Other (e.g. Cornflakes, Frosties) | 0.2118 | 0.0012 | 0.3465 | 1.55E-07 | -1.44751 | 0.325948 |
| 30120 | Lymphocyte count | 0.1455 | 0.0012 | 0.1123 | 0.0046 | 0.555249 | 0.544268 |
| 1369 | Beef intake | 0.2194 | 0.0013 | 0.3553 | 1.51E-08 | -1.46703 | 0.324501 |
| 6154_5 | Medication for pain relief, constipation, heartburn: Omeprazole (e.g. Zanprol) | 0.2994 | 0.0013 | 0.2936 | 0.0021 | 0.043533 | 0.507749 |
| 20517 | Trouble falling or staying asleep, or sleeping too much | 0.2541 | 0.0013 | 0.2363 | 0.0064 | 0.151755 | 0.523769 |
| 6152_9 | Blood clot, DVT, bronchitis, emphysema, asthma, rhinitis, eczema, allergy diagnosed by doctor: Hayfever, allergic rhinitis or eczema | -0.1587 | 0.0015 | -0.2106 | 4.32E-06 | 0.76542 | 0.568997 |
| 100370 | Intake of sugar added to coffee | 0.4058 | 0.0015 | 0.3281 | 0.0114 | 0.427218 | 0.602657 |
| 30500 | Microalbumin in urine | 0.3399 | 0.0015 | 0.2633 | 0.02 | 0.491978 | 0.601235 |
| 20003_1141194794 | Treatment/medication code: bendroflumethiazide | 0.2401 | 0.0015 | 0.1673 | 0.0388 | 0.657076 | 0.596314 |
| 2664_4 | Reason for reducing amount of alcohol drunk: Financial reasons | 0.4174 | 0.0015 | 0.2284 | 0.0391 | 1.09953 | 0.73663 |
| I9_CHD | Major coronary heart disease event | 0.2176 | 0.0017 | 0.2325 | 0.0004 | -0.15524 | 0.4801 |
| I9_CHD_NOREV | Major coronary heart disease event excluding revascularizations | 0.2176 | 0.0017 | 0.2325 | 0.0004 | -0.15524 | 0.4801 |
| 2227 | Other eye problems | 0.4108 | 0.0017 | 0.2587 | 0.0245 | 0.872931 | 0.694766 |
| 3526 | Mother's age at death | -0.7305 | 0.0018 | -0.3373 | 0.028 | -1.40334 | 0.093938 |
| 767 | Length of working week for main job | 0.2372 | 0.0019 | 0.7252 | 1.36E-18 | -4.34285 | 0.051089 |
| 1389 | Pork intake | 0.2147 | 0.0019 | 0.1704 | 0.0151 | 0.450379 | 0.558974 |
| 796 | Distance between home and job workplace | -0.4032 | 0.002 | 0.3491 | 0.0207 | -3.76718 | 0.005875 |
| I9_MI | Myocardial infarction | 0.2329 | 0.0021 | 0.2322 | 0.0015 | 0.00663 | 0.500935 |
| I9_MI_STRICT | Myocardial infarction, strict | 0.2329 | 0.0021 | 0.2322 | 0.0015 | 0.00663 | 0.500935 |
| 22502 | Cough on most days | 0.3389 | 0.0021 | 0.2364 | 0.0489 | 0.62913 | 0.634309 |
| M13_MENISCUSDERANGEMENTS | Meniscus derangement | 0.3594 | 0.0023 | 0.4586 | 0.0002 | -0.57782 | 0.369856 |
| 4294_1 | Final attempt correct: yes | -0.3764 | 0.0023 | -0.2406 | 0.0256 | -0.82802 | 0.324621 |
| 20002_1223 | Non-cancer illness code, self-reported: type 2 diabetes | 0.3591 | 0.0023 | 0.2327 | 0.0343 | 0.783538 | 0.663974 |
| K21 | Diagnoses - main ICD10: K21 Gastro-oesophageal reflux disease | 0.5545 | 0.0025 | 0.425 | 0.0014 | 0.57062 | 0.667752 |
| 20153 | Forced expiratory volume in 1-second (FEV1), predicted | -0.1455 | 0.0025 | -0.0867 | 0.0405 | -0.91798 | 0.421941 |
| M13_DORSALGIANAS | Other/unspecified dorsalgia | 0.4775 | 0.0026 | 0.3707 | 0.0091 | 0.501358 | 0.639712 |
| 3393 | Hearing aid user | 0.3504 | 0.0028 | 0.2339 | 0.0412 | 0.711033 | 0.651797 |
| K63 | Diagnoses - main ICD10: K63 Other diseases of intestine | 0.4007 | 0.0029 | 0.4946 | 0.0019 | -0.45027 | 0.376576 |
| 3731 | Former alcohol drinker | 0.4859 | 0.0029 | 0.4747 | 0.0021 | 0.049885 | 0.514961 |

| | | | | | | | |
|---|---|---|---|---|---|---|---|
| I21 | Diagnoses - main ICD10: I21 Acute myocardial infarction | 0.2323 | 0.0033 | 0.2771 | 0.0002 | -0.4149 | 0.440366 |
| I20 | Diagnoses - main ICD10: I20 Angina pectoris | 0.2766 | 0.0033 | 0.2318 | 0.0183 | 0.329226 | 0.559634 |
| 6160_4 | Leisure/social activities: Adult education class | -0.3711 | 0.0034 | -0.5928 | 1.67E-05 | 1.185654 | 0.77111 |
| 20508 | Recent trouble concentrating on things | 0.2884 | 0.0035 | 0.3041 | 0.0028 | -0.11084 | 0.479033 |
| 4291 | Number of attempts | 0.2261 | 0.0035 | 0.2138 | 0.007 | 0.110929 | 0.51643 |
| 1031 | Frequency of friend/family visits | -0.1875 | 0.0036 | -0.4114 | 2.69E-09 | 2.370385 | 0.773335 |
| I84 | Diagnoses - main ICD10: I84 Haemorrhoids | 0.2853 | 0.0037 | 0.2964 | 0.0017 | -0.08149 | 0.485172 |
| 22702 | Home location - east co-ordinate (rounded) | -0.3178 | 0.0038 | -0.3347 | 0.0029 | 0.107651 | 0.522568 |
| 20003_1140860806 | Treatment/medication code: ramipril | 0.2636 | 0.0038 | 0.2119 | 0.0292 | 0.387886 | 0.568734 |
| 6154_2 | Medication for pain relief, constipation, heartburn: Ibuprofen (e.g. Nurofen) | 0.2886 | 0.0039 | 0.4811 | 9.42E-06 | -1.30396 | 0.259557 |
| 20491 | Someone to take to doctor when needed as a child | -0.3767 | 0.0039 | -0.4096 | 0.0043 | 0.169801 | 0.54387 |
| 20497 | Repeated disturbing thoughts of stressful experience in past month | 0.2885 | 0.004 | 0.2843 | 0.0019 | 0.030966 | 0.505612 |
| 4294_0 | Final attempt correct: no | 0.3737 | 0.0043 | 0.2724 | 0.0162 | 0.585387 | 0.632797 |
| 20512 | Recent feelings of foreboding | 0.2786 | 0.0044 | 0.1791 | 0.0269 | 0.783066 | 0.630524 |
| D12 | Diagnoses - main ICD10: D12 Benign neoplasm of colon, rectum, anus and anal canal | 0.2802 | 0.0045 | 0.1945 | 0.0364 | 0.632627 | 0.612952 |
| G47 | Diagnoses - main ICD10: G47 Sleep disorders | 0.2687 | 0.0048 | 0.2893 | 0.0065 | -0.14429 | 0.472498 |
| 6154_4 | Medication for pain relief, constipation, heartburn: Ranitidine (e.g. Zantac) | 0.4257 | 0.0048 | 0.2732 | 0.0441 | 0.751172 | 0.695236 |
| 1359 | Poultry intake | 0.1771 | 0.0049 | 0.3073 | 3.72E-05 | -1.33536 | 0.331397 |
| 6157_2 | Why stopped smoking: Doctor's advice | 0.4209 | 0.005 | 0.3219 | 0.0165 | 0.491895 | 0.629892 |
| 6157_100 | Why stopped smoking: None of the above | 0.2094 | 0.0051 | 0.3091 | 0.0003 | -0.87234 | 0.369224 |
| K29 | Diagnoses - main ICD10: K29 Gastritis and duodenitis | 0.5864 | 0.0051 | 0.5542 | 0.0011 | 0.119585 | 0.54294 |
| E4_DM2 | Type 2 diabetes | 0.5665 | 0.0051 | 0.7719 | 0.006 | -0.59384 | 0.245753 |
| SLEEP | Sleep disorders (combined) | 0.2633 | 0.0051 | 0.289 | 0.0062 | -0.18197 | 0.465704 |
| 1767 | Adopted as a child | 0.5239 | 0.0052 | 0.6694 | 0.0026 | -0.50054 | 0.313022 |
| PULM_MEDICATIO_COMORB | Medication related adverse effects (Asthma/COPD) | 0.2713 | 0.0057 | 0.2307 | 0.0181 | 0.293392 | 0.55408 |
| I9_CORATHER | Coronary atherosclerosis | 0.1771 | 0.006 | 0.1703 | 0.0026 | 0.079251 | 0.509085 |
| 1747_6 | Hair colour (natural, before greying): Other | 0.3183 | 0.006 | 0.274 | 0.0055 | 0.291273 | 0.558974 |
| 20002_1452 | Non-cancer illness code, self-reported: eczema/dermatitis | -0.2341 | 0.0062 | -0.2573 | 0.0069 | 0.181309 | 0.530967 |
| 6160_1 | Leisure/social activities: Sports club or gym | -0.1602 | 0.0064 | 0.1231 | 0.0418 | -3.36075 | 0.171357 |
| 4979 | FI5 : family relationship calculation | -0.2939 | 0.0066 | -0.3591 | 0.0006 | 0.433447 | 0.586428 |
| 22617_8211 | Job SOC coding: Heavy goods vehicle drivers | 0.4843 | 0.0066 | 0.5356 | 0.0021 | -0.20621 | 0.431793 |
| E11 | Diagnoses - main ICD10: E11 Non-insulin-dependent diabetes mellitus | 0.5672 | 0.0067 | 0.7573 | 0.0063 | -0.54752 | 0.262169 |
| R31 | Diagnoses - main ICD10: R31 Unspecified haematuria | 0.439 | 0.0067 | 0.2616 | 0.0352 | 0.869386 | 0.723791 |
| 2664_3 | Reason for reducing amount of alcohol drunk: Health precaution | -0.2366 | 0.0073 | -0.2829 | 0.0002 | 0.399275 | 0.561615 |
| 680_4 | Own or rent accommodation lived in: Rent - from private landlord or letting agency | 0.5474 | 0.0074 | 0.4528 | 0.0297 | 0.324232 | 0.624314 |
| 20426 | Restless during period of worst anxiety | 0.4794 | 0.0076 | 0.468 | 0.0064 | 0.04588 | 0.515228 |
| 20074 | Home location at assessment - east co-ordinate (rounded) | -0.3147 | 0.0076 | -0.3137 | 0.008 | -0.00599 | 0.498664 |
| ASTHMA_MEDICATIO_COMORB | Medication related adverse effects | 0.2479 | 0.0076 | 0.1893 | 0.0393 | 0.448681 | 0.577797 |
| 6149_5 | Mouth/teeth dental problems: Toothache | 0.4722 | 0.0077 | 0.3247 | 0.0272 | 0.640682 | 0.689347 |
| 30510 | Creatinine (enzymatic) in urine | 0.1386 | 0.0081 | 0.2565 | 1.44E-07 | -1.64823 | 0.346471 |
| COLITNONINFNAS | Noninfectious colitis NAS | 0.7083 | 0.0082 | 0.5651 | 0.0153 | 0.403251 | 0.684244 |
| 20536_1 | Weight change during worst episode of depression: Gained weight | 0.3771 | 0.0083 | 0.3766 | 0.006 | 0.002525 | 0.500668 |
| 20528 | Diagnosed with life-threatening illness | 0.3817 | 0.0095 | 0.4368 | 0.0089 | -0.24767 | 0.426796 |

| | | | | | | | |
|---|---|---|---|---|---|---|---|
| 20154 | Forced expiratory volume in 1-second (FEV1), predicted percentage | -0.1344 | 0.0099 | -0.1719 | 0.0024 | 0.487467 | 0.549973 |
| M13_SPONDYLOPATHY | #Spondylopathies | 0.353 | 0.01 | 0.326 | 0.012 | 0.14301 | 0.536026 |
| 1598 | Average weekly spirits intake | 0.1834 | 0.0104 | 0.2271 | 0.0012 | -0.43581 | 0.441819 |
| 20536_0 | Weight change during worst episode of depression: Stayed about the same or was on a diet | -0.3204 | 0.0106 | -0.4717 | 3.71E-05 | 0.892093 | 0.693827 |
| XXI_HEALTHFACTORS | Factors influencing health status and contact with health services | 0.8232 | 0.0114 | 0.4287 | 0.0335 | 1.031051 | 0.90679 |
| 1508_4 | Coffee type: Other type of coffee | 0.2791 | 0.0115 | 0.3156 | 0.0173 | -0.21146 | 0.451353 |
| H26 | Diagnoses - main ICD10: H26 Other cataract | 0.2521 | 0.012 | 0.2354 | 0.0173 | 0.118558 | 0.522302 |
| M13_JOINTOTH | Other specific joint derangements/joint disorders | 0.9456 | 0.0124 | 0.8063 | 0.0168 | 0.275 | 0.679584 |
| 2907 | Ever stopped smoking for 6+ months | 0.6704 | 0.0124 | 0.4741 | 0.0243 | 0.575758 | 0.744551 |
| 3404 | Neck/shoulder pain for 3+ months | 0.6937 | 0.0125 | 0.5438 | 0.0136 | 0.422978 | 0.69218 |
| J34 | Diagnoses - main ICD10: J34 Other disorders of nose and nasal sinuses | 0.391 | 0.0128 | 0.4975 | 0.0288 | -0.38518 | 0.360664 |
| 22601_82112603 | Job coding: heavy goods vehicle (hgv) driver, lorry or truck driver, tanker driver, haulage driver | 0.4535 | 0.0131 | 0.5139 | 0.0025 | -0.2419 | 0.419845 |
| 6139_2 | Gas or solid-fuel cooking/heating: A gas fire that you use regularly in winter time | 0.2834 | 0.0144 | 0.3061 | 0.009 | -0.13784 | 0.469699 |
| I24 | Diagnoses - main ICD10: I24 Other acute ischaemic heart diseases | 0.3393 | 0.0147 | 0.3066 | 0.0448 | 0.158304 | 0.543604 |
| 20002_1138 | Non-cancer illness code, self-reported: gastro-oesophageal reflux (gord) / gastric reflux | 0.2499 | 0.0149 | 0.2892 | 0.0054 | -0.26914 | 0.447642 |
| 20003_1140879802 | Treatment/medication code: amlodipine | 0.1945 | 0.0149 | 0.1963 | 0.0076 | -0.01658 | 0.497595 |
| 22616_1 | Breathing problems during period of job: Yes | 0.4555 | 0.0152 | 0.3774 | 0.023 | 0.311778 | 0.603173 |
| 6145_3 | Illness, injury, bereavement, stress in last 2 years: Death of a close relative | 0.5599 | 0.0153 | 0.4148 | 0.0041 | 0.5328 | 0.686503 |
| K60 | Diagnoses - main ICD10: K60 Fissure and fistula of anal and rectal regions | 0.3051 | 0.0155 | 0.2672 | 0.0305 | 0.214814 | 0.550503 |
| 4674 | Private healthcare | 0.3194 | 0.0155 | -0.235 | 0.0378 | 3.188196 | 0.968328 |
| 20110_6 | Illnesses of mother: Chronic bronchitis/emphysema | 0.5483 | 0.016 | 0.6222 | 0.0021 | -0.24263 | 0.40226 |
| 1970 | Nervous feelings | -0.1351 | 0.0163 | -0.3068 | 2.39E-06 | 1.998209 | 0.717371 |
| 20498 | Felt very upset when reminded of stressful experience in past month | 0.2527 | 0.0164 | 0.2807 | 0.0089 | -0.18634 | 0.462643 |
| I25 | Diagnoses - main ICD10: I25 Chronic ischaemic heart disease | 0.1514 | 0.0173 | 0.1731 | 0.0022 | -0.25468 | 0.471032 |
| K11_OTHGASTR | Other gastritis (incl. Duodenitis) | 0.5331 | 0.0178 | 0.6525 | 0.0049 | -0.36986 | 0.344619 |
| S09 | Diagnoses - main ICD10: S09 Other and unspecified injuries of head | 0.3022 | 0.018 | 0.3833 | 0.0153 | -0.39893 | 0.392959 |
| 20003_1140879406 | Treatment/medication code: ranitidine | 0.3258 | 0.0181 | 0.3197 | 0.0182 | 0.031564 | 0.50815 |
| 3064 | Peak expiratory flow (PEF) | -0.1101 | 0.0184 | -0.1045 | 0.0388 | -0.08133 | 0.492518 |
| G6_EPIPAROX | Episodal and paroxysmal disorders | 0.2278 | 0.0186 | 0.359 | 0.0007 | -0.9121 | 0.330183 |
| M13_SYNOTEND | Disorders of synovium and tendon | 0.3033 | 0.0186 | 0.4278 | 0.0007 | -0.68882 | 0.33835 |
| 20513 | Recent thoughts of suicide or self-harm | 0.4833 | 0.0188 | 0.4989 | 0.0299 | -0.05058 | 0.479166 |
| M67 | Diagnoses - main ICD10: M67 Other disorders of synovium and tendon | 0.4394 | 0.019 | 0.4178 | 0.0366 | 0.078831 | 0.528835 |
| M25 | Diagnoses - main ICD10: M25 Other joint disorders, not elsewhere classified | 0.8963 | 0.0197 | 0.7376 | 0.018 | 0.320583 | 0.702468 |
| 6152_5 | Blood clot, DVT, bronchitis, emphysema, asthma, rhinitis, eczema, allergy diagnosed by doctor: Blood clot in the leg (DVT) | 0.2461 | 0.0224 | 0.275 | 0.0076 | -0.19374 | 0.461446 |
| H7_LENS | Disorders of lens | 0.2021 | 0.023 | 0.2295 | 0.0109 | -0.21647 | 0.463442 |
| COX_ARTHROSIS | Coxarthrosis [arthrosis of hip](FG) | 0.1831 | 0.0234 | 0.2786 | 0.005 | -0.74643 | 0.374543 |
| M16 | Diagnoses - main ICD10: M16 Coxarthrosis [arthrosis of hip] | 0.1831 | 0.0234 | 0.2786 | 0.005 | -0.74643 | 0.374543 |
| 20003_1141168318 | Treatment/medication code: clopidogrel | 0.329 | 0.0235 | 0.3152 | 0.0238 | 0.068512 | 0.518432 |

| | | | | | | | |
|---|---|---|---|---|---|---|---|
| II_NEOPLASM | Neoplasms | 0.2079 | 0.0237 | 0.3422 | 0.0003 | -1.0205 | 0.32643 |
| 3456 | Number of cigarettes currently smoked daily (current cigarette smokers) | 0.4774 | 0.0239 | 0.3561 | 0.0255 | 0.458149 | 0.657721 |
| 20515 | Recent trouble relaxing | 0.1684 | 0.0241 | 0.1975 | 0.0093 | -0.27326 | 0.46118 |
| I73 | Diagnoses - main ICD10: I73 Other peripheral vascular diseases | 0.3768 | 0.0251 | 0.4515 | 0.0281 | -0.28106 | 0.401223 |
| K52 | Diagnoses - main ICD10: K52 Other non-infective gastro-enteritis and colitis | 0.7653 | 0.0254 | 0.5765 | 0.0333 | 0.432504 | 0.736411 |
| 22611_1 | Workplace had a lot of cigarette smoke from other people smoking: Sometimes | 0.3454 | 0.0258 | 0.4111 | 0.003 | -0.31608 | 0.41292 |
| 1980 | Worrier / anxious feelings | -0.1048 | 0.0259 | -0.1527 | 0.0193 | 0.594928 | 0.563727 |
| I9_PAD | Peripheral artery disease | 0.5455 | 0.026 | 0.4982 | 0.0404 | 0.137017 | 0.562935 |
| 100170 | Fizzy drink intake | 0.3544 | 0.0262 | 0.3485 | 0.0219 | 0.026779 | 0.507883 |
| 20002_1094 | Non-cancer illness code, self-reported: deep venous thrombosis (dvt) | 0.2302 | 0.0265 | 0.2962 | 0.0032 | -0.45681 | 0.412529 |
| XII_SKIN_SUBCUTAN | Diseases of the skin and subcutaneous tissue | 0.3355 | 0.0282 | 0.5565 | 0.0014 | -0.95471 | 0.229601 |
| 1180 | Morning/evening person (chronotype) | -0.0844 | 0.0318 | -0.1033 | 0.0246 | 0.312386 | 0.525236 |
| K44 | Diagnoses - main ICD10: K44 Diaphragmatic hernia | 0.3889 | 0.032 | 0.5644 | 0.0045 | -0.65264 | 0.278341 |
| 22601_21133023 | Job coding: physicist, astronomer, geologist, geophysicist, meteorologist, oceanographer, seismologist | -0.3986 | 0.0338 | -0.4482 | 0.0229 | 0.182191 | 0.565968 |
| 22617_2311 | Job SOC coding: Higher education teaching professionals | -0.2547 | 0.0339 | -0.5023 | 0.0011 | 1.269878 | 0.796519 |
| M13_ROTATORCUFF | Rotator cuff syndrome | 0.3062 | 0.0342 | 0.3022 | 0.0285 | 0.020012 | 0.505344 |
| 6145_5 | Illness, injury, bereavement, stress in last 2 years: Marital separation/divorce | 0.5269 | 0.0356 | 0.9968 | 0.0255 | -0.91797 | 0.057771 |
| 22617_3311 | Job SOC coding: NCOs and other ranks | 0.8869 | 0.0393 | 0.5646 | 0.0054 | 0.677535 | 0.859802 |
| 22601_23113399 | Job coding: higher education teaching professional, university lecturer/professor (including college/university head/vice chancellor) | -0.2175 | 0.0396 | -0.409 | 0.0044 | 1.074475 | 0.739357 |
| 6162_1 | Types of transport used (excluding work): Car/motor vehicle | -0.1468 | 0.0408 | 0.3392 | 1.71E-07 | -5.02146 | 0.051795 |
| 20509 | Recent inability to stop or control worrying | 0.2227 | 0.041 | 0.2329 | 0.0259 | -0.06758 | 0.486374 |
| IV_ENDOCRIN_NUTRIT | Endocrine, nutritional and metabolic diseases | 0.7104 | 0.0436 | 0.5509 | 0.0222 | 0.373866 | 0.703395 |
| 6141_6 | How are people in household related to participant: Grandchild | 0.4643 | 0.0441 | 0.3769 | 0.0222 | 0.30836 | 0.61513 |
| 22617_5314 | Job SOC coding: Plumbers, heating and ventilating engineers | 0.3193 | 0.0461 | 0.4141 | 0.0147 | -0.40621 | 0.375432 |

**Table S27.** Differential gene set enrichment between autism spectrum disorder (ASD) and *PhoneUse* surviving Bonferroni correction ($p<4.69x10^{-6}$).

| Enrichment | N | ASD | | *PhoneUse* | | Difference | |
|---|---|---|---|---|---|---|---|
| | Genes | beta | P | beta | p | z | p |
| GO midgut development | 14 | -0.49 | 0.96387 | 1.01 | 0.00016 | -3.8427 | 3.703E-18 |
| Response to paclitaxel via mapk8 down (Want) | 5 | -0.46 | 0.8363 | 1.02 | 0.00814 | -2.3356 | 1.001E-17 |
| GO sphinGOid metabolic process | 13 | -0.796 | 0.99887 | 0.594 | 0.00924 | -3.8313 | 7.487E-16 |
| GO diol metabolic process | 11 | -0.677 | 0.99221 | 0.646 | 0.0078 | -3.4195 | 1.567E-14 |
| Response to sulindac up (Zerbini) | 9 | -0.243 | 0.76244 | 1.05 | 0.00036 | -2.8098 | 5.835E-14 |
| GO positive regulation of neuron migration | 10 | -0.278 | 0.84092 | 0.978 | 0.00261 | -2.8061 | 2.836E-13 |
| GO aspartate metabolic process | 11 | -0.582 | 0.99293 | 0.656 | 0.00466 | -3.5787 | 6.023E-13 |
| GO dna polymerase binding | 12 | -0.442 | 0.93006 | 0.792 | 0.00028 | -3.2765 | 7.11E-13 |
| chen hoxa5 targets 6hr up | 10 | -0.477 | 0.94478 | 0.694 | 0.014 | -2.6917 | 9.066E-12 |
| GO magnesium ion transport | 12 | -0.133 | 0.66252 | 0.975 | 0.00077 | -2.5069 | 1.017E-10 |
| kreppel cd99 targets up | 5 | -0.297 | 0.7889 | 0.792 | 0.01891 | -2.0505 | 2.056E-10 |
| GO cell migration involved in heart development | 14 | -0.596 | 0.99525 | 0.412 | 0.04465 | -3.0127 | 3.628E-09 |
| Il6 signaling scar down (Dasu) | 16 | -0.534 | 0.98382 | 0.44 | 0.03956 | -2.7549 | 1.137E-08 |
| Carcinogenesis by kras and stk11 up (Ji) | 12 | -0.542 | 0.98404 | 0.421 | 0.03466 | -2.8114 | 1.632E-08 |
| Liver cancer metastasis up (Budhu) | 8 | -0.367 | 0.88964 | 0.593 | 0.0393 | -2.1277 | 1.8E-08 |
| GO hemidesmosome assembly | 12 | -0.179 | 0.76522 | 0.768 | 0.00411 | -2.4818 | 2.743E-08 |
| Reactome sphinGOlipid de novo biosynthesis | 30 | -0.475 | 0.99763 | 0.439 | 0.00716 | -3.7232 | 7.796E-08 |
| GO membrane repolarization | 15 | -0.251 | 0.84961 | 0.658 | 0.00577 | -2.5592 | 9.106E-08 |
| GO host | 12 | -0.349 | 0.91698 | 0.556 | 0.01005 | -2.6057 | 1.03E-07 |
| Muc4 targets up (Bafna) | 6 | -0.255 | 0.76233 | 0.648 | 0.04625 | -1.7198 | 1.096E-07 |
| Escape from anoikis (Yan) | 20 | -0.541 | 0.99653 | 0.336 | 0.03709 | -3.195 | 2.413E-07 |
| Sp3 targets up (Vanloo) | 8 | -0.181 | 0.69932 | 0.694 | 0.02925 | -1.7324 | 2.562E-07 |
| Pten targets down (He) | 7 | 0.0681 | 0.4255 | 0.932 | 0.00836 | -1.6258 | 3.563E-07 |
| GO negative regulation of skeletal muscle tissue development | 13 | -0.212 | 0.82562 | 0.65 | 0.00625 | -2.5022 | 3.768E-07 |
| Reactome hyaluronan uptake and degradation | 10 | -0.34 | 0.93265 | 0.52 | 0.02601 | -2.454 | 3.997E-07 |
| GO cardiac cell fate commitment | 10 | -0.229 | 0.80628 | 0.6 | 0.02135 | -2.0866 | 9.795E-07 |
| Hras targets down (Castellano) | 6 | 0.0587 | 0.42053 | 0.886 | 0.00249 | -1.923 | 1.028E-06 |
| Reactome activation of the ap1 family of transcription factors | 10 | -0.233 | 0.79283 | 0.593 | 0.02789 | -1.9615 | 1.067E-06 |
| GO s100 protein binding | 12 | -0.284 | 0.8596 | 0.53 | 0.02185 | -2.1844 | 1.495E-06 |
| Reactome crmps in sema3a signaling | 14 | -0.14 | 0.73433 | 0.668 | 0.0025 | -2.4774 | 1.767E-06 |
| Reactome homologous recombination repair of replication independent double strand breaks | 15 | -0.382 | 0.95433 | 0.421 | 0.03545 | -2.4685 | 2.029E-06 |
| EGF response 40 mcf10a (Amit) | 19 | -0.412 | 0.97336 | 0.389 | 0.04328 | -2.5732 | 2.144E-06 |

**Table S28.** Linkage disequilibrium score regression (rg) results for 1,050 UK Biobank traits demonstrating nominally significant genetic correlations with autism spectrum disorder (ASD) and/or *PhoneUse*. Traits are listed in ascending order by most significant difference between ASD and *CompGaming*.

| Field ID | Trait | ASD | | *PhoneUse* | | Difference | |
|---|---|---|---|---|---|---|---|
| | | rg | p | rg | p | z | p |
| M13_LIMBPAIN | Pain in limb | -0.3665 | 0.134 | 0.7301 | 0.0017 | -3.24484 | 2.84E-08 |
| BRONCHITIS | Bronchitis | -0.4946 | 0.2152 | 0.5815 | 0.0035 | -2.41371 | 4.99E-08 |
| M13_JOINTOTH | Other specific joint derangements/joint disorders | -0.202 | 0.1249 | 0.7876 | 9.63E-09 | -5.2015 | 4.82E-07 |
| M25 | Diagnoses - main ICD10: M25 Other joint disorders, not elsewhere classified | -0.191 | 0.148 | 0.7912 | 1.94E-08 | -5.08722 | 5.81E-07 |
| 20003_1141168684 | Treatment/medication code: xenical 120mg capsule | -0.4219 | 0.2192 | 0.5359 | 0.0377 | -2.23055 | 1.06E-06 |
| 6145_5 | Illness, injury, bereavement, stress in last 2 years: Marital separation/divorce | -0.2452 | 0.0579 | 0.6656 | 1.37E-05 | -4.54503 | 3.26E-06 |
| M13_SOFTTISSUEOTH | Other soft tissue disorders, not elsewhere classified | -0.2163 | 0.1711 | 0.6912 | 2.97E-11 | -4.79763 | 3.52E-06 |
| M79 | Diagnoses - main ICD10: M79 Other soft tissue disorders, not elsewhere classified | -0.2163 | 0.1711 | 0.6912 | 2.97E-11 | -4.79763 | 3.52E-06 |
| 3741 | Stomach/abdominal pain for 3+ months | -0.3398 | 0.0893 | 0.5019 | 0.0013 | -3.32002 | 1.55E-05 |
| J34 | Diagnoses - main ICD10: J34 Other disorders of nose and nasal sinuses | -0.3763 | 0.0065 | 0.4236 | 0.0046 | -3.93178 | 3.75E-05 |
| 20523 | Physical violence by partner or ex-partner as an adult | -0.3737 | 1.26E-06 | 0.4241 | 6.13E-12 | -8.0791 | 3.92E-05 |
| 20002_99999 | Non-cancer illness code, self-reported: unclassifiable | -0.4785 | 0.0564 | 0.3139 | 0.0306 | -2.73512 | 4.38E-05 |
| K59 | Diagnoses - main ICD10: K59 Other functional intestinal disorders | -0.3029 | 0.0858 | 0.4714 | 0.0061 | -3.14457 | 6.33E-05 |
| 20548_5 | Manifestations of mania or irritability: I needed less sleep than usual | -0.3604 | 0.0003 | 0.3995 | 1.33E-05 | -5.59159 | 8.44E-05 |
| R06 | Diagnoses - main ICD10: R06 Abnormalities of breathing | -0.2546 | 0.1027 | 0.5044 | 0.0038 | -3.24683 | 8.59E-05 |
| R13 | Diagnoses - main ICD10: R13 Dysphagia | 0.0835 | 0.5311 | 0.8313 | 0.0047 | -2.3156 | 0.000107 |
| C_STROKE | STROKE | -0.2938 | 0.116 | 0.4411 | 0.0059 | -2.98701 | 0.000137 |
| R51 | Diagnoses - main ICD10: R51 Headache | -0.1005 | 0.5393 | 0.6305 | 9.25E-07 | -3.51123 | 0.000148 |
| K30 | Diagnoses - main ICD10: K30 Dyspepsia | -0.1868 | 0.1338 | 0.5391 | 1.58E-05 | -4.11454 | 0.000163 |
| K63 | Diagnoses - main ICD10: K63 Other diseases of intestine | -0.1508 | 0.2606 | 0.5728 | 0.0002 | -3.52787 | 0.00017 |
| 20521 | Belittlement by partner or ex-partner as an adult | -0.4577 | 9.62E-15 | 0.2655 | 4.06E-08 | -9.46726 | 0.000172 |
| PULMONARYDG | Other pulmonary diagnosis | -0.1701 | 0.0429 | 0.5488 | 9.45E-22 | -7.07006 | 0.000186 |
| X_RESPIRATORY | Diseases of the respiratory system | -0.1701 | 0.0429 | 0.5488 | 9.45E-22 | -7.07006 | 0.000186 |
| 22613_2 | Worked with paints, thinners or glues: Often | -0.3463 | 0.0911 | 0.3696 | 0.0032 | -2.98074 | 0.000197 |
| R04 | Diagnoses - main ICD10: R04 Haemorrhage from respiratory passages | -0.1995 | 0.2143 | 0.5122 | 8.83E-05 | -3.43689 | 0.000213 |
| 22610_2 | Workplace full of chemical or other fumes: Often | -0.1969 | 0.2026 | 0.5143 | 4.92E-05 | -3.55942 | 0.000215 |
| G43 | Diagnoses - main ICD10: G43 Migraine | -0.4472 | 0.0443 | 0.2602 | 0.0369 | -2.77535 | 0.000231 |
| 20003_1140865716 | Treatment/medication code: senna | -0.0237 | 0.8837 | 0.6834 | 0.0022 | -2.56278 | 0.000232 |
| 20518 | Recent changes in speed/amount of moving or speaking | -0.3933 | 5.84E-06 | 0.312 | 6.9E-06 | -6.34644 | 0.00024 |
| COPD_EXCL | COPD differential diagnosis | -0.1582 | 0.0483 | 0.5446 | 4.96E-23 | -7.22885 | 0.000252 |
| ILD_DIFF_DG | ILD differential diagnosis | -0.1582 | 0.0483 | 0.5446 | 4.96E-23 | -7.22885 | 0.000252 |
| E4_DM2NOCOMP | Type 2 diabetes without complications | -0.1496 | 0.3591 | 0.5521 | 0.0059 | -2.71344 | 0.000257 |
| XVIII_MISCFINDINGS | Symptoms, signs and abnormal clinical and laboratory findings, not elsewhere classified | -0.1661 | 0.0015 | 0.5343 | 1.24E-58 | -11.3006 | 0.000263 |
| N23 | Diagnoses - main ICD10: N23 Unspecified renal colic | -0.1967 | 0.348 | 0.5002 | 0.0406 | -2.16552 | 0.00028 |
| XXI_HEALTHFACTORS | Factors influencing health status and contact with health services | -0.1896 | 0.0354 | 0.507 | 3.14E-11 | -5.89684 | 0.000282 |
| 20553_4 | Methods of self-harm used: Ingesting a medication in excess of the normal dose | -0.4624 | 0.0001 | 0.2298 | 0.0152 | -4.55148 | 0.000306 |

| | | | | | | | |
|---|---|---|---|---|---|---|---|
| 20540 | Multiple worries during worst period of anxiety | -0.4964 | 0.0191 | 0.1844 | 0.1125 | -2.8181 | 0.000376 |
| 4642 | Ever manic/hyper for 2 days | -0.2348 | 0.0071 | 0.4444 | 2.3E-08 | -5.75591 | 0.000387 |
| 6157_2 | Why stopped smoking: Doctor's advice | -0.2284 | 0.1224 | 0.4481 | 2.43E-06 | -3.84917 | 0.000406 |
| 2844 | Had other major operations | -0.2679 | 0.0016 | 0.407 | 9.18E-17 | -6.87628 | 0.000417 |
| 6155_6 | Vitamin and mineral supplements: Folic acid or Folate (Vit B9) | -0.4173 | 0.0472 | 0.2563 | 0.1054 | -2.55907 | 0.000427 |
| 20548_2 | Manifestations of mania or irritability: I was more restless than usual | -0.3315 | 2.20E-05 | 0.342 | 3.64E-08 | -6.74986 | 0.000428 |
| 22605 | Work hours per week - exact value | -0.0846 | 0.615 | 0.5865 | 0.0008 | -2.76238 | 0.000447 |
| 20526 | Been in serious accident believed to be life-threatening | -0.4089 | 0.0004 | 0.2621 | 0.0008 | -4.81052 | 0.000447 |
| 6145_4 | Illness, injury, bereavement, stress in last 2 years: Death of a spouse or partner | 0.0001 | 0.9997 | 0.6701 | 0.0048 | -1.90023 | 0.000455 |
| IV_ENDOCRIN_NUTRIT | Endocrine, nutritional and metabolic diseases | -0.2075 | 0.1801 | 0.4606 | 8.88E-06 | -3.58568 | 0.000471 |
| T81 | Diagnoses - main ICD10: T81 Complications of procedures, not elsewhere classified | -0.1892 | 0.1356 | 0.4759 | 2.66E-05 | -3.91133 | 0.000497 |
| 41235 | Spells in hospital | -0.1496 | 0.0069 | 0.5149 | 1.44E-40 | -9.84135 | 0.000502 |
| 20002_1406 | Non-cancer illness code, self-reported: muscle or soft tissue injuries | -0.158 | 0.5148 | 0.5053 | 0.0265 | -1.99357 | 0.000513 |
| 20516 | Recent restlessness | -0.2971 | 7.21E-05 | 0.3659 | 1.47E-10 | -7.04544 | 0.000515 |
| M24 | Diagnoses - main ICD10: M24 Other specific joint derangements | 0.0698 | 0.687 | 0.7276 | 0.0071 | -2.04795 | 0.000564 |
| KRA_PSY_PERSON | Personality disorders | -0.3455 | 0.0958 | 0.3097 | 0.0219 | -2.64705 | 0.000591 |
| 20487 | Felt hated by family member as a child | -0.3736 | 2.30E-08 | 0.2815 | 5.17E-09 | -7.94493 | 0.000592 |
| 41248_1000 | Destinations on discharge from hospital (recoded): Usual Place of residence | -0.0868 | 0.1289 | 0.5678 | 3.56E-43 | -9.28601 | 0.000597 |
| 6145_6 | Illness, injury, bereavement, stress in last 2 years: Financial difficulties | -0.1436 | 0.0056 | 0.5104 | 8.96E-52 | -10.5686 | 0.000603 |
| 6150_3 | Vascular/heart problems diagnosed by doctor: Stroke | -0.2352 | 0.1084 | 0.4178 | 2.98E-05 | -3.68027 | 0.000614 |
| 20442 | Lifetime number of depressed periods | -0.5429 | 0.0002 | 0.1072 | 0.2994 | -3.65156 | 0.000645 |
| R10 | Diagnoses - main ICD10: R10 Abdominal and pelvic pain | -0.1223 | 0.0809 | 0.5273 | 3.16E-22 | -7.32744 | 0.000651 |
| 20003_1140860834 | Treatment/medication code: glyceryl trinitrate | -0.0817 | 0.651 | 0.5679 | 0.037 | -1.98808 | 0.000651 |
| 20488 | Physically abused by family as a child | -0.3003 | 4.92E-06 | 0.347 | 3.05E-12 | -7.85742 | 0.000677 |
| 20548_1 | Manifestations of mania or irritability: I was more talkative than usual | -0.4502 | 4.79E-07 | 0.1969 | 0.0031 | -5.80771 | 0.000679 |
| 41231_1 | Hospital episode type: General episode | -0.0907 | 0.1838 | 0.5497 | 4.42E-34 | -7.82707 | 0.000762 |
| 20002_1081 | Non-cancer illness code, self-reported: stroke | -0.1961 | 0.171 | 0.4434 | 9.67E-05 | -3.49591 | 0.000774 |
| I9_STR | Stroke, excluding SAH | -0.2137 | 0.4065 | 0.4212 | 0.0251 | -1.99137 | 0.000836 |
| ASTHMA_PNEUMONIA | Asthma-related pneumonia | -0.0719 | 0.6137 | 0.5603 | 1.57E-06 | -3.43237 | 0.000875 |
| PNEUMONIA | Pneumonias (Asthma/COPD co-morbidities) | -0.0719 | 0.6137 | 0.5603 | 1.57E-06 | -3.43237 | 0.000875 |
| 20548_9 | Manifestations of mania or irritability: I was more active than usual | -0.3253 | 0.0014 | 0.3046 | 3.44E-05 | -5.02643 | 0.00091 |
| 20002_1297 | Non-cancer illness code, self-reported: muscle/soft tissue problem | -0.4372 | 0.0121 | 0.1912 | 0.1764 | -2.79983 | 0.000933 |
| M84 | Diagnoses - main ICD10: M84 Disorders of continuity of bone | -0.3175 | 0.1901 | 0.3104 | 0.0446 | -2.18437 | 0.000941 |
| 20511 | Recent poor appetite or overeating | -0.2745 | 0.0015 | 0.3524 | 1.44E-12 | -6.28085 | 0.000957 |
| 6153_4 | Medication for cholesterol, blood pressure, diabetes, or take exogenous hormones: Hormone replacement therapy | -0.1581 | 0.1032 | 0.4654 | 1.25E-09 | -5.04456 | 0.001013 |
| ICDMAIN_ANY_ENTRY | Any ICDMAIN event in hilmo or causes of death | -0.0724 | 0.1844 | 0.5508 | 1.31E-50 | -9.46483 | 0.001018 |
| 20519 | Recent feelings of tiredness or low energy | -0.3569 | 1.60E-08 | 0.2646 | 9.06E-10 | -8.11848 | 0.001047 |
| 20417 | Tense, sore, or aching muscles during worst period of anxiety | -0.1469 | 0.1149 | 0.4732 | 1.08E-08 | -4.97402 | 0.001072 |
| 20003_1140923350 | Treatment/medication code: co-dydramol | 0.0651 | 0.6966 | 0.6852 | 0.0012 | -2.30308 | 0.001072 |

| | | | | | | | |
|---|---|---|---|---|---|---|---|
| 22601_71113328 | Job coding: sales assistant, retail/shop assistant, counter assistant, sub post office assistant, takeaway food worker | -0.1466 | 0.4873 | 0.4718 | 0.0081 | -2.23849 | 0.001102 |
| 20497 | Repeated disturbing thoughts of stressful experience in past month | -0.4046 | 6.27E-11 | 0.2129 | 1.19E-05 | -7.84633 | 0.001119 |
| FIBRO_COMORB | Fibromyalgia related co-morbidities | -0.283 | 0.0998 | 0.3338 | 0.0038 | -2.9779 | 0.001132 |
| J18 | Diagnoses - main ICD10: J18 Pneumonia, organism unspecified | -0.0527 | 0.7394 | 0.5621 | 2.25E-05 | -2.97726 | 0.00117 |
| G6_EPIPAROX | Episodal and paroxysmal disorders | -0.2662 | 0.0341 | 0.3467 | 3.45E-06 | -4.1916 | 0.001207 |
| 6146_2 | Attendance/disability/mobility allowance: Disability living allowance | -0.0375 | 0.5445 | 0.5751 | 3.54E-53 | -8.46448 | 0.001213 |
| 20003_1140923348 | Treatment/medication code: co-proxamol | -0.1632 | 0.4795 | 0.4486 | 0.0256 | -1.99948 | 0.001229 |
| 20536_3 | Weight change during worst episode of depression: Both gained and lost some weight during the episode | 0.0176 | 0.9198 | 0.6216 | 0.0034 | -2.1987 | 0.001395 |
| 22617_7111 | Job SOC coding: Sales and retail assistants | -0.151 | 0.4316 | 0.453 | 0.0053 | -2.40175 | 0.001395 |
| 20110_6 | Illnesses of mother: Chronic bronchitis/emphysema | -0.1389 | 0.0537 | 0.4644 | 1.79E-15 | -6.50761 | 0.001411 |
| 20126_3 | Bipolar and major depression status: Probable Recurrent major depression (severe) | -0.3502 | 0.0018 | 0.2529 | 0.0016 | -4.37663 | 0.001416 |
| 2080 | Frequency of tiredness / lethargy in last 2 weeks | -0.3338 | 1.17E-16 | 0.2685 | 4.53E-19 | -11.9741 | 0.001434 |
| N39 | Diagnoses - main ICD10: N39 Other disorders of urinary system | -0.0309 | 0.7067 | 0.5707 | 9.71E-14 | -5.35801 | 0.00145 |
| K20 | Diagnoses - main ICD10: K20 Oesophagitis | -0.0807 | 0.5692 | 0.518 | 0.0002 | -2.99077 | 0.00152 |
| 20003_1141162764 | Treatment/medication code: tolterodine l-tartrate | -0.2892 | 0.0838 | 0.3019 | 0.0393 | -2.6581 | 0.001716 |
| 20513 | Recent thoughts of suicide or self-harm | -0.2899 | 0.0127 | 0.2996 | 0.0013 | -3.96038 | 0.00176 |
| 2227 | Other eye problems | -0.3615 | 6.78E-05 | 0.2279 | 0.0006 | -5.24342 | 0.001763 |
| 20507 | Recent feelings of inadequacy | -0.3938 | 1.05E-07 | 0.1945 | 0.0007 | -6.27352 | 0.001794 |
| 22607_2 | Workplace very cold: Often | -0.1764 | 0.0346 | 0.4101 | 7.37E-09 | -5.3542 | 0.001846 |
| 22617_7212 | Job SOC coding: Customer care occupations | -0.4017 | 0.0455 | 0.1836 | 0.1632 | -2.43652 | 0.001881 |
| T40 | Diagnoses - main ICD10: T40 Poisoning by narcotics and psychodysleptics [hallucinogens] | -0.148 | 0.3068 | 0.4373 | 0.021 | -2.4542 | 0.001881 |
| 189 | Townsend deprivation index at recruitment | -0.291 | 1.44E-09 | 0.2934 | 2.89E-17 | -9.85328 | 0.001908 |
| 6146_3 | Attendance/disability/mobility allowance: Blue badge | -0.0285 | 0.6998 | 0.5558 | 3.03E-37 | -6.80978 | 0.001911 |
| 6141_8 | How are people in household related to participant: Other unrelated | -0.297 | 0.0384 | 0.2856 | 0.0639 | -2.76769 | 0.001963 |
| R07 | Diagnoses - main ICD10: R07 Pain in throat and chest | -0.0533 | 0.3956 | 0.5289 | 2.18E-34 | -7.63235 | 0.001976 |
| 20548_6 | Manifestations of mania or irritability: I was more creative or had more ideas than usual | -0.3409 | 0.0159 | 0.2412 | 0.0179 | -3.34136 | 0.001979 |
| K62 | Diagnoses - main ICD10: K62 Other diseases of anus and rectum | -0.161 | 0.1621 | 0.4209 | 4.34E-05 | -3.76901 | 0.001985 |
| 20498 | Felt very upset when reminded of stressful experience in past month | -0.3239 | 1.50E-06 | 0.2565 | 5.07E-09 | -7.22319 | 0.002032 |
| 2296 | Falls in the last year | -0.2445 | 1.17E-06 | 0.3344 | 2.96E-24 | -9.63163 | 0.00208 |
| 6157_1 | Why stopped smoking: Illness or ill health | -0.2131 | 0.0575 | 0.3654 | 0.0003 | -3.85078 | 0.002093 |
| 22618_1 | Breathing problems improved/stopped away from workplace or on holiday: Yes | -0.3773 | 0.0098 | 0.1989 | 0.0921 | -3.06839 | 0.00217 |
| 20003_1140865634 | Treatment/medication code: omeprazole | -0.1109 | 0.1332 | 0.4644 | 2.43E-18 | -6.31799 | 0.002201 |
| M13_RADICULOPATHY | Radiculopathy | -0.1273 | 0.5222 | 0.4476 | 0.0091 | -2.18848 | 0.002214 |
| 20527 | Been involved in combat or exposed to war-zone | -0.0623 | 0.6288 | 0.5117 | 2.4E-07 | -3.53204 | 0.002245 |
| 6164_100 | Types of physical activity in last 4 weeks: None of the above | -0.1288 | 0.0249 | 0.4446 | 2.72E-30 | -8.26945 | 0.002266 |
| 20548_7 | Manifestations of mania or irritability: I was easily distracted | -0.3022 | 0.0014 | 0.2702 | 0.0004 | -4.69758 | 0.002302 |
| 136 | Number of operations, self-reported | -0.1166 | 0.0307 | 0.4549 | 1.76E-48 | -9.17109 | 0.002334 |

| | | | | | | | |
|---|---|---|---|---|---|---|---|
| 20552_2 | Behavioural and miscellaneous addictions: A behaviour | -0.5202 | 0.046 | 0.0496 | 0.6536 | -2.01207 | 0.002396 |
| 20002_1093 | Non-cancer illness code, self-reported: pulmonary embolism +/- dvt | -0.0523 | 0.6164 | 0.5164 | 2.39E-09 | -4.19461 | 0.002437 |
| XI_DIGESTIVE | Diseases of the digestive system | -0.116 | 0.0083 | 0.4505 | 1.48E-39 | -10.1798 | 0.002521 |
| 20422 | More irritable than usual during worst period of anxiety | -0.2585 | 0.0674 | 0.3036 | 0.0041 | -3.18542 | 0.002697 |
| 20548_3 | Manifestations of mania or irritability: My thoughts were racing | -0.2004 | 0.0243 | 0.3613 | 4.64E-09 | -5.18674 | 0.002713 |
| 2188 | Long-standing illness, disability or infirmity | -0.2475 | 1.51E-09 | 0.3137 | 2.79E-28 | -11.252 | 0.002734 |
| 22506_111 | Tobacco smoking: Smokes on most or all days | -0.0925 | 0.4962 | 0.4675 | 0.0025 | -2.71858 | 0.002784 |
| 6152_7 | Blood clot, DVT, bronchitis, emphysema, asthma, rhinitis, eczema, allergy diagnosed by doctor: Blood clot in the lung | -0.0581 | 0.5696 | 0.5001 | 1.16E-09 | -4.25856 | 0.002862 |
| 6154_5 | Medication for pain relief, constipation, heartburn: Omeprazole (e.g. Zanprol) | -0.1428 | 0.022 | 0.4151 | 3.45E-19 | -7.18199 | 0.002875 |
| C_RESPIRATORY_INTRATHORACIC | NA | -0.1681 | 0.1017 | 0.3895 | 0.0006 | -3.65522 | 0.002888 |
| C3_RESPIRATORY_INTRATHORACIC | Malignant neoplasm of respiratory system and intrathoracic organs | -0.1681 | 0.1017 | 0.3895 | 0.0006 | -3.65522 | 0.002888 |
| 2100 | Seen a psychiatrist for nerves, anxiety, tension or depression | -0.3574 | 5.32E-15 | 0.1992 | 5.29E-09 | -9.76146 | 0.002932 |
| 20532 | Did your sleep change? | -0.196 | 0.1392 | 0.3597 | 0.0002 | -3.39624 | 0.002972 |
| 2335 | Chest pain or discomfort | -0.2424 | 2.55E-07 | 0.3116 | 4.78E-18 | -9.35763 | 0.003049 |
| 6159_8 | Pain type(s) experienced in last month: Pain all over the body | -0.0811 | 0.3474 | 0.4727 | 1.87E-14 | -5.22021 | 0.003058 |
| 20111_2 | Illnesses of siblings: Stroke | -0.1602 | 0.2726 | 0.3936 | 0.0028 | -2.81559 | 0.003058 |
| R31 | Diagnoses - main ICD10: R31 Unspecified haematuria | -0.27 | 0.0115 | 0.2836 | 0.0004 | -4.15987 | 0.003068 |
| 20494 | Felt irritable or had angry outbursts in past month | -0.3761 | 0.0051 | 0.1772 | 0.0948 | -3.23276 | 0.003081 |
| 20547_1 | Activities undertaken to treat depression: Talking therapies, such as psychotherapy, counselling, group therapy or CBT | -0.4217 | 3.59E-11 | 0.1295 | 0.0137 | -6.6723 | 0.00318 |
| E4_DM2 | Type 2 diabetes | -0.1315 | 0.2871 | 0.4195 | 3.85E-05 | -3.44134 | 0.00319 |
| F32 | Diagnoses - main ICD10: F32 Depressive episode | -0.2763 | 0.077 | 0.274 | 0.0117 | -2.89137 | 0.003224 |
| R32 | Diagnoses - main ICD10: R32 Unspecified urinary incontinence | -0.2315 | 0.2102 | 0.3183 | 0.0073 | -2.50479 | 0.003248 |
| 6159_2 | Pain type(s) experienced in last month: Facial pain | -0.3011 | 0.0027 | 0.2485 | 0.0042 | -4.13903 | 0.003258 |
| 20485 | Ever contemplated self-harm | -0.4968 | 1.01E-10 | 0.0524 | 0.3046 | -5.95718 | 0.003277 |
| 2149 | Lifetime number of sexual partners | -0.1549 | 0.004 | 0.3942 | 3.24E-52 | -9.19616 | 0.003282 |
| 20111_6 | Illnesses of siblings: Chronic bronchitis/emphysema | -0.1267 | 0.2598 | 0.4219 | 9.29E-06 | -3.72441 | 0.003307 |
| COLITNONINFNAS | Noninfectious colitis NAS | -0.096 | 0.3612 | 0.4524 | 3.93E-08 | -4.10821 | 0.003317 |
| 20500 | Ever suffered mental distress preventing usual activities | -0.4048 | 2.01E-11 | 0.1429 | 0.0007 | -7.45065 | 0.003351 |
| M13_SACROCOGG | Rotator cuff syndrome | -0.2451 | 0.1073 | 0.3024 | 0.037 | -2.60449 | 0.003361 |
| E66 | Diagnoses - main ICD10: E66 Obesity | -0.1874 | 0.0808 | 0.3587 | 0.0002 | -3.81763 | 0.003432 |
| 20502 | Ever had period extreme irritability | -0.3407 | 5.25E-07 | 0.2043 | 0.0002 | -6.27756 | 0.003489 |
| XIX_INJURY_POISON | Injury, poisoning and certain other consequences of external causes | -0.1148 | 0.0427 | 0.4288 | 3.73E-20 | -7.41456 | 0.003562 |
| N92 | Diagnoses - main ICD10: N92 Excessive, frequent and irregular menstruation | -0.1302 | 0.3129 | 0.4127 | 1.64E-07 | -3.59147 | 0.003599 |
| 680_4 | Own or rent accommodation lived in: Rent - from private landlord or letting agency | -0.0717 | 0.5109 | 0.4702 | 2.56E-08 | -3.92865 | 0.003653 |
| 20002_1267 | Non-cancer illness code, self-reported: spinal injury | -0.1749 | 0.5389 | 0.3657 | 0.0407 | -1.60827 | 0.003724 |
| 6159_3 | Pain type(s) experienced in last month: Neck or shoulder pain | -0.1438 | 0.0009 | 0.3962 | 6.85E-31 | -9.74796 | 0.003757 |

| | | | | | | | |
|---|---|---|---|---|---|---|---|
| E4_OBESITY | Obesity | -0.1883 | 0.089 | 0.3507 | 0.0002 | -3.73301 | 0.003813 |
| I10 | Diagnoses - main ICD10: I10 Essential (primary) hypertension | -0.2242 | 0.1372 | 0.3137 | 0.0239 | -2.62269 | 0.003875 |
| 20002_1442 | Non-cancer illness code, self-reported: helicobacter pylori | -0.2052 | 0.1283 | 0.3325 | 0.0064 | -2.95736 | 0.003887 |
| F5_PERSONALITY | Specific personality disorders | -0.2849 | 0.0934 | 0.2525 | 0.0286 | -2.6176 | 0.003904 |
| C34 | Diagnoses - main ICD10: C34 Malignant neoplasm of bronchus and lung | -0.1591 | 0.2155 | 0.3777 | 0.0073 | -2.81813 | 0.003938 |
| 20002_1398 | Non-cancer illness code, self-reported: pneumonia | -0.3496 | 0.0134 | 0.1868 | 0.0672 | -3.07697 | 0.003962 |
| K11_CHRONPANC | Chronic pancreatitis | -0.0474 | 0.8128 | 0.4887 | 0.0301 | -1.77872 | 0.003979 |
| 20002_1201 | Non-cancer illness code, self-reported: bladder problem (not cancer) | -0.2395 | 0.1271 | 0.2955 | 0.008 | -2.77912 | 0.004044 |
| 20471 | Ever seen an un-real vision | -0.204 | 0.1293 | 0.3306 | 0.0018 | -3.12515 | 0.004068 |
| 3089 | Caffeine drink within last hour | -0.0339 | 0.8617 | 0.4995 | 0.0066 | -1.9938 | 0.00414 |
| 20508 | Recent trouble concentrating on things | -0.3117 | 7.42E-06 | 0.2212 | 1.25E-05 | -6.19295 | 0.00417 |
| 6152_6 | Blood clot, DVT, bronchitis, emphysema, asthma, rhinitis, eczema, allergy diagnosed by doctor: Emphysema/chronic bronchitis | -0.1238 | 0.2687 | 0.4087 | 1.83E-12 | -4.22491 | 0.004194 |
| VI_NERVOUS | Diseases of the nervous system | -0.0303 | 0.6693 | 0.5002 | 1.79E-23 | -6.10496 | 0.004318 |
| 20546_4 | Substances taken for depression: Drugs or alcohol (more than once) | -0.3763 | 0.0003 | 0.1539 | 0.0316 | -4.18548 | 0.004337 |
| 20454 | Maximum frequency of taking cannabis | -0.1315 | 0.2754 | 0.3981 | 0.0003 | -3.2418 | 0.004375 |
| 20501 | Ever had period of mania / excitability | -0.2411 | 0.0127 | 0.2882 | 0.0002 | -4.29707 | 0.004394 |
| XIV_GENITOURINARY | Diseases of the genitourinary system | -0.0621 | 0.3043 | 0.4655 | 2.59E-26 | -7.06591 | 0.004504 |
| M13_OTHERJOINT | #Other joint disorders | -0.0415 | 0.5097 | 0.486 | 2.61E-22 | -6.56488 | 0.00451 |
| 6144_2 | Never eat eggs, dairy, wheat, sugar: Dairy products | -0.1747 | 0.1701 | 0.3523 | 0.0018 | -3.10086 | 0.004543 |
| 20003_1140879406 | Treatment/medication code: ranitidine | -0.1842 | 0.0825 | 0.3407 | 0.0004 | -3.67018 | 0.004683 |
| T39 | Diagnoses - main ICD10: T39 Poisoning by nonopioid analgesics, antipyretics and antirheumatics | -0.2089 | 0.199 | 0.3157 | 0.0072 | -2.615 | 0.004703 |
| 2020 | Loneliness, isolation | -0.2872 | 3.76E-09 | 0.2372 | 8.05E-10 | -8.43871 | 0.004717 |
| 22606_2 | Workplace very noisy: Often | -0.0778 | 0.2963 | 0.4466 | 2.2E-18 | -5.80469 | 0.004717 |
| 6149_2 | Mouth/teeth dental problems: Painful gums | -0.2395 | 0.0177 | 0.2848 | 0.0006 | -4.01839 | 0.004724 |
| 22620_1 | Job involved shift work: Yes | -0.0455 | 0.637 | 0.4786 | 6.75E-12 | -4.40575 | 0.004737 |
| 20002_1142 | Non-cancer illness code, self-reported: gastric/stomach ulcers | 0.0374 | 0.8055 | 0.5614 | 0.0232 | -1.80517 | 0.004744 |
| 2814 | Ever used hormone-replacement therapy (HRT) | -0.1302 | 0.0218 | 0.3937 | 1.69E-21 | -7.46002 | 0.004751 |
| 6159_5 | Pain type(s) experienced in last month: Stomach or abdominal pain | -0.2511 | 0.0002 | 0.2725 | 1.14E-07 | -6.23558 | 0.004772 |
| E4_OBESITYNAS | Obesity, other/unspecified | -0.1719 | 0.1025 | 0.3516 | 0.0001 | -3.77035 | 0.004778 |
| K29 | Diagnoses - main ICD10: K29 Gastritis and duodenitis | -0.1111 | 0.1106 | 0.4119 | 7.31E-13 | -5.7972 | 0.004813 |
| 20510 | Recent feelings of depression | -0.3458 | 3.30E-07 | 0.1766 | 0.0005 | -6.16327 | 0.004855 |
| 20002_1112 | Non-cancer illness code, self-reported: chronic obstructive airways disease/copd | -0.0981 | 0.3657 | 0.424 | 1.11E-05 | -3.59561 | 0.004876 |
| 6145_1 | Illness, injury, bereavement, stress in last 2 years: Serious illness, injury or assault to yourself | -0.2057 | 0.0044 | 0.3154 | 2.4E-09 | -5.82583 | 0.004946 |
| 3773 | Knee pain for 3+ months | -0.2014 | 0.0713 | 0.3192 | 3.93E-06 | -3.96456 | 0.004982 |
| 20003_1140883568 | Treatment/medication code: oxybutynin | -0.2286 | 0.1067 | 0.2913 | 0.0293 | -2.66863 | 0.005032 |
| M13_IMPINGEMENT | Impingement syndrome of shoulder | -0.0899 | 0.4322 | 0.4299 | 2.14E-07 | -3.67924 | 0.005039 |
| 4717 | Shortness of breath walking on level ground | -0.0975 | 0.1669 | 0.4218 | 1.4E-16 | -5.96253 | 0.005075 |
| 2060 | Frequency of unenthusiasm / disinterest in last 2 weeks | -0.2302 | 2.32E-06 | 0.2887 | 2.8E-13 | -8.27523 | 0.005104 |
| 4631 | Ever unenthusiastic/disinterested for a whole week | -0.3886 | 1.10E-07 | 0.1295 | 0.0056 | -5.96326 | 0.005163 |

| | | | | | | | |
|---|---|---|---|---|---|---|---|
| K44 | Diagnoses - main ICD10: K44 Diaphragmatic hernia | -0.0055 | 0.9598 | 0.512 | 1.04E-13 | -4.00791 | 0.005207 |
| 3404 | Neck/shoulder pain for 3+ months | -0.0996 | 0.4753 | 0.4178 | 3.3E-06 | -3.11866 | 0.005215 |
| 20496 | Felt distant from other people in past month | -0.4227 | 0.0013 | 0.0943 | 0.2274 | -3.37463 | 0.005244 |
| 4728 | Leg pain on walking | -0.0805 | 0.264 | 0.4358 | 5.93E-23 | -6.11118 | 0.005297 |
| 20455 | Age when last took cannabis | -0.0629 | 0.6634 | 0.4521 | 0.009 | -2.28396 | 0.005396 |
| 6162_100 | Types of transport used (excluding work): None of the above | -0.094 | 0.4361 | 0.4209 | 3.33E-05 | -3.2663 | 0.005403 |
| 41248_5001 | Destinations on discharge from hospital (recoded): Transfer to other NHS provider: General ward, young physically disabled, A&E | -0.2553 | 0.0547 | 0.259 | 0.0418 | -2.79568 | 0.005449 |
| 20003_1140884464 | Treatment/medication code: dihydrocodeine | 0.0819 | 0.586 | 0.5959 | 2.11E-05 | -2.50069 | 0.005473 |
| K52 | Diagnoses - main ICD10: K52 Other non-infective gastro-enteritis and colitis | -0.0639 | 0.5414 | 0.4496 | 9.13E-08 | -3.82592 | 0.005512 |
| 4548 | Health satisfaction | -0.2086 | 0.0033 | 0.3042 | 1.6E-13 | -6.24695 | 0.005566 |
| 22617_3311 | Job SOC coding: NCOs and other ranks | 0.0762 | 0.6374 | 0.588 | 1.64E-05 | -2.4202 | 0.005646 |
| 20002_1295 | Non-cancer illness code, self-reported: joint disorder | -0.3874 | 0.0414 | 0.1241 | 0.2676 | -2.3197 | 0.00567 |
| E11 | Diagnoses - main ICD10: E11 Non-insulin-dependent diabetes mellitus | -0.1094 | 0.3718 | 0.4015 | 3.56E-05 | -3.26838 | 0.005718 |
| 6151_7 | Fractured bone site(s): Other bones | -0.224 | 0.0133 | 0.2856 | 5.75E-05 | -4.43329 | 0.005824 |
| K81 | Diagnoses - main ICD10: K81 Cholecystitis | -0.1528 | 0.3964 | 0.3565 | 0.0198 | -2.15447 | 0.005848 |
| 2050 | Frequency of depressed mood in last 2 weeks | -0.3032 | 1.36E-09 | 0.2058 | 3.68E-06 | -7.60443 | 0.005873 |
| 22601_33113434 | Job coding: non-commissioned officers or other rank of armed forces | 0.0737 | 0.6523 | 0.5821 | 3.33E-05 | -2.35977 | 0.005923 |
| 2090 | Seen doctor (GP) for nerves, anxiety, tension or depression | -0.3366 | 9.70E-16 | 0.1716 | 3.59E-07 | -9.45122 | 0.005939 |
| 20003_1140864752 | Treatment/medication code: lansoprazole | -0.0789 | 0.2627 | 0.4288 | 5.18E-16 | -5.76015 | 0.005981 |
| 2473 | Other serious medical condition/disability diagnosed by doctor | -0.2961 | 8.75E-11 | 0.2114 | 1.6E-07 | -8.33936 | 0.005998 |
| 2070 | Frequency of tenseness / restlessness in last 2 weeks | -0.2422 | 1.43E-08 | 0.265 | 3.76E-12 | -8.86299 | 0.006023 |
| 20544_11 | Mental health problems ever diagnosed by a professional: Depression | -0.3594 | 3.08E-07 | 0.1462 | 0.0025 | -5.92955 | 0.00616 |
| 20554_1 | Actions taken following self-harm: See anyone from psychiatric or mental health services, including liaison services | -0.339 | 0.011 | 0.1666 | 0.0671 | -3.13099 | 0.00616 |
| 20003_1140879540 | Treatment/medication code: fluoxetine | -0.306 | 0.0204 | 0.1995 | 0.0154 | -3.24966 | 0.006168 |
| VIII_EAR_MASTOID | Diseases of the ear and mastoid process | -0.0936 | 0.4232 | 0.4118 | 4.21E-06 | -3.43279 | 0.006177 |
| 20002_1154 | Non-cancer illness code, self-reported: irritable bowel syndrome | -0.2536 | 0.0179 | 0.2513 | 0.0092 | -3.50069 | 0.00622 |
| 6142_4 | Current employment status: Unable to work because of sickness or disability | -0.0521 | 0.4348 | 0.4522 | 2.89E-27 | -6.40662 | 0.006273 |
| M47 | Diagnoses - main ICD10: M47 Spondylosis | 0.0068 | 0.9703 | 0.5107 | 1.23E-05 | -2.33375 | 0.006308 |
| 2040 | Risk taking | -0.0126 | 0.8041 | 0.491 | 1.07E-58 | -8.50656 | 0.006334 |
| R69 | Diagnoses - main ICD10: R69 Unknown and unspecified causes of morbidity | -0.1296 | 0.2736 | 0.3702 | 0.0005 | -3.13713 | 0.006678 |
| 20495 | Avoided activities or situations because of previous stressful experience in past month | -0.3928 | 1.02E-07 | 0.1066 | 0.0839 | -5.19158 | 0.006715 |
| 100160 | Low calorie drink intake | -0.1161 | 0.5038 | 0.3826 | 0.008 | -2.20903 | 0.00678 |
| 22610_1 | Workplace full of chemical or other fumes: Sometimes | -0.1349 | 0.1692 | 0.3637 | 1.86E-06 | -4.01194 | 0.00679 |
| K11_OTHGASTR | Other gastritis (incl. Duodenitis) | -0.0982 | 0.1969 | 0.3996 | 1.02E-09 | -4.96107 | 0.006865 |
| 20003_1140921600 | Treatment/medication code: citalopram | -0.2749 | 0.0455 | 0.2218 | 0.0228 | -2.94774 | 0.00697 |
| 20499 | Ever sought or received professional help for mental distress | -0.4183 | 3.52E-12 | 0.0777 | 0.0905 | -6.55197 | 0.007038 |
| R53 | Diagnoses - main ICD10: R53 Malaise and fatigue | -0.2624 | 0.0678 | 0.2332 | 0.0377 | -2.71838 | 0.007077 |
| 20480 | Ever self-harmed | -0.4309 | 1.36E-06 | 0.063 | 0.3482 | -4.42245 | 0.007244 |
| M13_SOFTTISSUENAS | Other specified/unspecified soft tissue disorders | -0.0579 | 0.7641 | 0.4354 | 8.25E-06 | -2.27948 | 0.007304 |

| | | | | | | | |
|---|---|---|---|---|---|---|---|
| S61 | Diagnoses - main ICD10: S61 Open wound of wrist and hand | -0.0164 | 0.9596 | 0.4748 | 0.0187 | -1.28639 | 0.007517 |
| 20110_3 | Illnesses of mother: Lung cancer | 0.1022 | 0.3873 | 0.5925 | 2.68E-06 | -2.83559 | 0.00761 |
| 6141_6 | How are people in household related to participant: Grandchild | -0.058 | 0.6662 | 0.4323 | 1.74E-05 | -2.92193 | 0.00761 |
| 135 | Number of self-reported non-cancer illnesses | -0.2074 | 5.53E-07 | 0.2794 | 7.6E-26 | -9.89251 | 0.007981 |
| 22609_2 | Workplace very dusty: Often | -0.0727 | 0.4305 | 0.4125 | 3.9E-10 | -4.27824 | 0.008156 |
| 20514 | Recent lack of interest or pleasure in doing things | -0.2778 | 0.0002 | 0.2072 | 0.0001 | -5.23671 | 0.008178 |
| 4581 | Financial situation satisfaction | -0.087 | 0.1974 | 0.398 | 2.37E-22 | -6.14512 | 0.008178 |
| Z41 | Diagnoses - main ICD10: Z41 Procedures for purposes other than remedying health state | 0.0774 | 0.7347 | 0.562 | 0.0481 | -1.32832 | 0.008222 |
| 3456 | Number of cigarettes currently smoked daily (current cigarette smokers) | -0.2292 | 0.0122 | 0.2545 | 0.0003 | -4.17426 | 0.008323 |
| 22599 | Number of jobs held | -0.2507 | 0.019 | 0.2328 | 0.001 | -3.77496 | 0.008345 |
| 20524 | Sexual interference by partner or ex-partner without consent as an adult | -0.2652 | 0.0074 | 0.2178 | 0.0015 | -4.01097 | 0.008402 |
| M13_MUSCULOSKELEOTH | Other disorders of the musculoskeletal system and connective tissue | -0.1333 | 0.5218 | 0.3481 | 0.0188 | -1.88474 | 0.008585 |
| 20003_1140927730 | Treatment/medication code: epaderm ointment | -0.2995 | 0.0392 | 0.1815 | 0.2691 | -2.19302 | 0.008631 |
| V_MENTAL_BEHAV | Mental and behavioural disorders | -0.2251 | 0.0171 | 0.2557 | 0.0005 | -4.01874 | 0.008654 |
| 20002_1113 | Non-cancer illness code, self-reported: emphysema/chronic bronchitis | -0.0713 | 0.5248 | 0.4088 | 1.18E-09 | -3.67332 | 0.008736 |
| CHRONNAS | Chron's disease NAS | -0.199 | 0.1695 | 0.2807 | 0.0205 | -2.54039 | 0.008783 |
| 6144_3 | Never eat eggs, dairy, wheat, sugar: Wheat products | 0.0559 | 0.7757 | 0.534 | 1.8E-05 | -2.05677 | 0.008973 |
| 20515 | Recent trouble relaxing | -0.2959 | 4.55E-06 | 0.1818 | 0.0004 | -5.7789 | 0.009021 |
| 2492 | Taking other prescription medications | -0.1388 | 0.0043 | 0.338 | 7.82E-34 | -8.50836 | 0.00913 |
| 20401 | Ever addicted to any substance or behaviour | -0.2067 | 0.1069 | 0.2686 | 0.0029 | -3.0344 | 0.009315 |
| G47 | Diagnoses - main ICD10: G47 Sleep disorders | -0.1626 | 0.1662 | 0.3127 | 9.19E-05 | -3.34695 | 0.009315 |
| 20118_5 | Home area population density - urban or rural: England/Wales - Urban - less sparse | -0.2836 | 0.0046 | 0.1916 | 0.0152 | -3.72832 | 0.009327 |
| 22608_2 | Workplace very hot: Often | -0.1225 | 0.1435 | 0.3518 | 3.97E-07 | -4.36221 | 0.00944 |
| LUNG_CANCER | Lung cancer and mesothelioma | -0.1139 | 0.2569 | 0.3582 | 0.0006 | -3.25297 | 0.009719 |
| LUNG_CANCER_MESOT | Lung cancer and mesothelioma | -0.1139 | 0.2569 | 0.3582 | 0.0006 | -3.25297 | 0.009719 |
| 20463 | Ever heard an un-real voice | -0.2428 | 0.0389 | 0.2293 | 0.0099 | -3.20413 | 0.009719 |
| C_BRONCHUS_LUNG | Malignant neoplasm of bronchus and lung | -0.1582 | 0.0734 | 0.3127 | 0.0007 | -3.68238 | 0.009875 |
| C3_BRONCHUS_LUNG | Malignant neoplasm of bronchus and lung | -0.1582 | 0.0734 | 0.3127 | 0.0007 | -3.68238 | 0.009875 |
| 2178 | Overall health rating | -0.1184 | 0.0073 | 0.3516 | 2.22E-36 | -9.00651 | 0.009993 |
| 680_3 | Own or rent accommodation lived in: Rent - from local authority, local council, housing association | -0.0752 | 0.1217 | 0.3944 | 8.06E-40 | -8.22977 | 0.010045 |
| 20002_1311 | Non-cancer illness code, self-reported: spine arthritis/spondylitis | 0.1577 | 0.1647 | 0.6261 | 5.51E-06 | -2.62373 | 0.010206 |
| 20512 | Recent feelings of foreboding | -0.3187 | 4.77E-06 | 0.1487 | 0.0052 | -5.33056 | 0.010341 |
| SLEEP | Sleep disorders (combined) | -0.1567 | 0.1725 | 0.3105 | 8.27E-05 | -3.35393 | 0.010368 |
| 20546_1 | Substances taken for depression: Unprescribed medication (more than once) | -0.3896 | 0.025 | 0.0765 | 0.4679 | -2.2931 | 0.010519 |
| 20534 | Sleeping too much | -0.3322 | 0.0063 | 0.1338 | 0.0614 | -3.3023 | 0.010532 |
| M13_POLYARTHROPATHIES | #Polyarthropathies | -0.1355 | 0.4017 | 0.3301 | 0.0003 | -2.51383 | 0.010588 |
| 20003_1141177600 | Treatment/medication code: rosiglitazone | -0.2275 | 0.0668 | 0.2354 | 0.0212 | -2.87934 | 0.010968 |
| 2316 | Wheeze or whistling in the chest in last year | -0.0802 | 0.1162 | 0.3823 | 4.77E-36 | -7.77178 | 0.011025 |
| 20003_1140876592 | Treatment/medication code: multivitamin+mineral preparations | -0.5056 | 0.0039 | -0.0432 | 0.6783 | -2.27088 | 0.01104 |
| M13_DORSALGIA | Dorsalgia | 0.0462 | 0.5504 | 0.5077 | 9.85E-24 | -4.99069 | 0.01117 |
| M54 | Diagnoses - main ICD10: M54 Dorsalgia | 0.0462 | 0.5504 | 0.5077 | 9.85E-24 | -4.99069 | 0.01117 |

| | | | | | | | |
|---|---|---|---|---|---|---|---|
| 6151_2 | Fractured bone site(s): Leg | -0.0992 | 0.4715 | 0.3617 | 0.0119 | -2.31411 | 0.011257 |
| 20107_6 | Illnesses of father: Chronic bronchitis/emphysema | -0.1868 | 0.0066 | 0.2739 | 3.18E-07 | -5.28715 | 0.011286 |
| 20544_6 | Mental health problems ever diagnosed by a professional: Panic attacks | -0.1826 | 0.0346 | 0.2781 | 1.86E-05 | -4.26337 | 0.011286 |
| 20553_5 | Methods of self-harm used: Self-injury such as self-cutting, scratching or hitting, etc. | -0.489 | 0.0067 | -0.03 | 0.799 | -2.12983 | 0.011538 |
| G6_SLEEPAPNO | Sleep apnoea | -0.194 | 0.1662 | 0.2642 | 0.0029 | -2.76237 | 0.011658 |
| I9_PAD | Peripheral artery disease | -0.0564 | 0.7038 | 0.4006 | 0.0038 | -2.25221 | 0.01184 |
| 6146_1 | Attendance/disability/mobility allowance: Attendance allowance | 0.0997 | 0.2708 | 0.5552 | 1.59E-13 | -3.8686 | 0.012071 |
| XIII_MUSCULOSKELET | Diseases of the musculoskeletal system and connective tissue | 0.0444 | 0.478 | 0.4998 | 7.93E-51 | -6.42259 | 0.012087 |
| 20447 | Depression possibly related to stressful or traumatic event | 0.0594 | 0.6159 | 0.5146 | 8.81E-05 | -2.57574 | 0.012118 |
| 2415 | Had major operations | -0.0955 | 0.1991 | 0.3595 | 1.93E-11 | -4.962 | 0.012149 |
| I74 | Diagnoses - main ICD10: I74 Arterial embolism and thrombosis | 0.0835 | 0.8493 | 0.5385 | 0.0045 | -0.95087 | 0.012149 |
| 22615_2 | Workplace had a lot of diesel exhaust: Often | 0.0626 | 0.6657 | 0.5171 | 6.61E-06 | -2.45751 | 0.012228 |
| M13_LOWBACKPAIN | Low back pain | 0.066 | 0.481 | 0.5202 | 2.14E-17 | -4.05643 | 0.012275 |
| 20002_1223 | Non-cancer illness code, self-reported: type 2 diabetes | -0.2844 | 0.0184 | 0.1691 | 0.0181 | -3.23263 | 0.012386 |
| 6149_5 | Mouth/teeth dental problems: Toothache | -0.1956 | 0.0291 | 0.2579 | 0.0001 | -4.04622 | 0.012386 |
| 20003_1141177526 | Treatment/medication code: esomeprazole | 0.0664 | 0.6558 | 0.5187 | 0.0018 | -2.02896 | 0.012578 |
| 20003_1141168318 | Treatment/medication code: clopidogrel | -0.0478 | 0.6776 | 0.4005 | 0.0002 | -2.82685 | 0.013236 |
| 20529 | Victim of physically violent crime | -0.4629 | 2.67E-08 | -0.0154 | 0.7855 | -4.44712 | 0.013372 |
| 20531 | Victim of sexual assault | -0.3382 | 1.39E-06 | 0.1085 | 0.0439 | -5.05514 | 0.013508 |
| I73 | Diagnoses - main ICD10: I73 Other peripheral vascular diseases | -0.0961 | 0.5282 | 0.3504 | 0.0334 | -1.99042 | 0.013542 |
| 20549_4 | Substances taken for anxiety: Drugs or alcohol (more than once) | -0.4314 | 0.0014 | 0.0148 | 0.8716 | -2.72806 | 0.013594 |
| 20546_3 | Substances taken for depression: Medication prescribed to you (for at least two weeks) | -0.3439 | 2.53E-07 | 0.1019 | 0.0391 | -5.37099 | 0.013663 |
| I9_ARTEMBTHR | Arterial embolism and thrombosis | 0.0831 | 0.8344 | 0.5288 | 0.004 | -1.01825 | 0.01368 |
| 6154_4 | Medication for pain relief, constipation, heartburn: Ranitidine (e.g. Zantac) | -0.1159 | 0.2462 | 0.3291 | 0.0007 | -3.18624 | 0.013802 |
| 2887 | Number of cigarettes previously smoked daily | -0.1626 | 0.008 | 0.2822 | 2.71E-08 | -5.58698 | 0.013837 |
| K76 | Diagnoses - main ICD10: K76 Other diseases of liver | -0.1896 | 0.0981 | 0.2552 | 0.0208 | -2.79526 | 0.013837 |
| 20003_99999 | Treatment/medication code: Free-text entry, unable to be coded | -0.017 | 0.9243 | 0.4275 | 0.0016 | -1.98489 | 0.01389 |
| 20544_7 | Mental health problems ever diagnosed by a professional: Obsessive compulsive disorder (OCD) | -0.3899 | 0.0483 | 0.0544 | 0.6543 | -1.91678 | 0.013925 |
| 4653 | Ever highly irritable/argumentative for 2 days | -0.0859 | 0.1961 | 0.3573 | 1.81E-10 | -5.09787 | 0.014119 |
| F5_MOOD | Mood [affective] disorders | -0.2664 | 0.0116 | 0.1763 | 0.0184 | -3.42097 | 0.014208 |
| KRA_PSY_MOOD | Mood disorders | -0.2664 | 0.0116 | 0.1763 | 0.0184 | -3.42097 | 0.014208 |
| 20428 | Professional informed about anxiety | -0.1429 | 0.1941 | 0.2995 | 0.0024 | -2.99465 | 0.014262 |
| 20003_1140864992 | Treatment/medication code: tramadol | 0.0478 | 0.5728 | 0.4891 | 3.32E-12 | -4.00866 | 0.014461 |
| 20479 | Ever thought that life not worth living | -0.4246 | 8.41E-16 | 0.0166 | 0.713 | -6.35967 | 0.014479 |
| 20528 | Diagnosed with life-threatening illness | -0.1584 | 0.1096 | 0.2824 | 0.0019 | -3.28302 | 0.014552 |
| 20549_3 | Substances taken for anxiety: Medication prescribed to you (for at least two weeks) | -0.3489 | 4.26E-06 | 0.0914 | 0.1274 | -4.55084 | 0.014643 |
| 4825 | Noisy workplace | -0.1102 | 0.0616 | 0.3296 | 6.24E-12 | -5.78714 | 0.014736 |
| 20003_1140883468 | Treatment/medication code: clonidine | -0.1679 | 0.2561 | 0.2707 | 0.0426 | -2.20136 | 0.014959 |
| 4598 | Ever depressed for a whole week | -0.3611 | 7.44E-09 | 0.0759 | 0.1149 | -5.54104 | 0.01526 |
| H02 | Diagnoses - main ICD10: H02 Other disorders of eyelid | -0.104 | 0.5949 | 0.3313 | 0.0254 | -1.77281 | 0.015587 |
| H7_EYELIDDIS | Other disorders of eyelid | -0.1019 | 0.601 | 0.3333 | 0.0256 | -1.77319 | 0.015606 |

| | | | | | | |
|---|---|---|---|---|---|---|
| 6155_5 | Vitamin and mineral supplements: Vitamin E | -0.1305 | 0.1697 | 0.3046 | 3.35E-05 | -3.62425 | 0.015626 |
| L12_ATROPHICSKIN | Atrophic disorders of skin | 0.1658 | 0.5029 | 0.6003 | 0.0341 | -1.15502 | 0.015742 |
| L90 | Diagnoses - main ICD10: L90 Atrophic disorders of skin | 0.1658 | 0.5029 | 0.6003 | 0.0341 | -1.15502 | 0.015742 |
| 20459 | General happiness with own health | -0.2354 | 0.0004 | 0.1978 | 2.95E-06 | -5.51423 | 0.015998 |
| 22616_1 | Breathing problems during period of job: Yes | -0.1894 | 0.0341 | 0.2426 | 0.0006 | -3.79022 | 0.016237 |
| S82 | Diagnoses - main ICD10: S82 Fracture of lower leg, including ankle | -0.1687 | 0.2255 | 0.2632 | 0.0459 | -2.25303 | 0.016258 |
| 4803_13 | Tinnitus: Yes, now some of the time | -0.3401 | 0.0051 | 0.0912 | 0.3169 | -2.83899 | 0.016378 |
| S02 | Diagnoses - main ICD10: S02 Fracture of skull and facial bones | -0.1524 | 0.3346 | 0.2786 | 0.0247 | -2.14608 | 0.016439 |
| 2463 | Fractured/broken bones in last 5 years | -0.1785 | 0.0062 | 0.252 | 5.99E-07 | -5.22008 | 0.016541 |
| 20117_1 | Alcohol drinker status: Previous | -0.0983 | 0.1486 | 0.3321 | 8.82E-11 | -5.05164 | 0.016561 |
| K02 | Diagnoses - main ICD10: K02 Dental caries | 0.0065 | 0.9697 | 0.4369 | 0.0158 | -1.72847 | 0.016561 |
| 20003_1140923346 | Treatment/medication code: co-codamol | 0.0448 | 0.5646 | 0.4748 | 5.92E-16 | -4.41205 | 0.016643 |
| F5_DEPRESSIO | Depression | -0.2104 | 0.0829 | 0.2173 | 0.0098 | -2.89765 | 0.01712 |
| KRA_PSY_ANYMENTAL | Any mental disorder | -0.193 | 0.0466 | 0.2345 | 0.0016 | -3.50452 | 0.017162 |
| 3571 | Back pain for 3+ months | -0.0775 | 0.3614 | 0.3496 | 1.77E-09 | -4.15157 | 0.017247 |
| IBD_PULM | Respiratory diseases principally affecting the interstitium, IBD co-morbidites | -0.0491 | 0.7732 | 0.3763 | 0.0274 | -1.76372 | 0.017609 |
| 20002_1330 | Non-cancer illness code, self-reported: iron deficiency anaemia | -0.4244 | 0.0063 | -0.0013 | 0.9879 | -2.37569 | 0.01811 |
| 2907 | Ever stopped smoking for 6+ months | -0.0549 | 0.5935 | 0.368 | 2.12E-05 | -3.14444 | 0.018154 |
| 20003_1140881856 | Treatment/medication code: salbutamol | -0.171 | 0.1618 | 0.2514 | 0.0014 | -2.90441 | 0.018264 |
| M75 | Diagnoses - main ICD10: M75 Shoulder lesions | 0.0237 | 0.7838 | 0.4455 | 9.73E-11 | -3.81634 | 0.018398 |
| 20450 | Feelings of worthlessness during worst period of depression | -0.3628 | 0.0002 | 0.0585 | 0.3559 | -3.66467 | 0.01851 |
| 2654_8 | Non-butter spread type details: Other low or reduced fat spread | -0.2581 | 0.0156 | 0.1627 | 0.0568 | -3.07724 | 0.018622 |
| 22609_1 | Workplace very dusty: Sometimes | 0.1102 | 0.2567 | 0.5308 | 4.65E-09 | -3.16534 | 0.018667 |
| M13_SHOULDER | Shoulder lesions | 0.0244 | 0.7772 | 0.4442 | 1.12E-10 | -3.80149 | 0.018849 |
| 20517 | Trouble falling or staying asleep, or sleeping too much | -0.2041 | 0.0012 | 0.215 | 4.76E-06 | -5.33204 | 0.019009 |
| 20003_1140871310 | Treatment/medication code: ibuprofen | -0.0599 | 0.3966 | 0.3586 | 4.04E-11 | -4.69455 | 0.019147 |
| 767 | Length of working week for main job | 0.0703 | 0.3696 | 0.4886 | 1.16E-20 | -4.4398 | 0.019194 |
| 5463 | Leg pain in calf/calves | -0.0799 | 0.5581 | 0.3378 | 0.0017 | -2.40257 | 0.019333 |
| 6159_4 | Pain type(s) experienced in last month: Back pain | -0.0506 | 0.3052 | 0.3667 | 7.63E-32 | -7.14216 | 0.019426 |
| 20441 | Ever had prolonged loss of interest in normal activities | -0.2826 | 3.92E-08 | 0.1346 | 0.0014 | -6.27329 | 0.01945 |
| I84 | Diagnoses - main ICD10: I84 Haemorrhoids | -0.1389 | 0.0715 | 0.2783 | 1.79E-07 | -4.45109 | 0.01945 |
| 20111_9 | Illnesses of siblings: Diabetes | -0.1651 | 0.0264 | 0.2519 | 2.12E-07 | -4.69241 | 0.019496 |
| 20086_10 | Type of special diet followed: Low calorie | 0.0732 | 0.5147 | 0.4892 | 5.81E-06 | -2.66995 | 0.019732 |
| 6149_4 | Mouth/teeth dental problems: Loose teeth | -0.1396 | 0.0667 | 0.2759 | 1.08E-07 | -4.50672 | 0.019851 |
| 20448 | Professional informed about depression | -0.2034 | 0.0364 | 0.212 | 0.0014 | -3.52717 | 0.019875 |
| 22615_1 | Workplace had a lot of diesel exhaust: Sometimes | 0.0675 | 0.5108 | 0.4819 | 4.35E-09 | -3.15175 | 0.020114 |
| 20435 | Difficulty concentrating during worst depression | -0.2206 | 0.0522 | 0.1937 | 0.0324 | -2.85095 | 0.020139 |
| PRIM_KNEEARTHROSIS | Primary gonarthrosis, bilateral | -0.2127 | 0.0954 | 0.2016 | 0.0453 | -2.55 | 0.020139 |
| M13_DORSALGIANAS | Other/unspecified dorsalgia | 0.0343 | 0.769 | 0.4479 | 2.72E-05 | -2.6132 | 0.020308 |
| 6155_2 | Vitamin and mineral supplements: Vitamin B | -0.1179 | 0.1795 | 0.2955 | 1.6E-05 | -3.71228 | 0.020357 |
| 20434 | Age at last episode of depression | 0.0304 | 0.756 | 0.4436 | 1.96E-06 | -3.05855 | 0.020405 |
| 6179_6 | Mineral and other dietary supplements: Selenium | -0.2116 | 0.0958 | 0.1993 | 0.003 | -2.8607 | 0.020973 |
| M13_ARTHROSIS_OTH | Other arthrosis | 0.1462 | 0.1726 | 0.5567 | 1.05E-09 | -2.91661 | 0.021073 |
| Z47 | Diagnoses - main ICD10: Z47 Other orthopaedic follow-up care | -0.1181 | 0.3229 | 0.2923 | 0.0027 | -2.66451 | 0.021098 |

| | | | | | | | |
|---|---|---|---|---|---|---|---|
| 6154_6 | Medication for pain relief, constipation, heartburn: Laxatives (e.g. Dulcolax, Senokot) | 0.0082 | 0.9089 | 0.4177 | 1.99E-12 | -4.38719 | 0.021324 |
| 20003_1140863152 | Treatment/medication code: diazepam | 0.0231 | 0.8613 | 0.4323 | 0.0018 | -2.13639 | 0.0214 |
| 137 | Number of treatments/medications taken | -0.0915 | 0.0365 | 0.3173 | 1.73E-31 | -7.94194 | 0.021502 |
| 20084_478 | Vitamin and/or mineral supplement use: Vitamin C | -0.1248 | 0.4281 | 0.2839 | 0.0231 | -2.0332 | 0.021528 |
| 20126_4 | Bipolar and major depression status: Probable Recurrent major depression (moderate) | -0.3484 | 0.0018 | 0.0596 | 0.4305 | -3.02866 | 0.021707 |
| 20509 | Recent inability to stop or control worrying | -0.3208 | 5.80E-06 | 0.087 | 0.1013 | -4.60791 | 0.021758 |
| K11_GASTRODUOULC | Gastroduodenal ulcer | 0.0557 | 0.7051 | 0.4632 | 0.0165 | -1.67732 | 0.021836 |
| 41215_0 | Detention categories: Informal, not formally detained | 0.071 | 0.3947 | 0.4784 | 6.46E-09 | -3.47491 | 0.021861 |
| 6151_1 | Fractured bone site(s): Ankle | -0.1673 | 0.0951 | 0.24 | 0.0038 | -3.13192 | 0.021887 |
| 6155_3 | Vitamin and mineral supplements: Vitamin C | -0.1306 | 0.1585 | 0.2759 | 6.37E-07 | -3.76713 | 0.022095 |
| 20418 | Impact on normal roles during worst period of anxiety | -0.18 | 0.1592 | 0.2263 | 0.0044 | -2.69892 | 0.022147 |
| Z09 | Diagnoses - main ICD10: Z09 Follow-up examination after treatment for conditions other than malignant neoplasms | -0.2798 | 0.036 | 0.1264 | 0.1328 | -2.57582 | 0.022173 |
| 6145_3 | Illness, injury, bereavement, stress in last 2 years: Death of a close relative | 0.0244 | 0.8069 | 0.4301 | 2.72E-08 | -3.20827 | 0.022304 |
| 20111_1 | Illnesses of siblings: Heart disease | -0.1153 | 0.0886 | 0.2896 | 4.93E-10 | -4.92652 | 0.022516 |
| 20550_1 | Activities undertaken to treat anxiety: Talking therapies, such as psychotherapy, counselling, group therapy or CBT | -0.3526 | 8.24E-06 | 0.0518 | 0.4078 | -4.00896 | 0.022648 |
| J84 | Diagnoses - main ICD10: J84 Other interstitial pulmonary diseases | -0.0718 | 0.5725 | 0.3308 | 0.0014 | -2.45578 | 0.023132 |
| 20505 | Recent easy annoyance or irritability | -0.2107 | 0.0058 | 0.1915 | 0.0009 | -4.2124 | 0.023241 |
| 20002_1190 | Non-cancer illness code, self-reported: peritonitis | -0.1117 | 0.4431 | 0.2905 | 0.0186 | -2.10732 | 0.023241 |
| 3476 | Difficulty not smoking for 1 day | -0.2114 | 0.0218 | 0.1894 | 0.0052 | -3.50212 | 0.023624 |
| 20446 | Ever had prolonged feelings of sadness or depression | -0.3965 | 6.60E-11 | 0.0041 | 0.9359 | -5.02834 | 0.023679 |
| 4620 | Number of depression episodes | -0.3584 | 0.0144 | 0.042 | 0.6843 | -2.23438 | 0.023735 |
| 22660_103 | Gap coding: Full-time or part-time education | -0.5284 | 3.28E-05 | -0.1287 | 0.1672 | -2.53472 | 0.023929 |
| 20002_1286 | Non-cancer illness code, self-reported: depression | -0.2558 | 0.0002 | 0.1428 | 0.0056 | -4.67724 | 0.024238 |
| 2277 | Frequency of solarium/sunlamp use | 0.1906 | 0.0044 | 0.5888 | 1.22E-49 | -5.12103 | 0.024351 |
| K50 | Diagnoses - main ICD10: K50 Crohn's disease [regional enteritis] | -0.2166 | 0.0909 | 0.1815 | 0.025 | -2.62667 | 0.024379 |
| 22617_7112 | Job SOC coding: Retail cashiers and check-out operators | -0.1362 | 0.4294 | 0.26 | 0.0457 | -1.8351 | 0.024922 |
| 670_2 | Type of accommodation lived in: A flat, maisonette or apartment | -0.304 | 1.27E-06 | 0.0909 | 0.0483 | -5.07816 | 0.0253 |
| M18 | Diagnoses - main ICD10: M18 Arthrosis of first carpometacarpal joint | 0.0731 | 0.6375 | 0.4667 | 0.0174 | -1.57416 | 0.025682 |
| 20003_1141146138 | Treatment/medication code: lipitor 10mg tablet | -0.0824 | 0.7246 | 0.3107 | 0.0315 | -1.42935 | 0.02583 |
| M19 | Diagnoses - main ICD10: M19 Other arthrosis | 0.1765 | 0.1739 | 0.5687 | 3.29E-08 | -2.36779 | 0.026099 |
| 20520 | Recent worrying too much about different things | -0.3278 | 4.33E-05 | 0.0636 | 0.3054 | -3.85874 | 0.02634 |
| 20437 | Thoughts of death during worst depression | -0.0918 | 0.4343 | 0.2986 | 0.0021 | -2.56463 | 0.026644 |
| 20002_1478 | Non-cancer illness code, self-reported: cervical spondylosis | -0.0852 | 0.5141 | 0.3048 | 0.0033 | -2.34061 | 0.026766 |
| M15 | Diagnoses - main ICD10: M15 Polyarthrosis | -0.0776 | 0.6018 | 0.3114 | 0.0092 | -2.03997 | 0.027074 |
| 20002_1266 | Non-cancer illness code, self-reported: head injury | -0.0866 | 0.4577 | 0.3021 | 0.0247 | -2.18365 | 0.027167 |
| H65 | Diagnoses - main ICD10: H65 Nonsuppurative otitis media | -0.3058 | 0.0368 | 0.0828 | 0.5021 | -2.03025 | 0.027198 |
| 20003_1140872228 | Treatment/medication code: gabapentin | 0.0434 | 0.7138 | 0.432 | 0.0005 | -2.25994 | 0.027198 |
| 4836 | Loud music exposure frequency | -0.1695 | 0.0413 | 0.2177 | 0.0003 | -3.78631 | 0.027635 |

| | | | | | | | |
|---|---|---|---|---|---|---|---|
| 20536_1 | Weight change during worst episode of depression: Gained weight | -0.1029 | 0.2862 | 0.2836 | 2.6E-06 | -3.39658 | 0.027856 |
| I20 | Diagnoses - main ICD10: I20 Angina pectoris | -0.0699 | 0.3982 | 0.316 | 3.31E-06 | -3.60169 | 0.028047 |
| 20003_1141174520 | Treatment/medication code: symbicort 100/6 turbohaler | -0.1576 | 0.2222 | 0.2283 | 0.0181 | -2.39333 | 0.028047 |
| RHEUMA_SEROPOS | Seropositive rheumatoid arthritis | -0.0603 | 0.7863 | 0.3249 | 0.0461 | -1.39688 | 0.02827 |
| 20002_1138 | Non-cancer illness code, self-reported: gastro-oesophageal reflux (gord) / gastric reflux | -0.0454 | 0.5272 | 0.3395 | 5.4E-10 | -4.26047 | 0.028367 |
| 20002_1464 | Non-cancer illness code, self-reported: rheumatoid arthritis | -0.1086 | 0.3209 | 0.2762 | 0.0006 | -2.8281 | 0.028399 |
| 6159_6 | Pain type(s) experienced in last month: Hip pain | -0.0393 | 0.5054 | 0.3453 | 3.57E-17 | -5.35304 | 0.028463 |
| 20003_1140910766 | Treatment/medication code: nicorandil | -0.0911 | 0.5065 | 0.2933 | 0.0047 | -2.23513 | 0.028528 |
| 2159 | Ever had same-sex intercourse | -0.2556 | 0.0021 | 0.1277 | 0.052 | -3.6156 | 0.028885 |
| 20003_1140883548 | Treatment/medication code: ipratropium | -0.0996 | 0.6016 | 0.283 | 0.0294 | -1.65696 | 0.029114 |
| 24004 | Nitrogen oxides air pollution; 2010 | -0.3048 | 3.49E-05 | 0.0759 | 0.1999 | -4.03052 | 0.029744 |
| 6179_4 | Mineral and other dietary supplements: Zinc | -0.2467 | 0.0059 | 0.1336 | 0.0277 | -3.51398 | 0.029878 |
| 2664_4 | Reason for reducing amount of alcohol drunk: Financial reasons | -0.0225 | 0.8602 | 0.3575 | 0.0001 | -2.39372 | 0.029979 |
| 2644 | Light smokers, at least 100 smokes in lifetime | 0.0773 | 0.2363 | 0.4562 | 6.06E-32 | -4.98832 | 0.030351 |
| 796 | Distance between home and job workplace | -0.1241 | 0.6423 | 0.2546 | 0.0018 | -1.35563 | 0.030419 |
| 22613_1 | Worked with paints, thinners or glues: Sometimes | -0.0913 | 0.3392 | 0.2871 | 8.11E-05 | -3.15114 | 0.030522 |
| K11_GALLBILPANC | Disorders of gallbladder, biliary tract and pancreas | -0.1714 | 0.0314 | 0.2068 | 0.0004 | -3.83228 | 0.03059 |
| 20111_12 | Illnesses of siblings: Severe depression | -0.4007 | 1.93E-08 | -0.0227 | 0.6311 | -4.42067 | 0.030659 |
| 20429 | Easily tired during worst period of anxiety | -0.1874 | 0.0909 | 0.1895 | 0.032 | -2.65903 | 0.031038 |
| IX_CIRCULATORY | Diseases of the circulatory system | -0.024 | 0.6445 | 0.3519 | 5.1E-22 | -5.91675 | 0.031386 |
| 826 | Job involves shift work | 0.0257 | 0.6879 | 0.4014 | 1.63E-16 | -4.67624 | 0.031456 |
| 6159_7 | Pain type(s) experienced in last month: Knee pain | -0.0374 | 0.4114 | 0.3378 | 9.2E-25 | -6.67264 | 0.031631 |
| 20506 | Recent feelings or nervousness or anxiety | -0.38 | 7.97E-07 | -0.0053 | 0.9302 | -3.82641 | 0.031808 |
| 24006 | Particulate matter air pollution (pm2.5); 2010 | -0.2778 | 0.0004 | 0.096 | 0.1126 | -3.7626 | 0.032127 |
| 6154_2 | Medication for pain relief, constipation, heartburn: Ibuprofen (e.g. Nurofen) | -0.0635 | 0.3159 | 0.31 | 3.14E-10 | -4.65518 | 0.032234 |
| 6179_5 | Mineral and other dietary supplements: Iron | -0.3689 | 0.0004 | 0.0029 | 0.9702 | -2.88082 | 0.032847 |
| 20116_1 | Smoking status: Previous | -0.0895 | 0.0534 | 0.2823 | 9.52E-25 | -6.90422 | 0.032847 |
| J45 | Diagnoses - main ICD10: J45 Asthma | -0.1329 | 0.405 | 0.2382 | 0.0157 | -1.97813 | 0.033102 |
| 20003_1140871462 | Treatment/medication code: naproxen | 0.0855 | 0.5239 | 0.4563 | 0.0364 | -1.44846 | 0.033211 |
| 22606_1 | Workplace very noisy: Sometimes | 0.0167 | 0.8662 | 0.3865 | 1.07E-06 | -2.91503 | 0.03358 |
| 2030 | Guilty feelings | -0.2561 | 5.80E-06 | 0.1133 | 0.0058 | -5.28715 | 0.033728 |
| 6159_1 | Pain type(s) experienced in last month: Headache | -0.1656 | 0.0011 | 0.2027 | 1.52E-06 | -5.5833 | 0.034138 |
| 2453 | Cancer diagnosed by doctor | -0.1671 | 0.0831 | 0.2003 | 0.0081 | -2.99747 | 0.034476 |
| 6142_7 | Current employment status: Full or part-time student | -0.4107 | 0.0478 | -0.0439 | 0.7791 | -1.41163 | 0.034704 |
| XII_SKIN_SUBCUTAN | Diseases of the skin and subcutaneous tissue | 0.1367 | 0.3462 | 0.5029 | 4.33E-07 | -2.08141 | 0.034932 |
| 20002_1202 | Non-cancer illness code, self-reported: urinary frequency / incontinence | -0.0731 | 0.6273 | 0.293 | 0.0074 | -1.96678 | 0.034971 |
| 21001 | Body mass index (BMI) | -0.0549 | 0.1272 | 0.31 | 1.08E-40 | -8.52012 | 0.035432 |
| 20439 | Frequency of depressed days during worst episode of depression | -0.4684 | 0.0001 | -0.1038 | 0.196 | -2.51306 | 0.035548 |
| 22604_4 | Work hours - lumped category: Over 40 hours | 0.0304 | 0.7966 | 0.3948 | 2.49E-05 | -2.41815 | 0.035625 |
| 20550_3 | Activities undertaken to treat anxiety: Other therapeutic activities such as mindfulness, yoga or art classes | -0.3861 | 7.85E-06 | -0.0224 | 0.727 | -3.38069 | 0.035898 |

| | | | | | | | |
|---|---|---|---|---|---|---|---|
| 23104 | Body mass index (BMI) | -0.0529 | 0.1447 | 0.3104 | 9.23E-40 | -8.4014 | 0.036054 |
| M05 | Diagnoses - main ICD10: M05 Seropositive rheumatoid arthritis | -0.0464 | 0.8248 | 0.3168 | 0.0355 | -1.40648 | 0.036094 |
| S09 | Diagnoses - main ICD10: S09 Other and unspecified injuries of head | -0.0709 | 0.5535 | 0.2922 | 0.0033 | -2.33484 | 0.036133 |
| 22504 | Bring up phlegm/sputum/mucus on most days | -0.1741 | 0.0512 | 0.1889 | 0.013 | -3.09562 | 0.036172 |
| N50 | Diagnoses - main ICD10: N50 Other disorders of male genital organs | 0.098 | 0.582 | 0.4604 | 0.0123 | -1.41558 | 0.036408 |
| 6152_5 | Blood clot, DVT, bronchitis, emphysema, asthma, rhinitis, eczema, allergy diagnosed by doctor: Blood clot in the leg (DVT) | 0.0463 | 0.6134 | 0.4084 | 3.76E-09 | -3.15032 | 0.036527 |
| M13_RHEUMATISM | Rheumatoid arthritis | 0.0773 | 0.6293 | 0.439 | 0.006 | -1.599 | 0.036686 |
| J44 | Diagnoses - main ICD10: J44 Other chronic obstructive pulmonary disease | 0.0304 | 0.7734 | 0.3916 | 9.06E-09 | -2.87456 | 0.036885 |
| 6142_5 | Current employment status: Unemployed | -0.1939 | 0.0764 | 0.1672 | 0.0281 | -2.70964 | 0.036925 |
| K80 | Diagnoses - main ICD10: K80 Cholelithiasis | -0.1766 | 0.0202 | 0.1834 | 0.0022 | -3.71723 | 0.037367 |
| 23112 | Leg fat mass (right) | -0.054 | 0.1107 | 0.3059 | 2.79E-39 | -8.74921 | 0.037407 |
| 24007 | Particulate matter air pollution (pm2.5) absorbance; 2010 | -0.2789 | 0.0015 | 0.0809 | 0.211 | -3.30386 | 0.037448 |
| 23116 | Leg fat mass (left) | -0.0524 | 0.1218 | 0.307 | 1.14E-39 | -8.73705 | 0.03761 |
| 4559 | Family relationship satisfaction | -0.3741 | 5.30E-09 | -0.0171 | 0.6803 | -4.67181 | 0.038594 |
| 24003 | Nitrogen dioxide air pollution; 2010 | -0.3425 | 7.23E-07 | 0.0141 | 0.8063 | -3.96969 | 0.03876 |
| 20490 | Sexually molested as a child | -0.1211 | 0.2268 | 0.2345 | 0.0007 | -2.92019 | 0.039177 |
| 2237 | Plays computer games | -0.1788 | 5.59E-06 | 0.1764 | 5.52E-09 | -7.14636 | 0.039345 |
| 20539 | Frequency of inability to stop worrying during worst period of anxiety | -0.1011 | 0.3431 | 0.2537 | 0.001 | -2.69928 | 0.039514 |
| 20002_1465 | Non-cancer illness code, self-reported: osteoarthritis | -0.0246 | 0.6821 | 0.3289 | 9.95E-16 | -4.8589 | 0.040066 |
| 3090 | Used an inhaler for chest within last hour | -0.1263 | 0.2889 | 0.2261 | 0.0346 | -2.20105 | 0.040538 |
| M23 | Diagnoses - main ICD10: M23 Internal derangement of knee | 0.0795 | 0.256 | 0.4317 | 1.78E-12 | -3.78788 | 0.040625 |
| M13 | Diagnoses - main ICD10: M13 Other arthritis | -0.0132 | 0.935 | 0.3389 | 0.0029 | -1.77806 | 0.040668 |
| RHEU_ARTHRITIS_OTH | Other arthritis (FG) | -0.0132 | 0.935 | 0.3389 | 0.0029 | -1.77806 | 0.040668 |
| 20530 | Witnessed sudden violent death | -0.0778 | 0.4019 | 0.2741 | 2.6E-05 | -3.10277 | 0.040754 |
| 23124 | Arm fat mass (left) | -0.07 | 0.0441 | 0.2797 | 5.63E-33 | -8.33896 | 0.041716 |
| COPD_EARLYANDLATER | COPD, early/later onset | 0.0445 | 0.6857 | 0.3941 | 8.62E-09 | -2.70138 | 0.04176 |
| AB1_INFECTIONS | Certain infectious and parasitic diseases | 0.088 | 0.5183 | 0.4376 | 0.0017 | -1.79118 | 0.04176 |
| I9_HYPTENS | Hypertension | -0.0461 | 0.7593 | 0.3031 | 0.0117 | -1.81241 | 0.041937 |
| G6_NERPLEX | Nerve, nerve root and plexus disorders | 0.0623 | 0.2861 | 0.4113 | 7.67E-21 | -4.77689 | 0.042026 |
| 20544_15 | Mental health problems ever diagnosed by a professional: Anxiety, nerves or generalized anxiety disorder | -0.3373 | 3.51E-06 | 0.011 | 0.8621 | -3.61076 | 0.042338 |
| M13_SPONDYLOPATHY | #Spondylopathies | 0.0223 | 0.8549 | 0.3699 | 1.69E-06 | -2.41097 | 0.042652 |
| 23120 | Arm fat mass (right) | -0.0719 | 0.0377 | 0.2755 | 1.84E-32 | -8.33931 | 0.042741 |
| 20002_1094 | Non-cancer illness code, self-reported: deep venous thrombosis (dvt) | 0.055 | 0.5524 | 0.4007 | 3.54E-09 | -3.01062 | 0.043512 |
| 22611_2 | Workplace had a lot of cigarette smoke from other people smoking: Often | -0.0132 | 0.8948 | 0.3325 | 5.06E-06 | -2.80265 | 0.043512 |
| 20116_2 | Smoking status: Current | 0.0838 | 0.101 | 0.429 | 5.42E-40 | -5.70522 | 0.043741 |
| 20003_1140925800 | Treatment/medication code: movicol oral powder | -0.0256 | 0.8707 | 0.3192 | 0.0169 | -1.66948 | 0.043925 |
| 3849 | Number of pregnancy terminations | -0.0862 | 0.3723 | 0.2584 | 2.49E-05 | -3.01202 | 0.044017 |
| M13_ARTHRITISNAS | Ohter specific/unspecified arthritis | -0.0604 | 0.7485 | 0.2836 | 0.0201 | -1.53262 | 0.044294 |
| 48 | Waist circumference | -0.0741 | 0.0133 | 0.269 | 8.49E-35 | -9.25737 | 0.044713 |
| M13_MENISCUSDERANGEMENTS | Meniscus derangement | 0.0777 | 0.2851 | 0.4208 | 4.06E-11 | -3.54959 | 0.044713 |

| | | | | | | | |
|---|---|---|---|---|---|---|---|
| 6154_1 | Medication for pain relief, constipation, heartburn: Aspirin | -0.0691 | 0.3091 | 0.2709 | 5.48E-08 | -4.03391 | 0.046179 |
| 20003_2038460150 | Treatment/medication code: paracetamol | -0.022 | 0.6632 | 0.317 | 7.06E-15 | -5.22669 | 0.04666 |
| 20003_1140876404 | Treatment/medication code: aqueous cream bp | -0.0329 | 0.8266 | 0.3056 | 0.0227 | -1.68182 | 0.046902 |
| 22618_0 | Breathing problems improved/stopped away from workplace or on holiday: No | -0.0816 | 0.4324 | 0.2563 | 0.0054 | -2.43251 | 0.047194 |
| 22502 | Cough on most days | -0.1856 | 0.0358 | 0.1499 | 0.0232 | -3.04115 | 0.048375 |
| 20003_1140874744 | Treatment/medication code: gliclazide | -0.0721 | 0.4001 | 0.2621 | 3.16E-06 | -3.261 | 0.049025 |
| 6153_3 | Medication for cholesterol, blood pressure, diabetes, or take exogenous hormones: Insulin | -0.1467 | 0.1639 | 0.1869 | 0.0295 | -2.45347 | 0.049328 |
| 20003_1141146188 | Treatment/medication code: latanoprost | 0.0321 | 0.8154 | 0.3652 | 0.0013 | -1.87189 | 0.049581 |
| 20547_3 | Activities undertaken to treat depression: Other therapeutic activities such as mindfulness, yoga or art classes | -0.3299 | 2.17E-05 | 0.0018 | 0.9754 | -3.3999 | 0.050295 |
| 20003_1140884600 | Treatment/medication code: metformin | -0.1203 | 0.0421 | 0.211 | 1.31E-08 | -4.74204 | 0.0505 |
| 23115 | Leg fat percentage (left) | -0.01 | 0.749 | 0.321 | 2.32E-42 | -8.45682 | 0.050655 |
| 20003_1141192410 | Treatment/medication code: rosuvastatin | -0.0072 | 0.9563 | 0.322 | 0.0015 | -1.98596 | 0.05159 |
| 20002_1136 | Non-cancer illness code, self-reported: liver/biliary/pancreas problem | -0.3645 | 0.0303 | -0.0358 | 0.7291 | -1.66497 | 0.051852 |
| 23100 | Whole body fat mass | -0.0515 | 0.1057 | 0.2769 | 8.1E-34 | -8.39275 | 0.05201 |
| 20160 | Ever smoked | -0.068 | 0.0973 | 0.2597 | 5.56E-22 | -6.68273 | 0.05238 |
| 23111 | Leg fat percentage (right) | -0.0124 | 0.6931 | 0.3139 | 1.06E-39 | -8.29839 | 0.053126 |
| 23119 | Arm fat percentage (right) | -0.0618 | 0.0556 | 0.2635 | 1.36E-30 | -8.21585 | 0.053664 |
| 2834 | Bilateral oophorectomy (both ovaries removed) | 0.0291 | 0.7205 | 0.3529 | 9.78E-11 | -3.30823 | 0.054479 |
| 22506_113 | Tobacco smoking: Ex-smoker | 0.0549 | 0.4625 | 0.3777 | 5.05E-16 | -3.66637 | 0.055028 |
| 20003_1140863202 | Treatment/medication code: temazepam | -0.0569 | 0.6663 | 0.2647 | 0.0318 | -1.78044 | 0.055692 |
| ASTHMA_HOSPITAL1 | Asthma, hospital admissions 1 | -0.055 | 0.7267 | 0.2649 | 0.0038 | -1.75695 | 0.056644 |
| 23123 | Arm fat percentage (left) | -0.0528 | 0.1071 | 0.2663 | 3.18E-30 | -7.94731 | 0.057096 |
| 30290 | High light scatter reticulocyte percentage | -0.1444 | 0.0005 | 0.1746 | 2.07E-13 | -6.66802 | 0.057153 |
| 100730 | Spirits intake | -0.0726 | 0.5921 | 0.2438 | 0.04 | -1.75642 | 0.058643 |
| 134 | Number of self-reported cancers | -0.1091 | 0.2419 | 0.2072 | 0.0034 | -2.70061 | 0.058701 |
| 6015 | Chest pain felt during physical activity | 0.0642 | 0.6916 | 0.3797 | 0.0045 | -1.50324 | 0.059167 |
| M70 | Diagnoses - main ICD10: M70 Soft tissue disorders related to use, overuse and pressure | 0.1888 | 0.2627 | 0.5032 | 0.015 | -1.17799 | 0.059811 |
| M13_SOFTOVERUSE | Soft tissue disorders related to use, overuse and pressure | 0.193 | 0.2576 | 0.5069 | 0.0156 | -1.16178 | 0.060106 |
| 4526 | Happiness | -0.4127 | 1.76E-14 | -0.0989 | 0.0132 | -4.68491 | 0.060165 |
| 20111_3 | Illnesses of siblings: Lung cancer | 0.2372 | 0.2425 | 0.5509 | 0.0001 | -1.2573 | 0.060224 |
| 5107 | 3mm strong meridian angle (right) | -0.1392 | 0.1096 | 0.1737 | 0.001 | -3.07932 | 0.060698 |
| 20003_1140868226 | Treatment/medication code: aspirin | -0.0443 | 0.4891 | 0.2681 | 5.94E-08 | -3.86113 | 0.060996 |
| 1990 | Tense / 'highly strung' | -0.2304 | 1.82E-05 | 0.0818 | 0.0738 | -4.41868 | 0.061116 |
| 22611_1 | Workplace had a lot of cigarette smoke from other people smoking: Sometimes | -0.043 | 0.7418 | 0.2685 | 0.004 | -1.94246 | 0.061536 |
| ASTHMA_CHILD | Childhood asthma (age<16) | -0.0579 | 0.7083 | 0.2536 | 0.0056 | -1.73312 | 0.061536 |
| J10_ASTHMA | Asthma | -0.0579 | 0.7083 | 0.2536 | 0.0056 | -1.73312 | 0.061536 |
| J10_ASTHMA_MAIN | Asthma | -0.0579 | 0.7083 | 0.2536 | 0.0056 | -1.73312 | 0.061536 |
| 22607_1 | Workplace very cold: Sometimes | 0.0727 | 0.3809 | 0.3839 | 1.63E-11 | -3.09328 | 0.061716 |
| K85 | Diagnoses - main ICD10: K85 Acute pancreatitis | -0.084 | 0.4798 | 0.2248 | 0.0202 | -2.01407 | 0.063177 |
| 20419 | Difficulty concentrating during worst period of anxiety | 0.026 | 0.8749 | 0.3341 | 0.0102 | -1.46575 | 0.063607 |
| D23 | Diagnoses - main ICD10: D23 Other benign neoplasms of skin | 0.0046 | 0.9748 | 0.3121 | 0.0089 | -1.62904 | 0.063979 |
| 20107_3 | Illnesses of father: Lung cancer | 0.041 | 0.5255 | 0.3476 | 1.03E-11 | -3.7259 | 0.064539 |
| 2926 | Number of unsuccessful stop-smoking attempts | -0.1935 | 0.0275 | 0.1129 | 0.0531 | -2.90568 | 0.064663 |
| 24024 | Average 24-hour sound level of noise pollution | -0.1043 | 0.452 | 0.2006 | 0.0291 | -1.83252 | 0.065606 |
| 24023 | Average 16-hour sound level of noise pollution | -0.1043 | 0.452 | 0.2002 | 0.0294 | -1.83011 | 0.06586 |
| 24020 | Average daytime sound level of noise pollution | -0.1046 | 0.451 | 0.1999 | 0.0302 | -1.82738 | 0.06586 |

| | | | | | | | |
|---|---|---|---|---|---|---|---|
| K21 | Diagnoses - main ICD10: K21 Gastro-oesophageal reflux disease | 0.0211 | 0.8153 | 0.3255 | 8.2E-10 | -2.90244 | 0.065923 |
| 24022 | Average night-time sound level of noise pollution | -0.1046 | 0.451 | 0.1998 | 0.0304 | -1.82617 | 0.065923 |
| 24021 | Average evening sound level of noise pollution | -0.1044 | 0.4517 | 0.1994 | 0.0302 | -1.82439 | 0.066305 |
| 2247_1 | Hearing difficulty/problems: Yes | -0.2275 | 3.21E-07 | 0.076 | 0.03 | -5.36078 | 0.066496 |
| F10 | Diagnoses - main ICD10: F10 Mental and behavioural disorders due to use of alcohol | -0.015 | 0.9007 | 0.2879 | 0.0065 | -1.89237 | 0.06688 |
| 20425 | Ever worried more than most people would in similar situation | -0.3923 | 1.87E-07 | -0.0906 | 0.1297 | -3.13758 | 0.067654 |
| I9_UAP | Unstable angina pectoris | -0.0178 | 0.8511 | 0.2829 | 0.0002 | -2.46943 | 0.068304 |
| 21002 | Weight | -0.0645 | 0.0436 | 0.236 | 2.31E-27 | -7.76084 | 0.068434 |
| 20110_1 | Illnesses of mother: Heart disease | -0.1639 | 0.0155 | 0.1361 | 0.0109 | -3.47925 | 0.068761 |
| 23098 | Weight | -0.0632 | 0.0489 | 0.2366 | 5.4E-27 | -7.70389 | 0.068893 |
| 20003_1140879616 | Treatment/medication code: amitriptyline | 0.0557 | 0.5554 | 0.3549 | 7.28E-07 | -2.52359 | 0.069287 |
| 6143_1 | Transport type for commuting to job workplace: Car/motor vehicle | 0.338 | 1.38E-06 | 0.6369 | 1.01E-34 | -3.4324 | 0.069485 |
| 23128 | Trunk fat mass | -0.0446 | 0.1476 | 0.254 | 1.44E-29 | -7.82843 | 0.069684 |
| 22147 | Age asthma diagnosed by doctor | -0.2176 | 0.0479 | 0.0809 | 0.3235 | -2.17565 | 0.06975 |
| 23099 | Body fat percentage | -0.0244 | 0.4222 | 0.274 | 1.19E-32 | -7.82784 | 0.069816 |
| 20421 | Ever felt worried, tense, or anxious for most of a month or longer | -0.3418 | 5.75E-07 | -0.0443 | 0.394 | -3.46245 | 0.070415 |
| M13_ROTATORCUFF | Rotator cuff syndrome | 0.014 | 0.8677 | 0.3107 | 2.19E-05 | -2.66652 | 0.070951 |
| 20003_1141182628 | Treatment/medication code: tiotropium | 0.1038 | 0.7135 | 0.3995 | 0.0071 | -0.92614 | 0.071625 |
| 20405_2 | Ever had known person concerned about, or recommend reduction of, alcohol consumption: Yes, during the last year | -0.266 | 0.0077 | 0.0288 | 0.6885 | -2.39784 | 0.072236 |
| 6144_4 | Never eat eggs, dairy, wheat, sugar: Sugar or foods/drinks containing sugar | -0.0194 | 0.7296 | 0.2733 | 2.4E-15 | -4.45007 | 0.073677 |
| 2664_1 | Reason for reducing amount of alcohol drunk: Illness or ill health | -0.0541 | 0.6402 | 0.2382 | 0.0231 | -1.87154 | 0.073954 |
| 20003_1140865396 | Treatment/medication code: buscopan 10mg tablet | 0.1608 | 0.3408 | 0.4527 | 0.0315 | -1.08183 | 0.074231 |
| 680_2 | Own or rent accommodation lived in: Own with a mortgage | 0.2442 | 0.0025 | 0.5359 | 4.16E-21 | -2.95587 | 0.074371 |
| 6155_4 | Vitamin and mineral supplements: Vitamin D | -0.3175 | 0.0025 | -0.0258 | 0.7569 | -2.17485 | 0.074371 |
| 6148_1 | Eye problems/disorders: Diabetes related eye disease | -0.0048 | 0.9582 | 0.2869 | 4.31E-05 | -2.52065 | 0.074371 |
| 49 | Hip circumference | -0.0809 | 0.0134 | 0.2108 | 6.76E-20 | -7.2859 | 0.074371 |
| M13_RECUDISLOCATIO | Recurrent dislocation and subluxation of joint | 0.1574 | 0.364 | 0.4476 | 0.0322 | -1.06861 | 0.075421 |
| E87 | Diagnoses - main ICD10: E87 Other disorders of fluid, electrolyte and acid-base balance | 0.008 | 0.959 | 0.2979 | 0.0167 | -1.44907 | 0.075632 |
| 20003_1141145660 | Treatment/medication code: valsartan | 0.0271 | 0.7945 | 0.3163 | 0.0001 | -2.17421 | 0.076127 |
| 30240 | Reticulocyte percentage | -0.129 | 0.0036 | 0.1596 | 2.9E-11 | -5.72807 | 0.076553 |
| 22617_8211 | Job SOC coding: Heavy goods vehicle drivers | 0.0866 | 0.4701 | 0.3752 | 0.0005 | -1.7877 | 0.076553 |
| 20110_12 | Illnesses of mother: Severe depression | -0.3261 | 0.0003 | -0.0376 | 0.5595 | -2.60882 | 0.076624 |
| 3591 | Ever had hysterectomy (womb removed) | 0.0721 | 0.4473 | 0.36 | 1.77E-09 | -2.56665 | 0.077052 |
| 2257 | Hearing difficulty/problems with background noise | -0.344 | 6.56E-15 | -0.0564 | 0.1035 | -5.13083 | 0.077267 |
| G56 | Diagnoses - main ICD10: G56 Mononeuropathies of upper limb | 0.0539 | 0.3356 | 0.3413 | 2.09E-14 | -4.0189 | 0.077411 |
| 3606 | Chest pain or discomfort walking normally | 0.187 | 0.0184 | 0.4738 | 1.47E-21 | -3.06452 | 0.077842 |
| G6_CARPTU | Carpal tunnel syndrome | 0.0399 | 0.4898 | 0.3259 | 2.83E-13 | -3.91744 | 0.078421 |
| 2443 | Diabetes diagnosed by doctor | -0.0977 | 0.0618 | 0.1875 | 3.46E-09 | -4.66341 | 0.079002 |
| 20003_1140911732 | Treatment/medication code: garlic product | 0.1558 | 0.3082 | 0.4401 | 0.003 | -1.33384 | 0.07966 |
| 6155_7 | Vitamin and mineral supplements: Multivitamins +/- minerals | -0.1348 | 0.0209 | 0.1474 | 0.0006 | -3.88483 | 0.081212 |
| 24016 | Nitrogen dioxide air pollution; 2005 | -0.3585 | 9.12E-09 | -0.0767 | 0.1763 | -3.34232 | 0.08151 |
| N20 | Diagnoses - main ICD10: N20 Calculus of kidney and ureter | -0.1898 | 0.0178 | 0.0913 | 0.156 | -2.73501 | 0.082034 |

| | | | | | | | |
|---|---|---|---|---|---|---|---|
| T84 | Diagnoses - main ICD10: T84 Complications of internal orthopaedic prosthetic devices, implants and grafts | 0.1268 | 0.2433 | 0.4079 | 6.83E-06 | -1.98558 | 0.082034 |
| 30250 | Reticulocyte count | -0.1244 | 0.0033 | 0.1557 | 1.84E-11 | -5.80585 | 0.082787 |
| 24017 | Nitrogen dioxide air pollution; 2006 | -0.3464 | 5.37E-08 | -0.0669 | 0.2331 | -3.29282 | 0.083241 |
| 6154_3 | Medication for pain relief, constipation, heartburn: Paracetamol | -0.0027 | 0.9532 | 0.2768 | 2.02E-13 | -4.64487 | 0.083241 |
| KNEE_ARTHROSIS | Gonarthrosis [arthrosis of knee](FG) | 0.0689 | 0.2305 | 0.3482 | 5.68E-18 | -3.98235 | 0.083392 |
| M17 | Diagnoses - main ICD10: M17 Gonarthrosis [arthrosis of knee] | 0.0689 | 0.2305 | 0.3482 | 5.68E-18 | -3.98235 | 0.083392 |
| I9_HYPERTENSION | Hypertensive diseases | -0.0071 | 0.962 | 0.2721 | 0.0153 | -1.49561 | 0.083468 |
| 2704 | Years since last cervical smear test | -0.077 | 0.4728 | 0.2012 | 0.005 | -2.15574 | 0.084231 |
| 30300 | High light scatter reticulocyte count | -0.1054 | 0.0031 | 0.1722 | 1.07E-13 | -6.53294 | 0.084691 |
| 20127 | Neuroticism score | -0.2245 | 2.47E-05 | 0.0522 | 0.2189 | -4.06736 | 0.085384 |
| Z42 | Diagnoses - main ICD10: Z42 Follow-up care involving plastic surgery | -0.048 | 0.7321 | 0.2272 | 0.0111 | -1.65452 | 0.08655 |
| N35 | Diagnoses - main ICD10: N35 Urethral stricture | 0.0487 | 0.7778 | 0.322 | 0.0129 | -1.2661 | 0.088043 |
| 20003_1140926606 | Treatment/medication code: salbutamol 100micrograms spacehaler | -0.0493 | 0.7371 | 0.2212 | 0.0088 | -1.59745 | 0.090278 |
| 40001_C349 | Underlying (primary) cause of death: ICD10: C34.9 Bronchus or lung, unspecified | -0.0119 | 0.9282 | 0.2584 | 0.0284 | -1.52723 | 0.090439 |
| 24019 | Particulate matter air pollution (pm10); 2007 | -0.3465 | 2.58E-08 | -0.0772 | 0.1688 | -3.21507 | 0.091248 |
| 23127 | Trunk fat percentage | -0.0229 | 0.4451 | 0.246 | 2.96E-27 | -7.14773 | 0.091574 |
| 20002_1453 | Non-cancer illness code, self-reported: psoriasis | -0.2296 | 0.0399 | 0.038 | 0.6196 | -1.97741 | 0.092637 |
| R63 | Diagnoses - main ICD10: R63 Symptoms and signs concerning food and fluid intake | 0.1526 | 0.4166 | 0.4198 | 0.0197 | -1.02688 | 0.092966 |
| 30530 | Sodium in urine | 0.0039 | 0.9255 | 0.2705 | 4.26E-16 | -4.97394 | 0.093461 |
| II_NEOPLASM | Neoplasms | -0.0824 | 0.2502 | 0.1842 | 0.0006 | -2.98209 | 0.093461 |
| 20003_1140888266 | Treatment/medication code: warfarin | -0.0614 | 0.5934 | 0.2051 | 0.0166 | -1.85894 | 0.093543 |
| 2345 | Ever had bowel cancer screening | -0.118 | 0.0791 | 0.1476 | 0.0064 | -3.07868 | 0.09429 |
| D17 | Diagnoses - main ICD10: D17 Benign lipomatous neoplasm | -0.0107 | 0.9283 | 0.2547 | 0.0231 | -1.62701 | 0.094457 |
| 20002_1291 | Non-cancer illness code, self-reported: mania/bipolar disorder/manic depression | -0.4231 | 0.0005 | -0.1578 | 0.0651 | -1.77622 | 0.09454 |
| 1080 | Time spent using computer | -0.2339 | 1.31E-08 | 0.031 | 0.3432 | -5.03615 | 0.094874 |
| 30280 | Immature reticulocyte fraction | -0.0896 | 0.009 | 0.1743 | 3.57E-12 | -6.20898 | 0.095713 |
| Z03 | Diagnoses - main ICD10: Z03 Medical observation and evaluation for suspected diseases and conditions | 0.1214 | 0.3876 | 0.3841 | 0.0014 | -1.41976 | 0.096726 |
| M48 | Diagnoses - main ICD10: M48 Other spondylopathies | -0.0477 | 0.6339 | 0.2147 | 0.0024 | -2.14218 | 0.096981 |
| 22612_2 | Worked with materials containing asbestos: Often | -0.0096 | 0.9311 | 0.2521 | 0.0129 | -1.7424 | 0.097577 |
| 6149_6 | Mouth/teeth dental problems: Dentures | 0.0981 | 0.0702 | 0.3593 | 3.19E-25 | -4.06206 | 0.098004 |
| K43 | Diagnoses - main ICD10: K43 Ventral hernia | 0.0979 | 0.416 | 0.3565 | 0.0001 | -1.69639 | 0.100248 |
| M13_SPINSTENOSIS | Spinal stenosis | -0.0552 | 0.5798 | 0.2021 | 0.0036 | -2.11712 | 0.101384 |
| 20107_9 | Illnesses of father: Diabetes | -0.1085 | 0.1549 | 0.1484 | 0.0028 | -2.82293 | 0.101736 |
| 20110_9 | Illnesses of mother: Diabetes | -0.1831 | 0.0029 | 0.0736 | 0.1004 | -3.37375 | 0.101912 |
| 3761 | Age hay fever, rhinitis or eczema diagnosed | -0.0391 | 0.5318 | 0.2165 | 1.07E-06 | -3.33396 | 0.102884 |
| 24018 | Nitrogen dioxide air pollution; 2007 | -0.3502 | 3.05E-09 | -0.0955 | 0.081 | -3.16284 | 0.103685 |
| N17 | Diagnoses - main ICD10: N17 Acute renal failure | -6.36E-05 | 0.9997 | 0.2545 | 0.0469 | -1.27082 | 0.103806 |
| 6153_1 | Medication for cholesterol, blood pressure, diabetes, or take exogenous hormones: Cholesterol lowering medication | -0.0348 | 0.4347 | 0.2197 | 1.79E-08 | -4.30107 | 0.103863 |
| 971 | Frequency of walking for pleasure in last 4 weeks | 0.025 | 0.6563 | 0.2783 | 2.38E-11 | -3.61541 | 0.104939 |

| | | | | | | | |
|---|---|---|---|---|---|---|---|
| 4440 | Average monthly spirits intake | 0.0849 | 0.4097 | 0.338 | 4.56E-05 | -1.91427 | 0.105119 |
| 22617_1112 | Job SOC coding: Directors and chief executives of major organisations | 0.1557 | 0.4161 | 0.4083 | 0.0163 | -0.98699 | 0.10557 |
| I42 | Diagnoses - main ICD10: I42 Cardiomyopathy | 0.0554 | 0.7184 | 0.308 | 0.0287 | -1.21184 | 0.10557 |
| 20002_1074 | Non-cancer illness code, self-reported: angina | 0.061 | 0.2794 | 0.313 | 5.5E-13 | -3.54104 | 0.106113 |
| M13_ARTHROSIS | #Arthrosis | 0.1327 | 0.0402 | 0.3838 | 4.56E-23 | -3.32837 | 0.106932 |
| 20002_1474 | Non-cancer illness code, self-reported: hiatus hernia | 0.0741 | 0.4313 | 0.3248 | 1.57E-06 | -2.16222 | 0.107297 |
| 3786 | Age asthma diagnosed | -0.0608 | 0.3548 | 0.1871 | 0.0019 | -2.77986 | 0.109879 |
| 3143 | Ankle spacing width | -0.0641 | 0.07 | 0.182 | 1.06E-14 | -5.79194 | 0.111563 |
| 24009 | Traffic intensity on the nearest road | -0.0278 | 0.8386 | 0.2183 | 0.0269 | -1.46011 | 0.111563 |
| 6150_2 | Vascular/heart problems diagnosed by doctor: Angina | 0.0664 | 0.231 | 0.3118 | 2.22E-12 | -3.45271 | 0.112222 |
| 2634 | Duration of heavy DIY | 0.0443 | 0.5139 | 0.2893 | 9.5E-08 | -2.82 | 0.1126 |
| 4100 | Ankle spacing width (left) | -0.0877 | 0.0266 | 0.157 | 3.72E-09 | -5.13843 | 0.112884 |
| 22660_105 | Gap coding: Looking after the home and/or family | -0.0169 | 0.9172 | 0.2276 | 0.0284 | -1.26689 | 0.113074 |
| I95 | Diagnoses - main ICD10: I95 Hypotension | 0.0194 | 0.8906 | 0.2636 | 0.0165 | -1.36492 | 0.113359 |
| OTHER_ILD_CVD_COMORB | Other ILD-related CVD-co-morbidities | -0.0889 | 0.4749 | 0.1549 | 0.0399 | -1.675 | 0.11374 |
| 22601_82112603 | Job coding: heavy goods vehicle (hgv) driver, lorry or truck driver, tanker driver, haulage driver | 0.1237 | 0.3048 | 0.3657 | 0.0006 | -1.50782 | 0.115465 |
| M13_SYNOTEND | Disorders of synovium and tendon | 0.1696 | 0.1767 | 0.4106 | 8.16E-05 | -1.47745 | 0.116432 |
| 20002_1075 | Non-cancer illness code, self-reported: heart attack/myocardial infarction | 0.0521 | 0.3119 | 0.2916 | 2.25E-11 | -3.54933 | 0.117892 |
| K60 | Diagnoses - main ICD10: K60 Fissure and fistula of anal and rectal regions | -0.0611 | 0.5304 | 0.1781 | 0.0074 | -2.02821 | 0.118186 |
| 20111_8 | Illnesses of siblings: High blood pressure | -0.0699 | 0.1811 | 0.1668 | 6.07E-05 | -3.54613 | 0.120653 |
| 20002_1220 | Non-cancer illness code, self-reported: diabetes | -0.0499 | 0.3539 | 0.1864 | 6.36E-08 | -3.6973 | 0.121051 |
| L03 | Diagnoses - main ICD10: L03 Cellulitis | 0.0538 | 0.6894 | 0.2901 | 0.0004 | -1.49778 | 0.121051 |
| 20003_1141146234 | Treatment/medication code: atorvastatin | -0.0429 | 0.51 | 0.1934 | 0.0003 | -2.80009 | 0.121051 |
| I9_IHD | Ischaemic heart disease, wide definition | -0.0067 | 0.8904 | 0.2288 | 9.96E-11 | -3.93263 | 0.12185 |
| 5474 | Leg pain when walking uphill or hurrying | 0.0866 | 0.4438 | 0.3212 | 0.001 | -1.57325 | 0.122753 |
| C18 | Diagnoses - main ICD10: C18 Malignant neoplasm of colon | -0.278 | 0.0084 | -0.0442 | 0.5573 | -1.80411 | 0.12356 |
| 23117 | Leg fat-free mass (left) | -0.0769 | 0.0181 | 0.1569 | 5.01E-12 | -5.89769 | 0.12356 |
| 23126 | Arm predicted mass (left) | -0.0477 | 0.1419 | 0.1851 | 3.38E-17 | -5.93181 | 0.124574 |
| K22 | Diagnoses - main ICD10: K22 Other diseases of oesophagus | 0.005 | 0.9672 | 0.2374 | 0.0018 | -1.62838 | 0.124981 |
| 23118 | Leg predicted mass (left) | -0.0764 | 0.0189 | 0.1556 | 8.08E-12 | -5.84383 | 0.125389 |
| I9_CHD | Major coronary heart disease event | 0.0047 | 0.9337 | 0.2361 | 7.61E-08 | -3.21631 | 0.126003 |
| I9_CHD_NOREV | Major coronary heart disease event excluding revascularizations | 0.0047 | 0.9337 | 0.2361 | 7.61E-08 | -3.21631 | 0.126003 |
| M20 | Diagnoses - main ICD10: M20 Acquired deformities of fingers and toes | -0.0615 | 0.3388 | 0.1696 | 0.002 | -2.73333 | 0.12631 |
| 23105 | Basal metabolic rate | -0.0651 | 0.0378 | 0.1655 | 3.68E-14 | -6.03653 | 0.126824 |
| 20458 | General happiness | -0.4258 | 1.79E-12 | -0.1953 | 7.05E-05 | -2.96123 | 0.126927 |
| 23125 | Arm fat-free mass (left) | -0.0498 | 0.1285 | 0.1805 | 2.57E-16 | -5.84343 | 0.127134 |
| 4119 | Ankle spacing width (right) | -0.0673 | 0.0901 | 0.1626 | 9.96E-10 | -4.81088 | 0.127546 |
| 20002_1408 | Non-cancer illness code, self-reported: alcohol dependency | 0.1107 | 0.3837 | 0.3396 | 0.0186 | -1.19037 | 0.128583 |
| 22601_11612669 | Job coding: road transport/traffic manager, haulage contractor/manager, taxi/courier service manager, bus/coach station manager, removals manager, shipping manager | 0.1562 | 0.3975 | 0.3832 | 0.0387 | -0.86787 | 0.130568 |
| 5201 | logMAR, final (right) | 0.0755 | 0.5541 | 0.2983 | 0.0003 | -1.46763 | 0.135031 |
| D12 | Diagnoses - main ICD10: D12 Benign neoplasm of colon, rectum, anus and anal canal | -0.0978 | 0.1255 | 0.1249 | 0.019 | -2.67879 | 0.135139 |
| K57 | Diagnoses - main ICD10: K57 Diverticular disease of intestine | -0.0244 | 0.6239 | 0.1972 | 1.74E-06 | -3.42857 | 0.136325 |
| 23114 | Leg predicted mass (right) | -0.0755 | 0.0188 | 0.146 | 1.27E-10 | -5.63393 | 0.136434 |
| 23113 | Leg fat-free mass (right) | -0.0755 | 0.0189 | 0.1456 | 1.27E-10 | -5.6203 | 0.136867 |
| 20107_2 | Illnesses of father: Stroke | -0.2982 | 0.0257 | -0.0775 | 0.4041 | -1.35559 | 0.137301 |

| | | | | | | | |
|---|---|---|---|---|---|---|---|
| I21 | Diagnoses - main ICD10: I21 Acute myocardial infarction | -0.0119 | 0.8496 | 0.2087 | 2.18E-05 | -2.77339 | 0.13741 |
| 20003_1140884444 | Treatment/medication code: codeine | -0.0583 | 0.57 | 0.1617 | 0.0489 | -1.67322 | 0.138064 |
| 20002_1294 | Non-cancer illness code, self-reported: back problem | 0.0014 | 0.9904 | 0.221 | 0.0098 | -1.4989 | 0.138501 |
| 6138_5 | Qualifications: NVQ or HND or HNC or equivalent | 0.0735 | 0.2691 | 0.2911 | 3.19E-10 | -2.68541 | 0.1407 |
| 20002_1265 | Non-cancer illness code, self-reported: migraine | -0.1354 | 0.0368 | 0.0813 | 0.076 | -2.72807 | 0.141698 |
| S42 | Diagnoses - main ICD10: S42 Fracture of shoulder and upper arm | -0.3057 | 0.0291 | -0.0893 | 0.4383 | -1.19307 | 0.142031 |
| 20003_1140874420 | Treatment/medication code: quinine | 0.1963 | 0.0688 | 0.412 | 4.85E-07 | -1.59233 | 0.142812 |
| 5087 | Cylindrical power (right) | -0.1809 | 0.0395 | 0.0335 | 0.5677 | -2.02842 | 0.144268 |
| 20002_1309 | Non-cancer illness code, self-reported: osteoporosis | -0.1961 | 0.0122 | 0.0182 | 0.759 | -2.18493 | 0.144381 |
| 22612_1 | Worked with materials containing asbestos: Sometimes | 0.0807 | 0.4395 | 0.2945 | 0.0002 | -1.63605 | 0.144944 |
| 23122 | Arm predicted mass (right) | -0.0352 | 0.2787 | 0.1784 | 1.34E-16 | -5.47367 | 0.14517 |
| R55 | Diagnoses - main ICD10: R55 Syncope and collapse | -0.035 | 0.7277 | 0.1782 | 0.0157 | -1.70879 | 0.145622 |
| 23121 | Arm fat-free mass (right) | -0.0354 | 0.2721 | 0.1776 | 2.07E-16 | -5.48167 | 0.145848 |
| 6150_1 | Vascular/heart problems diagnosed by doctor: Heart attack | 0.0749 | 0.158 | 0.2859 | 8.41E-11 | -3.0597 | 0.148126 |
| M51 | Diagnoses - main ICD10: M51 Other intervertebral disk disorders | 0.1536 | 0.1317 | 0.3633 | 1.28E-06 | -1.65738 | 0.149619 |
| 22601_11123211 | Job coding: director or chief executive of private organisation/company, company chairman or president, general manager | 0.1833 | 0.3625 | 0.392 | 0.0154 | -0.80808 | 0.150774 |
| 2654_4 | Non-butter spread type details: Soft (tub) margarine | 0.0386 | 0.5739 | 0.2471 | 9.11E-07 | -2.45107 | 0.151006 |
| 4230 | Signal-to-noise-ratio (SNR) of triplet (left) | -0.0708 | 0.4423 | 0.1367 | 0.0335 | -1.84597 | 0.152168 |
| I_INFECT_PARASIT | Certain infectious and parasitic diseases | 0.1174 | 0.3578 | 0.324 | 0.0057 | -1.19292 | 0.153219 |
| N81 | Diagnoses - main ICD10: N81 Female genital prolapse | -0.1308 | 0.0482 | 0.0734 | 0.1792 | -2.37964 | 0.156045 |
| 100010 | Portion size | -0.2044 | 0.0161 | -0.0006 | 0.9914 | -1.99623 | 0.15652 |
| 6138_100 | Qualifications: None of the above | 0.1748 | 4.41E-07 | 0.3776 | 1.51E-53 | -4.78348 | 0.15771 |
| 23101 | Whole body fat-free mass | -0.0646 | 0.035 | 0.1381 | 2.33E-10 | -5.3834 | 0.157829 |
| 22617_2131 | Job SOC coding: IT strategy and planning professionals | -0.3352 | 0.0405 | -0.133 | 0.2626 | -1.00008 | 0.158427 |
| 23102 | Whole body water mass | -0.0634 | 0.039 | 0.1383 | 2.07E-10 | -5.35685 | 0.159026 |
| 20002_1458 | Non-cancer illness code, self-reported: diverticular disease/diverticulitis | -0.1482 | 0.0169 | 0.0533 | 0.2622 | -2.57989 | 0.159266 |
| 1090 | Time spent driving | 0.3468 | 3.26E-12 | 0.5442 | 2.64E-59 | -3.28895 | 0.164239 |
| 20003_1140860954 | Treatment/medication code: isosorbide mononitrate | 0.0776 | 0.5043 | 0.2728 | 0.0072 | -1.26517 | 0.166949 |
| 20003_1140922174 | Treatment/medication code: alendronate sodium | -0.3585 | 0.0003 | -0.1652 | 0.0278 | -1.54863 | 0.169312 |
| 6179_3 | Mineral and other dietary supplements: Calcium | -0.2069 | 0.0021 | -0.016 | 0.7658 | -2.21562 | 0.172328 |
| M67 | Diagnoses - main ICD10: M67 Other disorders of synovium and tendon | 0.055 | 0.6247 | 0.2455 | 0.029 | -1.19737 | 0.172834 |
| K11_HERNIA | Hernia | 0.0293 | 0.6291 | 0.2196 | 1.03E-06 | -2.51581 | 0.173087 |
| 20003_1140884488 | Treatment/medication code: diclofenac | 0.1479 | 0.1487 | 0.3363 | 6.32E-07 | -1.53613 | 0.175505 |
| 22601_21313058 | Job coding: it strategy or planning professional, computer consultant, software consultant | -0.2862 | 0.0353 | -0.0981 | 0.3762 | -1.07228 | 0.175889 |
| 6158_1 | Why reduced smoking: Illness or ill health | 0.0412 | 0.7667 | 0.2291 | 0.0202 | -1.1022 | 0.176145 |
| J47 | Diagnoses - main ICD10: J47 Bronchiectasis | 0.0847 | 0.6282 | 0.2726 | 0.027 | -0.87807 | 0.176145 |
| 3393 | Hearing aid user | 0.0064 | 0.9319 | 0.1933 | 0.0009 | -1.97332 | 0.17743 |
| 41248_6000 | Destinations on discharge from hospital (recoded): Transfer within NHS provider | 0.124 | 0.5455 | 0.309 | 0.0233 | -0.75141 | 0.179886 |
| 6155_1 | Vitamin and mineral supplements: Vitamin A | 0.101 | 0.4645 | 0.2838 | 0.0043 | -1.0747 | 0.182757 |
| 20003_1141176832 | Treatment/medication code: seretide 50 evohaler | 0.0104 | 0.9074 | 0.1926 | 0.0016 | -1.68693 | 0.183545 |

| | | | | | | | |
|---|---|---|---|---|---|---|---|
| 6149_3 | Mouth/teeth dental problems: Bleeding gums | -0.0205 | 0.7664 | 0.1599 | 0.0012 | -2.12525 | 0.185921 |
| 6152_8 | Blood clot, DVT, bronchitis, emphysema, asthma, rhinitis, eczema, allergy diagnosed by doctor: Asthma | -0.0809 | 0.091 | 0.099 | 0.0013 | -3.15603 | 0.186585 |
| 20003_1140864070 | Treatment/medication code: kapake tablet | 0.0245 | 0.8603 | 0.2043 | 0.0406 | -1.05024 | 0.186717 |
| 20002_1111 | Non-cancer illness code, self-reported: asthma | -0.0767 | 0.1091 | 0.1019 | 0.0009 | -3.13621 | 0.188316 |
| 20107_12 | Illnesses of father: Severe depression | -0.2868 | 0.0008 | -0.1083 | 0.1101 | -1.63582 | 0.18845 |
| 20003_1140861998 | Treatment/medication code: ventolin 100micrograms inhaler | -0.0067 | 0.9171 | 0.1711 | 0.0002 | -2.24396 | 0.189387 |
| 20002_1162 | Non-cancer illness code, self-reported: cholelithiasis/gall stones | -0.006 | 0.9515 | 0.1711 | 0.0194 | -1.44594 | 0.190327 |
| 30000 | White blood cell (leukocyte) count | -0.04 | 0.2717 | 0.1368 | 7.45E-09 | -4.0704 | 0.190731 |
| COPD_OPPORTUNIST_INFECTIONS | COPD related to chronic (opportunist) infections | 0.0895 | 0.6096 | 0.266 | 0.0361 | -0.81588 | 0.191135 |
| M13_TRIGGERFINGER | Trigger finger | 0.1416 | 0.2681 | 0.3169 | 0.0008 | -1.10249 | 0.192757 |
| 22617_5314 | Job SOC coding: Plumbers, heating and ventilating engineers | 0.2172 | 0.2864 | 0.3919 | 0.0073 | -0.69691 | 0.193571 |
| 6149_1 | Mouth/teeth dental problems: Mouth ulcers | -0.1359 | 0.0176 | 0.0384 | 0.322 | -2.51876 | 0.194115 |
| I9_CORATHER | Coronary atherosclerosis | 0.033 | 0.5147 | 0.2063 | 1.68E-07 | -2.7023 | 0.195479 |
| 103250 | Vegetarian alternatives intake | -0.3203 | 0.0247 | -0.1477 | 0.2208 | -0.92386 | 0.196437 |
| M13_GANGLION | Ganglion | 0.0685 | 0.5753 | 0.2407 | 0.0269 | -1.05198 | 0.196986 |
| I25 | Diagnoses - main ICD10: I25 Chronic ischaemic heart disease | 0.0354 | 0.4832 | 0.2071 | 3.55E-07 | -2.64727 | 0.197673 |
| 23130 | Trunk predicted mass | -0.059 | 0.0469 | 0.1118 | 1.85E-07 | -4.65836 | 0.198914 |
| 23129 | Trunk fat-free mass | -0.0576 | 0.052 | 0.1124 | 1.49E-07 | -4.64396 | 0.200021 |
| N47 | Diagnoses - main ICD10: N47 Redundant prepuce, phimosis and paraphimosis | 0.0745 | 0.6614 | 0.2443 | 0.0226 | -0.84452 | 0.200299 |
| 30500 | Microalbumin in urine | -0.0153 | 0.9284 | 0.1533 | 0.0483 | -0.9009 | 0.201967 |
| I9_MI_STRICT | Myocardial infarction, strict | 0.0233 | 0.7026 | 0.1918 | 6.84E-05 | -2.16735 | 0.202107 |
| 20003_1140861958 | Treatment/medication code: simvastatin | -0.0154 | 0.7561 | 0.1496 | 0.0002 | -2.59264 | 0.207023 |
| M13_HAMMERTOE | Other hammer toe(s) (acquired) | 0.1965 | 0.2351 | 0.3604 | 0.0157 | -0.73556 | 0.208583 |
| 30140 | Neutrophill count | -0.0561 | 0.15 | 0.1064 | 1.94E-05 | -3.51192 | 0.210578 |
| 3751 | Chest pain or discomfort when walking uphill or hurrying | 0.156 | 0.1224 | 0.3175 | 4.83E-06 | -1.31788 | 0.21201 |
| ENDOMETRIOSIS_UTERUS | Endometriosis of uterus | 0.2277 | 0.3413 | 0.3878 | 0.035 | -0.53038 | 0.214024 |
| N83 | Diagnoses - main ICD10: N83 Noninflammatory disorders of ovary, Fallopian tube and broad ligament | 0.2481 | 0.394 | 0.4076 | 0.0174 | -0.47216 | 0.21489 |
| 5208 | logMAR, final (left) | 0.0022 | 0.984 | 0.1611 | 0.0385 | -1.16123 | 0.215759 |
| G6_ULLNLE | Lesion of ulnar nerve | 0.1358 | 0.4383 | 0.2932 | 0.0416 | -0.69401 | 0.217939 |
| 2306 | Weight change compared with 1 year ago | 0.2528 | 0.0016 | 0.4099 | 1.42E-10 | -1.5332 | 0.218377 |
| 1050 | Time spend outdoors in summer | 0.2642 | 1.22E-10 | 0.4211 | 7.91E-45 | -3.08837 | 0.218669 |
| 4570 | Friendships satisfaction | -0.3637 | 2.66E-11 | -0.2068 | 1.56E-07 | -2.33026 | 0.218669 |
| 20416 | Frequency of consuming six or more units of alcohol | 0.0279 | 0.6656 | 0.1841 | 6.72E-05 | -1.96876 | 0.219693 |
| H26 | Diagnoses - main ICD10: H26 Other cataract | -0.0274 | 0.7317 | 0.1259 | 0.0089 | -1.64226 | 0.223963 |
| 20003_1140879802 | Treatment/medication code: amlodipine | -0.0386 | 0.5564 | 0.1143 | 0.0117 | -1.91596 | 0.224556 |
| 22617_2111 | Job SOC coding: Chemists | -0.3367 | 0.0167 | -0.1854 | 0.109 | -0.83058 | 0.226936 |
| 6157_4 | Why stopped smoking: Financial reasons | -0.3192 | 0.0235 | -0.1686 | 0.1071 | -0.85781 | 0.227982 |
| 6177_1 | Medication for cholesterol, blood pressure or diabetes: Cholesterol lowering medication | 0.0141 | 0.7751 | 0.1646 | 8.77E-06 | -2.4416 | 0.228131 |
| 104900 | Time spent doing vigorous physical activity | 0.147 | 0.2204 | 0.296 | 0.0019 | -0.97284 | 0.230382 |
| C_COLON | Malignant neoplasm of colon | -0.2738 | 0.0102 | -0.1252 | 0.1203 | -1.1126 | 0.230984 |
| C3_COLON | Malignant neoplasm of colon | -0.2738 | 0.0102 | -0.1252 | 0.1203 | -1.1126 | 0.230984 |
| 2395_1 | Hair/balding pattern: Pattern 1 | -0.1173 | 0.001 | 0.0306 | 0.2438 | -3.34602 | 0.23204 |
| 20003_1140916356 | Treatment/medication code: losartan | 0.0526 | 0.6676 | 0.2001 | 0.0062 | -1.03398 | 0.232645 |
| VII_EYE_ADNEXA | Diseases of the eye and adnexa | -0.0174 | 0.8216 | 0.1259 | 0.0178 | -1.53072 | 0.239046 |

| | | | | | | | |
|---|---|---|---|---|---|---|---|
| 30520 | Potassium in urine | -0.0647 | 0.1943 | 0.0774 | 0.0261 | -2.33893 | 0.240893 |
| 22601_21113020 | Job coding: chemist (analytic, developmental, industrial, research) | -0.3422 | 0.0254 | -0.2016 | 0.1194 | -0.70116 | 0.243212 |
| 4803_14 | Tinnitus: Yes, but not now, but have in the past | 0.0272 | 0.796 | 0.163 | 0.0248 | -1.06175 | 0.250714 |
| 20003_1140860840 | Treatment/medication code: nitrolingual 400micrograms spray | 0.2139 | 0.1643 | 0.3486 | 0.0259 | -0.61388 | 0.25245 |
| 6160_2 | Leisure/social activities: Pub or social club | 0.0846 | 0.1212 | 0.2193 | 9.3E-09 | -2.02142 | 0.25245 |
| 3731 | Former alcohol drinker | 0.0609 | 0.5155 | 0.1922 | 0.0069 | -1.11705 | 0.257856 |
| 20003_1140909708 | Treatment/medication code: furosemide | 0.1995 | 0.1457 | 0.3284 | 0.0015 | -0.75037 | 0.261708 |
| 22608_1 | Workplace very hot: Sometimes | 0.1989 | 0.0713 | 0.3277 | 5.66E-07 | -1.00404 | 0.261869 |
| 1070 | Time spent watching television (TV) | 0.0985 | 0.0061 | 0.2251 | 7.53E-15 | -2.74324 | 0.265427 |
| M13_ENTESOPA THYOTH | Other enthesopathies | 0.1801 | 0.2454 | 0.3066 | 0.0268 | -0.60855 | 0.265589 |
| 816 | Job involves heavy manual or physical work | 0.1796 | 2.08E-05 | 0.304 | 2.69E-23 | -2.38649 | 0.269009 |
| 24008 | Particulate matter air pollution 2.5-10um; 2010 | 0.1357 | 0.6731 | 0.2596 | 0.0251 | -0.36224 | 0.269826 |
| 41248_1001 | Destinations on discharge from hospital (recoded): Usual Place of residence: Living with relatives | 0.0781 | 0.4444 | 0.2019 | 0.0219 | -0.91802 | 0.26999 |
| M77 | Diagnoses - main ICD10: M77 Other enthesopathies | 0.1817 | 0.2376 | 0.3049 | 0.0241 | -0.60141 | 0.270973 |
| 4291 | Number of attempts | 0.1904 | 0.0018 | 0.3135 | 5.72E-13 | -1.64305 | 0.271137 |
| 20002_1473 | Non-cancer illness code, self-reported: high cholesterol | 0.0106 | 0.8122 | 0.1332 | 0.0001 | -2.16654 | 0.271958 |
| 2405 | Number of children fathered | 0.3635 | 1.59E-09 | 0.4855 | 1.15E-25 | -1.60512 | 0.272944 |
| 4537 | Work/job satisfaction | -0.2748 | 0.0015 | -0.1529 | 0.0013 | -1.23515 | 0.273109 |
| 20003_1140875408 | Treatment/medication code: allopurinol | 0.0485 | 0.3417 | 0.1701 | 0.0002 | -1.78264 | 0.273603 |
| 6138_4 | Qualifications: CSEs or equivalent | 0.1011 | 0.0777 | 0.2224 | 5.8E-07 | -1.67194 | 0.274097 |
| 4803_11 | Tinnitus: Yes, now most or all of the time | -0.2126 | 0.0133 | -0.0914 | 0.1074 | -1.17787 | 0.274262 |
| 20110_2 | Illnesses of mother: Stroke | -0.2104 | 0.0385 | -0.0897 | 0.2943 | -0.90896 | 0.275088 |
| 30130 | Monocyte count | -0.059 | 0.137 | 0.0615 | 0.0236 | -2.50394 | 0.275418 |
| 5855_1 | Which eye(s) affected by astigmatism: Right eye | 0.1617 | 0.2099 | 0.2794 | 0.0067 | -0.71301 | 0.280066 |
| 5256 | Corneal hysteresis (right) | -0.0374 | 0.4697 | 0.08 | 0.0122 | -1.93253 | 0.280566 |
| 100370 | Intake of sugar added to coffee | 0.0833 | 0.359 | 0.2006 | 0.0105 | -0.97718 | 0.280733 |
| 30120 | Lymphocyte count | 0.0221 | 0.559 | 0.1365 | 1.51E-09 | -2.59759 | 0.285591 |
| 6144_1 | Never eat eggs, dairy, wheat, sugar: Eggs or foods containing eggs | 0.0398 | 0.6005 | 0.1542 | 0.0158 | -1.15214 | 0.285591 |
| PULM_MEDICAT IO_COMORB | Medication related adverse effects (Asthma/COPD) | 0.0083 | 0.9285 | 0.1223 | 0.0184 | -1.08013 | 0.286265 |
| 2624 | Frequency of heavy DIY in last 4 weeks | 0.0993 | 0.275 | 0.2111 | 0.0034 | -0.96296 | 0.289981 |
| 5106 | 6mm strong meridian angle (right) | -0.003 | 0.9734 | 0.108 | 0.0475 | -1.06188 | 0.291338 |
| 6177_2 | Medication for cholesterol, blood pressure or diabetes: Blood pressure medication | 0.0267 | 0.621 | 0.1358 | 0.0002 | -1.66811 | 0.294573 |
| 22601_42153302 | Job coding: all other personal assistants and secretaries | 0.1335 | 0.2383 | 0.2424 | 0.0068 | -0.75432 | 0.294914 |
| 20002_1466 | Non-cancer illness code, self-reported: gout | 0.0044 | 0.9304 | 0.1108 | 0.0102 | -1.60813 | 0.299198 |
| 1060 | Time spent outdoors in winter | 0.2153 | 1.83E-05 | 0.3216 | 1.61E-17 | -1.69159 | 0.29937 |
| 20002_1452 | Non-cancer illness code, self-reported: eczema/dermatitis | -0.2394 | 0.0033 | -0.1332 | 0.0417 | -1.01631 | 0.299542 |
| 6147_4 | Reason for glasses/contact lenses: For 'astigmatism' | -0.3464 | 2.67E-05 | -0.2416 | 0.0001 | -1.0096 | 0.301954 |
| 2724 | Had menopause | -0.0079 | 0.9071 | 0.0968 | 0.0456 | -1.25563 | 0.302127 |
| 894 | Duration of moderate activity | 0.1048 | 0.0591 | 0.2094 | 2.8E-07 | -1.51674 | 0.3023 |
| 20403 | Amount of alcohol drunk on a typical drinking day | 0.0604 | 0.3549 | 0.1639 | 0.0002 | -1.31396 | 0.304202 |
| 20003_1140860806 | Treatment/medication code: ramipril | 0.0177 | 0.8291 | 0.1198 | 0.0443 | -1.00558 | 0.306631 |
| 20107_1 | Illnesses of father: Heart disease | -0.1201 | 0.0278 | -0.0182 | 0.639 | -1.5213 | 0.306979 |
| 20002_1065 | Non-cancer illness code, self-reported: hypertension | -0.0017 | 0.9714 | 0.0996 | 0.0012 | -1.80448 | 0.308023 |
| 1100 | Drive faster than motorway speed limit | 0.0874 | 0.0734 | 0.1884 | 1.46E-09 | -1.74537 | 0.308546 |
| 20001_1061 | Cancer code, self-reported: basal cell carcinoma | -0.1839 | 0.0462 | -0.0841 | 0.1658 | -0.90409 | 0.31064 |
| 20003_1141192736 | Treatment/medication code: ezetimibe | 0.0917 | 0.3708 | 0.1883 | 0.0111 | -0.76341 | 0.316255 |

| | | | | | | | |
|---|---|---|---|---|---|---|---|
| 6150_4 | Vascular/heart problems diagnosed by doctor: High blood pressure | -0.0034 | 0.9436 | 0.0928 | 0.0026 | -1.69929 | 0.31696 |
| 20003_1140879760 | Treatment/medication code: bisoprolol | 0.1238 | 0.1368 | 0.2182 | 0.0034 | -0.84477 | 0.320141 |
| ASTHMA_OPPORTUNIST_INFECTIONS | NA | 0.1388 | 0.3955 | 0.2308 | 0.0499 | -0.45685 | 0.324402 |
| 777 | Frequency of travelling from home to job workplace | 0.0656 | 0.3796 | 0.1565 | 0.0026 | -1.00024 | 0.326363 |
| 991 | Frequency of strenuous sports in last 4 weeks | 0.1892 | 0.1271 | 0.2791 | 0.004 | -0.57081 | 0.328149 |
| 100150 | Drinking water intake | 0.0685 | 0.4498 | 0.1525 | 0.0254 | -0.74074 | 0.33877 |
| 22617_4215 | Job SOC coding: Personal assistants and other secretaries | 0.1345 | 0.2003 | 0.2179 | 0.0071 | -0.6289 | 0.339857 |
| 6153_2 | Medication for cholesterol, blood pressure, diabetes, or take exogenous hormones: Blood pressure medication | 0.014 | 0.8207 | 0.0936 | 0.016 | -1.09006 | 0.346775 |
| 20003_1140888560 | Treatment/medication code: perindopril | 0.0729 | 0.5043 | 0.1518 | 0.0403 | -0.5985 | 0.348055 |
| I9_DVTANDPULM | DVT of lower extremities and pulmonary embolism | 0.0503 | 0.506 | 0.1282 | 0.0103 | -0.85998 | 0.349887 |
| 100250 | Instant coffee intake | 0.1751 | 0.2028 | 0.2526 | 0.0089 | -0.4612 | 0.350621 |
| 806 | Job involves mainly walking or standing | 0.2384 | 1.86E-07 | 0.3146 | 1.88E-22 | -1.36163 | 0.353009 |
| 4294_0 | Final attempt correct: no | 0.3133 | 0.0008 | 0.3877 | 2.41E-06 | -0.5987 | 0.356325 |
| M65 | Diagnoses - main ICD10: M65 Synovitis and tenosynovitis | 0.2389 | 0.0717 | 0.311 | 0.001 | -0.44226 | 0.360578 |
| I9_DISVEINLYMPH | Diseases of veins, lymphatic vessels and lymph nodes, not elsewhere classified | 0.1101 | 0.1227 | 0.1818 | 0.0002 | -0.82605 | 0.361319 |
| 22617_4216 | Job SOC coding: Receptionists | 0.1772 | 0.1858 | 0.2453 | 0.0415 | -0.37817 | 0.368016 |
| I9_VTE | Venous thromboembolism | 0.0599 | 0.4123 | 0.1277 | 0.009 | -0.77164 | 0.368576 |
| 6160_100 | Leisure/social activities: None of the above | 0.0408 | 0.5085 | 0.1083 | 0.0044 | -0.93261 | 0.369136 |
| I83 | Diagnoses - main ICD10: I83 Varicose veins of lower extremities | 0.0945 | 0.1785 | 0.1578 | 0.0014 | -0.73693 | 0.377007 |
| 904 | Number of days/week of vigorous physical activity 10+ minutes | 0.0196 | 0.7034 | 0.0819 | 0.0211 | -0.99732 | 0.378888 |
| 4979 | FI5 : family relationship calculation | -0.2541 | 0.0004 | -0.1951 | 0.0004 | -0.65756 | 0.385118 |
| 2000 | Worry too long after embarrassment | -0.2387 | 3.67E-06 | -0.1799 | 9.32E-08 | -0.95408 | 0.385497 |
| 20003_1141194794 | Treatment/medication code: bendroflumethiazide | 0.0669 | 0.4045 | 0.1218 | 0.0109 | -0.58802 | 0.3929 |
| 1021 | Duration of light DIY | 0.0571 | 0.3585 | 0.109 | 0.0051 | -0.70745 | 0.398621 |
| 5084 | Spherical power (right) | 0.1229 | 0.0057 | 0.1731 | 1.99E-10 | -0.96253 | 0.401872 |
| 22601_21323059 | Job coding: software professional, analyst-programmer, computer programmer, software analyst or engineer, systems designer or programmer | -0.4193 | 0.0107 | -0.3702 | 0.0011 | -0.24619 | 0.40398 |
| I50 | Diagnoses - main ICD10: I50 Heart failure | 0.1441 | 0.1165 | 0.1909 | 0.0313 | -0.36682 | 0.408396 |
| HEARTFAIL | Heart failure | 0.1269 | 0.1502 | 0.1736 | 0.0415 | -0.38082 | 0.408588 |
| I9_HEARTFAIL | Heart failure,strict | 0.1269 | 0.1502 | 0.1736 | 0.0415 | -0.38082 | 0.408588 |
| I9_HEARTFAIL_NS | Heart failure, not strict | 0.1269 | 0.1502 | 0.1736 | 0.0415 | -0.38082 | 0.408588 |
| 4674 | Private healthcare | -0.1991 | 0.0158 | -0.1539 | 0.0193 | -0.42833 | 0.411475 |
| 914 | Duration of vigorous activity | 0.2082 | 0.0033 | 0.2525 | 1.8E-07 | -0.51555 | 0.413209 |
| 5085 | Spherical power (left) | 0.1269 | 0.0071 | 0.1711 | 1E-10 | -0.81787 | 0.413402 |
| COX_ARTHROSIS | Coxarthrosis [arthrosis of hip](FG) | 0.1356 | 0.0862 | 0.177 | 0.0007 | -0.43672 | 0.418809 |
| M16 | Diagnoses - main ICD10: M16 Coxarthrosis [arthrosis of hip] | 0.1356 | 0.0862 | 0.177 | 0.0007 | -0.43672 | 0.418809 |
| 6152_9 | Blood clot, DVT, bronchitis, emphysema, asthma, rhinitis, eczema, allergy diagnosed by doctor: Hayfever, allergic rhinitis or eczema | -0.1164 | 0.0023 | -0.0773 | 0.0028 | -0.84975 | 0.423262 |
| 874 | Duration of walks | 0.2116 | 2.97E-05 | 0.2507 | 2.05E-12 | -0.63056 | 0.423262 |
| 22617_2132 | Job SOC coding: Software professionals | -0.4032 | 0.0088 | -0.3674 | 0.001 | -0.18816 | 0.429668 |
| 6148_2 | Eye problems/disorders: Glaucoma | 0.0959 | 0.2105 | 0.1314 | 0.023 | -0.36994 | 0.430251 |
| 5133 | 6mm strong meridian (right) | 0.036 | 0.4367 | 0.0683 | 0.0496 | -0.55767 | 0.436482 |
| 6157_100 | Why stopped smoking: None of the above | 0.1387 | 0.0492 | 0.1684 | 0.0053 | -0.3197 | 0.441557 |
| 22617_2329 | Job SOC coding: Researchers n.e.c. | -0.2896 | 0.1332 | -0.2652 | 0.042 | -0.10479 | 0.45193 |
| 4290 | Duration screen displayed | 0.1666 | 0.0074 | 0.1898 | 6.22E-08 | -0.32484 | 0.454284 |
| 22617_2211 | Job SOC coding: Medical practitioners | -0.1682 | 0.0722 | -0.1471 | 0.0339 | -0.18108 | 0.458406 |

| | | | | | | | |
|---|---|---|---|---|---|---|---|
| 21021 | Pulse wave Arterial Stiffness index | 0.1073 | 0.2066 | 0.1273 | 0.0169 | -0.19934 | 0.460567 |
| G6_PLANTAR | Lesion of plantar nerve | 0.328 | 0.2171 | 0.3458 | 0.0222 | -0.05823 | 0.464893 |
| 6162_3 | Types of transport used (excluding work): Public transport | -0.3071 | 2.25E-07 | -0.2921 | 3E-13 | -0.2097 | 0.470404 |
| 20003_1140866738 | Treatment/medication code: atenolol | 0.1509 | 0.0471 | 0.1641 | 0.0017 | -0.14299 | 0.47395 |
| 22608_0 | Workplace very hot: Rarely/never | -0.1793 | 0.0369 | -0.168 | 0.0074 | -0.10625 | 0.477696 |
| 5096 | 3mm weak meridian (left) | 0.0657 | 0.1904 | 0.0734 | 0.0345 | -0.12618 | 0.484797 |
| 20128 | Number of fluid intelligence questions attempted within time limit | -0.1461 | 0.002 | -0.1391 | 0.0006 | -0.11228 | 0.486179 |
| 5099 | 3mm weak meridian (right) | 0.0641 | 0.188 | 0.0698 | 0.0394 | -0.09606 | 0.488745 |
| 22601_23293029 | Job coding: university research fellow or assistant/associate | -0.3631 | 0.2119 | -0.3591 | 0.0212 | -0.01212 | 0.492101 |
| 3144 | Heel Broadband ultrasound attenuation, direct entry | 0.0733 | 0.2033 | 0.0771 | 0.0367 | -0.05555 | 0.492496 |
| 6160_5 | Leisure/social activities: Other group activity | -0.2325 | 0.0004 | -0.2309 | 3.7E-06 | -0.01949 | 0.49684 |
| 2217 | Age started wearing glasses or contact lenses | 0.1859 | 5.69E-06 | 0.1842 | 2.87E-08 | 0.032224 | 0.503357 |
| 5012 | FI8 : chained arithmetic | -0.2093 | 5.03E-05 | -0.2115 | 1.73E-07 | 0.033539 | 0.504345 |
| 30070 | Red blood cell (erythrocyte) distribution width | 0.0641 | 0.2679 | 0.0601 | 0.0467 | 0.061253 | 0.507899 |
| 20003_1141200322 | Treatment/medication code: solifenacin | 0.3541 | 0.1714 | 0.3473 | 0.0489 | 0.021706 | 0.513427 |
| 20084_475 | Vitamin and/or mineral supplement use: Vitamin A | 0.323 | 0.1775 | 0.3143 | 0.0418 | 0.030522 | 0.517176 |
| 100280 | Added milk to filtered coffee | -0.2031 | 0.0338 | -0.213 | 0.0031 | 0.082665 | 0.519543 |
| 20002_1387 | Non-cancer illness code, self-reported: hayfever/allergic rhinitis | -0.0973 | 0.0856 | -0.1114 | 0.006 | 0.202424 | 0.527823 |
| 100018 | Retinol | -0.2523 | 0.0519 | -0.2667 | 0.0153 | 0.084706 | 0.528414 |
| 398 | Number of correct matches in round | -0.0856 | 0.2132 | -0.1011 | 0.0423 | 0.182498 | 0.53058 |
| 2744 | Birth weight of first child | -0.0569 | 0.2039 | -0.0741 | 0.023 | 0.310437 | 0.533927 |
| 30090 | Platelet crit | 0.0622 | 0.066 | 0.0448 | 0.0478 | 0.427944 | 0.53432 |
| IBD_ENDOMETRIOSIS | Endometriosis, IBD co-morbidity | 0.1918 | 0.0567 | 0.1705 | 0.0238 | 0.169316 | 0.541987 |
| N80 | Diagnoses - main ICD10: N80 Endometriosis | 0.1918 | 0.0567 | 0.1705 | 0.0238 | 0.169316 | 0.541987 |
| 4195 | Pulse wave reflection index | 0.1396 | 0.297 | 0.1057 | 0.0207 | 0.239764 | 0.566635 |
| 4277 | Volume level set by participant (right) | 0.2941 | 0.0002 | 0.2582 | 1.71E-05 | 0.36283 | 0.570526 |
| 30180 | Lymphocyte percentage | 0.0768 | 0.0166 | 0.0387 | 0.1017 | 0.95486 | 0.574799 |
| 5699 | FI10 : arithmetic sequence recognition | -0.1993 | 0.0084 | -0.2408 | 0.0001 | 0.421709 | 0.581384 |
| 3064 | Peak expiratory flow (PEF) | -0.0423 | 0.3601 | -0.0842 | 0.0105 | 0.738752 | 0.582158 |
| 22601_23143401 | Job coding: secondary school teacher or teaching professional (including head teacher) | -0.1108 | 0.3023 | -0.1564 | 0.0432 | 0.344605 | 0.589295 |
| 22617_2112 | Job SOC coding: Biological scientists and biochemists | -0.3205 | 0.0213 | -0.3665 | 0.0004 | 0.265553 | 0.590065 |
| 50 | Standing height | -0.0286 | 0.294 | -0.0768 | 0.0001 | 1.425156 | 0.594293 |
| 6162_1 | Types of transport used (excluding work): Car/motor vehicle | 0.3247 | 6.04E-09 | 0.2746 | 1.05E-11 | 0.727249 | 0.597936 |
| 3647 | Duration of other exercises | 0.1887 | 0.0106 | 0.1359 | 0.0387 | 0.533968 | 0.603098 |
| 3496 | Wants to stop smoking | -0.1607 | 0.3134 | -0.2203 | 0.0121 | 0.327506 | 0.616017 |
| 2664_5 | Reason for reducing amount of alcohol drunk: Other reason | 0.1691 | 0.0042 | 0.1051 | 0.0277 | 0.842682 | 0.624309 |
| 20153 | Forced expiratory volume in 1-second (FEV1), predicted | -0.0083 | 0.7981 | -0.0781 | 0.0034 | 1.659489 | 0.635151 |
| 100003 | Protein | -0.161 | 0.1661 | -0.2329 | 0.0139 | 0.4794 | 0.639051 |
| ENDOMETRIOSIS_PELVICPERITONEUM | Endometriosis of pelvic peritoneum | 0.3808 | 0.0154 | 0.3058 | 0.0149 | 0.372882 | 0.644782 |
| 22601_41143216 | Job coding: officer of ngo, trade union organiser/official, charity administrator, secretary of research or charity or political or professional or trade association or trade union | -0.1515 | 0.2399 | -0.2274 | 0.0243 | 0.463493 | 0.646439 |
| 20016 | Fluid intelligence score | -0.2085 | 1.62E-07 | -0.2852 | 2.8E-20 | 1.522218 | 0.647911 |
| 6143_3 | Transport type for commuting to job workplace: Public transport | -0.349 | 1.60E-07 | -0.4268 | 1.45E-16 | 0.922767 | 0.64993 |
| 728 | Number of vehicles in household | 0.3508 | 1.39E-14 | 0.2716 | 4.57E-14 | 1.363217 | 0.652493 |

| | | | | | | | |
|---|---|---|---|---|---|---|---|
| 5263 | Intra-ocular pressure, Goldmann-correlated (left) | -0.004 | 0.9216 | -0.0836 | 0.0088 | 1.543999 | 0.653225 |
| 2267 | Use of sun/uv protection | 0.1902 | 9.52E-07 | 0.1085 | 0.0009 | 1.610112 | 0.657054 |
| 6147_1 | Reason for glasses/contact lenses: For short-sightedness, i.e. only or mainly for distance viewing such as driving, cinema etc (called 'myopia') | -0.1734 | 0.0068 | -0.2551 | 5.09E-10 | 1.074907 | 0.657054 |
| 100270 | Filtered coffee intake | -0.132 | 0.1244 | -0.214 | 0.0019 | 0.745077 | 0.6576 |
| 20003_1140874930 | Treatment/medication code: prednisolone | 0.2774 | 0.0372 | 0.1951 | 0.047 | 0.497324 | 0.658145 |
| 6160_4 | Leisure/social activities: Adult education class | -0.2996 | 2.32E-06 | -0.383 | 1.02E-15 | 1.051171 | 0.660143 |
| 100460 | Added milk to standard tea | -0.1115 | 0.3218 | -0.1976 | 0.0155 | 0.619163 | 0.665025 |
| 22617_2113 | Job SOC coding: Physicists, geologists and meteorologists | -0.2368 | 0.024 | -0.324 | 3.33E-05 | 0.666764 | 0.667007 |
| 20154 | Forced expiratory volume in 1-second (FEV1), predicted percentage | -0.0261 | 0.6058 | -0.1143 | 0.0005 | 1.458704 | 0.668804 |
| 2664_3 | Reason for reducing amount of alcohol drunk: Health precaution | -0.1612 | 0.0047 | -0.2512 | 6.02E-08 | 1.223231 | 0.672029 |
| 30270 | Mean sphered cell volume | 0.0862 | 0.0071 | -0.0063 | 0.7732 | 2.387305 | 0.676488 |
| 3063 | Forced expiratory volume in 1-second (FEV1) | -0.0366 | 0.3261 | -0.1358 | 6.56E-07 | 2.149862 | 0.68831 |
| H33 | Diagnoses - main ICD10: H33 Retinal detachments and breaks | -0.0233 | 0.7552 | -0.1226 | 0.0356 | 1.046388 | 0.688485 |
| 20150 | Forced expiratory volume in 1-second (FEV1), Best measure | -0.0338 | 0.3628 | -0.1347 | 1.08E-06 | 2.182077 | 0.69128 |
| 5001 | FI7 : synonym | -0.2661 | 0.0005 | -0.3677 | 3.62E-11 | 1.075248 | 0.692499 |
| 943 | Frequency of stair climbing in last 4 weeks | -0.0336 | 0.4878 | -0.1369 | 7.24E-05 | 1.737962 | 0.695451 |
| I9_PHLETHROMBDVTLOW | DVT of lower extremities | 0.217 | 0.0204 | 0.1133 | 0.0713 | 0.920015 | 0.696144 |
| 4935 | FI1 : numeric addition test | -0.1146 | 0.4555 | -0.2191 | 0.0499 | 0.550294 | 0.697528 |
| 2365 | Ever had prostate specific antigen (PSA) test | 0.1265 | 0.0412 | 0.0219 | 0.686 | 1.271193 | 0.6977 |
| 22617_4114 | Job SOC coding: Officers of non-governmental organisations | -0.1292 | 0.3024 | -0.2344 | 0.0177 | 0.659354 | 0.698736 |
| 2395_3 | Hair/balding pattern: Pattern 3 | 0.0168 | 0.77 | -0.0889 | 0.0301 | 1.496733 | 0.699597 |
| 3062 | Forced vital capacity (FVC) | -0.0347 | 0.3159 | -0.1408 | 2.32E-08 | 2.478728 | 0.700286 |
| 6138_6 | Qualifications: Other professional qualifications eg: nursing, teaching | -0.1687 | 0.0002 | -0.2751 | 8.8E-17 | 1.907429 | 0.700802 |
| 5110 | 6mm asymmetry angle (left) | 0.1522 | 0.0439 | 0.0457 | 0.507 | 1.041944 | 0.700974 |
| 4294_1 | Final attempt correct: yes | -0.2674 | 0.0032 | -0.376 | 1.38E-06 | 0.908321 | 0.704573 |
| 22617_2311 | Job SOC coding: Higher education teaching professionals | -0.253 | 0.0107 | -0.3625 | 5.95E-06 | 0.859337 | 0.706109 |
| 400 | Time to complete round | -0.0161 | 0.6747 | -0.1256 | 1.81E-06 | 2.352664 | 0.706109 |
| 47 | Hand grip strength (right) | 0.0287 | 0.3751 | -0.0808 | 0.0017 | 2.647795 | 0.706109 |
| 4282 | Maximum digits remembered correctly | -0.0893 | 0.226 | -0.1993 | 0.0003 | 1.19228 | 0.706961 |
| 22601_21123021 | Job coding: biologist, botanist, entomologist, zoologist, ecologist, microbiologist/pathologist/virologist/geneticist (not medically qualified) | -0.2638 | 0.0682 | -0.3747 | 0.0024 | 0.583587 | 0.708492 |
| 46 | Hand grip strength (left) | 0.0448 | 0.1734 | -0.0666 | 0.0102 | 2.660523 | 0.709341 |
| 5255 | Intra-ocular pressure, Goldmann-correlated (right) | 0.0209 | 0.6332 | -0.0924 | 0.0085 | 2.018569 | 0.712555 |
| 4270 | Volume level set by participant (left) | 0.3916 | 4.16E-06 | 0.2781 | 2E-05 | 1.058713 | 0.712893 |
| I80 | Diagnoses - main ICD10: I80 Phlebitis and thrombophlebitis | 0.209 | 0.0269 | 0.0952 | 0.1344 | 0.999045 | 0.713398 |
| 6157_3 | Why stopped smoking: Health precaution | -0.1518 | 0.0063 | -0.267 | 6.14E-08 | 1.551842 | 0.715753 |
| 4260 | Round of numeric memory test | -0.0592 | 0.4334 | -0.1749 | 0.0022 | 1.223038 | 0.716592 |
| 2395_4 | Hair/balding pattern: Pattern 4 | 0.1105 | 0.0037 | -0.0064 | 0.8075 | 2.528166 | 0.718599 |
| 20151 | Forced vital capacity (FVC), Best measure | -0.0238 | 0.4803 | -0.1435 | 1.49E-08 | 2.840531 | 0.723258 |
| 4283 | Number of rounds of numeric memory test performed | -0.0567 | 0.4517 | -0.1771 | 0.0019 | 1.275686 | 0.724416 |
| 6160_1 | Leisure/social activities: Sports club or gym | 0.1039 | 0.0257 | -0.0168 | 0.6151 | 2.103091 | 0.724912 |
| 5779 | FI11 : antonym | -0.2172 | 0.12 | -0.3388 | 0.002 | 0.684593 | 0.726397 |
| 22606_0 | Workplace very noisy: Rarely/never | -0.1393 | 0.1024 | -0.2631 | 0.0002 | 1.118066 | 0.73001 |
| 6150_100 | Vascular/heart problems diagnosed by doctor: None of the above | 0.0061 | 0.8943 | -0.1211 | 5.35E-05 | 2.309127 | 0.735546 |
| 2110 | Able to confide | 0.1251 | 0.0045 | -0.0035 | 0.9144 | 2.343285 | 0.737809 |

| | | | | | | | |
|---|---|---|---|---|---|---|---|
| 2385 | Relative age voice broke | 0.039 | 0.4678 | -0.0939 | 0.0167 | 1.99893 | 0.744696 |
| 100024 | Calcium | -0.1577 | 0.0972 | -0.292 | 7.41E-05 | 1.116237 | 0.746917 |
| 6138_1 | Qualifications: College or University degree | -0.2436 | 1.09E-15 | -0.3801 | 1.7E-58 | 3.546807 | 0.750388 |
| C_MALE_GENITAL | NA | 0.1761 | 0.0267 | 0.0391 | 0.4943 | 1.398827 | 0.751173 |
| C3_MALE_GENITAL | malignant neoplasm of male genital organs | 0.1761 | 0.0267 | 0.0391 | 0.4943 | 1.398827 | 0.751173 |
| 1031 | Frequency of friend/family visits | -0.2936 | 6.82E-11 | -0.4317 | 1.29E-41 | 2.503632 | 0.752896 |
| 4990 | FI6 : conditional arithmetic | -0.2219 | 0.0002 | -0.3608 | 3.45E-19 | 1.912953 | 0.754145 |
| 100007 | Polyunsaturated fat | -0.2293 | 0.1376 | -0.3689 | 0.0034 | 0.700719 | 0.755235 |
| 20460 | Belief that own life is meaningful | 0.3574 | 3.78E-09 | 0.2089 | 4.94E-06 | 1.956515 | 0.768865 |
| 100006 | Saturated fat | -0.1278 | 0.2506 | -0.2767 | 0.002 | 1.042672 | 0.769468 |
| 2207 | Wears glasses or contact lenses | 0.0019 | 0.9758 | -0.1527 | 0.0012 | 1.965419 | 0.777957 |
| 100390 | Tea consumed | -0.0051 | 0.9631 | -0.1603 | 0.0308 | 1.164578 | 0.77884 |
| 6177_100 | Medication for cholesterol, blood pressure or diabetes: None of the above | 0.003 | 0.9543 | -0.1547 | 2.02E-05 | 2.458039 | 0.782498 |
| 100004 | Fat | -0.1612 | 0.1232 | -0.3193 | 0.0003 | 1.153879 | 0.78308 |
| 6148_100 | Eye problems/disorders: None of the above | 0.017 | 0.8434 | -0.1427 | 0.0464 | 1.427113 | 0.785399 |
| 6152_100 | Blood clot, DVT, bronchitis, emphysema, asthma, rhinitis, eczema, allergy diagnosed by doctor: None of the above | 0.1193 | 0.0039 | -0.0424 | 0.1168 | 3.277089 | 0.788277 |
| 3581 | Age at menopause (last menstrual period) | -0.002 | 0.9687 | -0.1658 | 1.41E-05 | 2.567395 | 0.791275 |
| 4968 | FI4 : positional arithmetic | -0.1369 | 0.0179 | -0.3082 | 3.07E-14 | 2.425168 | 0.801776 |
| 5983 | ECG, heart rate | -0.0051 | 0.9487 | -0.1801 | 0.0002 | 1.895573 | 0.806836 |
| 2654_6 | Non-butter spread type details: Olive oil based spread (eg: Bertolli) | -0.0673 | 0.584 | -0.2448 | 0.0042 | 1.184483 | 0.81021 |
| 6143_2 | Transport type for commuting to job workplace: Walk | -0.2584 | 0.001 | -0.436 | 1.92E-10 | 1.707129 | 0.810345 |
| 4196 | Pulse wave peak to peak time | 0.0159 | 0.8685 | -0.1654 | 0.0022 | 1.644705 | 0.815269 |
| 22501 | Year ended full time education | -0.2402 | 1.77E-07 | -0.4257 | 2.61E-33 | 3.195826 | 0.820762 |
| 100920_2104 | Type milk consumed: wholemilk | -0.0225 | 0.8686 | -0.2103 | 0.0367 | 1.110832 | 0.823726 |
| 6142_6 | Current employment status: Doing unpaid or voluntary work | -0.1363 | 0.0777 | -0.327 | 1.89E-06 | 1.845184 | 0.827419 |
| 6138_2 | Qualifications: A levels/AS levels or equivalent | -0.2108 | 7.16E-10 | -0.4058 | 2.14E-50 | 4.462488 | 0.832803 |
| 22601_23113399 | Job coding: higher education teaching professional, university lecturer/professor (including college/university head/vice chancellor) | -0.1439 | 0.1124 | -0.3394 | 7.37E-05 | 1.567573 | 0.833422 |
| 6139_3 | Gas or solid-fuel cooking/heating: An open solid fuel fire that you use regularly in winter time | 0.0928 | 0.1507 | -0.104 | 0.0239 | 2.481583 | 0.835024 |
| 6160_3 | Leisure/social activities: Religious group | -0.089 | 0.1187 | -0.2862 | 8.5E-19 | 3.009971 | 0.835515 |
| 22617_2321 | Job SOC coding: Scientific researchers | -0.2904 | 0.0356 | -0.4886 | 1.86E-06 | 1.151907 | 0.836739 |
| 3872 | Age of primiparous women at birth of child | -0.2116 | 0.0065 | -0.4114 | 3.77E-07 | 1.778983 | 0.838684 |
| 20002_1463 | Non-cancer illness code, self-reported: ulcerative colitis | 0.0229 | 0.8122 | -0.1775 | 0.0137 | 1.664447 | 0.839409 |
| 5262 | Intra-ocular pressure, corneal-compensated (left) | 0.0707 | 0.1478 | -0.1302 | 0.0004 | 3.286965 | 0.840012 |
| 709 | Number in household | 0.2297 | 0.0005 | 0.0283 | 0.5816 | 2.418436 | 0.840614 |
| 5254 | Intra-ocular pressure, corneal-compensated (right) | 0.0618 | 0.2149 | -0.1397 | 8.63E-05 | 3.291623 | 0.840734 |
| 5556 | FI9 : concept interpolation | -0.2209 | 0.0009 | -0.4229 | 5.37E-15 | 2.356327 | 0.841334 |
| 5855_3 | Which eye(s) affected by astigmatism: Both eyes | -0.1362 | 0.3544 | -0.3424 | 0.0154 | 1.010586 | 0.846312 |
| 20405_0 | Ever had known person concerned about, or recommend reduction of, alcohol consumption: No | 0.206 | 0.0118 | -0.0019 | 0.9752 | 2.039816 | 0.848298 |
| 100002 | Energy | -0.1269 | 0.1869 | -0.3353 | 3.95E-06 | 1.728302 | 0.848879 |
| 6164_2 | Types of physical activity in last 4 weeks: Other exercises (eg: swimming, cycling, keep fit, bowling) | 0.0697 | 0.1156 | -0.1396 | 7.38E-06 | 3.862748 | 0.84992 |
| 738 | Average total household income before tax | 0.0393 | 0.2935 | -0.1705 | 8.1E-11 | 4.59443 | 0.850497 |

| | | | | | | | |
|---|---|---|---|---|---|---|---|
| 22601_23213026 | Job coding: scientific researcher, scientific officer, medical research associate, experimental officer | -0.2918 | 0.0489 | -0.5019 | 3.45E-06 | 1.145357 | 0.850842 |
| 23108 | Impedance of leg (left) | 0.0731 | 0.0607 | -0.139 | 3.93E-08 | 4.562514 | 0.85313 |
| 23109 | Impedance of arm (right) | -0.0205 | 0.5941 | -0.2334 | 1.07E-23 | 4.739956 | 0.854038 |
| 100009 | Englyst dietary fibre | -0.0268 | 0.8136 | -0.2409 | 0.0008 | 1.591245 | 0.855394 |
| 6138_3 | Qualifications: O levels/GCSEs or equivalent | -0.1148 | 0.0238 | -0.3307 | 1.25E-23 | 3.564026 | 0.857412 |
| 20003_1140876136 | Treatment/medication code: beconase 50micrograms nasal spray | -0.2014 | 0.3221 | -0.4186 | 0.0031 | 0.876188 | 0.858857 |
| 23107 | Impedance of leg (right) | 0.0719 | 0.0558 | -0.1466 | 4.35E-09 | 4.839142 | 0.860292 |
| 100025 | Vitamin E | -0.139 | 0.2631 | -0.3603 | 2.97E-05 | 1.463244 | 0.86335 |
| 6162_4 | Types of transport used (excluding work): Cycle | -0.0511 | 0.3547 | -0.2747 | 9.77E-13 | 3.322437 | 0.865827 |
| 4957 | FI3 : word interpolation | -0.2067 | 0.0005 | -0.4306 | 6.8E-26 | 3.097506 | 0.866148 |
| 6032 | Maximum workload during fitness test | 0.0837 | 0.3607 | -0.1404 | 0.0156 | 2.065971 | 0.866361 |
| 6143_4 | Transport type for commuting to job workplace: Cycle | -0.0776 | 0.1628 | -0.3021 | 4.42E-13 | 3.230216 | 0.866788 |
| 22601_21133023 | Job coding: physicist, astronomer, geologist, geophysicist, meteorologist, oceanographer, seismologist | -0.2246 | 0.0921 | -0.4505 | 0.0004 | 1.222828 | 0.868273 |
| 22607_0 | Workplace very cold: Rarely/never | -0.0616 | 0.4627 | -0.2899 | 1.75E-05 | 2.120127 | 0.870793 |
| M62 | Diagnoses - main ICD10: M62 Other disorders of muscle | 0.3038 | 0.0465 | 0.0739 | 0.5903 | 1.119958 | 0.872454 |
| 20522 | Been in a confiding relationship as an adult | 0.1586 | 0.0296 | -0.0729 | 0.1832 | 2.540048 | 0.8741 |
| 4803_0 | Tinnitus: No, never | 0.2015 | 0.0009 | -0.0306 | 0.5018 | 3.062813 | 0.874713 |
| 5843_3 | Which eye(s) affected by myopia (short sight): Both eyes | -0.0827 | 0.5983 | -0.3167 | 0.0154 | 1.145542 | 0.876642 |
| 23110 | Impedance of arm (left) | 0.0073 | 0.8477 | -0.2302 | 1.93E-23 | 5.361228 | 0.88014 |
| 12340 | QRS duration | 0.3208 | 0.0302 | 0.0817 | 0.3964 | 1.353703 | 0.881716 |
| 23106 | Impedance of whole body | 0.0455 | 0.2357 | -0.1953 | 1.53E-16 | 5.336309 | 0.883374 |
| 100016 | Potassium | 0.0179 | 0.8664 | -0.2234 | 0.0006 | 1.935071 | 0.883859 |
| 6153_100 | Medication for cholesterol, blood pressure, diabetes, or take exogenous hormones: None of the above | 0.0141 | 0.8173 | -0.2293 | 3.88E-09 | 3.364302 | 0.885878 |
| 6164_4 | Types of physical activity in last 4 weeks: Light DIY (eg: pruning, watering the lawn) | -0.1092 | 0.0351 | -0.3554 | 1.03E-24 | 3.952299 | 0.888531 |
| 102800 | Cheese consumers | -0.0362 | 0.7558 | -0.284 | 0.0045 | 1.616273 | 0.890028 |
| 22617_3513 | Job SOC coding: Ship and hovercraft officers | 0.2929 | 0.037 | 0.0432 | 0.6975 | 1.394659 | 0.891786 |
| 6141_1 | How are people in household related to participant: Husband, wife or partner | 0.2462 | 1.57E-06 | -0.0087 | 0.8239 | 3.951819 | 0.896494 |
| 20107_100 | Illnesses of father: None of the above (group 1) | 0.1926 | 0.0324 | -0.0637 | 0.2517 | 2.423945 | 0.897735 |
| 20003_1140909674 | Treatment/medication code: cod liver oil capsule | 0.2545 | 0.0484 | -0.0024 | 0.981 | 1.575888 | 0.898264 |
| 20433 | Age at first episode of depression | 0.4867 | 3.15E-06 | 0.2296 | 0.0009 | 2.052666 | 0.89844 |
| 100017 | Magnesium | -0.0754 | 0.4023 | -0.3331 | 2.91E-07 | 2.321245 | 0.898966 |
| 2764 | Age at last live birth | -0.1322 | 0.0205 | -0.3901 | 2.4E-24 | 3.75098 | 0.899141 |
| 6033 | Maximum heart rate during fitness test | -0.0367 | 0.7513 | -0.2956 | 2.56E-07 | 2.003164 | 0.900013 |
| 924 | Usual walking pace | -0.0752 | 0.0509 | -0.3368 | 2.52E-33 | 5.495204 | 0.902338 |
| 20118_11 | Home area population density - urban or rural: Scotland - Large Urban Area | 0.0811 | 0.3581 | -0.1833 | 0.008 | 2.356816 | 0.904707 |
| 100014 | Folate | 0.0341 | 0.7384 | -0.2371 | 0.0005 | 2.212265 | 0.910285 |
| 4407 | Average monthly red wine intake | 0.0827 | 0.4504 | -0.1911 | 0.0381 | 1.912555 | 0.912352 |
| 5117 | 6mm cylindrical power (right) | 0.2408 | 0.0343 | -0.0347 | 0.622 | 2.059613 | 0.913684 |
| 103990 | Vegetable consumers | -0.1384 | 0.2981 | -0.4187 | 0.0001 | 1.621636 | 0.917364 |
| 100011 | Iron | -0.0805 | 0.4101 | -0.3613 | 6.98E-07 | 2.304651 | 0.917741 |
| 5984 | ECG, load | 0.1278 | 0.2132 | -0.155 | 0.0125 | 2.356364 | 0.919234 |
| 3536 | Age started hormone-replacement therapy (HRT) | -0.1957 | 0.0187 | -0.4826 | 1.34E-22 | 2.966616 | 0.92223 |
| 4288 | Time to answer | 0.0872 | 0.3077 | -0.2012 | 0.0008 | 2.76411 | 0.923305 |
| 100008 | Total sugars | 0.1007 | 0.2641 | -0.1884 | 0.0013 | 2.686342 | 0.923802 |
| 20525 | Able to pay rent/mortgage as an adult | -0.0742 | 0.4612 | -0.3746 | 8.15E-07 | 2.379587 | 0.9315 |
| 6155_100 | Vitamin and mineral supplements: None of the above | 0.1544 | 0.0118 | -0.1467 | 0.0003 | 4.104382 | 0.931957 |

| | | | | | | | |
|---|---|---|---|---|---|---|---|
| 110001 | Invitation to complete online 24-hour recall dietary questionnaire, acceptance | -0.1196 | 0.0253 | -0.4214 | 6.79E-25 | 4.481545 | 0.932411 |
| 2247_0 | Hearing difficulty/problems: No | 0.2291 | 2.68E-07 | -0.078 | 0.0263 | 5.418443 | 0.935773 |
| 100023 | Starch | -0.2572 | 0.0372 | -0.5666 | 2.63E-05 | 1.692372 | 0.937191 |
| 3436 | Age started smoking in current smokers | -0.0581 | 0.6003 | -0.3682 | 5.68E-08 | 2.384127 | 0.937618 |
| 6162_2 | Types of transport used (excluding work): Walk | -0.171 | 0.003 | -0.4826 | 2.81E-41 | 4.585279 | 0.938524 |
| 6164_5 | Types of physical activity in last 4 weeks: Heavy DIY (eg: weeding, lawn mowing, carpentry, digging) | -4.05E-05 | 0.9994 | -0.315 | 9.56E-20 | 4.98931 | 0.940518 |
| 981 | Duration walking for pleasure | -0.0272 | 0.6918 | -0.3427 | 8.76E-14 | 3.823681 | 0.940833 |
| 20003_1141175684 | Treatment/medication code: risedronate sodium | 0.3929 | 0.0398 | 0.0746 | 0.571 | 1.371807 | 0.942449 |
| C_MESOTHELIOMA | Mesothelioma | 0.3531 | 0.0433 | 0.0326 | 0.7856 | 1.513407 | 0.943694 |
| C3_MESOTHELIOMA | Mesothelioma | 0.3531 | 0.0433 | 0.0326 | 0.7856 | 1.513407 | 0.943694 |
| 22601_35132648 | Job coding: ship's engineer, ship's purser | 0.3606 | 0.0215 | 0.0393 | 0.701 | 1.71539 | 0.944141 |
| 20007 | Interpolated Age of participant when cancer first diagnosed | -0.2097 | 0.283 | -0.5343 | 0.029 | 1.03679 | 0.945957 |
| 100005 | Carbohydrate | -0.062 | 0.5406 | -0.3907 | 9.73E-07 | 2.548925 | 0.948148 |
| 6020_33 | Completion status of test: Heart rate reached safety level | 0.0686 | 0.5533 | -0.2719 | 0.0089 | 2.188704 | 0.95406 |
| 699 | Length of time at current address | 0.0265 | 0.7331 | -0.3238 | 3.15E-14 | 3.945388 | 0.958548 |
| 6144_5 | Never eat eggs, dairy, wheat, sugar: I eat all of the above | 0.0336 | 0.5594 | -0.3168 | 4.72E-19 | 5.185282 | 0.958592 |
| 22506_114 | Tobacco smoking: Never smoked | -0.0314 | 0.6672 | -0.3866 | 3.46E-19 | 4.187457 | 0.960655 |
| 845 | Age completed full time education | -0.0163 | 0.6553 | -0.3804 | 1.67E-47 | 8.09323 | 0.964258 |
| 6149_100 | Mouth/teeth dental problems: None of the above | 0.0048 | 0.9328 | -0.3597 | 2E-21 | 5.30348 | 0.964413 |
| 20110_10 | Illnesses of mother: Alzheimer's disease/dementia | 0.1459 | 0.3873 | -0.2225 | 0.0451 | 1.823787 | 0.9659 |
| 104460 | Banana intake | 0.4131 | 0.0005 | 0.0435 | 0.7039 | 2.241032 | 0.966346 |
| 100013 | Vitamin B12 | 0.0812 | 0.7079 | -0.2956 | 0.0382 | 1.452526 | 0.968928 |
| 2867 | Age started smoking in former smokers | -0.079 | 0.284 | -0.4579 | 6.85E-24 | 4.372945 | 0.969649 |
| 84 | Cancer year/age first occurred | -0.1266 | 0.6683 | -0.5157 | 0.0415 | 1.000202 | 0.972957 |
| 20111_100 | Illnesses of siblings: None of the above (group 1) | 0.1423 | 0.0056 | -0.2476 | 2.23E-10 | 6.050461 | 0.973203 |
| 22610_0 | Workplace full of chemical or other fumes: Rarely/never | 0.192 | 0.1089 | -0.2052 | 0.0318 | 2.591521 | 0.975365 |
| 6154_100 | Medication for pain relief, constipation, heartburn: None of the above | 0.0245 | 0.5497 | -0.3734 | 2.28E-29 | 7.54221 | 0.975564 |
| 20536_0 | Weight change during worst episode of depression: Stayed about the same or was on a diet | -0.0298 | 0.7279 | -0.4285 | 2.32E-11 | 3.722683 | 0.97579 |
| 20117_2 | Alcohol drinker status: Current | 0.1386 | 0.0268 | -0.2622 | 9.49E-09 | 5.171182 | 0.976376 |
| 6038 | Number of trend entries | 0.182 | 0.1037 | -0.2207 | 0.0037 | 2.978275 | 0.976895 |
| 670_1 | Type of accommodation lived in: A house or bungalow | 0.2973 | 4.81E-07 | -0.1087 | 0.0147 | 5.483498 | 0.977774 |
| 6156_100 | Manic/hyper symptoms: None of the above | 0.0366 | 0.7846 | -0.3699 | 0.002 | 2.264263 | 0.977905 |
| 6142_2 | Current employment status: Retired | 0.0037 | 0.9735 | -0.4051 | 5.77E-11 | 3.243255 | 0.978498 |
| 20491 | Someone to take to doctor when needed as a child | 0.0464 | 0.6067 | -0.3711 | 1.16E-09 | 3.839984 | 0.98062 |
| 3526 | Mother's age at death | -0.0073 | 0.9388 | -0.4264 | 5.05E-12 | 3.681617 | 0.980991 |
| 20116_0 | Smoking status: Never | 0.0339 | 0.4111 | -0.3863 | 6.93E-56 | 8.766179 | 0.981242 |
| 6164_1 | Types of physical activity in last 4 weeks: Walking for pleasure (not as a means of transport) | 0.0902 | 0.0598 | -0.3367 | 5.82E-25 | 7.367828 | 0.982711 |
| 757 | Time employed in main current job | 0.1496 | 0.2517 | -0.284 | 6.16E-05 | 2.919547 | 0.984081 |
| 2754 | Age at first live birth | -0.016 | 0.7302 | -0.4658 | 2.52E-56 | 8.193173 | 0.987014 |
| 6153_5 | Medication for cholesterol, blood pressure, diabetes, or take exogenous hormones: Oral contraceptive pill or minipill | -0.0057 | 0.9723 | -0.4627 | 0.0005 | 2.161172 | 0.98816 |
| 6034 | Target heart rate achieved | 0.1521 | 0.0958 | -0.325 | 6.51E-05 | 3.900495 | 0.990906 |
| 100760 | Breakfast cereal consumed | 0.1243 | 0.2575 | -0.3562 | 0.0002 | 3.310877 | 0.991311 |
| 2139 | Age first had sexual intercourse | -0.0882 | 0.0579 | -0.5791 | 6.2E-132 | 9.405768 | 0.992452 |
| 6159_100 | Pain type(s) experienced in last month: None of the above | 0.1019 | 0.0112 | -0.3898 | 4.99E-47 | 10.14203 | 0.992534 |

| | | | | | | | |
|---|---|---|---|---|---|---|---|
| 20489 | Felt loved as a child | 0.389 | 1.15E-10 | -0.1236 | 0.0026 | 7.024357 | 0.994418 |
| 20126_0 | Bipolar and major depression status: No Bipolar or Depression | 0.3745 | 8.52E-09 | -0.1496 | 0.0038 | 6.315118 | 0.995263 |
| 2654_7 | Non-butter spread type details: Polyunsaturated/sunflower oil based spread (eg: Flora) | 0.2054 | 0.2245 | -0.3223 | 0.0045 | 2.592512 | 0.995503 |
| 100890 | Milk added to cereal | 0.1391 | 0.2563 | -0.3921 | 0.001 | 3.104019 | 0.995725 |
| 2794 | Age started oral contraceptive pill | 0.0195 | 0.7222 | -0.5123 | 4.37E-43 | 8.029166 | 0.995763 |
| 680_1 | Own or rent accommodation lived in: Own outright (by you or someone in your household) | -0.0708 | 0.2225 | -0.6071 | 2.7E-79 | 8.073612 | 0.996033 |
| 5507 | Leg pain on walking : action taken | 0.0789 | 0.5075 | -0.4582 | 1.76E-06 | 3.514302 | 0.996079 |
| 20111_101 | Illnesses of siblings: None of the above (group 2) | 0.4398 | 2.90E-08 | -0.1185 | 0.0479 | 5.617798 | 0.997143 |
| 3446_3 | Type of tobacco currently smoked: Cigars or pipes | 0.181 | 0.4094 | -0.3828 | 0.0212 | 2.04882 | 0.997372 |
| 104400 | Fruit consumers | 0.2534 | 0.0152 | -0.3259 | 0.0002 | 4.254717 | 0.997933 |
| 6146_100 | Attendance/disability/mobility allowance: None of the above | 0.0381 | 0.5701 | -0.5784 | 3.15E-56 | 8.075184 | 0.998863 |
| 6142_3 | Current employment status: Looking after home and/or family | 0.0645 | 0.5499 | -0.5637 | 2.21E-07 | 4.099683 | 0.999064 |
| 6017 | Able to walk or cycle unaided for 10 minutes | 0.1642 | 0.2862 | -0.4731 | 0.0013 | 2.996432 | 0.999197 |
| 6145_100 | Illness, injury, bereavement, stress in last 2 years: None of the above | 0.1544 | 0.0072 | -0.4924 | 4.83E-29 | 8.94309 | 0.999317 |
| 22660_108 | Gap coding: Retired | 0.1865 | 0.4031 | -0.4795 | 0.0105 | 2.285304 | 0.999511 |
